# Mapping and modeling the semantic space of math concepts

**DOI:** 10.1101/2024.05.27.596021

**Authors:** Samuel Debray, Stanislas Dehaene

## Abstract

Mathematics is an underexplored domain of human cognition. While many studies have focused on subsets of math concepts such as numbers, fractions, or geometric shapes, few have ventured beyond these elementary domains. Here, we attempted to map out the full space of math concepts and to answer two specific questions: can distributed semantic models, such a GloVe, provide a satisfactory fit to human semantic judgments in mathematics? And how does this fit vary with education? We first analyzed all of the French and English Wikipedia pages with math contents, and used a semi-automatic procedure to extract the 1,000 most frequent math terms in both languages. In a second step, we collected extensive behavioral judgments of familiarity and semantic similarity between them. About half of the variance in human similarity judgments was explained by vector embeddings that attempt to capture latent semantic structures based on cooccurence statistics. Participants’ self-reported level of education modulated familiarity and similarity, allowing us to create a partial hierarchy among high-level math concepts. Our results converge onto the proposal of a map of math space, organized as a database of math terms with information about their frequency, familiarity, grade of acquisition, and entanglement with other concepts.

## 1. Introduction

Mathematical cognition is a vast domain of human knowledge, essential to daily life as well as to scientific inquiry, and yet vastly underexplored compared to other domains of language or culture. Most cognitive studies tackle specific and narrow subdomains of elementary mathematics such as integers (Dehaene, 2011; Eger, 2016; Kutter et al., 2018; Shepard et al., 1975), fractions (Behr et al., 1983; Ni & Zhou, 2005; Siegler et al., 2013) or geometric shapes (Dillon et al., 2013; Izard et al., 2011; Sablé-Meyer et al., 2022), and only a few researchers have explored higher-level mathematical concepts in professional mathematicians (Amalric & Dehaene, 2016; Zeki et al., 2014). Without attempting to perform an extensive review, it seems fair to say that, in between those two extremes lies a vast space of mathematical concepts that remains largely unexplored from the cognitive viewpoint, to such an extent that they are not even systematically listed. Although a few dictionaries of mathematical concepts are available (Clapham & Nicholson, 2009; *Dictionnaire des mathématiques*, 2019), they are often technical, aimed for experts, and therefore uneasy to use as a lexical source for cognitive research.

Here, our goal is to take a first step towards a systematic study of the basic vocabulary of math cognition and how it varies with education, ranging from primary school to university level concepts. Inspired by the *THINGS* initiative (Hebart et al., 2019), where researchers from different labs collectively gathered a large database on object recognition including behavioral similarity judgements, fMRI, MEG and EEG (in humans and non-human primates) studies as well as modeling by deep neural networks, we started by creating a dataset of 1,000 mathematical words that could then be used for future studies using behavioral and brain imaging methods. Importantly, we used an unbiased computational method, GloVe (Pennington et al., 2014), to analyze a large mathematical corpus and extract the most frequent words for mathematical concepts, with the goal to obtain an objective picture of the space of math concepts.

From this first step, we obtained a lexicon of the most frequent math concepts, their frequency, their tentative age of acquisition, and a vector-based representation of their putative semantics, based on distributional cooccurence statistics. We then aimed to provide a first test of the cognitive validity of these measures. To this aim, we collected ratings of semantic similarity, which are often used as a marker of the organization of mental representations. This idea dates back to the seminal work of Shepard and Chipman (Shepard & Chipman, 1970). Using the principle of second order isomorphism – which states that there is a relation between the similarities of the internal representation of two objects and the similarities of the corresponding external objects – one can construct psychological representations of the structure of a set of stimuli by collecting subjective similarity measures and analyzing them, for instance using multi-dimensional scaling (Shepard, 1980) or more sophisticated feature-reconstruction methods (Hebart et al., 2020). This approach was first applied to numbers by Shepard et al. (1975), who presented pairs of numbers to participants in various notations (e.g. Arabic numerals, number words, dot patterns) and asked them to rate their conceptual similarity. They showed that the similarity ratings only depended on the judgment task, not the original notation, and that MDS enabled retrieving a conceptual space for the numbers, organized by interpretable semantic dimensions such as number magnitude and odd-even status.

The idea that mental objects, including high-level concepts, are encoded by vectors in a multi-dimensional mental space later became influential in both neuroscience (Ebitz & Hayden, 2021) and computational semantics (Bhatia et al., 2019). Latent Semantic Analysis (Landauer & Dumais, 1997) was one of the earliest methods to obtain semantic vector embedding for words. Since then, many algorithms and similarity measures have been proposed in order to compare human judgements with distributed semantic representations (Hebart et al., 2020; Kramer et al., 2023; Pereira et al., 2016; Richie & Bhatia, 2021). Importantly, distributed semantic representations have also been shown to capture context-dependent information about specific semantic dimensions such as size or dangerosity, which also compares favorably to human judgements (Grand et al., 2022). Finally, beyond behavioral judgements, such embeddings have also been shown to predict neuroimaging data (Huth et al., 2016; Millet et al., 2022; Mitchell et al., 2008; Pereira et al., 2018; Tong et al., 2022). Recently, the use of context-dependent embeddings from large language models has permitted an even better fit to brain activity during sentence processing (Caucheteux & King, 2022; Goldstein et al., 2022; Kumar et al., 2022; Pasquiou et al., 2022; Schrimpf et al., 2021).

Here, we extend this logic to math concepts. We asked people to provide similarity ratings between pairs of math words, and compared these similarities with those predicted by semantic embeddings learnt by the GloVe algorithm on a large math corpus. Although it may seem obvious that math concepts should behave similarly to other concepts, brain imaging indicates that math sentences activate a network of brain areas entirely different from classical language regions (Amalric & Dehaene, 2016, 2018, 2019), thus leaving open the possibility that a different logic or language may be needed to account for the mental organization of math concepts (Dehaene et al., 2022a).

A third goal of our study was to evaluate the impact of math education on such similarity ratings and their putative underlying vector representations. The mental representation of math concepts changes dramatically with education (Carey, 1988, 2009; Dehaene, 2011; Siegler & Opfer, 2003). Using brain imaging, Amalric and Dehaene (Amalric & Dehaene, 2016) showed a drastic enhancement of brain activity in a large-scale math-responsive network in professional mathematicians compared to other adults with more limited math education. Longitudinal, cross-sectional, and cross-cultural comparisons indicate that even the representation of simple integers exhibits massive change in the course of education to counting and the number line (Halberda & Feigenson, 2008; Opfer & Siegler, 2007; Piazza et al., 2013, 2018; Pica et al., 2004). Here, we tested the possibility that similarity ratings would also show a sensitivity to education and could therefore serve as a sensitive marker of the expansion of the conceptual space of mathematics.

In summary, we aimed to provide a 1,000-word vocabulary of math concepts covering all levels and domains. Anticipating on the results, we showed that similarity ratings collected during an online experiment are well captured by GloVe embeddings of the concepts of the vocabulary, and that the quality of the fit increases with participants’ education. In addition, we showed that the spatial layout of the embeddings makes sense and can be used to make predictions about how humans organize their mental map of math concepts.

## 2. Results

### 2.1. Creation of a vocabulary of 1,000 words in French and English

We first created a math vocabulary of 1,000 French words by manually reviewing the 39,345 most frequent words in the French Wikipedia math articles. Then, using GloVe (Pennington et al., 2014), we obtained vector embeddings for the words of this vocabulary. Three distinct embeddings were obtained from three different corpora: the above math corpus (math embedding); a non-math corpus consisting of all non-math pages of the French Wikipedia (non-math embedding); and their concatenation (global embedding) (Fig 1A). Our logic was that many words have both math and non-math meanings. For a given word, we hypothesized that its math embedding would primarily carry information about its math meaning (e.g. “ring” in the context of commutative algebra), while its non-math embedding would carry information about its meaning in everyday language (e.g. “ring” as in engagement ring). We also estimated, for each word of the vocabulary, its school grade of acquisition (hereafter referred to as “word grade”) by examining in which class the corresponding concept was introduced according to the national French curriculum. Finally, we used the above corpora to obtain estimates of each word’s log frequency per million in math and non-math corpora. A summary of the main characteristics of the vocabulary is provided in Table 1. The vocabulary was also translated into English, for which the same measures were obtained. The math vocabulary is available online from https://osf.io/dxg2w.

**Figure 1.**
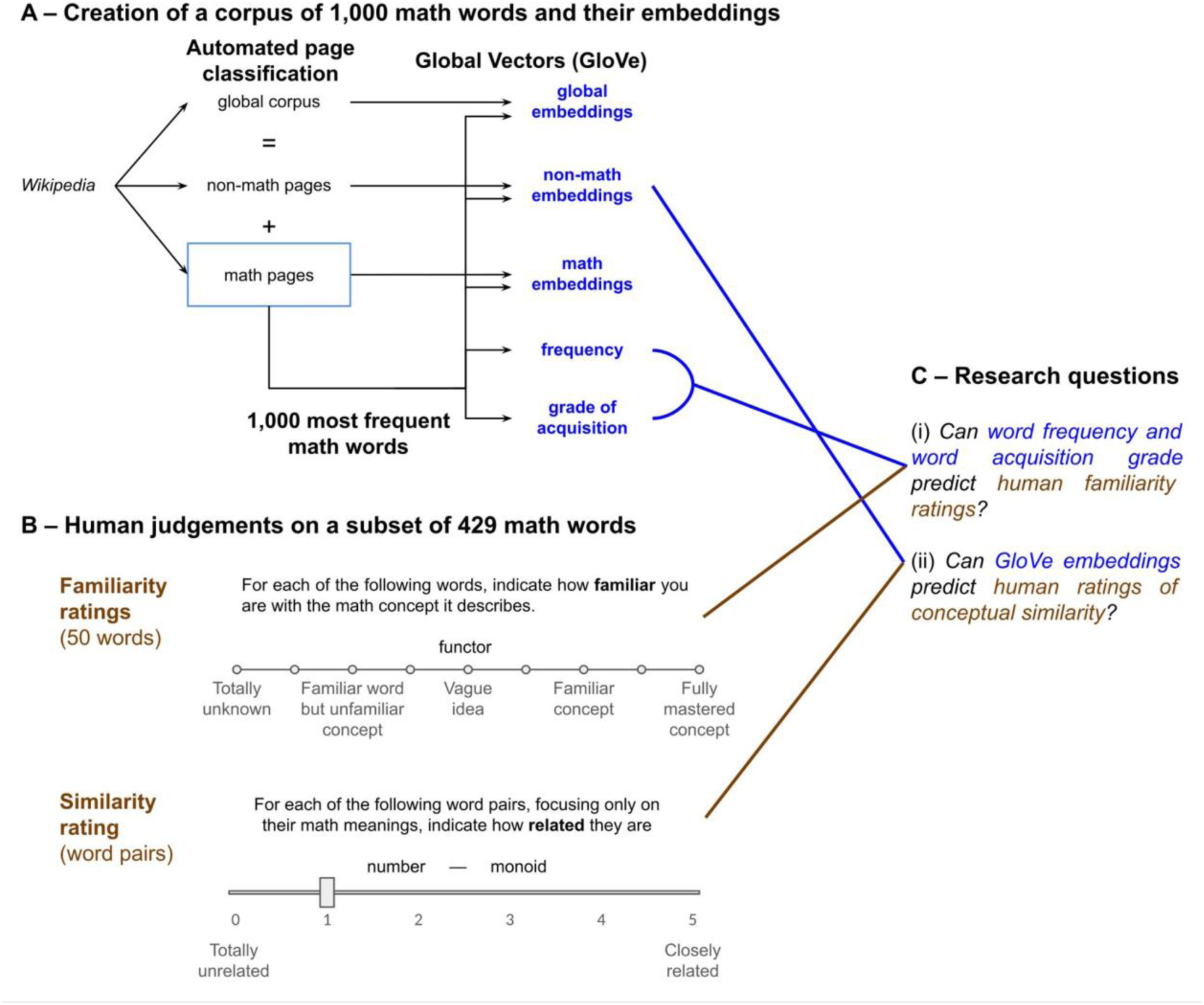
Experimental design. (A) Creation of corpora and embedding. (B) Behavioral experiment with a subset of words: familiarity and similarity ratings. (C) Relation to research questions.

**Table 1.** Summary of the main characteristics of the French vocabulary. Note: Frequency is expressed in Log10 units per million.

| Category |  | Count | Mean math freq | STD math freq | Mean non math freq | STD non math freq |
| --- | --- | --- | --- | --- | --- | --- |
| Word grade | *primary* | 214 | 1.90 | 0.63 | 1.24 | 0.91 |
|  | *6-7th grade* | 81 | 2.04 | 0.58 | 0.76 | 1.78 |
|  | *8-9th grade* | 52 | 1.89 | 0.70 | 0.44 | 1.13 |
|  | *10th grade* | 60 | 1.83 | 0.68 | 0.44 | 1.23 |
|  | *11-12th grade* | 110 | 1.80 | 0.51 | 0.31 | 1.06 |
|  | *bachelor* | 245 | 1.58 | 0.50 | 0.12 | 1.19 |
|  | *licence* | 121 | 1.38 | 0.40 | -1.08 | 2.84 |
|  | *master* | 55 | 1.40 | 0.43 | -1.61 | 3.85 |
| Grammatical category | *adjective* | 301 | 1.65 | 0.55 | -0.13 | 2.39 |
|  | *noun* | 534 | 1.82 | 0.62 | 0.45 | 1.88 |
|  | *verb* | 56 | 1.62 | 0.49 | 0.84 | 0.79 |
|  | *proper noun* | 126 | 1.39 | 0.38 | -0.52 | 1.35 |
|  | *adverb* | 10 | 1.45 | 0.42 | -0.16 | 1.01 |
|  | *number* | 24 | 1.83 | 0.85 | 1.91 | 0.77 |
|  | *symbol* | 9 | 1.87 | 1.00 | 0.91 | 1.31 |

### 2.2. Behavioral data collection

We ran a massive online experiment (n = 1230 participants) to collect familiarity ratings for a subset of 429 words of the vocabulary and similarity ratings for 3,756 pairs of these words (Fig 1B). The selected words were nouns, numbers or symbols which have a clear and precise mathematical meaning (see Methods). Our goal was to probe whether (1) the familiarity ratings from humans with different levels of education could be predicted by our estimated word grade and frequency in the math corpus; and (2) the similarity of GloVe embeddings is a good predictor of human similarity ratings (Fig 1C). All participants gave an estimate of their math education level, which ranged from primary school (n = 4) to PhD (n = 133) (see Methods). This experiment was also run in English for the English vocabulary (n = 174 participants).

Before anything else, participants answered a short survey and provided their last grade of education in mathematics and a self-assessment of their current math skills on a scale from 1 to 10. We found a moderate relation between these two variables, both in French (Spearman’s *r_S_*(1228) = .54, *p* < .001) and in English (Spearman’s *r_S_*(172) = .37, *p* < .001). For this reason, in the rest of this work, we used participants’ education as an indicator of their math level.

### 2.3. Familiarity ratings are predicted by the estimated grade of acquisition

Each participant first rated their familiarity with 50 words drawn randomly from the 429 selected for the experiment. Ratings were provided on a discrete 9-level Likert scale from 0 (totally unknown concept) to 8 (fully mastered concept) (see Fig 1 for the full calibration labels).

The results, shown in Fig 2, support two conclusions: (1) for a given participant’s math education level, familiarity ratings decrease as word grade increases; and (2) for a given word grade, familiarity ratings increase as participant education increases.

**Figure 2.**
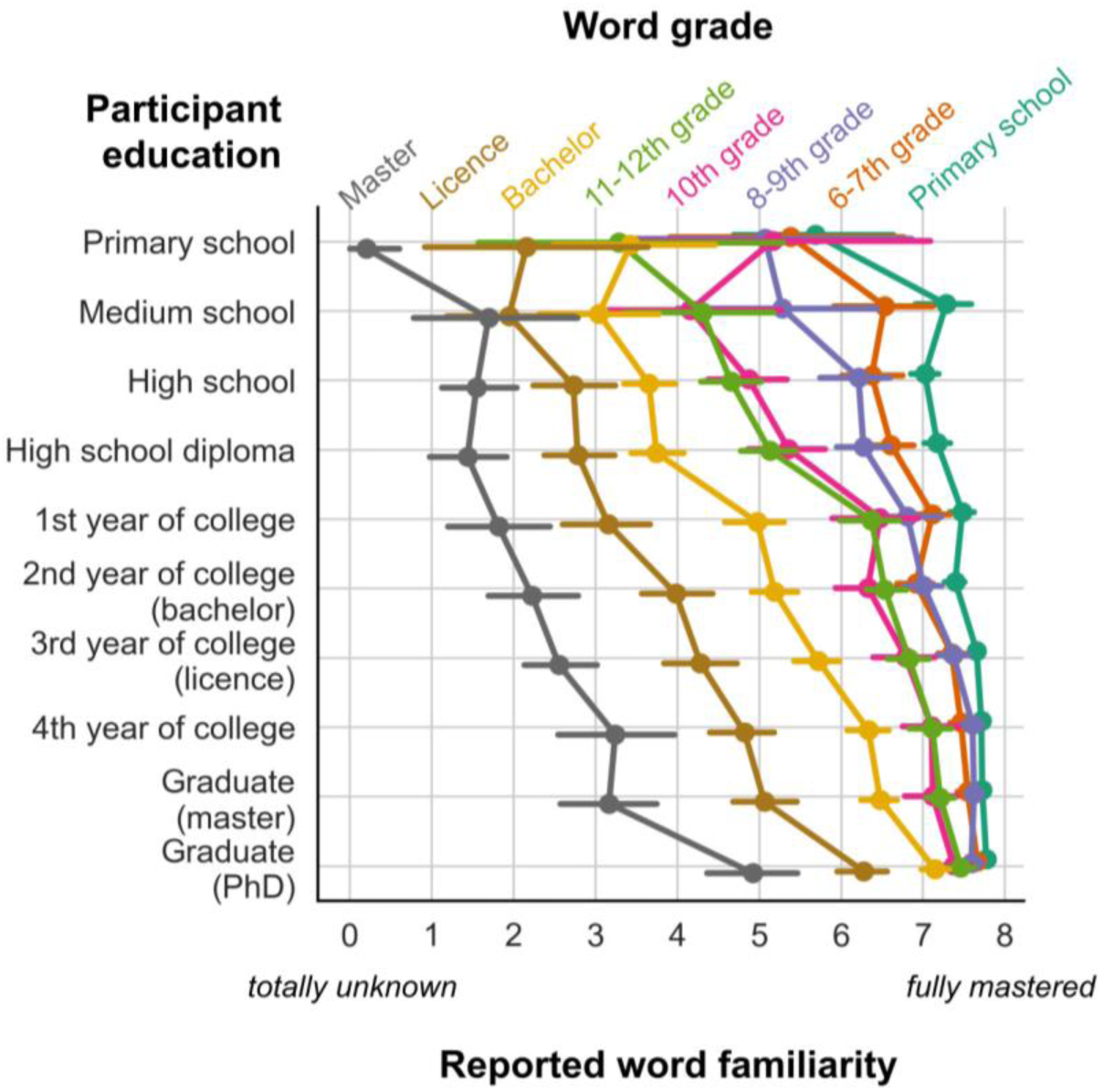
Familiarity ratings as a function of participant education and word grade. Note that, for greater readability, the dependent variable (rating of word familiarity) is on the x axis. Colors represent different categories of words, sorted according to the grade at which they are introduced in the French curriculum.

We confirmed those conclusions statistically by entering all of the participants’ familiarity ratings into a rank-order linear regression with participant education, word grade, word log frequency per million (in the math corpus), and their interactions as independent variables (see Methods). We found that these variables predicted familiarity ratings (*R^2^* = .35, *F*(7, 61792) = 4673, *p* < .001). All variables and their two-ways interactions were significant predictors (*p* < .001), and the three-ways interaction also was significant (*p* = .02). Out of all the predictors, word grade had the largest effect (*β* = -1.01, *t*(61792) = -116.42, *p* < .001): as seen on Fig 2, familiarity ratings decreased monotonically as the predicted word grade increased. This effect was modulated by a main effect of participant education (*β* = 0.67, *t*(61792) = 77.90, *p* < .001) indicating that familiarity ratings increased with education, and an interaction of education and word grade (*β* = 0.32, *t*(61792) = 36.33, *p* < .001), revealing that the effect of word grade was steeper in less educated participants. As can be seen on Fig 2, familiarity ratings became much flatter, although still increasing with education, when participants’ education fell above the postulated word grade. For instance, the familiarity rating of words assumed to be learned in 11-12th grade (green curve in Fig 2) reached a plateau around 7 once participants’ education reached or exceeded college level.

Finally, word frequency also had an effect on familiarity ratings (*β* = 0.33, *t*(61792) = 41.81, *p* < .001), but it was smaller than that of participant education and word grade. This effect indicated that, over and above word grade, the more frequent a word was in the math corpus, the more participants declared to be familiar with it. An interaction with word grade (*β* = 0.36, *t*(61792) = 43.53, *p* < .001) indicated that the effect of frequency on familiarity ratings was higher for more advanced math words, which is unsurprising because among the advanced words, the least frequent tend to be niche words, applicable only to a narrower domain of math, and therefore less familiar on average. There was also a small but significant interaction of frequency with participant education, indicating that higher education attenuated the impact of frequency (*β* = -0.05, *t*(61792) = -5.74, *p* < .001). This suggests that, while familiarity with math concepts is driven by frequency for participants with little math education, higher-level participants gained enough experience to mitigate the effect of frequency when judging their familiarity with a given concept: for them, even rare terms can be highly familiar.

These observations were replicated in English (*R^2^* = .28, *F*(7, 8792) = 498.7, *p* < .001), except for the interaction of word frequency and participant education which did not reach significance.

### 2.4. Item Response Theory

Mean familiarity ratings provide only a coarse idea of how advanced a math concept is. Likewise, a self-reported education level provides only a coarse estimate of a person’s math knowledge. Furthermore, those two parameters are linked: mean familiarity depends on the education level of the specific participants tested. We reasoned that item-response theory (IRT; Cai et al., 2016) could disentangle those parameters and provide a more refined estimation of them. IRT can jointly infer a latent ability for each participant, roughly capturing a person’s math knowledge, as well as a difficulty and a discrimination parameter for each word, evaluating respectively the overall likelihood that a word is judged familiar, and the amount of variation in this familiarity rating as a function of participants’ math knowledge.

Because the IRT package that we used (`mirt` in R) requires dichotomous outputs, we dichotomized the familiarity ratings into “unknown” (0, 1, 2 and 3 ratings) versus known (above 3). We reviewed manually all predictions made by the IRT algorithm and discarded 20% of all items for which the algorithm had extreme values of discrimination or difficulty parameters which did not fit with the data. Indeed, IRT cannot converge when human ratings are very skewed toward one value (either always known, or always unknown). Manual review of the data led us to only keep the first nine deciles of IRT discrimination parameters and the central 95% of IRT difficulty parameters (2.5th to 97.5th percentiles) for further analyses. After this curation phase, we were left with 289 words out of our original 363 for which the IRT algorithm converged. The resulting estimates are reported in the vocabulary database, and plots are provided in File S1.

For each word tested, we obtained two difficulty estimates, based either on mean familiarity rating or on IRT, and two discrimination estimates: the standard deviation (STD) of familiarity ratings across participants and the IRT discrimination estimate. We then compared the two approaches and correlated the descriptive parameters with those derived by the IRT algorithm. We found that IRT difficulty parameter and mean familiarity rating were strongly correlated across words (Spearman’s *r_S_*(287) = .90, *p* < .001). IRT difficulty parameter was also correlated with word grade (Spearman’s *r_S_*(287) = .51, p < .001) and, to a lesser extent, to word frequency (Spearman’s *r_S_*(287) = .14, p < .001). Likewise, the IRT-estimated participant latent ability and the reported participant education were highly correlated across participants (Spearman’s *r_S_*(1228) = .61, *p* < .001). However, the IRT discrimination parameter was not well estimated by the STD of familiarity ratings (*p* = .57), suggesting that IRT indeed provided a finer-grained approach.

We used the IRT results to manually select a subset of 80 words covering various difficulty levels, whose discriminability was high, and which were therefore highly selective of a given participant’s math knowledge (they are flagged in the database as “80 high-discriminability items”). In the future, we propose that collecting familiarity ratings on those words and analyzing them with IRT could serve as a quick assessment of a participant’s math knowledge. Similarly, the word difficulty and discrimination parameters that we collected could be used to select, within our vocabulary, a subsample of words adequate to participants of a given education range.

### 2.5. GloVe embeddings predict human math similarity ratings

We then turned to the similarity rating part of the experiment. Note that, to avoid asking ratings for unknown words, each participant only rated on a continuous Likert scale the similarity of word pairs at or below their grade level. Thus, more educated participants were asked to rate a large number of word pairs (20 pairs per grade level). Furthermore, for each word, in order to cover a large spectrum of similarities, we selected 3 pairs within each of four categories: (1) very similar pairs, as predicted by their GloVe embeddings; (2) moderately similar pairs, close to the mean similarity; (3) pairs of words with orthogonal embeddings; (4) pairs of words with opposite embeddings, i.e. negative cosine similarity (see Methods). This manipulation was adopted after piloting showed that a random choice of word pairs would have generated an excessive number of “unrelated” judgements, thus, making the task uninteresting for participants. Furthermore, it allowed for a first test of GloVe similarity: non-parametric Kruskal-Wallis (*H*(3, n = 3756) = 1520.11, *p* < .001) followed by pairwise Dunn’s tests with Bonferroni correction showed that similarity ratings showed systematic pairwise differences across those four categories (mean ratings = 3.2, 1.9, 1.2 and 1.0 respectively; all *p*’s < .001). In particular, even the negative cosine pairs were rated as slightly, but significantly less similar than orthogonal items. These observations were also replicated in English except for the negative cosine versus orthogonal comparison (mean ratings = 2.9, 1.9, 1.5 and 1.4 respectively; non-parametric Kruskal-Wallis: *H*(3, n = 2421) = 531.93, *p* < .001; Dunn’s test: all *p*’s < .001 except for the negative cosine versus orthogonal categories).

The next question we asked was whether human similarity ratings could be continuously predicted by the similarity of their GloVe embeddings. Following Pereira et al. (Pereira et al., 2016), we tested both euclidean distance and cosine similarity, which are respectively defined as

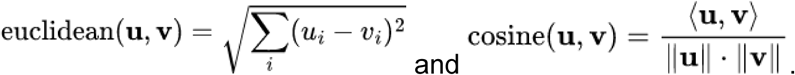

In all following work, as a measure of fit, we computed Spearman’s rank-order correlation *rS* between the similarity of 50-dimensional GloVe embeddings and the mean human similarity rating for each pair of words. Rank-order correlation seemed more appropriate, given that the Likert scale used to collect similarity ratings is not necessarily linear (see below).

When computing the correlation for word pairs aggregated over percentile bins of predicted similarity (Fig 3A), we found a strong correlation for both the cosine similarity and the euclidean distance (euclidean: Spearman’s *r_S_*(3754) = -0.56, *p* < .001; cosine: Spearman’s *r_S_*(3754) = .66, *p* < .001). Note that the correlation was positive for the cosine similarity while it was negative for the euclidean distance as the latter is a measure of dissimilarity, so it should decrease as the similarity ratings increase. The correlation was slightly better for cosine, so it is the measure we used for subsequent analyses (ΔAIC = 734.31, *p* < .001). However, similarly to what was observed by Pereira et al. (Pereira et al., 2016), the difference between the cosine similarity and euclidean distance was small.

**Figure 3.**
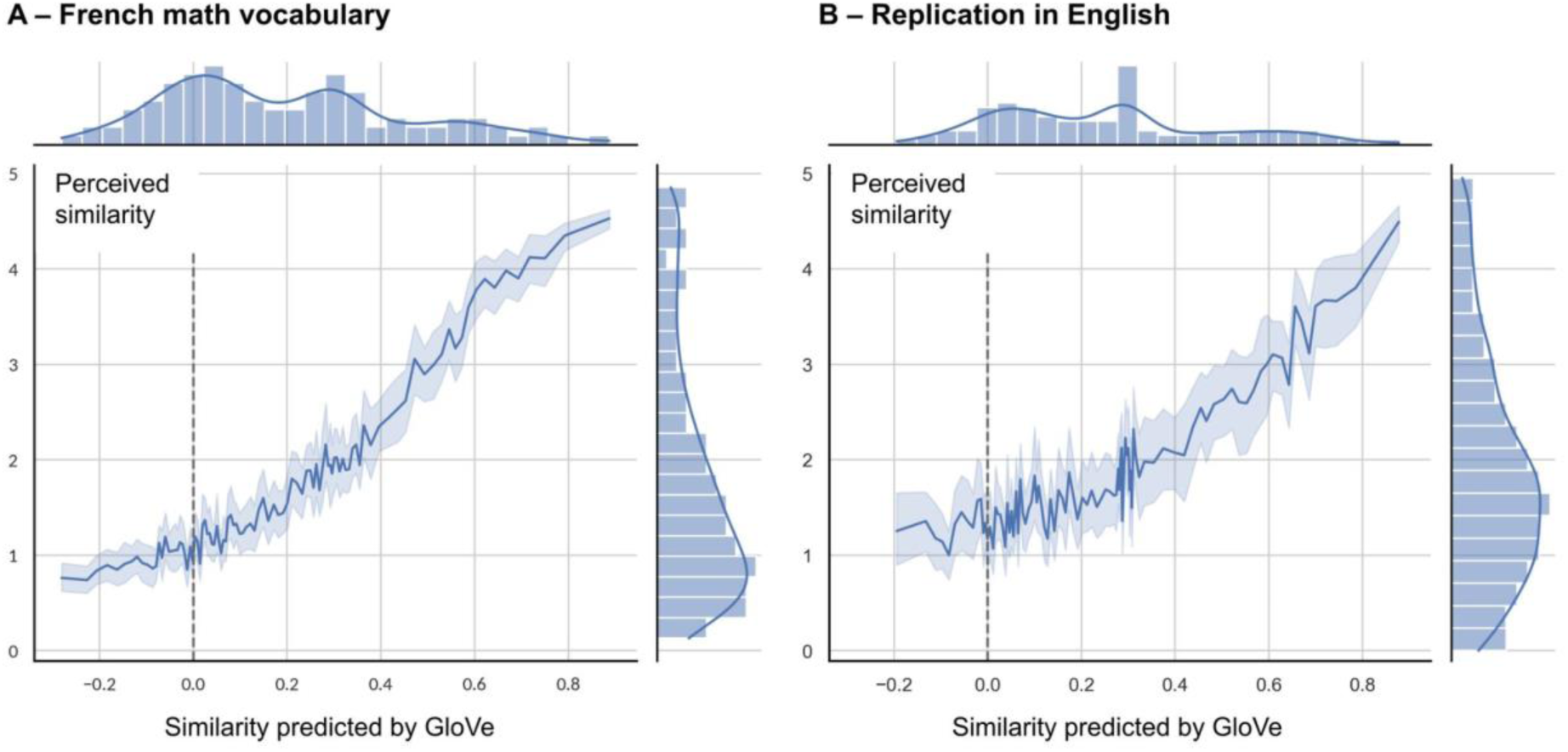
The perceived similarity between two math concepts is well predicted by the similarity of their GloVe 50-dimensional embeddings. The plot shows human similarity rating (on a scale from 0 to 5, y axis) as a function of their predicted similarity based on the cosine of the angle between their 50-dimensional vector embeddings derived from the math corpus (x axis), (A) in French (n = 3756); and (B) in English (n = 2421). Data were averaged over percentile bins. Marginal distributions of the x and y variables are also shown.

The results were also replicated in English (Fig 3B). Again, the correlation for cosine similarity was slightly better than that for euclidean distance (euclidean: Spearman’s *r_S_*(2419) = -0.42, *p* < .001; cosine: Spearman’s *r_S_*(2419) = .48, *p* < .001; ΔAIC = 174.92, *p* < .001).

An interesting observation, visible in Fig 3, is that human similarity ratings were not linearly related to GloVe cosine similarity. Indeed the curves for French and English were both convex, meaning that, for small values of similarities predicted by GloVe (between 0 and 0.4), human ratings varied little with GloVe similarities (though still significantly: Spearman’s *r_S_*(2061) = .37, *p* < .001), whereas for larger GloVe cosine similarities, they changed in a much steeper fashion.

We compared those GloVe fits with an estimate of the noise ceiling in our data (see Methods), in order to get an idea of how good those fits were relative to the explainable variance. To this end, we recomputed the correlations on the unaggregated, trial-by-trial data, without averaging the ratings for each pair of words across participants. This noise ceiling can be thought of as the fraction of variance in the data from one participant that could be accounted for by all the other participants (leave-one-out cross validation). We found that (1) the single-trial similarity ratings were highly reliable, exhibiting an explainable variance (*R²*) or noise ceiling of 42.84% across participants; (2) the cosine similarity of 50-dimensional GloVe embeddings, on the other hand, led to a single-trial correlation of 20.28%, i.e. about half of the explainable variance. Thus, while GloVe provides a first-approximation model of math concepts, it still leaves much to be explained.

### 2.6. Comparing different embeddings for math concepts

We examined the impact of the dimensionality of GloVe embeddings (Fig 4A). The above analyses were based on 50-dimensional embeddings, which is GloVe’s default, but we wondered whether increasing the dimensionality of the embeddings would also increase the correlation of their cosine similarities with human ratings.

**Figure 4.**
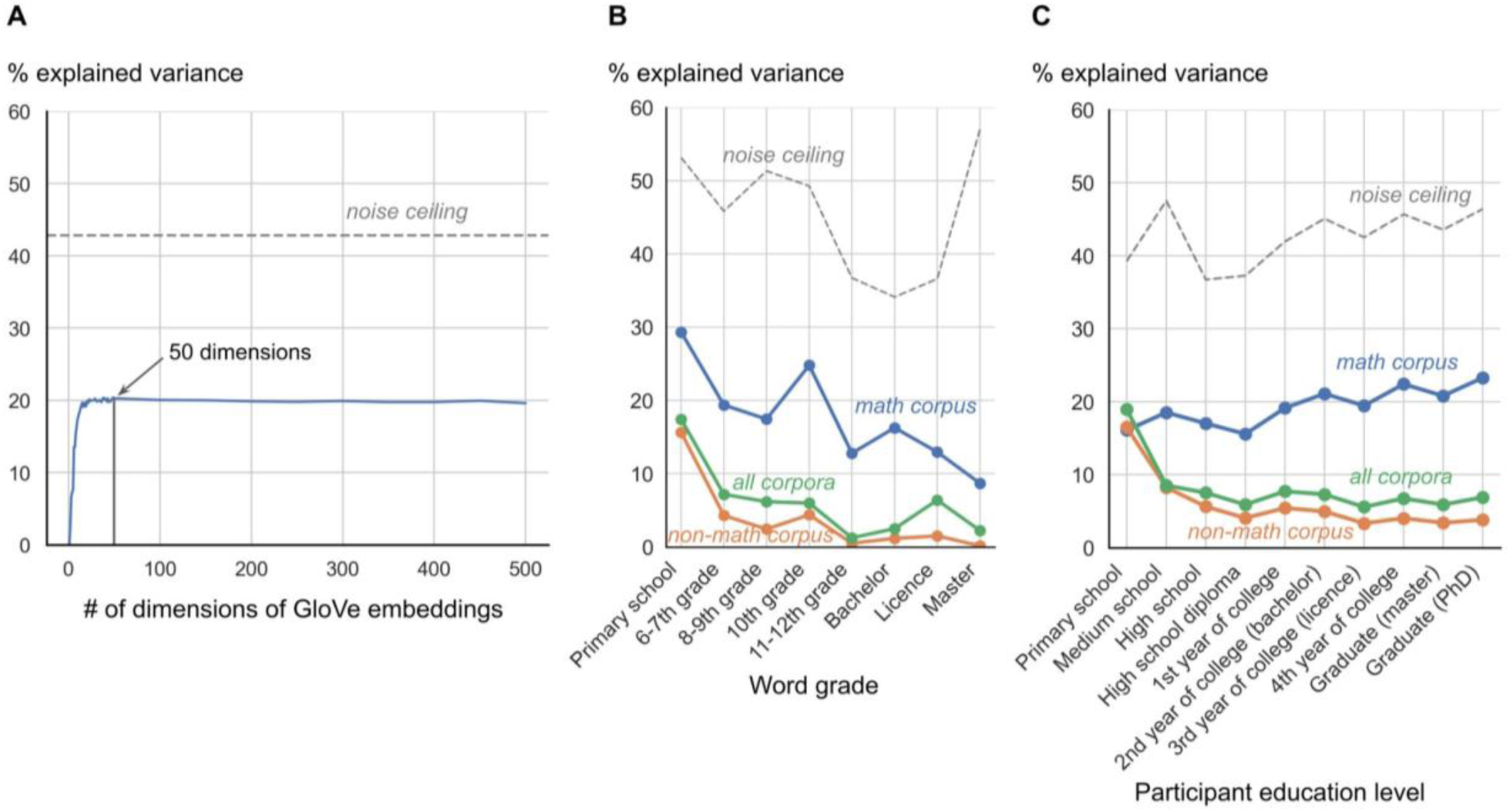
Factors affecting the correlation between human similarity ratings and the similarities predicted by GloVe embeddings. (A) Percentage of variance explained by a rank correlation of human similarity ratings against the cosine angle of GloVe math embeddings, as a function of the dimensionality of those embeddings. The dashed line shows the noise ceiling, estimated as the fraction of variance in the data from one participant that could be accounted for by all the other participants (leave-one-out cross validation, see Methods). 50-dimensional embeddings were used in the rest of this work. (B) Variation in the percentage of explained variance depending on word grade of acquisition and the corpus used to compute the embeddings; (C) Same, depending on participant education level.

The results showed that, for French words, the Spearman correlation coefficient increased with the number of dimensions of GloVe embeddings up to approximately 50 dimensions, with a sharp initial increase for dimensionality 1 to ∼15, and then reached a plateau for 50 to 500-dimensional embeddings. Therefore, the choice of 50-dimensional embeddings was felicitous. We observed the same trend for English words, although both the noise ceiling and the correlation coefficients were lower.

We also probed whether embeddings derived from the math corpus were better at predicting human similarity ratings than those derived from the non-math and global corpora. We therefore computed the rank correlation between the similarity ratings and the cosine similarity of the 50-dimensional embeddings derived from the math, non-math and global corpora. We indeed found that the math corpus was a better predictor, followed by the global and the non-math corpora (math: Spearman’s *r_S_*(97593) = .45, *p* < .001; global: Spearman’s *r_S_*(97593) = .25, *p* < .001; non-math: Spearman’s *r_S_*(97593) = .20, *p* < .001; ΔAIC_math-global_ = 15792.95, *p* < .001; ΔAIC_math-nonmath_ = 18264.42, *p* < .001). These findings were replicated in English (math: Spearman’s *r_S_*(13611) = .35, *p* < .001; global: Spearman’s *r_S_*(13611) = .21, *p* < .001; non-math: Spearman’s *r_S_*_(_13611) = .19, *p* < .001; ΔAIC_math-global_ = 1049.85, *p* < .001; ΔAIC_math-nonmath_ = 1347.46, *p* < .001).

To get a finer-grained view of the differences between the different corpora, we repeated the same analysis for each word grade of acquisition. We expected that (1) basic concepts might be modeled equally well by all corpora, but (2) as word grade increases, embeddings derived from the math corpus would become better predictors of human similarity ratings while correlations with embeddings derived from the other corpora would drop. These predictions were partly confirmed (Fig 4B): the correlation of the cosine of embeddings derived from the non-math corpus with human ratings decreased as the word grade increased (Spearman’s *r_S_*(6) = -0.83, *p* = .010), but the Spearman correlation between cosines from global corpus and word grade was not significant. Furthermore, embeddings derived from the non-math and global corpora were consistently poorer predictors of human ratings than those derived from the math corpus (exact one-sided Wilcoxon signed-rank test: math vs non-math: *W* = 36, *p* = .004; math vs global: *W* = 36, *p* = .004), even for very basic math concepts studied in primary school. In addition, the correlation between the cosine of math embeddings and human ratings also decreased as the word grade increased (Spearman’s *r_S_*(6) = -0.87, *p* = .007), probably due to the fact that advanced concepts were only rated by a smaller number of highly educated participants and, as a consequence, the estimation of their human similarity rating might be noisy.

To counter this possibility, we also analyzed the quality of GloVe fits as a function of education level. We hypothesized that the math corpus would be a better predictor of expert mathematicians than of people with a lower level of math education. Indeed, the math corpus comprised very advanced math content such as category theory or algebraic topology. Conversely, the non-math corpus should well predict the ratings of mathematically uneducated people, but not those of more advanced mathematicians, as the math words in the non-math corpus were either very low level (e.g. fractions or basic shapes) or advanced concepts which have several meanings outside the math domain (e.g. “ring” or “field”). We ran the same fine-grain analysis as for word grade (Fig 4C) and found that these predictions were verified: all corpora predicted ratings of non-educated participants equally well, and then diverged for educated participants. The correlation for the math corpus increased with participants’ level of education (Spearman’s *r_S_*(8) = .85, *p* = .002), while that for the non-math and global corpora decreased as participants’ level of education increased (global: Spearman’s *r_S_*(8) = -0.65, *p* = .043; non-math: Spearman’s *r_S_*(8) = -0.88, *p* < .001).

### 2.7. Visualizations of the semantic space of math concepts

Given the relatively good fit of the GloVe vectors to human judgments, we close our work with an analysis of the geometry of those vectors, as a proxy for human representations of the semantic space of math concepts (in the future, this model could be compared to actual measurements using fMRI or MEG; Kriegeskorte, Mur, & Bandettini, 2008; Kriegeskorte, Mur, Ruff, et al., 2008).

We first tried to obtain a global visualization of the semantic space of math concepts. To this end, we created a 2D map following the procedure in (Pereira et al., 2018). First, we separated the 1,000 vectors in 18 clusters using spectral clustering (von Luxburg, 2007). Then we projected all vectors along with the cluster centers in two dimensions using t-SNE (van der Maaten & Hinton, 2008). Finally, we used Voronoi tessellation around the projected centers to visualize the boundaries between the clusters. The number of clusters (18) was chosen out of a range from 2 to 100 using the elbow method. The resulting map is shown on Fig 5. We were able to label all clusters with a tentative semantic description. Different regions were dedicated to analysis, algebra, arithmetic, geometry, and, more tentatively, for arithmetics and to linear algebra. Thus, numbers had a dedicated cluster (Fig 5B). Likewise, an entire area of the map was dedicated to proper nouns, which were separate from the rest of the concepts (for instance, “Burnside” was closer to “Weierstrass” than they were respectively to other group theory and analysis concepts, see Fig 5B). However, some proper nouns, especially those that can be used as adjectives were also sometimes integrated to the cluster related to their field (e.g. “Poisson” is located in a probability cluster, as shown on Fig 5B).

**Figure 5.**
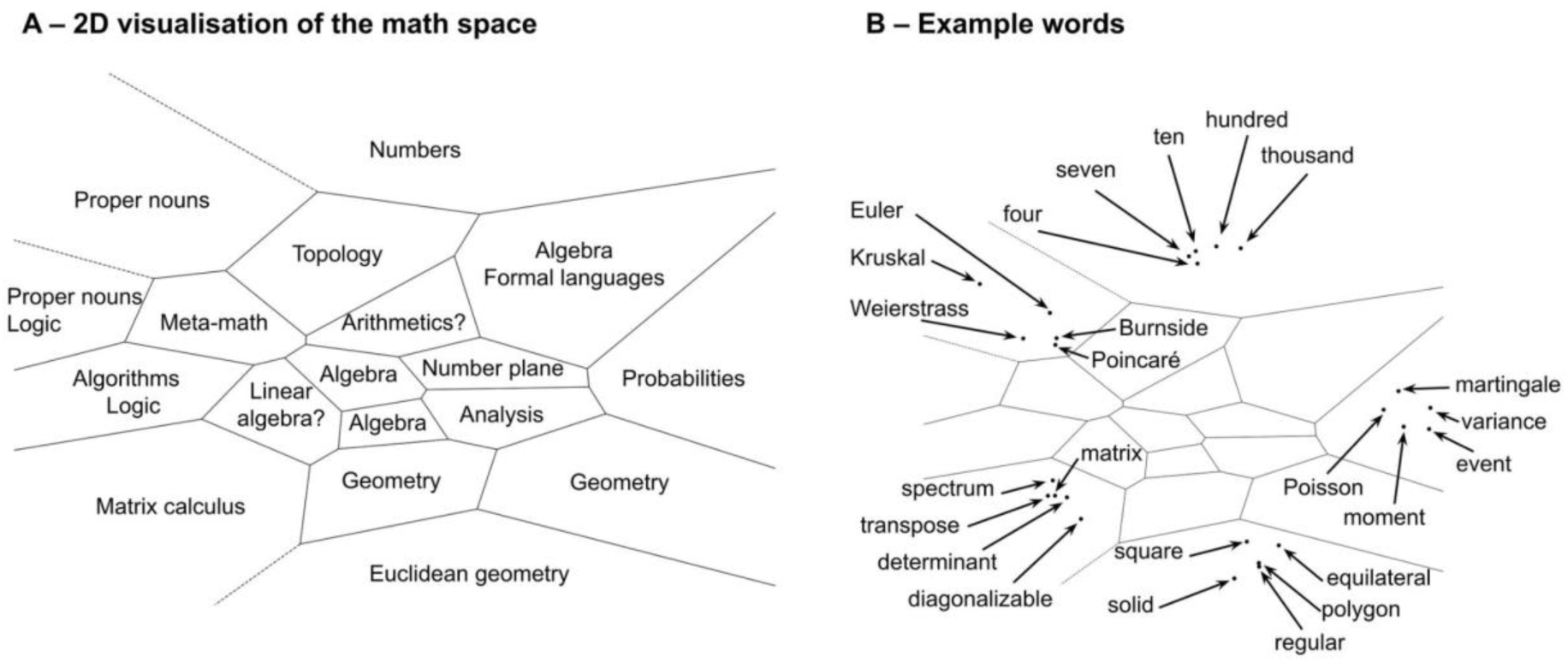
Global visualizations of the vector space for math semantics. We used the GloVe embeddings of the French math vocabulary to visualize the organization of the putative semantic space by projecting it in two dimensions using spectral clustering and t-SNE. (A) Each region was associated with a tentative label describing the words contained in the cluster. (B) We also show five words contained in different regions: numbers, proper nouns, matrix calculus, euclidean geometry and probabilities.

Interestingly, all terms relating to geometry occupied several clusters in a large and distinct sector of the map (bottom right-hand corner), very distant to those dedicated to numbers (top right-hand corner) or logic (left-hand corner) (Fig 5A). This observation parallels Amalric and Dehaene’s (2016, 2019) brain-imaging finding that, when judging the truth of math sentences, partially distinct cortical sites were found for sentences bearing on geometry relative to other domains such as algebra, arithmetic, or topology, within an otherwise highly integrated and overlapping cortical network for all math concepts. Similar to GloVe, the human brain may group together words that frequently cooccur, thus grouping together terms of geometry because they involve a partially distinct language of shapes with a strong visuospatial content (Amalric et al., 2017; Dehaene et al., 2022a; Sablé-Meyer et al., 2022).

It is worth noting that the map continued to make sense when we increased the number of clusters. Indeed, we reproduced this methodology with 100 clusters, and found that the above mentioned clusters were refined in an interesting way. For instance, proper nouns were also classified into sub-clusters depending on the area of math that they relate to (e.g. “Fermat” and “Markov” fell in different clusters, as the former worked on number theory and the later worked on probability). Similarly, the number cluster was split into smaller adjacent clusters distinguishing between and small numbers, multiples of ten and powers of ten.

We then turned to more local visualizations of the semantic space to get a better idea of the organizations of specific domains. The first things we looked into were numbers. We projected the embeddings of numbers from 1 to 100 (only those whose French name only consists of one word, which excluded for instance 70 or 21) on their first principal component (PC1) (Fig 6A). We found that this projection ordered numbers by their magnitude. Furthermore, larger numbers were grouped together, suggesting a logarithmic organization (correlation with log(n): Pearson’s *r*(20) = .90, *p* < .001). Following earlier work indicating that high-dimensional embeddings can be projected onto oriented axes for properties such size or ferocity (Grand et al., 2022), we also looked at the line joining the embeddings of “one” to “billion” and projected the embeddings of the other numbers on this axis. Indeed, this projection revealed a systematic organization of other numbers based on their relative size (Fig 6B). Again, numbers were roughly organized by magnitude, in a compressive manner. Overall, these findings indicate that GloVe recovers a compressive, quasi-logarithmic representation of numerical magnitude similar to the one that behavioral and neuroscience research has identified as lying at the core of intuitions of number in both humans and animals (Dehaene, 2003, 2011; Dehaene & Marques, 2002; Kutter et al., 2018; Nieder, 2021; Piazza et al., 2004; Siegler & Opfer, 2003). The fact that even very large numbers such as “hundred”, “thousand”, “million” or “billion” were placed appropriately in the same alignment fits with developmental research indicating that 6-year-old children can already tell which of two such numbers is larger (Cheung & Ansari, 2023), and suggests that statistical word distributional properties may suffice to develop such an extension of magnitude knowledge to very large numbers.

**Figure 6.**
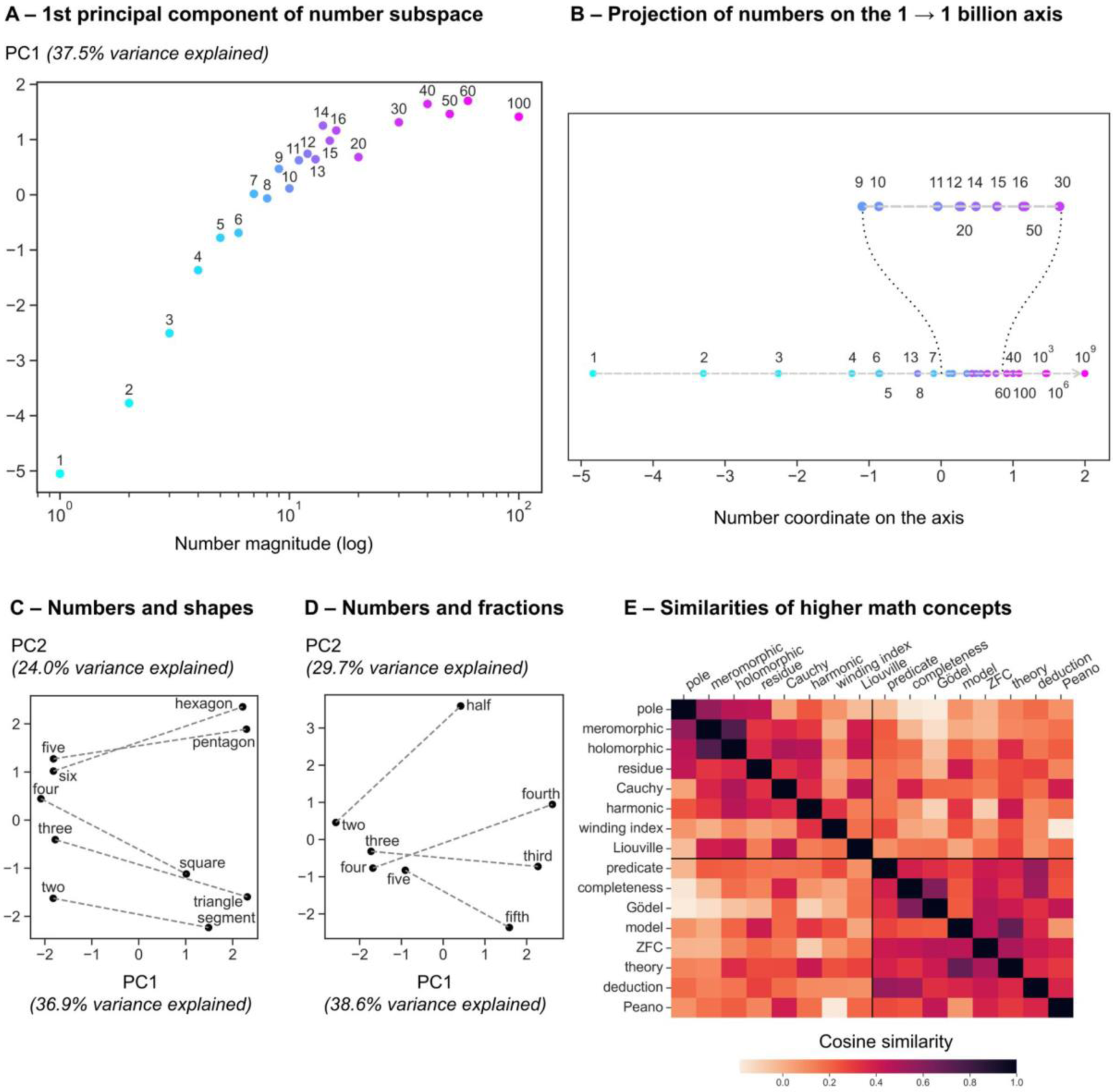
Partial visualizations of the vector space for math semantics. We used the GloVe embeddings of the French math vocabulary to visualize the organization of specific concepts in this putative semantic space. (A-B) Projection of the vector embeddings of numbers from 1 to 100 on their first principal component (PC1) (A), or on the line joining the two vectors for “one” and for “billion” (B). In panel A, the x axis is the log of the number magnitude in base 10, which shows a tight though non-linear correspondence with PC1. (C-D) Projection of the embeddings of two groups of related concepts on their first two principal components: numbers and shapes (C), and numbers and fractions (D). (E) Cosine similarity matrix for two domains of advanced math: complex analysis and logic (8 words each).

We then looked into the association between numbers and other interrelated math concepts. GloVe is known to capture the meaning specified by the juxtaposition of two words (e.g. the difference between “king” and “queen” is the same as that between “man” and “woman”) (Mikolov et al., 2013). We first focused on the correspondence between numbers and shapes (e.g. “three” and “triangle”, “four” and “square”; Fig 6C). In Fig 6C-D, we projected the embeddings on their first two principal components and drew a line to connect related concepts. The vectors for numbers and shapes were indeed systematically related (Fig 6C): In GloVe embeddings, a similar vector is needed to go from “three” to “triangle” as to go from “four” to “square”. In addition, PC1 made a clear distinction between numbers and shapes, while PC2 ordered numbers by their magnitude and shapes by their number of sides.

A similar analysis for numbers and fractions (e.g. “two” and “half”, “three” and “third”, etc) revealed different results (Fig 6D). Note that, contrary to English, the French for the fractions ⅓ (“tiers”) and ¼ (“quart”) are different from those for the ordinals 3rd (“troisième”) and 4th (“quatrième”). On Fig 6D, we see that the correspondence was not well captured by GloVe. PC1 still made a clear distinction between numbers and fractions, but along PC2, numbers were grouped together (still ordered by magnitude), while fractions were separated from each other. More work will be required to examine if a systematic but non-linear relationship between numbers and their corresponding fractions can be found using more sophisticated methods such as tensor decomposition (McCoy et al., 2019).

Finally, we wanted to probe whether higher-level math concepts were also accurately represented in the math semantic space computed by GloVe. We selected sixteen concepts from distinct, advanced math fields, namely complex analysis and logic, and asked whether the concepts from the same domain were consistently more similar than concepts from different domains. The correlation matrix between the sixteen concepts is shown on Fig 6E. We found that the intra-domain GloVe similarity was indeed greater than the inter-domain similarity (intra: μ = 0.34, σ = 0.17; inter: μ =0.13, σ = 0.14; *t*(118) = 7.34, *p* < .001).

## 3. Discussion

We created a comprehensive 1,000-word vocabulary of math concepts in French, covering basic and advanced levels and provided vector embeddings for words in this vocabulary. We then validated the embeddings by showing that they explain up to 47% of the explainable variance from human similarity ratings. In addition, we proved that education affects both familiarity and similarity ratings: the higher the math education level, the more accurate the prediction of similarity ratings by GloVe embeddings extracted from Wikipedia math pages. We also showed that the spatial layout of the vectors makes sense and captures important regularities in number concepts, geometric words, and other higher-level concepts. Importantly, we provided a translation of the vocabulary into English and proved that the main results were replicated in both languages.

### 3.1. Possible use of the vocabulary and vectors

The purpose of this database is to enable a more systematic exploration of math cognition, beyond the elementary concepts of numbers, geometry and algebra on which the vast majority of current cognitive neuroscience research is concentrated. Our approach may help provide a standardized approach to this field, cover the mathematical domain in an unbiased manner, and increase the comparability between different studies. Indeed, it could be desirable that studies tackling the cognition of advanced math all share a common subset of stimuli. This would be useful not only for the purpose of reducing bias inherent in the choice of stimuli, but also to join efforts to bring multiple converging data to bear on the same problem (e.g. behavioral, developmental, brain-imaging, intracranial recordings…), in a similar fashion to the *THINGS* initiative (Hebart et al., 2019). We believe that the dataset we propose would be useful in this regard, as we endeavored to make it exhaustive and unbiased.

Our dataset may also be used to devise benchmarks, test sets, and stimuli – for instance, we provide a reduced set of 80 high-discriminability items that may suffice to quickly determine a participant’s level of math knowledge. Following a method similar to Pereira et al. (Pereira et al., 2018), one could use the embeddings we provide to ensure a full coverage of the entire math semantic space.

### 3.2. Limitations of this work

The first limitation of this work concerns the vocabulary itself. A subset of mathematical words were reviewed and manually selected. The threshold of 1,000 words was arbitrary, and rare words could have been missed, although we tried to improve the vocabulary at different stages in the process. The grade level we assigned is necessarily only approximate and, since it was based on the French national curriculum, could be different in other countries. Furthermore, the very notion of a grade level may not be appropriate for ambiguous words (e.g. “order”) or very frequent words (e.g. “line”), as the depth of their understanding and, indeed, their very meaning evolves with education. Furthermore, our focus on single words may not do full justice to the combinatorial nature of mathematical concepts. To take just one example, the concept of “vector space”, being expressed by two words, was not represented in the present embeddings. In spite of these limits, the fact that we observed meaningful variations of both familiarity and similarity with grade and education indicates that the present work provides a useful first approximation to an exhaustive dictionary of the most frequent math concepts.

The second limitation concerns the embeddings. As explained above, GloVe vectors only accounted for 43% of the noise ceiling of participants’ similarity ratings. Further work will be needed to increase the percentage of explained variance. One possible solution could be to use deep neural networks instead of embeddings derived from single-word cooccurence statistics. In recent years, Transformer models (Vaswani et al., 2017) have drawn a lot of attention and have been shown to be able to predict brain activations in fMRI studies (Caucheteux & King, 2022; Pasquiou et al., 2022; Schrimpf et al., 2021), although their capacity to capture even elementary mathematical knowledge remains highly debated, and specialized math models may be required (Anand et al., 2024; Peng et al., 2021). Further investigation is needed in this direction. Another option would also be to obtain distributed semantic representations from a larger corpus. Indeed, math articles are only a small fraction of Wikipedia, and adding more content to the corpus (Bourbaki textbooks for instance) may be beneficial to the estimation of the embeddings, at least for mathematically advanced participants. Language-of-thought approaches to mathematics (Dehaene et al., 2022b; Goodman et al., 2014; Piantadosi et al., 2012; Sablé-Meyer et al., 2022), which focus on the hierarchical compositional nature of mathematical concepts, may also provide a more appropriate cognitive foundation for the construction of higher level concepts in the course of education.

Regarding the visualizations of the semantic space of GloVe vectors, the 2-dimensional views that we obtained with t-SNE or PCA should only be taken as indicative, as the projections of a high-dimensional space may vary with the tool used, the subset of words under consideration, or even the random seed inputted to the projection algorithms. It should always be remembered that the full data lives in high dimensions. Finally, it must also be noted that our behavioral study of conceptual familiarity and similarity focused only on nouns. The psychological validity of the GloVe representations obtained for verbs, adjectives or proper names remains to be evaluated.

### 3.3. Future directions

We provided a comparison between human similarity ratings and GloVe embeddings. A natural continuation would be to compare the present embeddings and similarity ratings to vectors obtained from the human brain using brain imaging or intracranial recording techniques. We would then be able to leverage representational similarity analysis (Kriegeskorte, Mur, & Bandettini, 2008) tools to gain insight in the organization of math concepts in the brain, including their topographic organization on the cortical surface of individual subjects, going beyond existing group fMRI studies of mathematicians (Amalric & Dehaene, 2016, 2019). Ultimately, this could lead to the creation of a math-specific brain viewer, similar to that proposed by Huth et al. (2016).

## 4. Material and methods

### 4.1. Creation of the corpora

The corpora were extracted from HuggingFace’s Wikipedia `20220301.fr` dataset. In order to divide the dataset into a math and a non-math corpus, we parsed all the pages and located them in the `wikipedia_fr_all_maxi_2022-04.zim` dump. A bot then decided whether each page was math or not. To do so, it reached the bottom of the page and searched for an occurence of one of the following strings: “Portail des mathématiques”; “Portail de la géométrie”; “Portail de l’analyse”; “Portail de l’algèbre”; “Portail des probabilités et de la statistique”; “Arithmétique et théorie des nombres”; “Portail de la logique”; “Portail de l’informatique théorique”. These labels indicate that a page belongs to the portal of mathematics or theoretical computer science.

The math (resp. non-math) pages were aggregated to form the math (resp. non-math) corpus, and the two corpora were also merged to form the global corpus.

In total, the math corpus comprised 16,455 pages out of the 2,402,095 included in the dataset, and the non-math corpus comprised 2,236,840 pages (the remaining pages were redirections and disambiguation articles).

### 4.2. Extraction of the vocabulary and generation of the embeddings

In order to ensure that the vocabulary did not contain several occurrences of the same word (e.g. singular and plural for a noun, or infinitive and conjugated forms for a verb), we performed a lemmatization step on each corpus (math, non-math and global). Two lemmatization passes were carried out using Python `spacy`’s `fr_core_news_md` model. The lemmatized math corpus was then provided as an input to the GloVe pipeline (Pennington et al., 2014). The pipeline has three steps: it first creates a 50,000-word vocabulary sorted by decreasing frequency, then builds a cooccurrence matrix on the vocabulary words in the corpus and finally runs GloVe on this matrix. Roughly, GloVe uses the cooccurrence matrix of a vocabulary in a corpus to derive semantic vectors for each word by taking into account ratios of cooccurrences. The window size for cooccurrence was set to 15 words. As reported in the main text, the number of dimensions of the embedding vectors was varied from 1 to 500 (all values between 1 and 50, and from 50 to 500 with an increment of 50).

The vocabulary output by GloVe on the math corpus actually contained only 39,345 words, because the GloVe pipeline did not count words with fewer than five occurrences in the corpus. We then applied the same procedure to the non-math and global corpora, while imposing the vocabulary to correspond to the output from the math corpus.

Words of the vocabulary were reviewed manually in decreasing order of frequency by a person with extensive math training. Only the 1,000 first words which belonged to the math domain (whether elementary or advanced) were kept, thereby constituting the final thousand-word math vocabulary.

The vocabulary was then complemented with word frequency in the math and non-math corpora. We also estimated words’ school grade of acquisition (“word grade”) by examining when the corresponding concept was introduced in the French national curricula (the following levels were distinguished: primary school, 6-7th grade, 8-9th grade, 10th grade, 11-12th grade, bachelor, *licence*, master), grammatical category (number, symbol, noun, name, adverb, adjective, verb or any combination of these when applicable) and meta-information such as whether the words are meta-mathematical meta-math (e.g. “theory” or “example”) and whether they have several different math meanings (polysemy, e.g. “tangent”).

### 4.3. Familiarity ratings and similarity judgements

#### 4.3.1. Selection of the pairs

For the behavioral experiment, we only kept those numbers, nouns and symbols which were not flagged as too polysemic or meta-math, leaving us with 429 words. As these numbers would have yielded 91,378 different pairs, we did not attempt to measure their full similarity matrix. Furthermore, a random sampling of this large matrix would have yielded a vast majority of unrelated words, thus making the experiment quite monotonous for participants. Instead, to select target pairs for a given participant, we computed the distribution of cosine similarities between pairs of words with the same grade of acquisition using 50-dimensional GloVe vectors obtained from the math corpus. For each of the 429 target words, we identified the three furthest words (cosine similarity close to -1), closest words (cosine similarity close to 1), most orthogonal words (cosine similarity close to 0) and words whose similarity was closest to the average similarity between words of the same grade (n = 14,790, μ = 0.24, σ = 0.20). With this procedure, we obtained a total of 3,756 pairs that we used for the similarity rating experiment (see below, and Table S2).

#### 4.3.2. Design of the online experiment

The experiment was coded in PHP (server side) and JavaScript (client side) using the jsPsych library. It was hosted on the secure server of NeuroSpin and no personal data was collected.

In the design of this experiment, pairs of words from 6-7th grade and 8-9th grade, 10th grade and 11-12th grade, and license and master were merged. This yielded five groups of grade of acquisition for the words of our vocabulary.

The experiment consisted of three different parts. The first was a short demographic survey, in which we asked participants, among other things, their last grade of education in mathematics and a self-assessment of their current math skills on a scale from 1 to 10. In the second part, participants were asked to judge how familiar they were with 50 words chosen at random from our pool of 429 words. The familiarity ratings were made on a discrete scale from 0 to 8. To make the scale as objective as possible for participants 0 was labeled as “totally unknown”, 2 as “familiar word but unfamiliar concept”, 4 as “vague idea”, 6 as “familiar concept”, and 8 as “fully mastered concept”.

Finally, in the last part, participants were shown 20 pairs of words at each grade of education at or below theirs (e.g. participants who reported a high-school level of math education were shown 20 word pairs from primary school, 20 pairs of 6-9th grade and 20 pairs of 10-12th grade). The similarity judgements were made on a continuous scale from 0 to 5, and labels indicated that 0 meant “totally unrelated” and 5 “‘closely related”. The similarity rating block was preceded by a training block consisting of 8 pairs of words from everyday life (“father -mother”, “boat - car”, “table - god”, “toothbrush - television”, “Gandhi - Hitler”, “stool - crown”, “clock - Julius Caesar”, “carrot - asparagus”) covering the whole scale of possible similarities, presented in a randomized order, and 1 pair from the sailing jargon (“jib - capstan”), always presented last. This training period ensured that scale was calibrated in the same way for all participants. The order of presentation of words within each pair was randomized across participants.

In total, we recruited 1,230 participants (809 males and 421 females; age 18-25 : n = 309, 25-40: n = 385, 40-60: n = 468, > 60: n = 68) on X (Twitter at the time). The math education of participants varied between primary school to PhD (primary school: n = 4; medium school: n = 8; high school: n = 232; 1st year college: n = 58; 2nd year college: n = 170; 3rd year college: n = 199; 4th year college: n = 122; master: n = 304; PhD: n = 133).

### 4.4. Analysis pipeline

All analyses were run in Python 3.11 using the `numpy`, `statsmodels`, `scipy` and `scikit-learn` libraries. The plots were obtained using the `matplotlib` and `seaborn` modules.

#### 4.4.1. Rank linear regressions

Because many of our variables were not quantitative nor linearly ordered, we first transformed them into ranks before entering them into regressions or general linear models (GLMs). We used the `scipy.stats.rankdata` function in Python, with the default option which assigns the mean rank to ties. We then used the `statsmodels.regression.linear_model.OLS` class to fit the GLMs, and manually added an intercept (as it is not added by default) using the `statsmodels.tools.add_constant` function.

#### 4.4.2. Noise ceiling

In order to get an approximation of the amount of noise in the behavioral data, we computed a noise ceiling. The ceiling captures the average amount of variance present in one participant’s responses that cannot be explained by the average responses of the other participants.

To compute the noise ceiling, we performed a leave-one-out cross validation. For each individual participant, we computed the correlation between its answers and the average answers of the other participants. We then computed the average *R^2^*.

In the case of noise ceiling in subgroups (for instance for groups of participants with the same level of math education), there sometimes was not enough overlap between the stimuli seen by one particular participant and all the others. Therefore, when computing noise ceiling in any given subgroup, we correlated the answers of each participant of the subgroup with the average answers of all participants (and not only those belonging to the same subgroup). It must be noted that the noise ceiling is an approximation of the level of noise in the data, and our measure probably overestimates this level.

#### 4.4.3. Comparison of similarities judged by humans with those predicted by GloVe

When computing the correlation between human judgements and GloVe-predicted similarities, we first averaged human judgements for a given pair over all trials and then computed the correlation across pairs (Fig 3). However, to compare with the within-participant noise ceiling, we also recomputed correlations across all individual trials, without performing the averaging step (Fig 4).

#### 4.4.4. Item Response Theory (IRT)

In order to apply Item Response Theory (Cai et al., 2016) to the familiarity ratings provided by participants, we dichotomised the answers: familiarity ratings comprised between 0 and 3 (included) were considered grouped in a low category (“unknown”), and ratings equal to or above 4 meant that the word was known to participant. This dichotomization step was needed since we wanted to model participants’ responses as a single sigmoid, and not as several logistic curves as standard methods for polytomous data would do (Samejima, 1997).

IRT aims to estimate jointly a latent ability for participants and a discrimination and difficulty parameter for each item. This is done using expectation-maximization algorithms. Given an item (i.e. a word in the vocabulary) *i* and a participant with latent ability *θ*, the probability *T_i_* that the participant answers that they know the item (familiarity rating equal to 4, 5, 6, 7 or 8) is modeled as

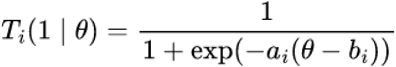

where *a_i_* is the item’s discrimination parameter and *b_i_* is its difficulty parameter.

Note that, for this particular analysis only, we used R 4.1 and its mirt package.

#### 4.4.5. Model comparison

Whenever we compared two or more statistical models, we used Akaike Information Criterion (AIC). For each model *i*, we report the ΔAIC_i_, which is defined as: ΔAIC_i_ = AIC_i_ - AIC_min_, where AIC_min_ is the lowest AIC obtained among the models under consideration (i.e. the AIC of the preferred model). The *p*-value associated with ΔAIC_i_ is given by *p_i_ =* exp(- ΔAICi/2) (Burnham & Anderson, 2004).

### 4.5. Replication in English

We applied a similar method as in French to create a math, non-math and global corpora from Wikipedia pages in English. This time, we used HuggingFace’s Wikipedia `20220301.en` dataset and the classification of math versus non-math pages was done using the `wikipedia_en_mathematics_nopic_2023-09.zim` dump, which was already curated to only contain the math pages from Wikipedia. The corpora gather 22,889 math pages and 6,435,781 non-math pages.

We also obtained GloVe embeddings using the same methodology, after two passes of lemmatization using spacy’s en_core_web_lg model, and used the vocabulary output by the GloVe pipeline on the math corpus to create a 1,000-word vocabulary of math words.

We then manually translated every single word of the French math vocabulary into English. For the online experiment, out of the original 429 stimuli tested with French participants, we selected only the 296 which had a unique translation in the English math vocabulary. Following the same procedure as in French, these stimuli yielded 2,458 pairs for the similarity rating task.

We translated the behavioral experiment in English and ran it online. In total, we recruited 174 participants (88 males and 86 females; age 18-25: n = 14, 25-40: n = 64, 40-60: n = 71, > 60: n = 25). The math education of participants varied from medium school to PhD (medium school: n = 1; high school: n = 37; 1st year college: n = 26; 2nd year college: n = 19; 3rd year college: n = 22; 4th year college: n = 12; master: n = 39; PhD: n = 18). The analysis pipeline was exactly the same in English as in French, but because of the lower number of participants, we did not attempt to perform within-level analyses.

## 5. Data availability statement

All human data, statistical analysis code, GloVe embeddings and online experiments source code are available on OSF at link https://osf.io/dxg2w. The math vocabulary and its 50-dimensional embeddings are also available in the OSF repository. Due their large size, the corpora used to train GloVe models could not be shared online but are available from the corresponding author SaD on request.

## 6. Acknowledgements

We are grateful to Alexandre Pasquiou for his daily support in the creation of the corpora and the extraction of the embeddings. We also thank Mathias Sablé-Meyer for his tutorial on how to get started with jsPsych, and all participants who took part in our online experiment.

## 8. Author contributions

● Conceptualization: SaD and StD
● Data curation: SaD
● Formal analysis: SaD and StD
● Funding acquisition: StD
● Investigation: SaD and StD
● Methodology: SaD and StD
● Supervision: StD
● Validation: StD
● Visualization: SaD
● Writing – original draft: SaD
● Writing – review and editing: StD

## 10. Supporting information

**S1 PDF File. Individual IRT plots for the 429 words selected in the online experiment based on participants’ familiarity ratings.**

**S2 Table. Number of pairs selected for the similarity rating experiment for each word grade.**

