## Supplementary Table 1 for "Mapping and modeling the semantic space of math concepts"

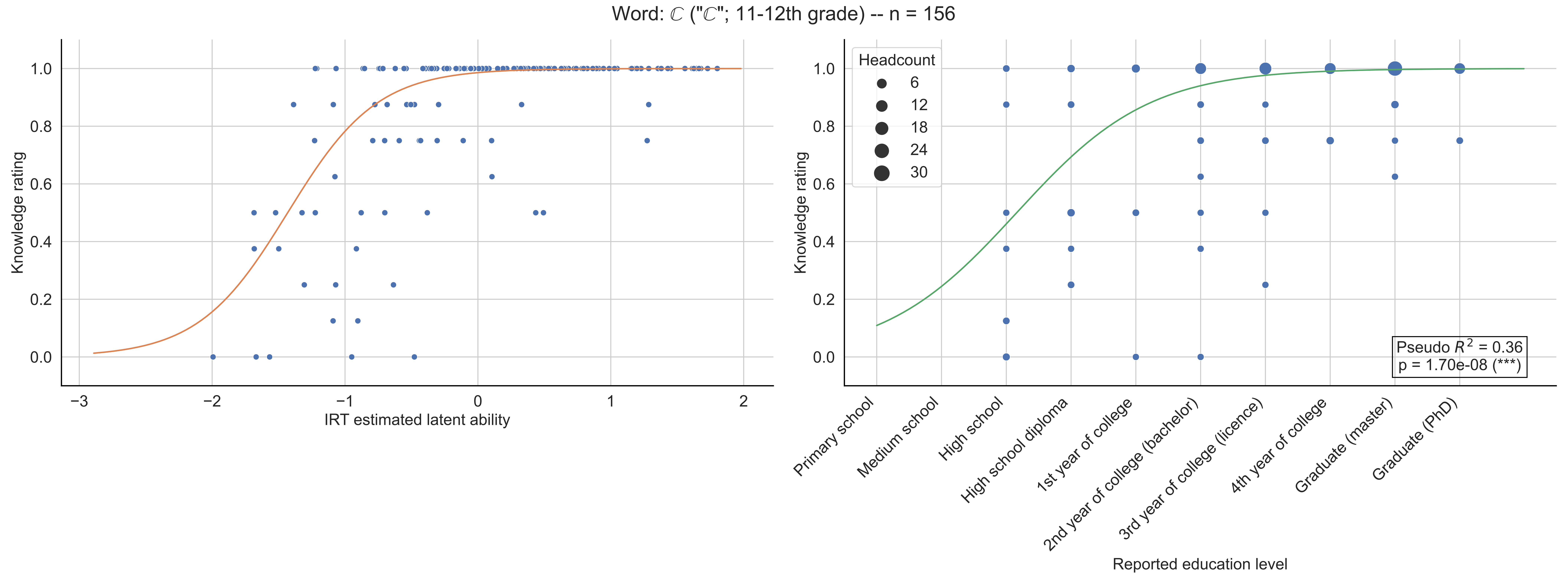

Word: ℂ ("ℂ"; 11-12th grade) -- n = 156
Knowledge rating
IRT estimated latent ability
Headcount
6
12
18
24
30
Pseudo $R^2$ = 0.36
p = 1.70e-08 (***)
Primary school
Medium school
High school
High school diploma
1st year of college
2nd year of college (bachelor)
3rd year of college (licence)
4th year of college
Graduate (master)
Graduate (PhD)
Reported education level

Word: ℕ ("ℕ"; 10th grade) -- n = 146

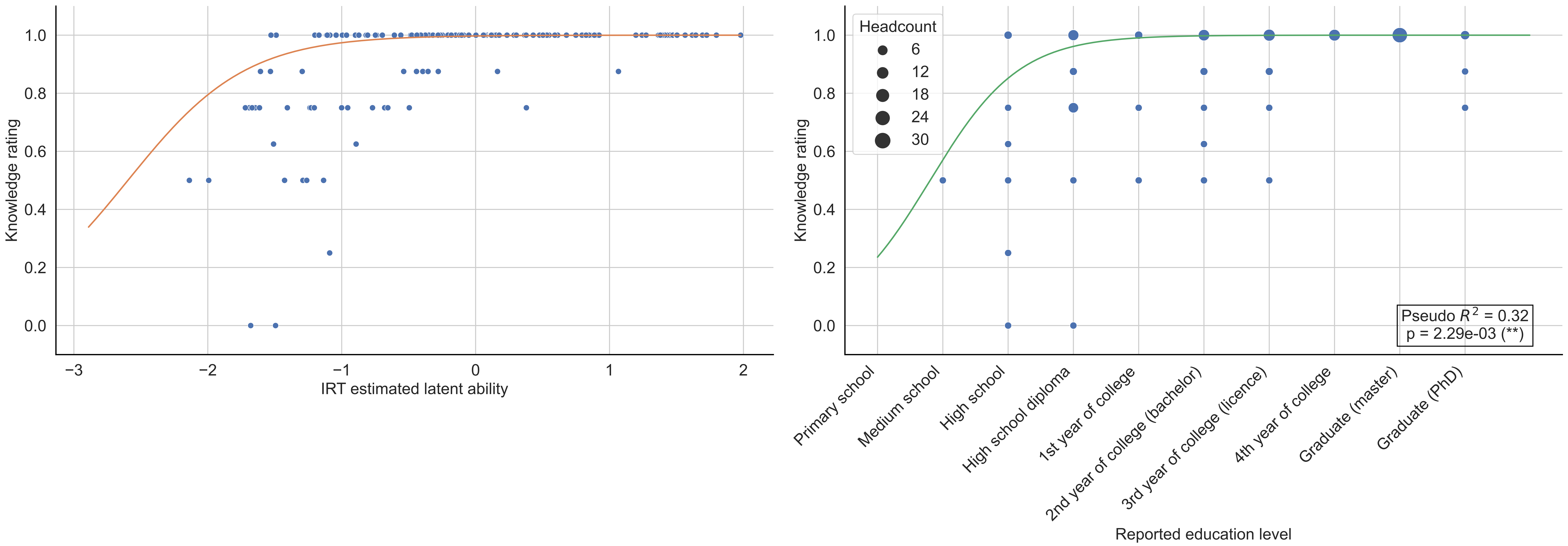

Word: ℚ ("ℚ"; 10th grade) -- n = 152

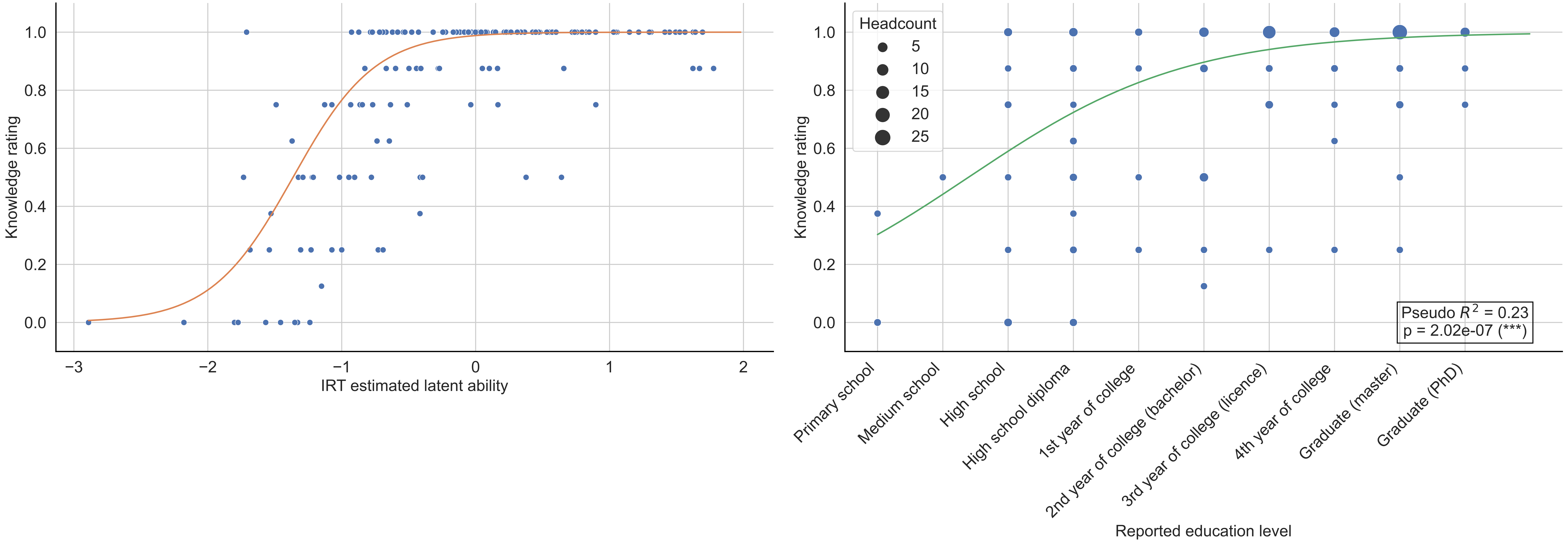

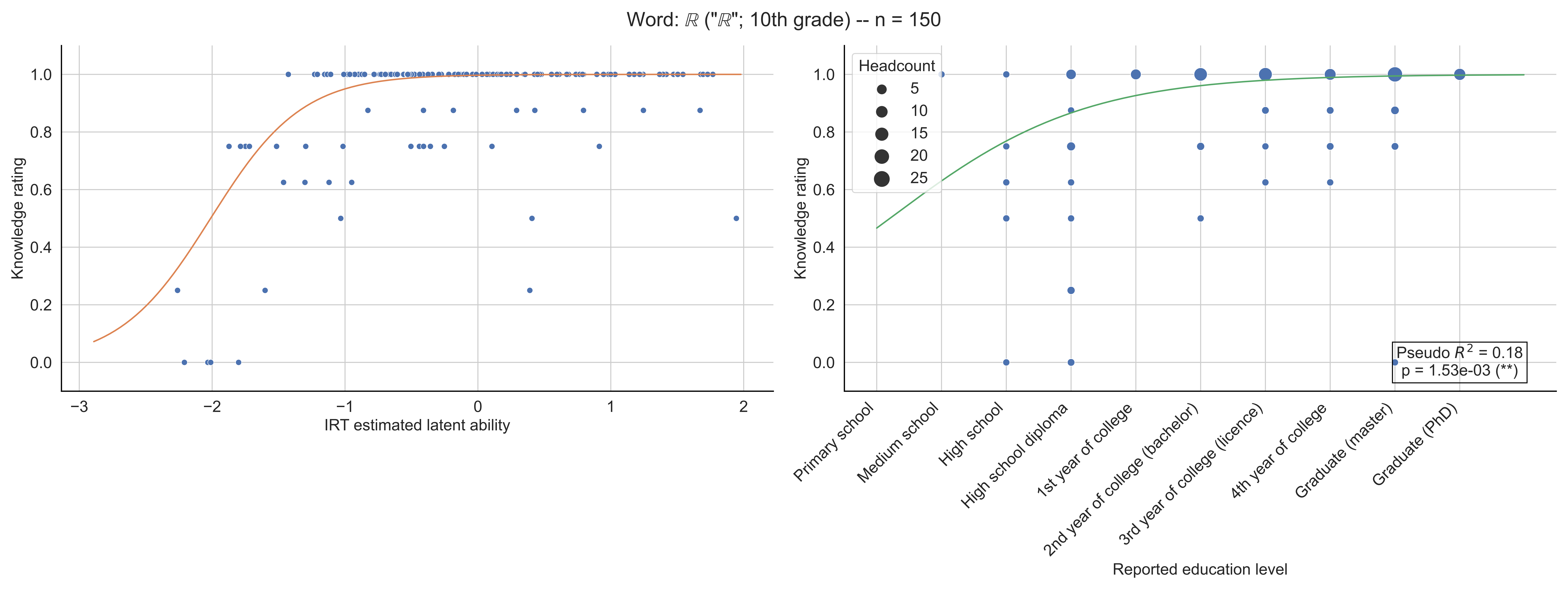

Word: ℝ ("ℝ"; 10th grade) -- n = 150
Knowledge rating
IRT estimated latent ability
Headcount
5
10
15
20
25
Pseudo $R^2$ = 0.18
p = 1.53e-03 (**)
Primary school
Medium school
High school
High school diploma
1st year of college
2nd year of college (bachelor)
3rd year of college (licence)
4th year of college
Graduate (master)
Graduate (PhD)
Reported education level

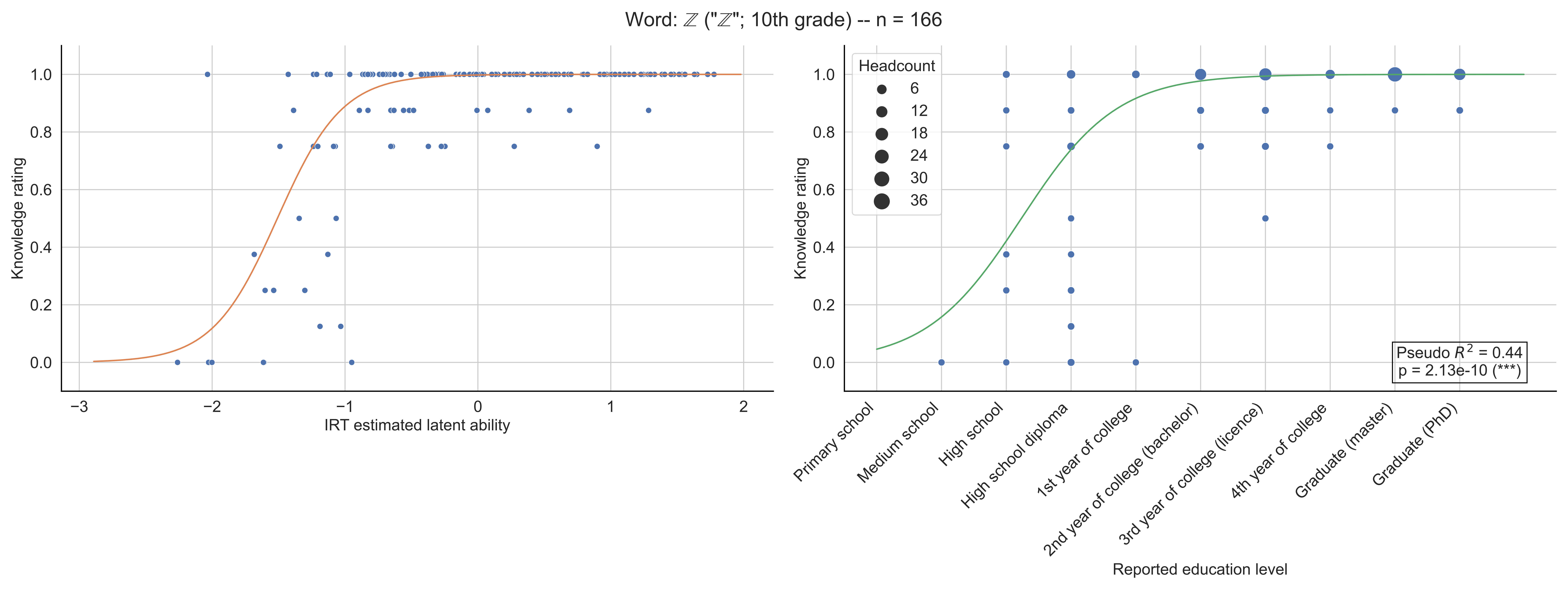

Word: ℤ ("ℤ"; 10th grade) -- n = 166
Knowledge rating
IRT estimated latent ability
Headcount
6
12
18
24
30
36
Pseudo $R^2$ = 0.44
p = 2.13e-10 (***)
Primary school
Medium school
High school
High school diploma
1st year of college
2nd year of college (bachelor)
3rd year of college (licence)
4th year of college
Graduate (master)
Graduate (PhD)
Reported education level

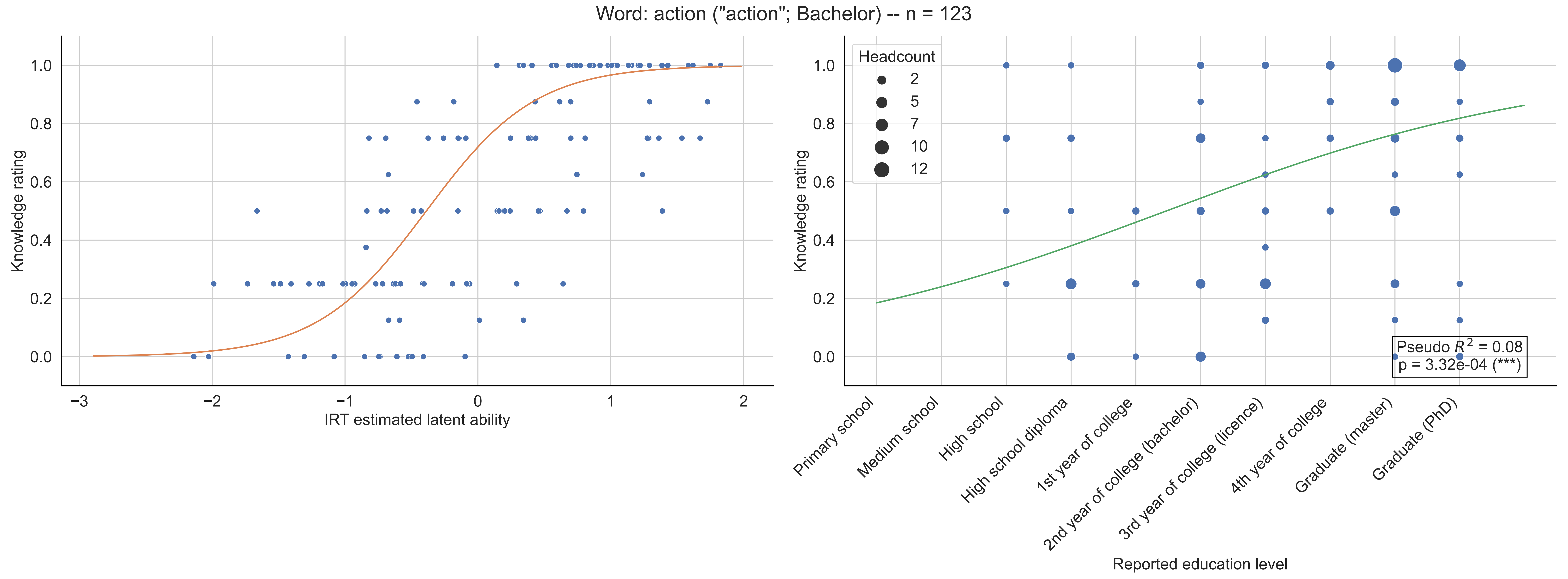

Word: action ("action"; Bachelor) -- n = 123
Knowledge rating
IRT estimated latent ability
Headcount
2
5
7
10
12
Primary school
Medium school
High school
High school diploma
1st year of college
2nd year of college (bachelor)
3rd year of college (licence)
4th year of college
Graduate (master)
Graduate (PhD)
Reported education level
Pseudo $R^2$ = 0.08
p = 3.32e-04 (***)

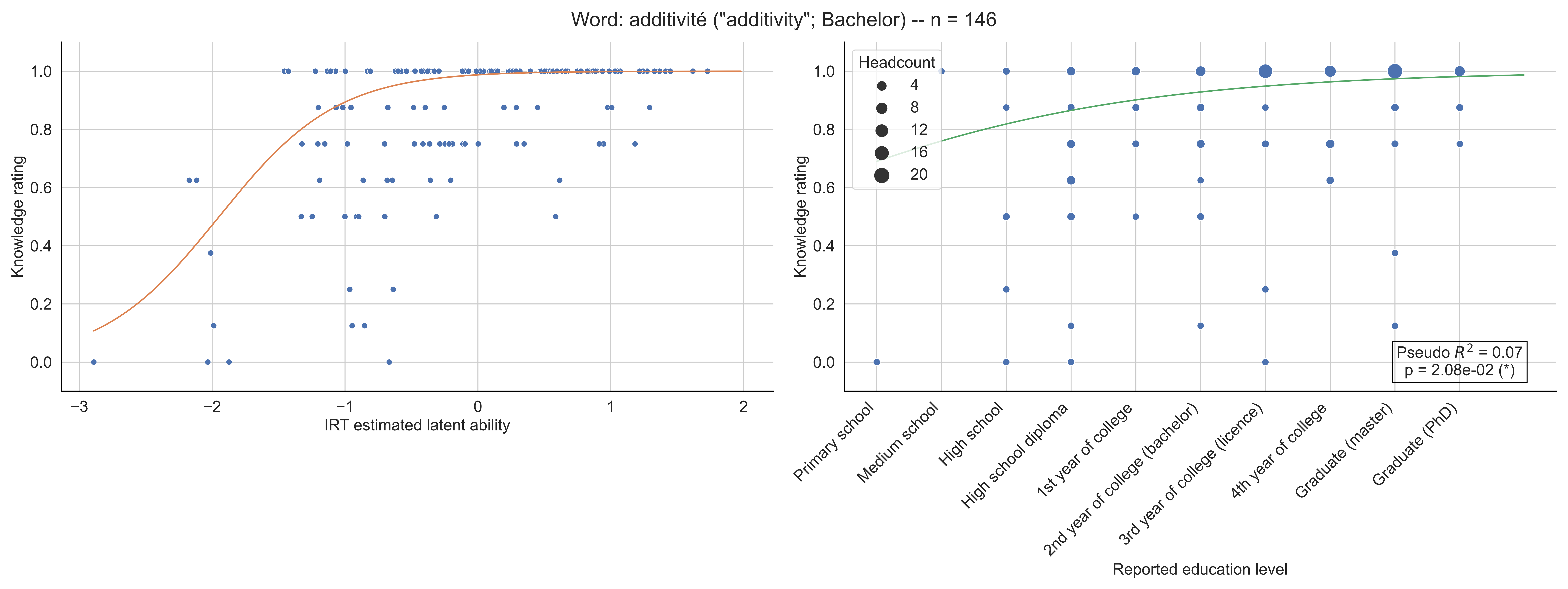

Word: additivité ("additivity"; Bachelor) -- n = 146
Knowledge rating
IRT estimated latent ability
Headcount
4
8
12
16
20
Pseudo $R^2$ = 0.07
p = 2.08e-02 (*)
Primary school
Medium school
High school
High school diploma
1st year of college
2nd year of college (bachelor)
3rd year of college (licence)
4th year of college
Graduate (master)
Graduate (PhD)
Reported education level

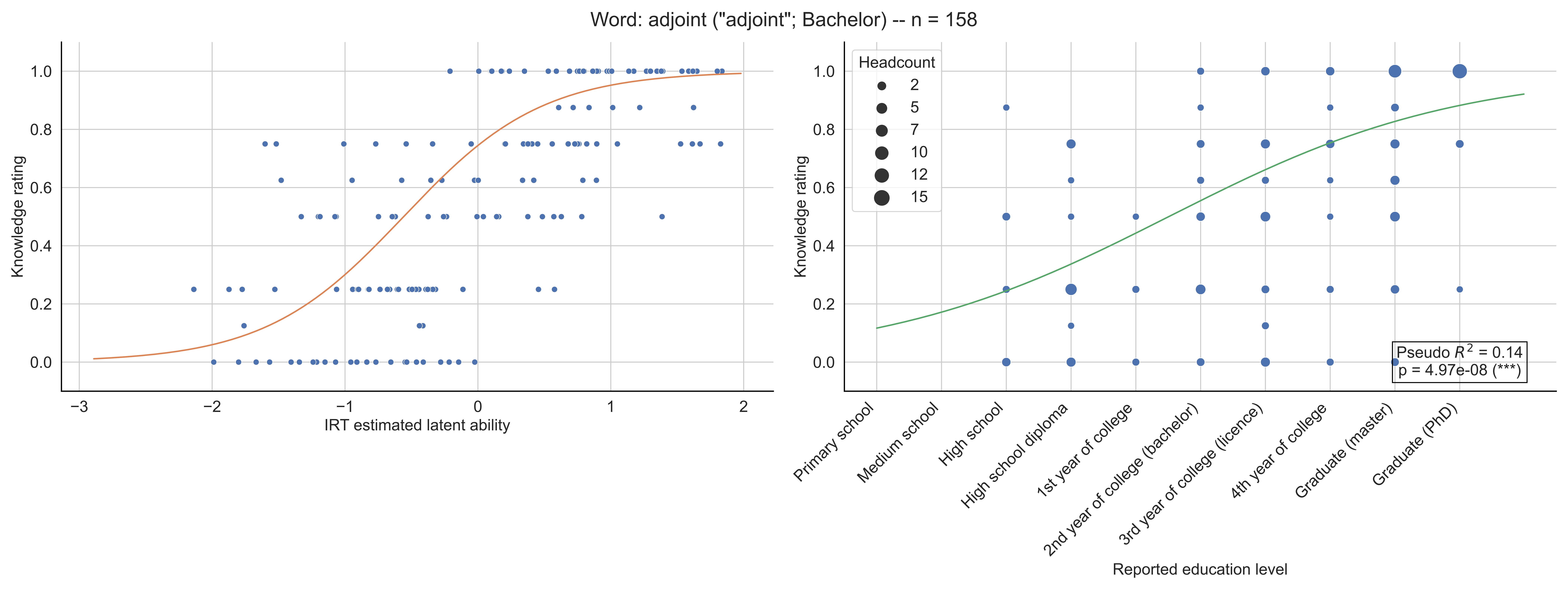
Word: adjoint ("adjoint"; Bachelor) -- n = 158
Knowledge rating
IRT estimated latent ability
Headcount
2
5
7
10
12
15
Primary school
Medium school
High school
High school diploma
1st year of college
2nd year of college (bachelor)
3rd year of college (licence)
4th year of college
Graduate (master)
Graduate (PhD)
Reported education level
Pseudo $R^2$ = 0.14
p = 4.97e-08 (***)

Word: affixe (11-12th grade) -- n = 181

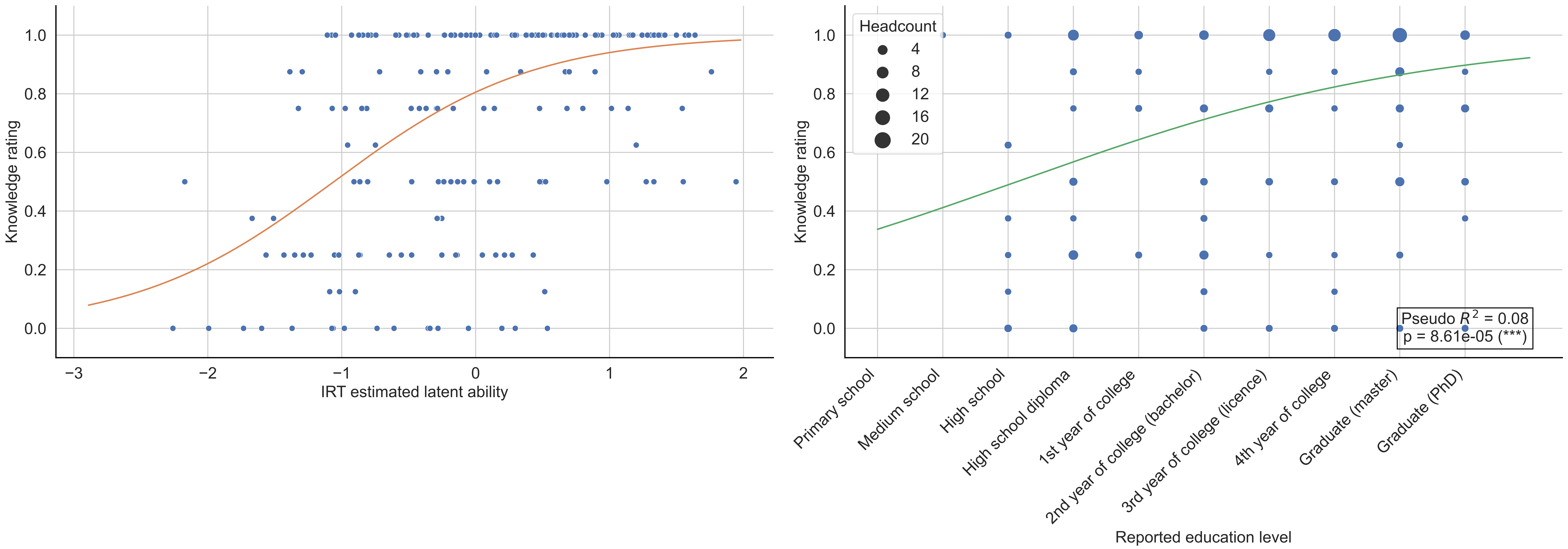

Word: algorithmique ("algorithmic"; 6-7th grade) -- n = 139

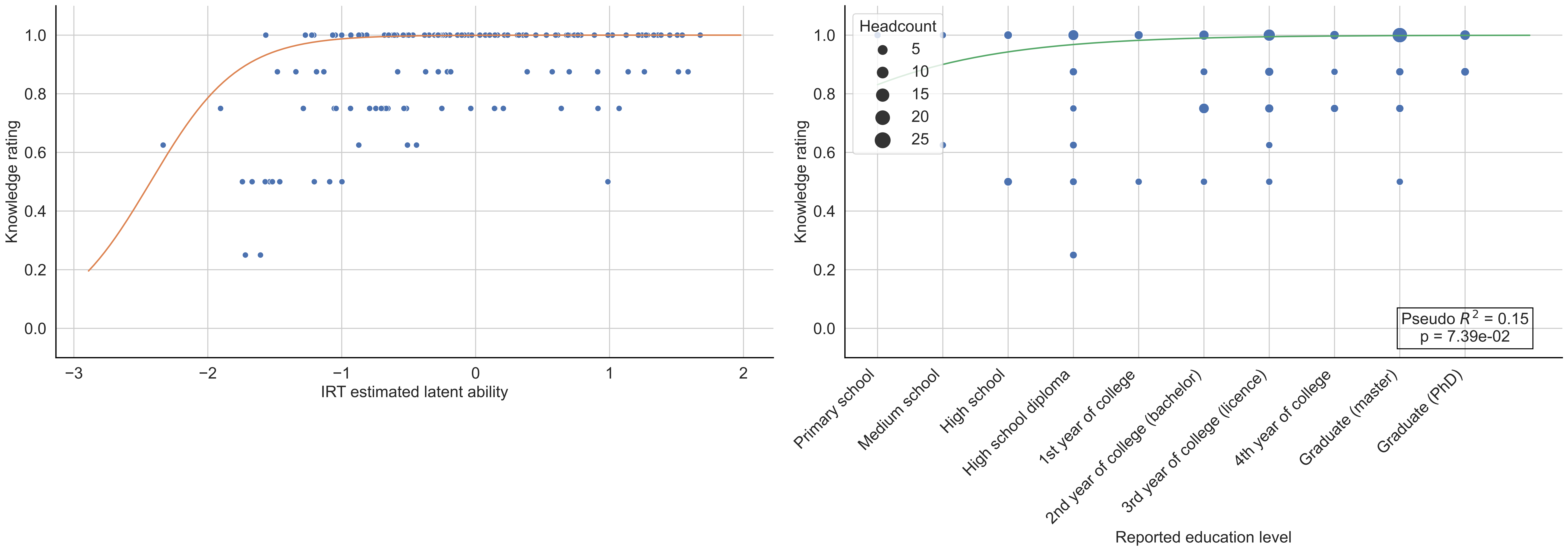

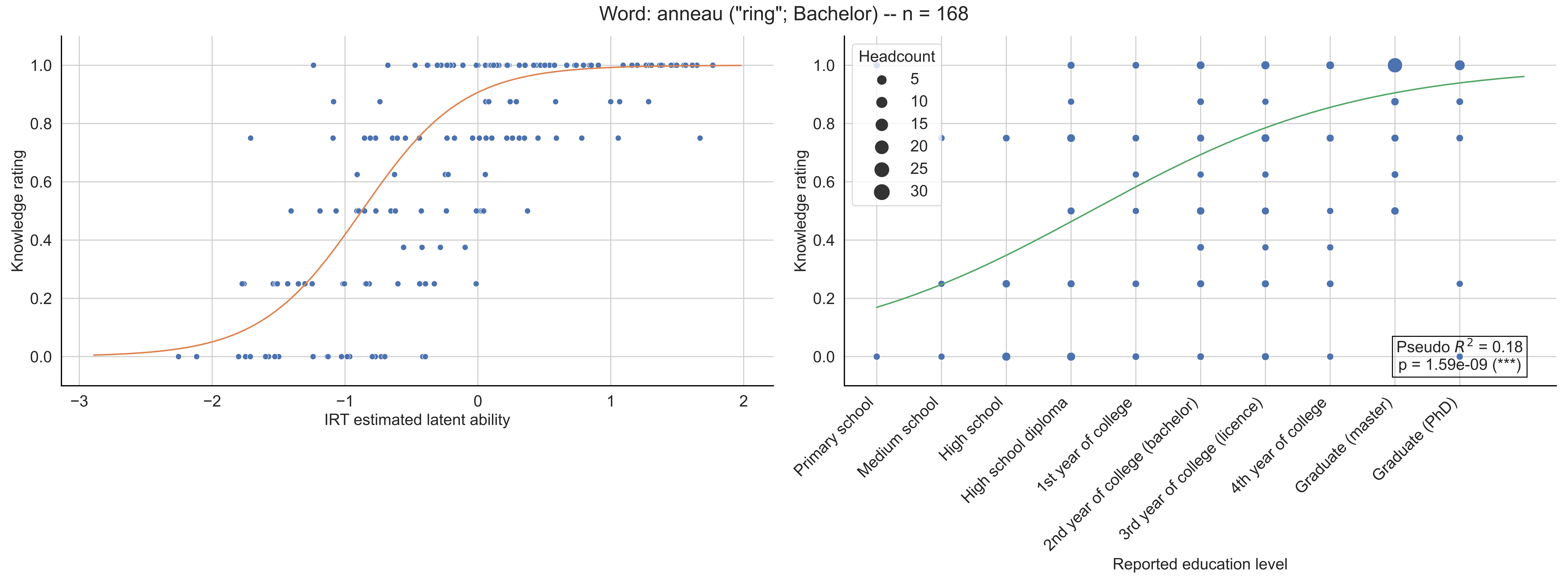
Word: anneau ("ring"; Bachelor) -- n = 168
Knowledge rating
IRT estimated latent ability
−3
−2
−1
0
1
2
0.0
0.2
0.4
0.6
0.8
1.0
Headcount
5
10
15
20
25
30
Pseudo $R^2$ = 0.18
p = 1.59e-09 (***)
Primary school
Medium school
High school
High school diploma
1st year of college
2nd year of college (bachelor)
3rd year of college (licence)
4th year of college
Graduate (master)
Graduate (PhD)
Reported education level

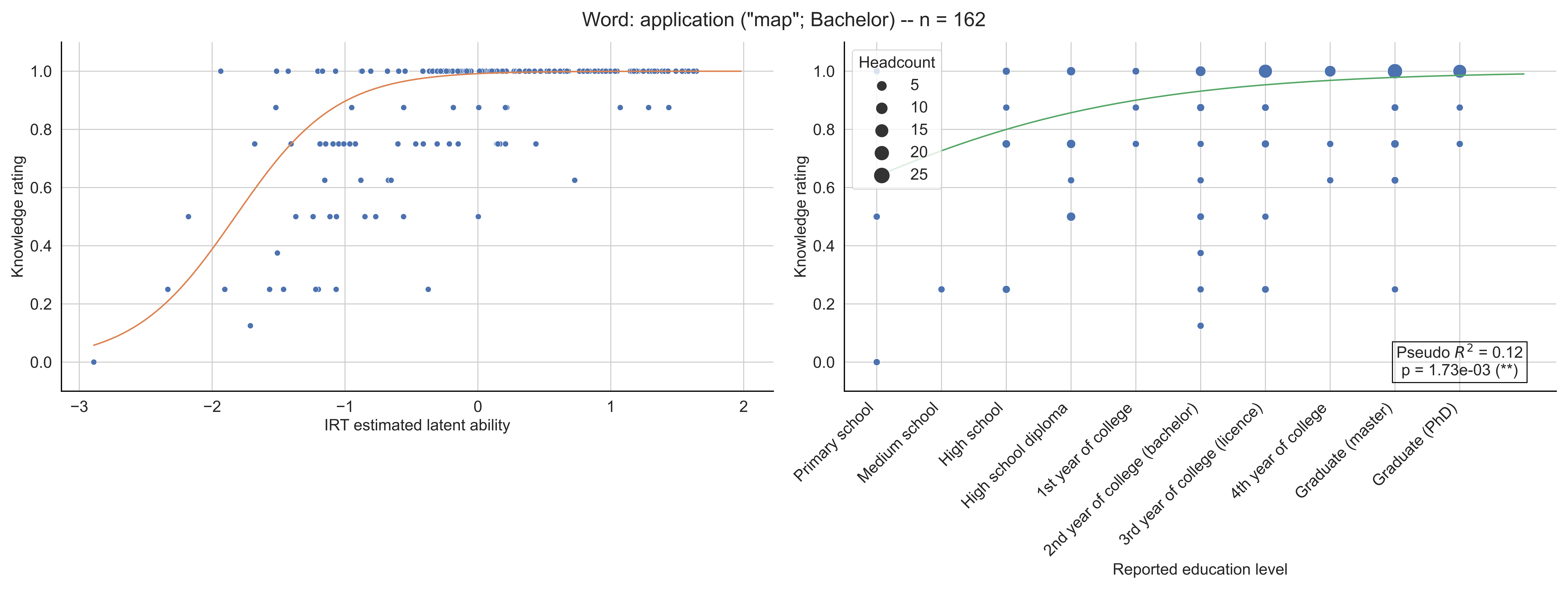
Word: application ("map"; Bachelor) -- n = 162
Knowledge rating
IRT estimated latent ability
Headcount
5
10
15
20
25
Pseudo $R^2$ = 0.12
p = 1.73e-03 (**)
Primary school
Medium school
High school
High school diploma
1st year of college
2nd year of college (bachelor)
3rd year of college (licence)
4th year of college
Graduate (master)
Graduate (PhD)
Reported education level

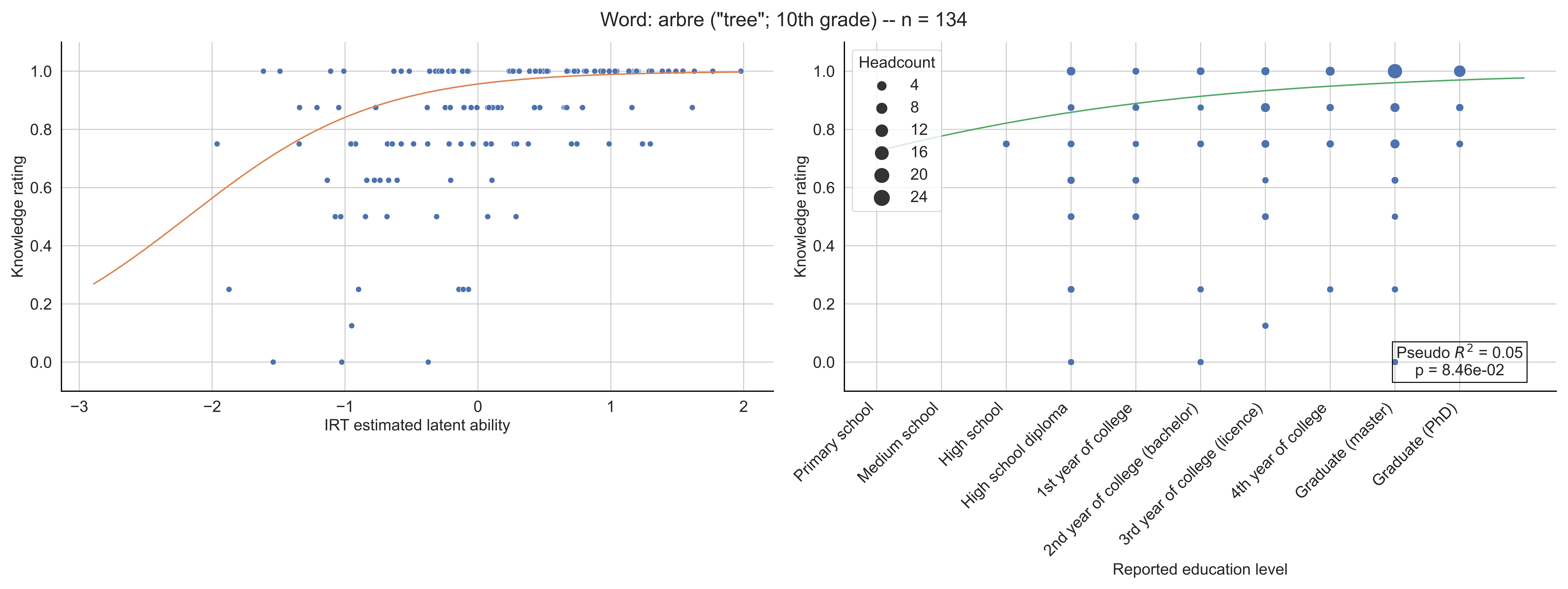

Word: arbre ("tree"; 10th grade) -- n = 134
Knowledge rating
IRT estimated latent ability
Headcount
4
8
12
16
20
24
Pseudo $R^2$ = 0.05
p = 8.46e-02
Primary school
Medium school
High school
High school diploma
1st year of college
2nd year of college (bachelor)
3rd year of college (licence)
4th year of college
Graduate (master)
Graduate (PhD)
Reported education level

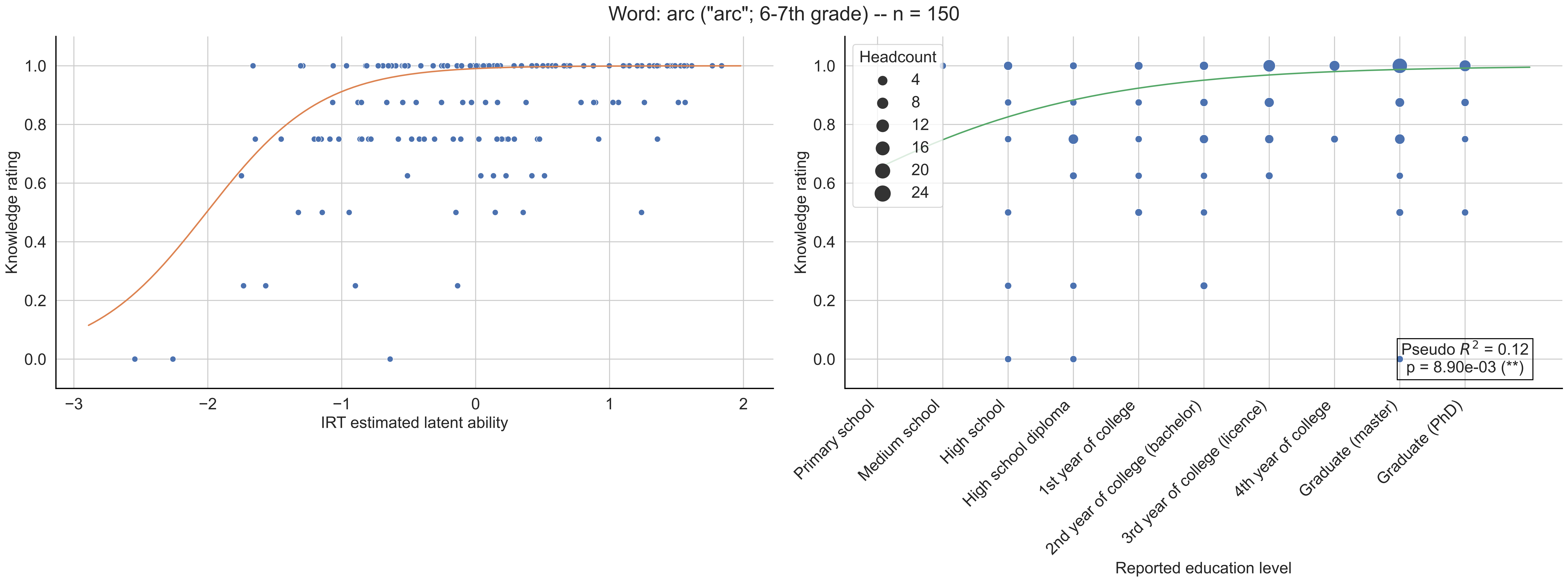

Word: arc ("arc"; 6-7th grade) -- n = 150
Knowledge rating
IRT estimated latent ability
Headcount
4
8
12
16
20
24
Pseudo $R^2$ = 0.12
p = 8.90e-03 (**)
Primary school
Medium school
High school
High school diploma
1st year of college
2nd year of college (bachelor)
3rd year of college (licence)
4th year of college
Graduate (master)
Graduate (PhD)
Reported education level

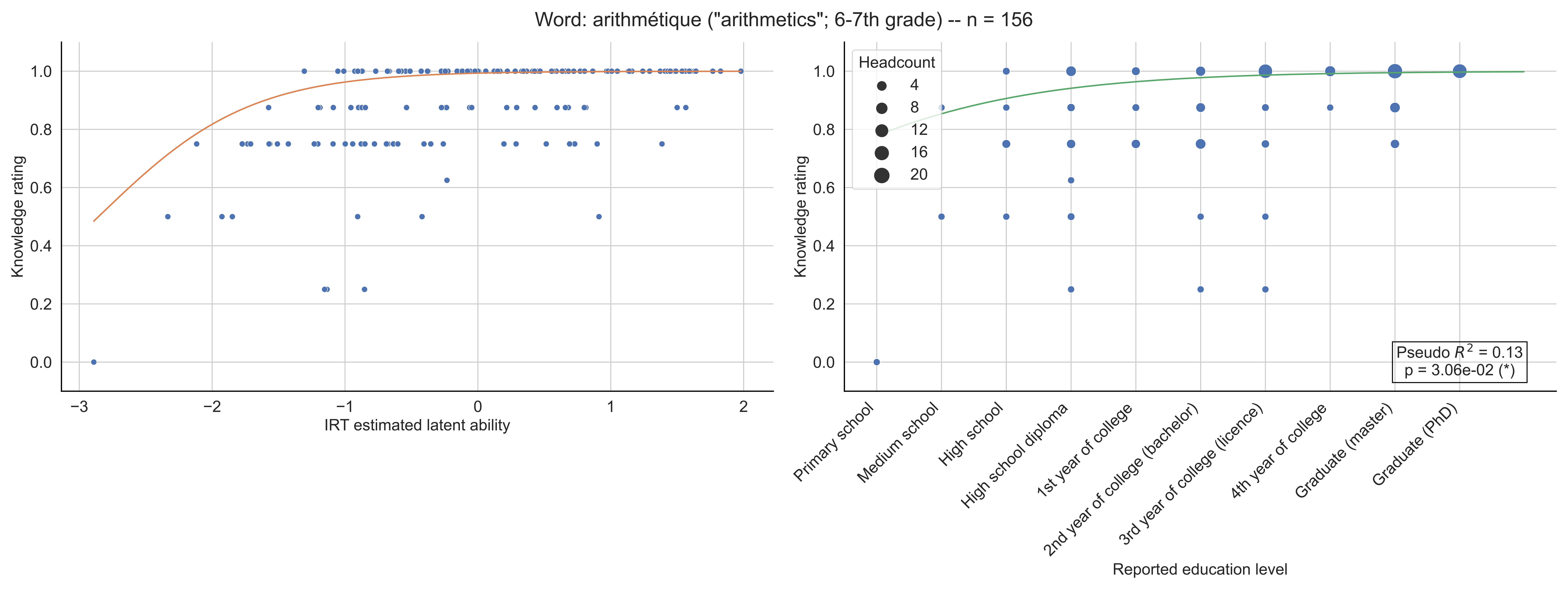

Word: arithmétique ("arithmetics"; 6-7th grade) -- n = 156
Knowledge rating
IRT estimated latent ability
Headcount
4
8
12
16
20
Pseudo $R^2$ = 0.13
p = 3.06e-02 (*)
Primary school
Medium school
High school
High school diploma
1st year of college
2nd year of college (bachelor)
3rd year of college (licence)
4th year of college
Graduate (master)
Graduate (PhD)
Reported education level

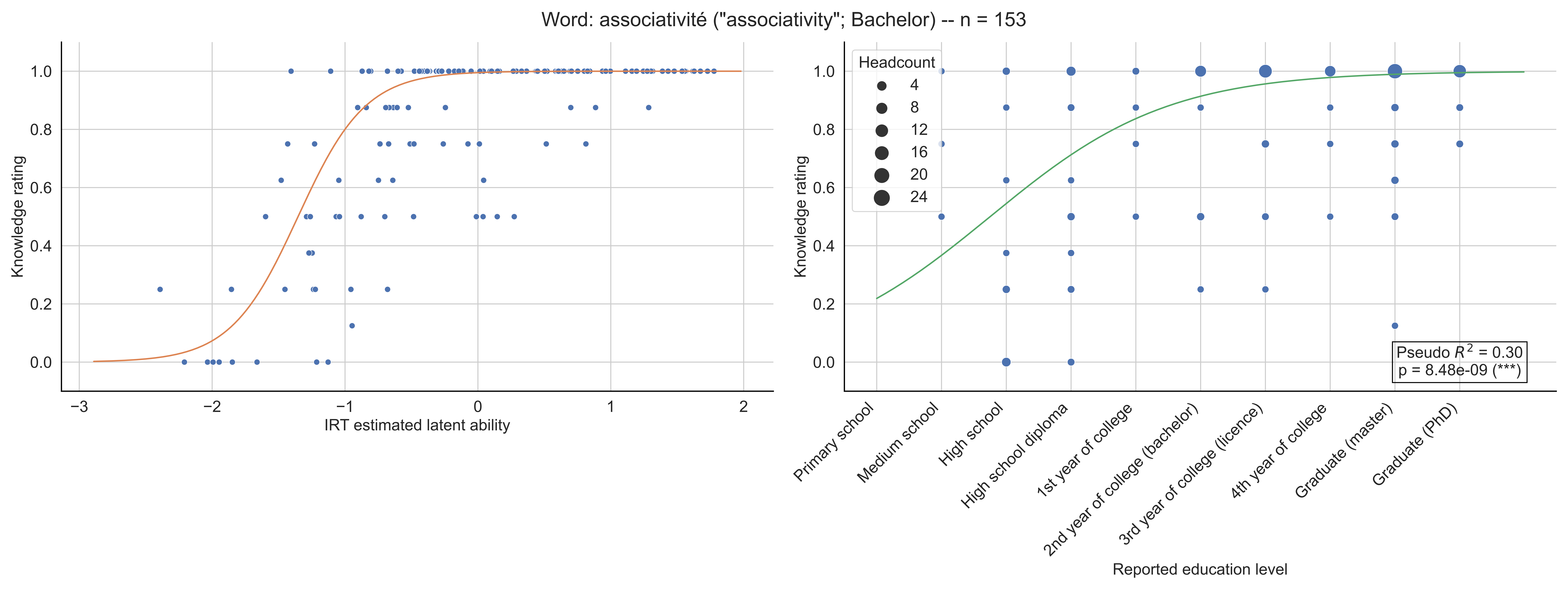

Word: associativité ("associativity"; Bachelor) -- n = 153
Knowledge rating
IRT estimated latent ability
Headcount
4
8
12
16
20
24
Pseudo $R^2$ = 0.30
p = 8.48e-09 (***)
Primary school
Medium school
High school
High school diploma
1st year of college
2nd year of college (bachelor)
3rd year of college (licence)
4th year of college
Graduate (master)
Graduate (PhD)
Reported education level

Word: asymptote ("asymptote"; 11-12th grade) -- n = 158

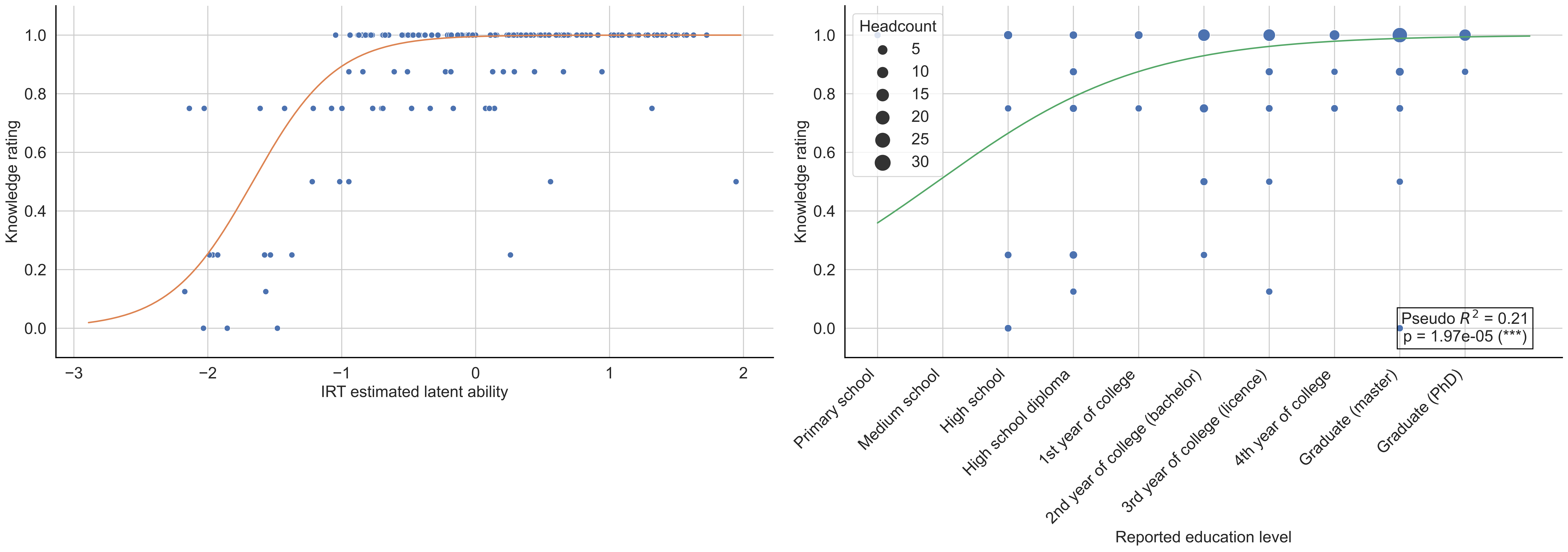

Word: attracteur ("attractor"; Licence) -- n = 146

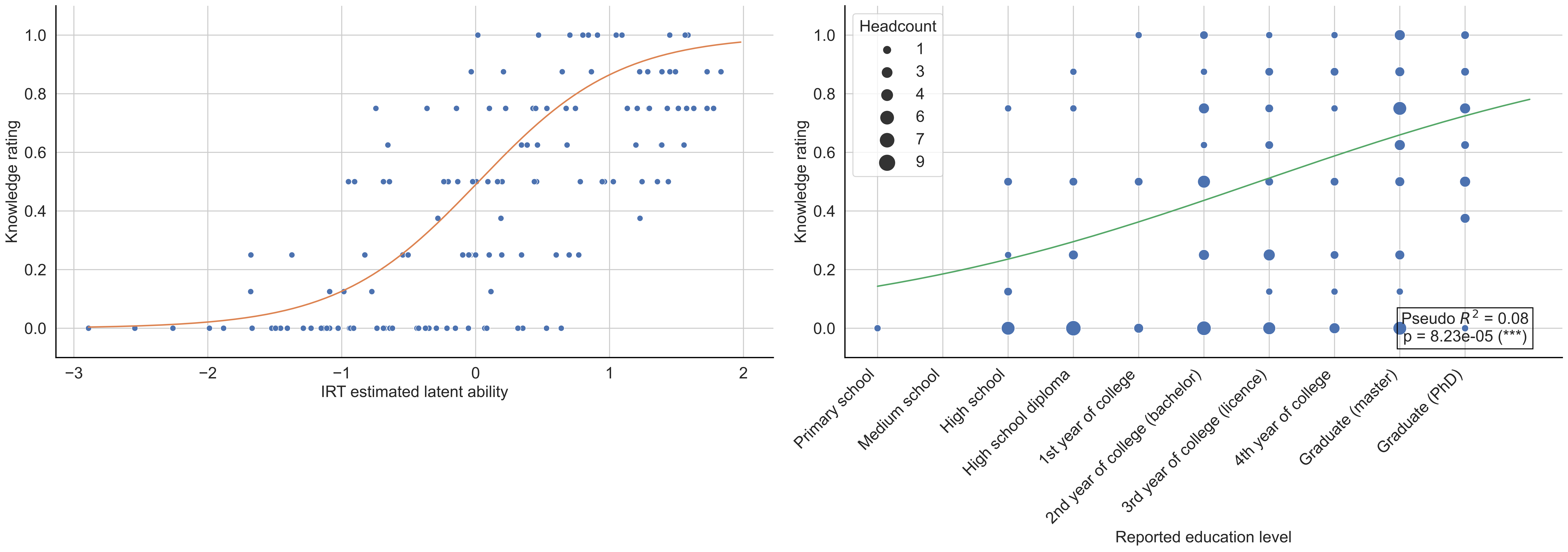

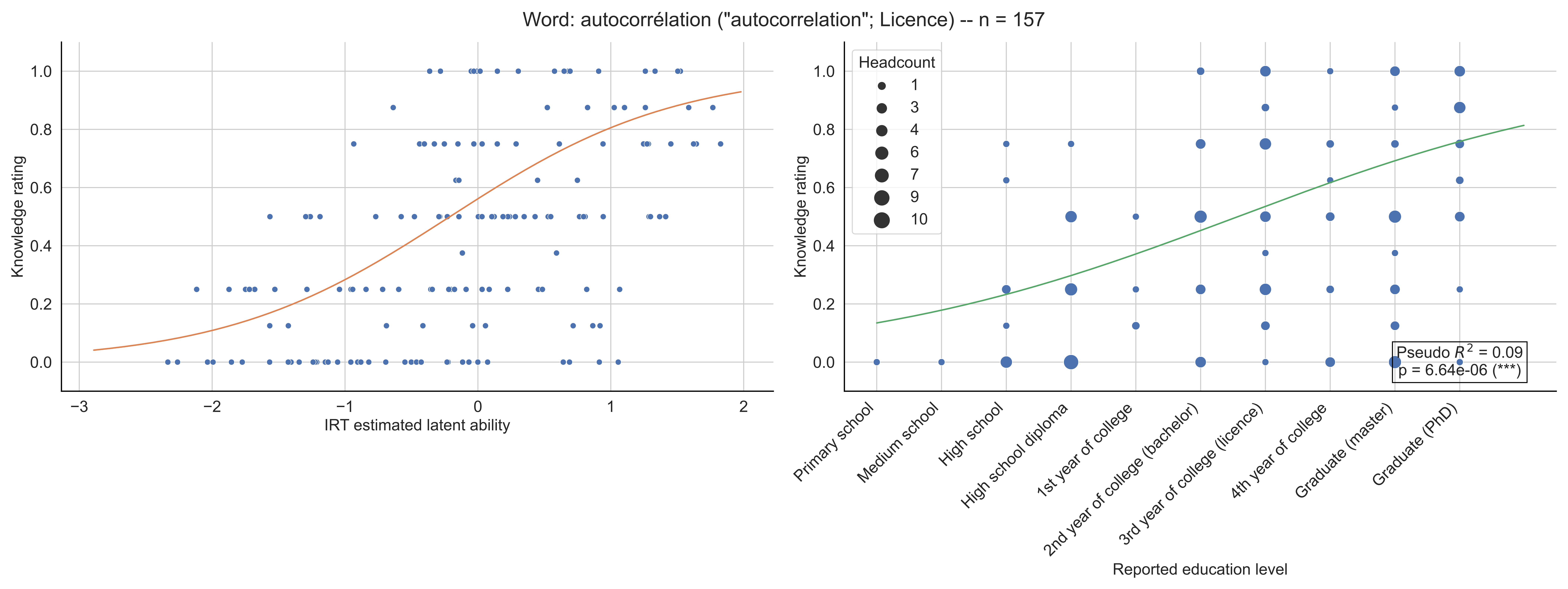

Word: autocorrélation ("autocorrelation"; Licence) -- n = 157
Knowledge rating
IRT estimated latent ability
Headcount
1
3
4
6
7
9
10
Pseudo $R^2$ = 0.09
p = 6.64e-06 (***)
Primary school
Medium school
High school
High school diploma
1st year of college
2nd year of college (bachelor)
3rd year of college (licence)
4th year of college
Graduate (master)
Graduate (PhD)
Reported education level

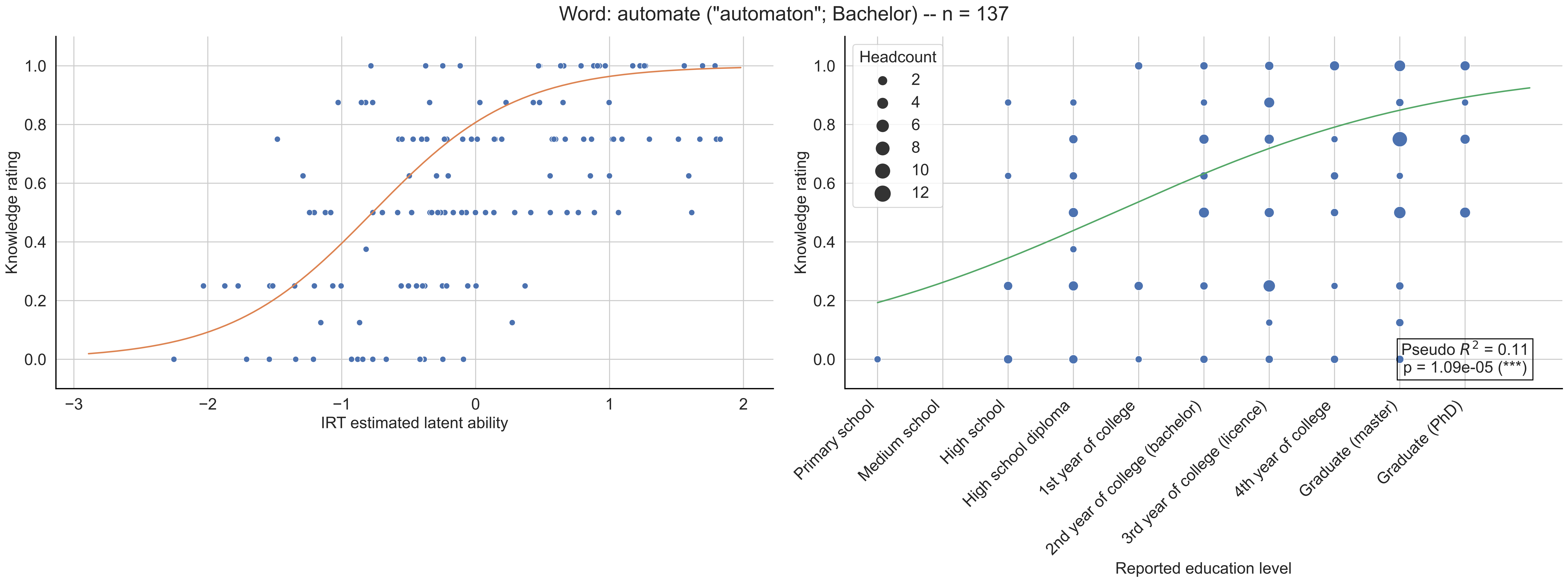
Word: automate ("automaton"; Bachelor) -- n = 137
Knowledge rating
IRT estimated latent ability
Headcount
2
4
6
8
10
12
Pseudo $R^2$ = 0.11
p = 1.09e-05 (***)
Primary school
Medium school
High school
High school diploma
1st year of college
2nd year of college (bachelor)
3rd year of college (licence)
4th year of college
Graduate (master)
Graduate (PhD)
Reported education level

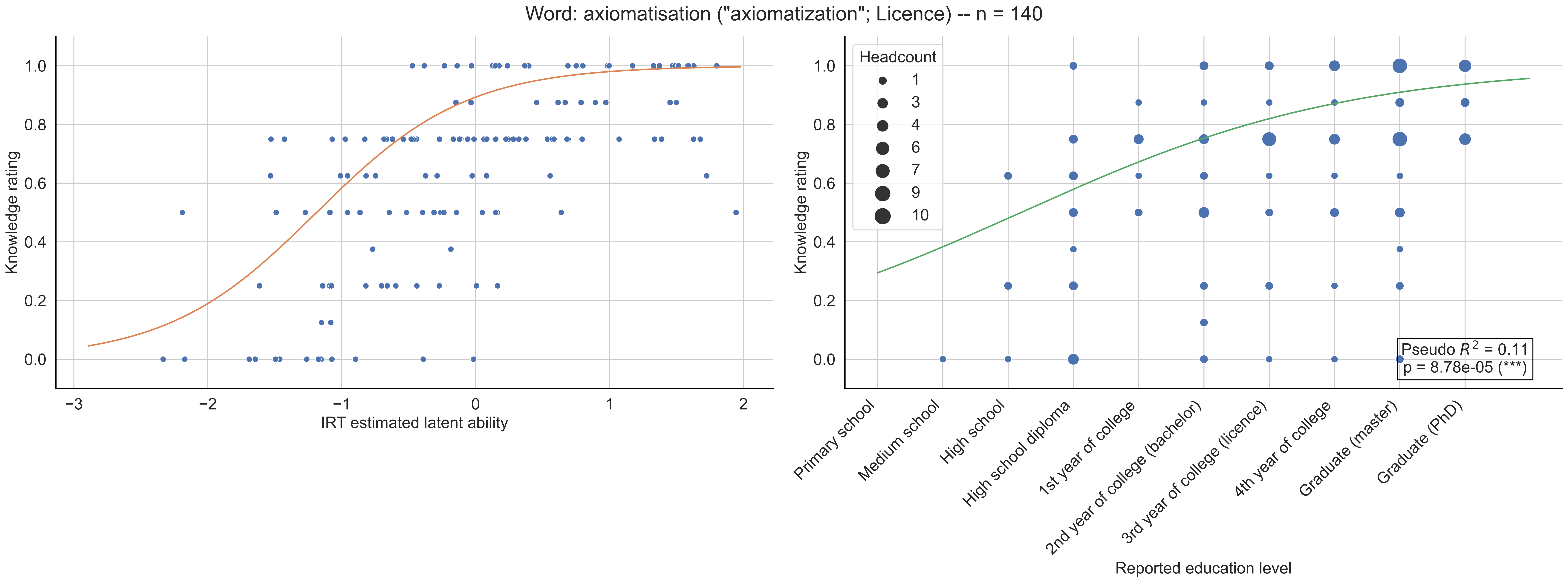

Word: axiomatisation ("axiomatization"; Licence) -- n = 140
Knowledge rating
IRT estimated latent ability
Headcount
1
3
4
6
7
9
10
Pseudo $R^2$ = 0.11
p = 8.78e-05 (***)
Primary school
Medium school
High school
High school diploma
1st year of college
2nd year of college (bachelor)
3rd year of college (licence)
4th year of college
Graduate (master)
Graduate (PhD)
Reported education level

Word: barycentre ("barycenter"; 6-7th grade) -- n = 147

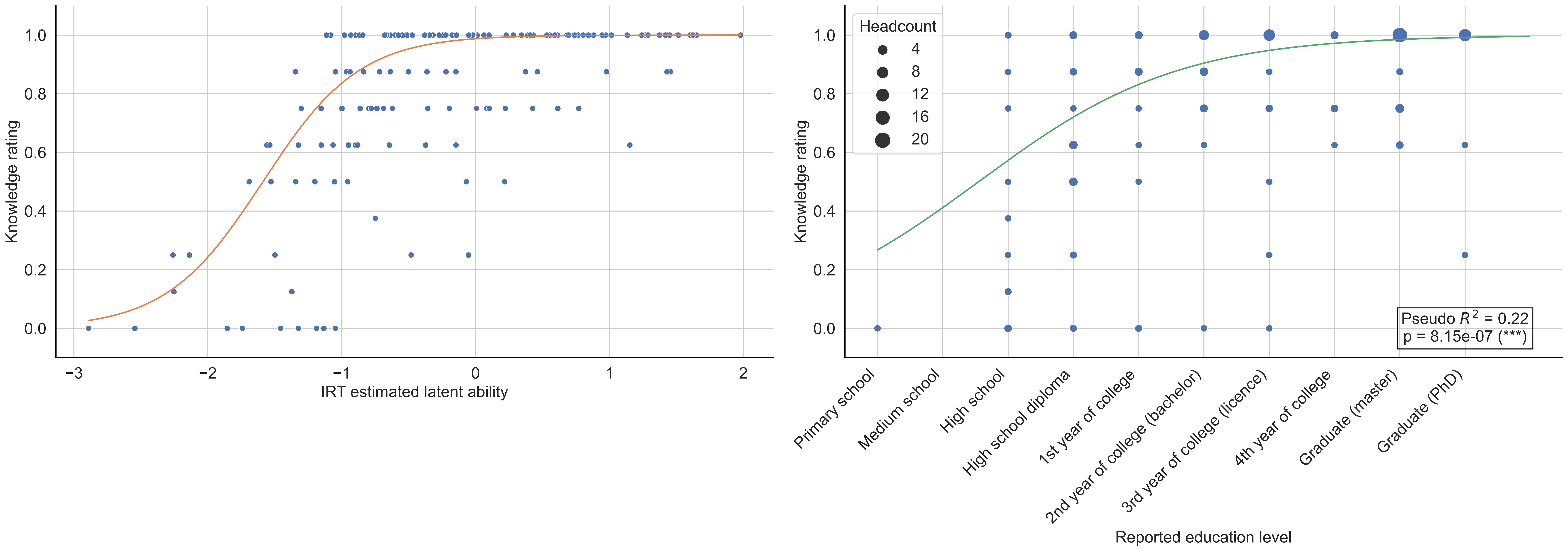

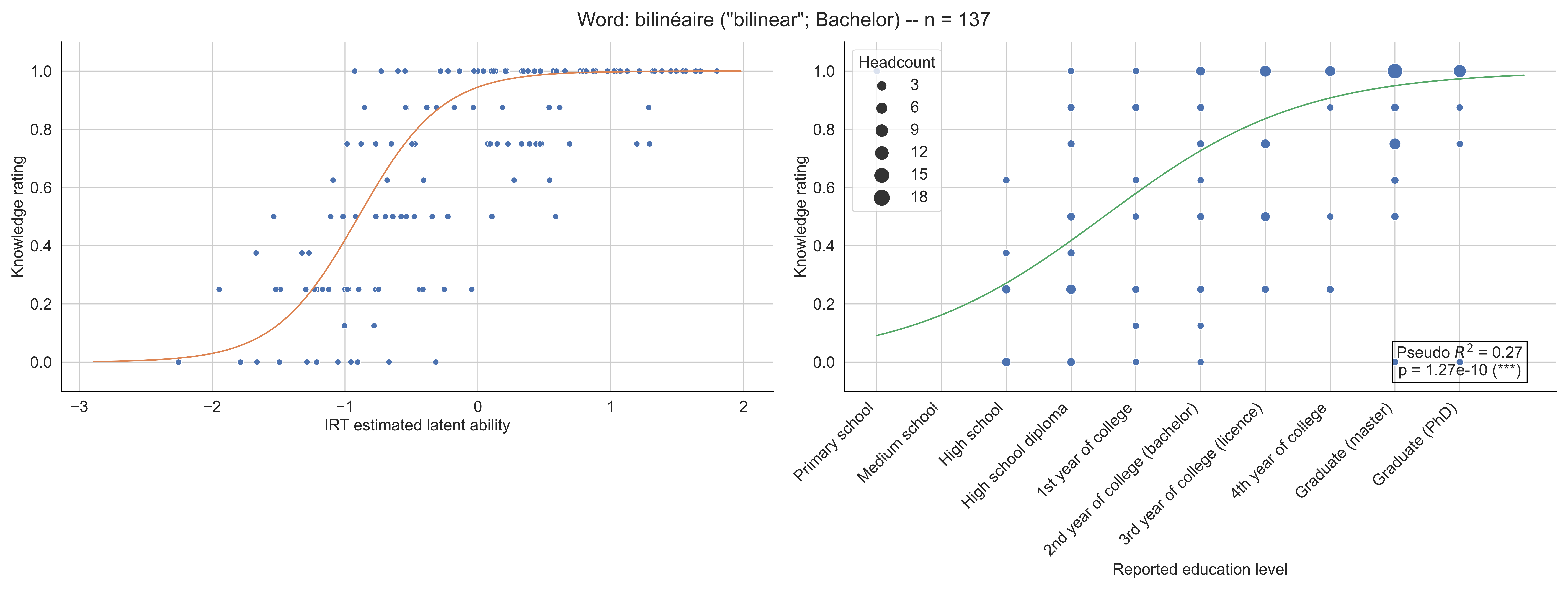

Word: bilinéaire ("bilinear"; Bachelor) -- n = 137
Knowledge rating
IRT estimated latent ability
Headcount
3
6
9
12
15
18
Pseudo $R^2$ = 0.27
p = 1.27e-10 (***)
Primary school
Medium school
High school
High school diploma
1st year of college
2nd year of college (bachelor)
3rd year of college (licence)
4th year of college
Graduate (master)
Graduate (PhD)
Reported education level

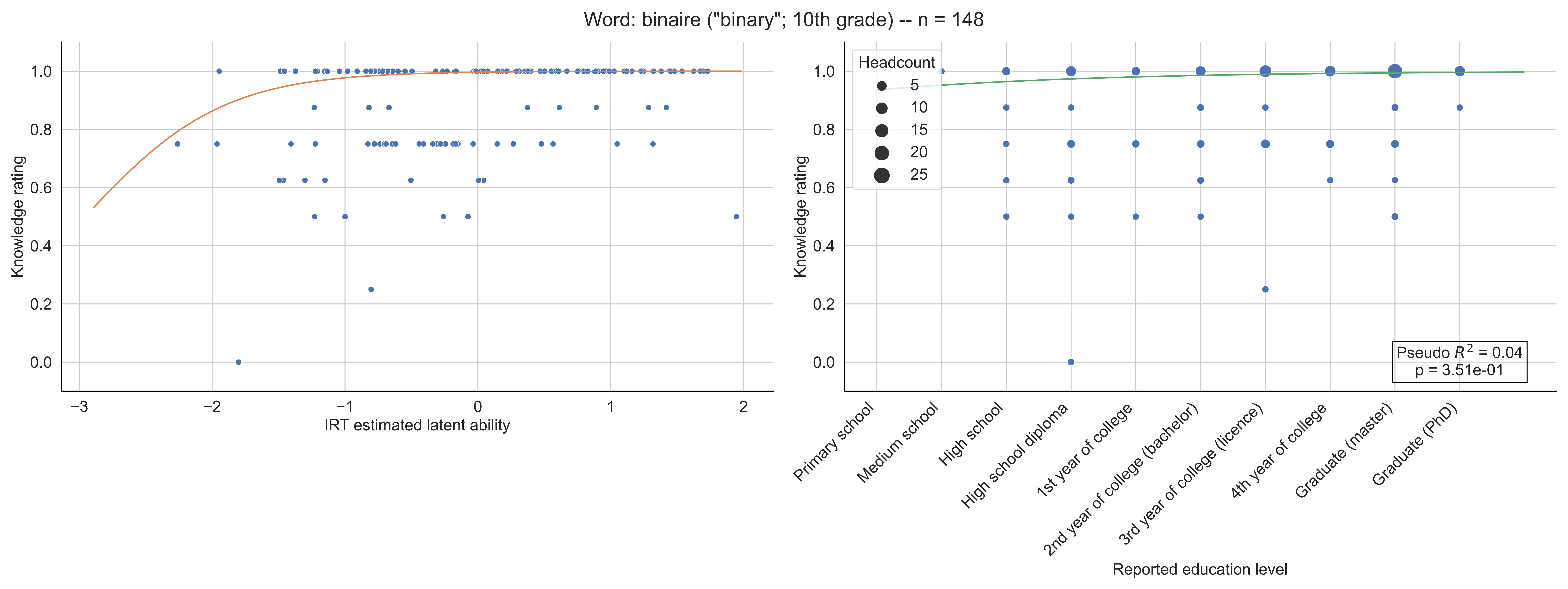

Word: binaire ("binary"; 10th grade) -- n = 148
Knowledge rating
IRT estimated latent ability
Headcount
5
10
15
20
25
Pseudo $R^2$ = 0.04
p = 3.51e-01
Primary school
Medium school
High school
High school diploma
1st year of college
2nd year of college (bachelor)
3rd year of college (licence)
4th year of college
Graduate (master)
Graduate (PhD)
Reported education level

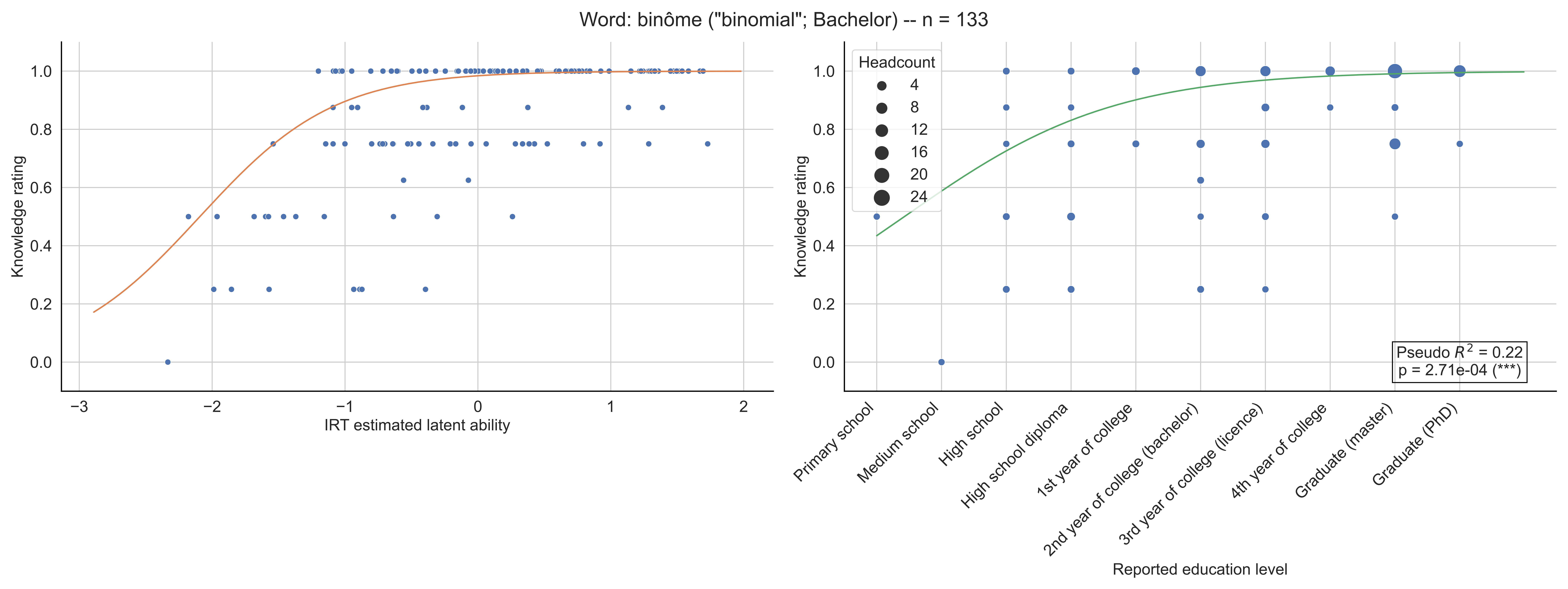
Word: binôme ("binomial"; Bachelor) -- n = 133
Knowledge rating
IRT estimated latent ability
Headcount
4
8
12
16
20
24
Pseudo $R^2$ = 0.22
p = 2.71e-04 (***)
Primary school
Medium school
High school
High school diploma
1st year of college
2nd year of college (bachelor)
3rd year of college (licence)
4th year of college
Graduate (master)
Graduate (PhD)
Reported education level

Word: bissectrice ("bisector"; Primary school) -- n = 136

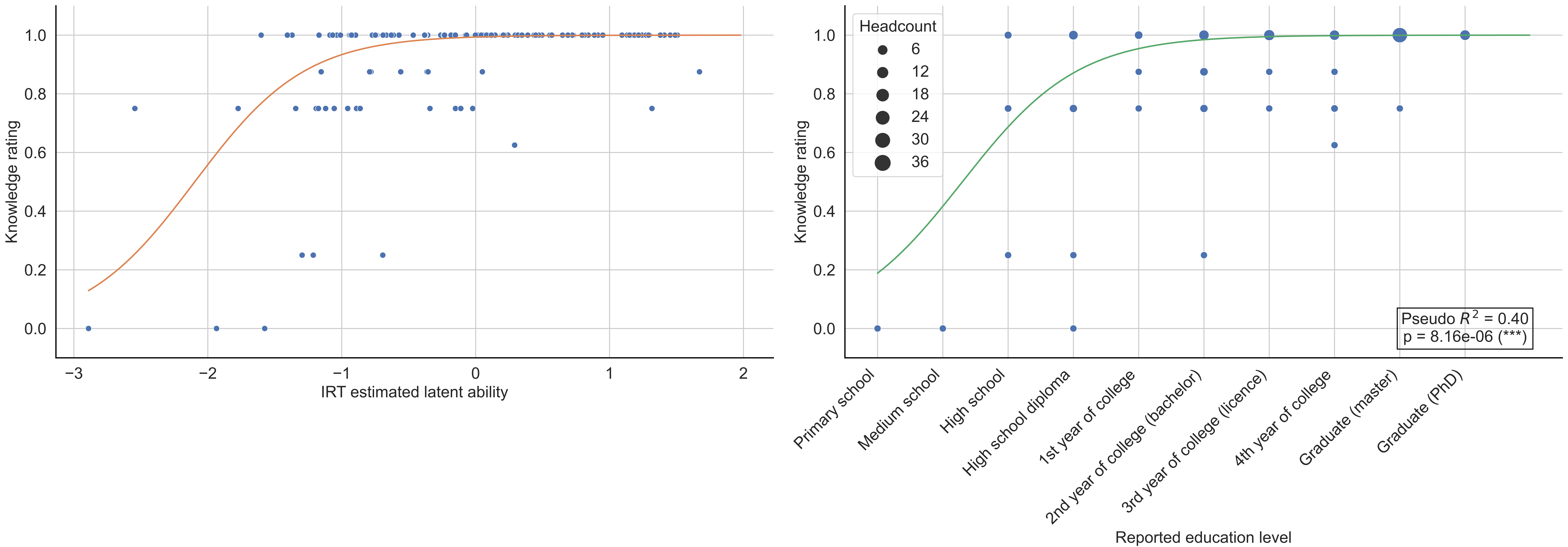

Word: booléen ("boolean"; 10th grade) -- n = 151

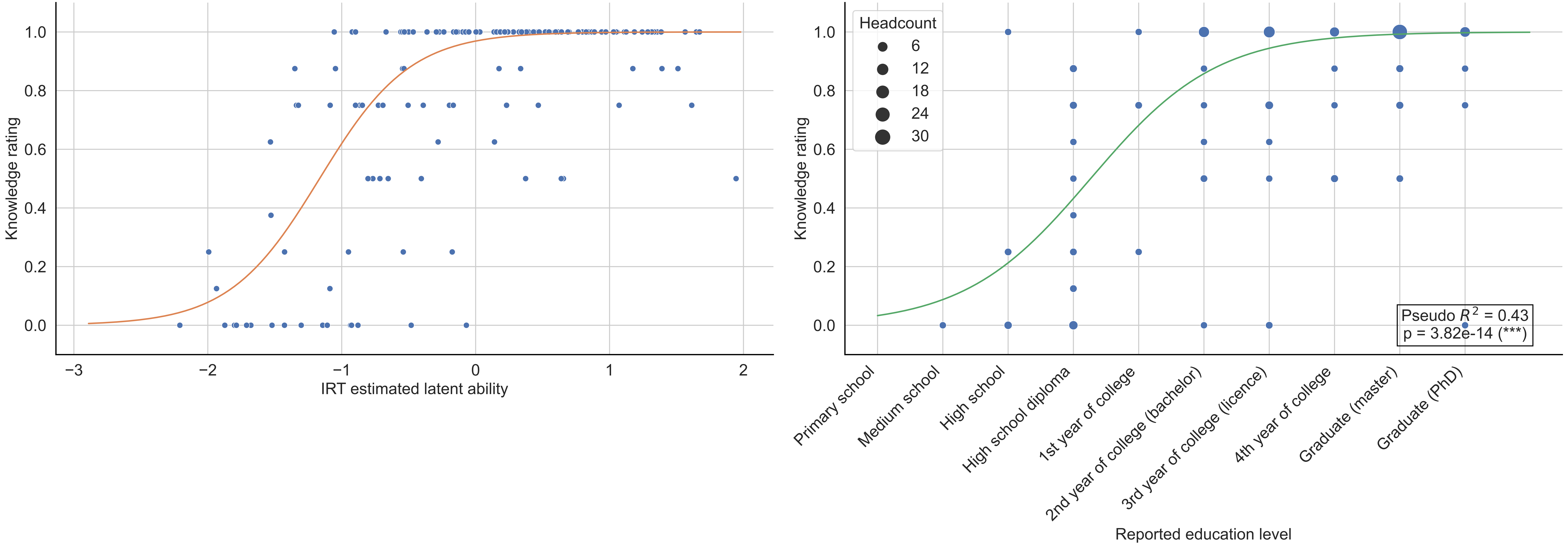

Word: borne ("bound"; 11-12th grade) -- n = 159

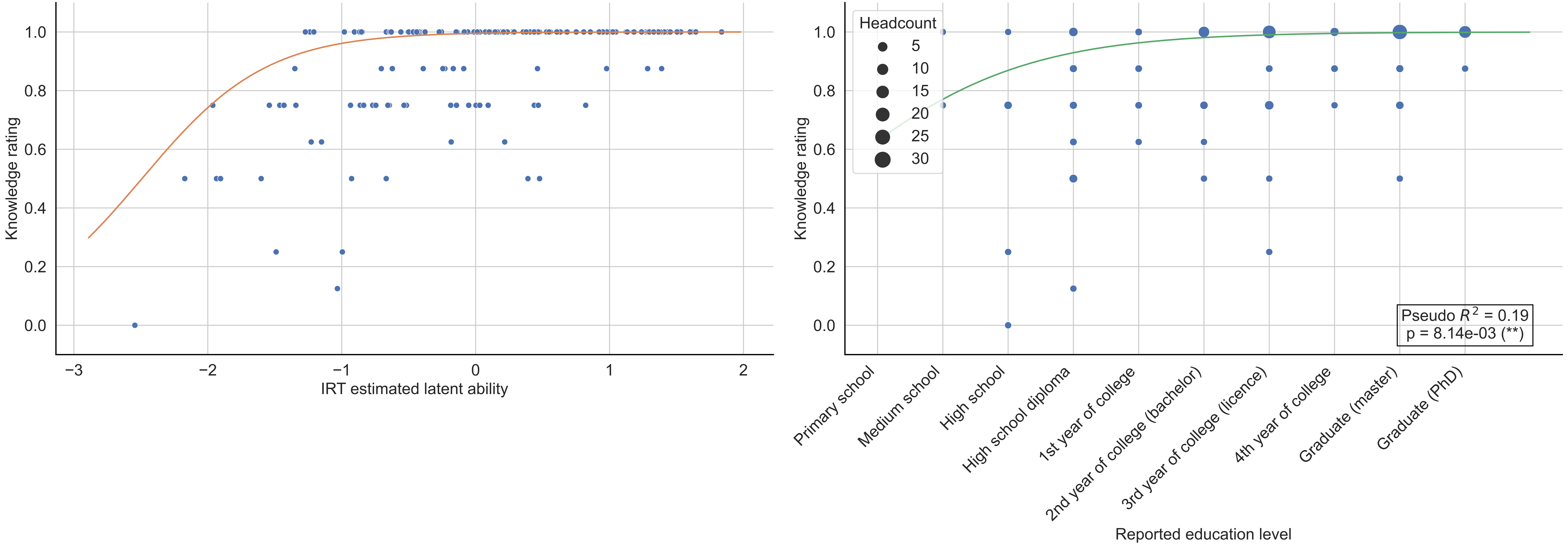

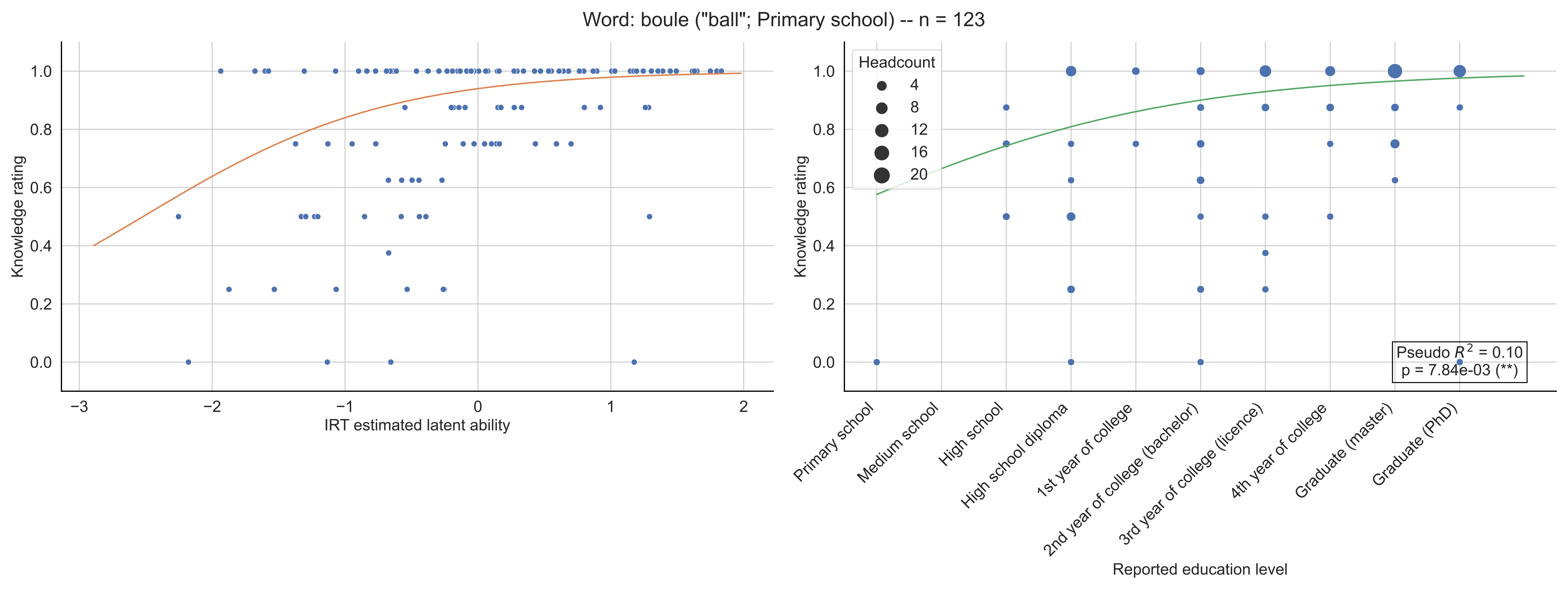

Word: boule ("ball"; Primary school) -- n = 123
Knowledge rating
IRT estimated latent ability
Headcount
4
8
12
16
20
Pseudo $R^2$ = 0.10
p = 7.84e-03 (**)
Primary school
Medium school
High school
High school diploma
1st year of college
2nd year of college (bachelor)
3rd year of college (licence)
4th year of college
Graduate (master)
Graduate (PhD)
Reported education level

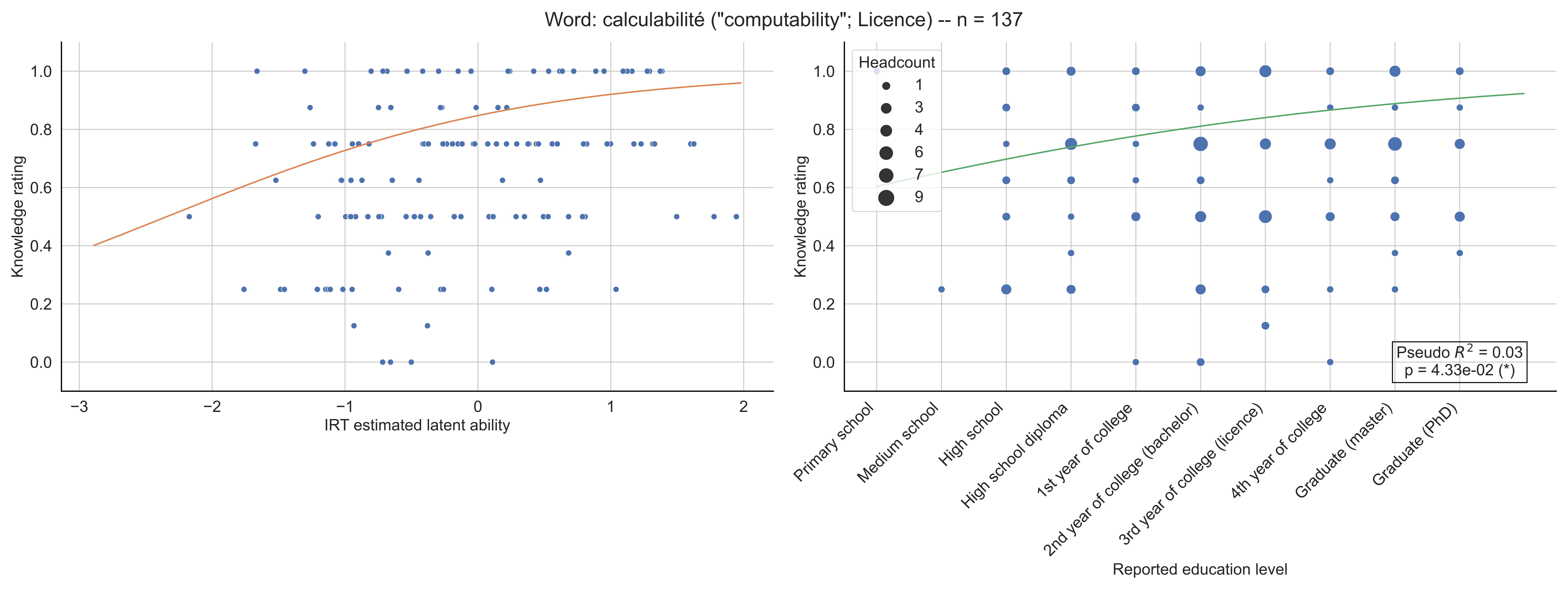
Word: calculabilité ("computability"; Licence) -- n = 137
Knowledge rating
IRT estimated latent ability
Headcount
1
3
4
6
7
9
Primary school
Medium school
High school
High school diploma
1st year of college
2nd year of college (bachelor)
3rd year of college (licence)
4th year of college
Graduate (master)
Graduate (PhD)
Reported education level
Pseudo $R^2$ = 0.03
p = 4.33e-02 (*)

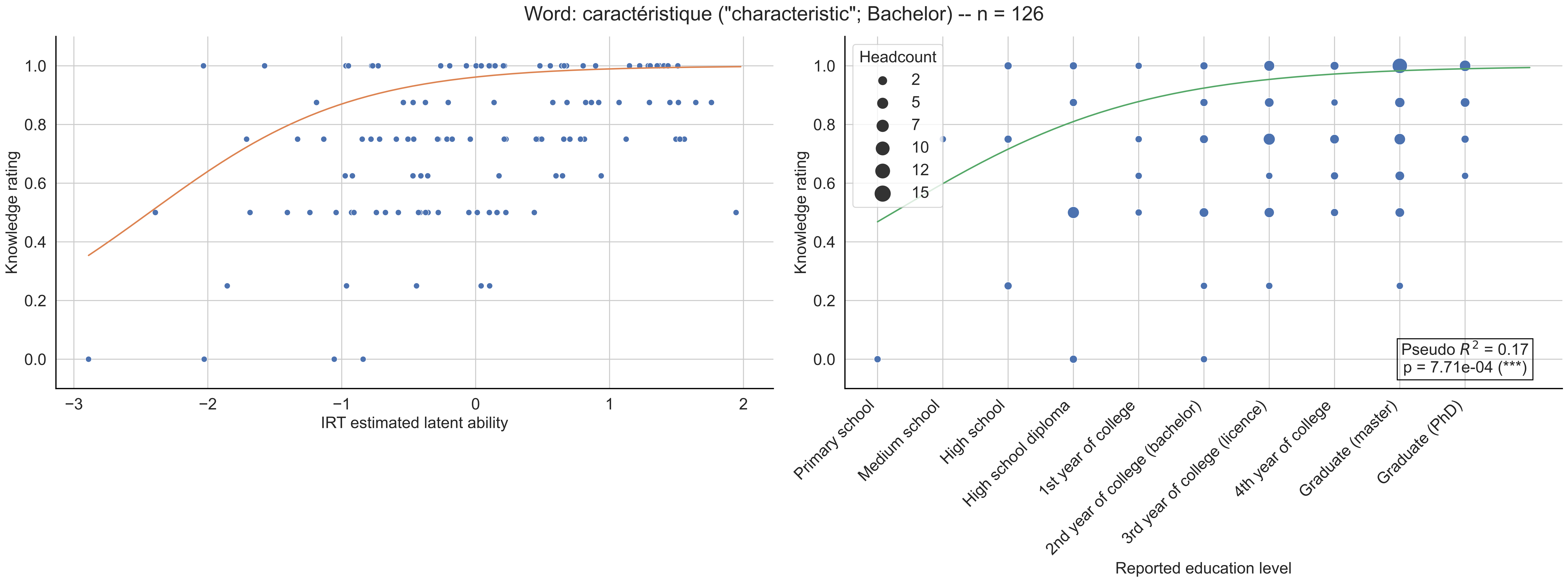

Word: caractéristique ("characteristic"; Bachelor) -- n = 126
Knowledge rating
IRT estimated latent ability
Headcount
2
5
7
10
12
15
Pseudo $R^2$ = 0.17
p = 7.71e-04 (***)
Primary school
Medium school
High school
High school diploma
1st year of college
2nd year of college (bachelor)
3rd year of college (licence)
4th year of college
Graduate (master)
Graduate (PhD)
Reported education level

Word: cardinal ("cardinal"; Bachelor) -- n = 157

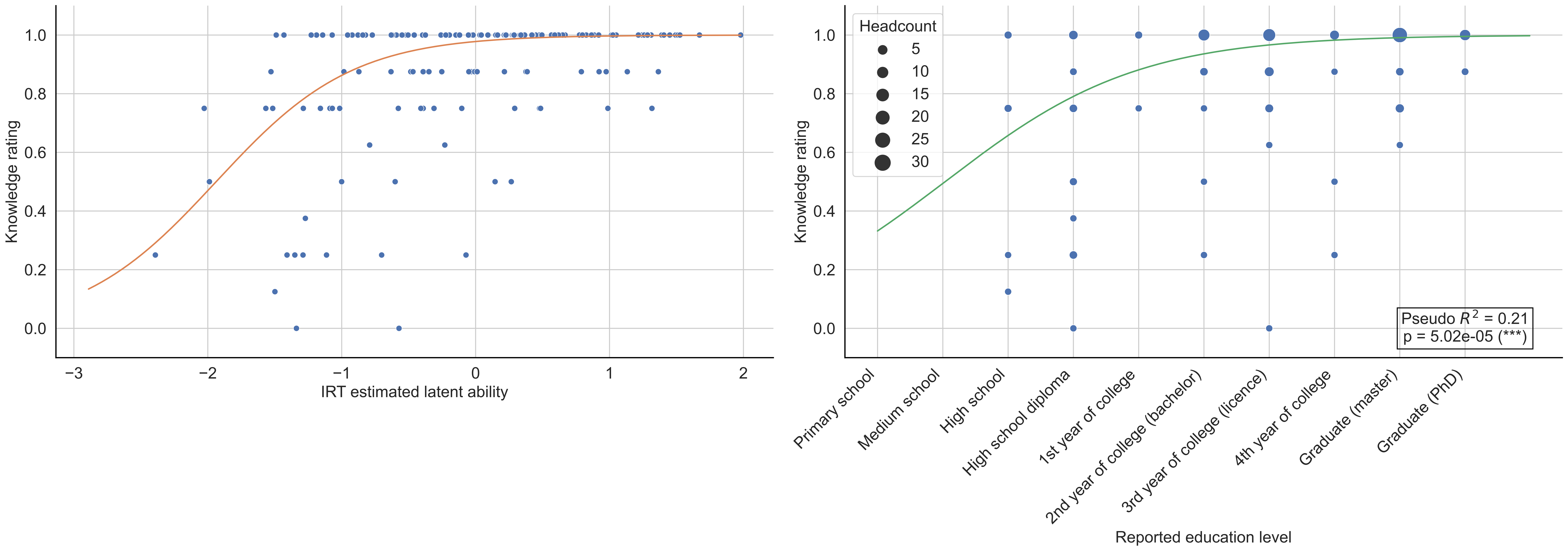

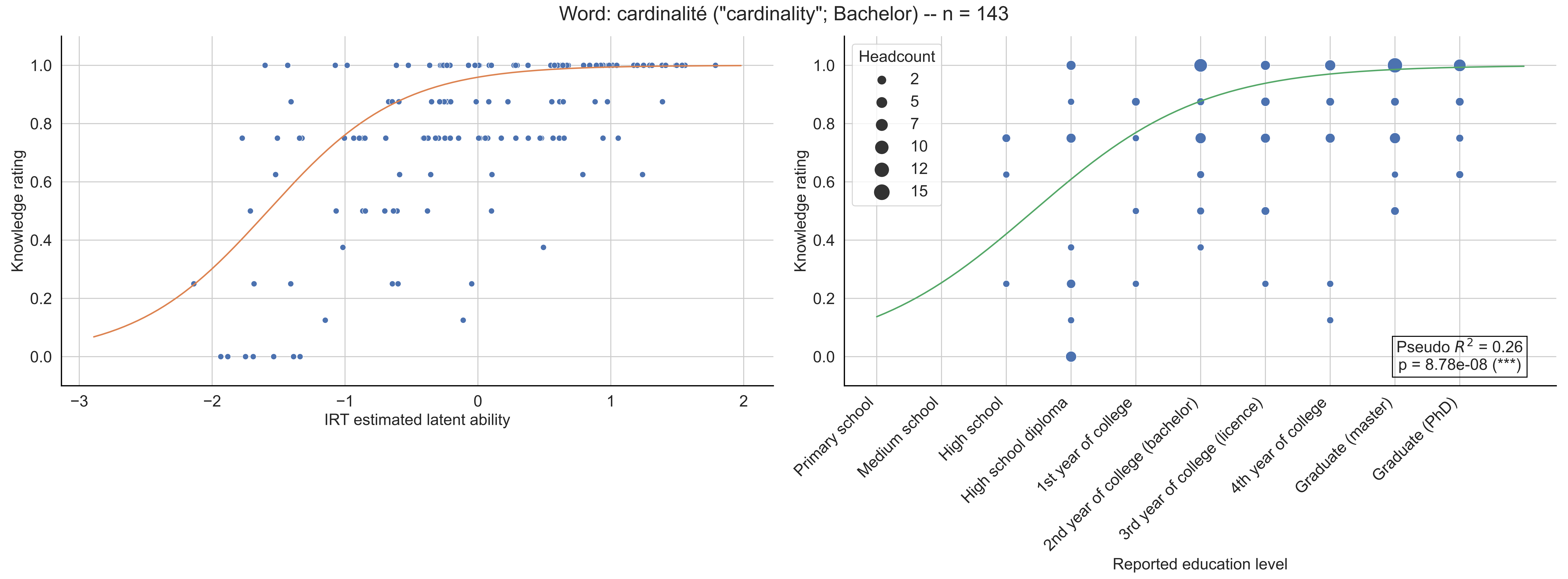

Word: cardinalité ("cardinality"; Bachelor) -- n = 143
Knowledge rating
IRT estimated latent ability
Headcount
2
5
7
10
12
15
Pseudo $R^2$ = 0.26
p = 8.78e-08 (***)
Primary school
Medium school
High school
High school diploma
1st year of college
2nd year of college (bachelor)
3rd year of college (licence)
4th year of college
Graduate (master)
Graduate (PhD)
Reported education level

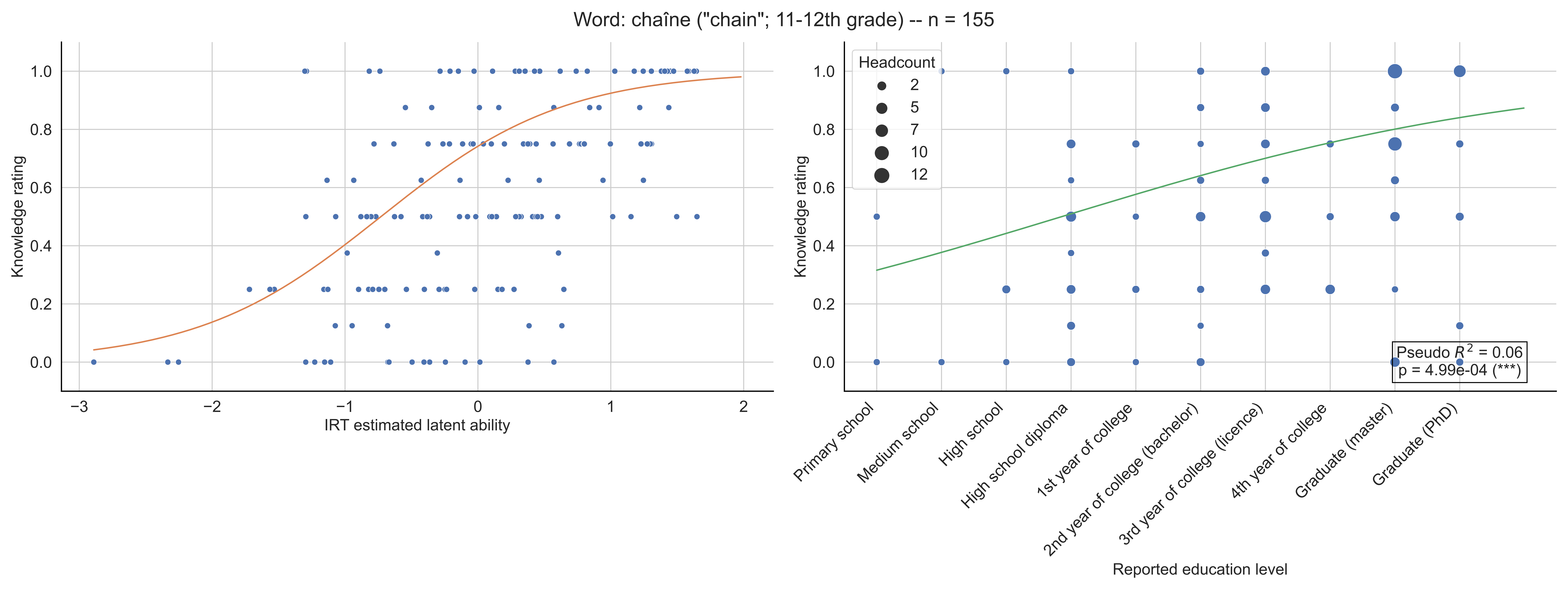

Word: chaîne ("chain"; 11-12th grade) -- n = 155
Knowledge rating
IRT estimated latent ability
Headcount
2
5
7
10
12
Primary school
Medium school
High school
High school diploma
1st year of college
2nd year of college (bachelor)
3rd year of college (licence)
4th year of college
Graduate (master)
Graduate (PhD)
Reported education level
Pseudo $R^2$ = 0.06
p = 4.99e-04 (***)

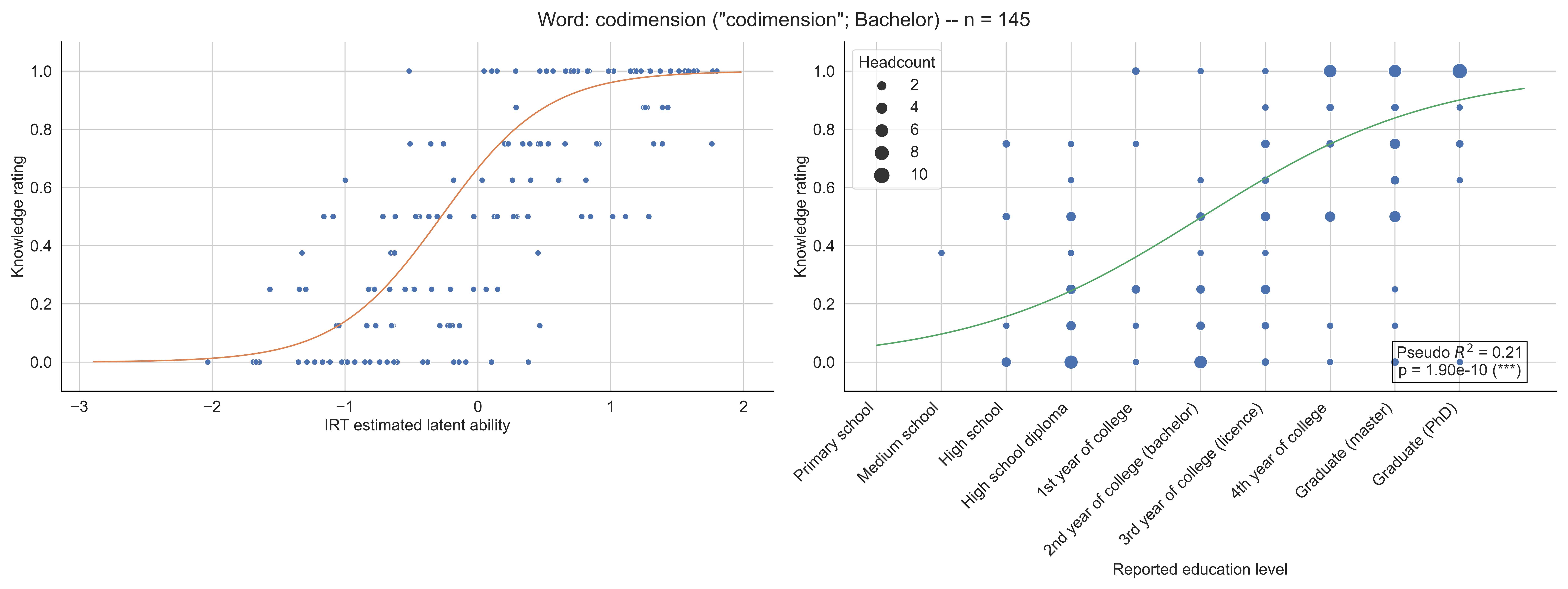

Word: codimension ("codimension"; Bachelor) -- n = 145
Knowledge rating
IRT estimated latent ability
Headcount
2
4
6
8
10
Pseudo $R^2$ = 0.21
p = 1.90e-10 (***)
Primary school
Medium school
High school
High school diploma
1st year of college
2nd year of college (bachelor)
3rd year of college (licence)
4th year of college
Graduate (master)
Graduate (PhD)
Reported education level

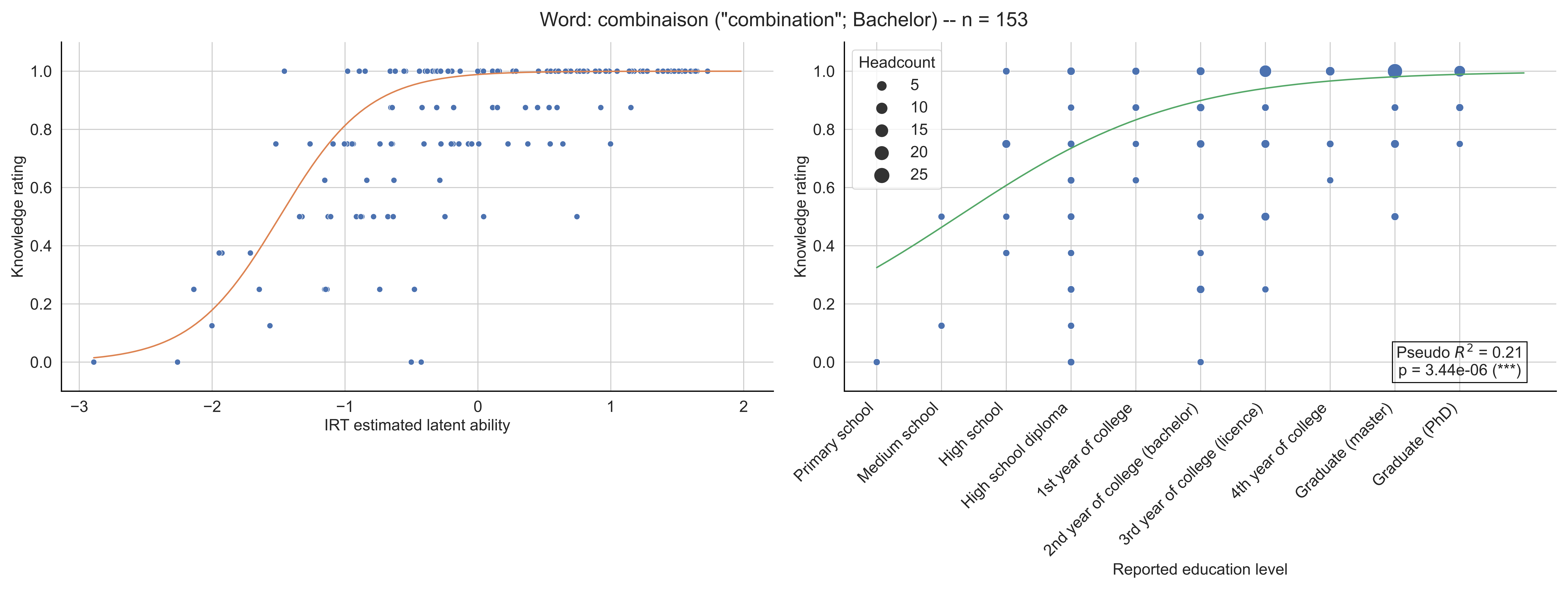

Word: combinaison ("combination"; Bachelor) -- n = 153
Knowledge rating
IRT estimated latent ability
Headcount
5
10
15
20
25
Pseudo $R^2$ = 0.21
p = 3.44e-06 (***)
Primary school
Medium school
High school
High school diploma
1st year of college
2nd year of college (bachelor)
3rd year of college (licence)
4th year of college
Graduate (master)
Graduate (PhD)
Reported education level

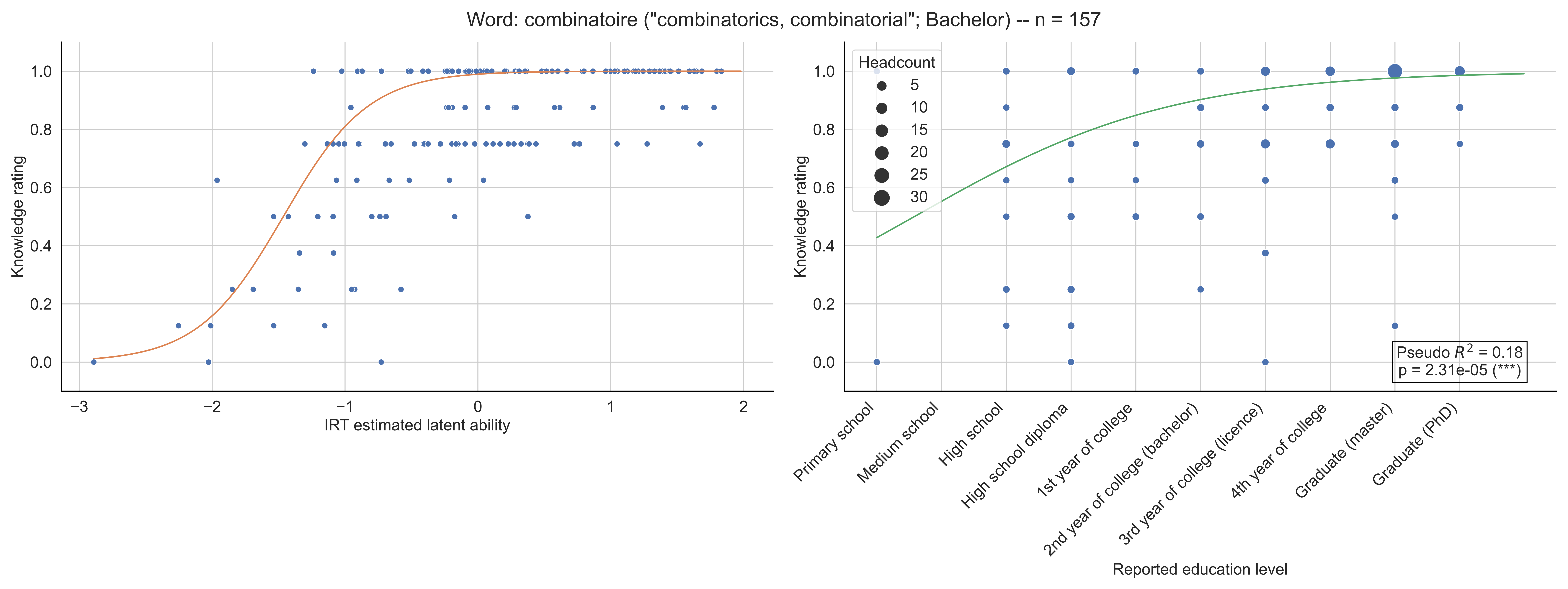

Word: combinatoire ("combinatorics, combinatorial"; Bachelor) -- n = 157
Knowledge rating
IRT estimated latent ability
Headcount
5
10
15
20
25
30
Pseudo $R^2$ = 0.18
p = 2.31e-05 (***)
Primary school
Medium school
High school
High school diploma
1st year of college
2nd year of college (bachelor)
3rd year of college (licence)
4th year of college
Graduate (master)
Graduate (PhD)
Reported education level

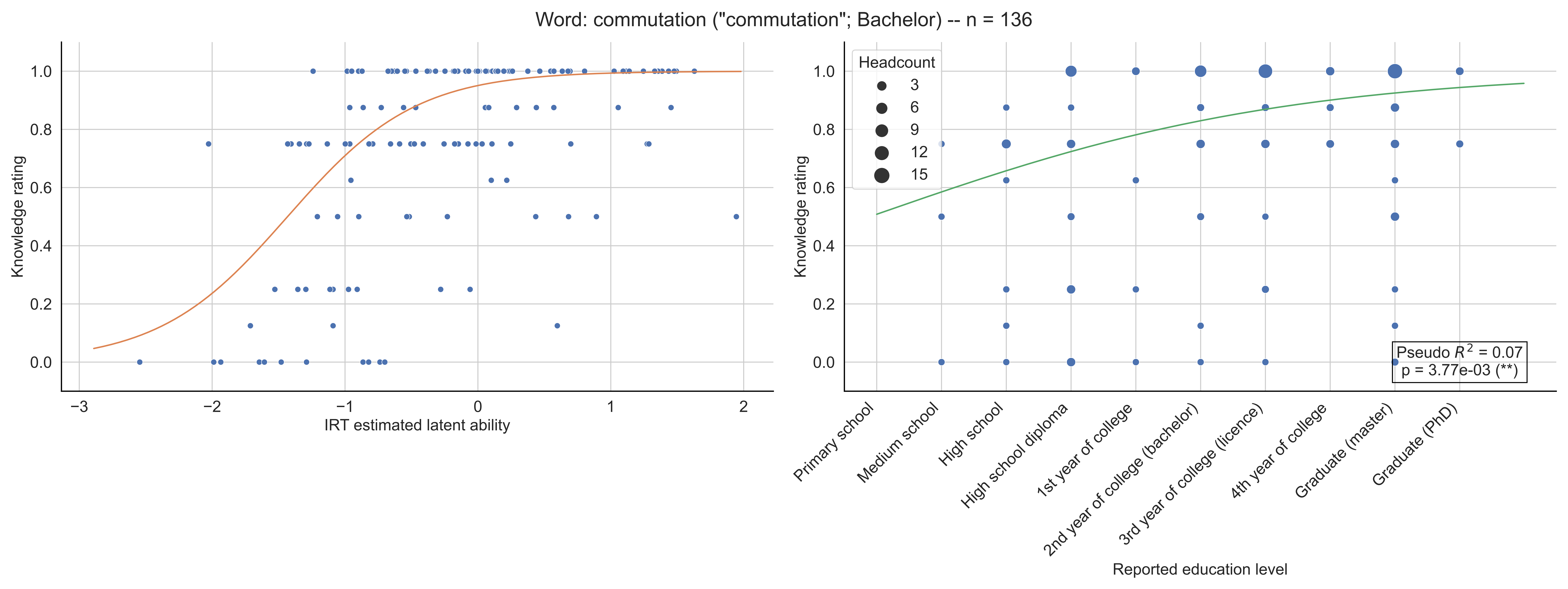
Word: commutation ("commutation"; Bachelor) -- n = 136
Knowledge rating
IRT estimated latent ability
Headcount
3
6
9
12
15
Pseudo $R^2$ = 0.07
p = 3.77e-03 (**)
Primary school
Medium school
High school
High school diploma
1st year of college
2nd year of college (bachelor)
3rd year of college (licence)
4th year of college
Graduate (master)
Graduate (PhD)
Reported education level

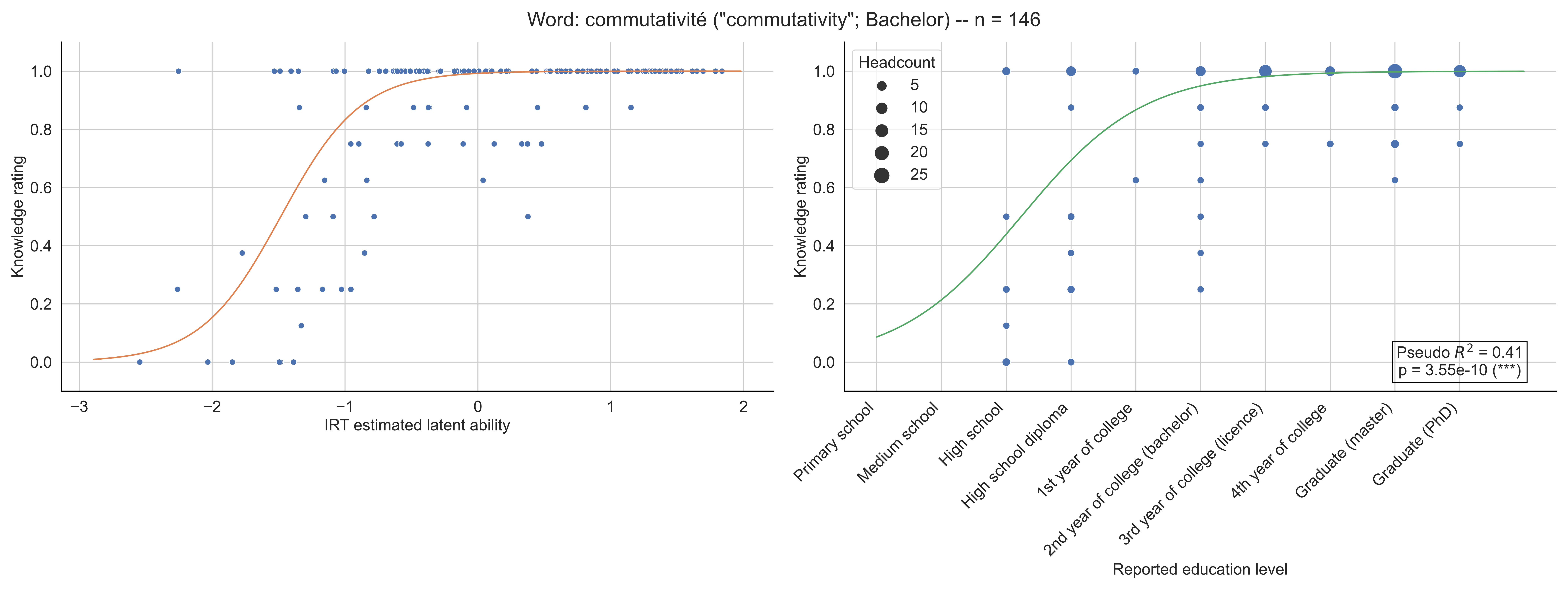

Word: commutativité ("commutativity"; Bachelor) -- n = 146
Knowledge rating
IRT estimated latent ability
Headcount
5
10
15
20
25
Pseudo $R^2$ = 0.41
p = 3.55e-10 (***)
Primary school
Medium school
High school
High school diploma
1st year of college
2nd year of college (bachelor)
3rd year of college (licence)
4th year of college
Graduate (master)
Graduate (PhD)
Reported education level

Word: compacité ("compactness"; Bachelor) -- n = 139

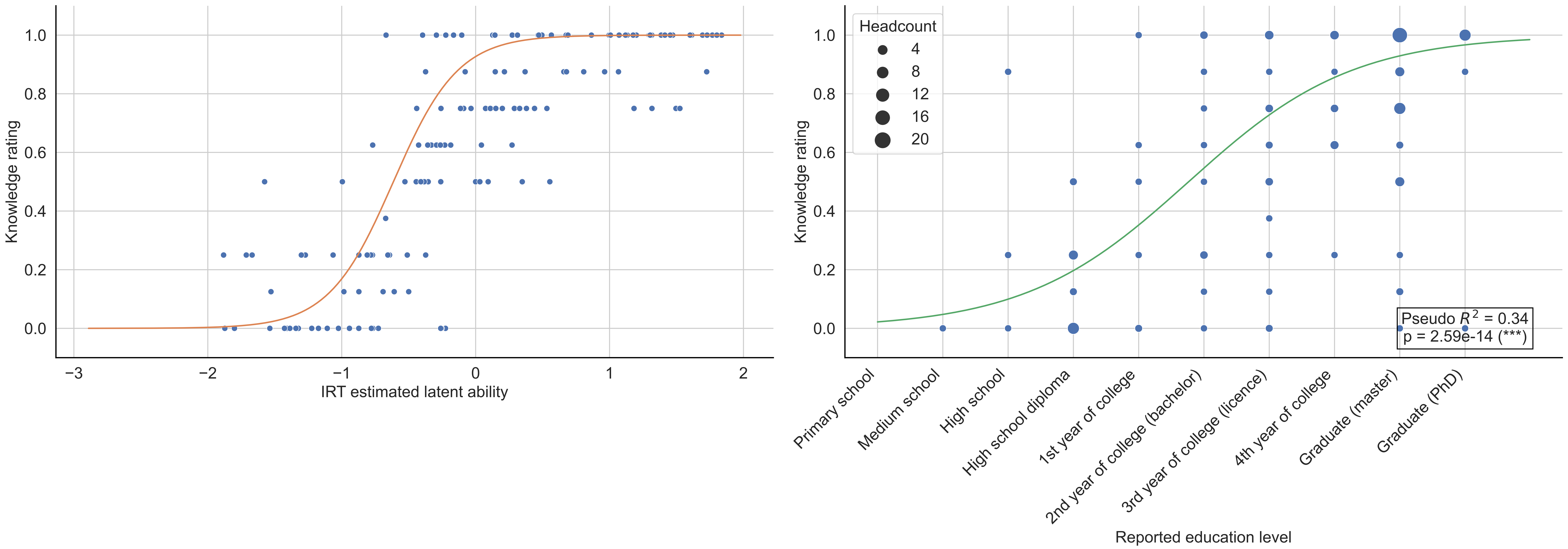

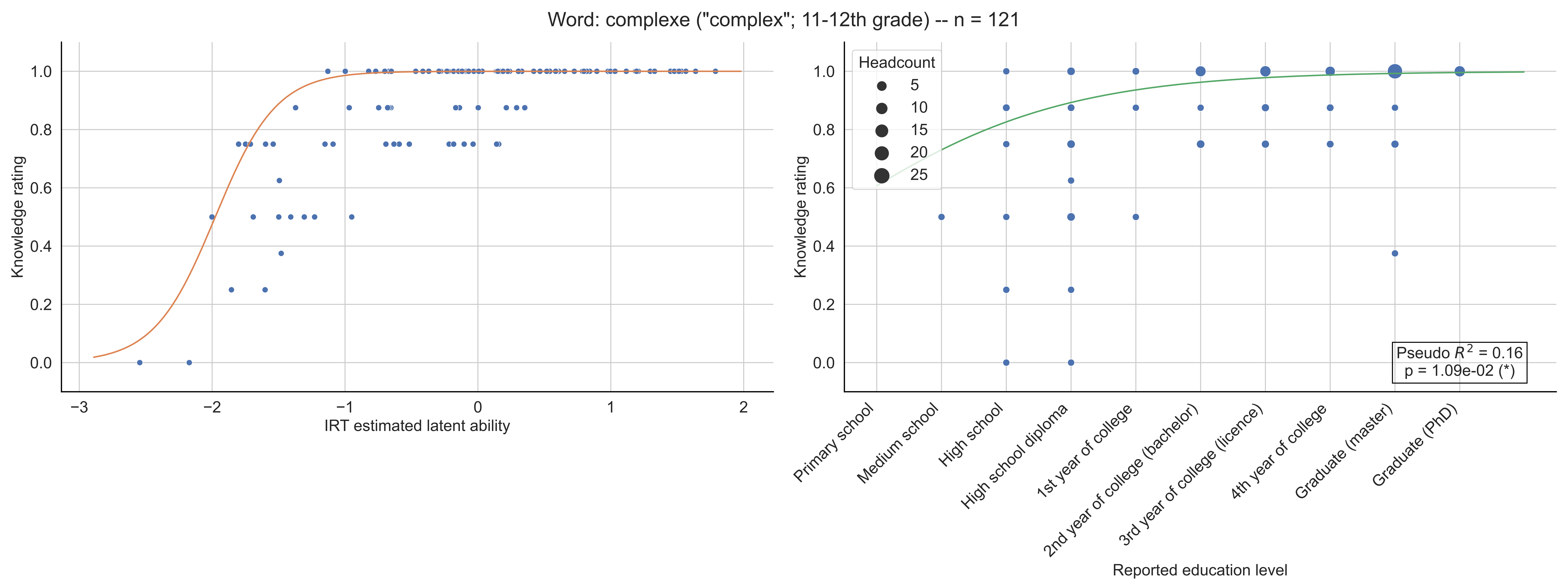

Word: complexe ("complex"; 11-12th grade) -- n = 121
Knowledge rating
IRT estimated latent ability
Headcount
5
10
15
20
25
Pseudo $R^2$ = 0.16
p = 1.09e-02 (*)
Primary school
Medium school
High school
High school diploma
1st year of college
2nd year of college (bachelor)
3rd year of college (licence)
4th year of college
Graduate (master)
Graduate (PhD)
Reported education level

Word: complexité ("complexity"; Licence) -- n = 139

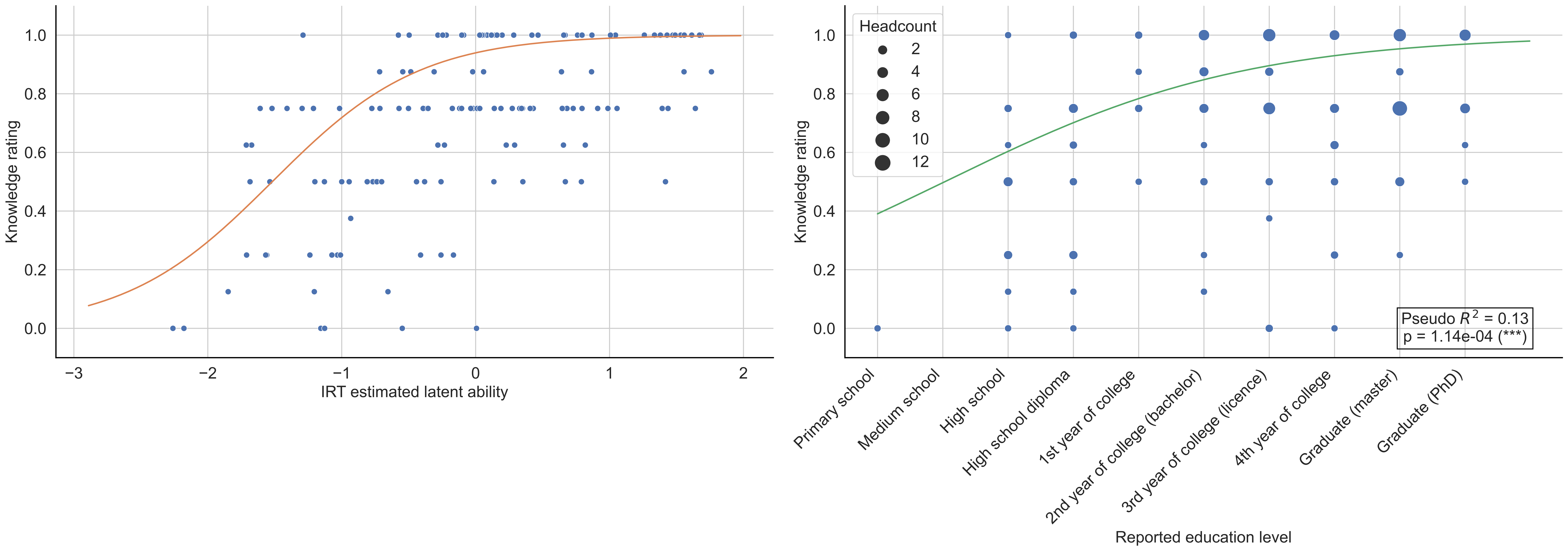

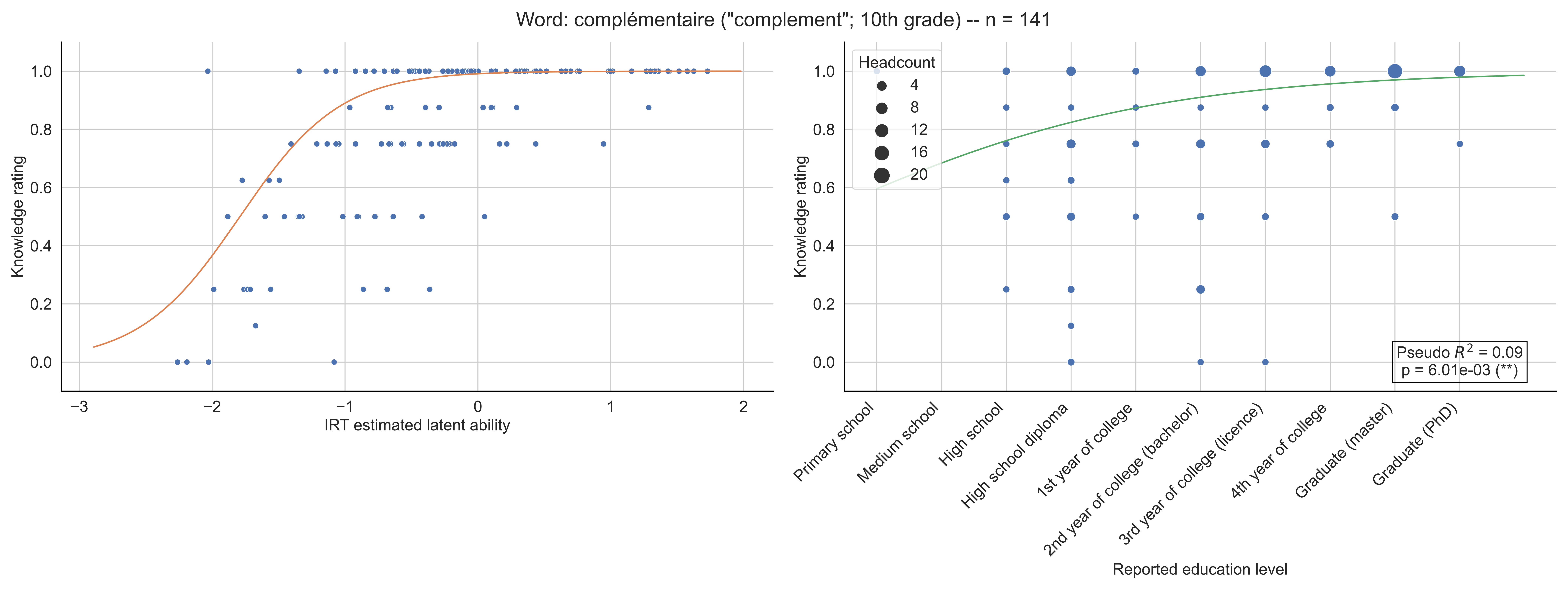
Word: complémentaire ("complement"; 10th grade) -- n = 141
Knowledge rating
IRT estimated latent ability
Headcount
4
8
12
16
20
Pseudo $R^2$ = 0.09
p = 6.01e-03 (**)
Primary school
Medium school
High school
High school diploma
1st year of college
2nd year of college (bachelor)
3rd year of college (licence)
4th year of college
Graduate (master)
Graduate (PhD)
Reported education level

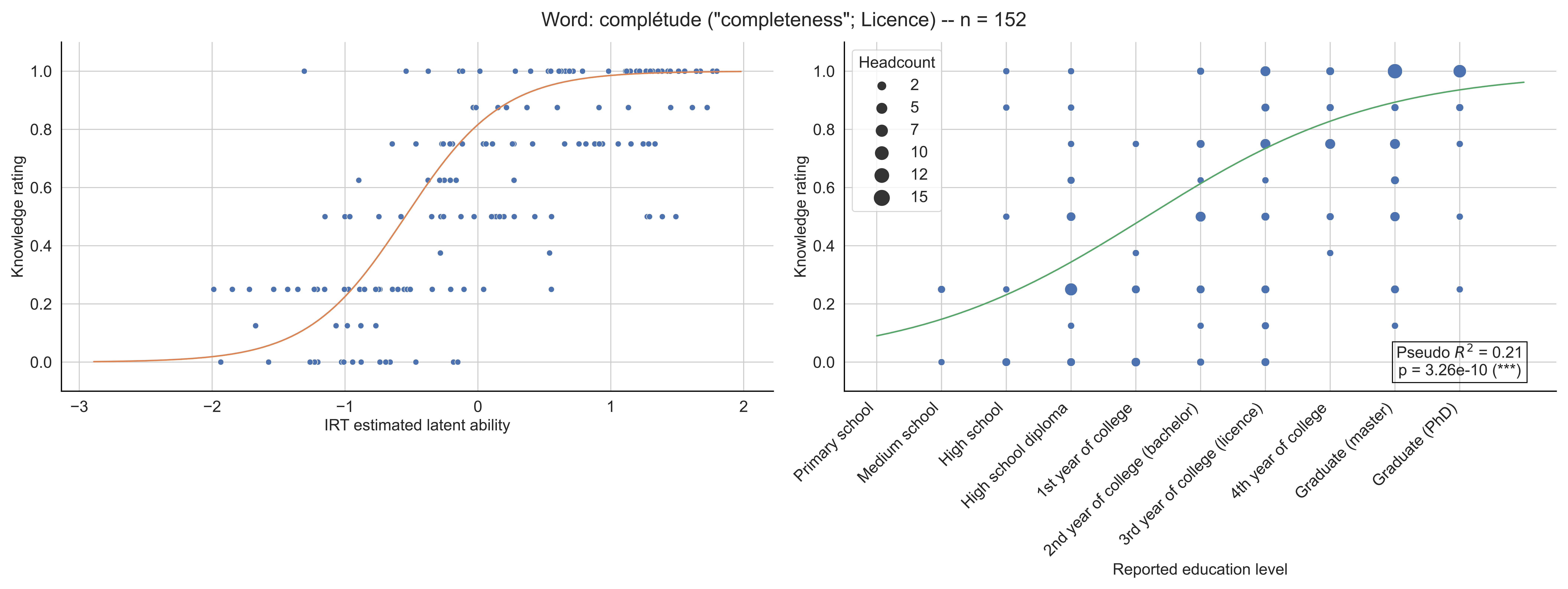

Word: complétude ("completeness"; Licence) -- n = 152
Knowledge rating
IRT estimated latent ability
Headcount
2
5
7
10
12
15
Pseudo $R^2$ = 0.21
p = 3.26e-10 (***)
Primary school
Medium school
High school
High school diploma
1st year of college
2nd year of college (bachelor)
3rd year of college (licence)
4th year of college
Graduate (master)
Graduate (PhD)
Reported education level

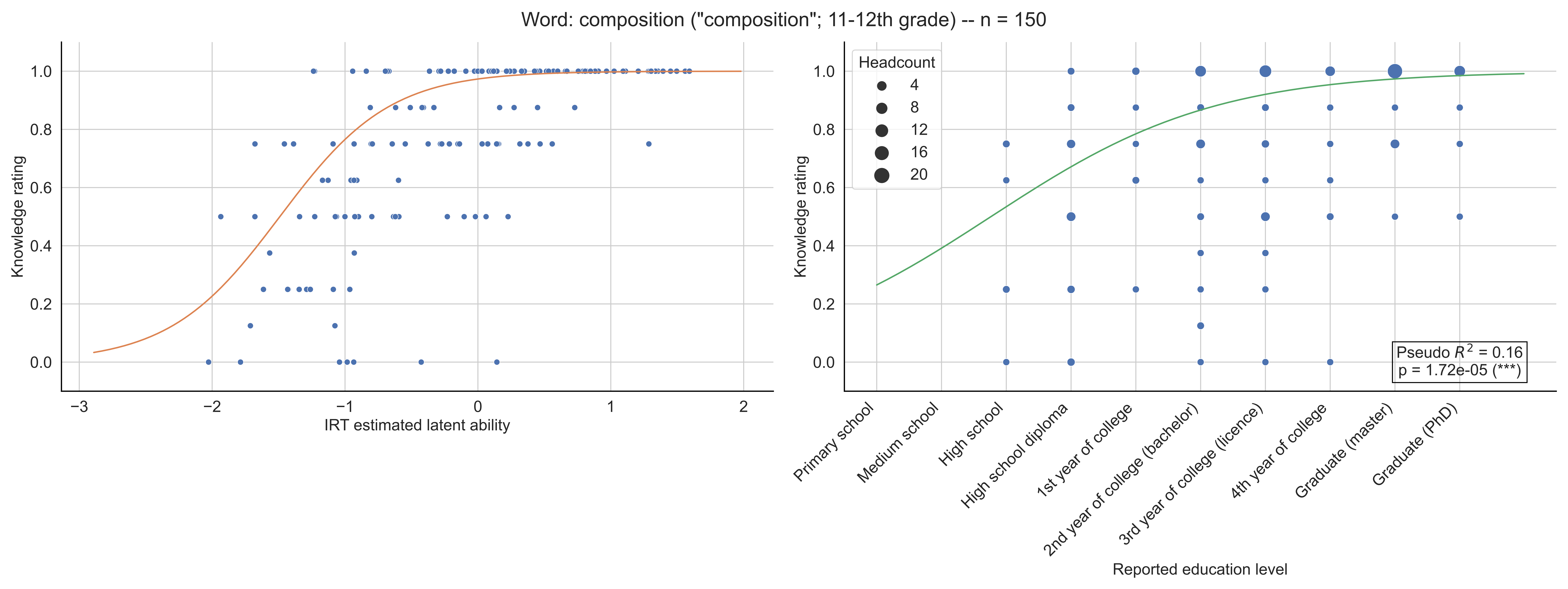

Word: composition ("composition"; 11-12th grade) -- n = 150
Knowledge rating
IRT estimated latent ability
Headcount
4
8
12
16
20
Pseudo $R^2$ = 0.16
p = 1.72e-05 (***)
Primary school
Medium school
High school
High school diploma
1st year of college
2nd year of college (bachelor)
3rd year of college (licence)
4th year of college
Graduate (master)
Graduate (PhD)
Reported education level

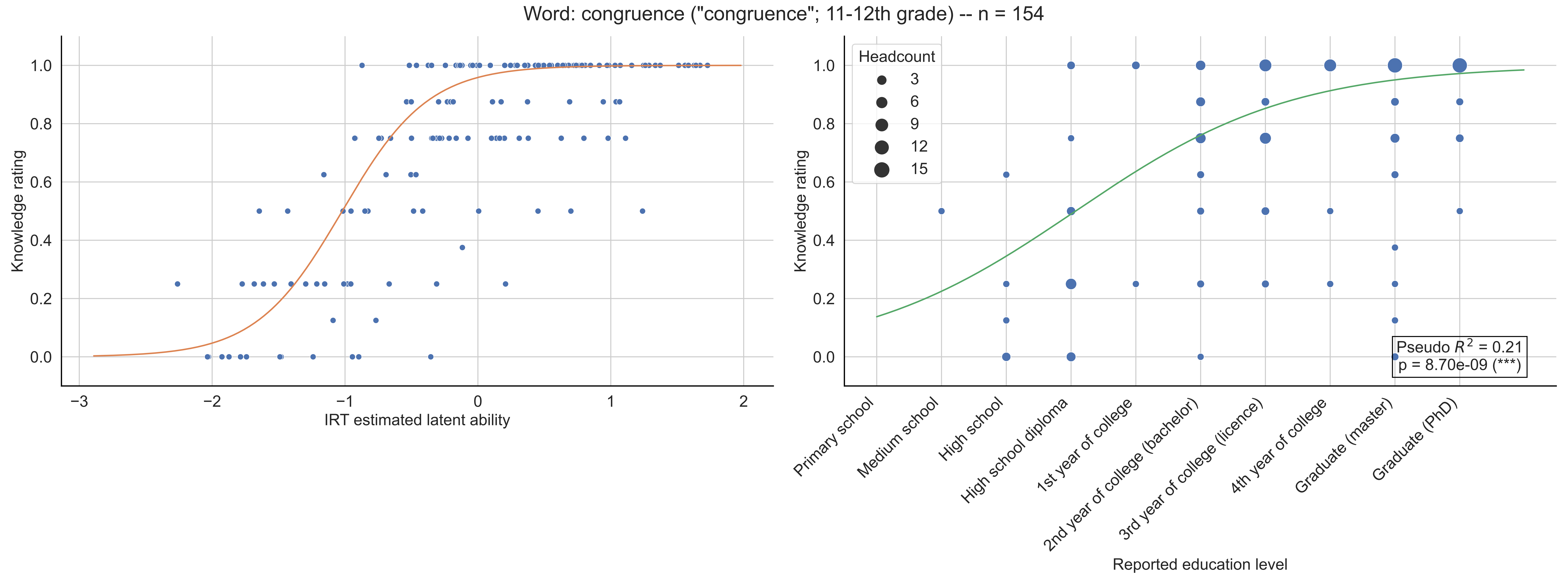

Word: congruence ("congruence"; 11-12th grade) -- n = 154
Knowledge rating
IRT estimated latent ability
Headcount
3
6
9
12
15
Pseudo $R^2$ = 0.21
p = 8.70e-09 (***)
Primary school
Medium school
High school
High school diploma
1st year of college
2nd year of college (bachelor)
3rd year of college (licence)
4th year of college
Graduate (master)
Graduate (PhD)
Reported education level

Word: conique ("conic"; Licence) -- n = 130

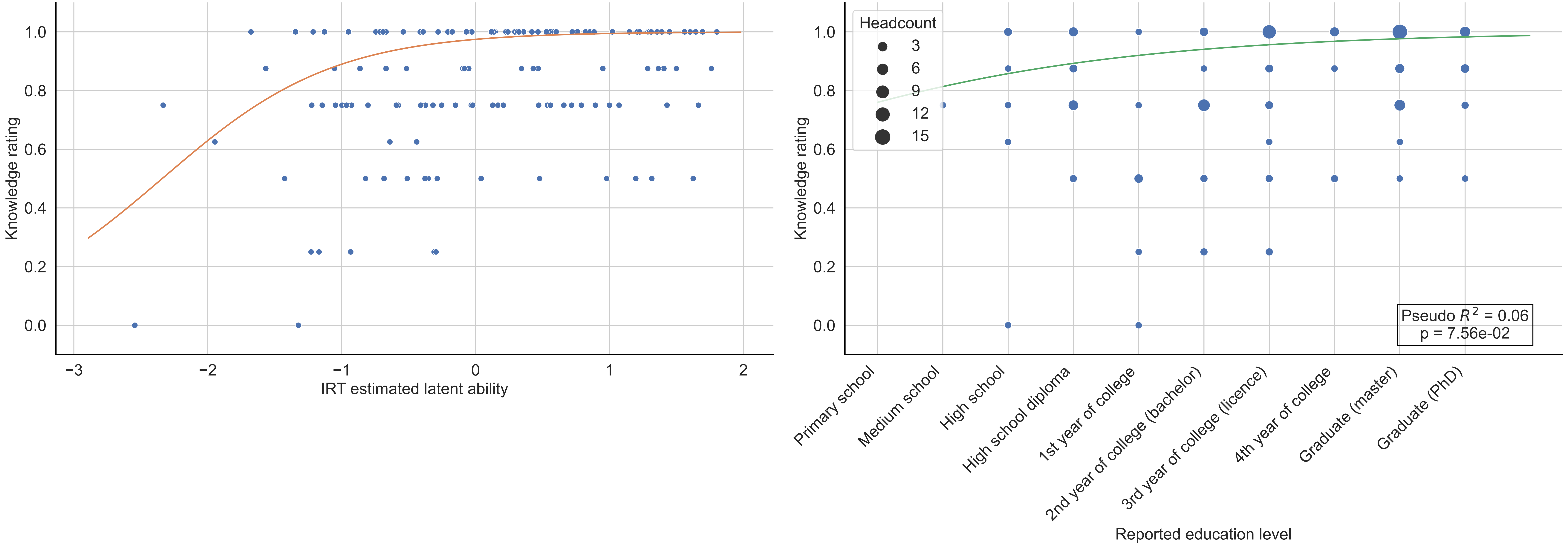

Word: connexité ("connectedness"; Bachelor) -- n = 122

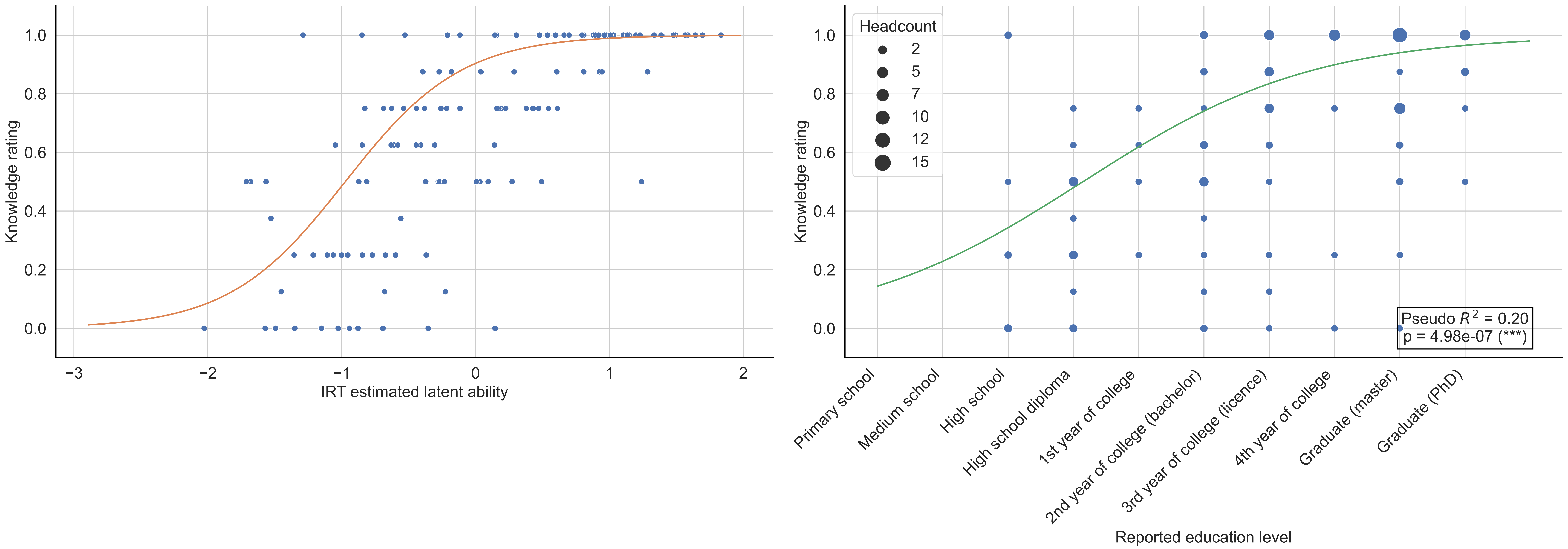

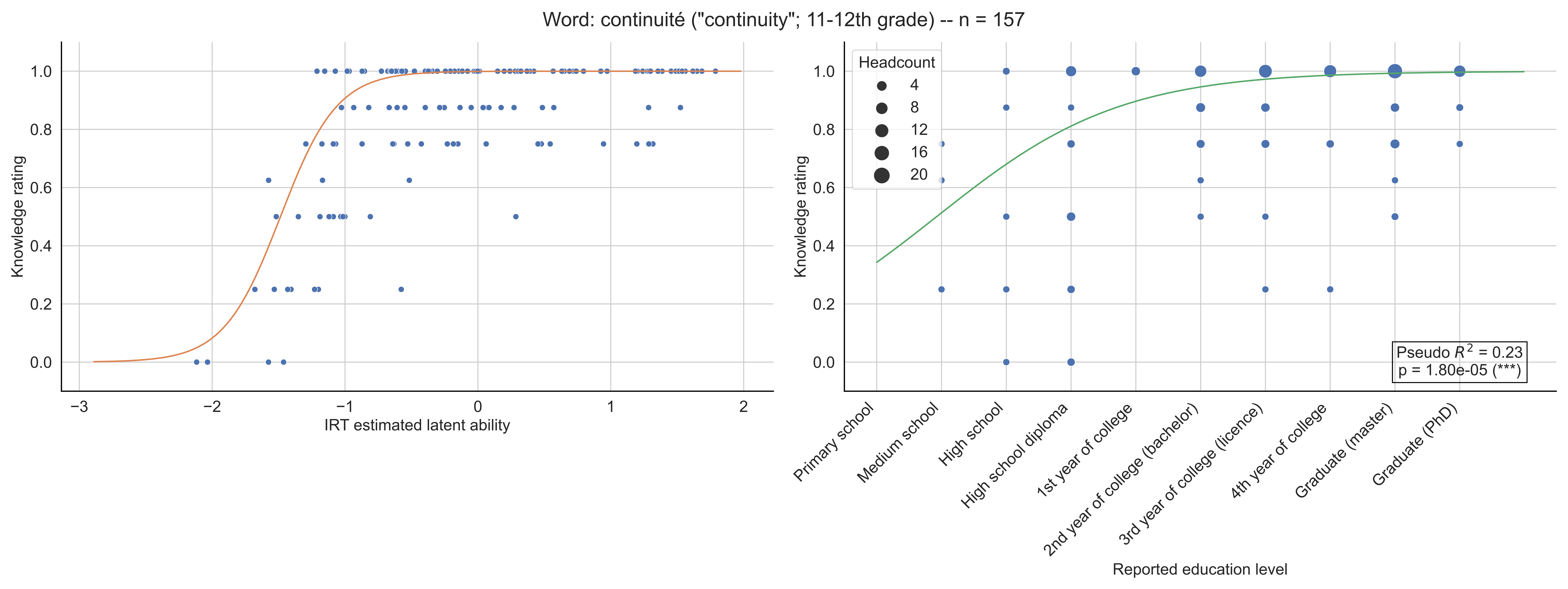
Word: continuité ("continuity"; 11-12th grade) -- n = 157
Knowledge rating
IRT estimated latent ability
Headcount
4
8
12
16
20
Pseudo $R^2$ = 0.23
p = 1.80e-05 (***)
Primary school
Medium school
High school
High school diploma
1st year of college
2nd year of college (bachelor)
3rd year of college (licence)
4th year of college
Graduate (master)
Graduate (PhD)
Reported education level

Word: convexité ("convexity"; 11-12th grade) -- n = 141

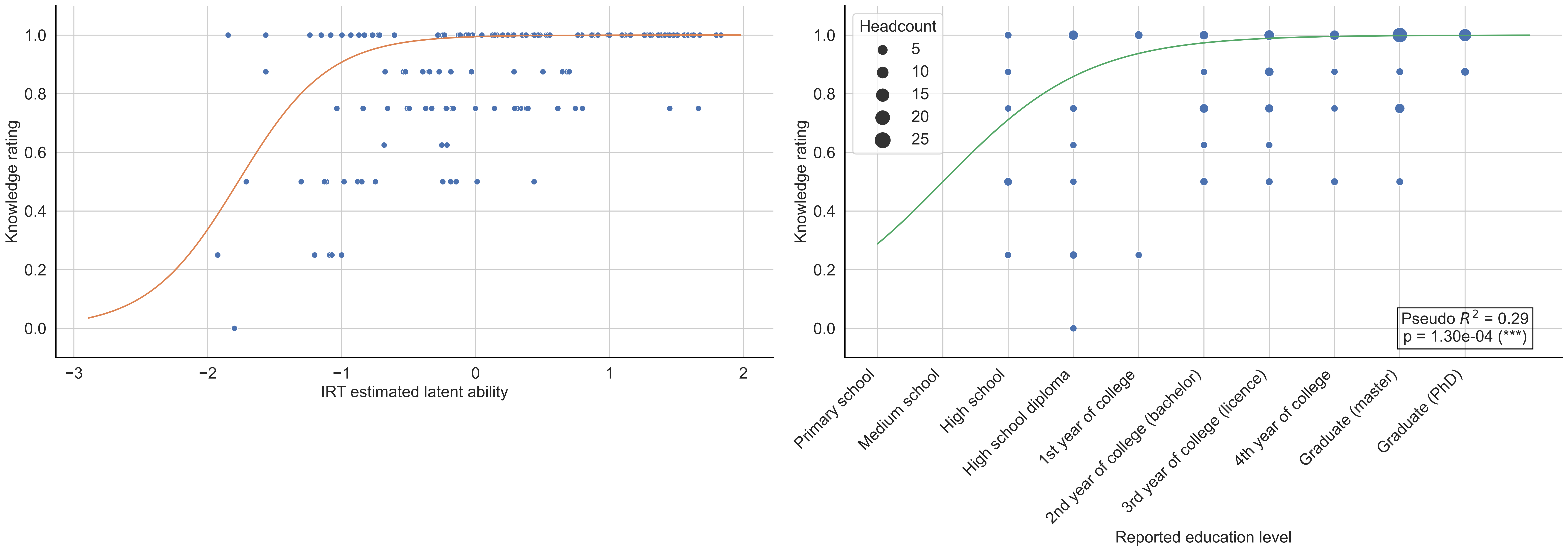

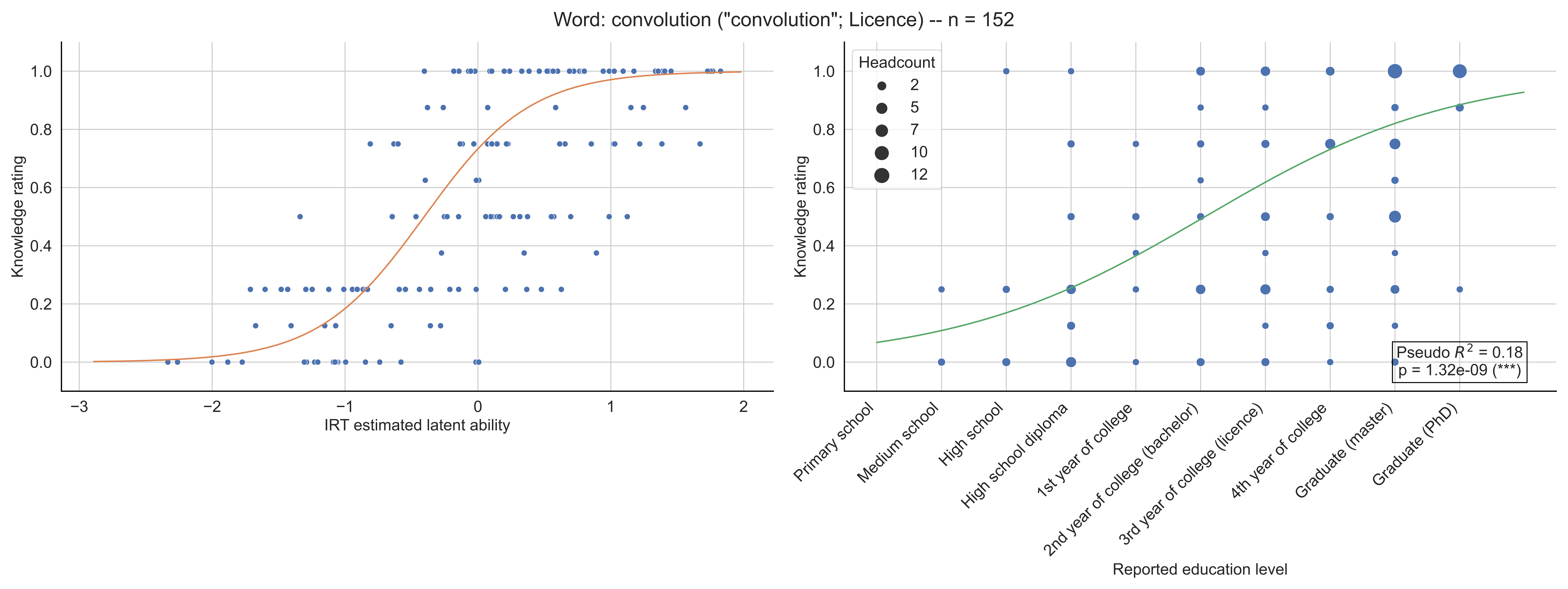

Word: convolution ("convolution"; Licence) -- n = 152
Knowledge rating
IRT estimated latent ability
Headcount
2
5
7
10
12
Pseudo $R^2$ = 0.18
p = 1.32e-09 (***)
Primary school
Medium school
High school
High school diploma
1st year of college
2nd year of college (bachelor)
3rd year of college (licence)
4th year of college
Graduate (master)
Graduate (PhD)
Reported education level

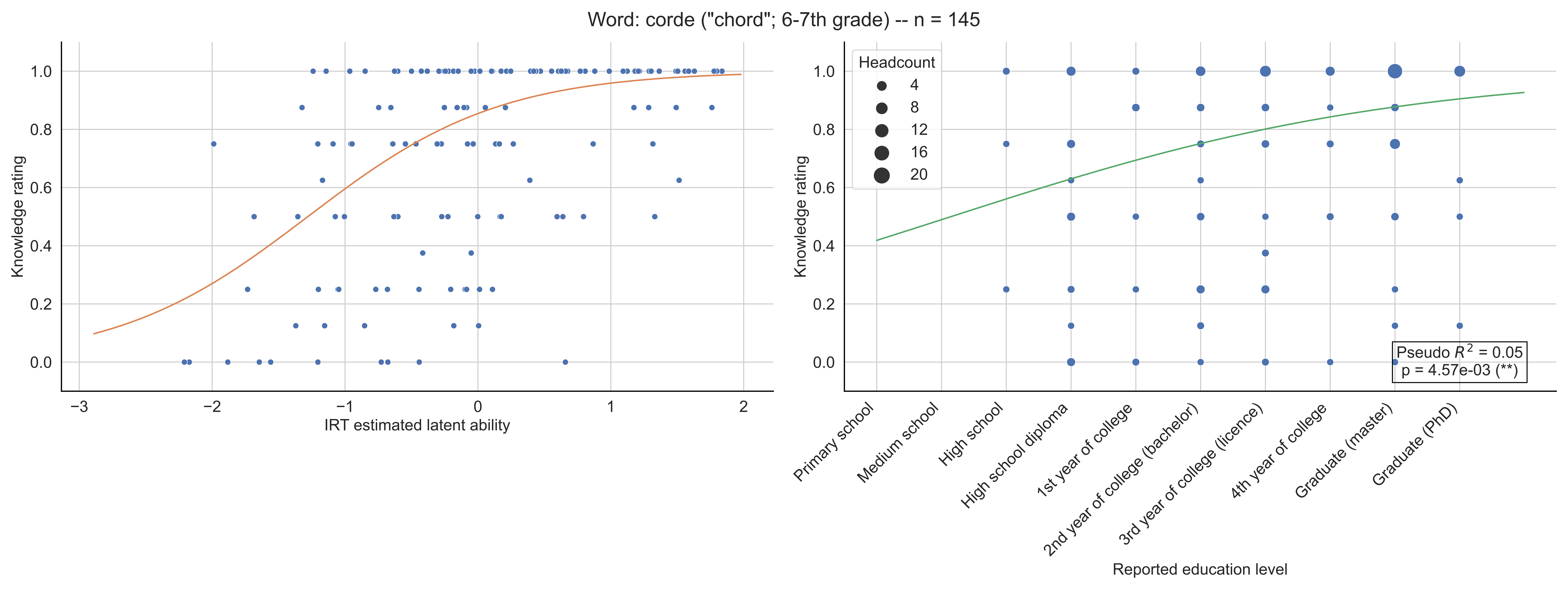
Word: corde ("chord"; 6-7th grade) -- n = 145
Knowledge rating
IRT estimated latent ability
Headcount
4
8
12
16
20
Pseudo $R^2$ = 0.05
p = 4.57e-03 (**)
Primary school
Medium school
High school
High school diploma
1st year of college
2nd year of college (bachelor)
3rd year of college (licence)
4th year of college
Graduate (master)
Graduate (PhD)
Reported education level

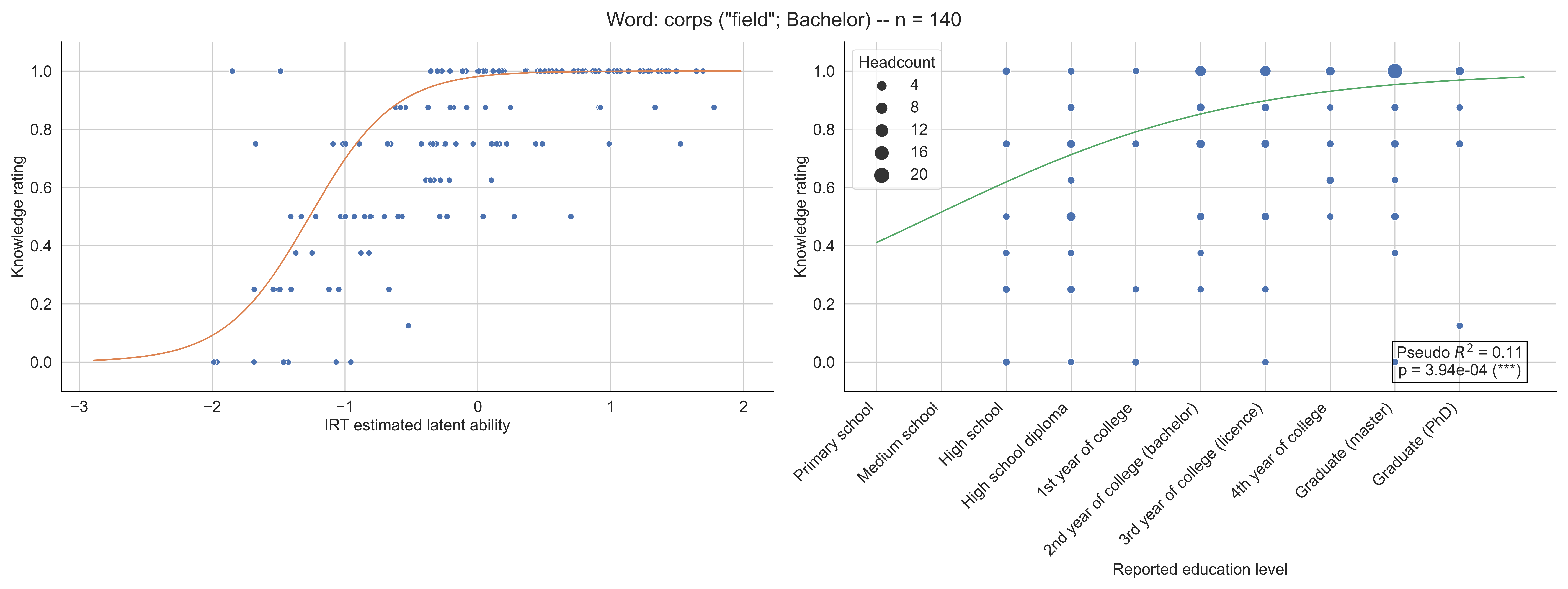
Word: corps ("field"; Bachelor) -- n = 140
Knowledge rating
IRT estimated latent ability
Headcount
4
8
12
16
20
Pseudo $R^2$ = 0.11
p = 3.94e-04 (***)
Primary school
Medium school
High school
High school diploma
1st year of college
2nd year of college (bachelor)
3rd year of college (licence)
4th year of college
Graduate (master)
Graduate (PhD)
Reported education level

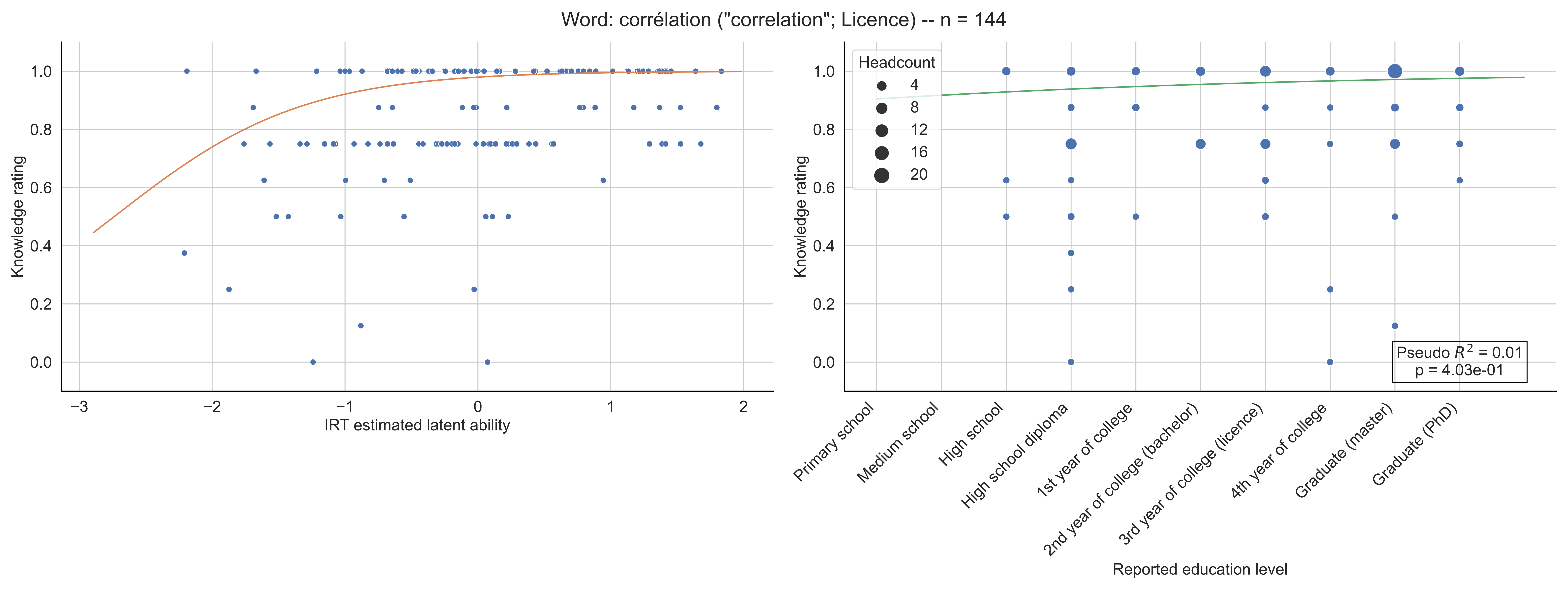

Word: corrélation ("correlation"; Licence) -- n = 144
Knowledge rating
IRT estimated latent ability
Headcount
4
8
12
16
20
Pseudo $R^2$ = 0.01
p = 4.03e-01
Primary school
Medium school
High school
High school diploma
1st year of college
2nd year of college (bachelor)
3rd year of college (licence)
4th year of college
Graduate (master)
Graduate (PhD)
Reported education level

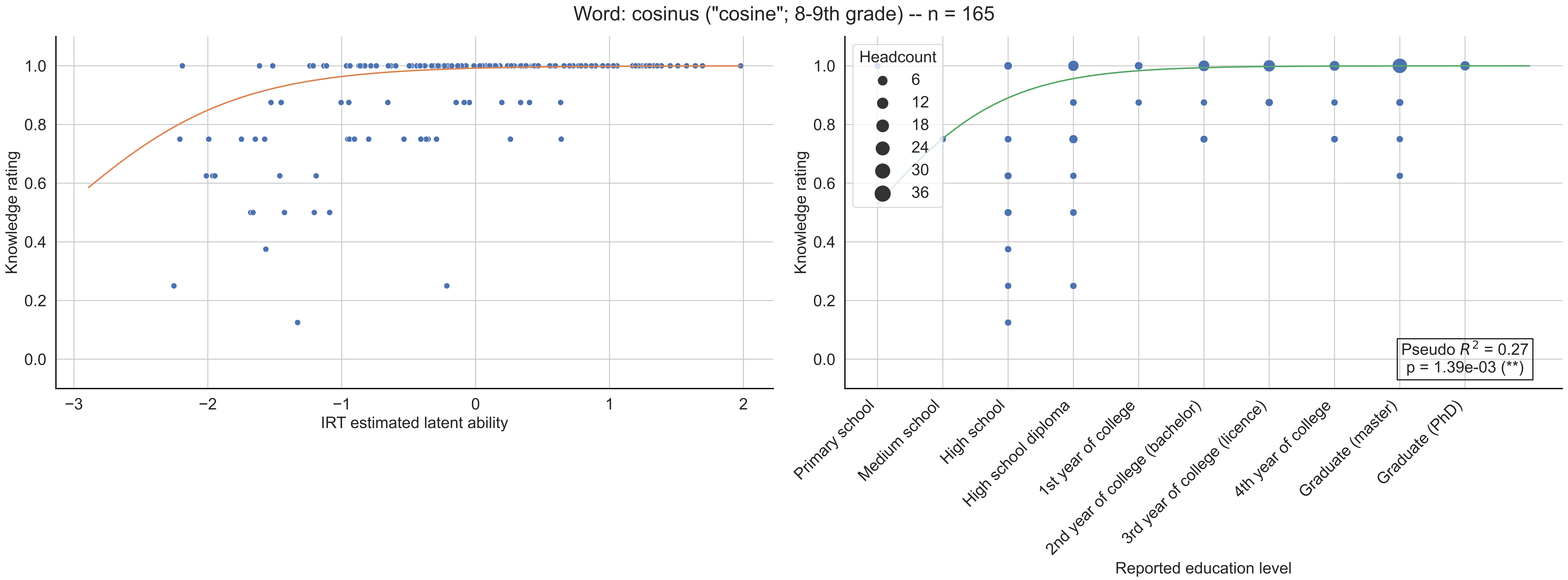
Word: cosinus ("cosine"; 8-9th grade) -- n = 165
Knowledge rating
IRT estimated latent ability
Headcount
6
12
18
24
30
36
Pseudo $R^2$ = 0.27
p = 1.39e-03 (**)
Primary school
Medium school
High school
High school diploma
1st year of college
2nd year of college (bachelor)
3rd year of college (licence)
4th year of college
Graduate (master)
Graduate (PhD)
Reported education level

Word: courbure ("curvature"; 10th grade) -- n = 151

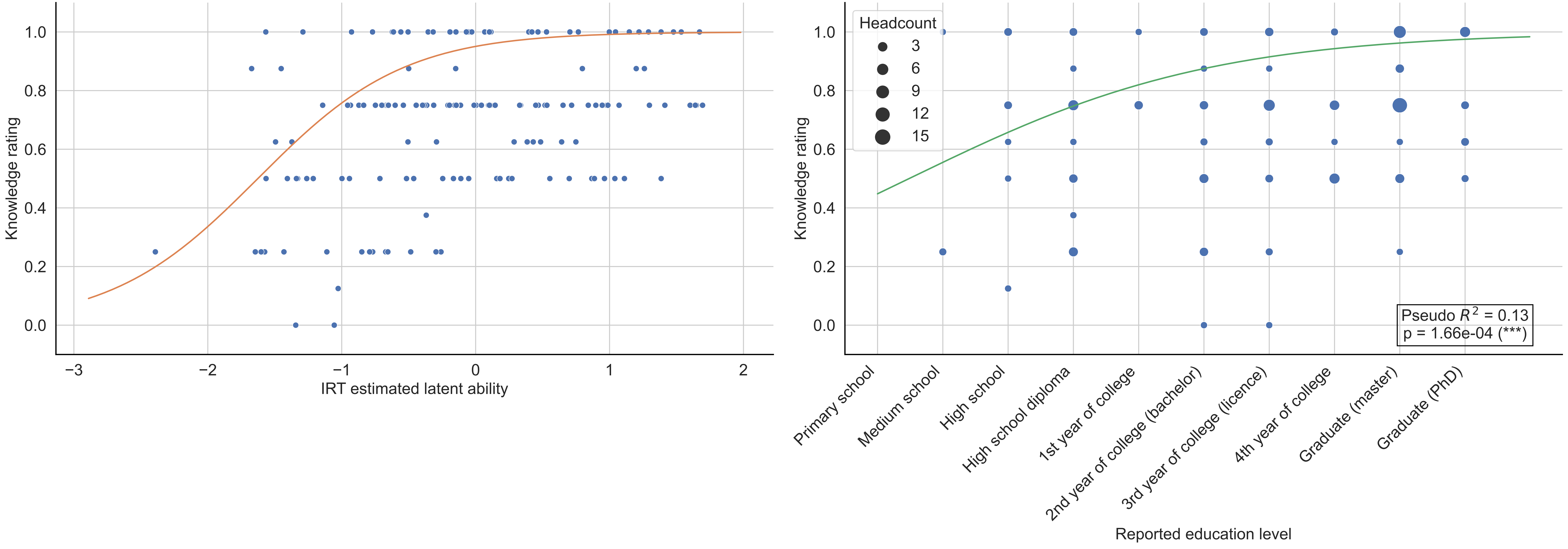

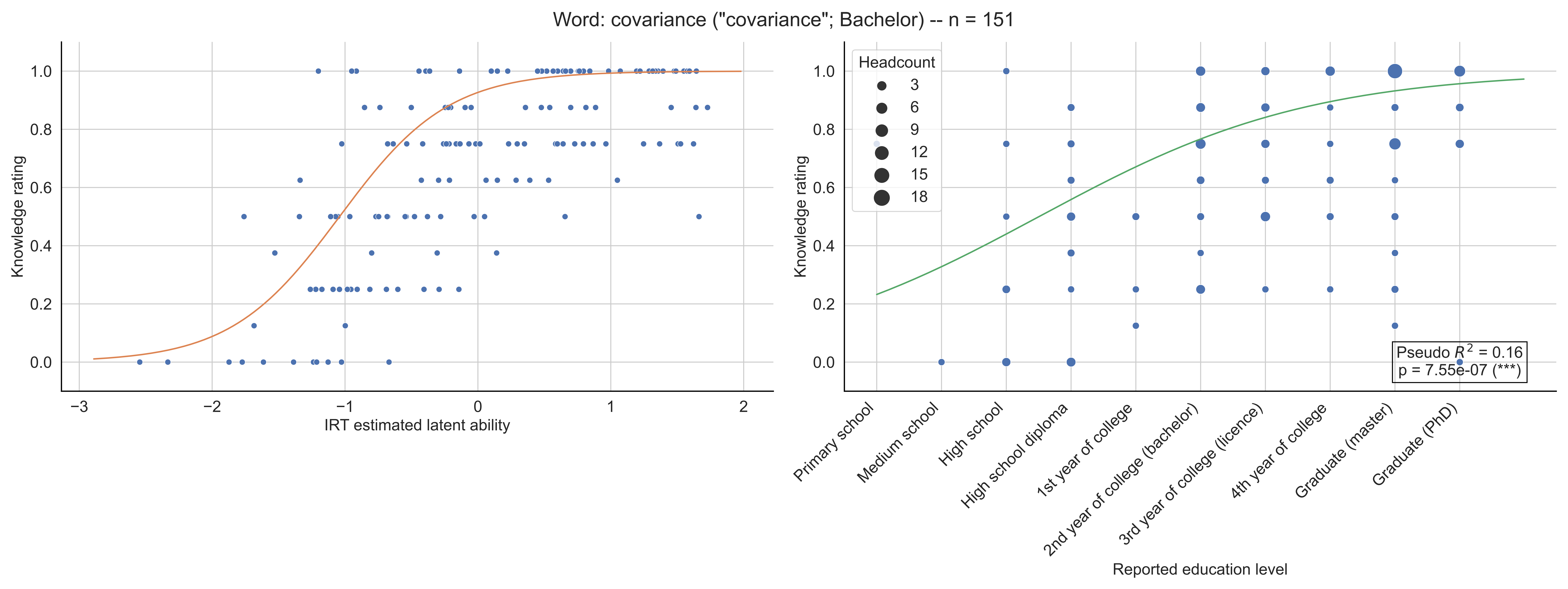

Word: covariance ("covariance"; Bachelor) -- n = 151
Knowledge rating
IRT estimated latent ability
Headcount
3
6
9
12
15
18
Pseudo $R^2$ = 0.16
p = 7.55e-07 (***)
Primary school
Medium school
High school
High school diploma
1st year of college
2nd year of college (bachelor)
3rd year of college (licence)
4th year of college
Graduate (master)
Graduate (PhD)
Reported education level

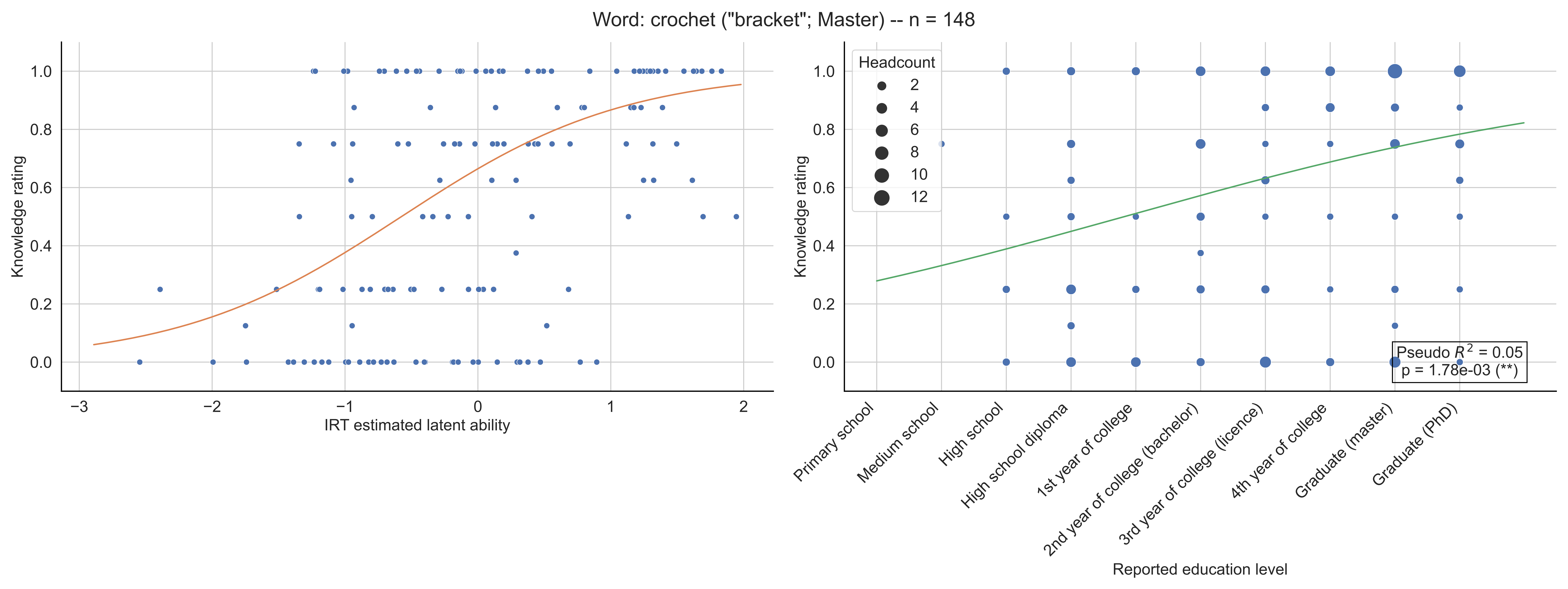

Word: crochet ("bracket"; Master) -- n = 148
Knowledge rating
IRT estimated latent ability
Headcount
2
4
6
8
10
12
Pseudo $R^2$ = 0.05
p = 1.78e-03 (**)
Primary school
Medium school
High school
High school diploma
1st year of college
2nd year of college (bachelor)
3rd year of college (licence)
4th year of college
Graduate (master)
Graduate (PhD)
Reported education level

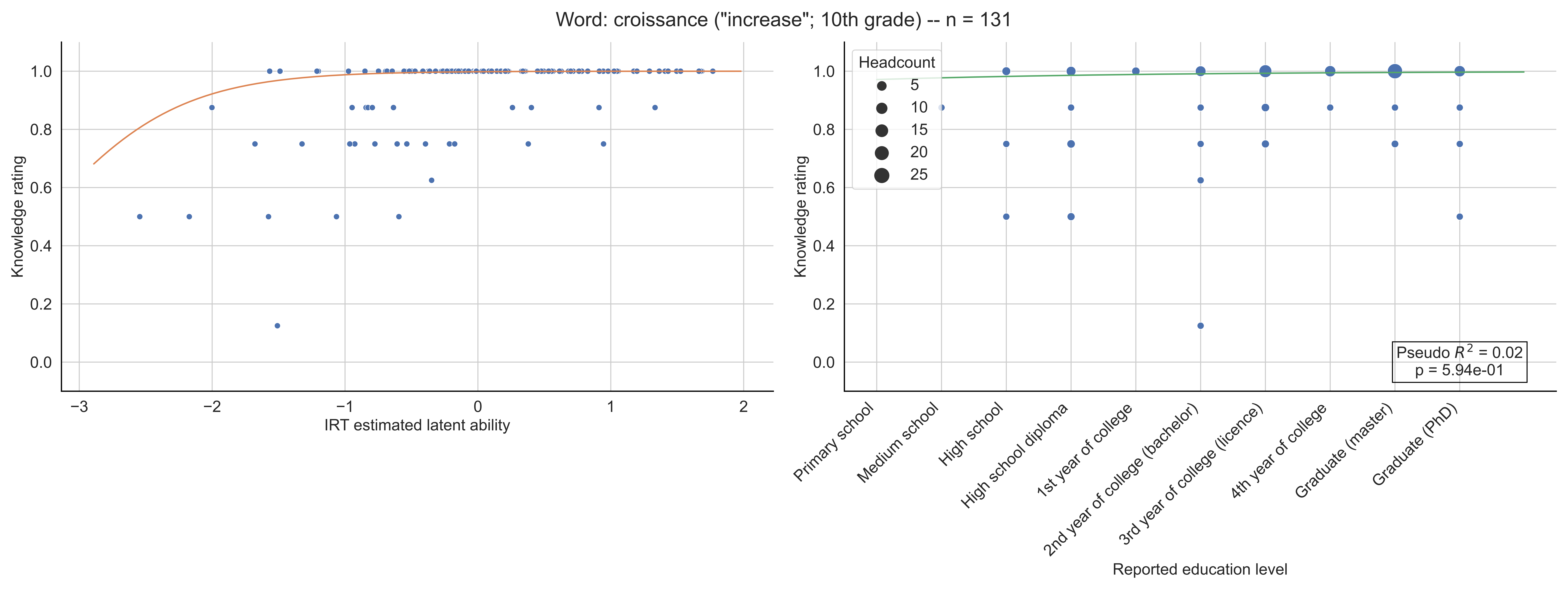

Word: croissance ("increase"; 10th grade) -- n = 131
Knowledge rating
IRT estimated latent ability
−3
−2
−1
0
1
2
0.0
0.2
0.4
0.6
0.8
1.0
Headcount
5
10
15
20
25
Pseudo $R^2$ = 0.02
p = 5.94e-01
Primary school
Medium school
High school
High school diploma
1st year of college
2nd year of college (bachelor)
3rd year of college (licence)
4th year of college
Graduate (master)
Graduate (PhD)
Reported education level

Word: cryptographie ("cryptography"; 10th grade) -- n = 130

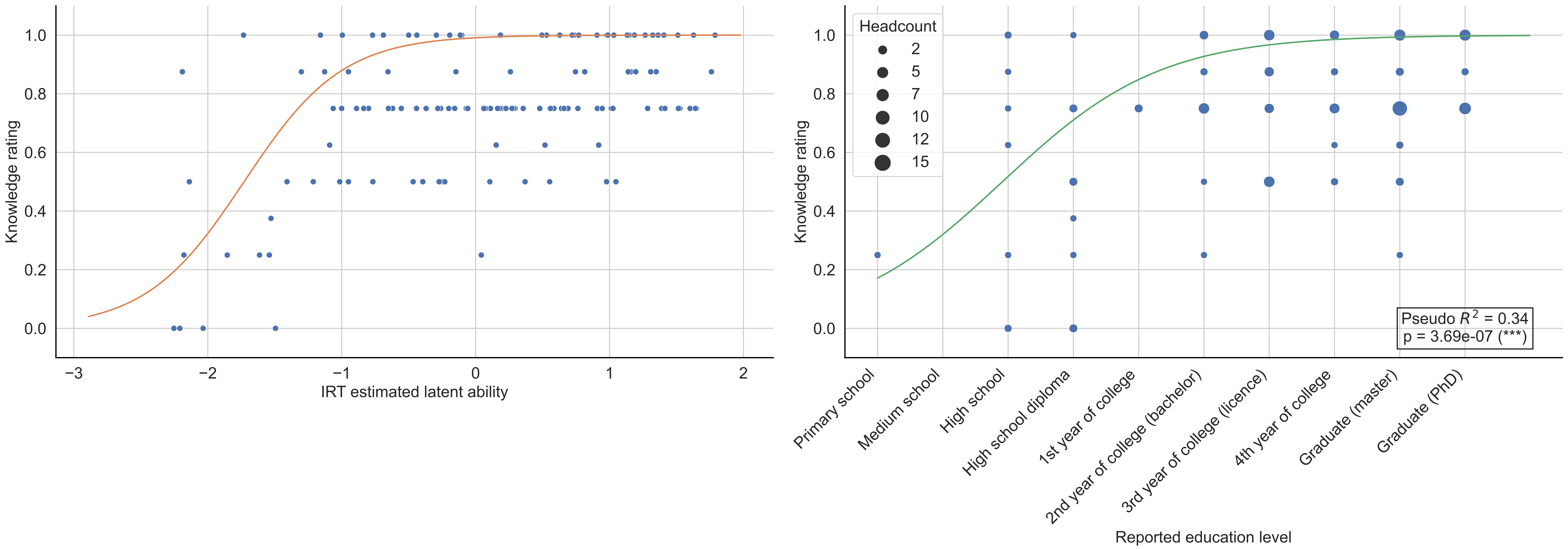

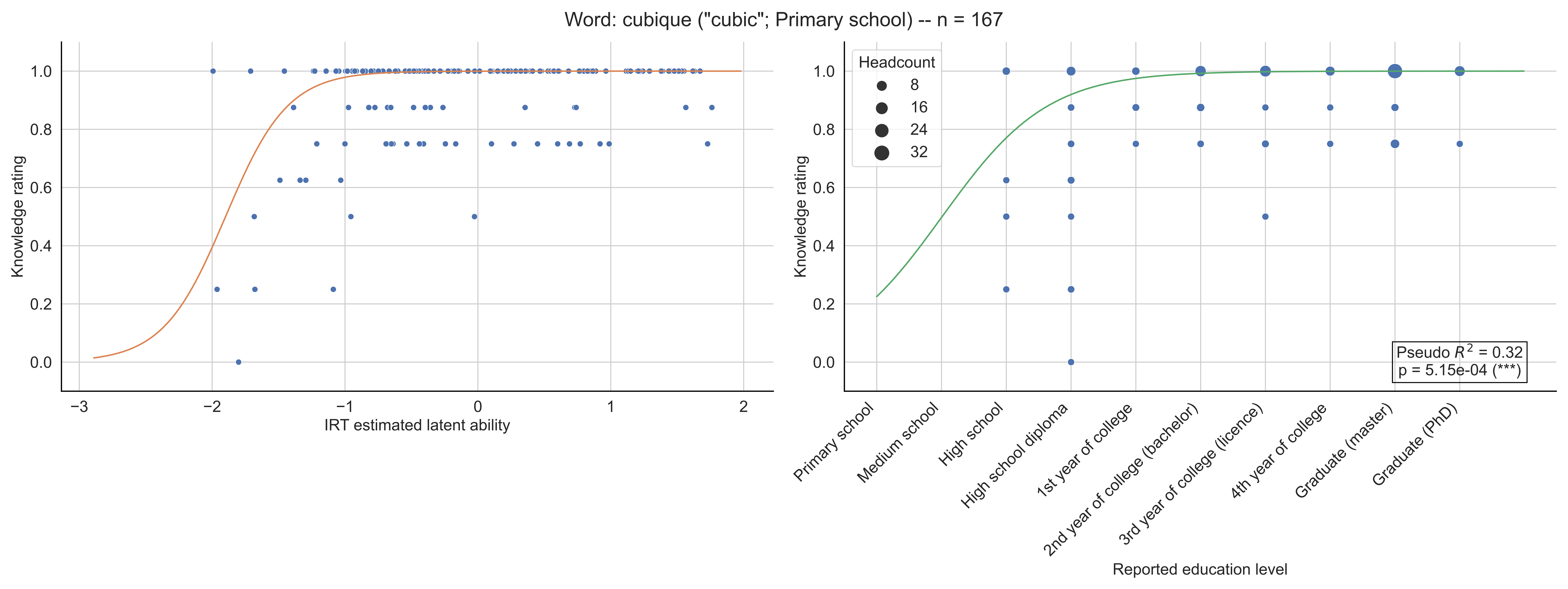
Word: cubique ("cubic"; Primary school) -- n = 167
Knowledge rating
IRT estimated latent ability
Headcount
8
16
24
32
Pseudo $R^2$ = 0.32
p = 5.15e-04 (***)
Primary school
Medium school
High school
High school diploma
1st year of college
2nd year of college (bachelor)
3rd year of college (licence)
4th year of college
Graduate (master)
Graduate (PhD)
Reported education level

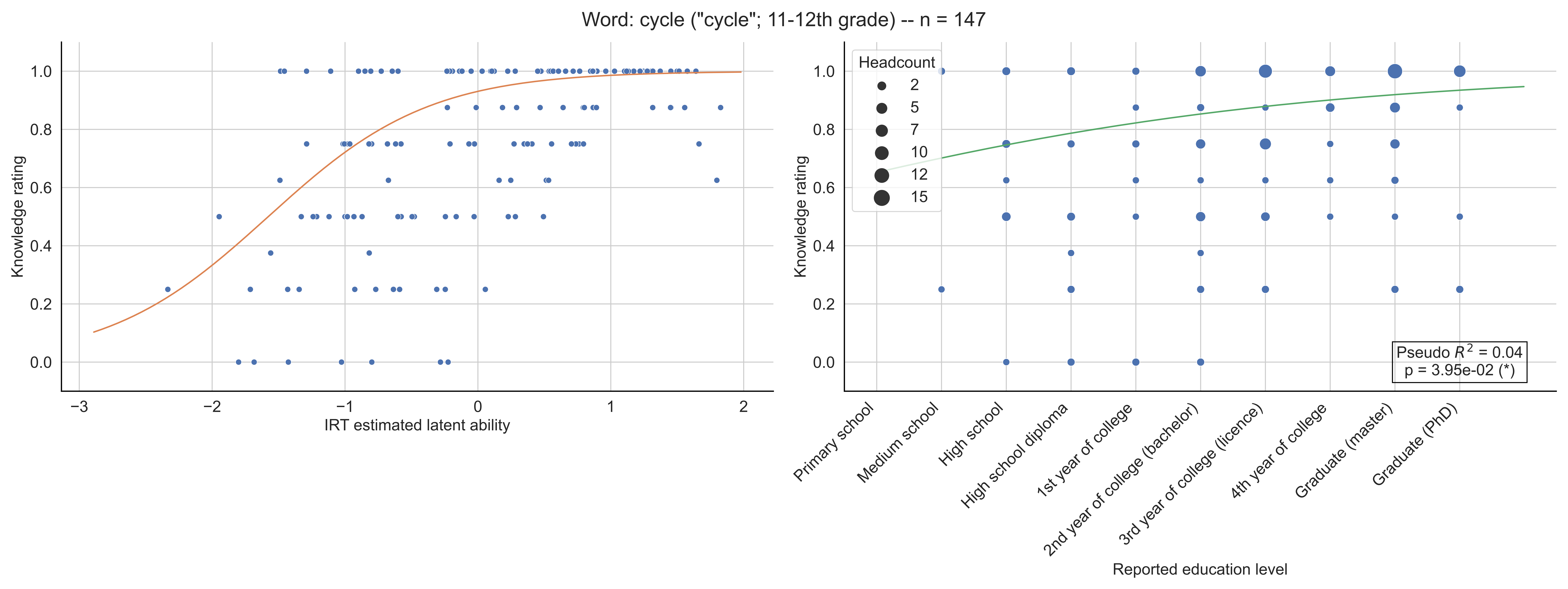

Word: cycle ("cycle"; 11-12th grade) -- n = 147
Knowledge rating
IRT estimated latent ability
Headcount
2
5
7
10
12
15
Pseudo $R^2$ = 0.04
p = 3.95e-02 (*)
Primary school
Medium school
High school
High school diploma
1st year of college
2nd year of college (bachelor)
3rd year of college (licence)
4th year of college
Graduate (master)
Graduate (PhD)
Reported education level

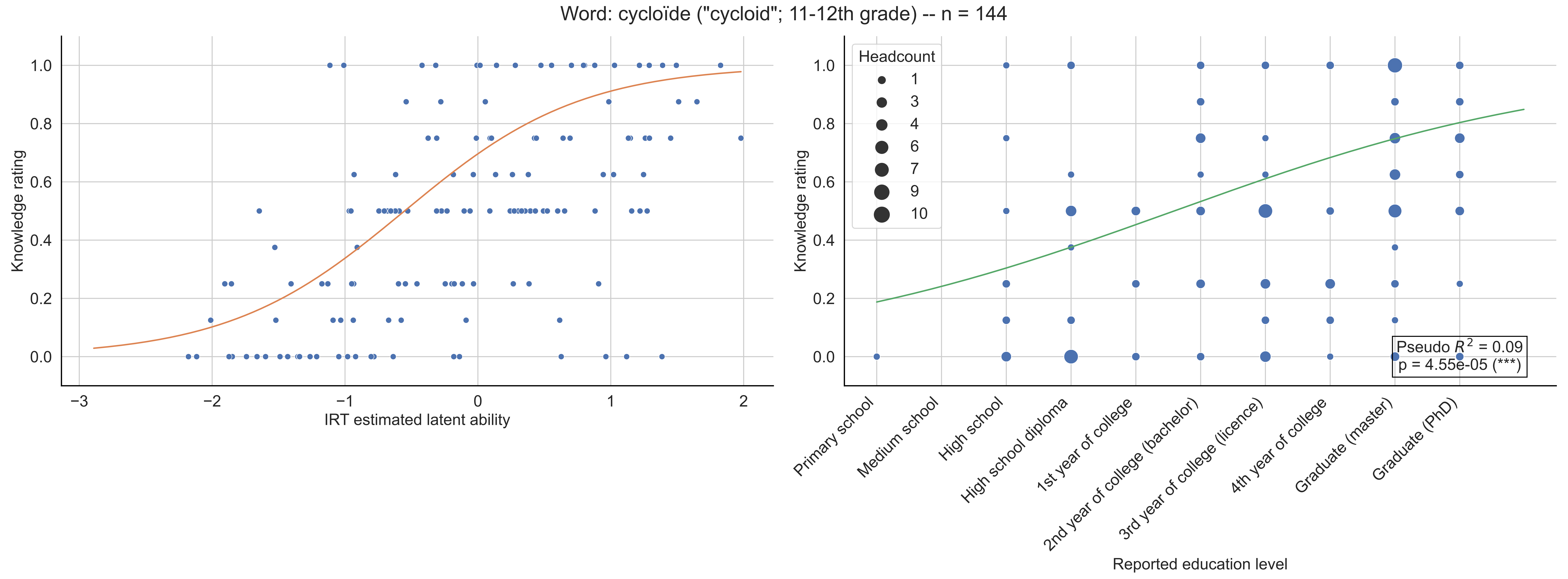

Word: cycloïde ("cycloid"; 11-12th grade) -- n = 144
Knowledge rating
IRT estimated latent ability
Headcount
1
3
4
6
7
9
10
Pseudo $R^2$ = 0.09
p = 4.55e-05 (***)
Primary school
Medium school
High school
High school diploma
1st year of college
2nd year of college (bachelor)
3rd year of college (licence)
4th year of college
Graduate (master)
Graduate (PhD)
Reported education level

Word: cône ("cone"; Primary school) -- n = 161

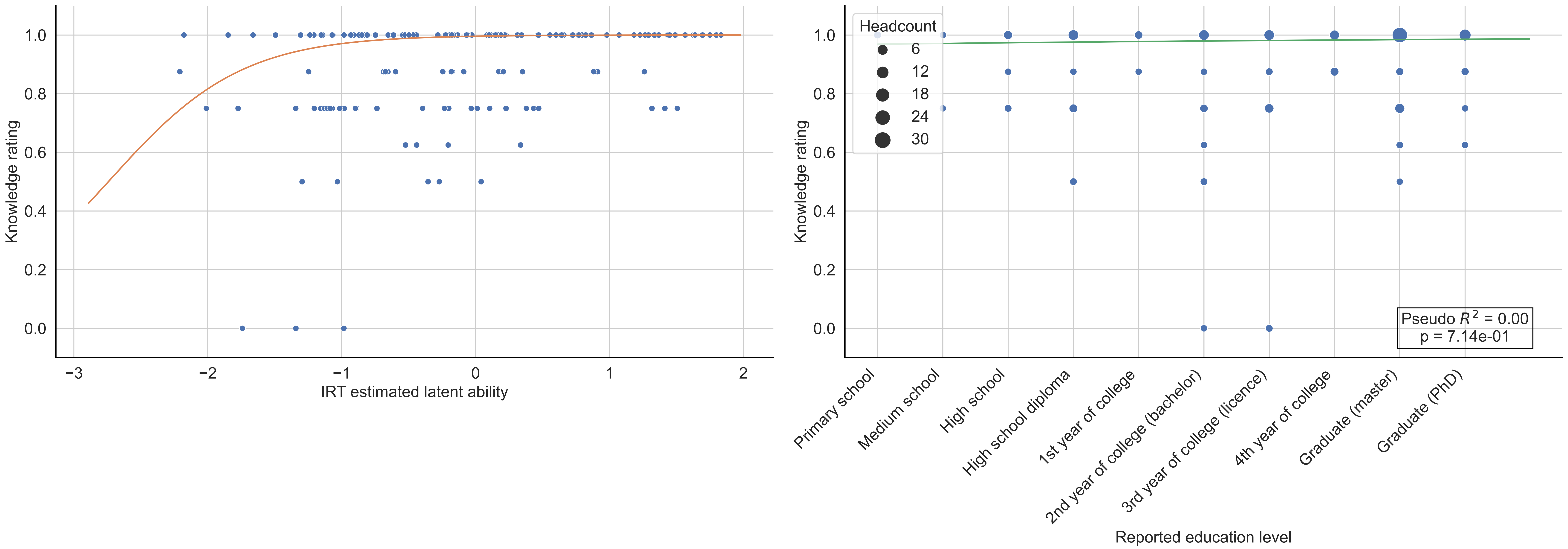

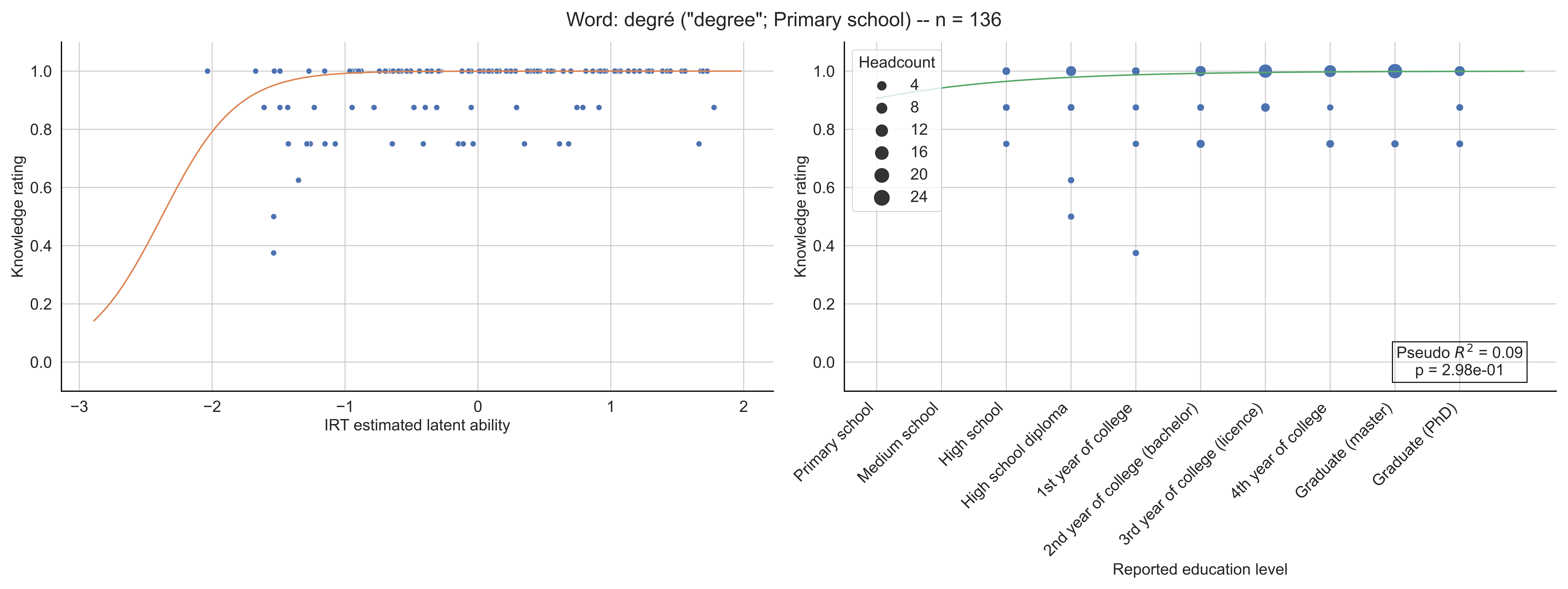

Word: degré ("degree"; Primary school) -- n = 136
Knowledge rating
IRT estimated latent ability
Headcount
4
8
12
16
20
24
Pseudo $R^2$ = 0.09
p = 2.98e-01
Primary school
Medium school
High school
High school diploma
1st year of college
2nd year of college (bachelor)
3rd year of college (licence)
4th year of college
Graduate (master)
Graduate (PhD)
Reported education level

Word: densité ("density"; 11-12th grade) -- n = 132

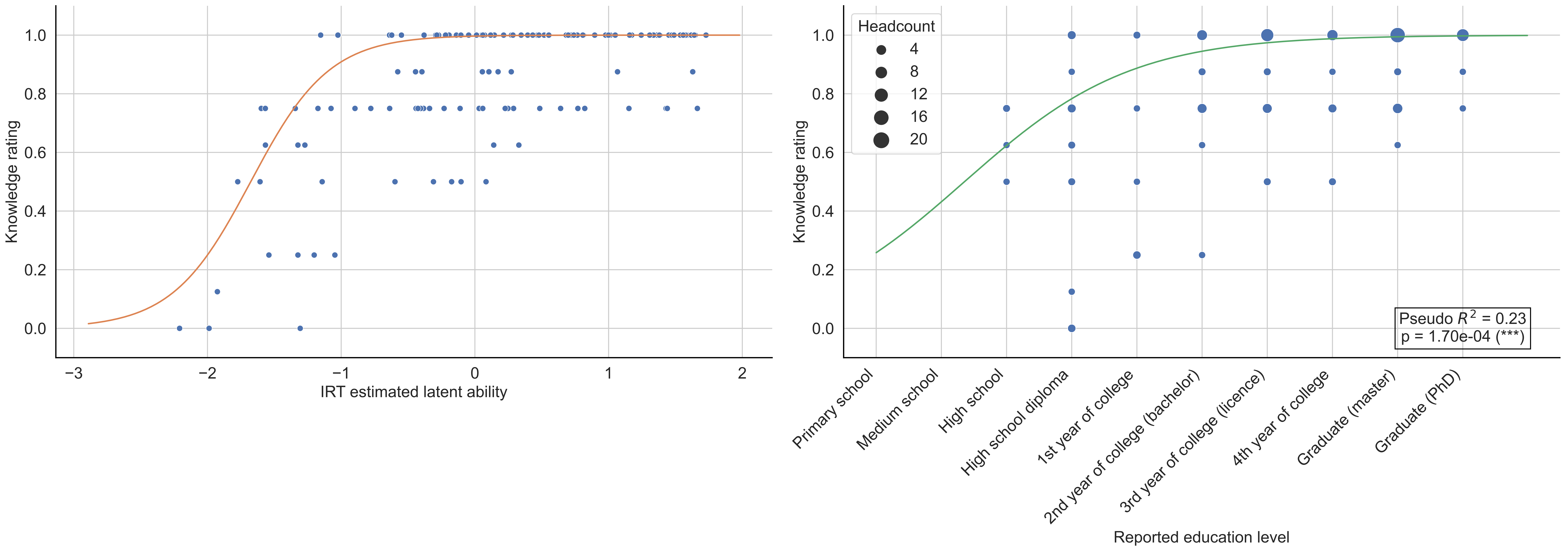

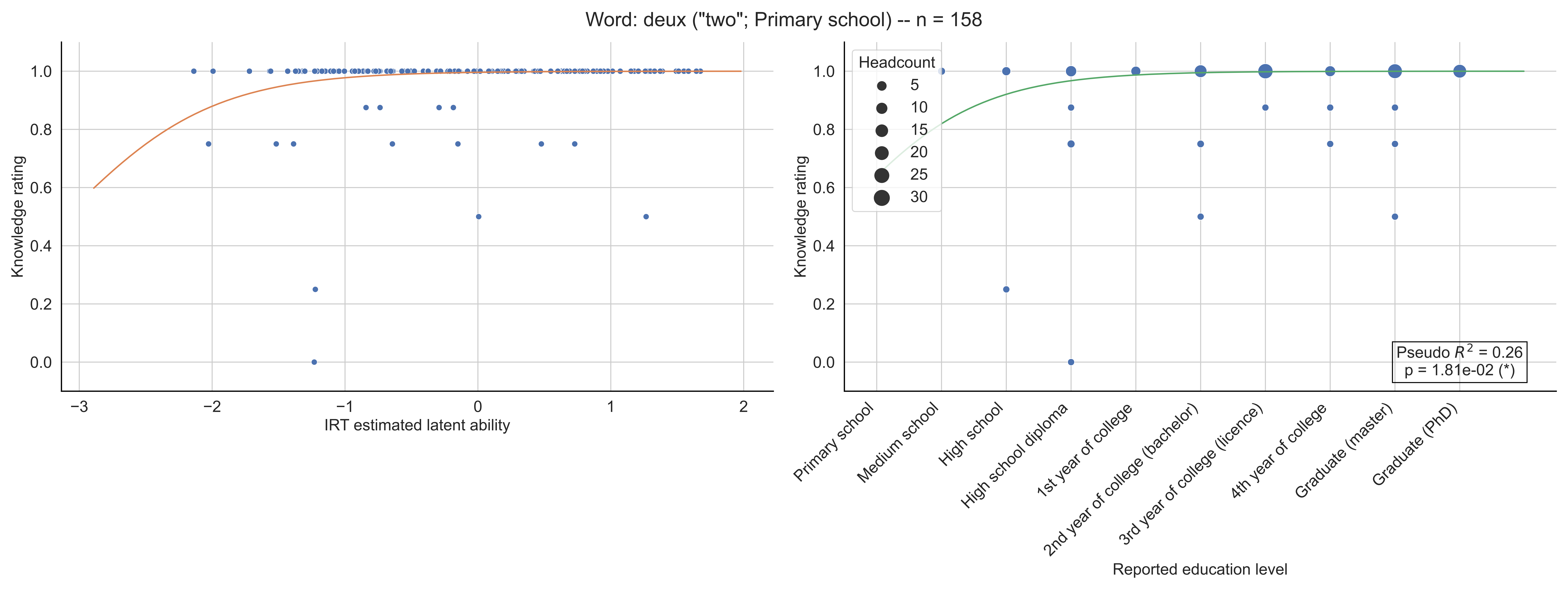
Word: deux ("two"; Primary school) -- n = 158
Knowledge rating
IRT estimated latent ability
Headcount
5
10
15
20
25
30
Pseudo $R^2$ = 0.26
p = 1.81e-02 (*)
Primary school
Medium school
High school
High school diploma
1st year of college
2nd year of college (bachelor)
3rd year of college (licence)
4th year of college
Graduate (master)
Graduate (PhD)
Reported education level

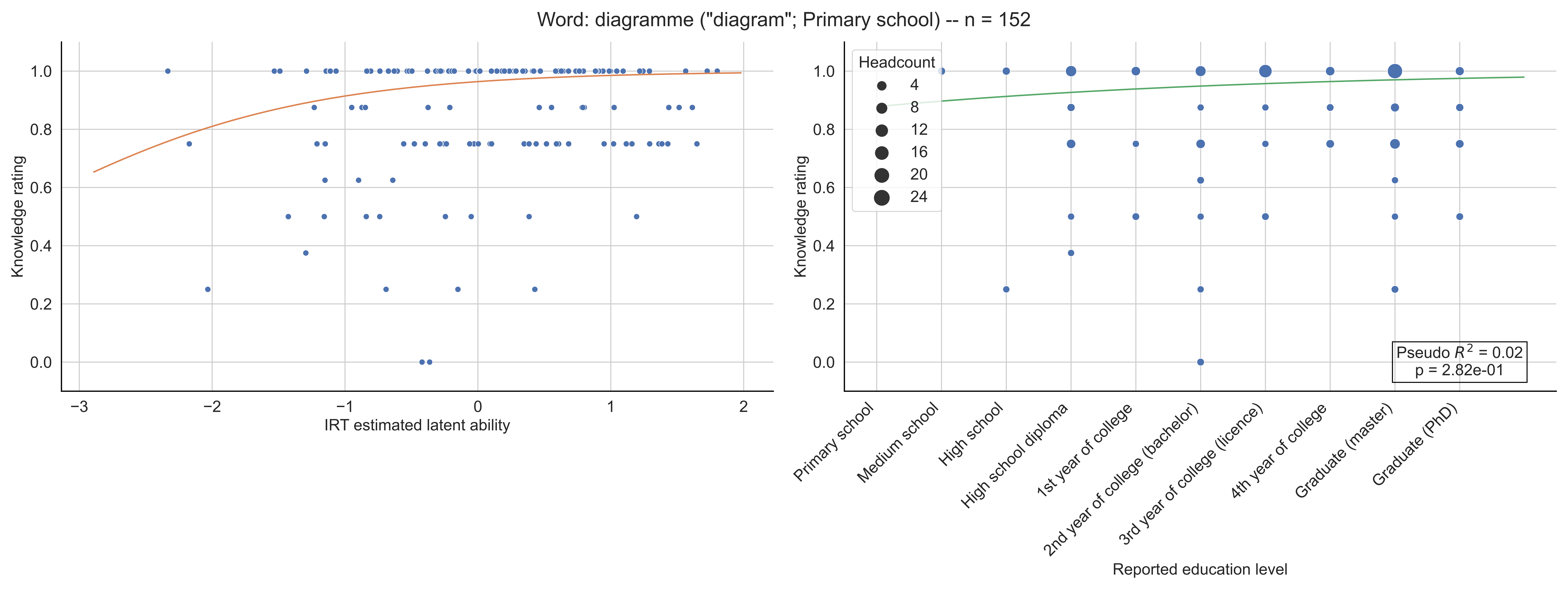
Word: diagramme ("diagram"; Primary school) -- n = 152
Knowledge rating
IRT estimated latent ability
Headcount
4
8
12
16
20
24
Pseudo $R^2$ = 0.02
p = 2.82e-01
Primary school
Medium school
High school
High school diploma
1st year of college
2nd year of college (bachelor)
3rd year of college (licence)
4th year of college
Graduate (master)
Graduate (PhD)
Reported education level

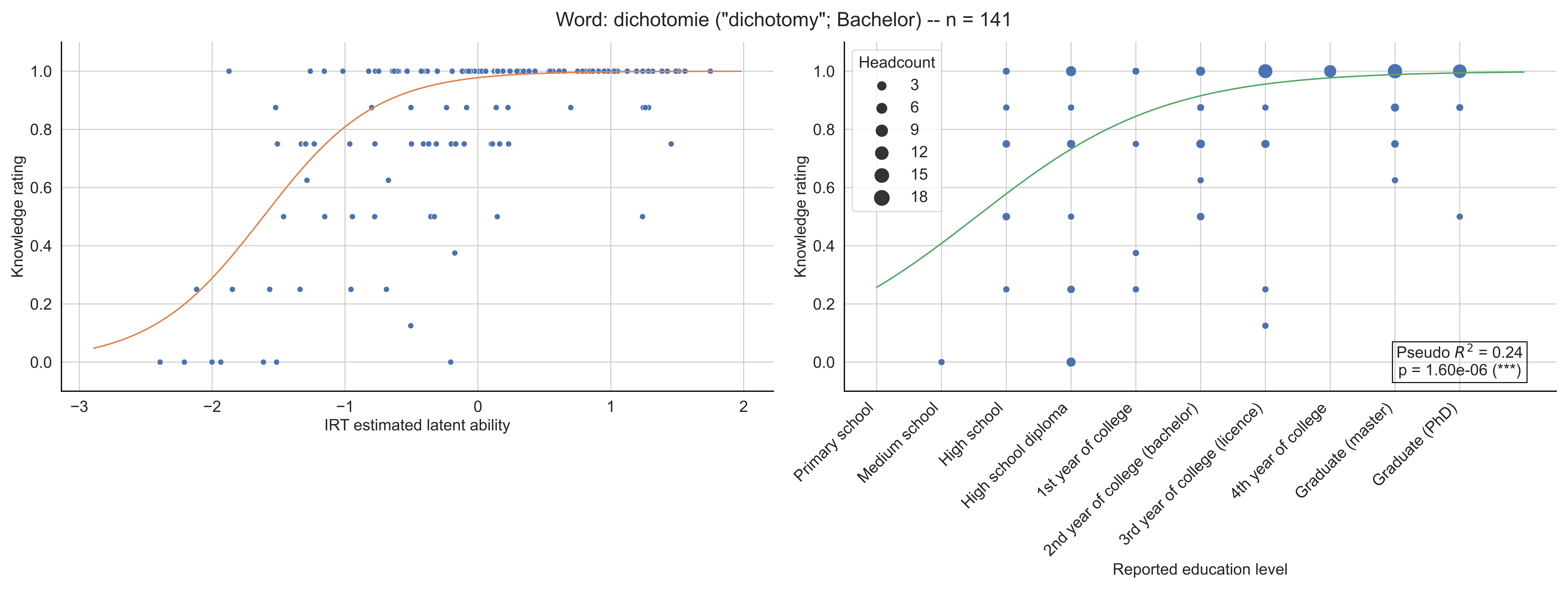

Word: dichotomie ("dichotomy"; Bachelor) -- n = 141
Knowledge rating
IRT estimated latent ability
Headcount
3
6
9
12
15
18
Pseudo $R^2$ = 0.24
p = 1.60e-06 (***)
Primary school
Medium school
High school
High school diploma
1st year of college
2nd year of college (bachelor)
3rd year of college (licence)
4th year of college
Graduate (master)
Graduate (PhD)
Reported education level

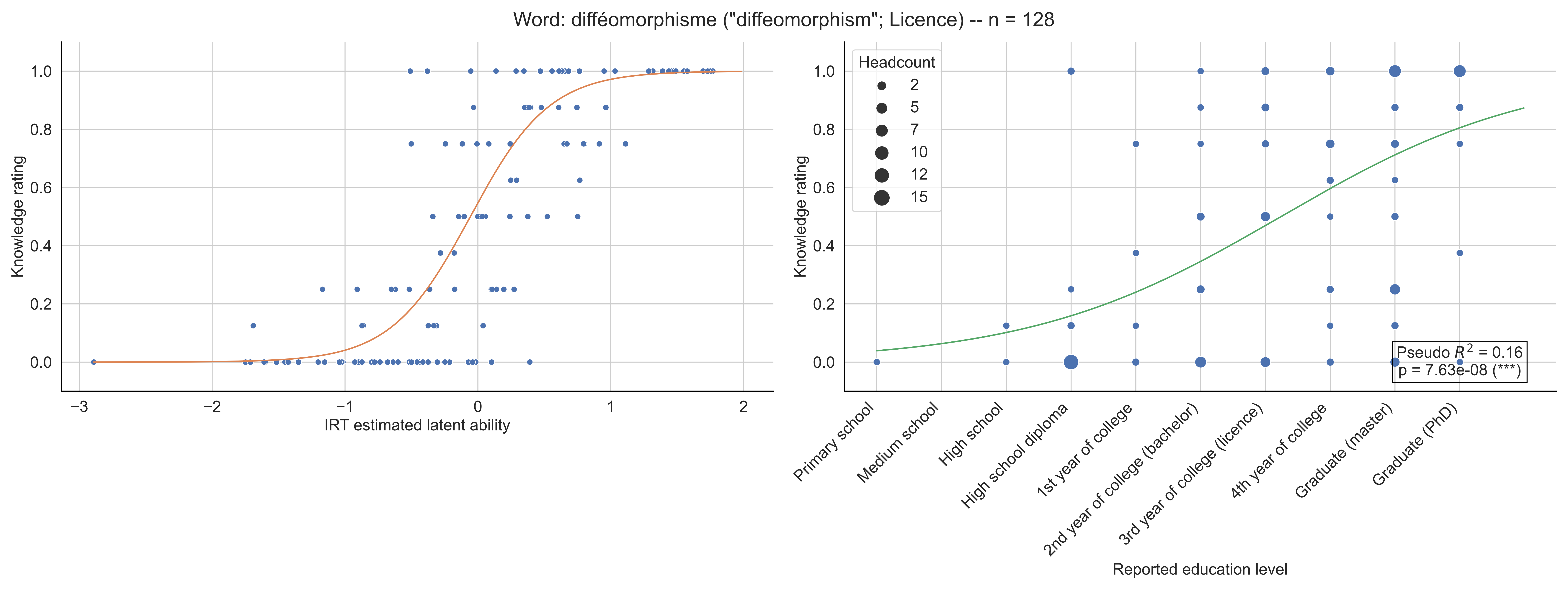

Word: difféomorphisme ("diffeomorphism"; Licence) -- n = 128
Knowledge rating
IRT estimated latent ability
Headcount
2
5
7
10
12
15
Pseudo $R^2$ = 0.16
p = 7.63e-08 (***)
Primary school
Medium school
High school
High school diploma
1st year of college
2nd year of college (bachelor)
3rd year of college (licence)
4th year of college
Graduate (master)
Graduate (PhD)
Reported education level

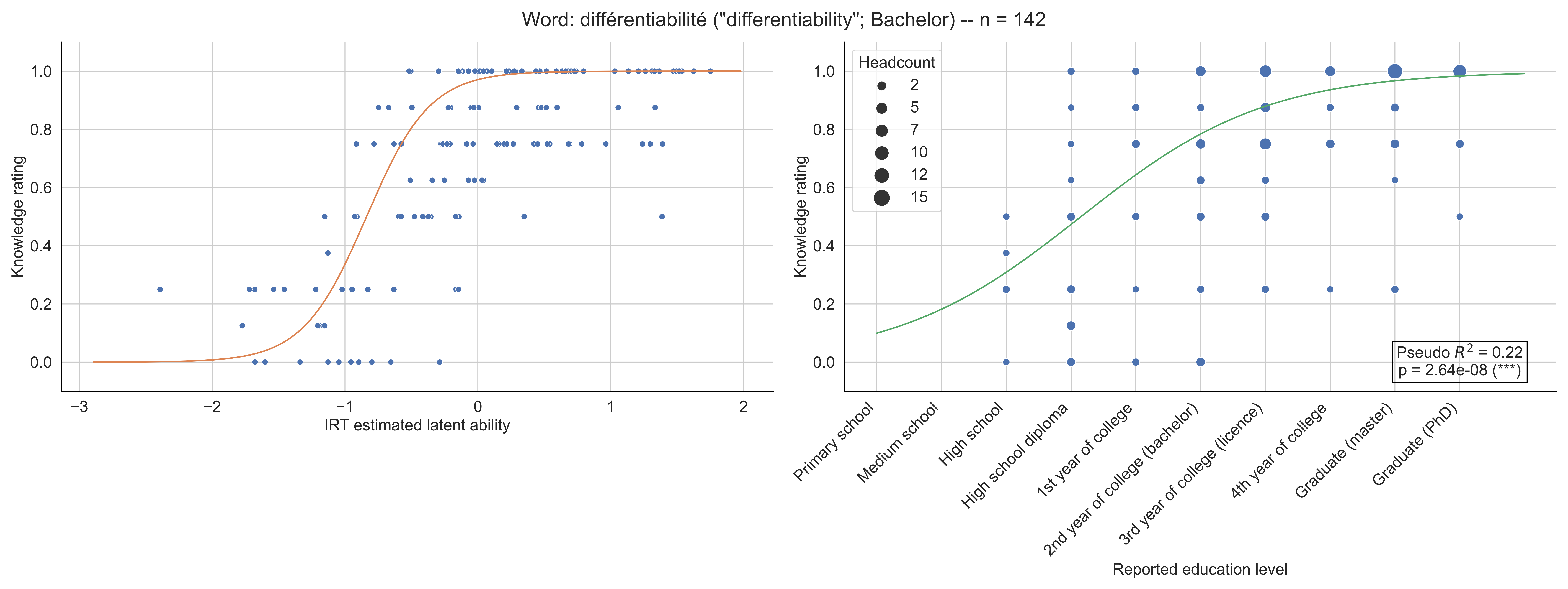
Word: différentiabilité ("differentiability"; Bachelor) -- n = 142
Knowledge rating
IRT estimated latent ability
Headcount
2
5
7
10
12
15
Pseudo $R^2$ = 0.22
p = 2.64e-08 (***)
Primary school
Medium school
High school
High school diploma
1st year of college
2nd year of college (bachelor)
3rd year of college (licence)
4th year of college
Graduate (master)
Graduate (PhD)
Reported education level

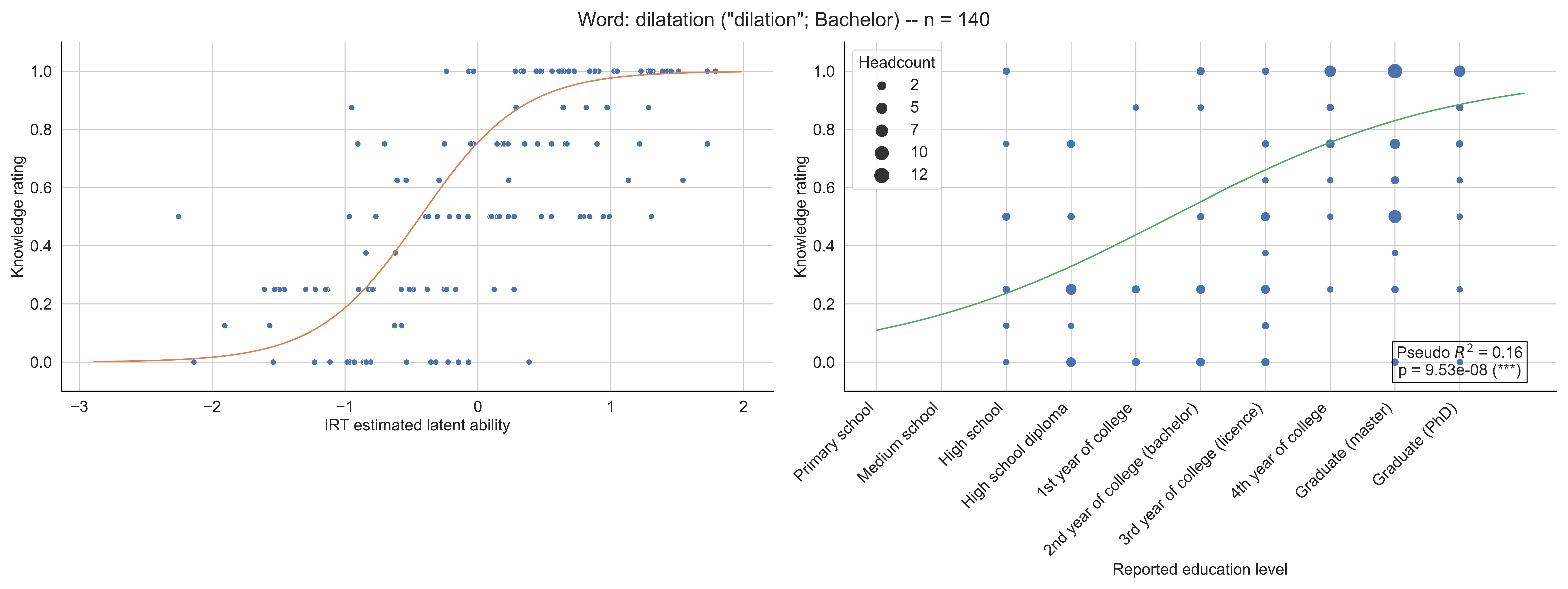

Word: dilatation ("dilation"; Bachelor) -- n = 140
Knowledge rating
IRT estimated latent ability
Headcount
2
5
7
10
12
Pseudo $R^2$ = 0.16
p = 9.53e-08 (***)
Primary school
Medium school
High school
High school diploma
1st year of college
2nd year of college (bachelor)
3rd year of college (licence)
4th year of college
Graduate (master)
Graduate (PhD)
Reported education level

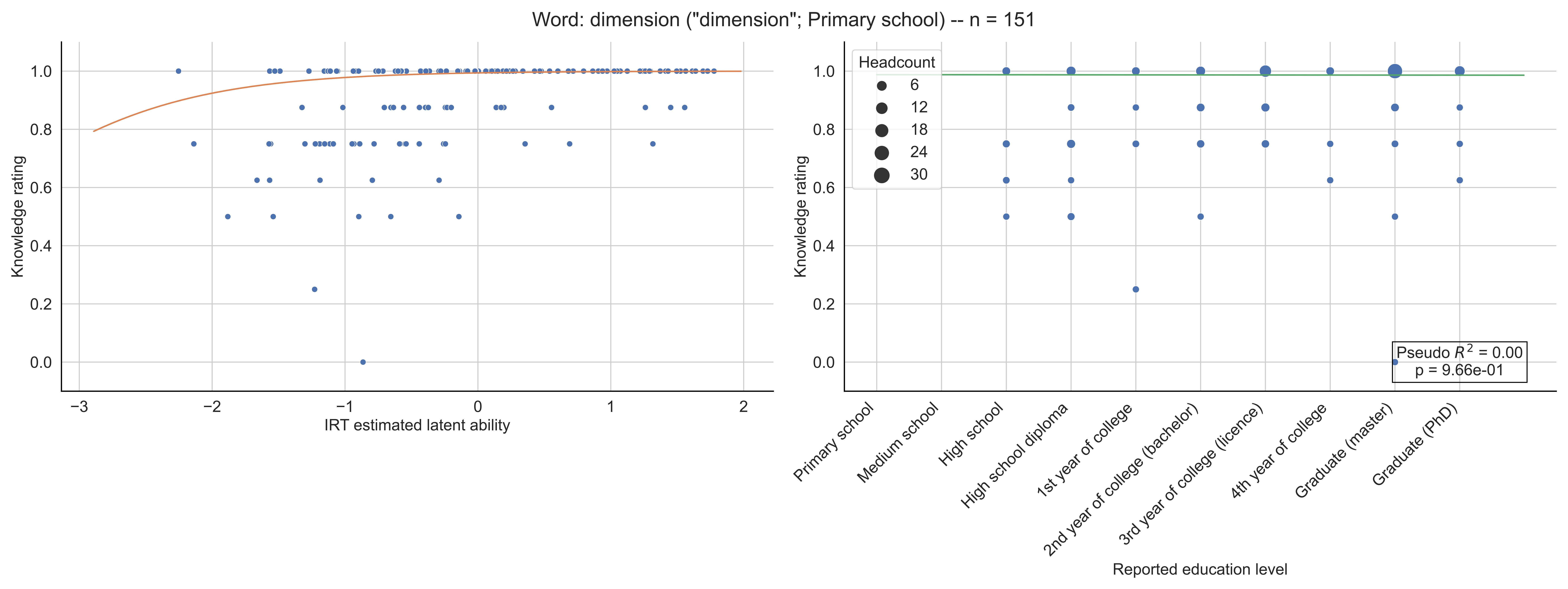

Word: dimension ("dimension"; Primary school) -- n = 151
Knowledge rating
IRT estimated latent ability
Headcount
6
12
18
24
30
Pseudo $R^2$ = 0.00
p = 9.66e-01
Primary school
Medium school
High school
High school diploma
1st year of college
2nd year of college (bachelor)
3rd year of college (licence)
4th year of college
Graduate (master)
Graduate (PhD)
Reported education level

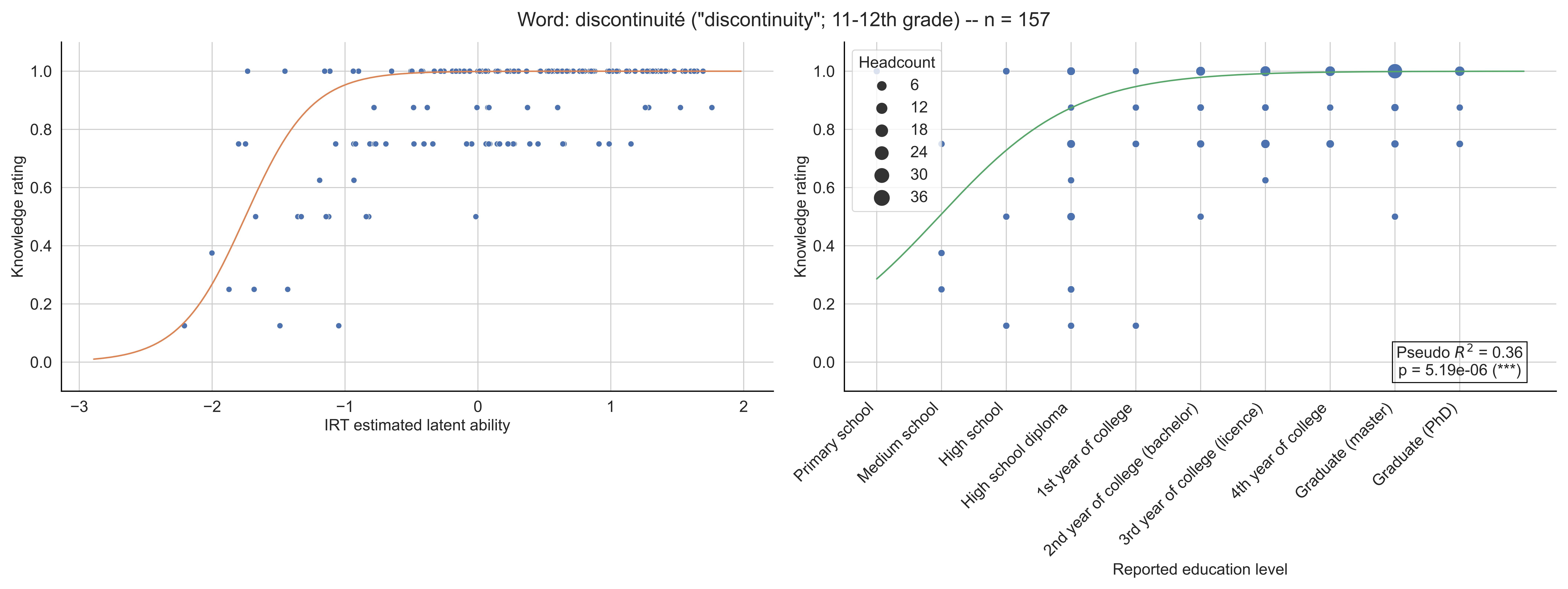
Word: discontinuité ("discontinuity"; 11-12th grade) -- n = 157
Knowledge rating
IRT estimated latent ability
−3
−2
−1
0
1
2
0.0
0.2
0.4
0.6
0.8
1.0
Headcount
6
12
18
24
30
36
Pseudo $R^2$ = 0.36
p = 5.19e-06 (***)
Primary school
Medium school
High school
High school diploma
1st year of college
2nd year of college (bachelor)
3rd year of college (licence)
4th year of college
Graduate (master)
Graduate (PhD)
Reported education level

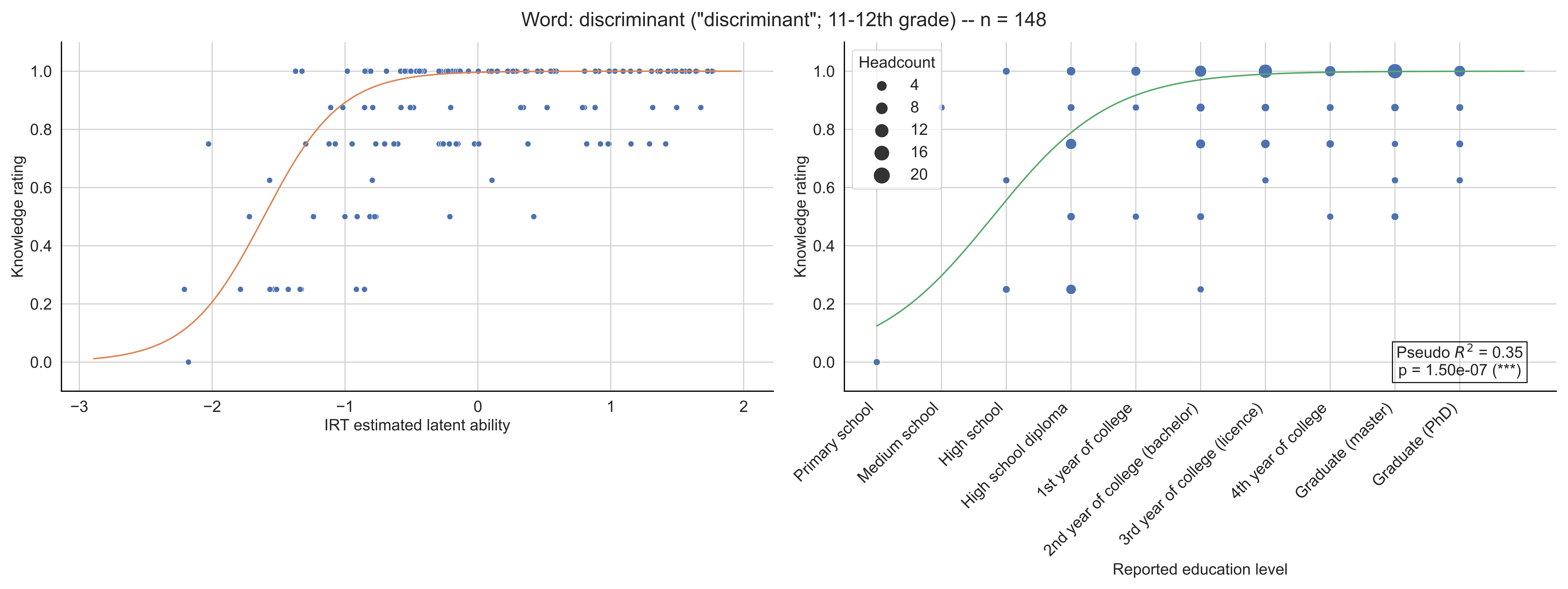

Word: discriminant ("discriminant"; 11-12th grade) -- n = 148
Knowledge rating
IRT estimated latent ability
Headcount
4
8
12
16
20
Pseudo $R^2$ = 0.35
p = 1.50e-07 (***)
Primary school
Medium school
High school
High school diploma
1st year of college
2nd year of college (bachelor)
3rd year of college (licence)
4th year of college
Graduate (master)
Graduate (PhD)
Reported education level

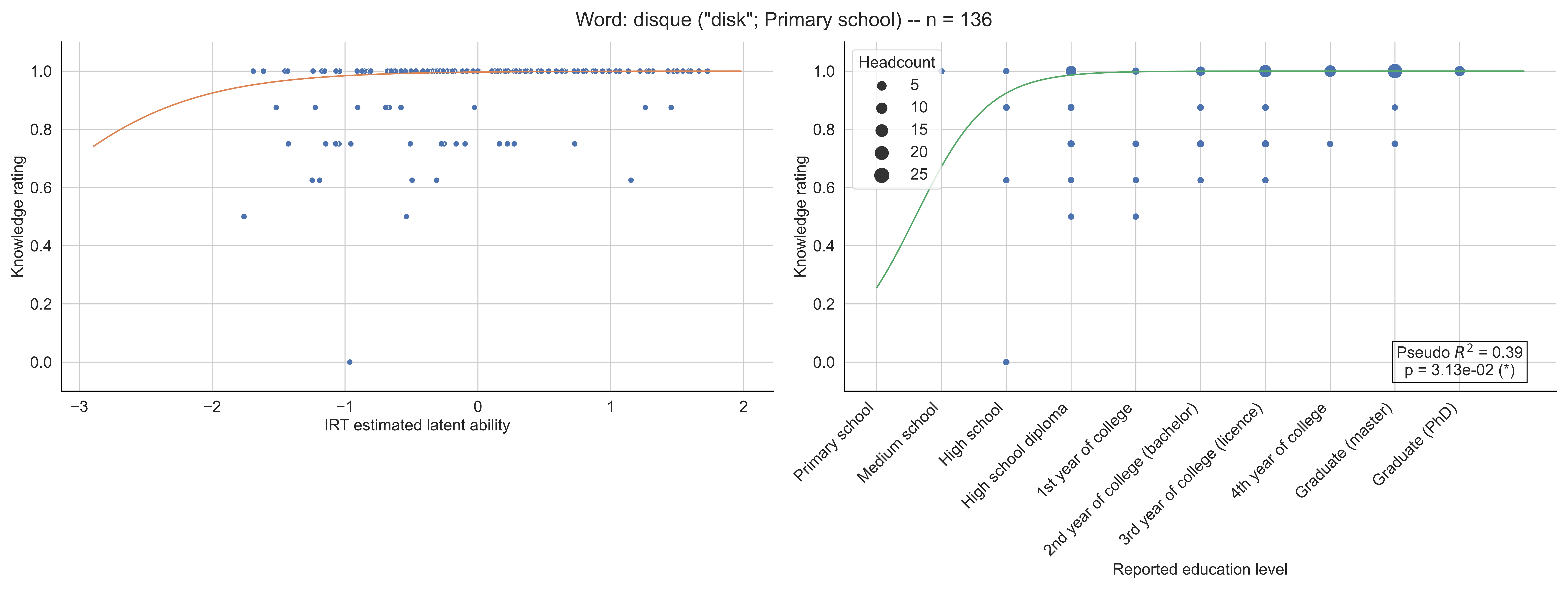

Word: disque ("disk"; Primary school) -- n = 136
Knowledge rating
IRT estimated latent ability
Headcount
5
10
15
20
25
Pseudo $R^2$ = 0.39
p = 3.13e-02 (*)
Primary school
Medium school
High school
High school diploma
1st year of college
2nd year of college (bachelor)
3rd year of college (licence)
4th year of college
Graduate (master)
Graduate (PhD)
Reported education level

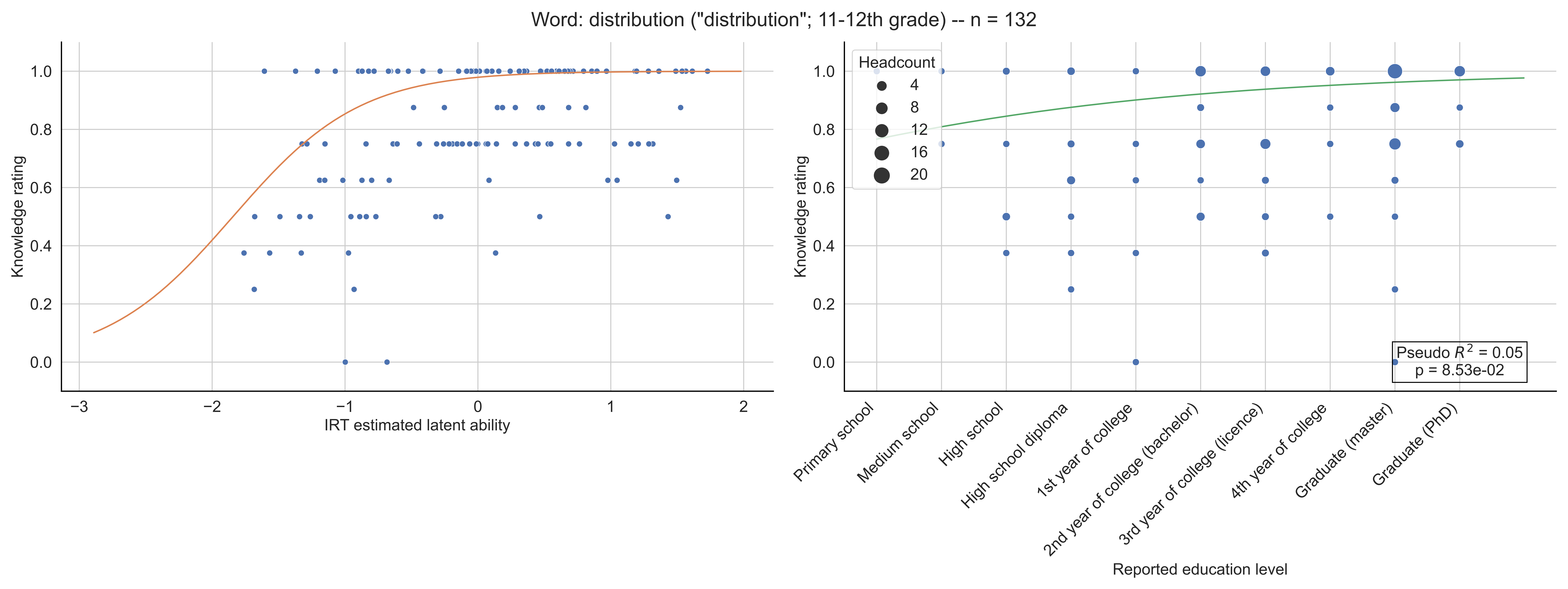
Word: distribution ("distribution"; 11-12th grade) -- n = 132
Knowledge rating
IRT estimated latent ability
Headcount
4
8
12
16
20
Pseudo $R^2$ = 0.05
p = 8.53e-02
Primary school
Medium school
High school
High school diploma
1st year of college
2nd year of college (bachelor)
3rd year of college (licence)
4th year of college
Graduate (master)
Graduate (PhD)
Reported education level

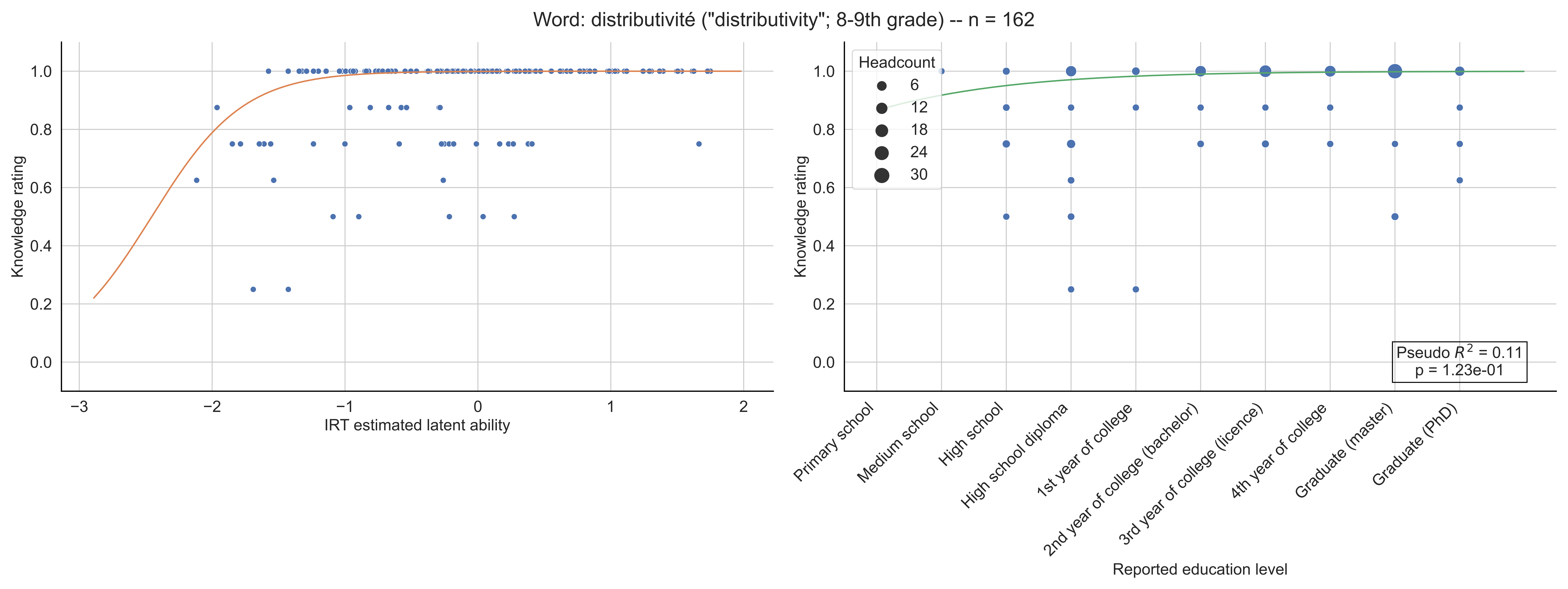
Word: distributivité ("distributivity"; 8-9th grade) -- n = 162
Knowledge rating
IRT estimated latent ability
Headcount
6
12
18
24
30
Pseudo $R^2$ = 0.11
p = 1.23e-01
Primary school
Medium school
High school
High school diploma
1st year of college
2nd year of college (bachelor)
3rd year of college (licence)
4th year of college
Graduate (master)
Graduate (PhD)
Reported education level

Word: dièdre ("dihedron"; Licence) -- n = 148

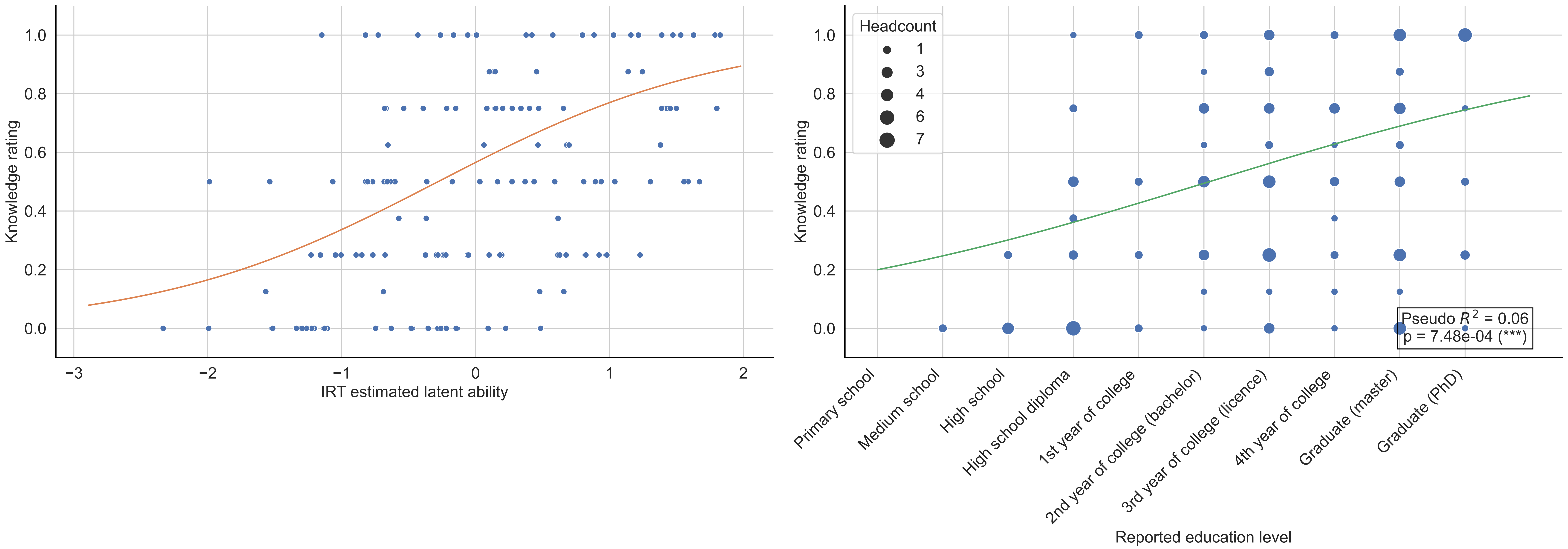

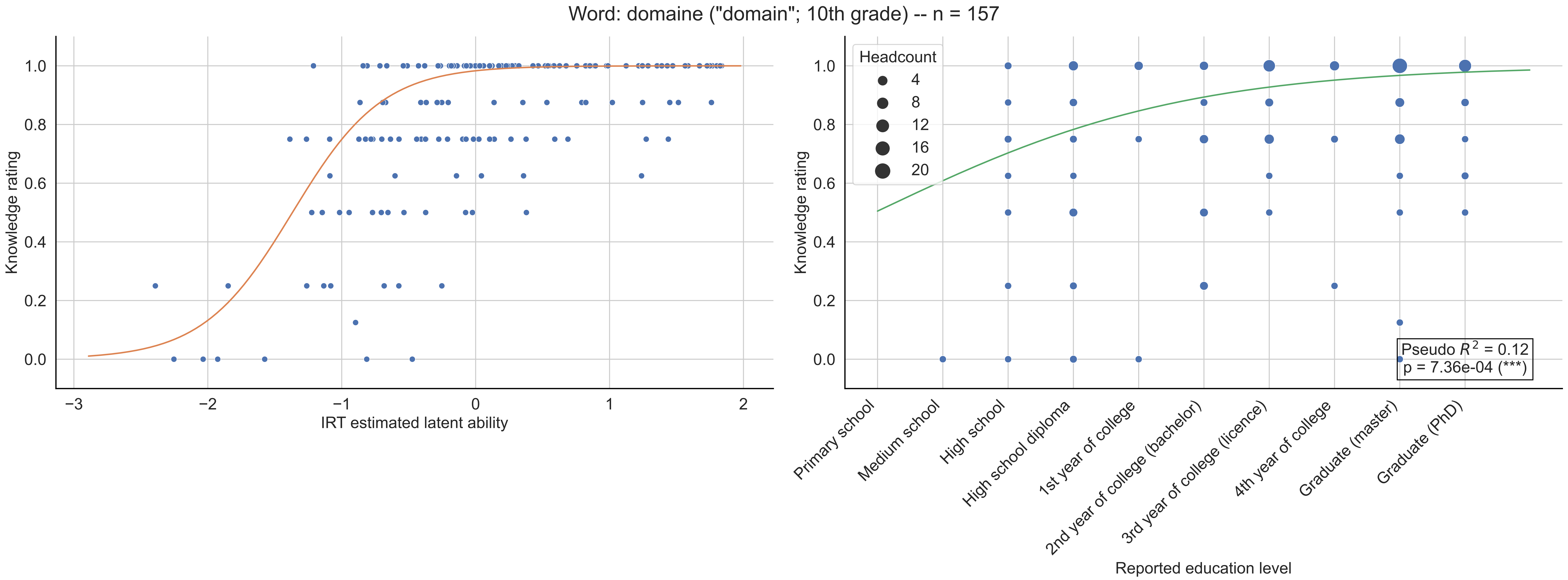

Word: domaine ("domain"; 10th grade) -- n = 157
Knowledge rating
IRT estimated latent ability
Headcount
4
8
12
16
20
Primary school
Medium school
High school
High school diploma
1st year of college
2nd year of college (bachelor)
3rd year of college (licence)
4th year of college
Graduate (master)
Graduate (PhD)
Reported education level
Pseudo $R^2$ = 0.12
p = 7.36e-04 (***)

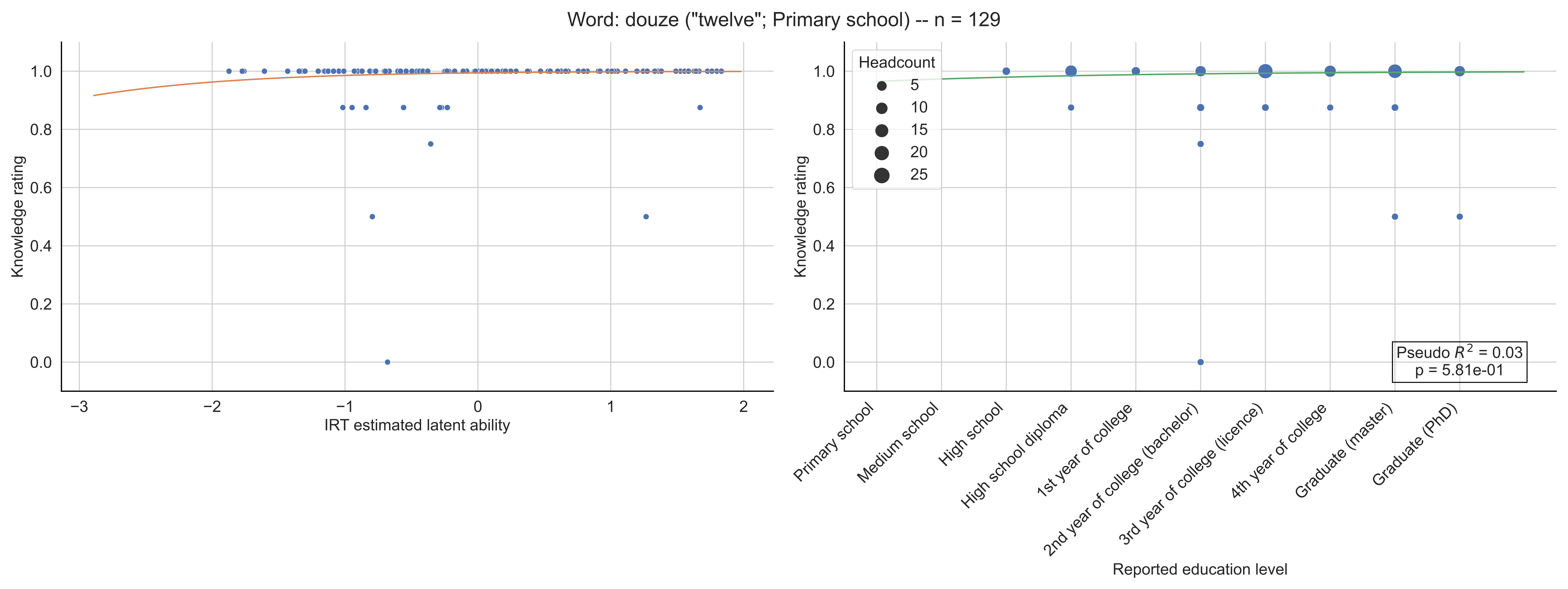

Word: douze ("twelve"; Primary school) -- n = 129
Knowledge rating
IRT estimated latent ability
Headcount
5
10
15
20
25
Pseudo $R^2$ = 0.03
p = 5.81e-01
Primary school
Medium school
High school
High school diploma
1st year of college
2nd year of college (bachelor)
3rd year of college (licence)
4th year of college
Graduate (master)
Graduate (PhD)
Reported education level

Word: dual ("dual"; Bachelor) -- n = 129

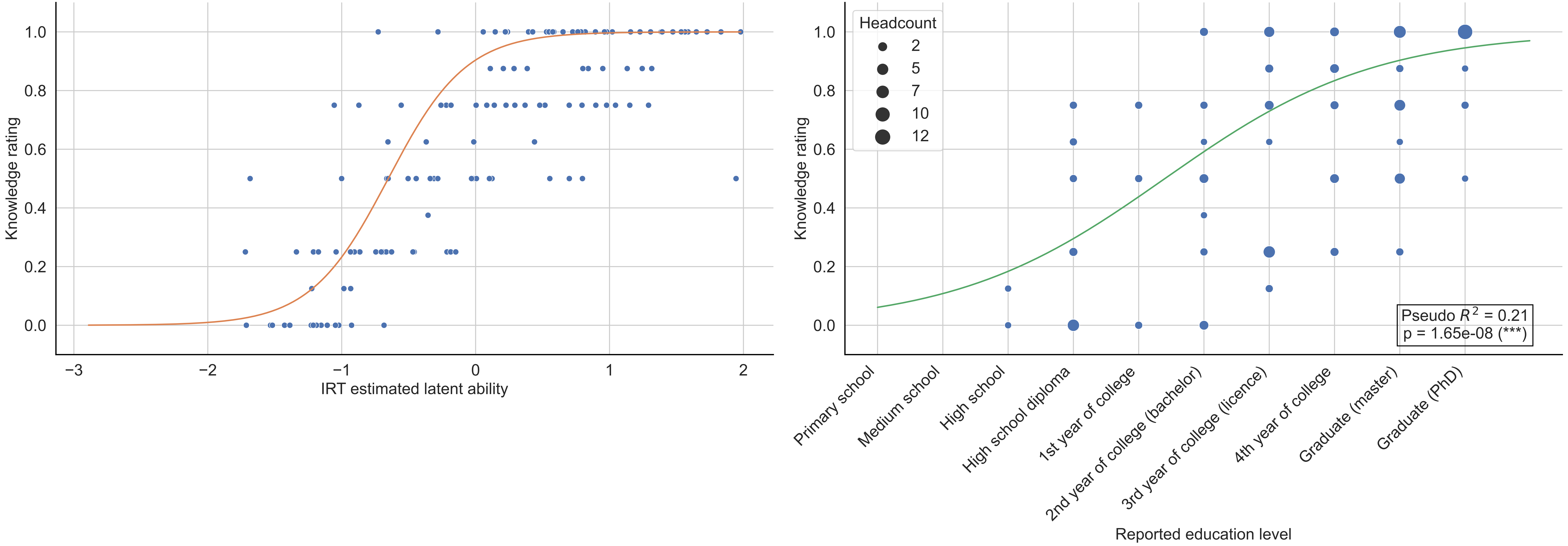

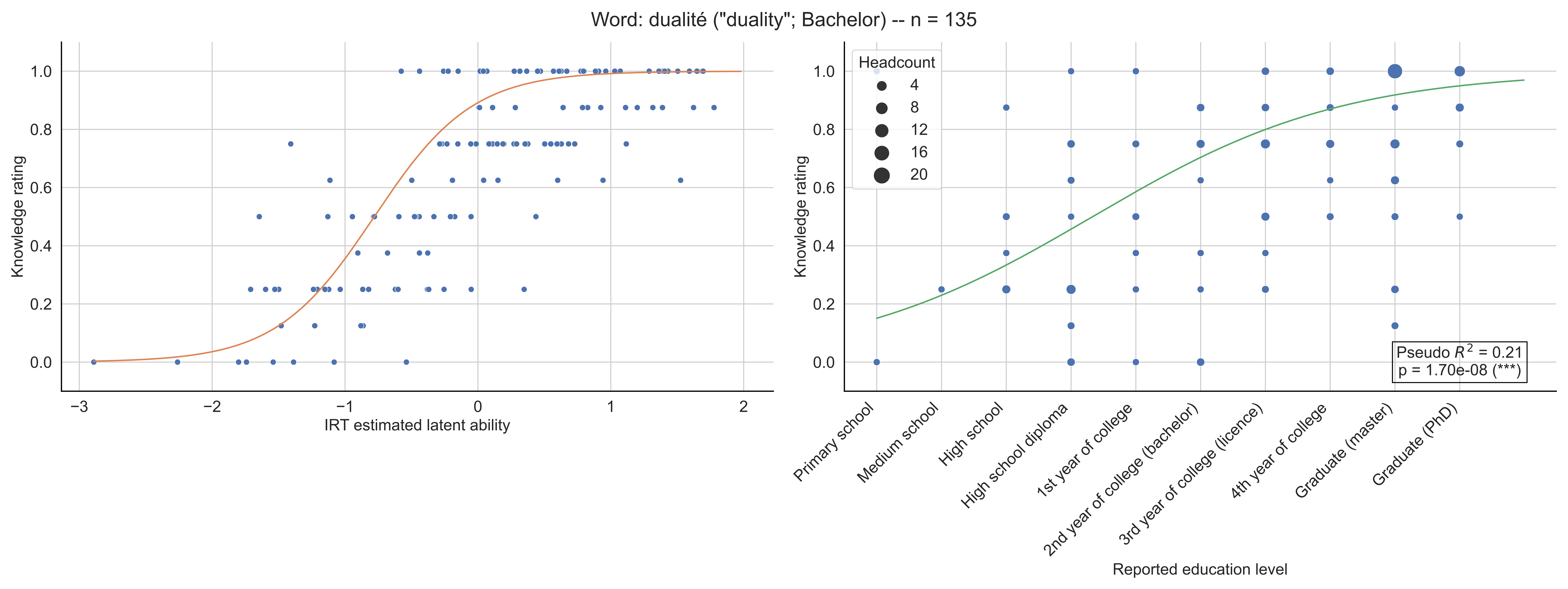

Word: dualité ("duality"; Bachelor) -- n = 135
Knowledge rating
IRT estimated latent ability
Headcount
4
8
12
16
20
Pseudo $R^2$ = 0.21
p = 1.70e-08 (***)
Primary school
Medium school
High school
High school diploma
1st year of college
2nd year of college (bachelor)
3rd year of college (licence)
4th year of college
Graduate (master)
Graduate (PhD)
Reported education level

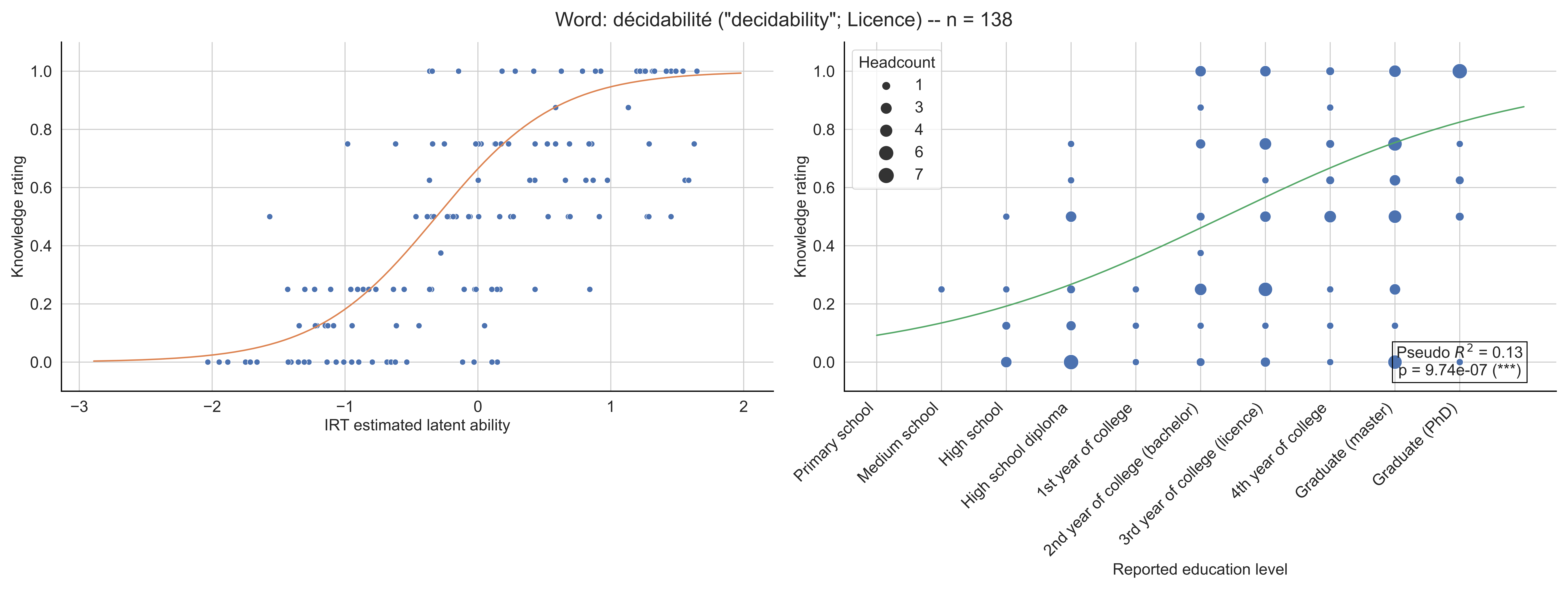

Word: décidabilité ("decidability"; Licence) -- n = 138
Knowledge rating
IRT estimated latent ability
Headcount
1
3
4
6
7
Pseudo $R^2$ = 0.13
p = 9.74e-07 (***)
Primary school
Medium school
High school
High school diploma
1st year of college
2nd year of college (bachelor)
3rd year of college (licence)
4th year of college
Graduate (master)
Graduate (PhD)
Reported education level

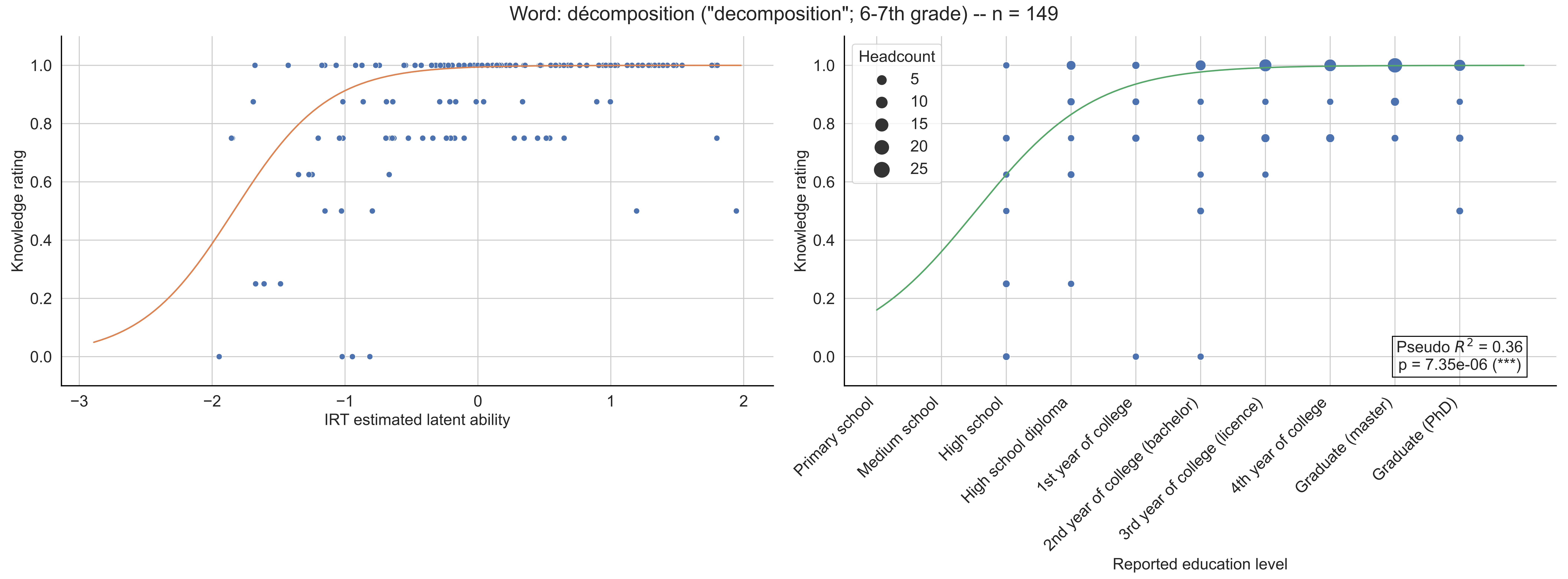

Word: décomposition ("decomposition"; 6-7th grade) -- n = 149
Knowledge rating
IRT estimated latent ability
Headcount
5
10
15
20
25
Pseudo $R^2$ = 0.36
p = 7.35e-06 (***)
Primary school
Medium school
High school
High school diploma
1st year of college
2nd year of college (bachelor)
3rd year of college (licence)
4th year of college
Graduate (master)
Graduate (PhD)
Reported education level

Word: dénombrement ("counting"; 8-9th grade) -- n = 164

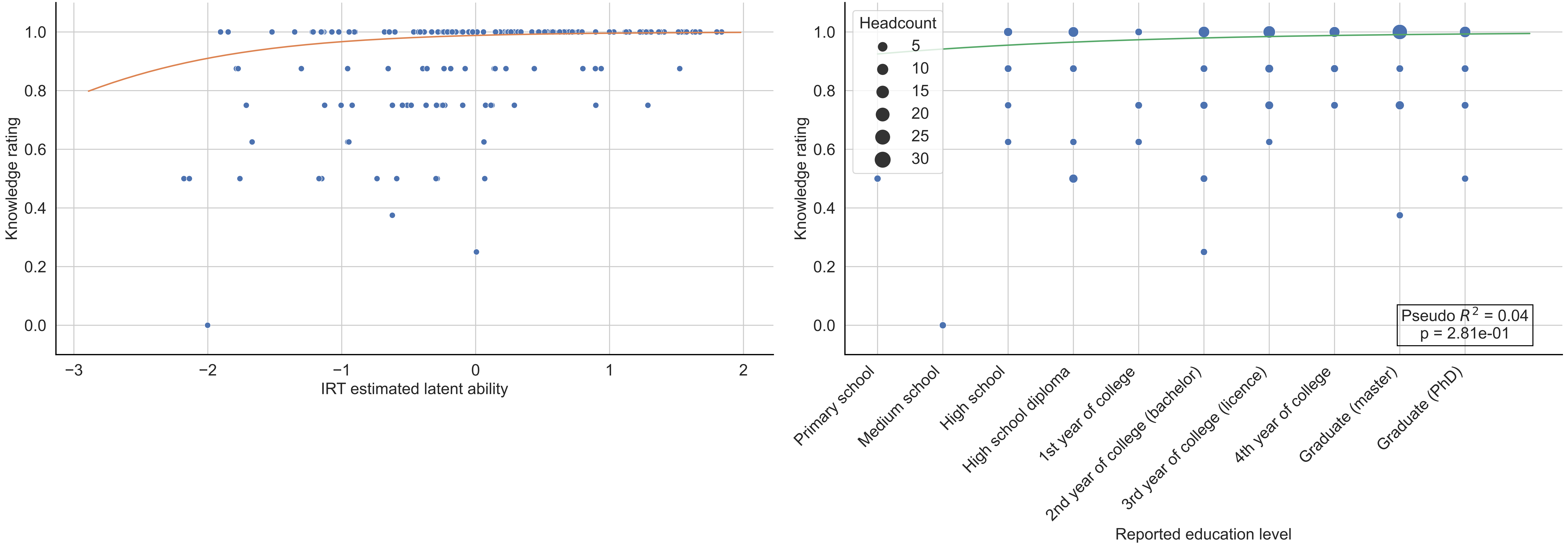

Word: dérivation ("derivation, differentiation"; 11-12th grade) -- n = 148

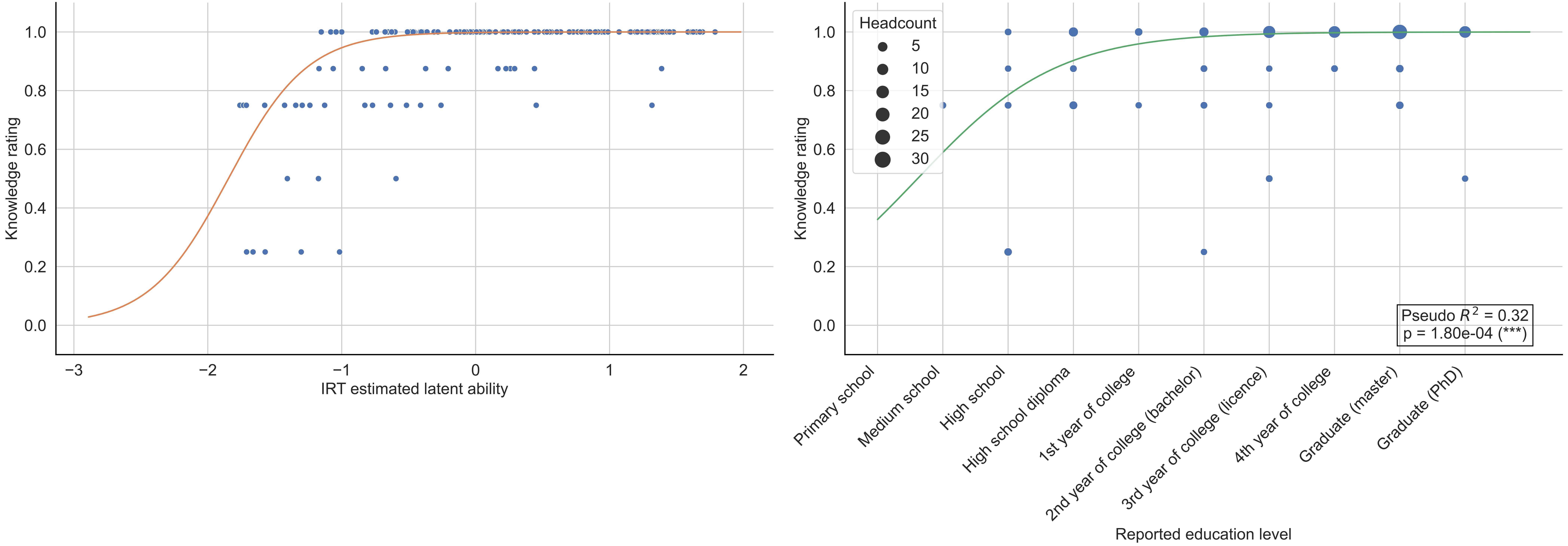

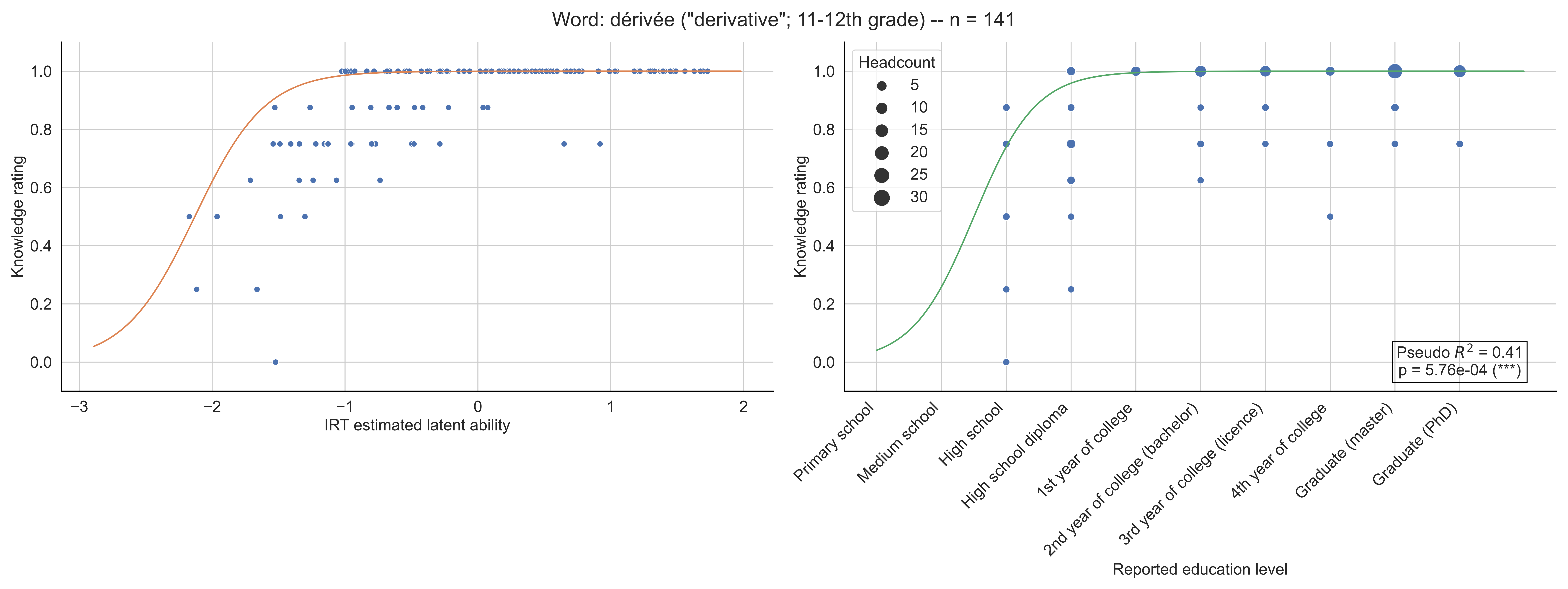

Word: dérivée ("derivative"; 11-12th grade) -- n = 141
Knowledge rating
IRT estimated latent ability
Headcount
5
10
15
20
25
30
Pseudo $R^2$ = 0.41
p = 5.76e-04 (***)
Primary school
Medium school
High school
High school diploma
1st year of college
2nd year of college (bachelor)
3rd year of college (licence)
4th year of college
Graduate (master)
Graduate (PhD)
Reported education level

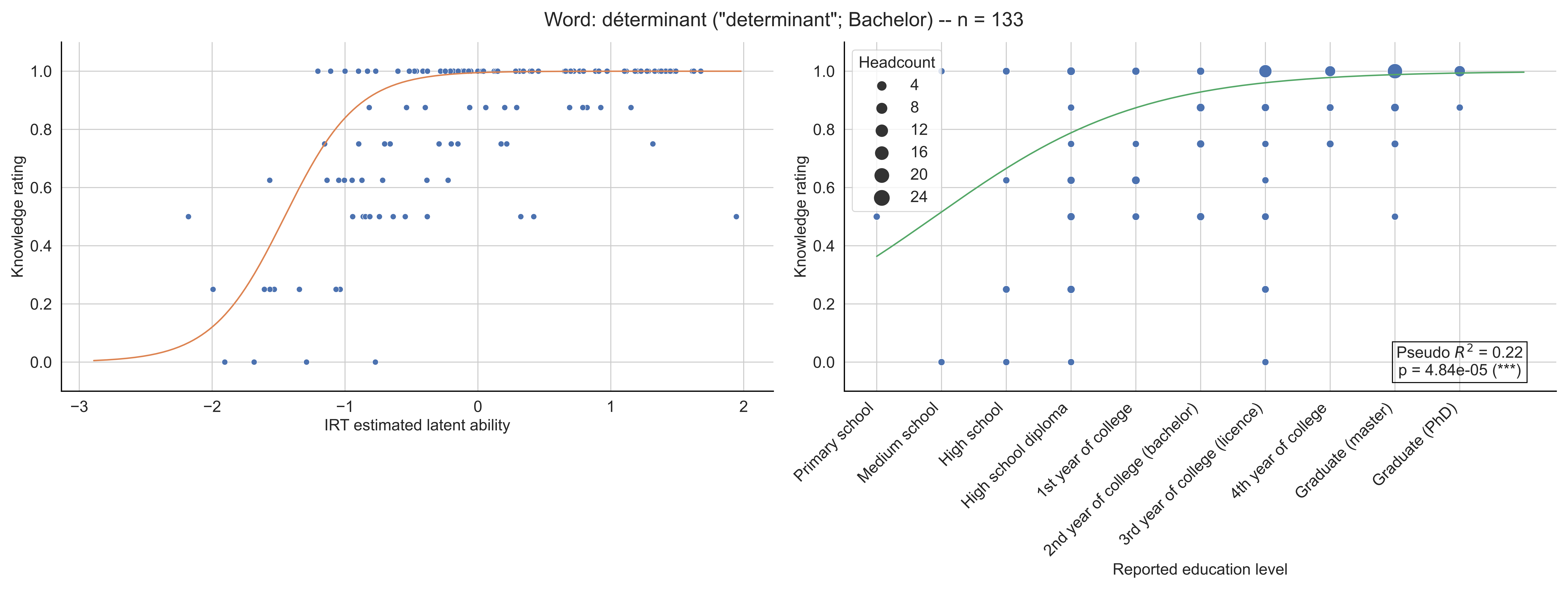

Word: déterminant ("determinant"; Bachelor) -- n = 133
Knowledge rating
IRT estimated latent ability
Headcount
4
8
12
16
20
24
Pseudo $R^2$ = 0.22
p = 4.84e-05 (***)
Primary school
Medium school
High school
High school diploma
1st year of college
2nd year of college (bachelor)
3rd year of college (licence)
4th year of college
Graduate (master)
Graduate (PhD)
Reported education level

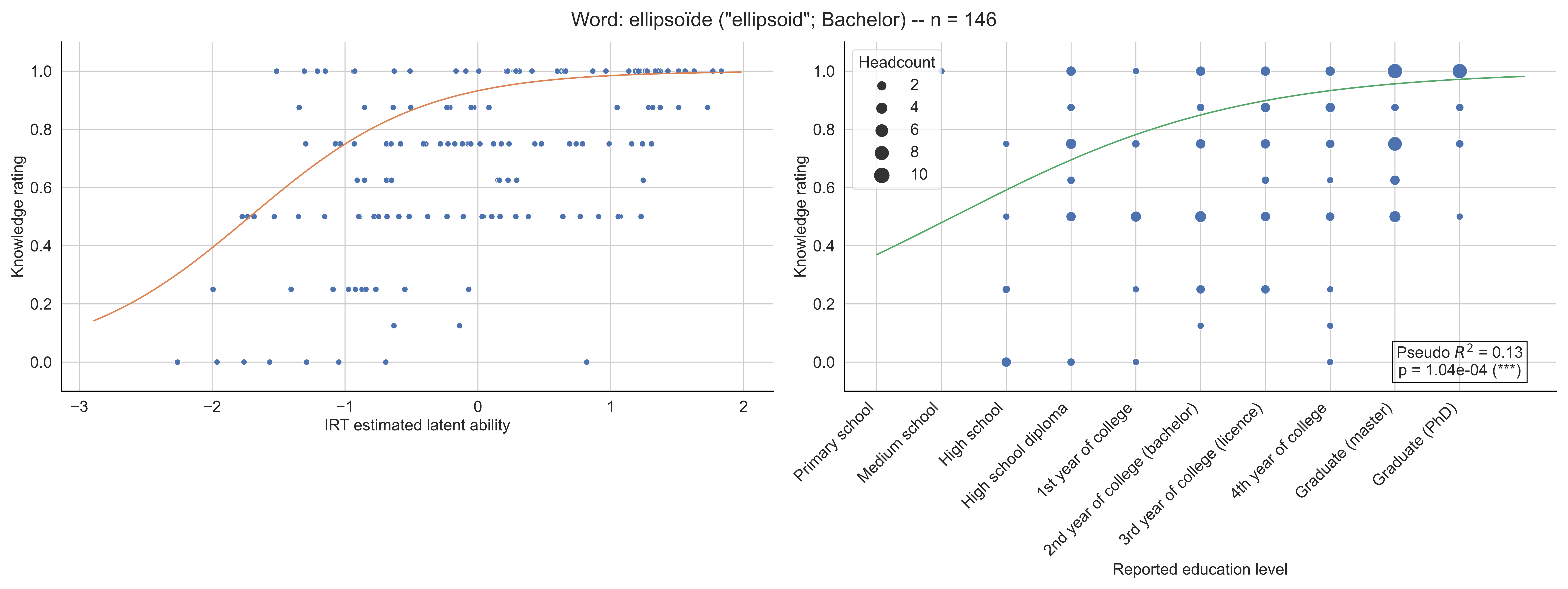
Word: ellipsoïde ("ellipsoid"; Bachelor) -- n = 146
Knowledge rating
IRT estimated latent ability
Headcount
2
4
6
8
10
Pseudo $R^2$ = 0.13
p = 1.04e-04 (***)
Primary school
Medium school
High school
High school diploma
1st year of college
2nd year of college (bachelor)
3rd year of college (licence)
4th year of college
Graduate (master)
Graduate (PhD)
Reported education level

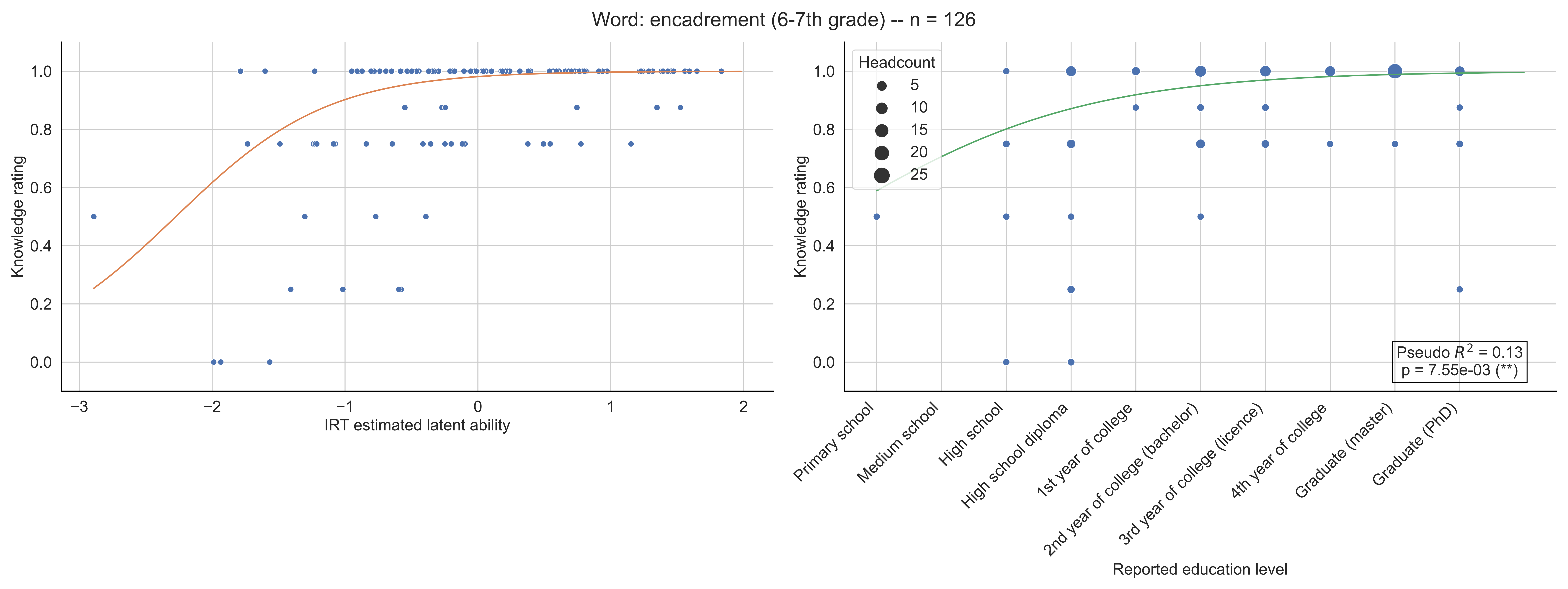
Word: encadrement (6-7th grade) -- n = 126
Knowledge rating
IRT estimated latent ability
Headcount
5
10
15
20
25
Pseudo $R^2$ = 0.13
p = 7.55e-03 (**)
Primary school
Medium school
High school
High school diploma
1st year of college
2nd year of college (bachelor)
3rd year of college (licence)
4th year of college
Graduate (master)
Graduate (PhD)
Reported education level

Word: endomorphisme ("endomorphism"; Bachelor) -- n = 119

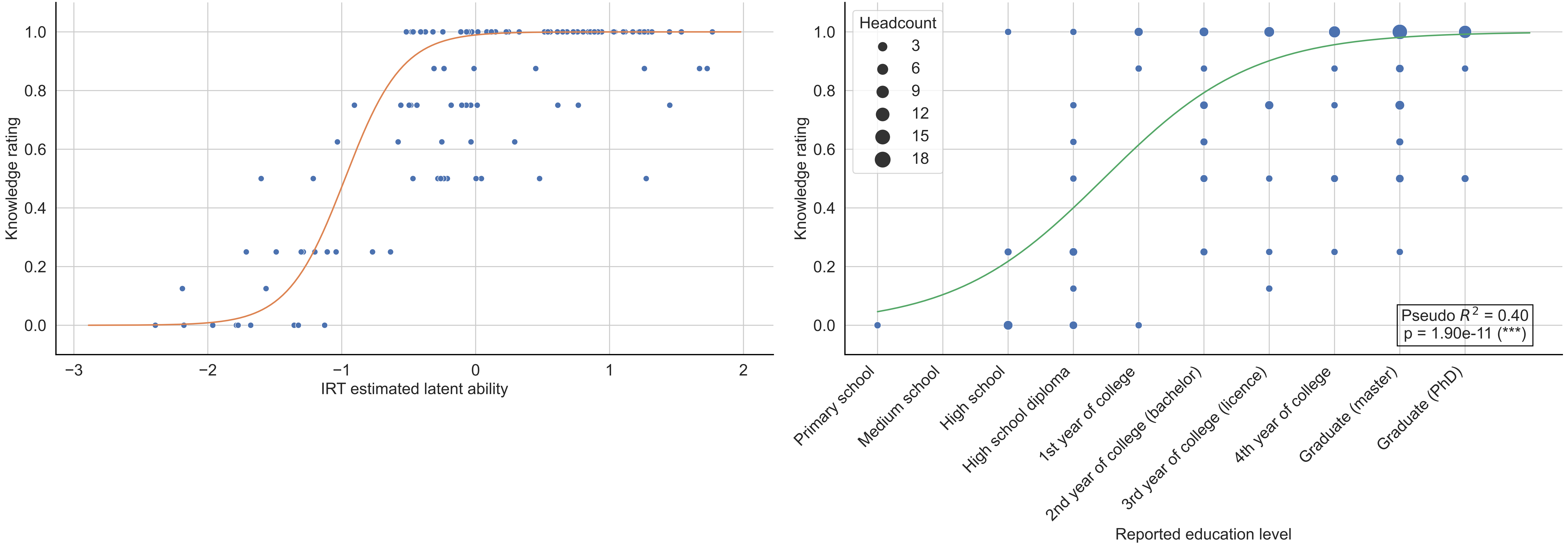

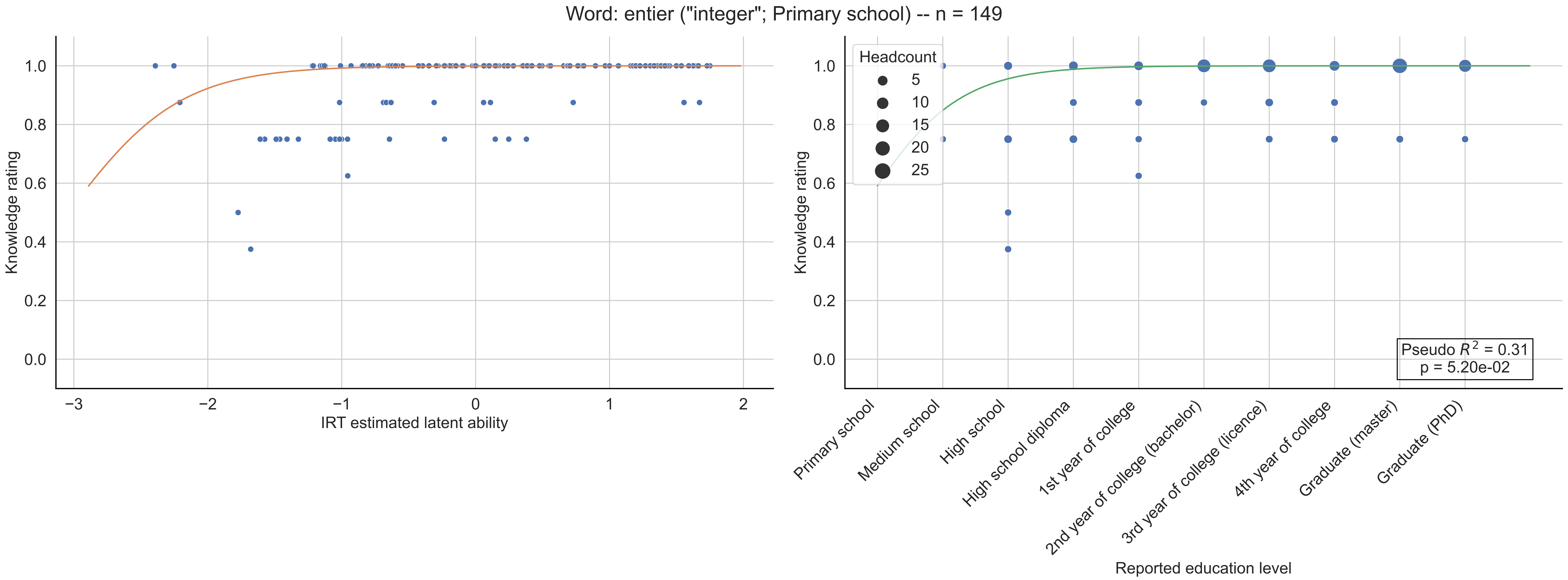

Word: entier ("integer"; Primary school) -- n = 149
Knowledge rating
IRT estimated latent ability
Headcount
5
10
15
20
25
Pseudo $R^2$ = 0.31
p = 5.20e-02
Primary school
Medium school
High school
High school diploma
1st year of college
2nd year of college (bachelor)
3rd year of college (licence)
4th year of college
Graduate (master)
Graduate (PhD)
Reported education level

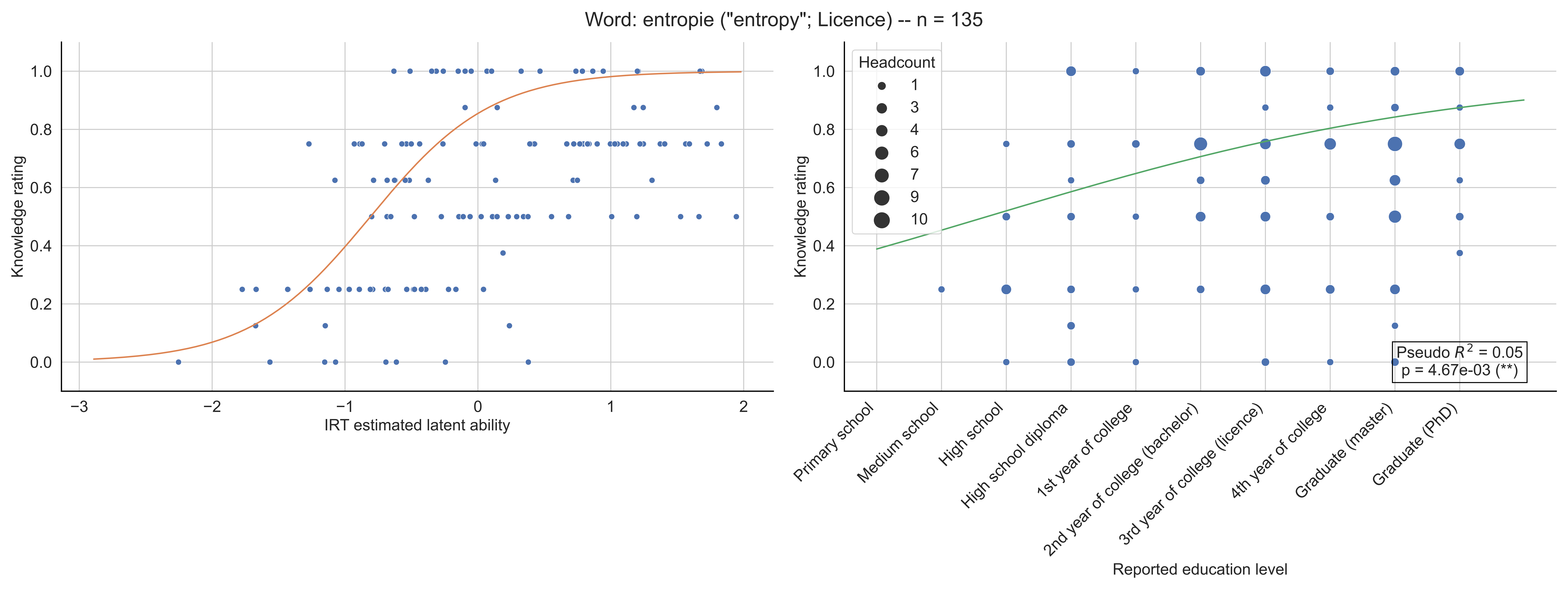

Word: entropie ("entropy"; Licence) -- n = 135
Knowledge rating
IRT estimated latent ability
Headcount
1
3
4
6
7
9
10
Pseudo $R^2$ = 0.05
p = 4.67e-03 (**)
Primary school
Medium school
High school
High school diploma
1st year of college
2nd year of college (bachelor)
3rd year of college (licence)
4th year of college
Graduate (master)
Graduate (PhD)
Reported education level

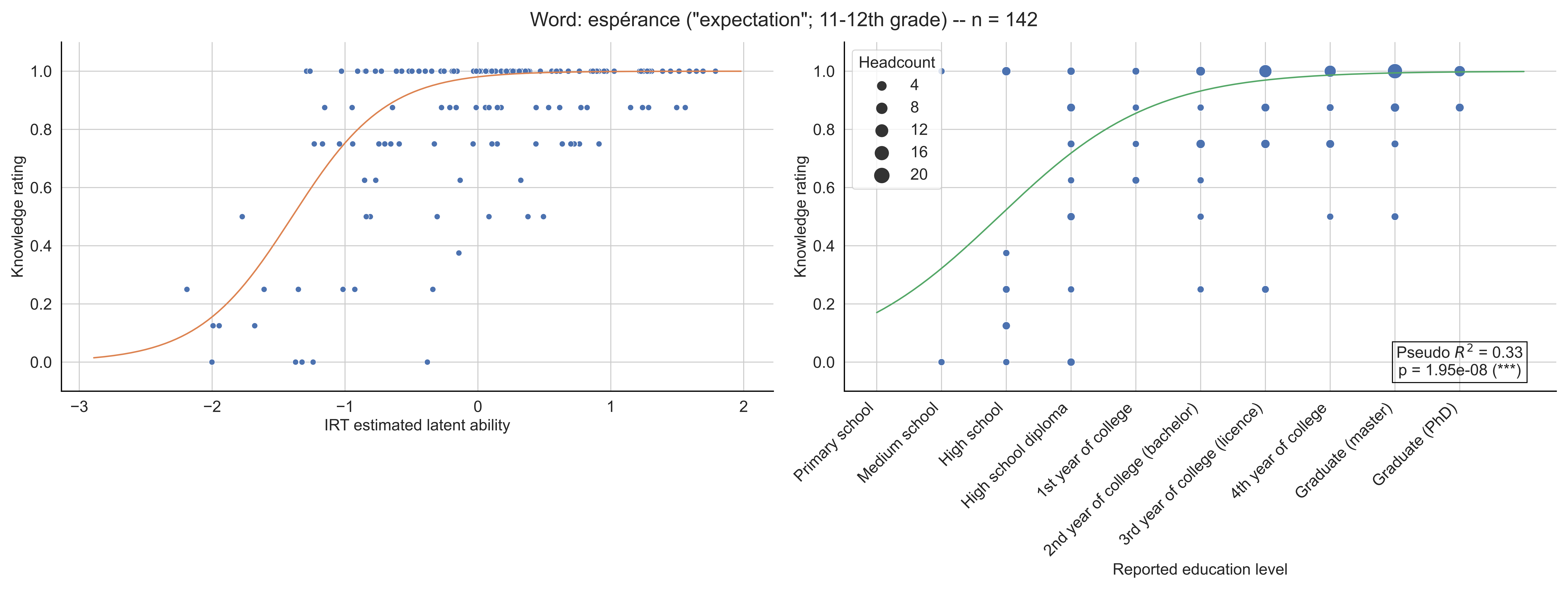

Word: espérance ("expectation"; 11-12th grade) -- n = 142
Knowledge rating
IRT estimated latent ability
Headcount
4
8
12
16
20
Pseudo $R^2$ = 0.33
p = 1.95e-08 (***)
Primary school
Medium school
High school
High school diploma
1st year of college
2nd year of college (bachelor)
3rd year of college (licence)
4th year of college
Graduate (master)
Graduate (PhD)
Reported education level

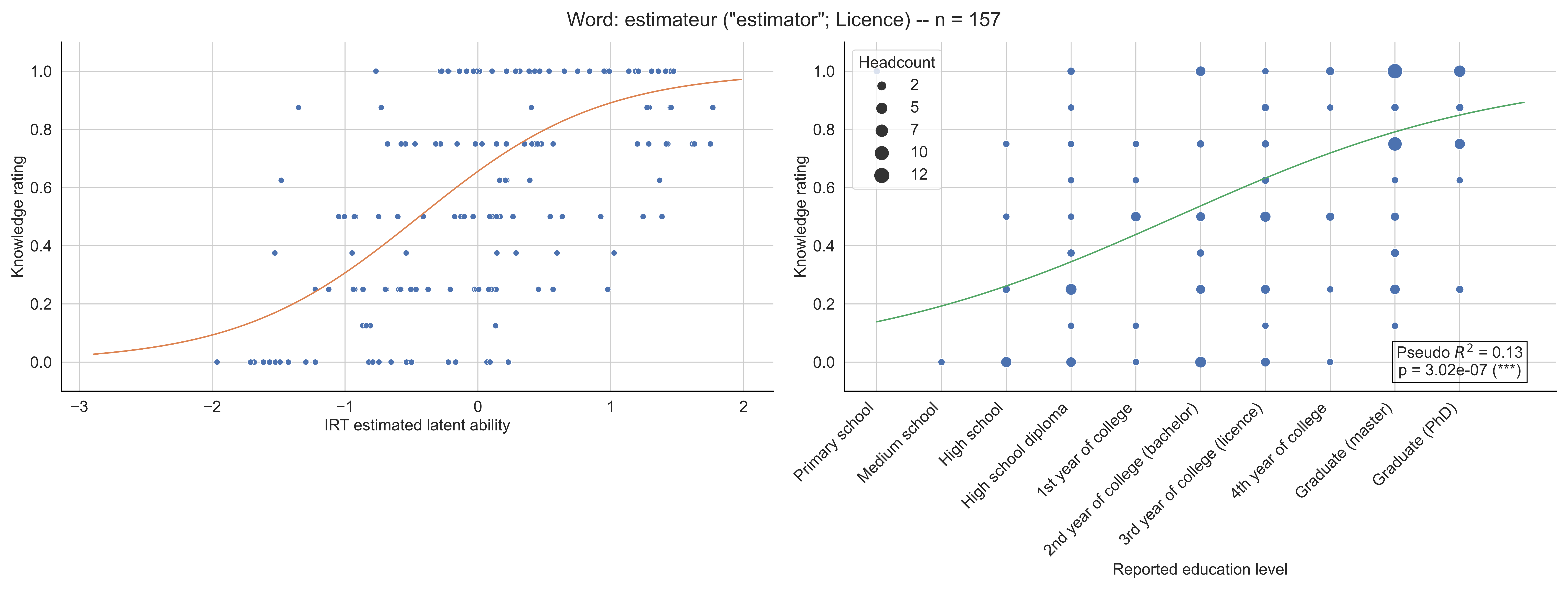
Word: estimateur ("estimator"; Licence) -- n = 157
Knowledge rating
IRT estimated latent ability
Headcount
2
5
7
10
12
Pseudo $R^2$ = 0.13
p = 3.02e-07 (***)
Primary school
Medium school
High school
High school diploma
1st year of college
2nd year of college (bachelor)
3rd year of college (licence)
4th year of college
Graduate (master)
Graduate (PhD)
Reported education level

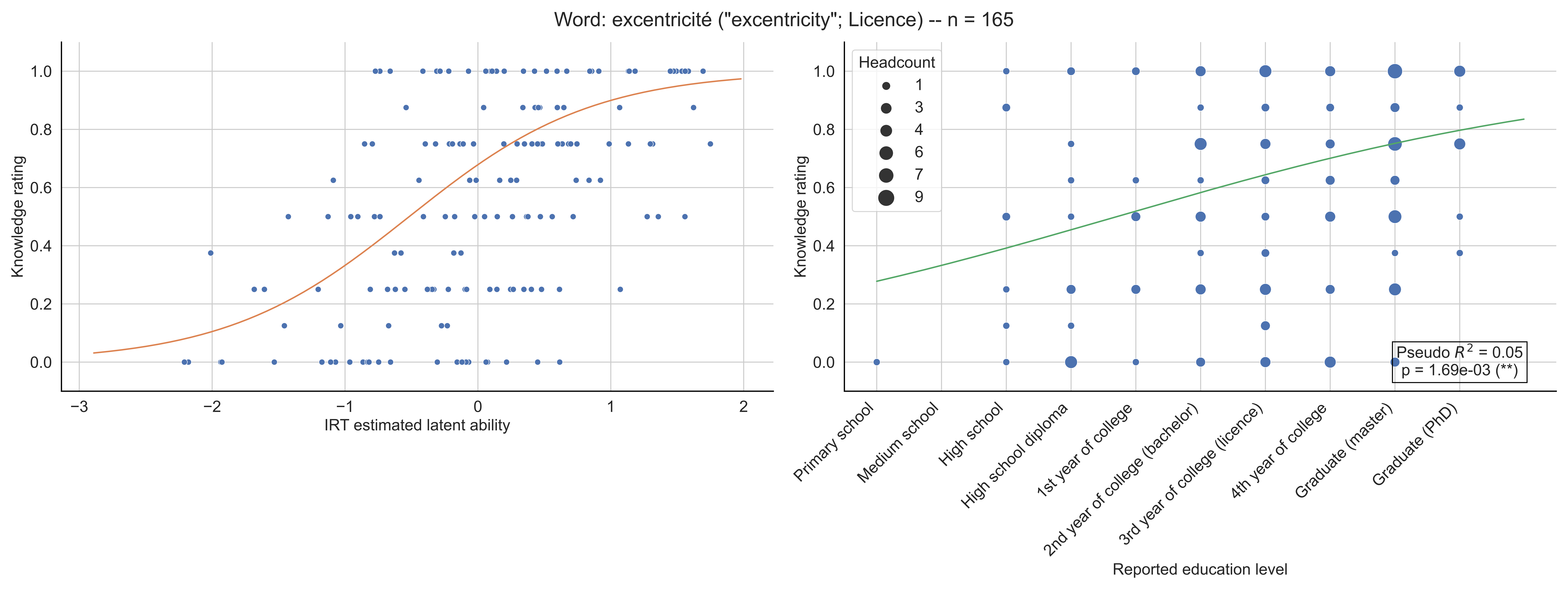

Word: excentricité ("excentricity"; Licence) -- n = 165
Knowledge rating
IRT estimated latent ability
Headcount
1
3
4
6
7
9
Pseudo $R^2$ = 0.05
p = 1.69e-03 (**)
Primary school
Medium school
High school
High school diploma
1st year of college
2nd year of college (bachelor)
3rd year of college (licence)
4th year of college
Graduate (master)
Graduate (PhD)
Reported education level

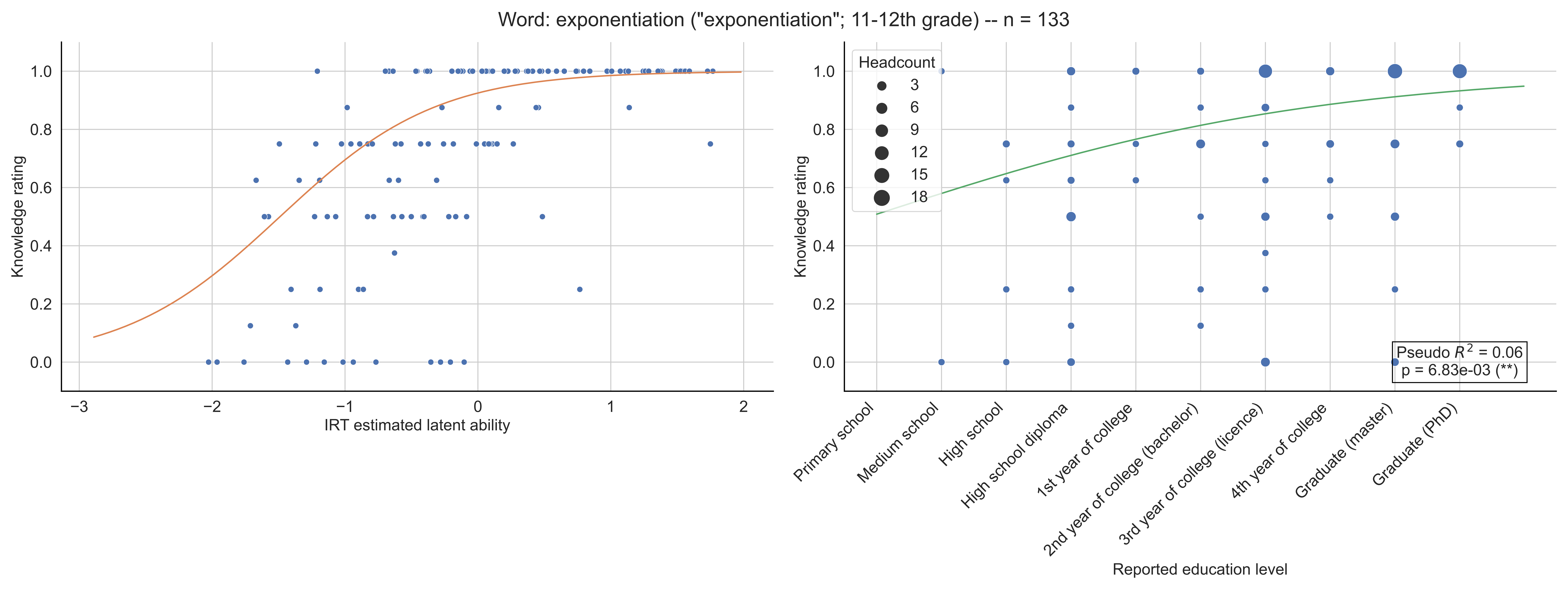

Word: exponentiation ("exponentiation"; 11-12th grade) -- n = 133
Knowledge rating
IRT estimated latent ability
Headcount
3
6
9
12
15
18
Primary school
Medium school
High school
High school diploma
1st year of college
2nd year of college (bachelor)
3rd year of college (licence)
4th year of college
Graduate (master)
Graduate (PhD)
Reported education level
Pseudo $R^2$ = 0.06
p = 6.83e-03 (**)

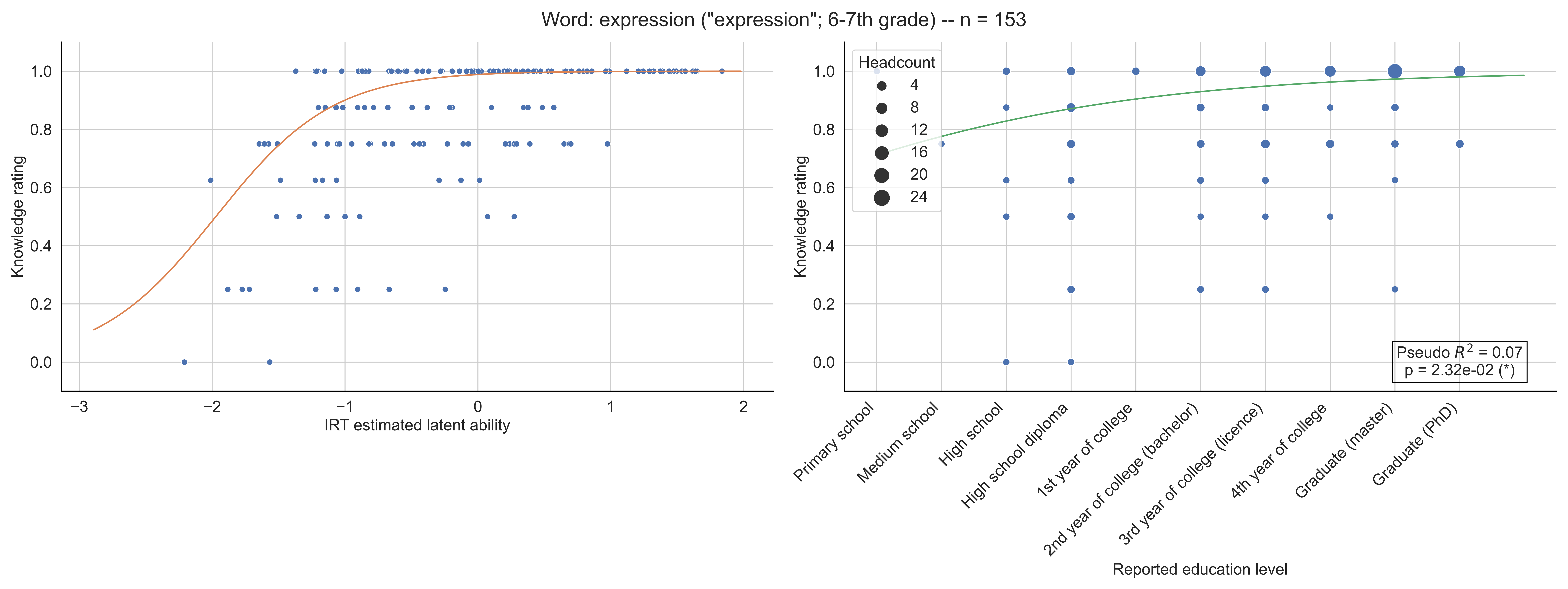

Word: expression ("expression"; 6-7th grade) -- n = 153
Knowledge rating
IRT estimated latent ability
Headcount
4
8
12
16
20
24
Primary school
Medium school
High school
High school diploma
1st year of college
2nd year of college (bachelor)
3rd year of college (licence)
4th year of college
Graduate (master)
Graduate (PhD)
Reported education level
Pseudo $R^2$ = 0.07
p = 2.32e-02 (*)

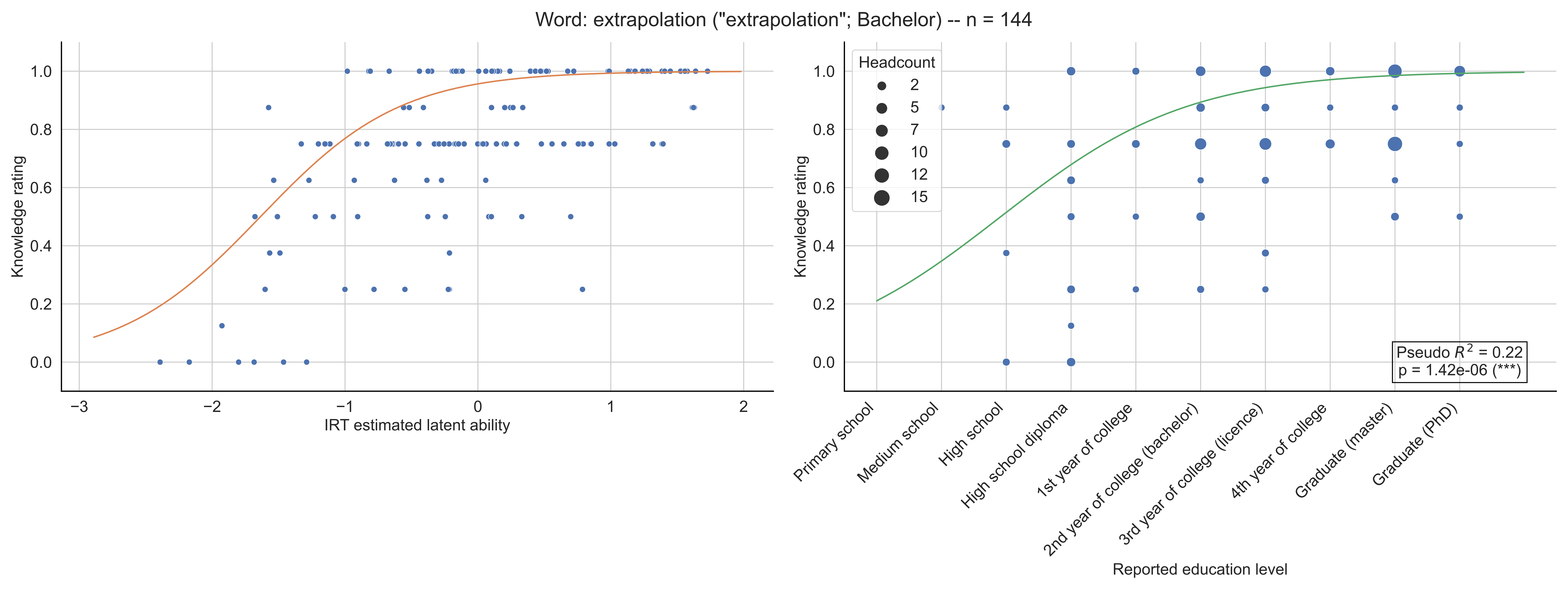

Word: extrapolation ("extrapolation"; Bachelor) -- n = 144
Knowledge rating
IRT estimated latent ability
Headcount
2
5
7
10
12
15
Pseudo $R^2$ = 0.22
p = 1.42e-06 (***)
Primary school
Medium school
High school
High school diploma
1st year of college
2nd year of college (bachelor)
3rd year of college (licence)
4th year of college
Graduate (master)
Graduate (PhD)
Reported education level

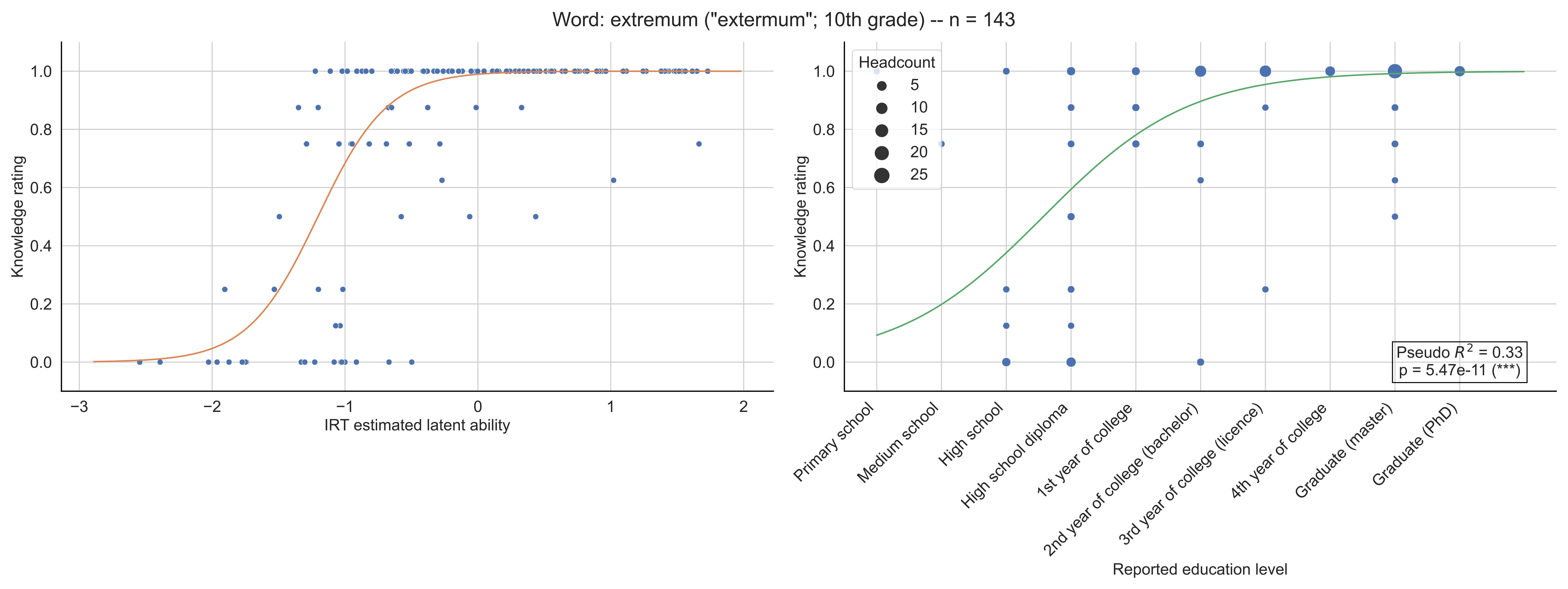

Word: extremum ("extermum"; 10th grade) -- n = 143
Knowledge rating
IRT estimated latent ability
Headcount
5
10
15
20
25
Pseudo $R^2$ = 0.33
p = 5.47e-11 (***)
Primary school
Medium school
High school
High school diploma
1st year of college
2nd year of college (bachelor)
3rd year of college (licence)
4th year of college
Graduate (master)
Graduate (PhD)
Reported education level

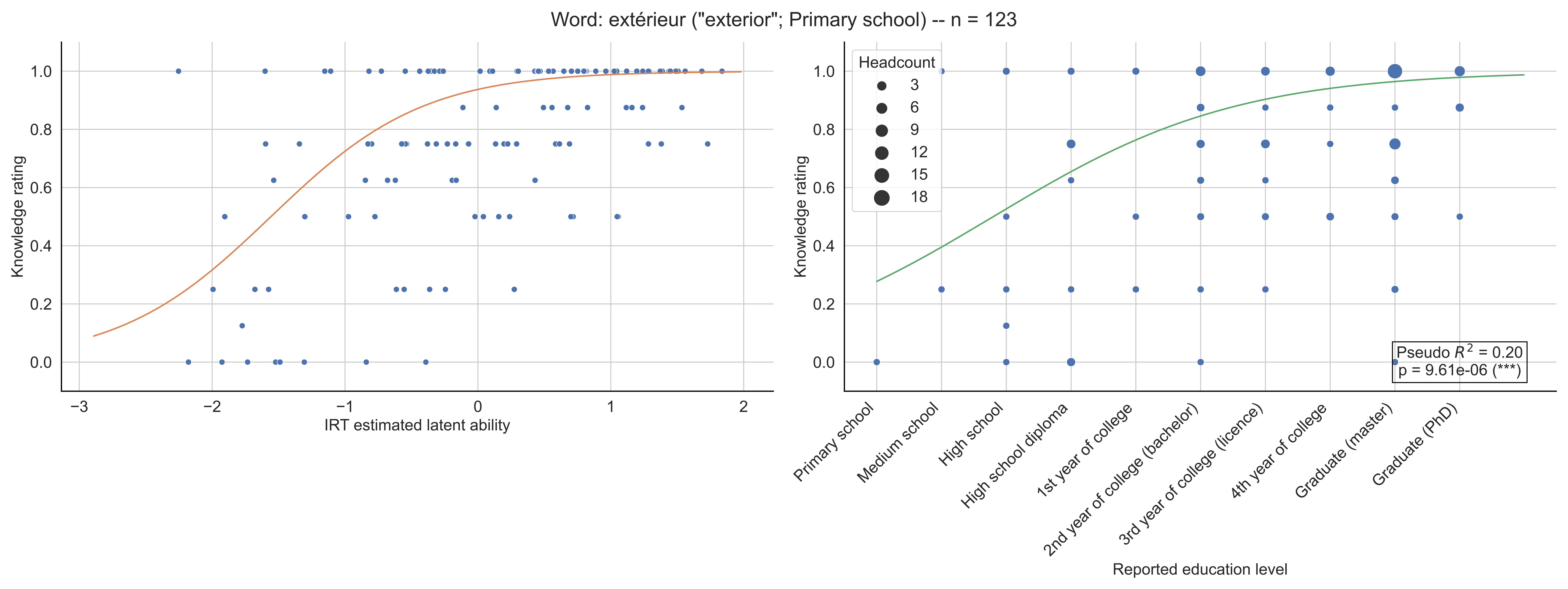
Word: extérieur ("exterior"; Primary school) -- n = 123
Knowledge rating
IRT estimated latent ability
Headcount
3
6
9
12
15
18
Pseudo $R^2$ = 0.20
p = 9.61e-06 (***)
Primary school
Medium school
High school
High school diploma
1st year of college
2nd year of college (bachelor)
3rd year of college (licence)
4th year of college
Graduate (master)
Graduate (PhD)
Reported education level

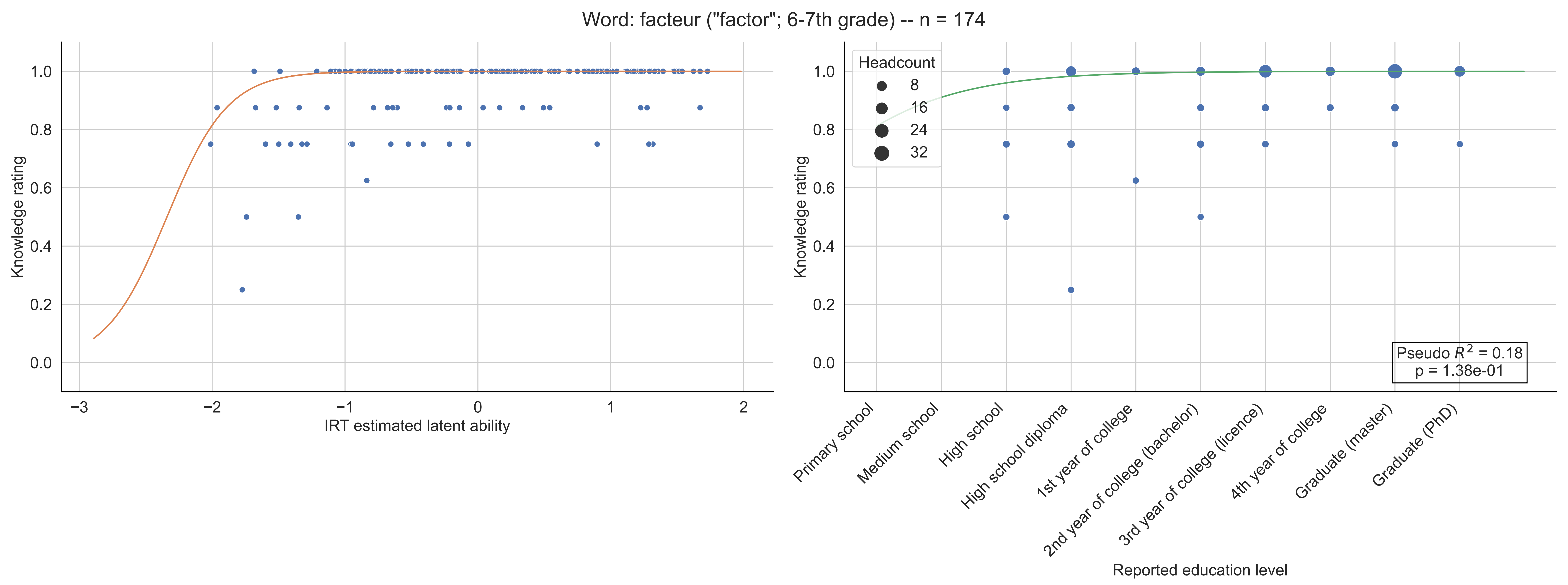
Word: facteur ("factor"; 6-7th grade) -- n = 174
Knowledge rating
IRT estimated latent ability
Headcount
8
16
24
32
Pseudo $R^2$ = 0.18
p = 1.38e-01
Primary school
Medium school
High school
High school diploma
1st year of college
2nd year of college (bachelor)
3rd year of college (licence)
4th year of college
Graduate (master)
Graduate (PhD)
Reported education level

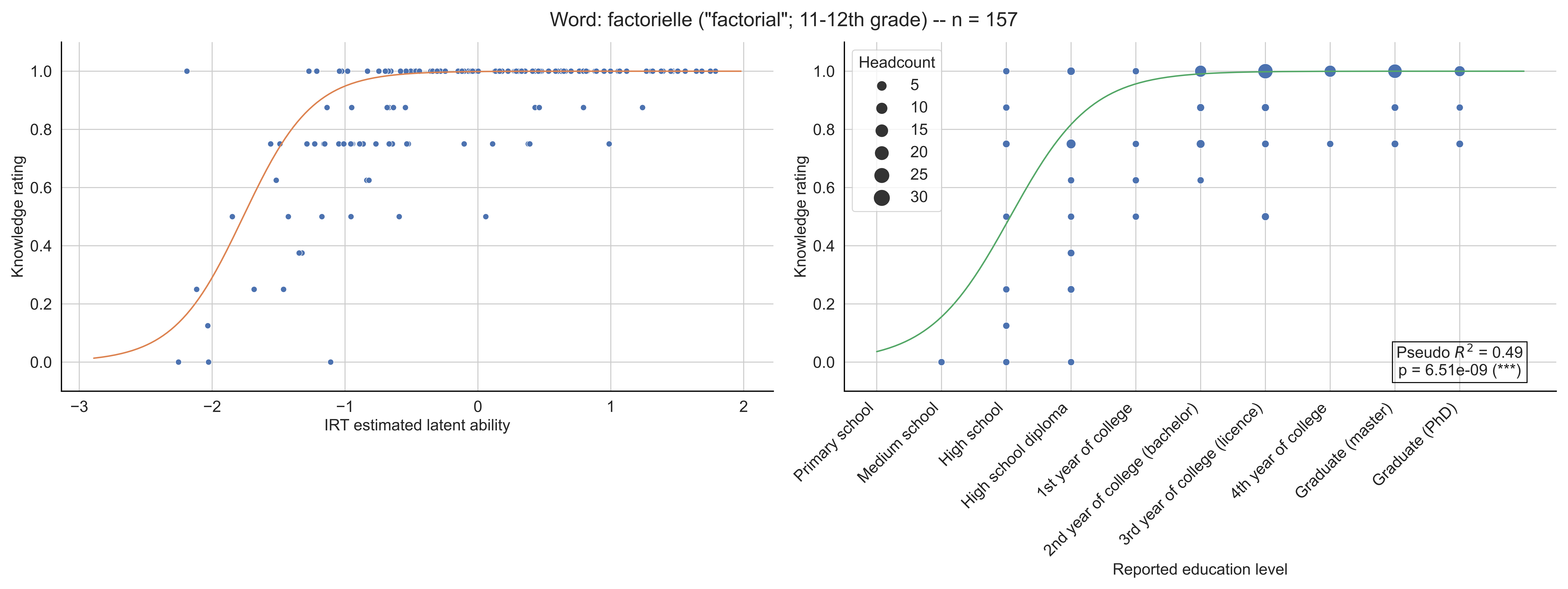

Word: factorielle ("factorial"; 11-12th grade) -- n = 157
Knowledge rating
1.0
0.8
0.6
0.4
0.2
0.0
−3
−2
−1
0
1
2
IRT estimated latent ability
Headcount
5
10
15
20
25
30
Primary school
Medium school
High school
High school diploma
1st year of college
2nd year of college (bachelor)
3rd year of college (licence)
4th year of college
Graduate (master)
Graduate (PhD)
Reported education level
Pseudo $R^2$ = 0.49
p = 6.51e-09 (***)

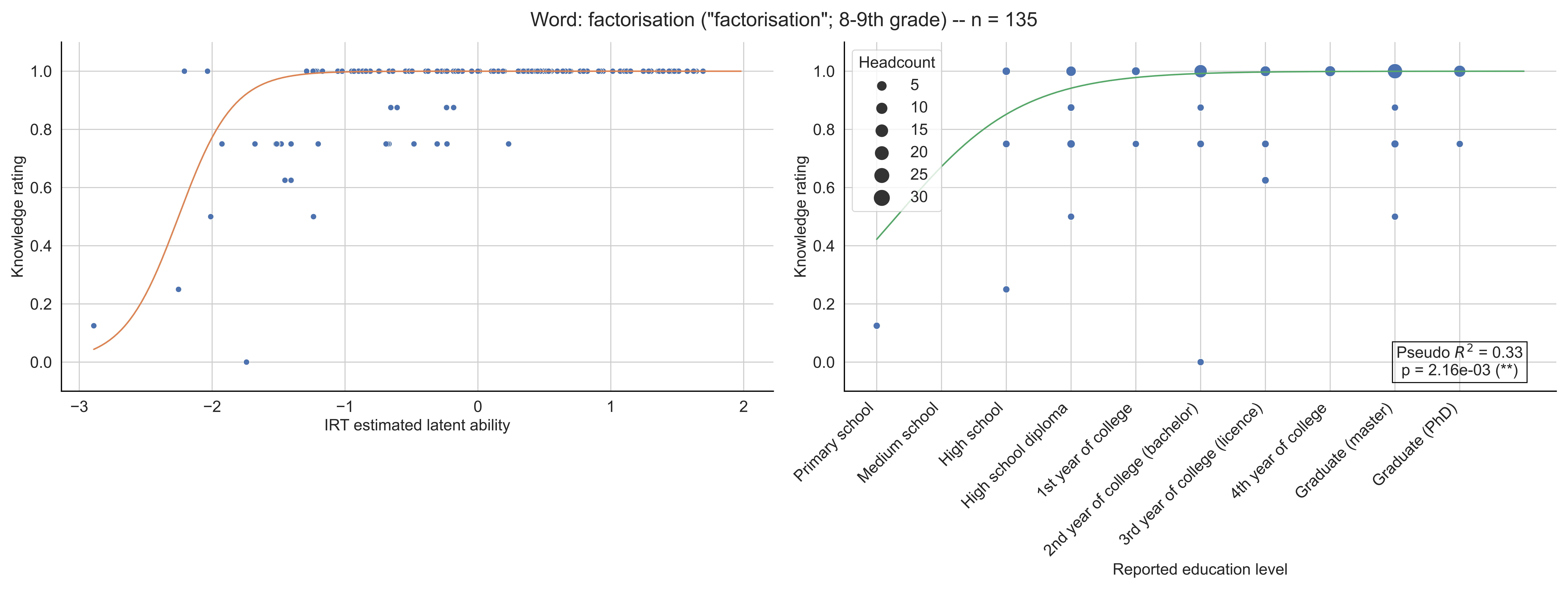

Word: factorisation ("factorisation"; 8-9th grade) -- n = 135
Knowledge rating
IRT estimated latent ability
Headcount
5
10
15
20
25
30
Pseudo $R^2$ = 0.33
p = 2.16e-03 (**)
Primary school
Medium school
High school
High school diploma
1st year of college
2nd year of college (bachelor)
3rd year of college (licence)
4th year of college
Graduate (master)
Graduate (PhD)
Reported education level

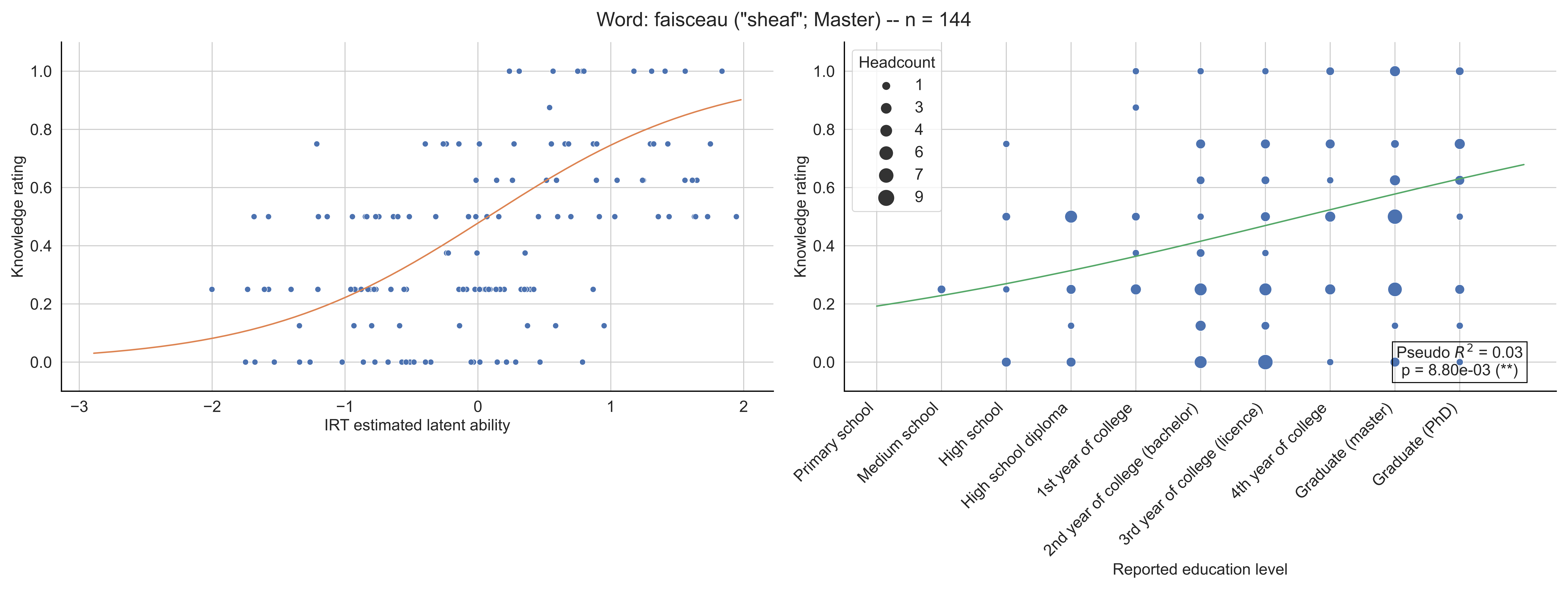

Word: faisceau ("sheaf"; Master) -- n = 144
Knowledge rating
IRT estimated latent ability
Headcount
1
3
4
6
7
9
Primary school
Medium school
High school
High school diploma
1st year of college
2nd year of college (bachelor)
3rd year of college (licence)
4th year of college
Graduate (master)
Graduate (PhD)
Reported education level
Pseudo $R^2$ = 0.03
p = 8.80e-03 (**)

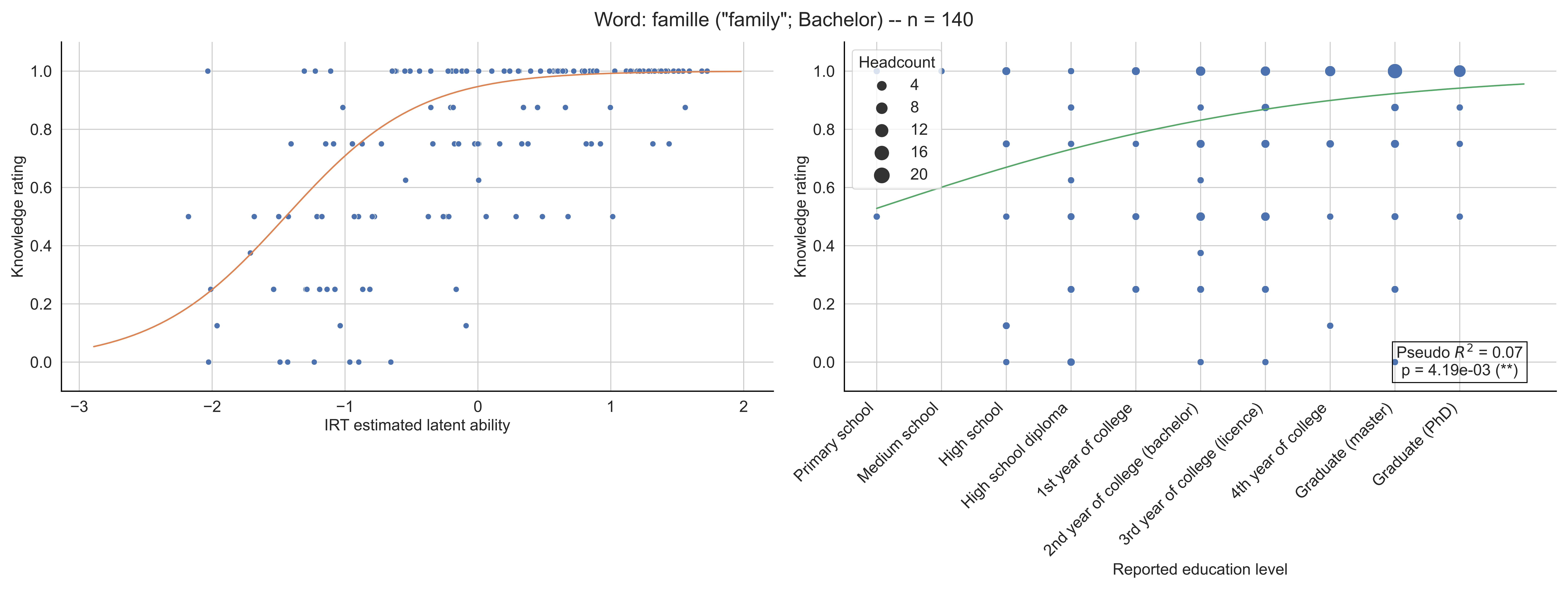

Word: famille ("family"; Bachelor) -- n = 140
Knowledge rating
IRT estimated latent ability
Headcount
4
8
12
16
20
Pseudo $R^2$ = 0.07
p = 4.19e-03 (**)
Primary school
Medium school
High school
High school diploma
1st year of college
2nd year of college (bachelor)
3rd year of college (licence)
4th year of college
Graduate (master)
Graduate (PhD)
Reported education level

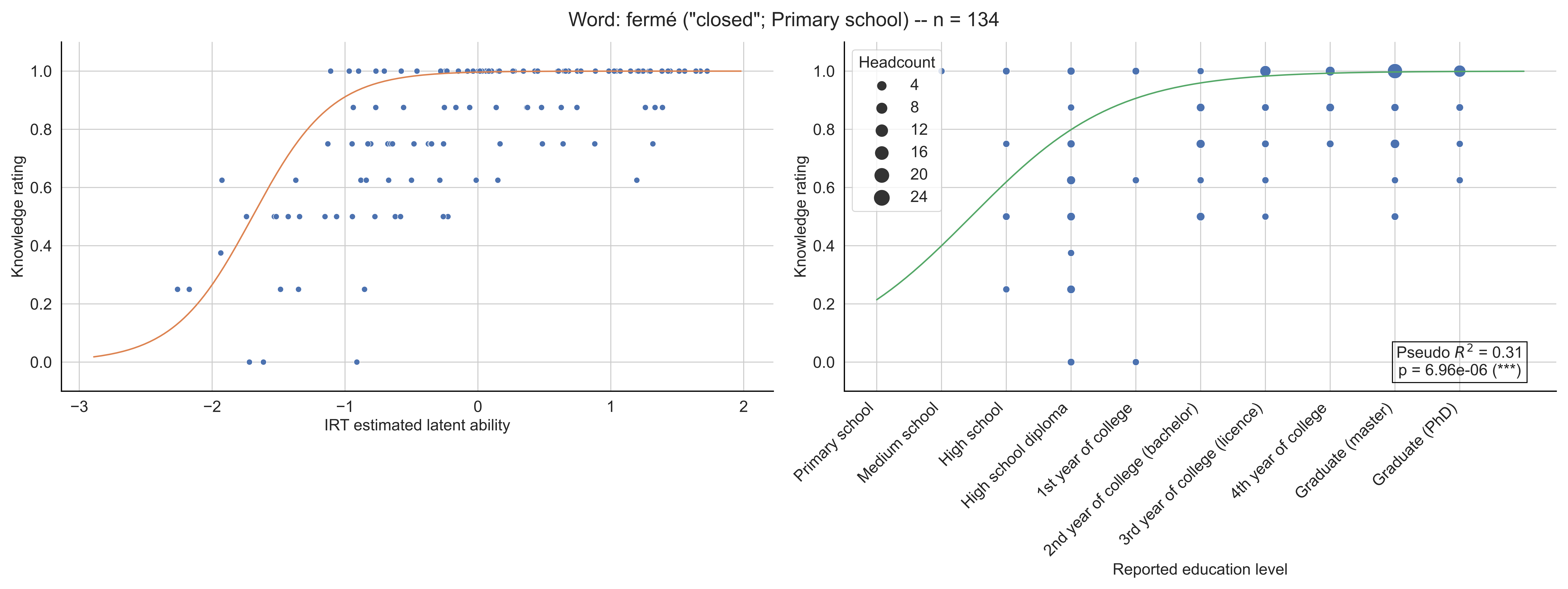
Word: fermé ("closed"; Primary school) -- n = 134
Knowledge rating
IRT estimated latent ability
Headcount
4
8
12
16
20
24
Pseudo $R^2$ = 0.31
p = 6.96e-06 (***)
Primary school
Medium school
High school
High school diploma
1st year of college
2nd year of college (bachelor)
3rd year of college (licence)
4th year of college
Graduate (master)
Graduate (PhD)
Reported education level

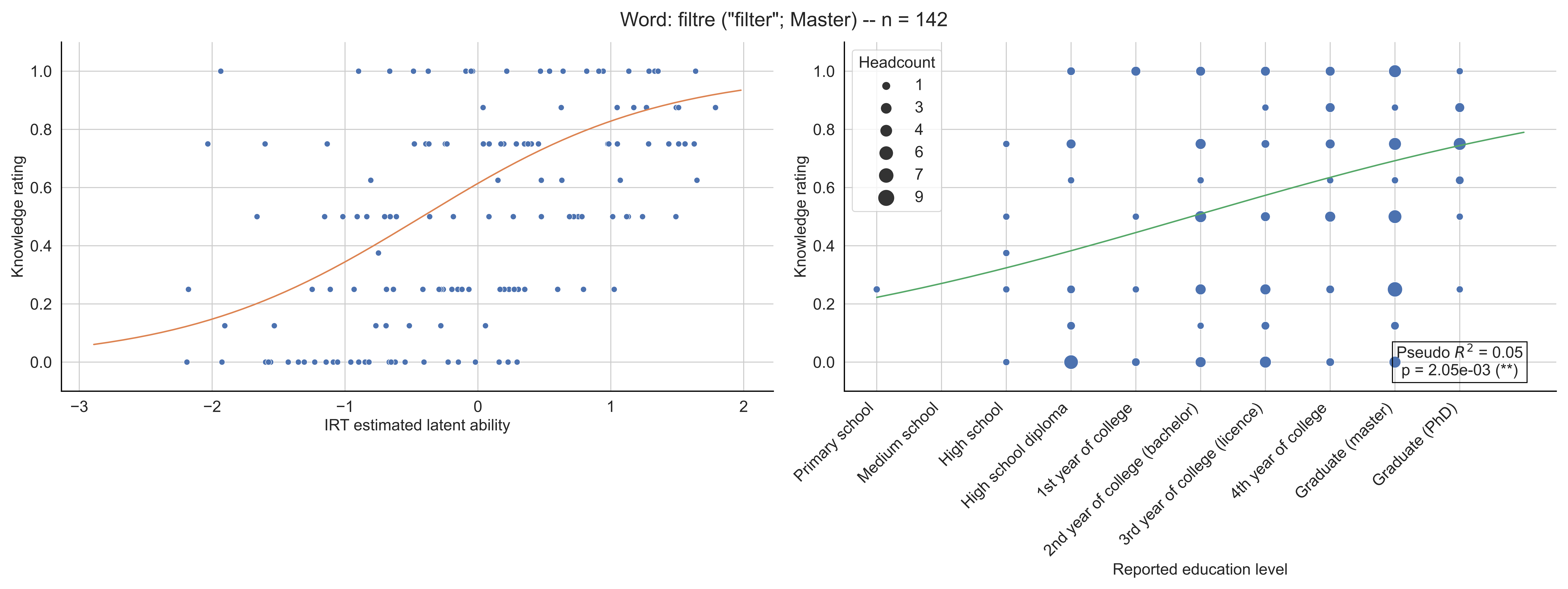

Word: filtre ("filter"; Master) -- n = 142
Knowledge rating
IRT estimated latent ability
Headcount
1
3
4
6
7
9
Pseudo $R^2$ = 0.05
p = 2.05e-03 (**)
Primary school
Medium school
High school
High school diploma
1st year of college
2nd year of college (bachelor)
3rd year of college (licence)
4th year of college
Graduate (master)
Graduate (PhD)
Reported education level

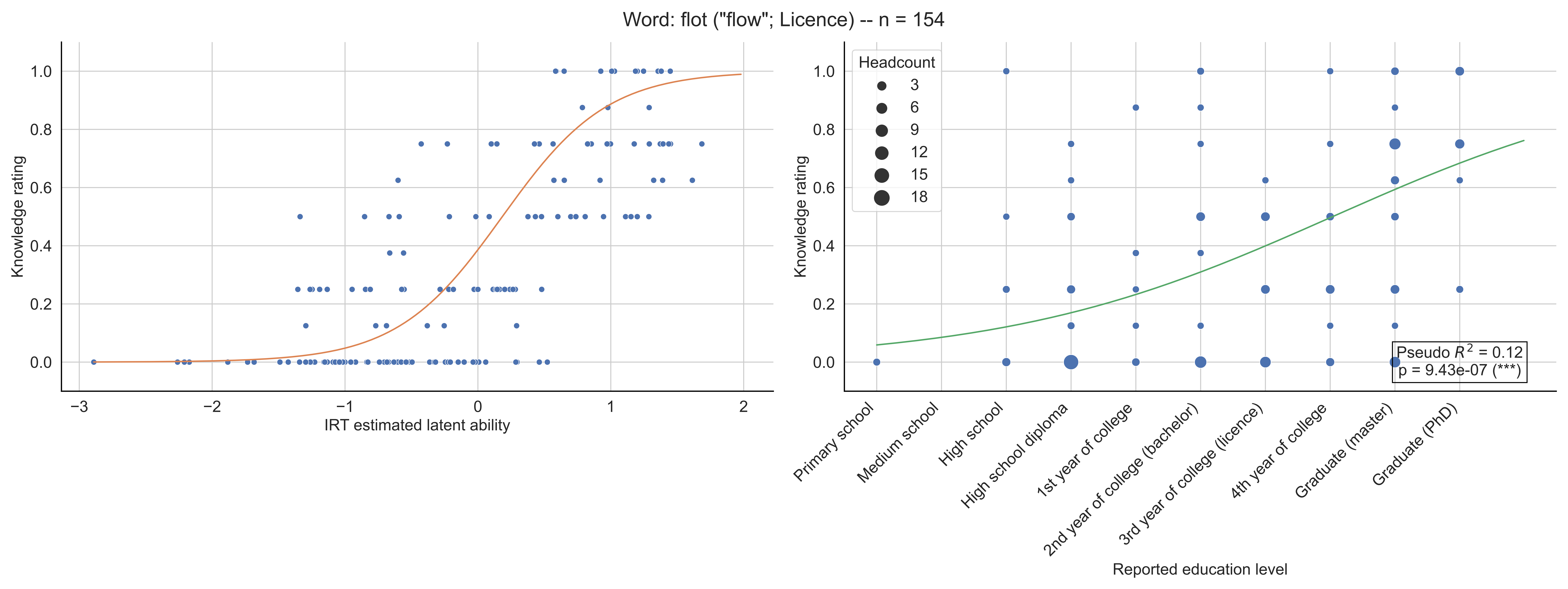

Word: flot ("flow"; Licence) -- n = 154
Knowledge rating
IRT estimated latent ability
Headcount
3
6
9
12
15
18
Pseudo $R^2$ = 0.12
p = 9.43e-07 (***)
Primary school
Medium school
High school
High school diploma
1st year of college
2nd year of college (bachelor)
3rd year of college (licence)
4th year of college
Graduate (master)
Graduate (PhD)
Reported education level

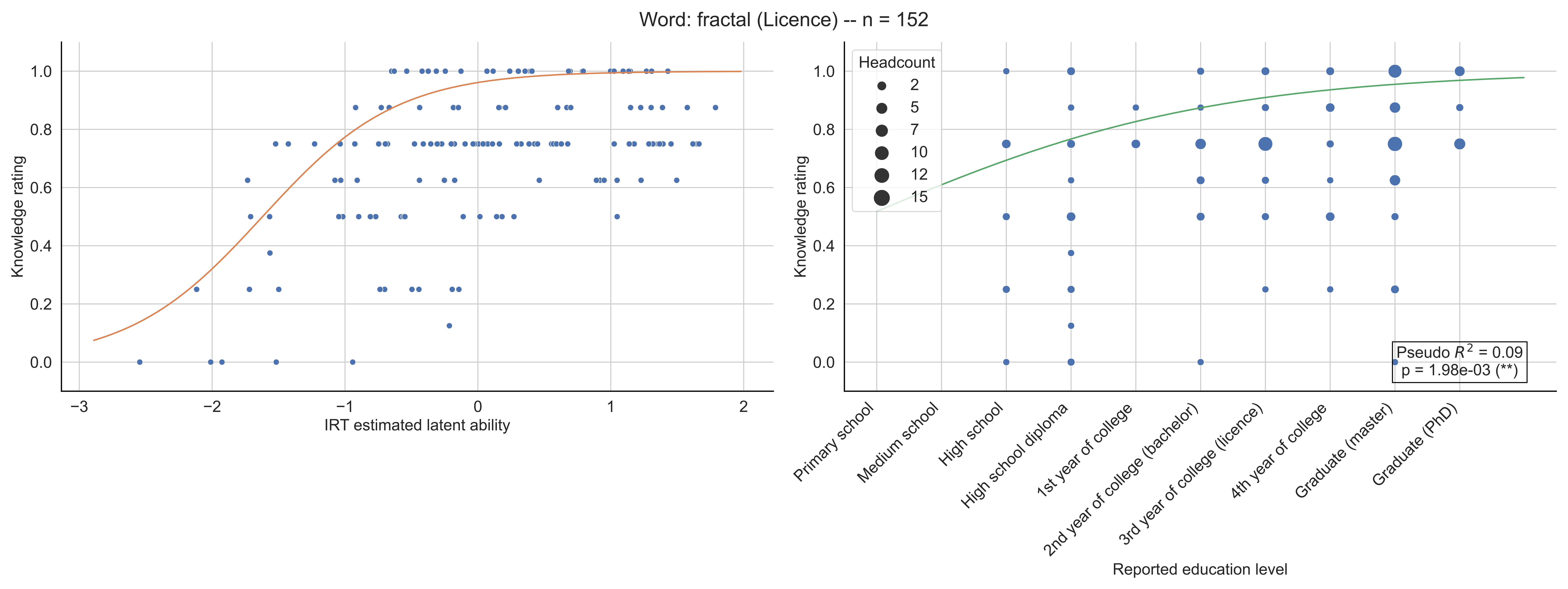
Word: fractal (Licence) -- n = 152
Knowledge rating
IRT estimated latent ability
Headcount
2
5
7
10
12
15
Pseudo $R^2$ = 0.09
p = 1.98e-03 (**)
Primary school
Medium school
High school
High school diploma
1st year of college
2nd year of college (bachelor)
3rd year of college (licence)
4th year of college
Graduate (master)
Graduate (PhD)
Reported education level

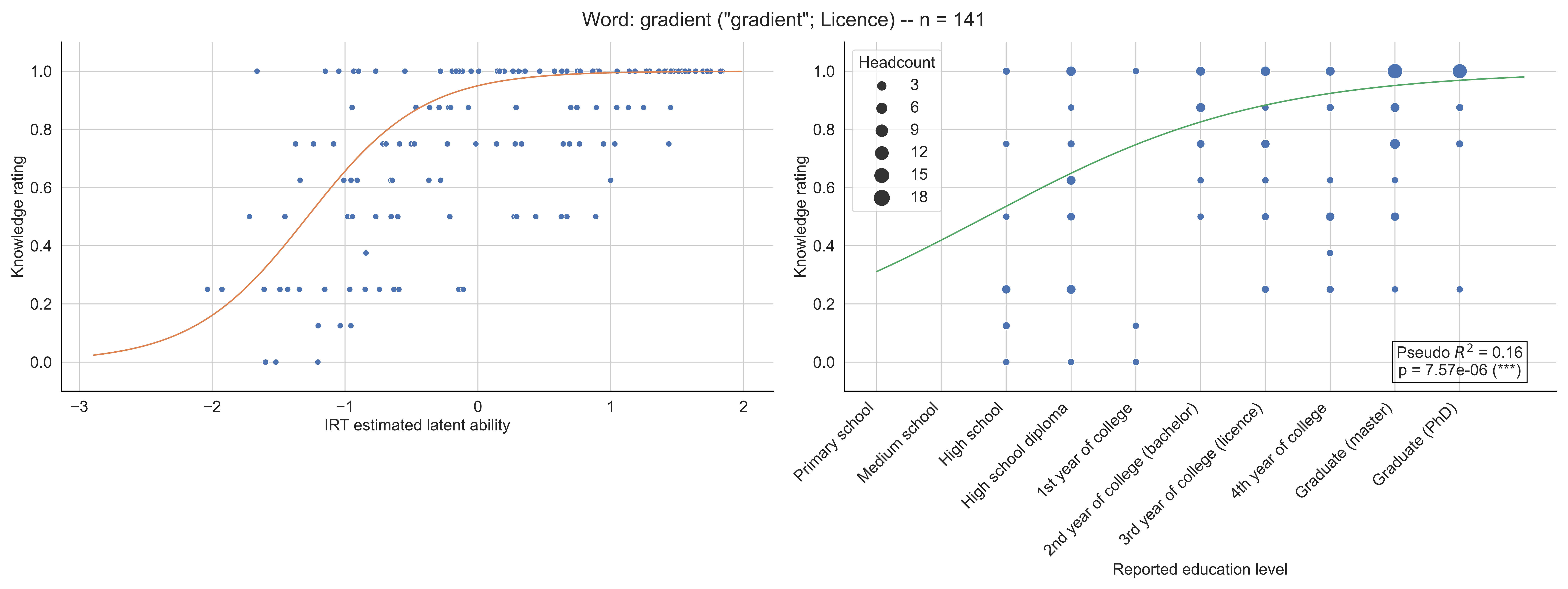

Word: gradient ("gradient"; Licence) -- n = 141
Knowledge rating
IRT estimated latent ability
Headcount
3
6
9
12
15
18
Pseudo $R^2$ = 0.16
p = 7.57e-06 (***)
Primary school
Medium school
High school
High school diploma
1st year of college
2nd year of college (bachelor)
3rd year of college (licence)
4th year of college
Graduate (master)
Graduate (PhD)
Reported education level

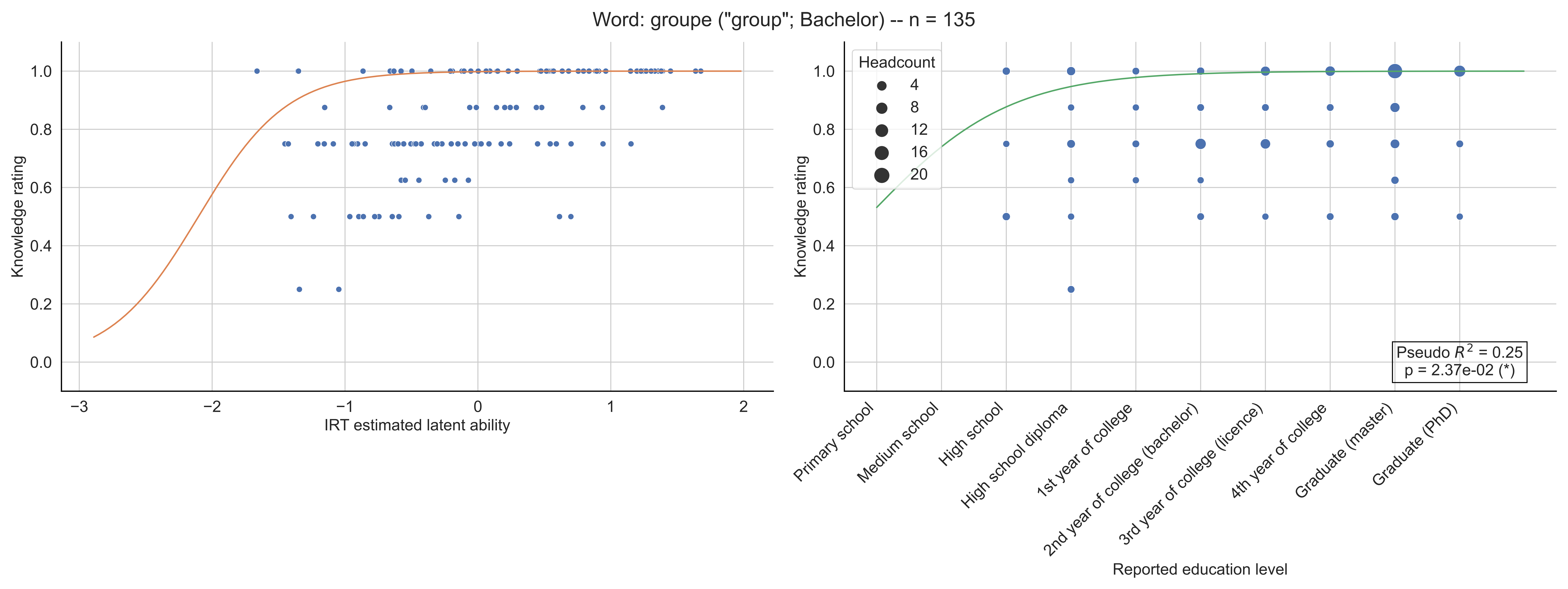

Word: groupe ("group"; Bachelor) -- n = 135
Knowledge rating
IRT estimated latent ability
Headcount
4
8
12
16
20
Pseudo $R^2$ = 0.25
p = 2.37e-02 (*)
Primary school
Medium school
High school
High school diploma
1st year of college
2nd year of college (bachelor)
3rd year of college (licence)
4th year of college
Graduate (master)
Graduate (PhD)
Reported education level

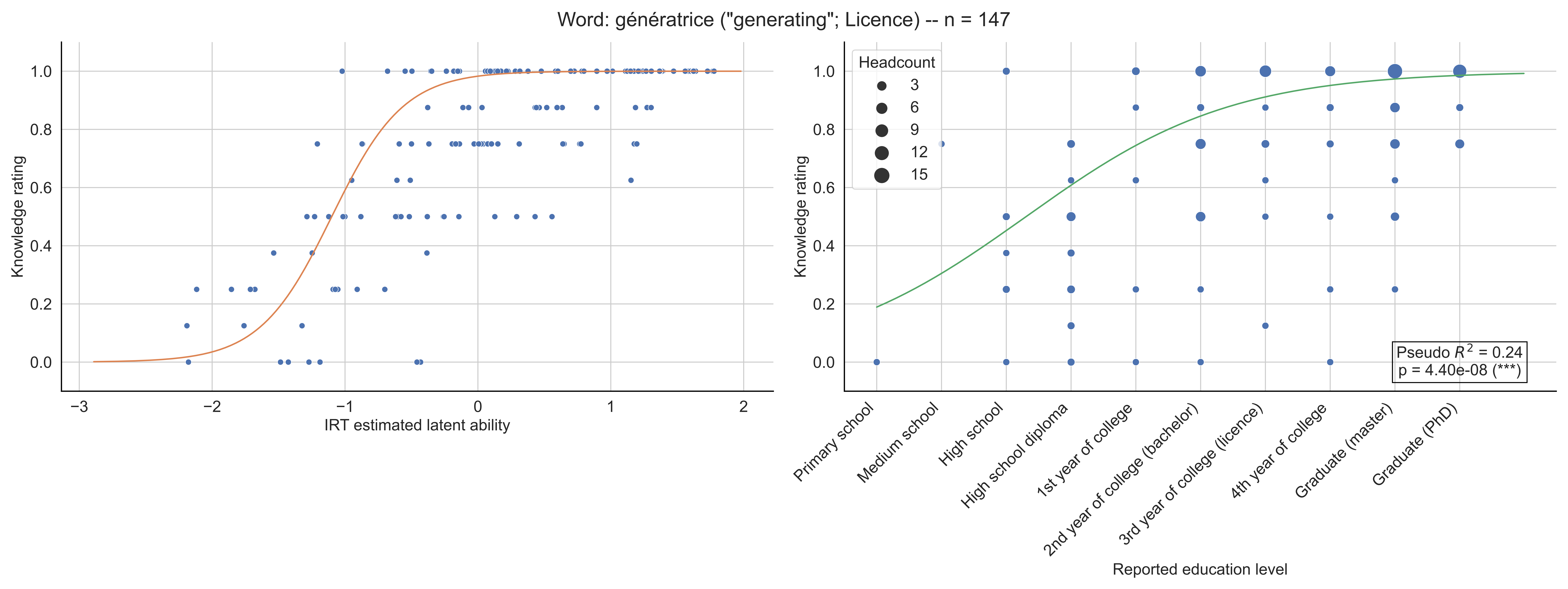

Word: génératrice ("generating"; Licence) -- n = 147
Knowledge rating
IRT estimated latent ability
Headcount
3
6
9
12
15
Pseudo $R^2$ = 0.24
p = 4.40e-08 (***)
Primary school
Medium school
High school
High school diploma
1st year of college
2nd year of college (bachelor)
3rd year of college (licence)
4th year of college
Graduate (master)
Graduate (PhD)
Reported education level

Word: hamiltonien ("hamiltonian"; Licence) -- n = 151

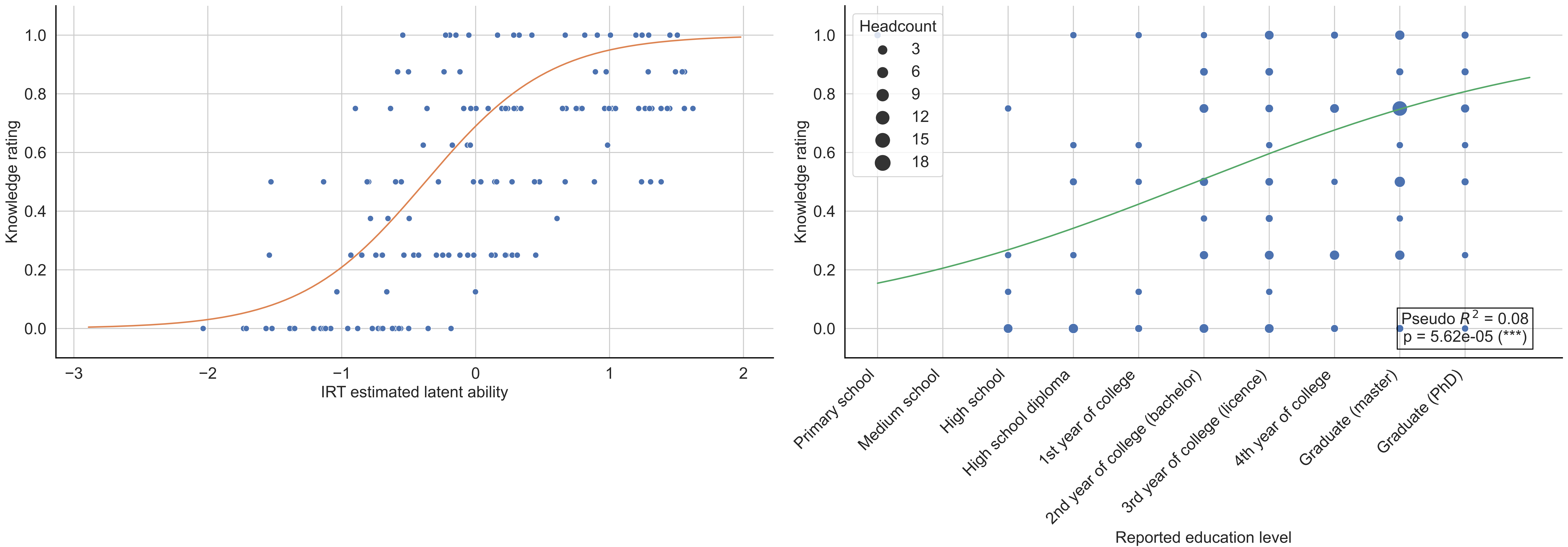

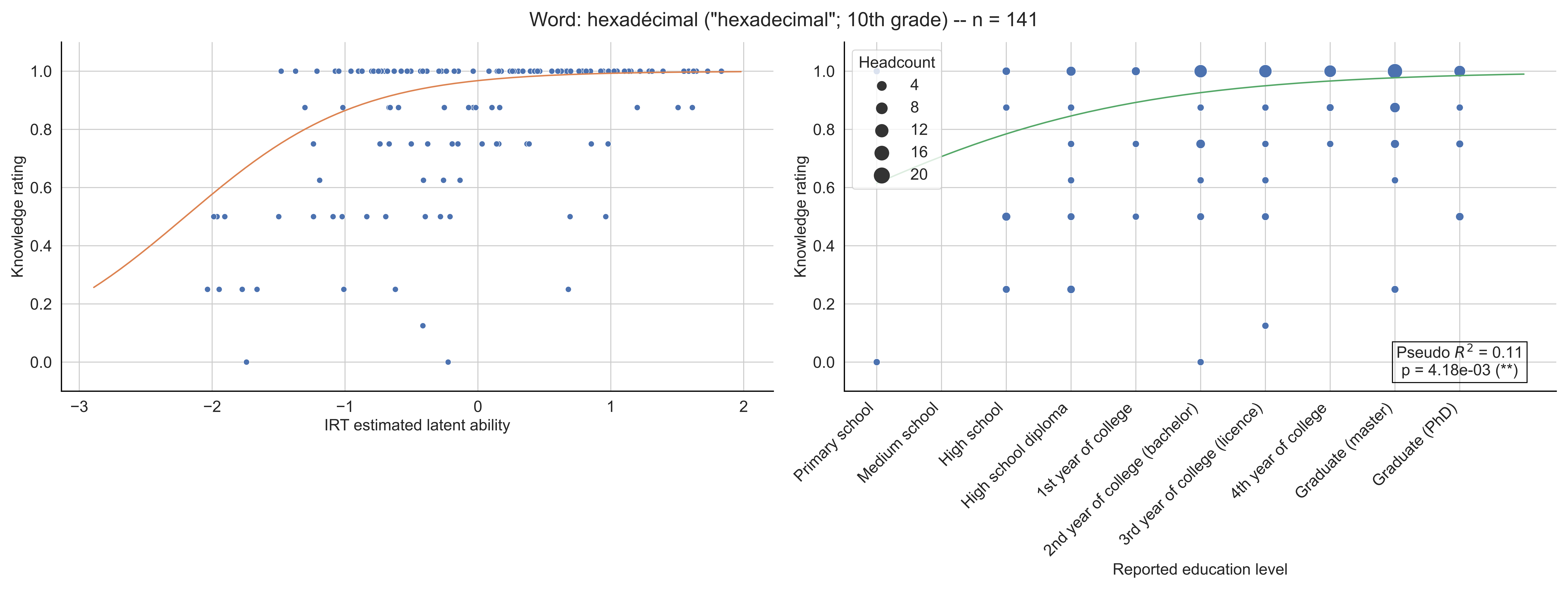

Word: hexadécimal ("hexadecimal"; 10th grade) -- n = 141
Knowledge rating
IRT estimated latent ability
Headcount
4
8
12
16
20
Pseudo $R^2$ = 0.11
p = 4.18e-03 (**)
Primary school
Medium school
High school
High school diploma
1st year of college
2nd year of college (bachelor)
3rd year of college (licence)
4th year of college
Graduate (master)
Graduate (PhD)
Reported education level

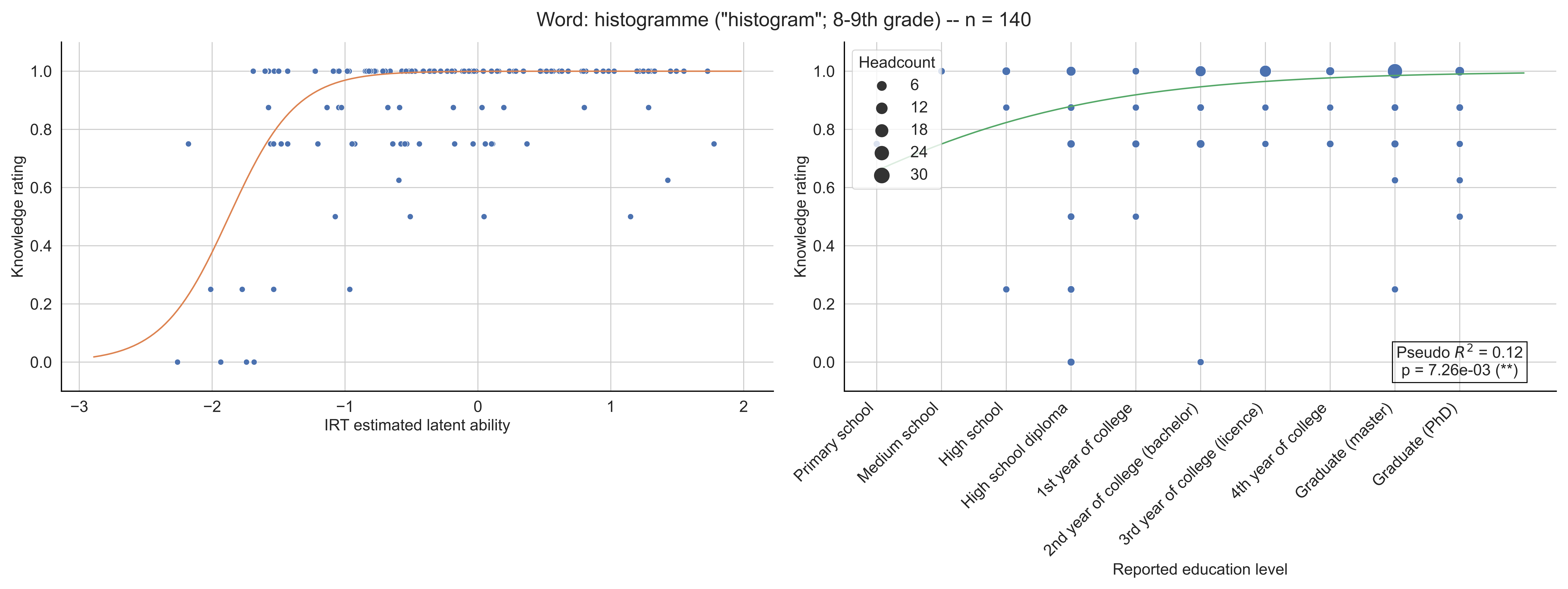

Word: histogramme ("histogram"; 8-9th grade) -- n = 140
Knowledge rating
IRT estimated latent ability
Headcount
6
12
18
24
30
Pseudo $R^2$ = 0.12
p = 7.26e-03 (**)
Primary school
Medium school
High school
High school diploma
1st year of college
2nd year of college (bachelor)
3rd year of college (licence)
4th year of college
Graduate (master)
Graduate (PhD)
Reported education level

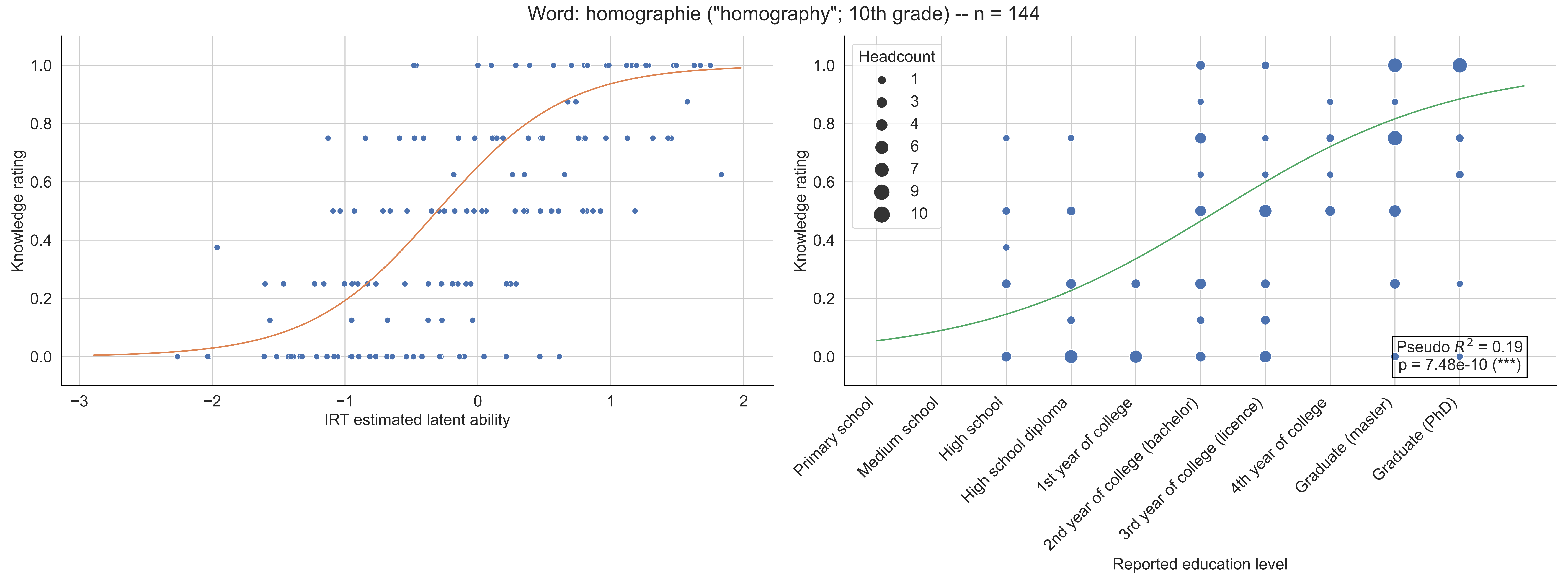

Word: homographie ("homography"; 10th grade) -- n = 144
Knowledge rating
IRT estimated latent ability
Headcount
1
3
4
6
7
9
10
Pseudo $R^2$ = 0.19
p = 7.48e-10 (***)
Primary school
Medium school
High school
High school diploma
1st year of college
2nd year of college (bachelor)
3rd year of college (licence)
4th year of college
Graduate (master)
Graduate (PhD)
Reported education level

Word: homologie ("homology"; Licence) -- n = 152

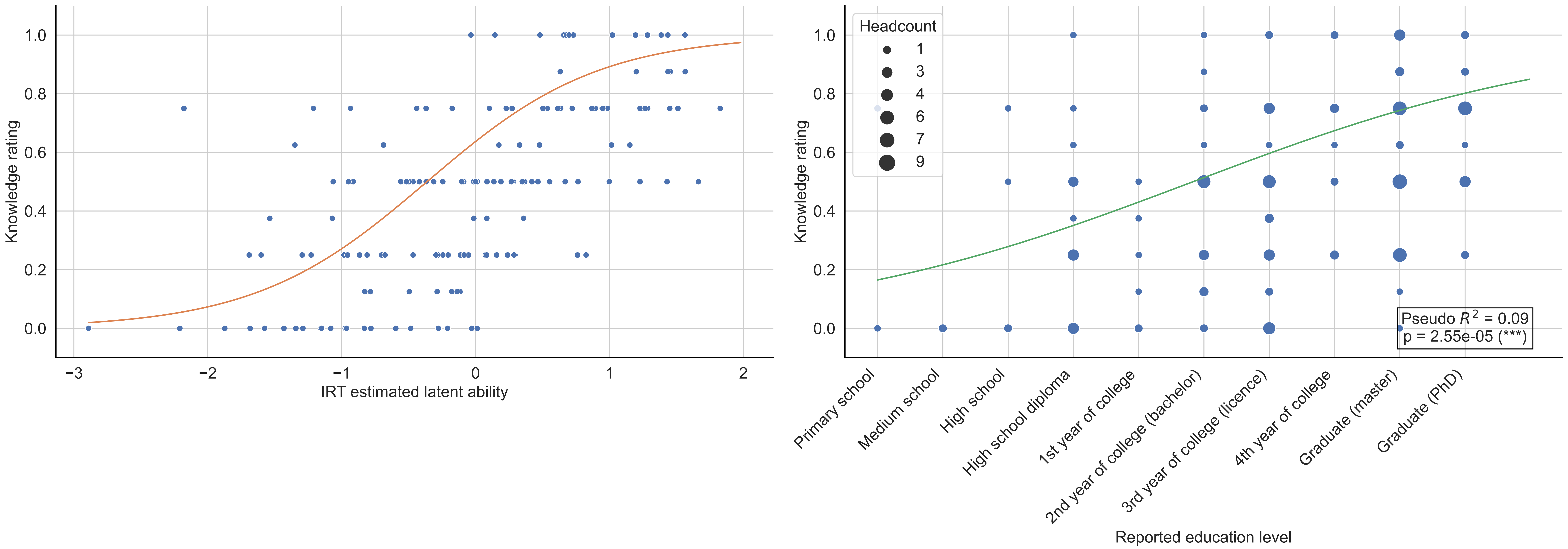

Word: homomorphisme ("homomorphism"; Bachelor) -- n = 131

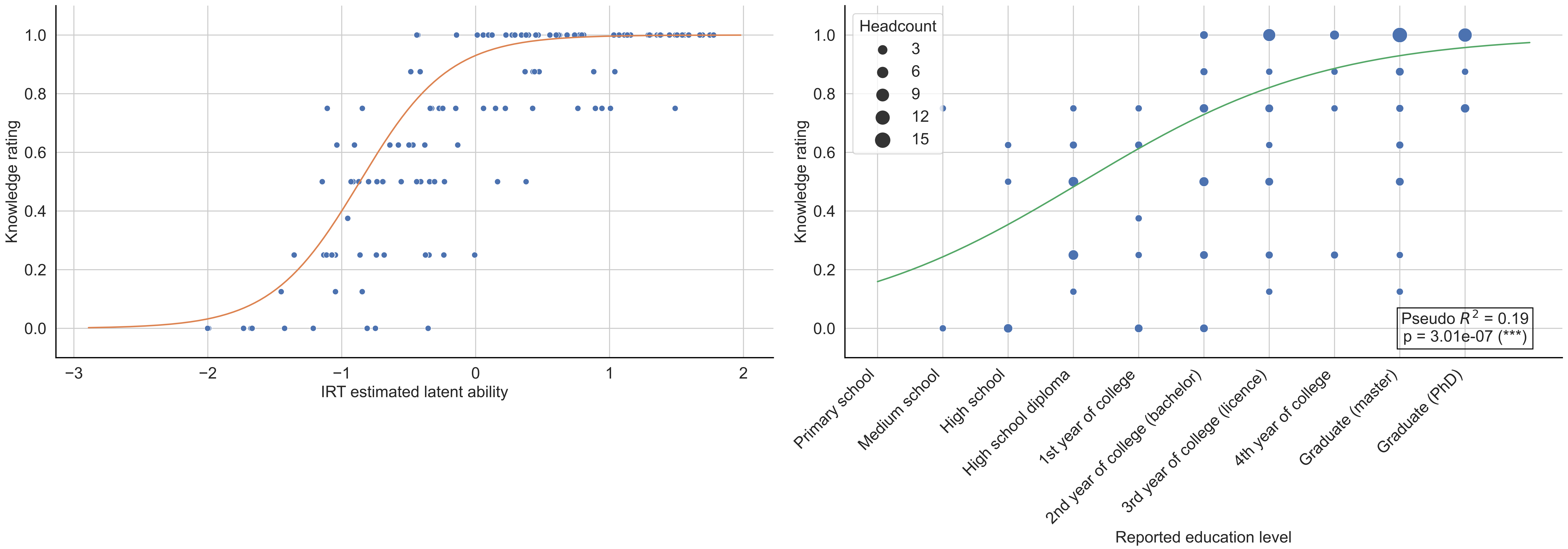

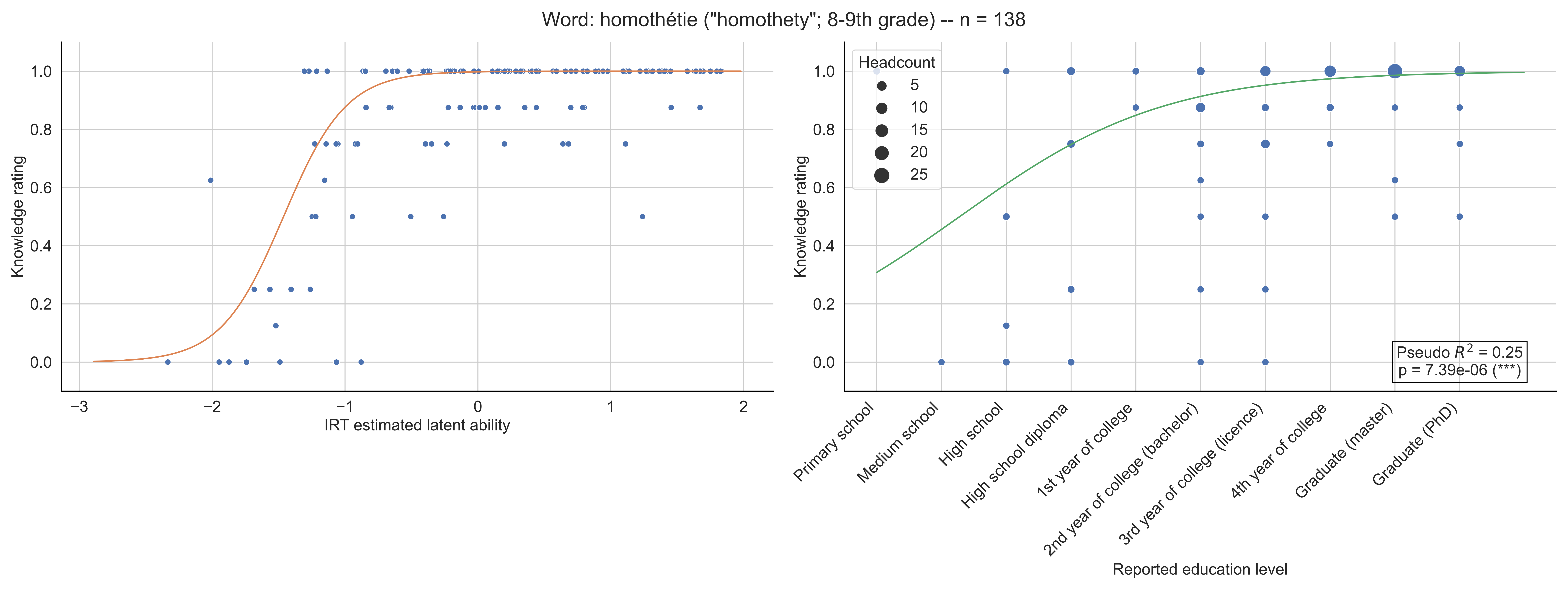
Word: homothétie ("homothety"; 8-9th grade) -- n = 138
Knowledge rating
IRT estimated latent ability
Headcount
5
10
15
20
25
Pseudo $R^2$ = 0.25
p = 7.39e-06 (***)
Primary school
Medium school
High school
High school diploma
1st year of college
2nd year of college (bachelor)
3rd year of college (licence)
4th year of college
Graduate (master)
Graduate (PhD)
Reported education level

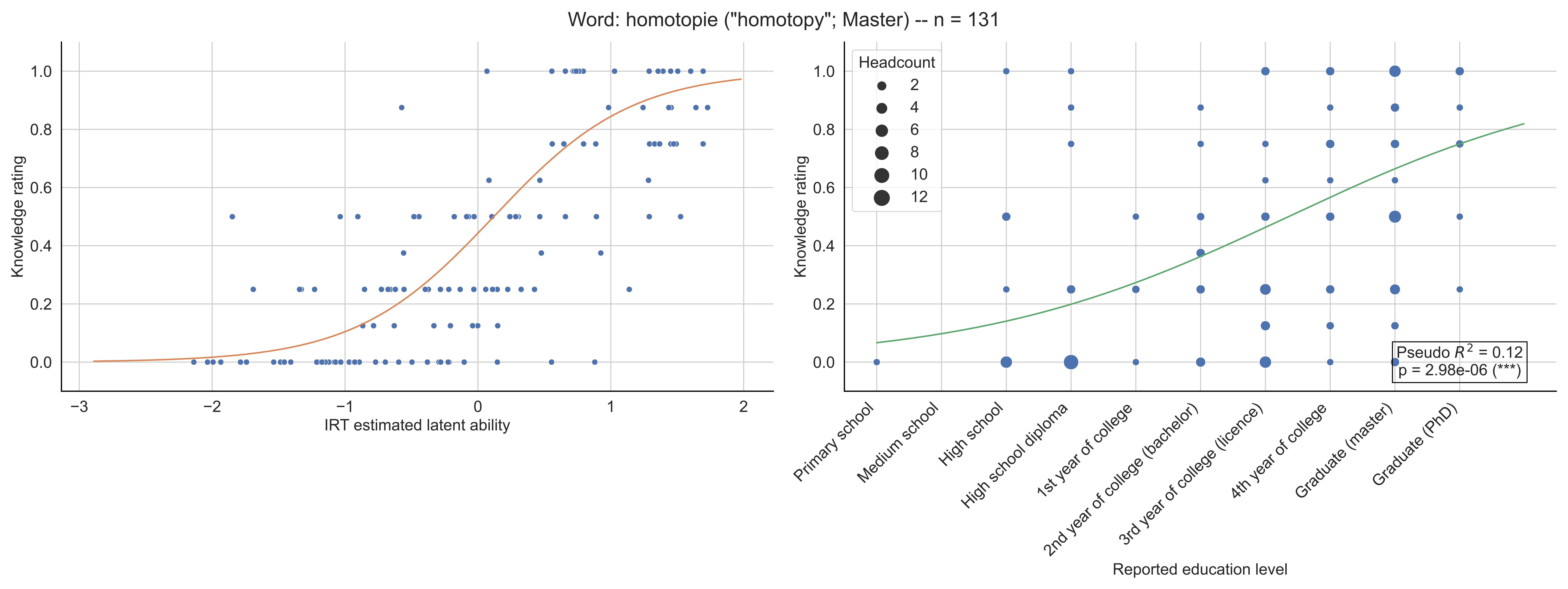

Word: homotopie ("homotopy"; Master) -- n = 131
Knowledge rating
IRT estimated latent ability
Headcount
2
4
6
8
10
12
Pseudo $R^2$ = 0.12
p = 2.98e-06 (***)
Primary school
Medium school
High school
High school diploma
1st year of college
2nd year of college (bachelor)
3rd year of college (licence)
4th year of college
Graduate (master)
Graduate (PhD)
Reported education level

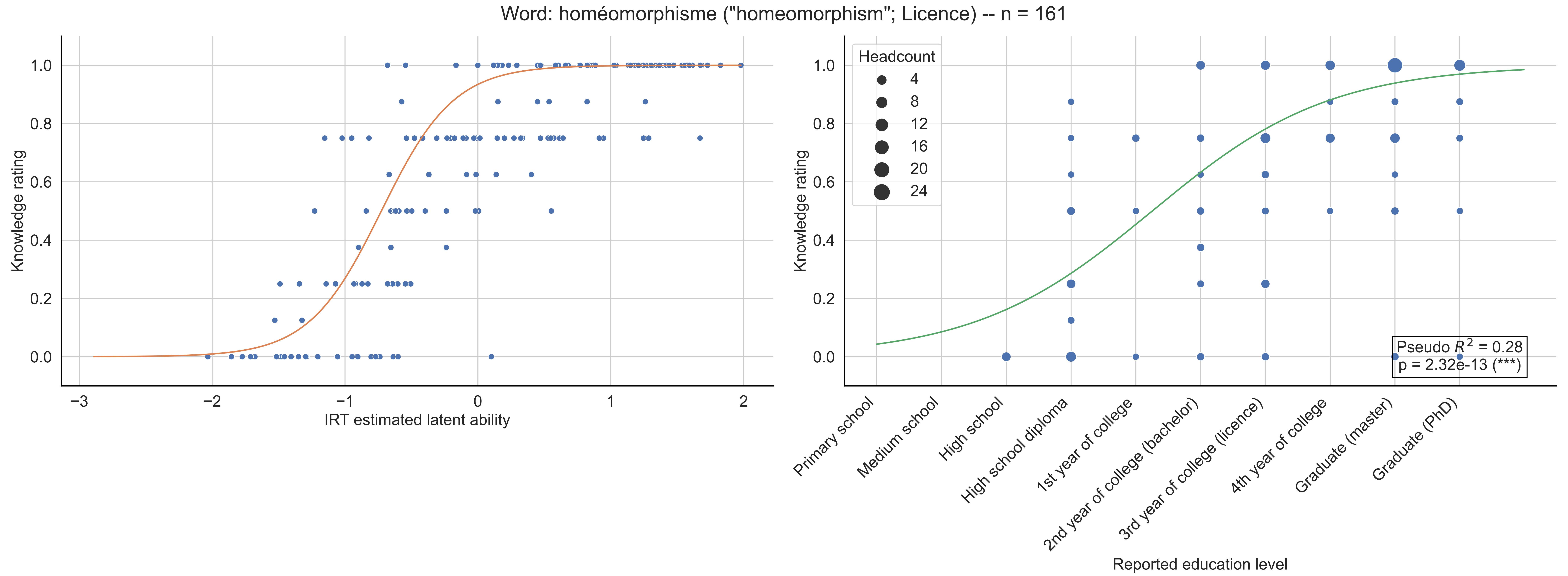
Word: homéomorphisme ("homeomorphism"; Licence) -- n = 161
Knowledge rating
IRT estimated latent ability
Headcount
4
8
12
16
20
24
Pseudo $R^2$ = 0.28
p = 2.32e-13 (***)
Primary school
Medium school
High school
High school diploma
1st year of college
2nd year of college (bachelor)
3rd year of college (licence)
4th year of college
Graduate (master)
Graduate (PhD)
Reported education level

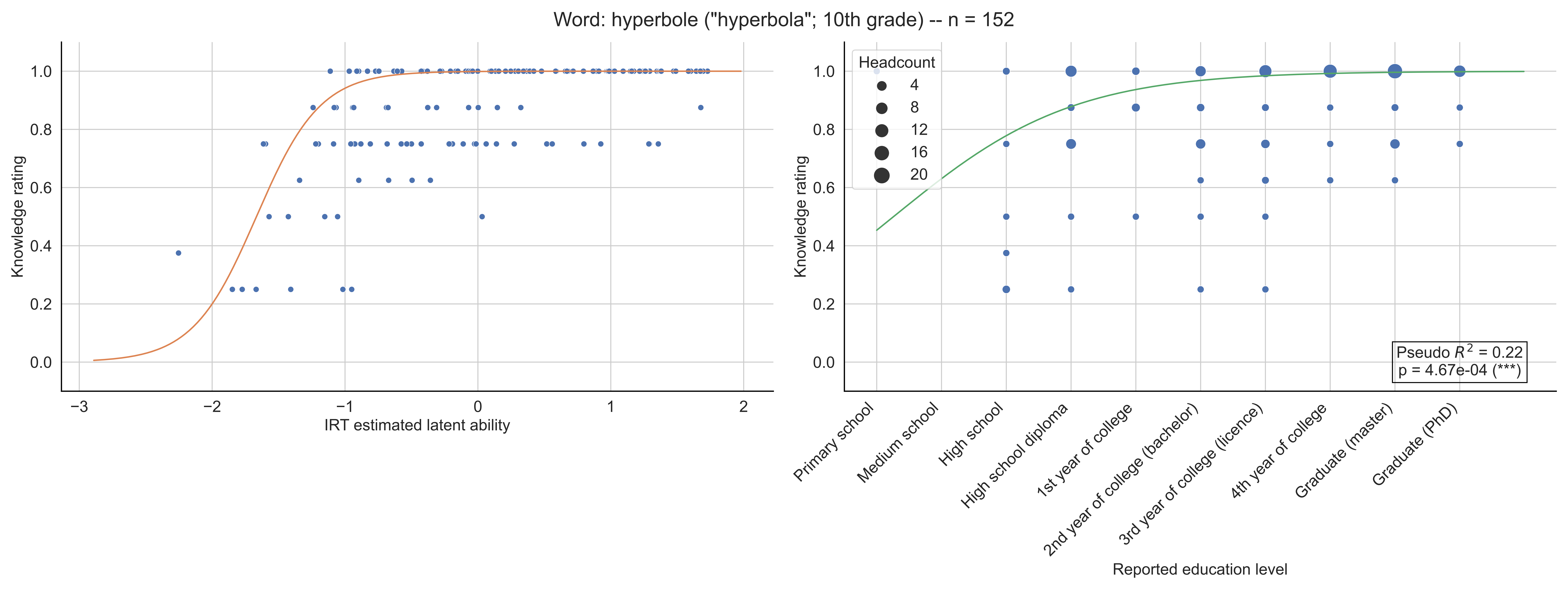

Word: hyperbole ("hyperbola"; 10th grade) -- n = 152
Knowledge rating
IRT estimated latent ability
Headcount
4
8
12
16
20
Pseudo $R^2$ = 0.22
p = 4.67e-04 (***)
Primary school
Medium school
High school
High school diploma
1st year of college
2nd year of college (bachelor)
3rd year of college (licence)
4th year of college
Graduate (master)
Graduate (PhD)
Reported education level

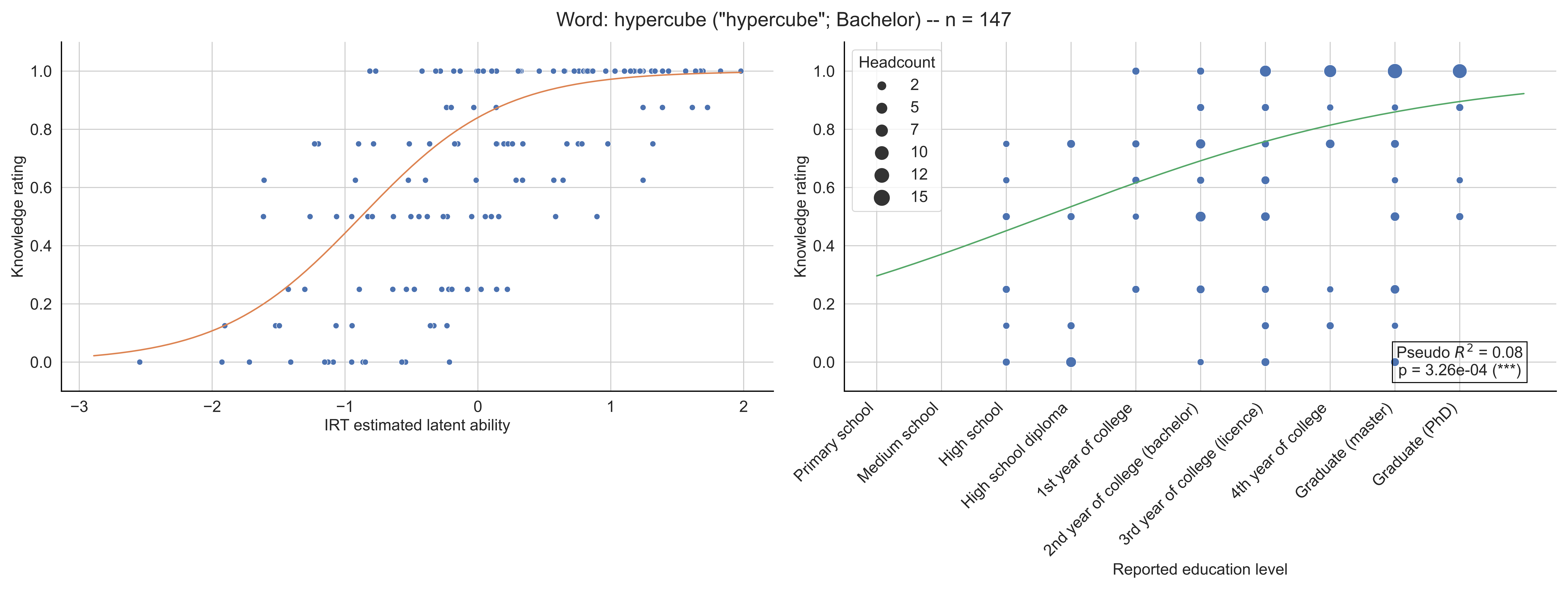

Word: hypercube ("hypercube"; Bachelor) -- n = 147
Knowledge rating
IRT estimated latent ability
Headcount
2
5
7
10
12
15
Pseudo $R^2$ = 0.08
p = 3.26e-04 (***)
Primary school
Medium school
High school
High school diploma
1st year of college
2nd year of college (bachelor)
3rd year of college (licence)
4th year of college
Graduate (master)
Graduate (PhD)
Reported education level

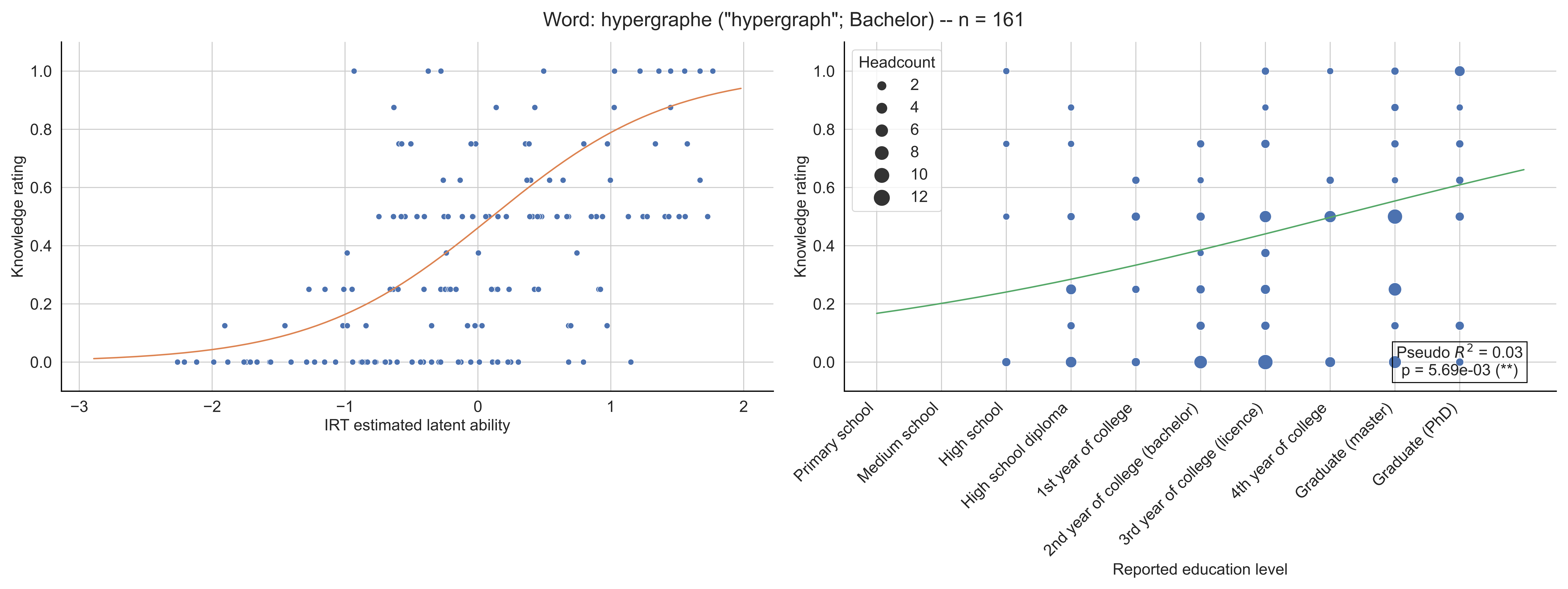
Word: hypergraphe ("hypergraph"; Bachelor) -- n = 161
Knowledge rating
IRT estimated latent ability
Headcount
2
4
6
8
10
12
Pseudo $R^2$ = 0.03
p = 5.69e-03 (**)
Reported education level
Primary school
Medium school
High school
High school diploma
1st year of college
2nd year of college (bachelor)
3rd year of college (licence)
4th year of college
Graduate (master)
Graduate (PhD)

Word: hypersurface ("hypersurface"; Bachelor) -- n = 151

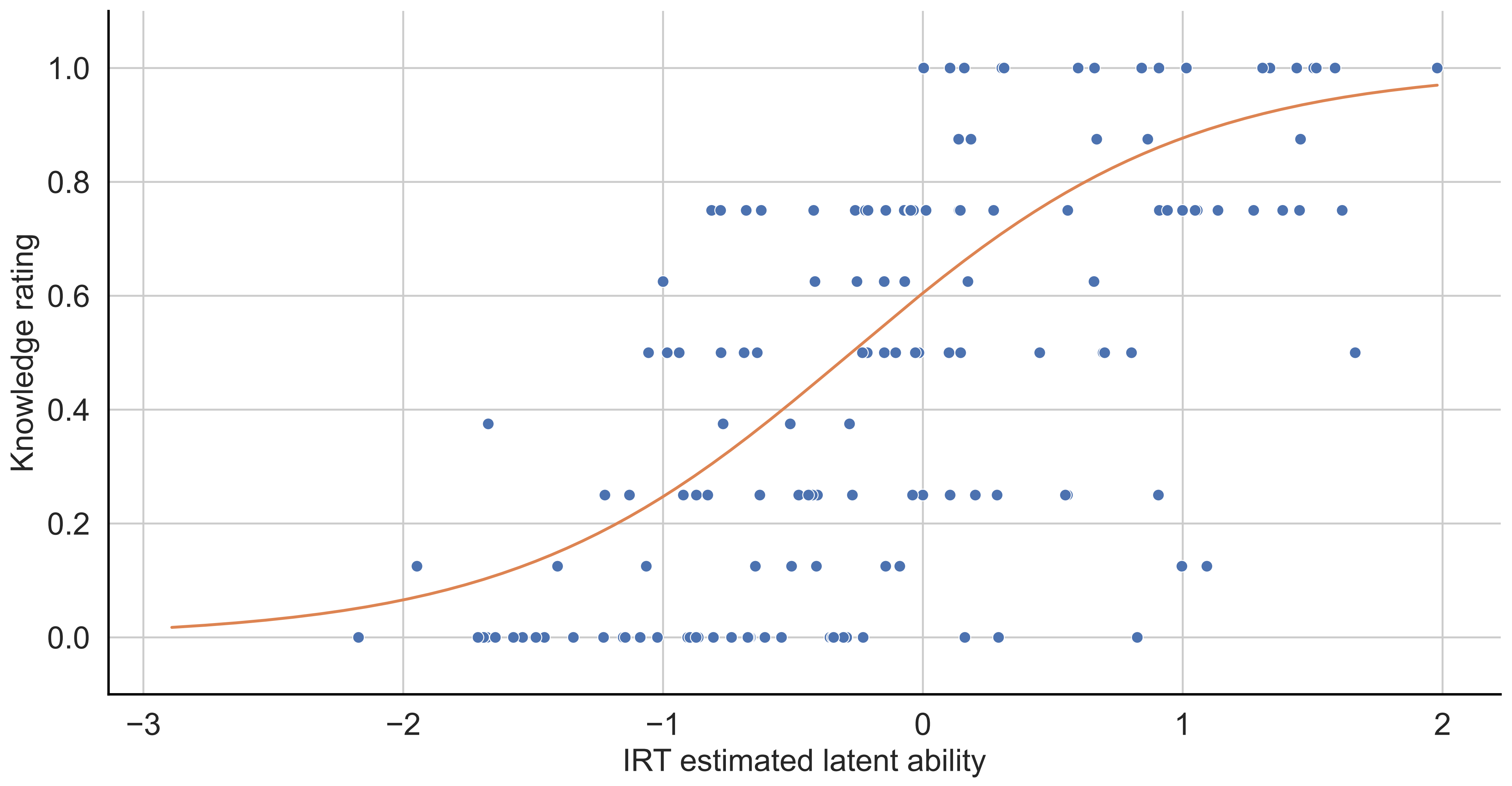


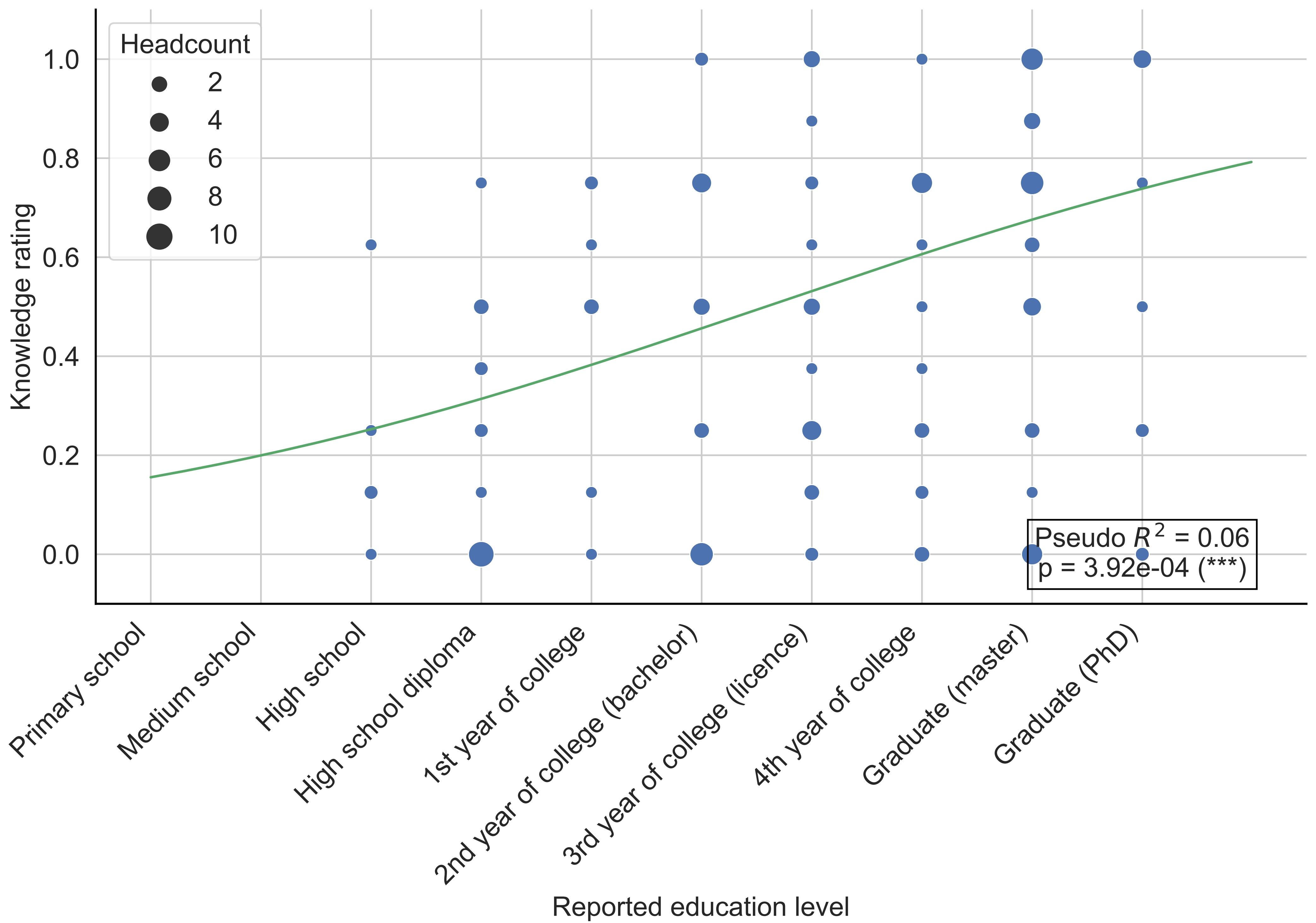

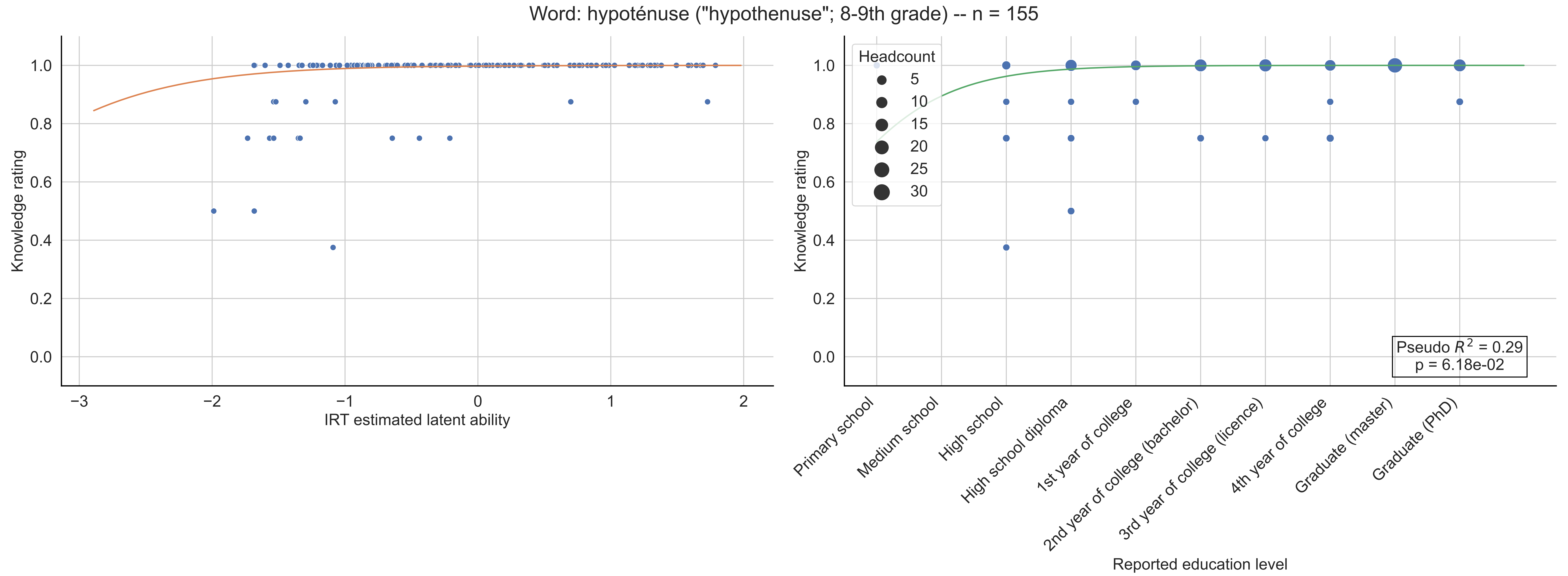

Word: hypoténuse ("hypothenuse"; 8-9th grade) -- n = 155
Knowledge rating
IRT estimated latent ability
Headcount
5
10
15
20
25
30
Pseudo $R^2$ = 0.29
p = 6.18e-02
Primary school
Medium school
High school
High school diploma
1st year of college
2nd year of college (bachelor)
3rd year of college (licence)
4th year of college
Graduate (master)
Graduate (PhD)
Reported education level

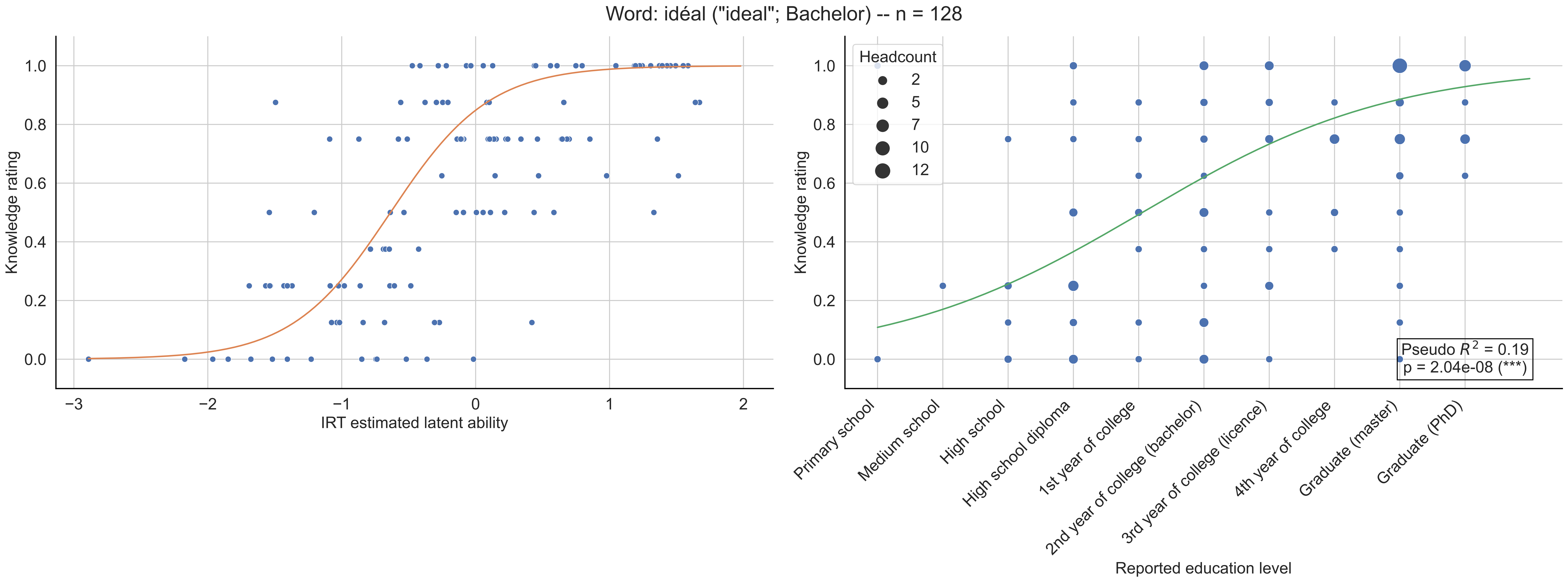

Word: idéal ("ideal"; Bachelor) -- n = 128
Knowledge rating
IRT estimated latent ability
Headcount
2
5
7
10
12
Pseudo $R^2$ = 0.19
p = 2.04e-08 (***)
Primary school
Medium school
High school
High school diploma
1st year of college
2nd year of college (bachelor)
3rd year of college (licence)
4th year of college
Graduate (master)
Graduate (PhD)
Reported education level

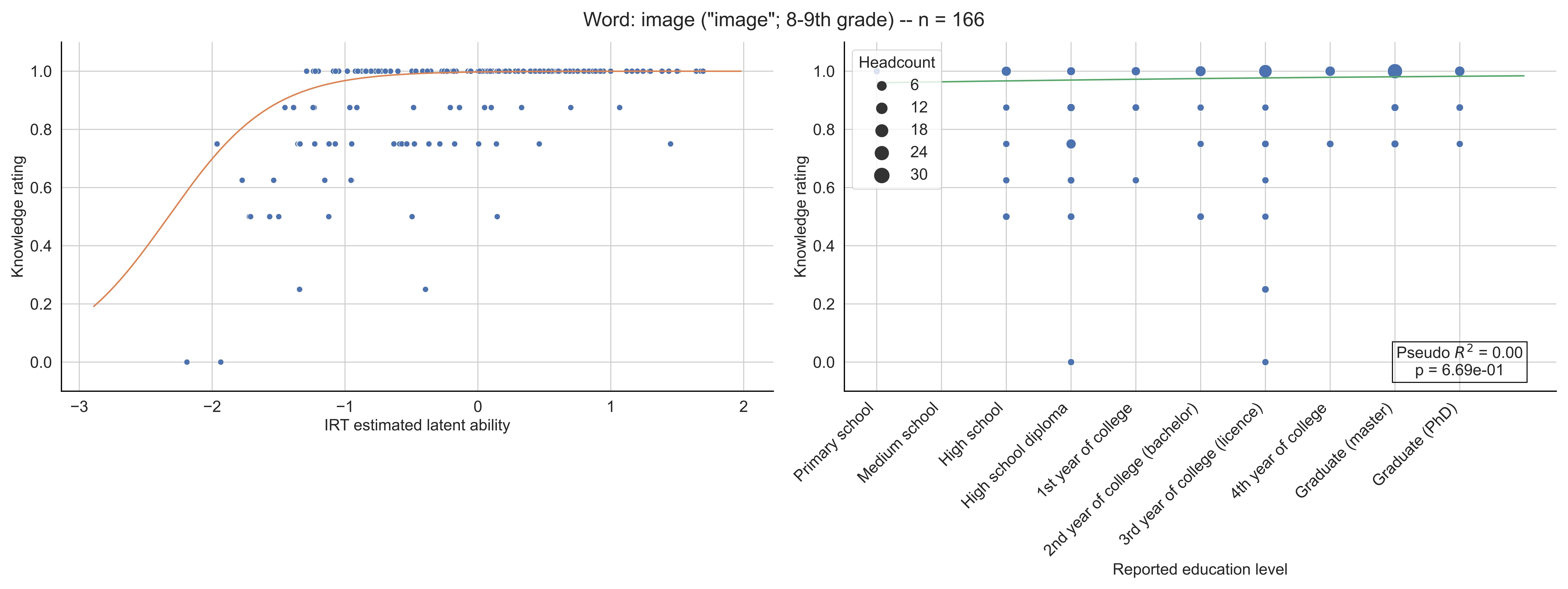
Word: image ("image"; 8-9th grade) -- n = 166
Knowledge rating
IRT estimated latent ability
Headcount
6
12
18
24
30
Pseudo $R^2$ = 0.00
p = 6.69e-01
Primary school
Medium school
High school
High school diploma
1st year of college
2nd year of college (bachelor)
3rd year of college (licence)
4th year of college
Graduate (master)
Graduate (PhD)
Reported education level

Word: incomplétude ("incompleteness"; Licence) -- n = 131

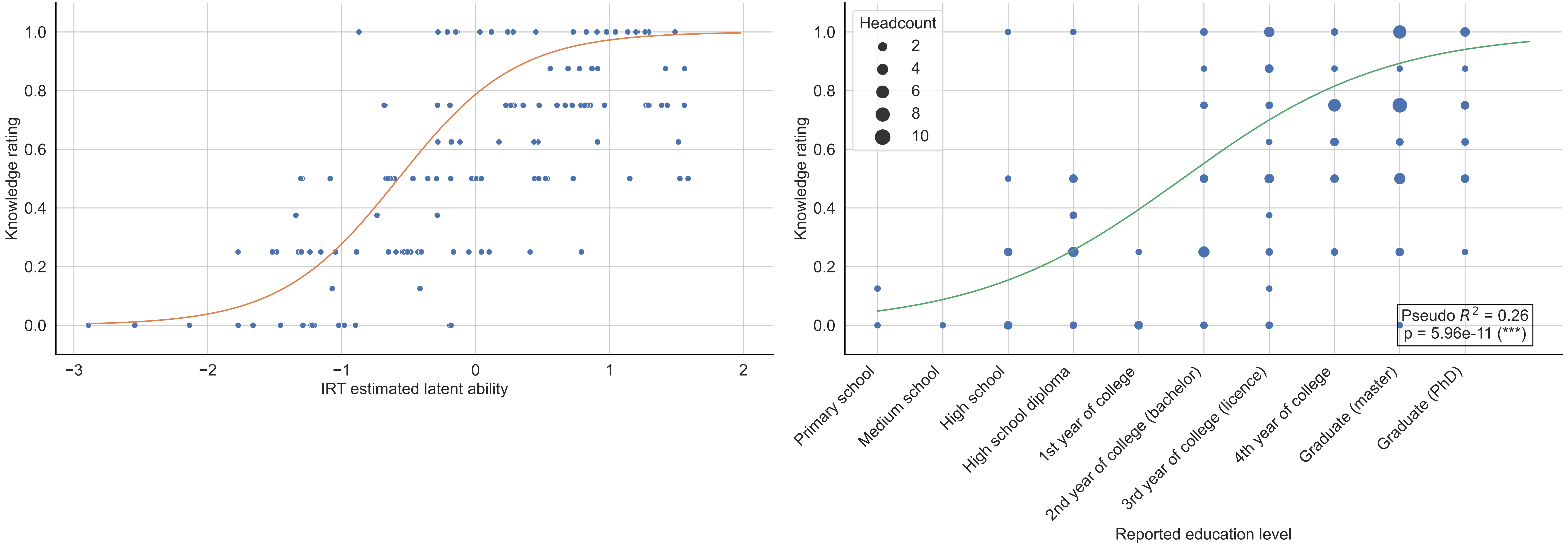

Word: indice ("index"; 11-12th grade) -- n = 133

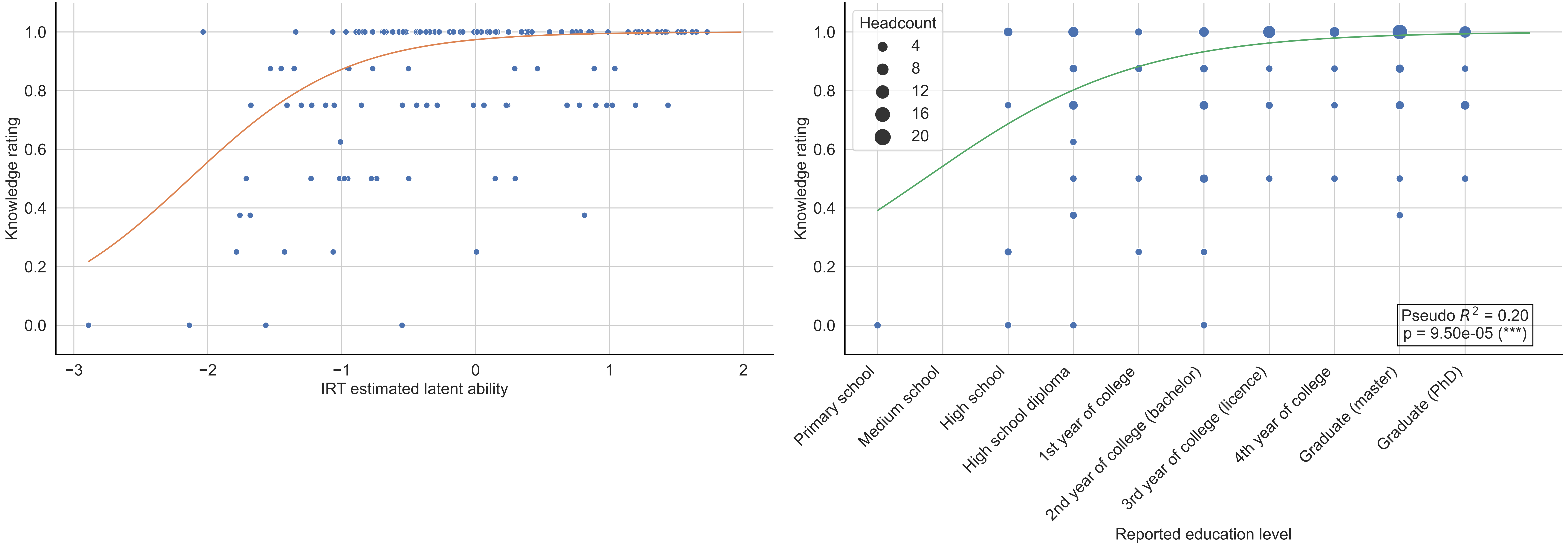

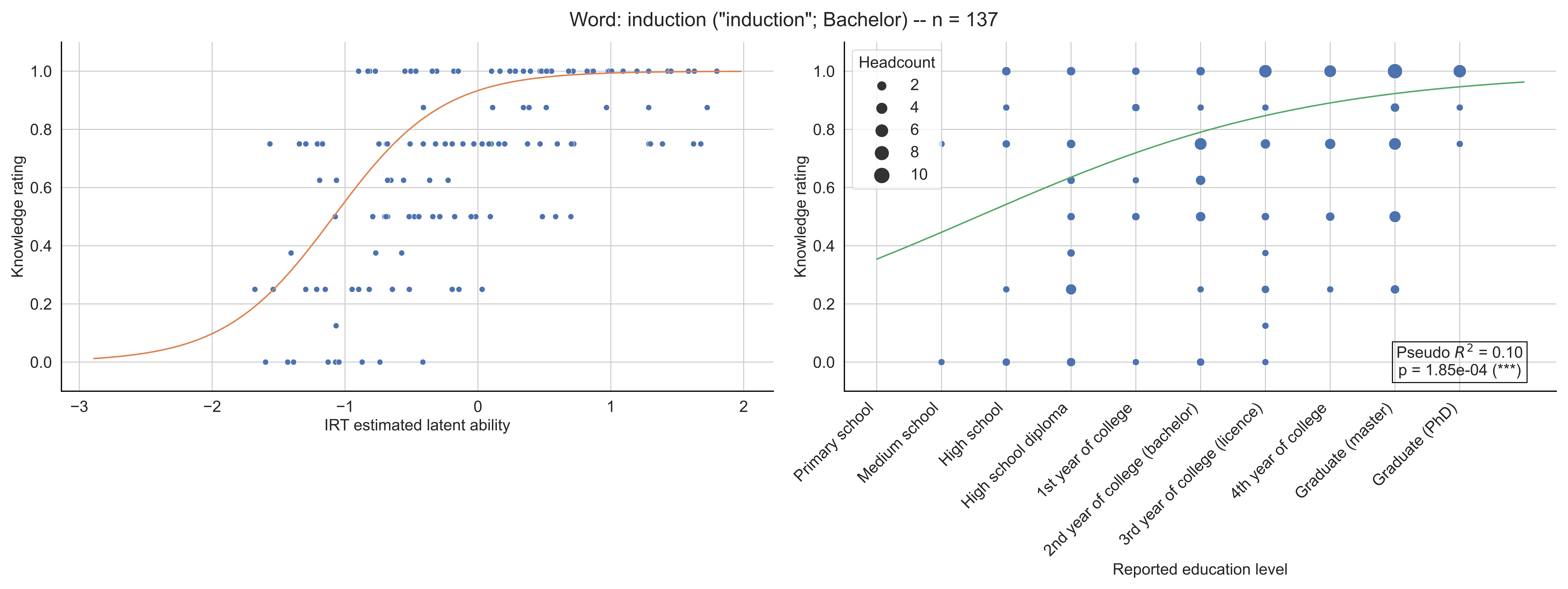
Word: induction ("induction"; Bachelor) -- n = 137
Knowledge rating
IRT estimated latent ability
Headcount
2
4
6
8
10
Pseudo $R^2$ = 0.10
p = 1.85e-04 (***)
Primary school
Medium school
High school
High school diploma
1st year of college
2nd year of college (bachelor)
3rd year of college (licence)
4th year of college
Graduate (master)
Graduate (PhD)
Reported education level

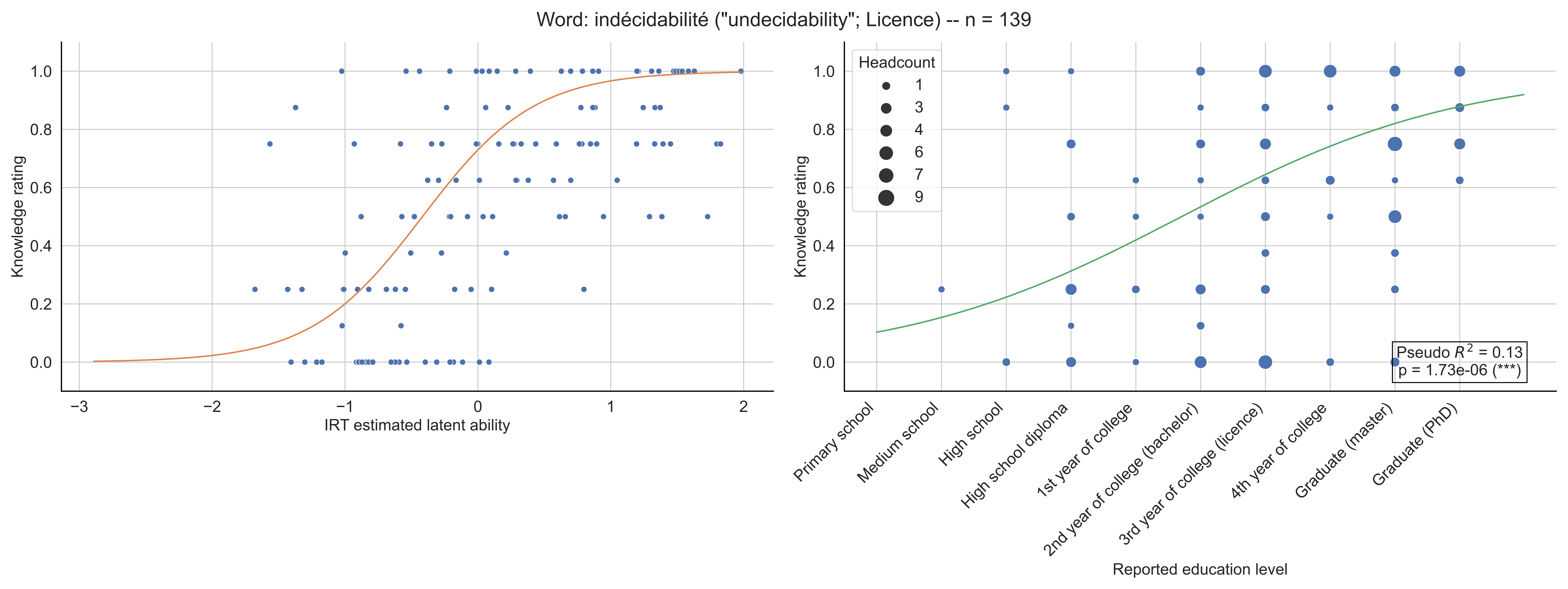

Word: indécidabilité ("undecidability"; Licence) -- n = 139
Knowledge rating
IRT estimated latent ability
Headcount
1
3
4
6
7
9
Pseudo $R^2$ = 0.13
p = 1.73e-06 (***)
Primary school
Medium school
High school
High school diploma
1st year of college
2nd year of college (bachelor)
3rd year of college (licence)
4th year of college
Graduate (master)
Graduate (PhD)
Reported education level

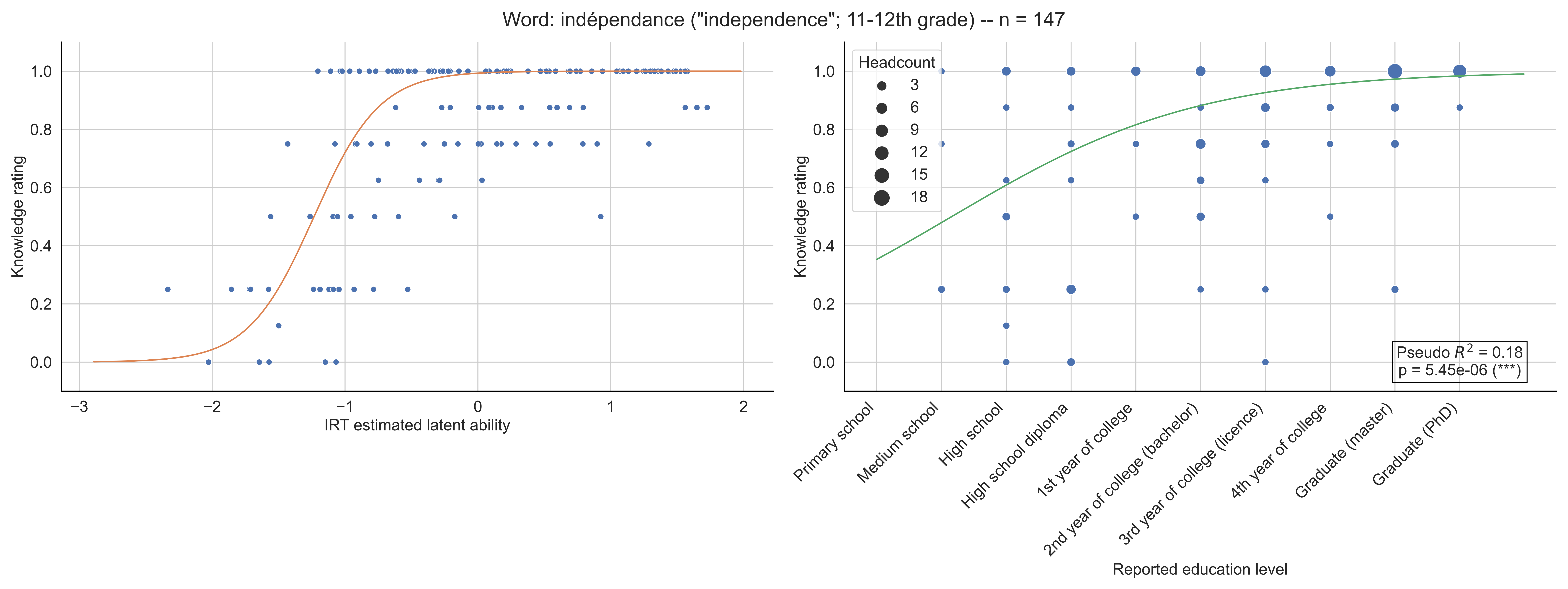

Word: indépendance ("independence"; 11-12th grade) -- n = 147
Knowledge rating
IRT estimated latent ability
Headcount
3
6
9
12
15
18
Pseudo $R^2$ = 0.18
p = 5.45e-06 (***)
Primary school
Medium school
High school
High school diploma
1st year of college
2nd year of college (bachelor)
3rd year of college (licence)
4th year of college
Graduate (master)
Graduate (PhD)
Reported education level

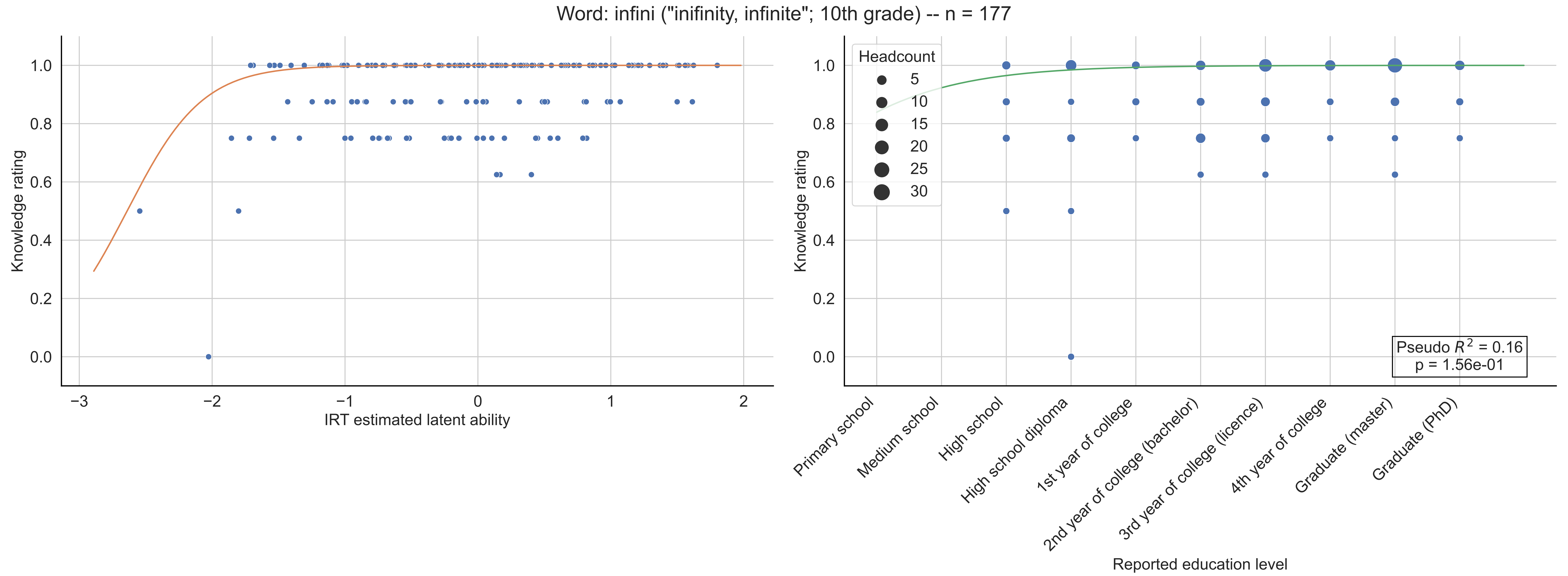

Word: infini ("inifinity, infinite"; 10th grade) -- n = 177
Knowledge rating
IRT estimated latent ability
Headcount
5
10
15
20
25
30
Pseudo $R^2$ = 0.16
p = 1.56e-01
Primary school
Medium school
High school
High school diploma
1st year of college
2nd year of college (bachelor)
3rd year of college (licence)
4th year of college
Graduate (master)
Graduate (PhD)
Reported education level

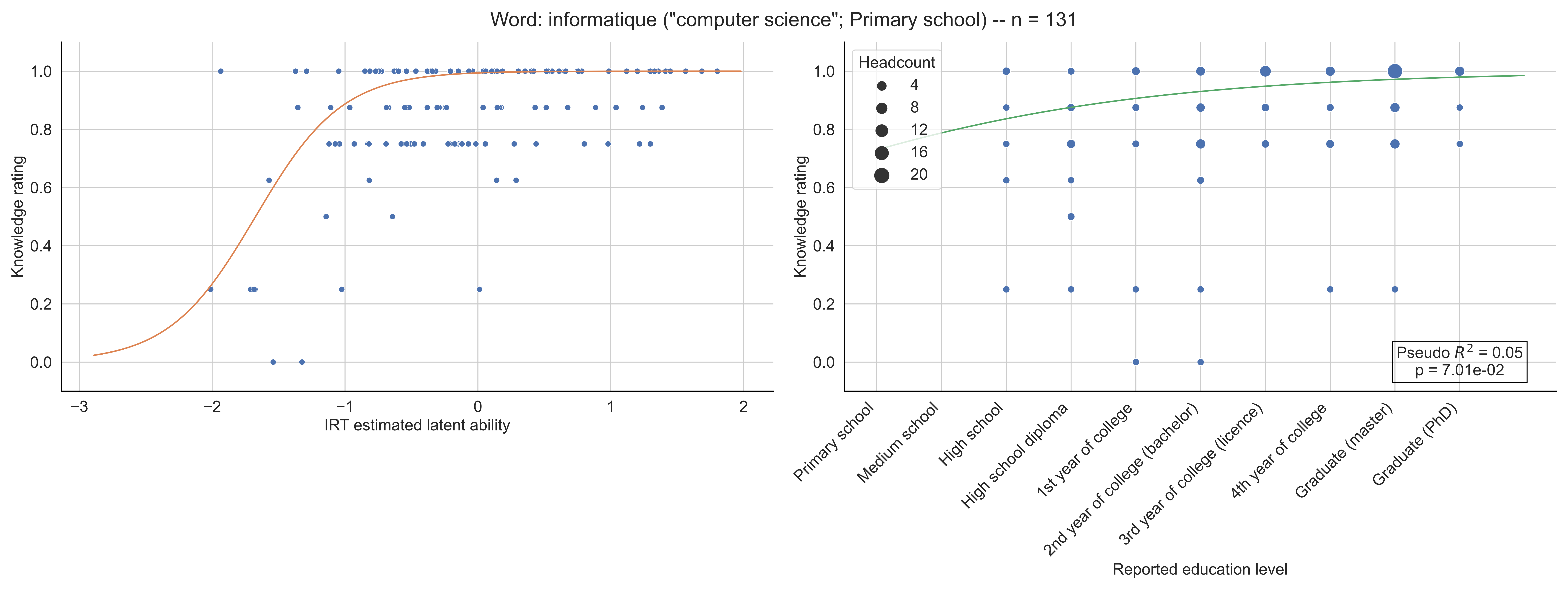
Word: informatique ("computer science"; Primary school) -- n = 131
Knowledge rating
IRT estimated latent ability
Headcount
4
8
12
16
20
Pseudo $R^2$ = 0.05
p = 7.01e-02
Primary school
Medium school
High school
High school diploma
1st year of college
2nd year of college (bachelor)
3rd year of college (licence)
4th year of college
Graduate (master)
Graduate (PhD)
Reported education level

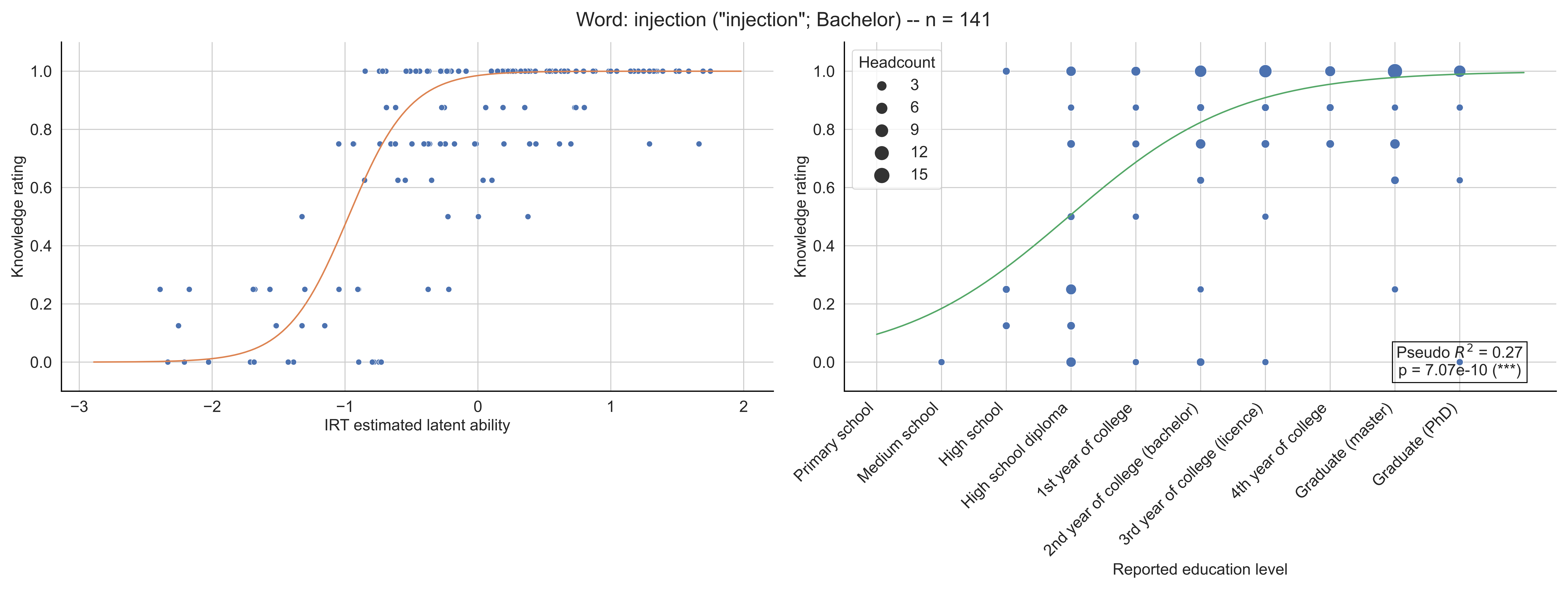

Word: injection ("injection"; Bachelor) -- n = 141
Knowledge rating
IRT estimated latent ability
Headcount
3
6
9
12
15
Primary school
Medium school
High school
High school diploma
1st year of college
2nd year of college (bachelor)
3rd year of college (licence)
4th year of college
Graduate (master)
Graduate (PhD)
Reported education level
Pseudo $R^2$ = 0.27
p = 7.07e-10 (***)

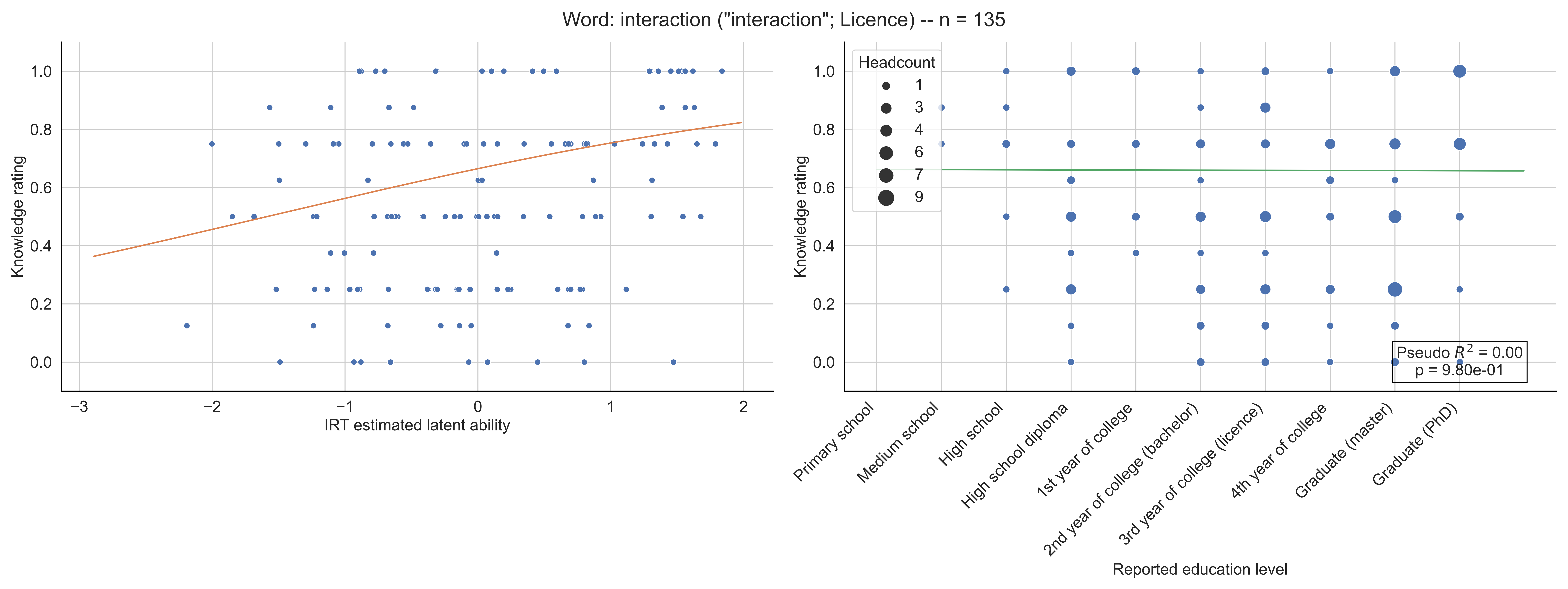

Word: interaction ("interaction"; Licence) -- n = 135
Knowledge rating
IRT estimated latent ability
Headcount
1
3
4
6
7
9
Pseudo $R^2$ = 0.00
p = 9.80e-01
Primary school
Medium school
High school
High school diploma
1st year of college
2nd year of college (bachelor)
3rd year of college (licence)
4th year of college
Graduate (master)
Graduate (PhD)
Reported education level

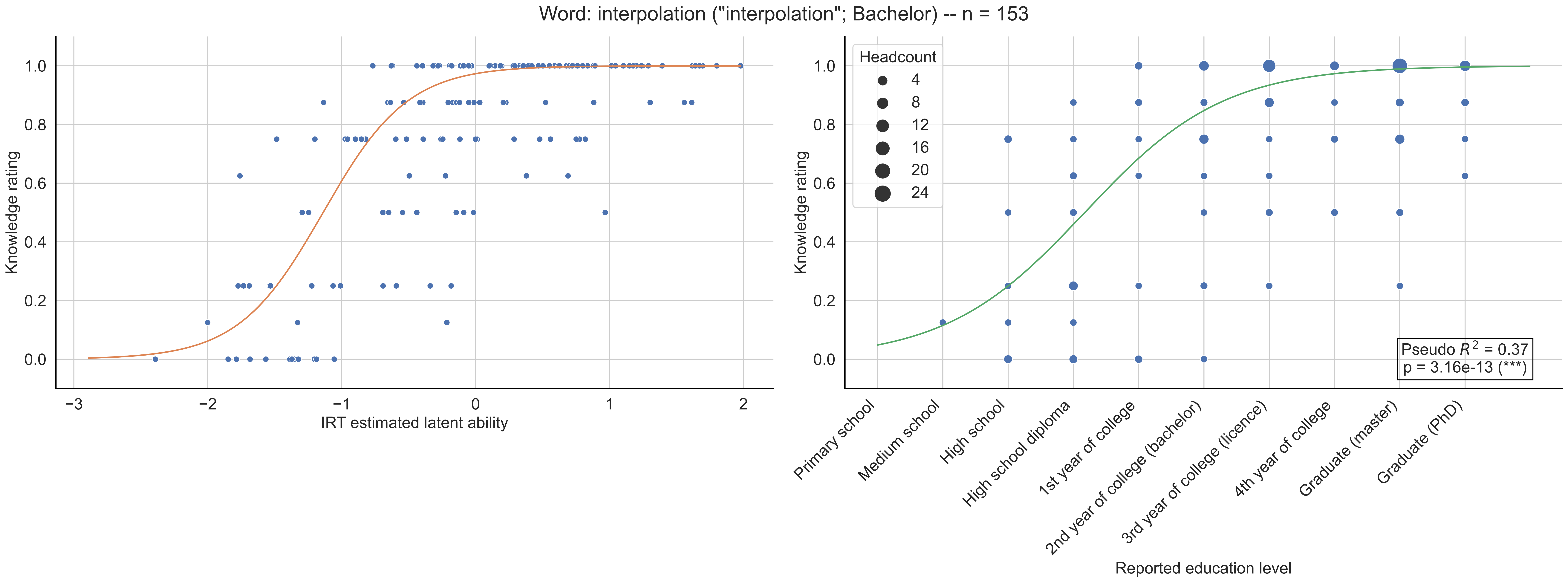
Word: interpolation ("interpolation"; Bachelor) -- n = 153
Knowledge rating
IRT estimated latent ability
Headcount
4
8
12
16
20
24
Pseudo $R^2$ = 0.37
p = 3.16e-13 (***)
Primary school
Medium school
High school
High school diploma
1st year of college
2nd year of college (bachelor)
3rd year of college (licence)
4th year of college
Graduate (master)
Graduate (PhD)
Reported education level

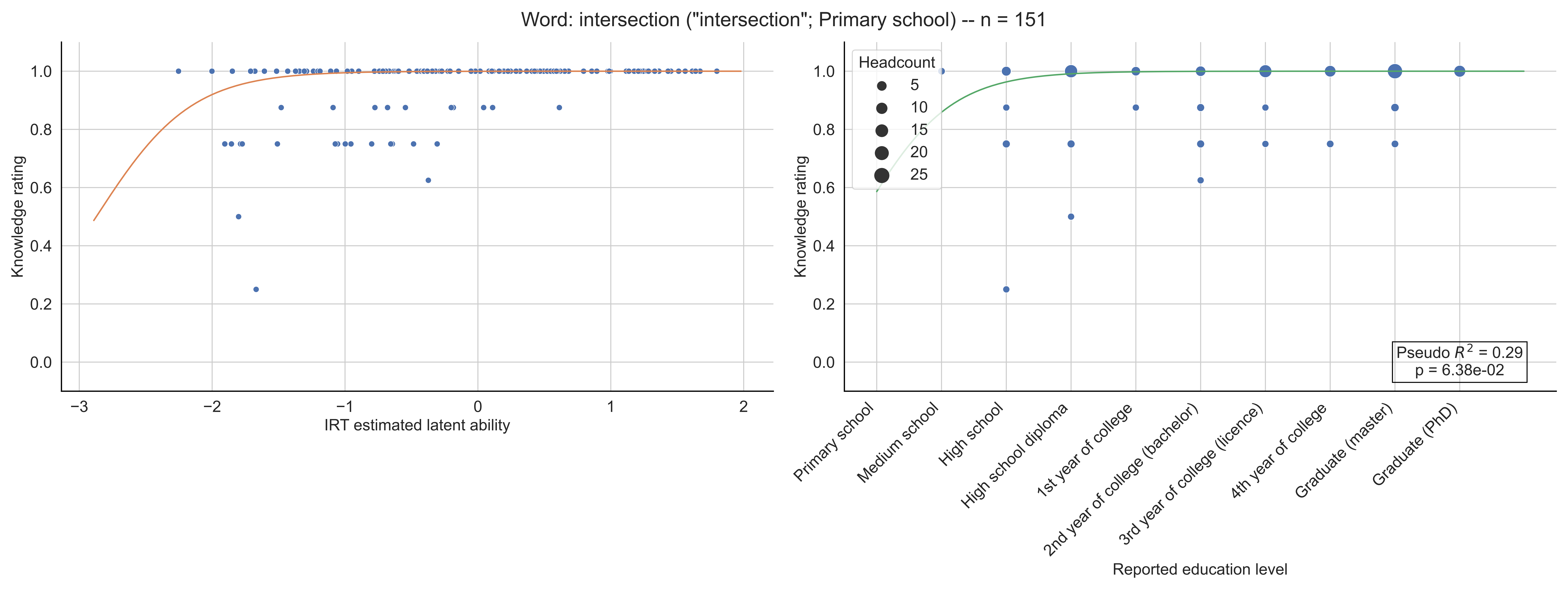
Word: intersection ("intersection"; Primary school) -- n = 151
Knowledge rating
IRT estimated latent ability
Headcount
5
10
15
20
25
Pseudo $R^2$ = 0.29
p = 6.38e-02
Primary school
Medium school
High school
High school diploma
1st year of college
2nd year of college (bachelor)
3rd year of college (licence)
4th year of college
Graduate (master)
Graduate (PhD)
Reported education level

Word: intégrale ("integral"; 11-12th grade) -- n = 152

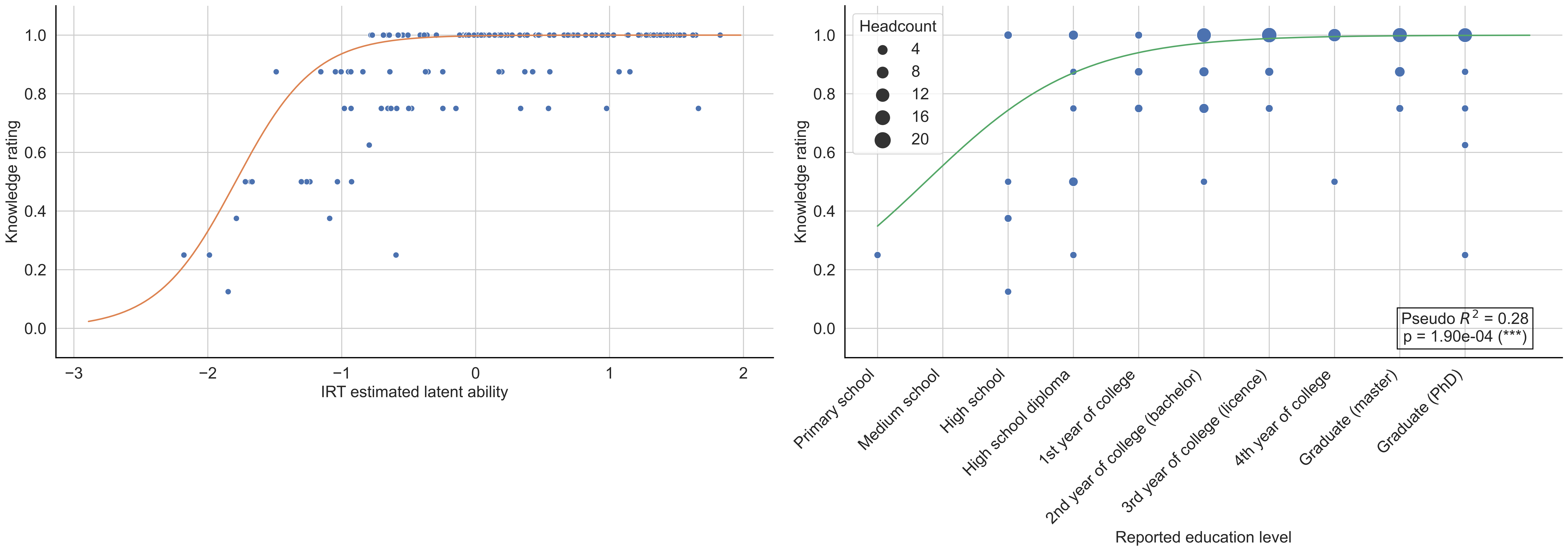

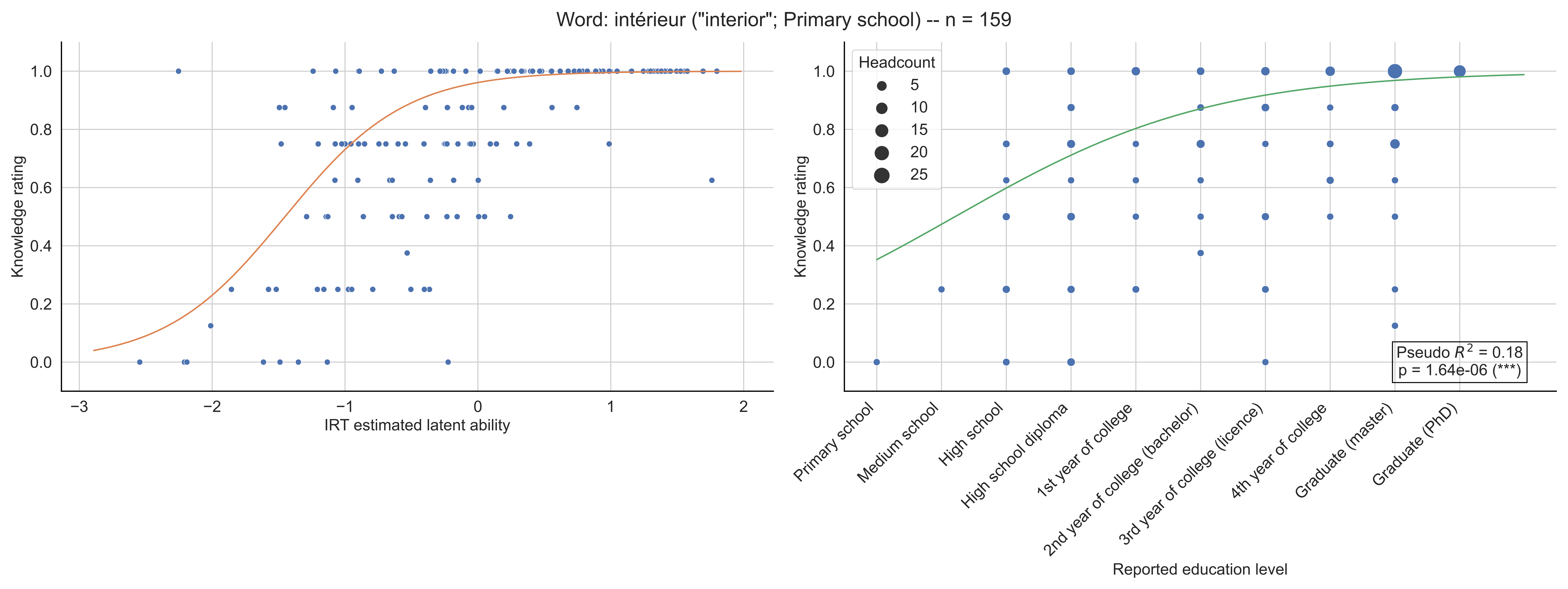

Word: intérieur ("interior"; Primary school) -- n = 159
Knowledge rating
IRT estimated latent ability
Headcount
5
10
15
20
25
Pseudo $R^2$ = 0.18
p = 1.64e-06 (***)
Primary school
Medium school
High school
High school diploma
1st year of college
2nd year of college (bachelor)
3rd year of college (licence)
4th year of college
Graduate (master)
Graduate (PhD)
Reported education level

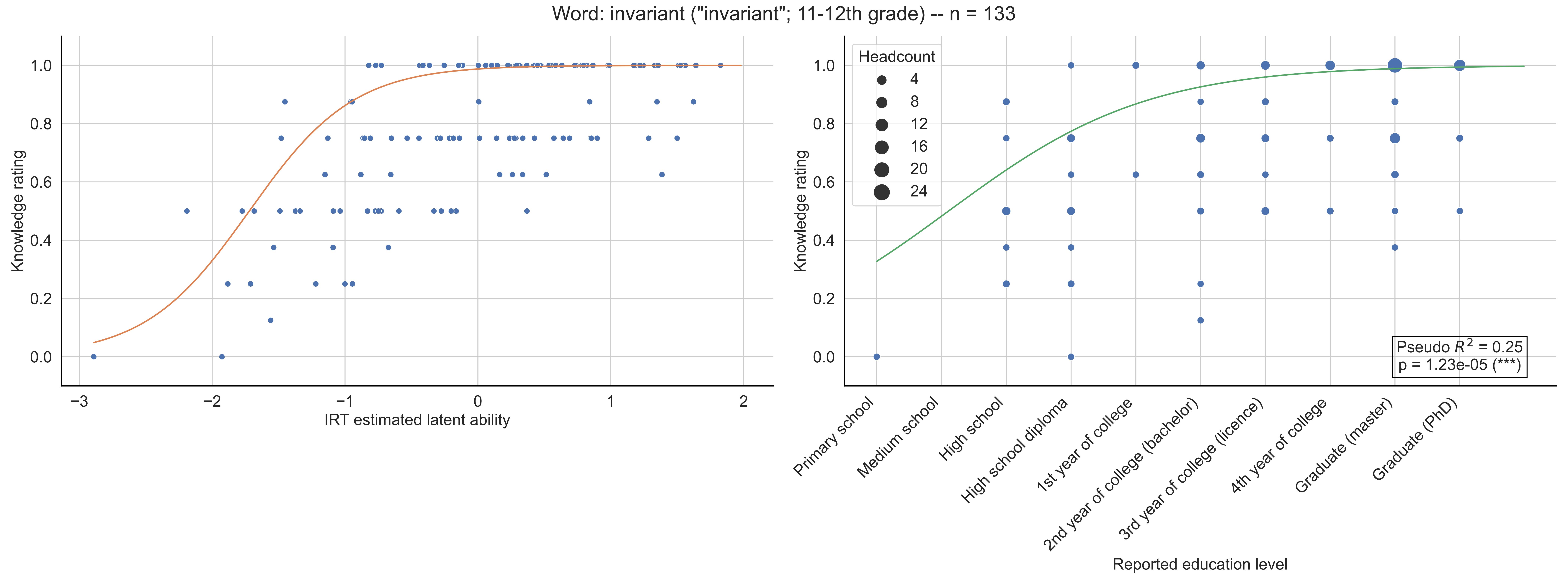

Word: invariant ("invariant"; 11-12th grade) -- n = 133
Knowledge rating
IRT estimated latent ability
Headcount
4
8
12
16
20
24
Pseudo $R^2$ = 0.25
p = 1.23e-05 (***)
Primary school
Medium school
High school
High school diploma
1st year of college
2nd year of college (bachelor)
3rd year of college (licence)
4th year of college
Graduate (master)
Graduate (PhD)
Reported education level

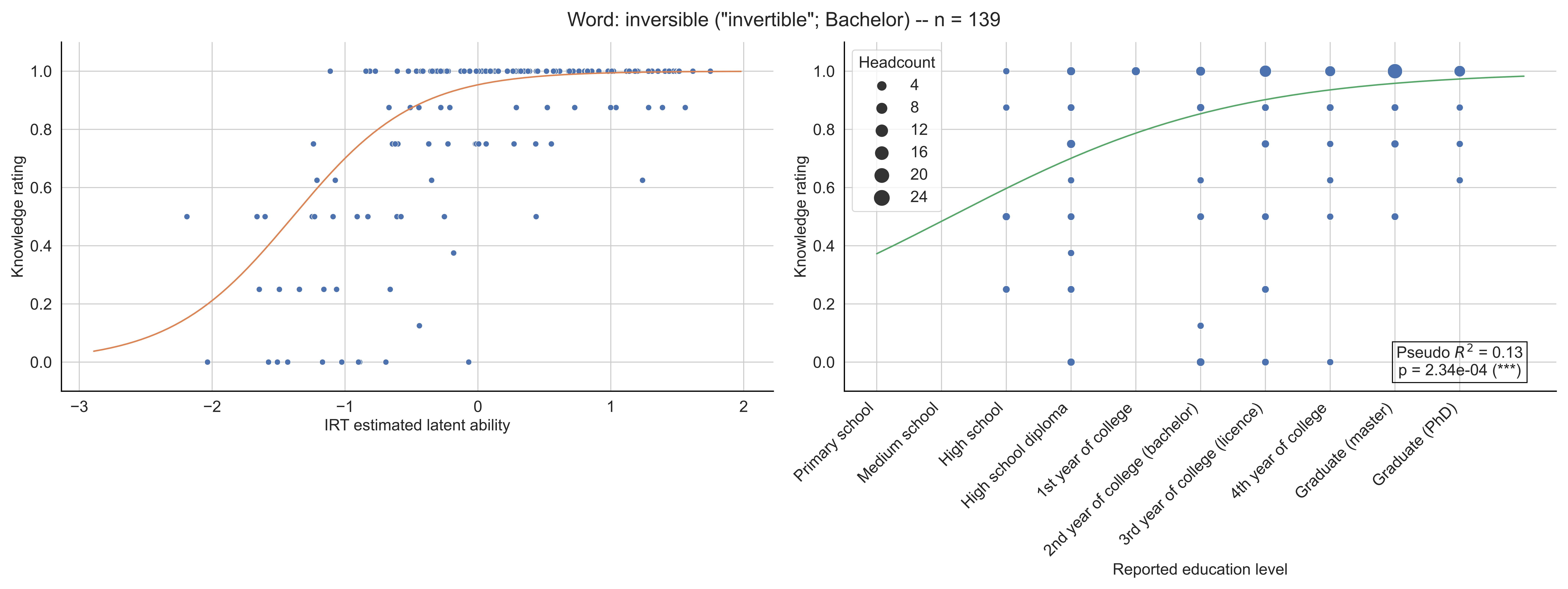

Word: inversible ("invertible"; Bachelor) -- n = 139
Knowledge rating
1.0
0.8
0.6
0.4
0.2
0.0
−3
−2
−1
0
1
2
IRT estimated latent ability
Headcount
4
8
12
16
20
24
Pseudo $R^2$ = 0.13
p = 2.34e-04 (***)
Primary school
Medium school
High school
High school diploma
1st year of college
2nd year of college (bachelor)
3rd year of college (licence)
4th year of college
Graduate (master)
Graduate (PhD)
Reported education level

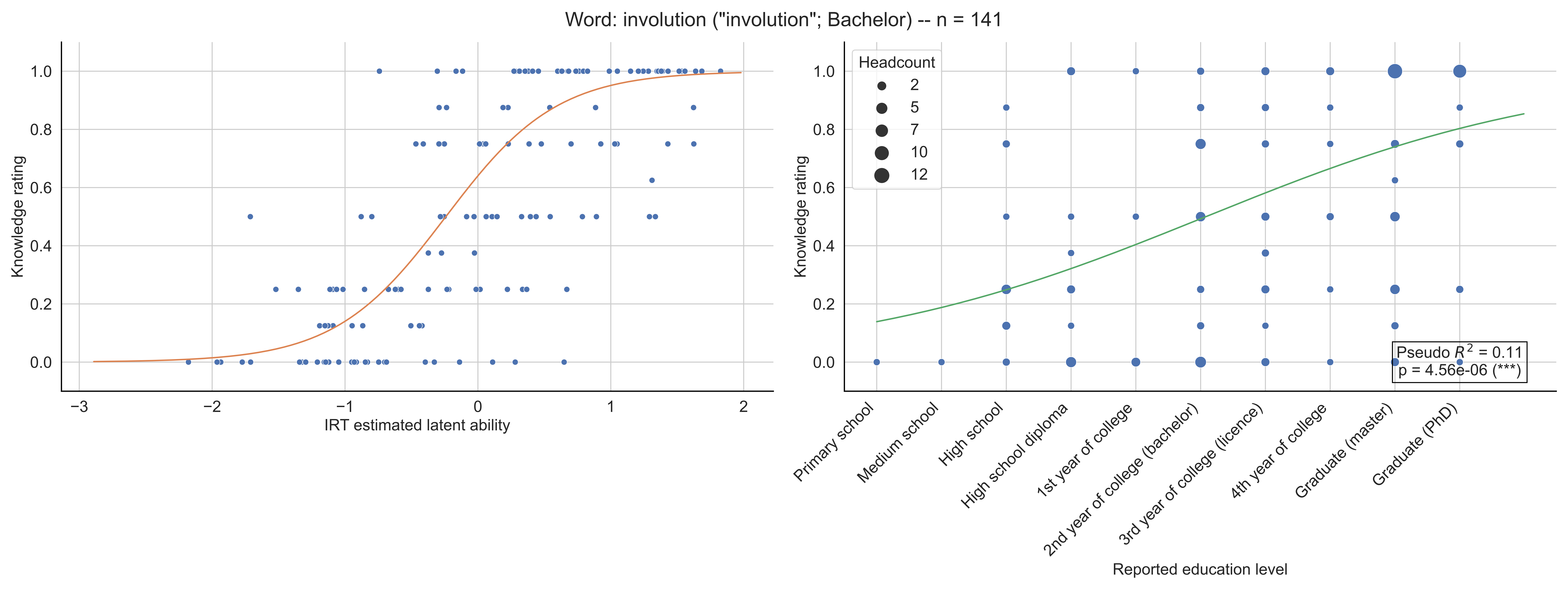

Word: involution ("involution"; Bachelor) -- n = 141
Knowledge rating
IRT estimated latent ability
Headcount
2
5
7
10
12
Pseudo $R^2$ = 0.11
p = 4.56e-06 (***)
Primary school
Medium school
High school
High school diploma
1st year of college
2nd year of college (bachelor)
3rd year of college (licence)
4th year of college
Graduate (master)
Graduate (PhD)
Reported education level

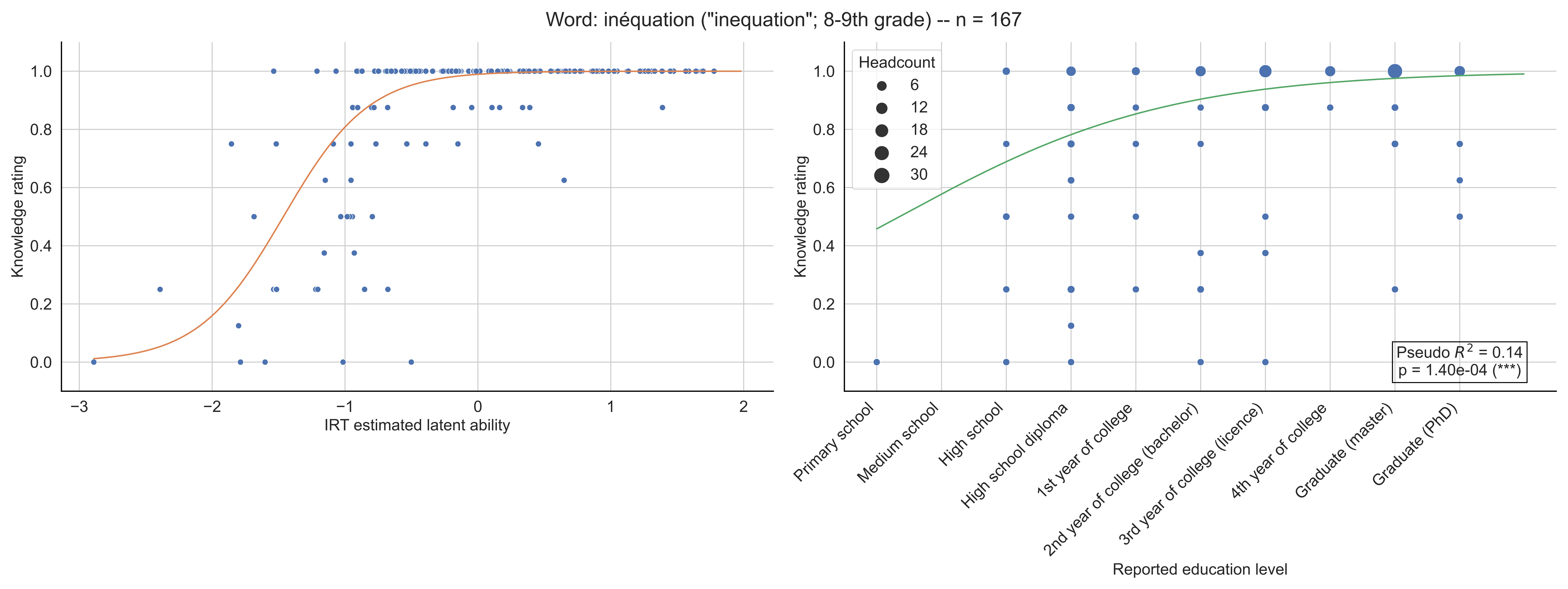

Word: inéquation ("inequation"; 8-9th grade) -- n = 167
Knowledge rating
IRT estimated latent ability
Headcount
6
12
18
24
30
Pseudo $R^2$ = 0.14
p = 1.40e-04 (***)
Primary school
Medium school
High school
High school diploma
1st year of college
2nd year of college (bachelor)
3rd year of college (licence)
4th year of college
Graduate (master)
Graduate (PhD)
Reported education level

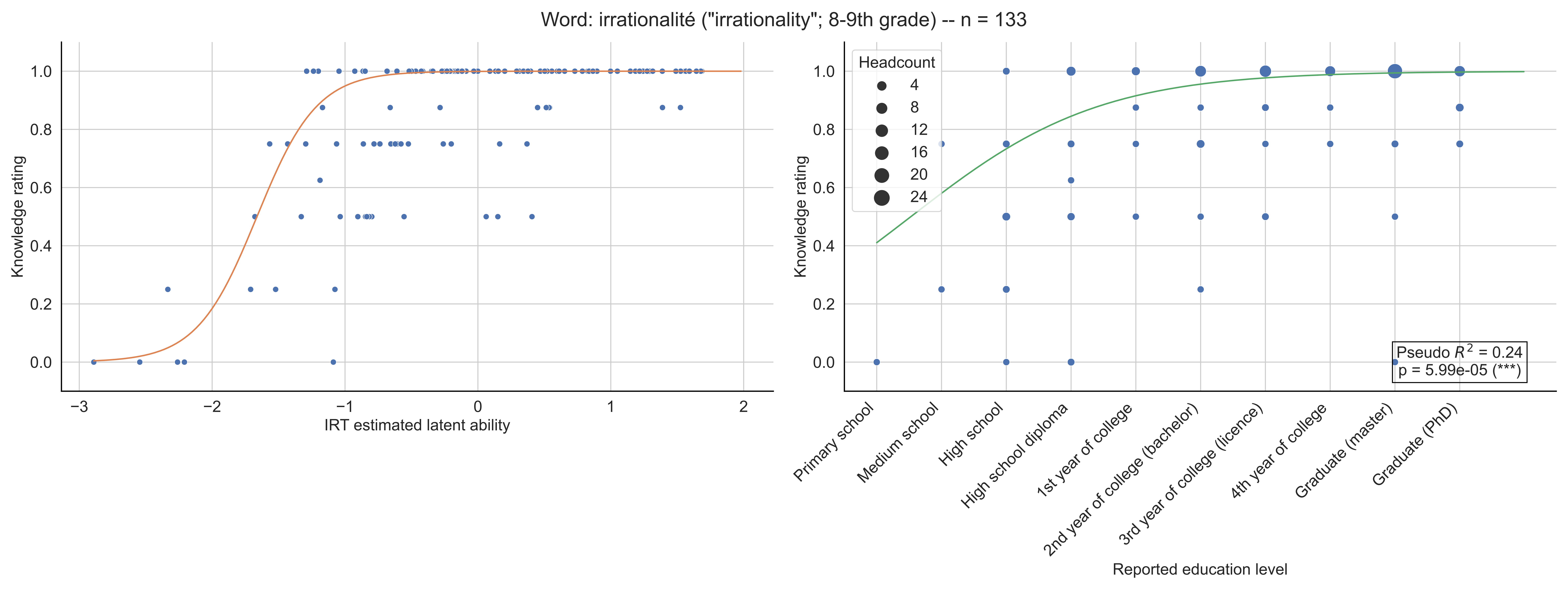
Word: irrationalité ("irrationality"; 8-9th grade) -- n = 133
Knowledge rating
IRT estimated latent ability
Headcount
4
8
12
16
20
24
Pseudo $R^2$ = 0.24
p = 5.99e-05 (***)
Primary school
Medium school
High school
High school diploma
1st year of college
2nd year of college (bachelor)
3rd year of college (licence)
4th year of college
Graduate (master)
Graduate (PhD)
Reported education level

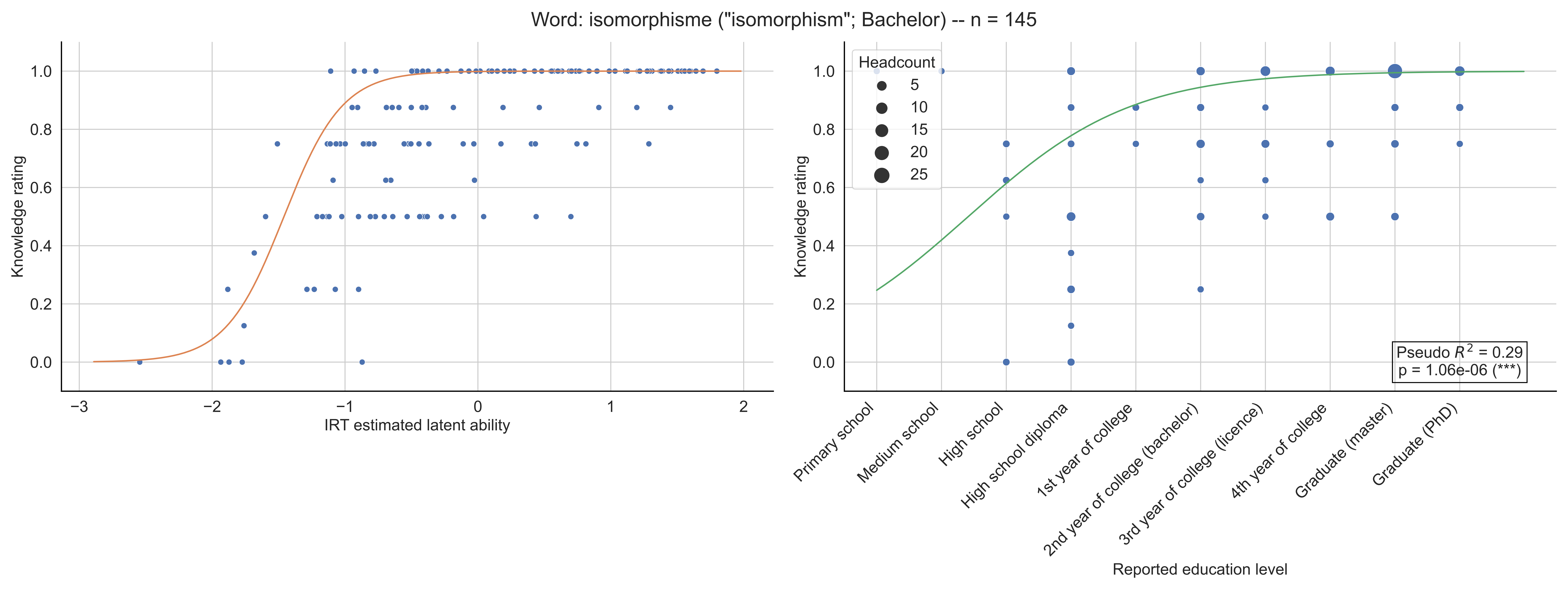

Word: isomorphisme ("isomorphism"; Bachelor) -- n = 145
Knowledge rating
IRT estimated latent ability
Headcount
5
10
15
20
25
Pseudo $R^2$ = 0.29
p = 1.06e-06 (***)
Primary school
Medium school
High school
High school diploma
1st year of college
2nd year of college (bachelor)
3rd year of college (licence)
4th year of college
Graduate (master)
Graduate (PhD)
Reported education level

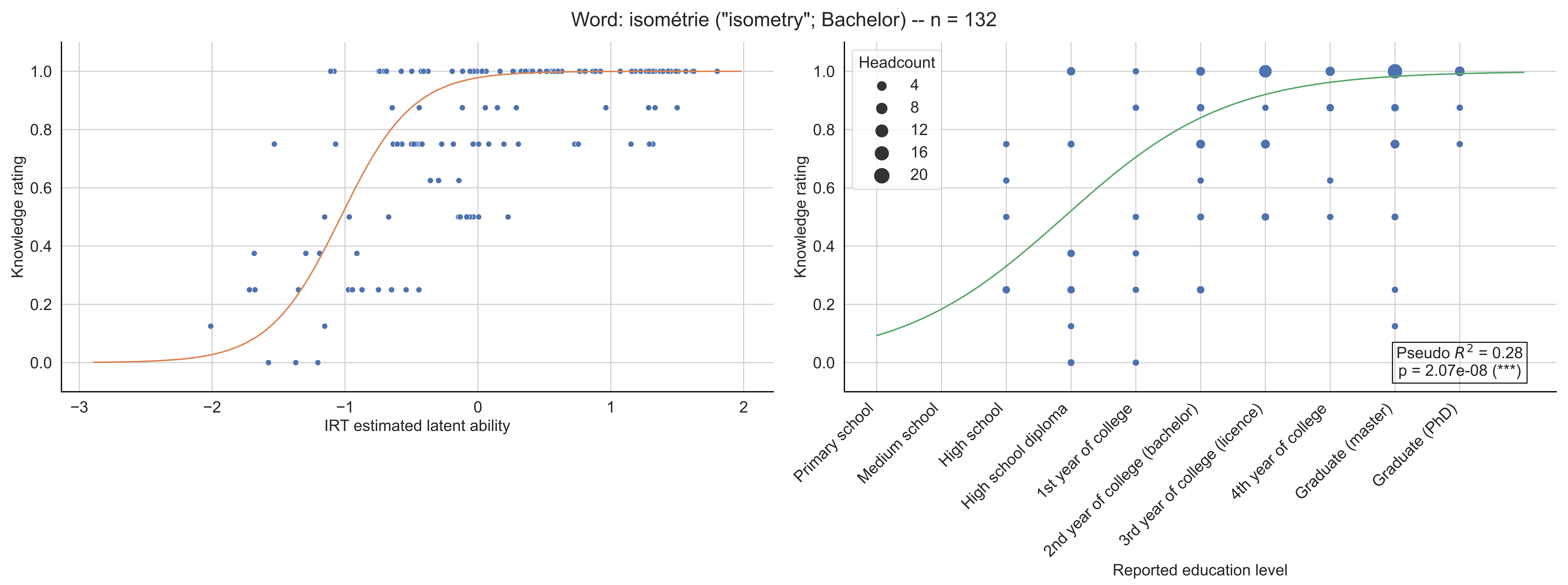
Word: isométrie ("isometry"; Bachelor) -- n = 132
Knowledge rating
IRT estimated latent ability
Headcount
4
8
12
16
20
Primary school
Medium school
High school
High school diploma
1st year of college
2nd year of college (bachelor)
3rd year of college (licence)
4th year of college
Graduate (master)
Graduate (PhD)
Pseudo $R^2$ = 0.28
p = 2.07e-08 (***)
Reported education level

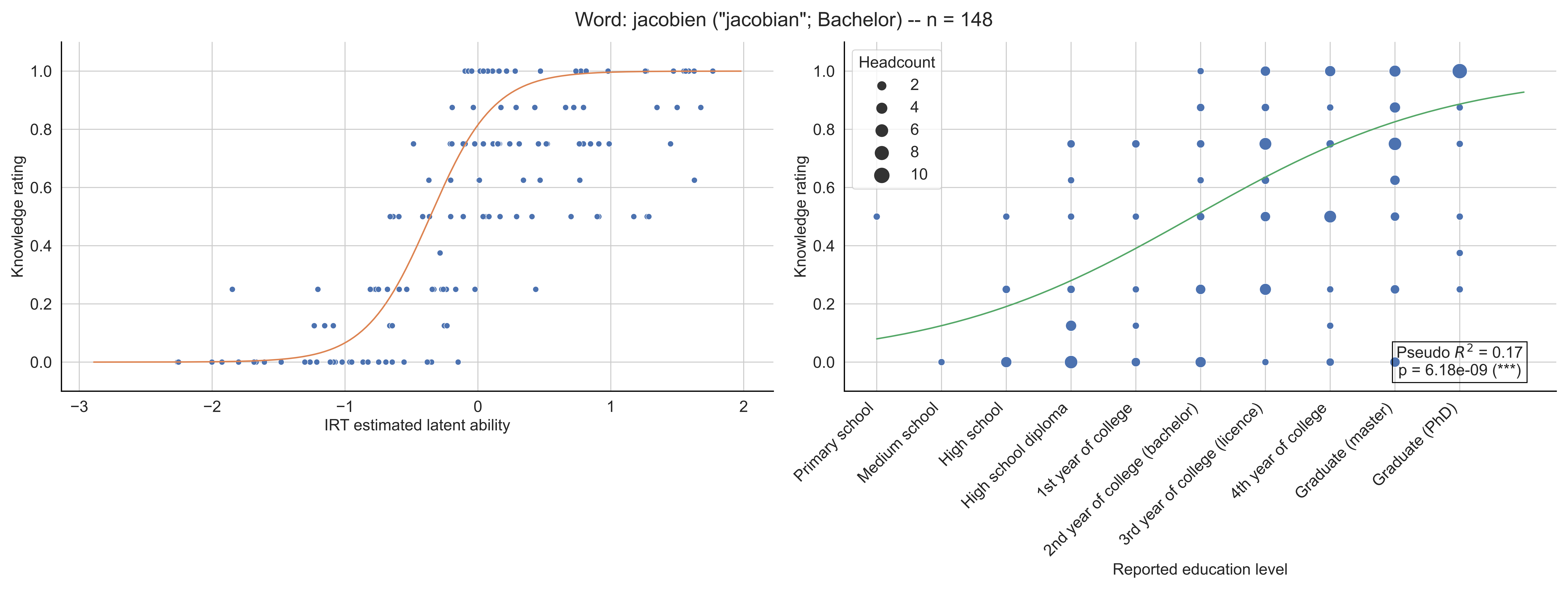

Word: jacobien ("jacobian"; Bachelor) -- n = 148
Knowledge rating
IRT estimated latent ability
Headcount
2
4
6
8
10
Pseudo $R^2$ = 0.17
p = 6.18e-09 (***)
Primary school
Medium school
High school
High school diploma
1st year of college
2nd year of college (bachelor)
3rd year of college (licence)
4th year of college
Graduate (master)
Graduate (PhD)
Reported education level

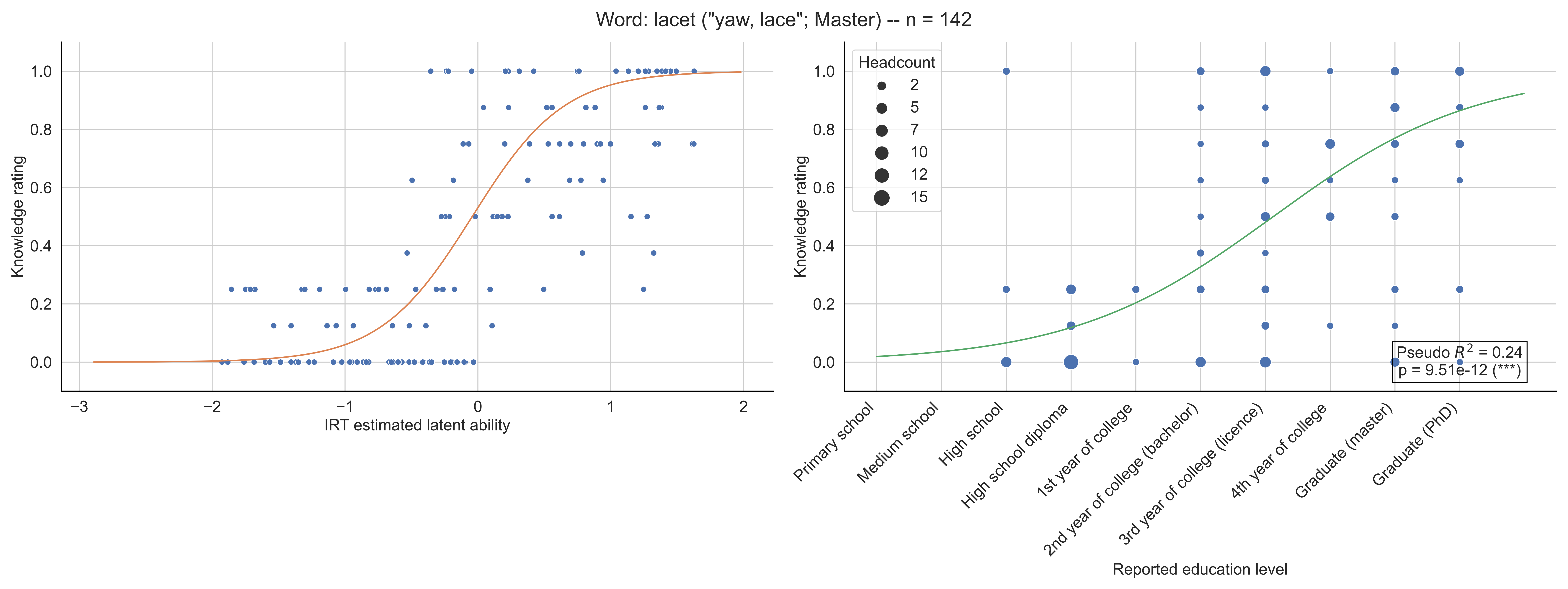
Word: lacet ("yaw, lace"; Master) -- n = 142
Knowledge rating
IRT estimated latent ability
Headcount
2
5
7
10
12
15
Pseudo $R^2$ = 0.24
p = 9.51e-12 (***)
Primary school
Medium school
High school
High school diploma
1st year of college
2nd year of college (bachelor)
3rd year of college (licence)
4th year of college
Graduate (master)
Graduate (PhD)
Reported education level

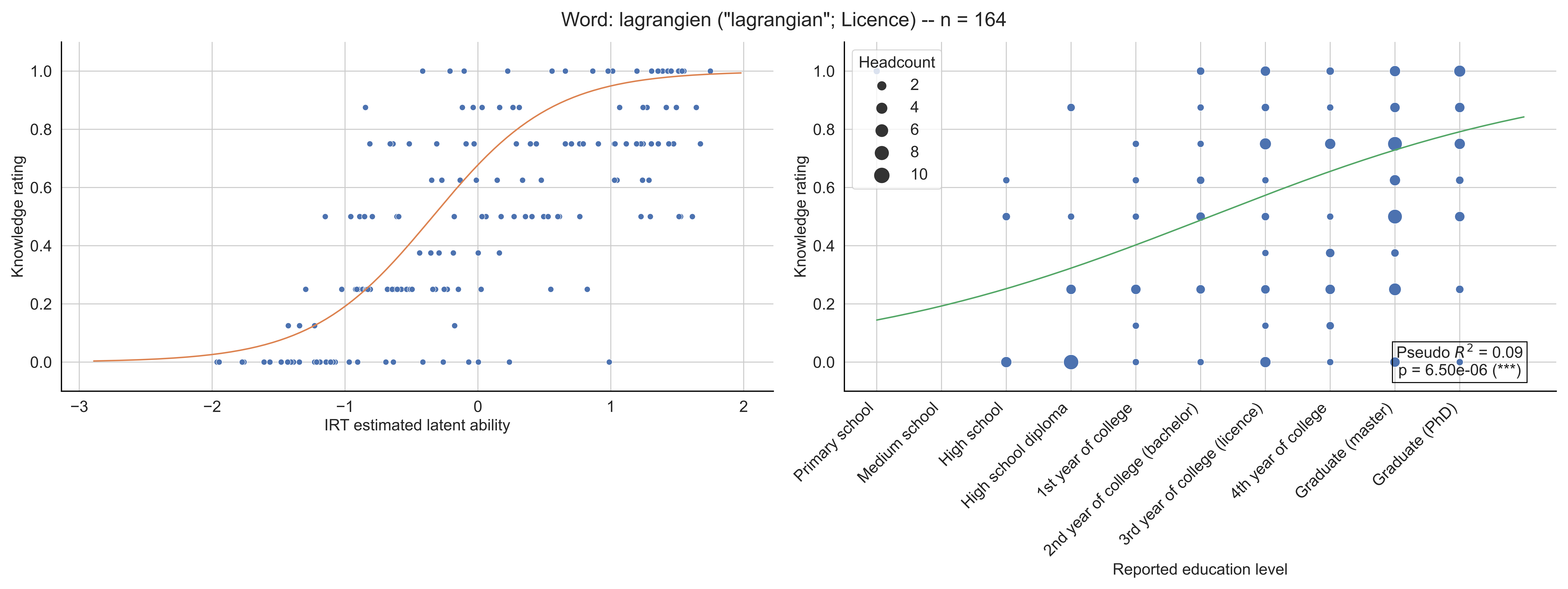

Word: lagrangien ("lagrangian"; Licence) -- n = 164
Knowledge rating
IRT estimated latent ability
Headcount
2
4
6
8
10
Pseudo $R^2$ = 0.09
p = 6.50e-06 (***)
Primary school
Medium school
High school
High school diploma
1st year of college
2nd year of college (bachelor)
3rd year of college (licence)
4th year of college
Graduate (master)
Graduate (PhD)
Reported education level

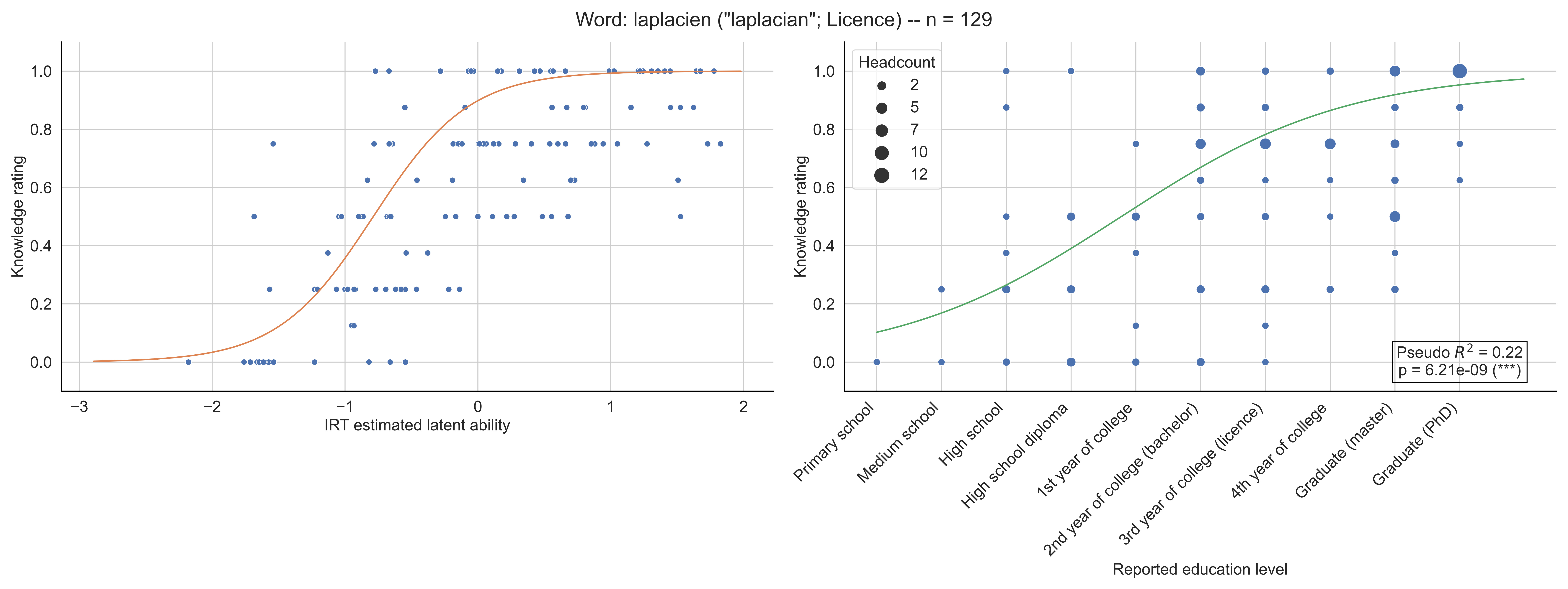

Word: laplacien ("laplacian"; Licence) -- n = 129
Knowledge rating
IRT estimated latent ability
Headcount
2
5
7
10
12
Pseudo $R^2$ = 0.22
p = 6.21e-09 (***)
Primary school
Medium school
High school
High school diploma
1st year of college
2nd year of college (bachelor)
3rd year of college (licence)
4th year of college
Graduate (master)
Graduate (PhD)
Reported education level

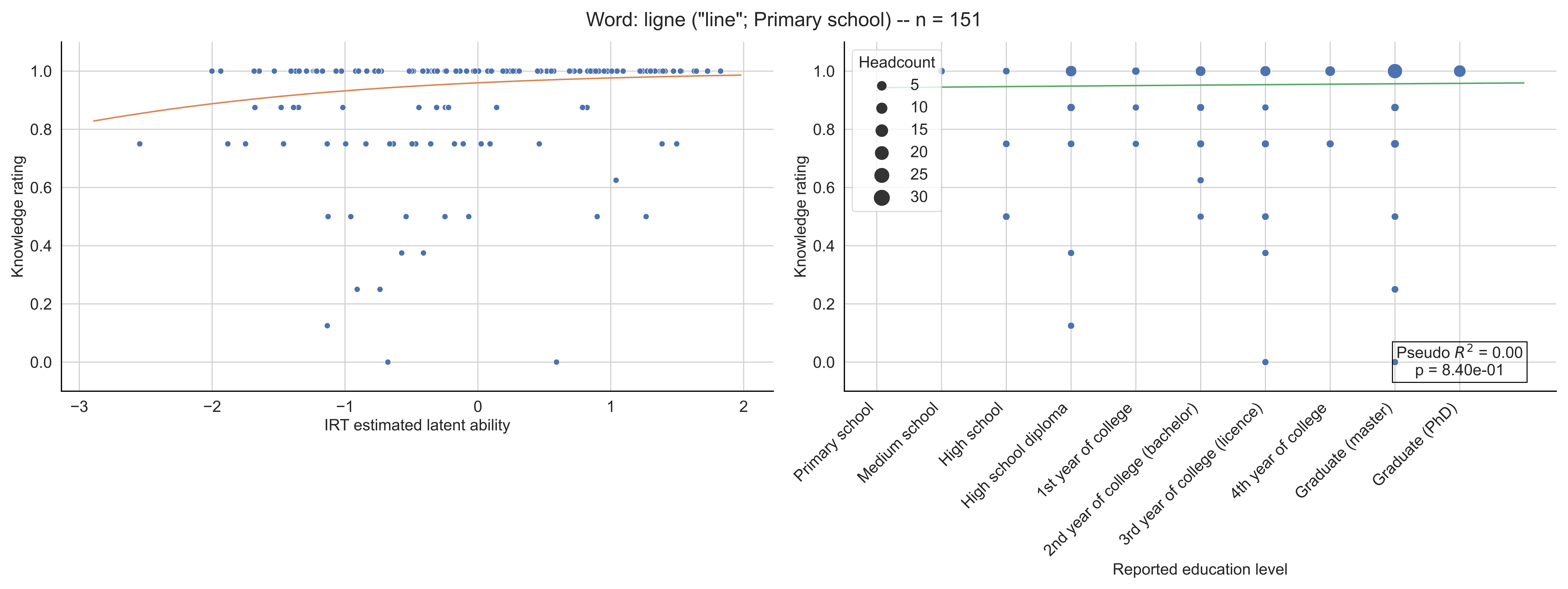

Word: ligne ("line"; Primary school) -- n = 151
Knowledge rating
IRT estimated latent ability
Headcount
5
10
15
20
25
30
Pseudo $R^2$ = 0.00
p = 8.40e-01
Primary school
Medium school
High school
High school diploma
1st year of college
2nd year of college (bachelor)
3rd year of college (licence)
4th year of college
Graduate (master)
Graduate (PhD)
Reported education level

Word: linéarité ("linearity"; 6-7th grade) -- n = 168

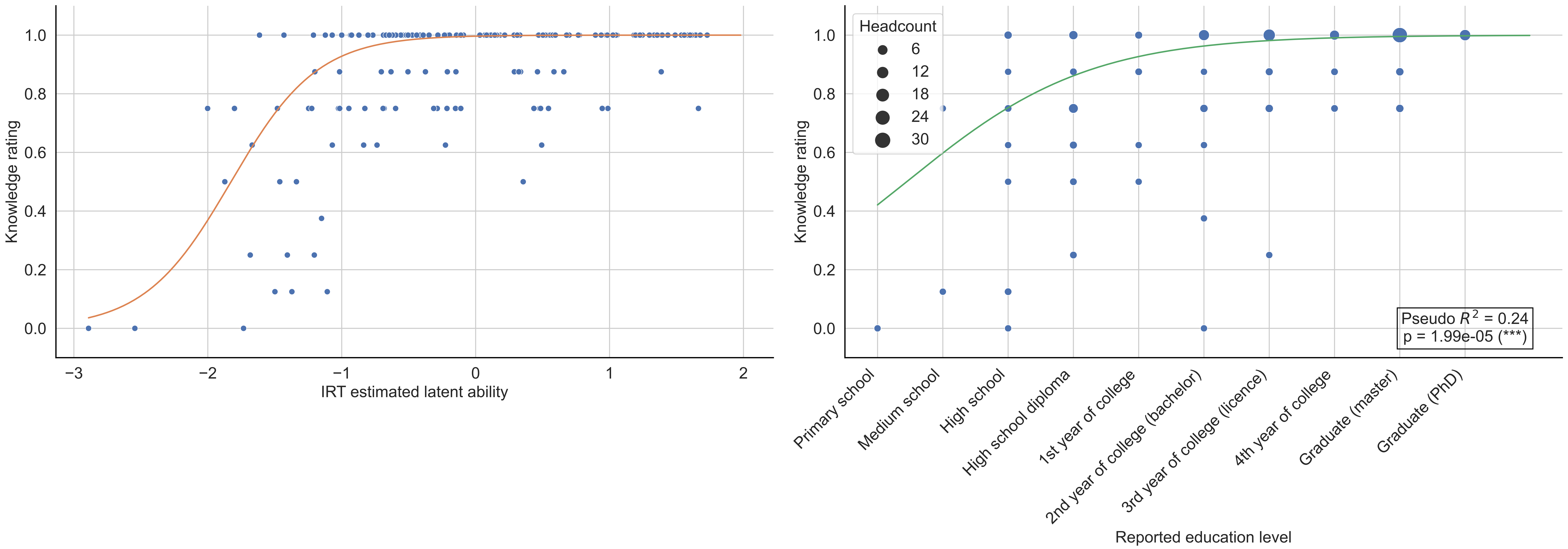

Word: logique ("logic"; 6-7th grade) -- n = 169

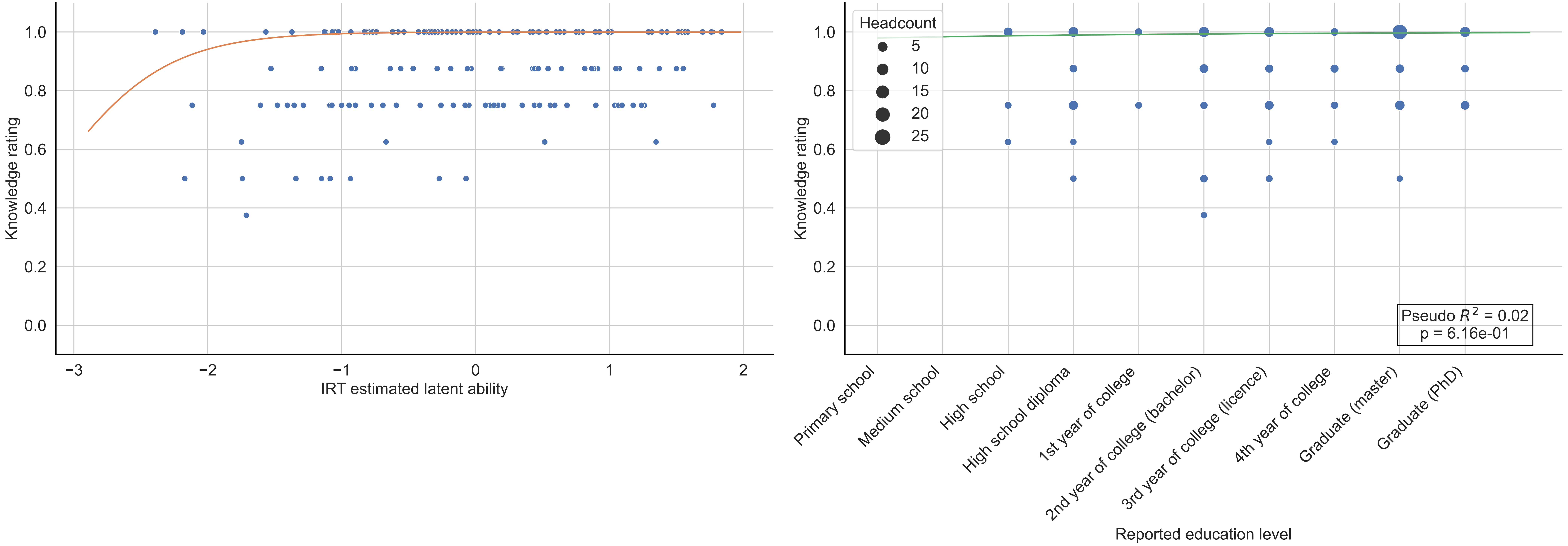

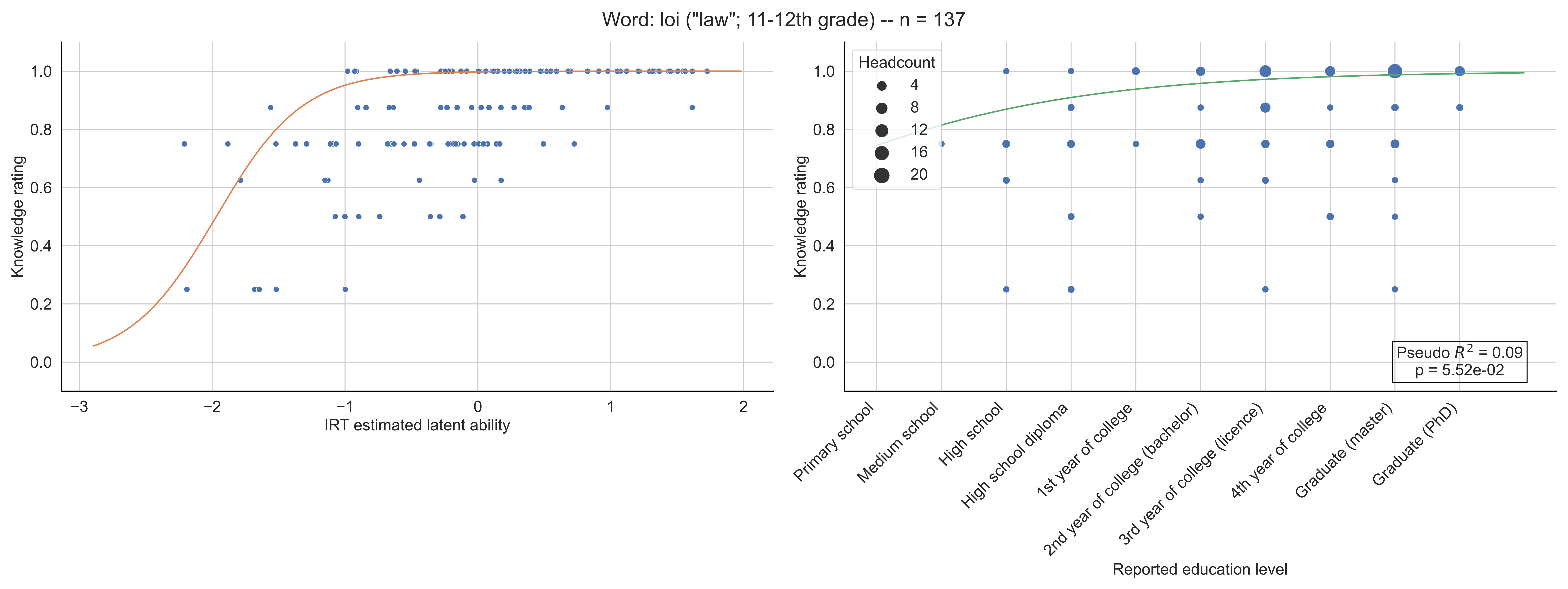

Word: loi ("law"; 11-12th grade) -- n = 137
Knowledge rating
IRT estimated latent ability
Headcount
4
8
12
16
20
Pseudo $R^2$ = 0.09
p = 5.52e-02
Primary school
Medium school
High school
High school diploma
1st year of college
2nd year of college (bachelor)
3rd year of college (licence)
4th year of college
Graduate (master)
Graduate (PhD)
Reported education level

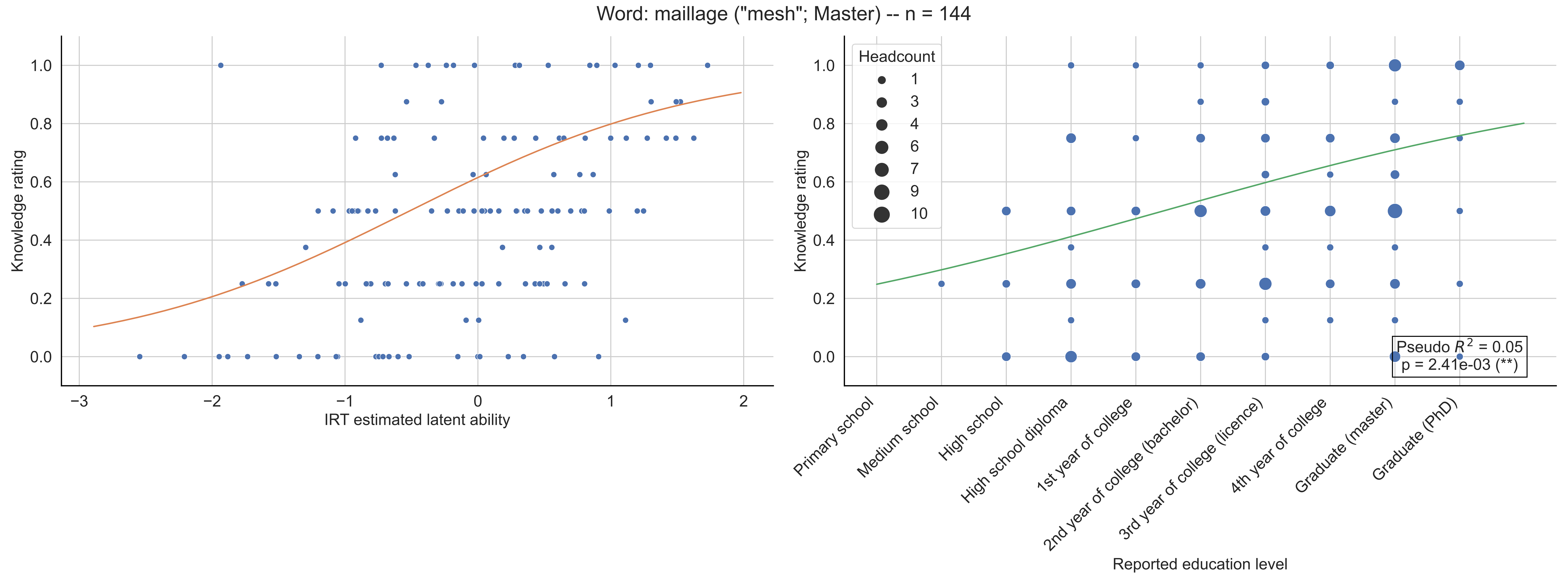
Word: maillage ("mesh"; Master) -- n = 144
Knowledge rating
IRT estimated latent ability
Headcount
1
3
4
6
7
9
10
Pseudo $R^2$ = 0.05
p = 2.41e-03 (**)
Primary school
Medium school
High school
High school diploma
1st year of college
2nd year of college (bachelor)
3rd year of college (licence)
4th year of college
Graduate (master)
Graduate (PhD)
Reported education level

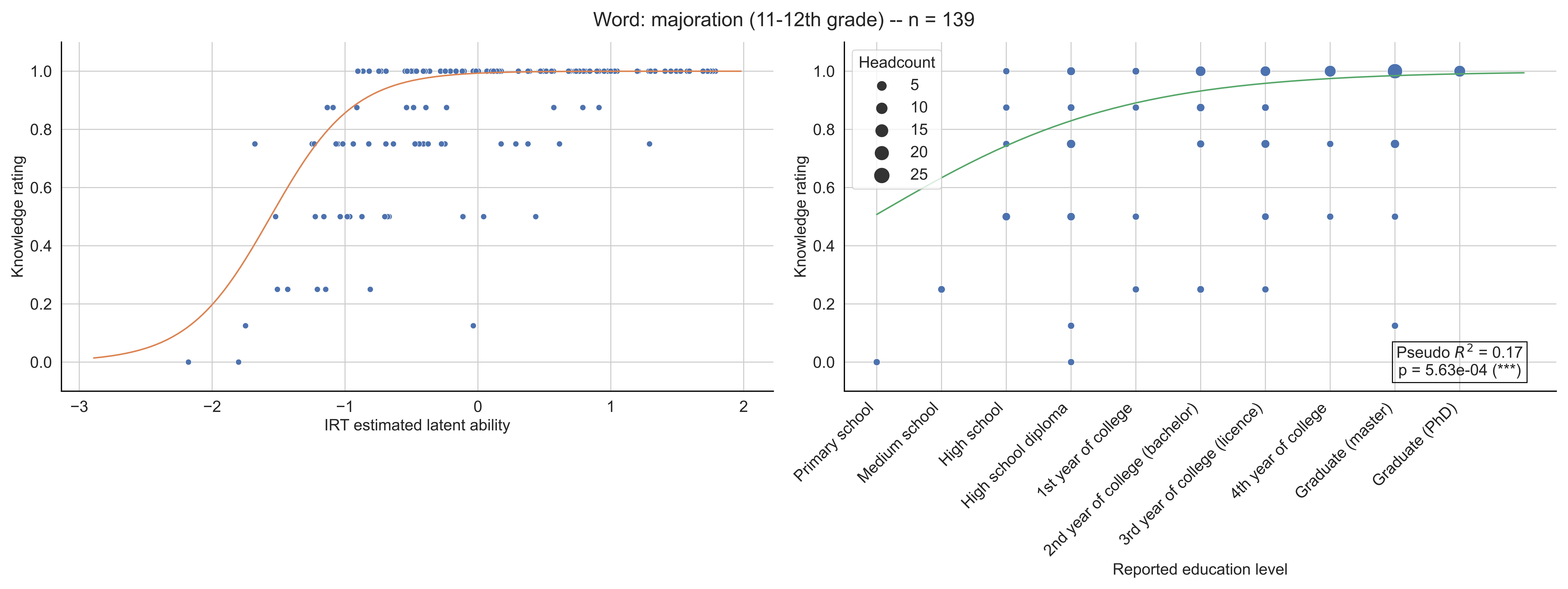

Word: majoration (11-12th grade) -- n = 139
Knowledge rating
IRT estimated latent ability
Headcount
5
10
15
20
25
Pseudo $R^2$ = 0.17
p = 5.63e-04 (***)
Primary school
Medium school
High school
High school diploma
1st year of college
2nd year of college (bachelor)
3rd year of college (licence)
4th year of college
Graduate (master)
Graduate (PhD)
Reported education level

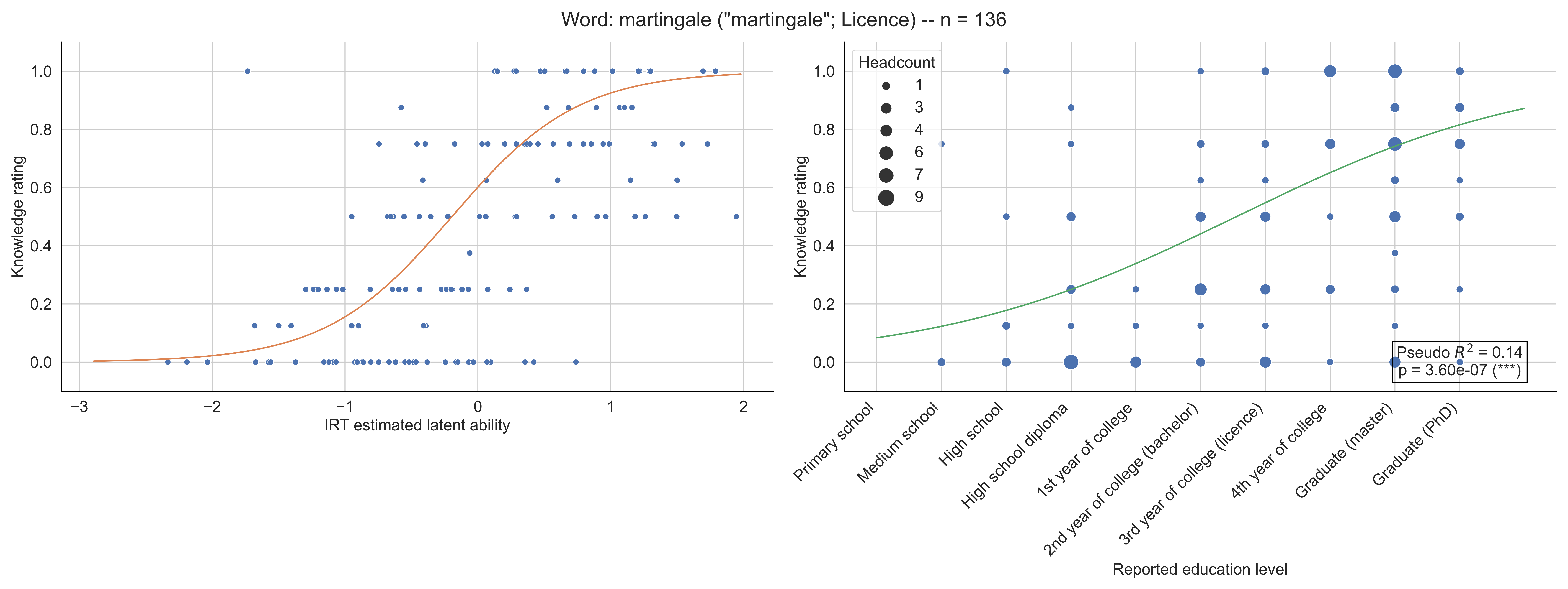
Word: martingale ("martingale"; Licence) -- n = 136
Knowledge rating
IRT estimated latent ability
Headcount
1
3
4
6
7
9
Pseudo $R^2$ = 0.14
p = 3.60e-07 (***)
Primary school
Medium school
High school
High school diploma
1st year of college
2nd year of college (bachelor)
3rd year of college (licence)
4th year of college
Graduate (master)
Graduate (PhD)
Reported education level

Word: matrice ("matrix"; 11-12th grade) -- n = 148

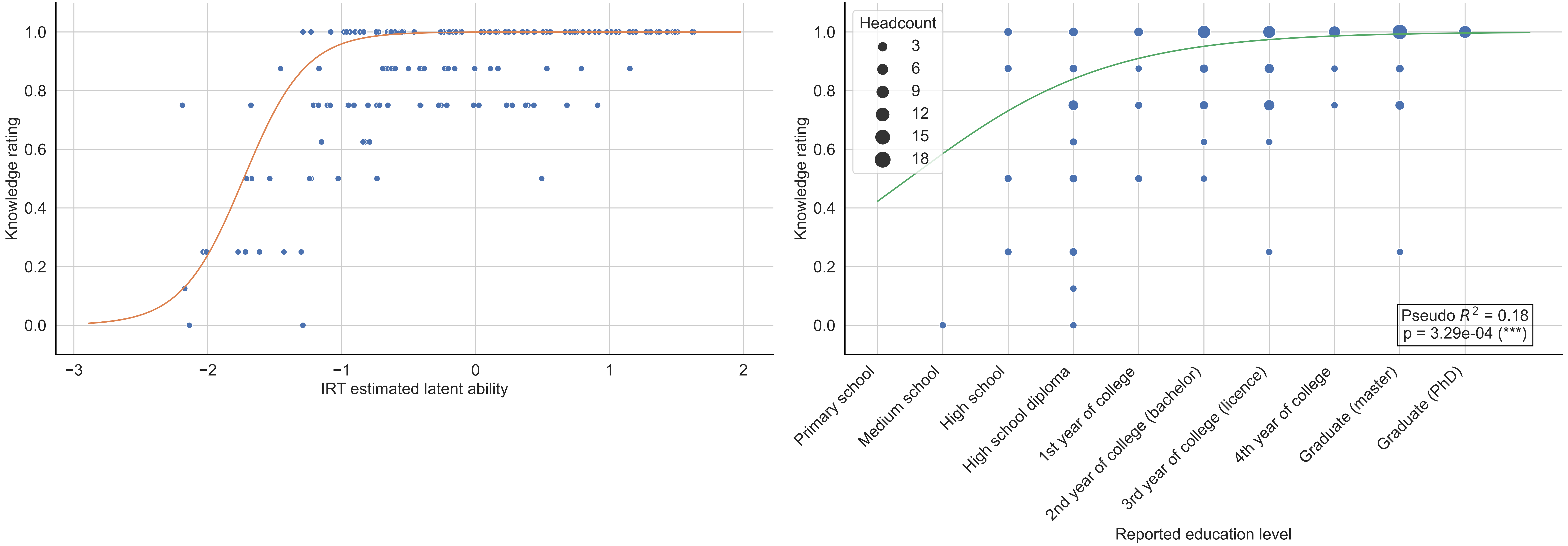

Word: mesure ("measure"; Primary school) -- n = 148

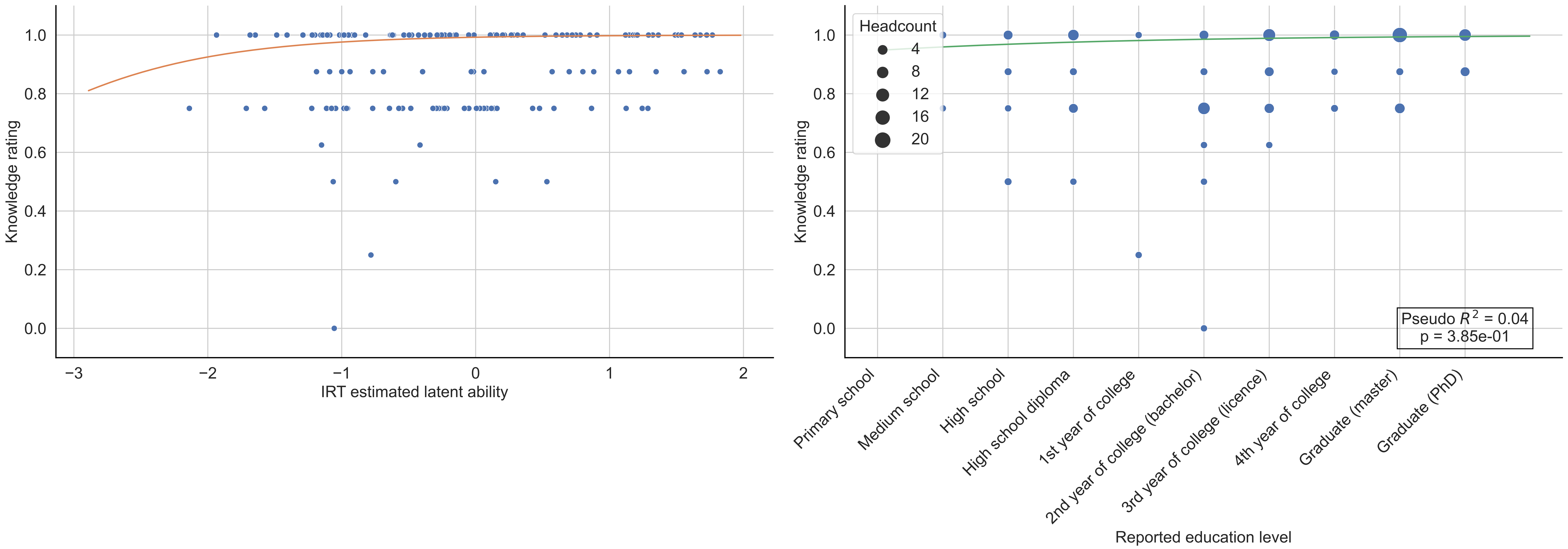

Word: milieu ("midpoint"; Primary school) -- n = 144

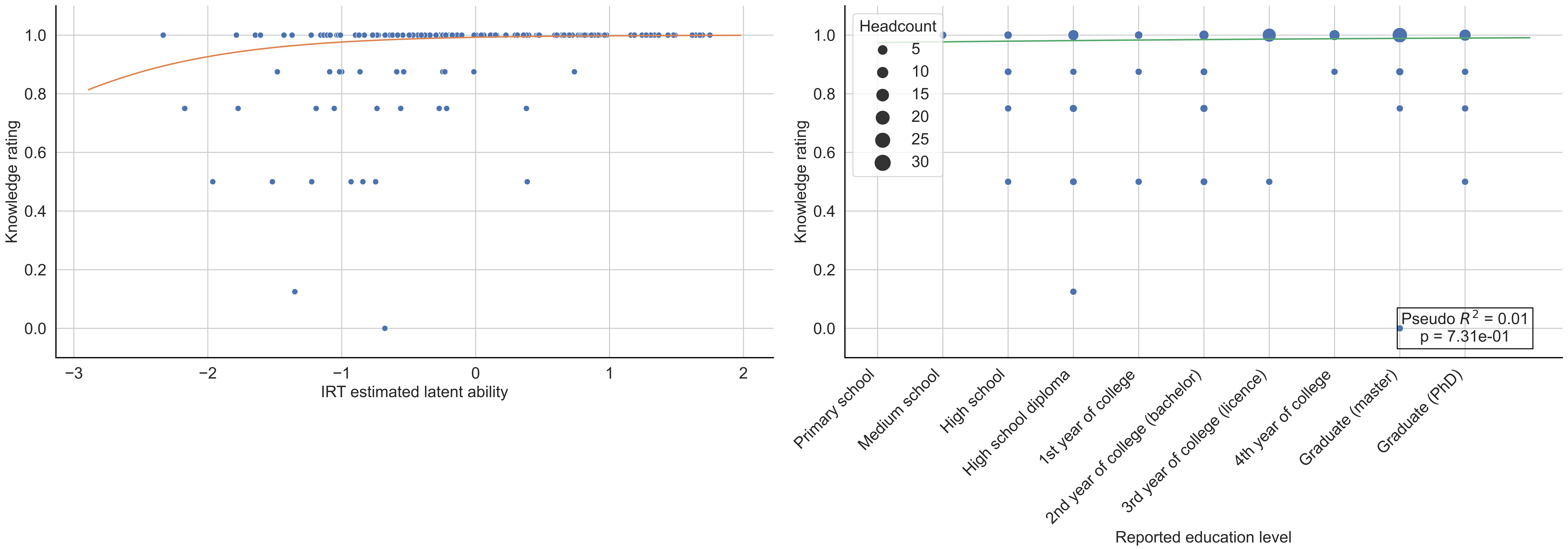

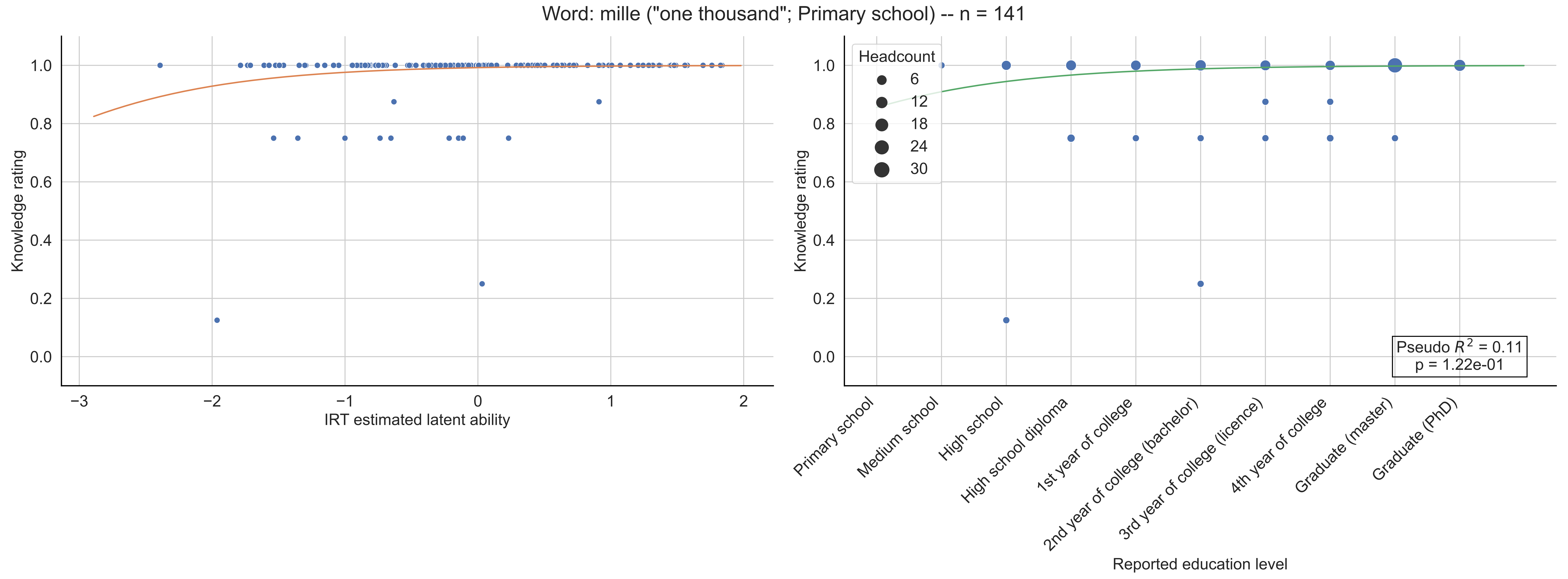

Word: mille ("one thousand"; Primary school) -- n = 141
Knowledge rating
IRT estimated latent ability
Headcount
6
12
18
24
30
Pseudo $R^2$ = 0.11
p = 1.22e-01
Primary school
Medium school
High school
High school diploma
1st year of college
2nd year of college (bachelor)
3rd year of college (licence)
4th year of college
Graduate (master)
Graduate (PhD)
Reported education level

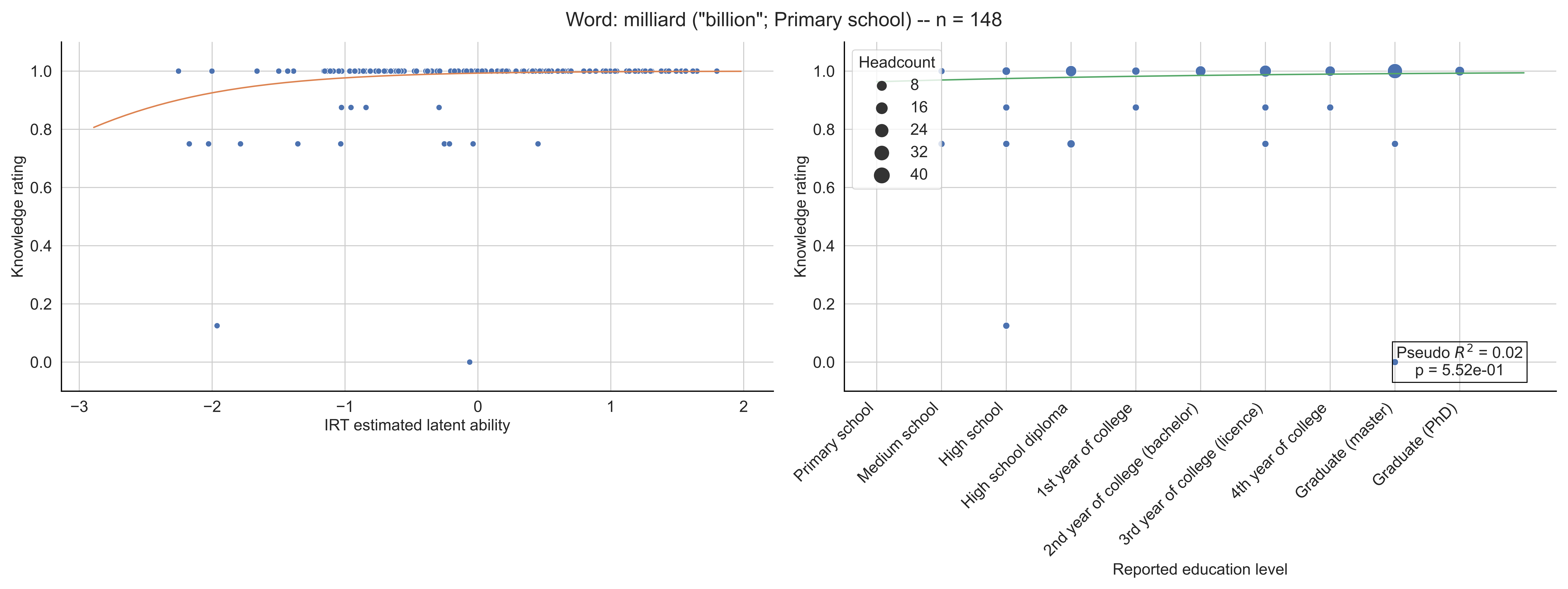
Word: milliard ("billion"; Primary school) -- n = 148
Knowledge rating
IRT estimated latent ability
Headcount
8
16
24
32
40
Pseudo $R^2$ = 0.02
p = 5.52e-01
Primary school
Medium school
High school
High school diploma
1st year of college
2nd year of college (bachelor)
3rd year of college (licence)
4th year of college
Graduate (master)
Graduate (PhD)
Reported education level

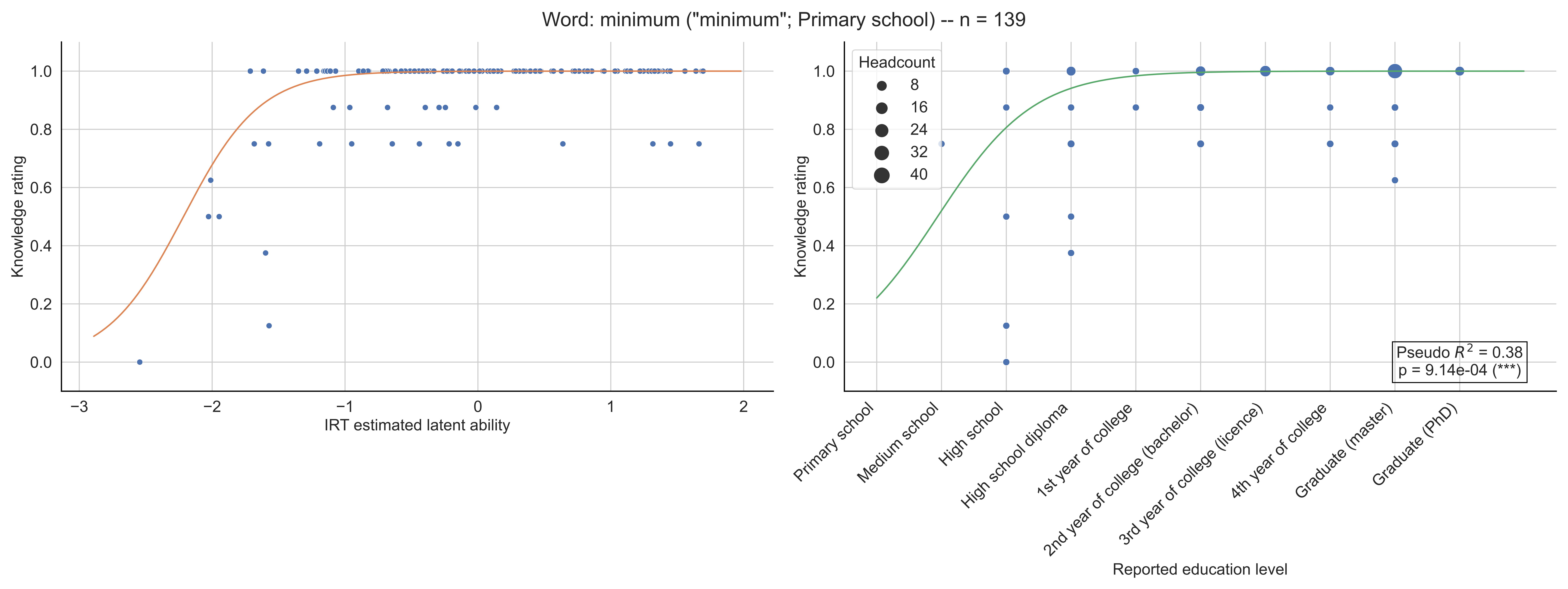

Word: minimum ("minimum"; Primary school) -- n = 139
Knowledge rating
IRT estimated latent ability
−3
−2
−1
0
1
2
0.0
0.2
0.4
0.6
0.8
1.0
Headcount
8
16
24
32
40
Primary school
Medium school
High school
High school diploma
1st year of college
2nd year of college (bachelor)
3rd year of college (licence)
4th year of college
Graduate (master)
Graduate (PhD)
Reported education level
Pseudo $R^2$ = 0.38
p = 9.14e-04 (***)

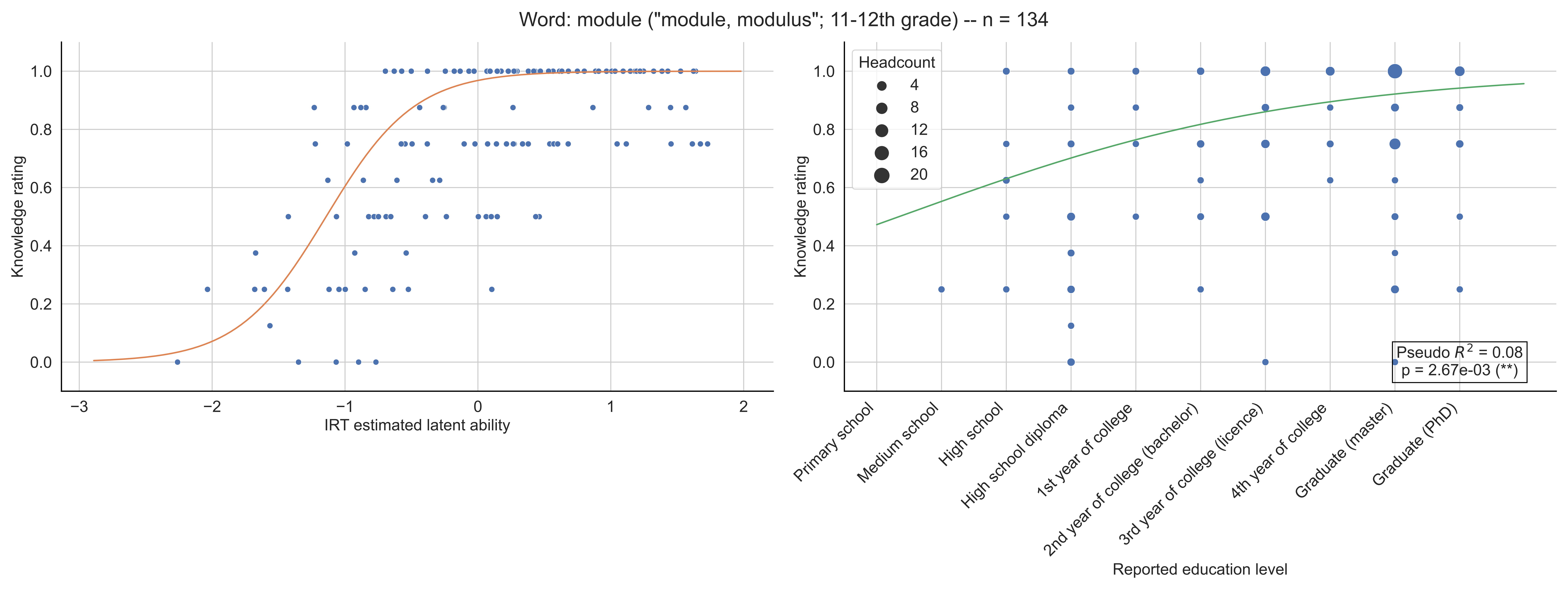
Word: module ("module, modulus"; 11-12th grade) -- n = 134
Knowledge rating
IRT estimated latent ability
Headcount
4
8
12
16
20
Pseudo $R^2$ = 0.08
p = 2.67e-03 (**)
Primary school
Medium school
High school
High school diploma
1st year of college
2nd year of college (bachelor)
3rd year of college (licence)
4th year of college
Graduate (master)
Graduate (PhD)
Reported education level

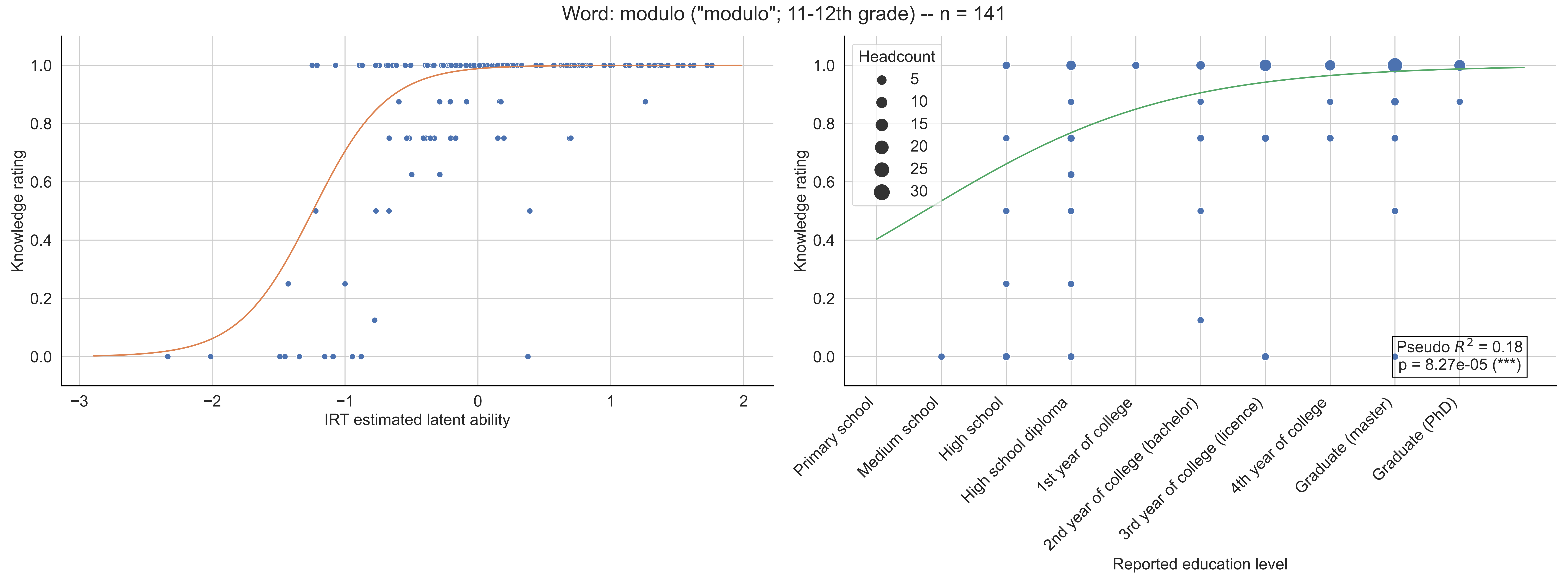

Word: modulo ("modulo"; 11-12th grade) -- n = 141
Knowledge rating
IRT estimated latent ability
Headcount
5
10
15
20
25
30
Pseudo $R^2$ = 0.18
p = 8.27e-05 (***)
Primary school
Medium school
High school
High school diploma
1st year of college
2nd year of college (bachelor)
3rd year of college (licence)
4th year of college
Graduate (master)
Graduate (PhD)
Reported education level

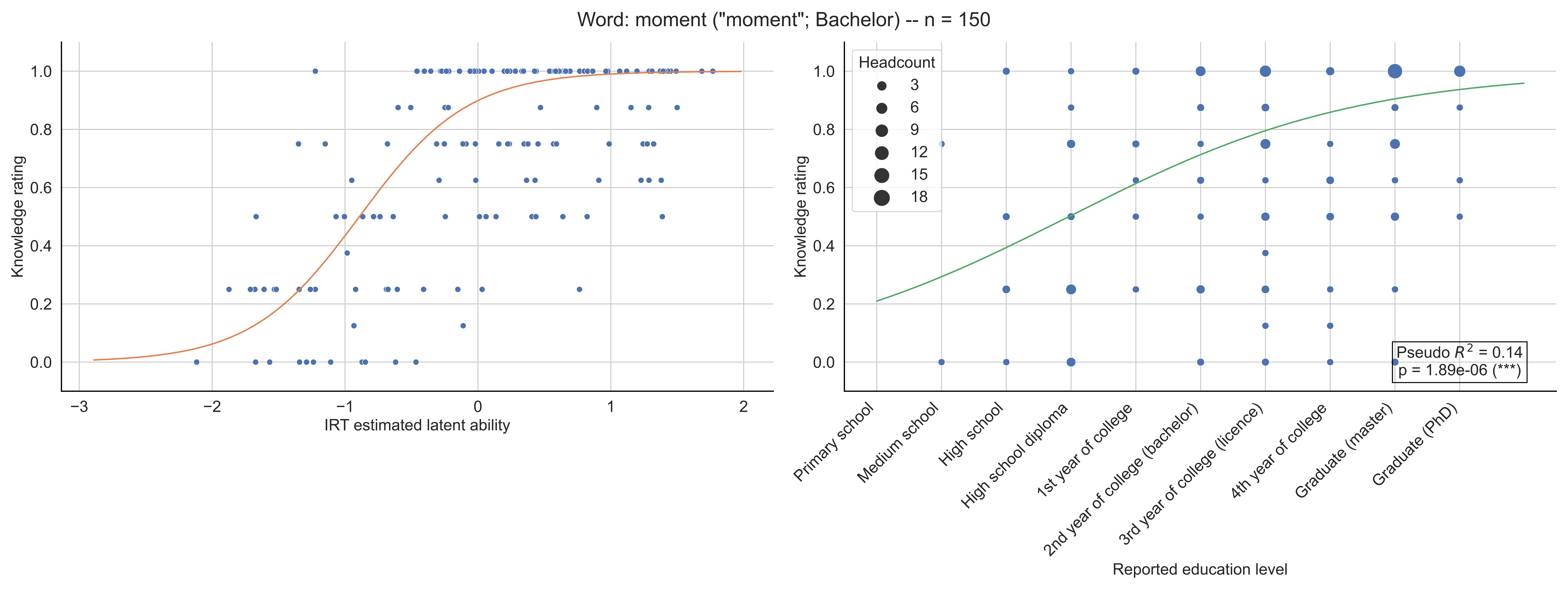
Word: moment ("moment"; Bachelor) -- n = 150
Knowledge rating
IRT estimated latent ability
Headcount
3
6
9
12
15
18
Pseudo $R^2$ = 0.14
p = 1.89e-06 (***)
Primary school
Medium school
High school
High school diploma
1st year of college
2nd year of college (bachelor)
3rd year of college (licence)
4th year of college
Graduate (master)
Graduate (PhD)
Reported education level

Word: monotonie ("monotonicity"; 10th grade) -- n = 141

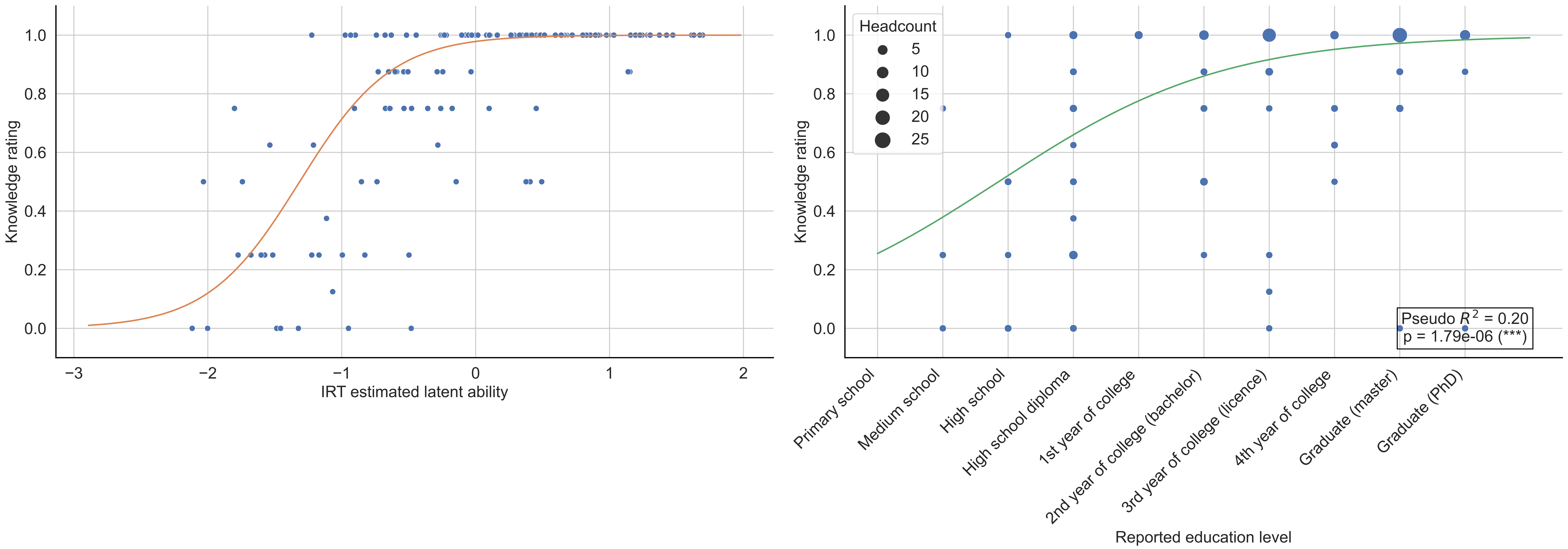

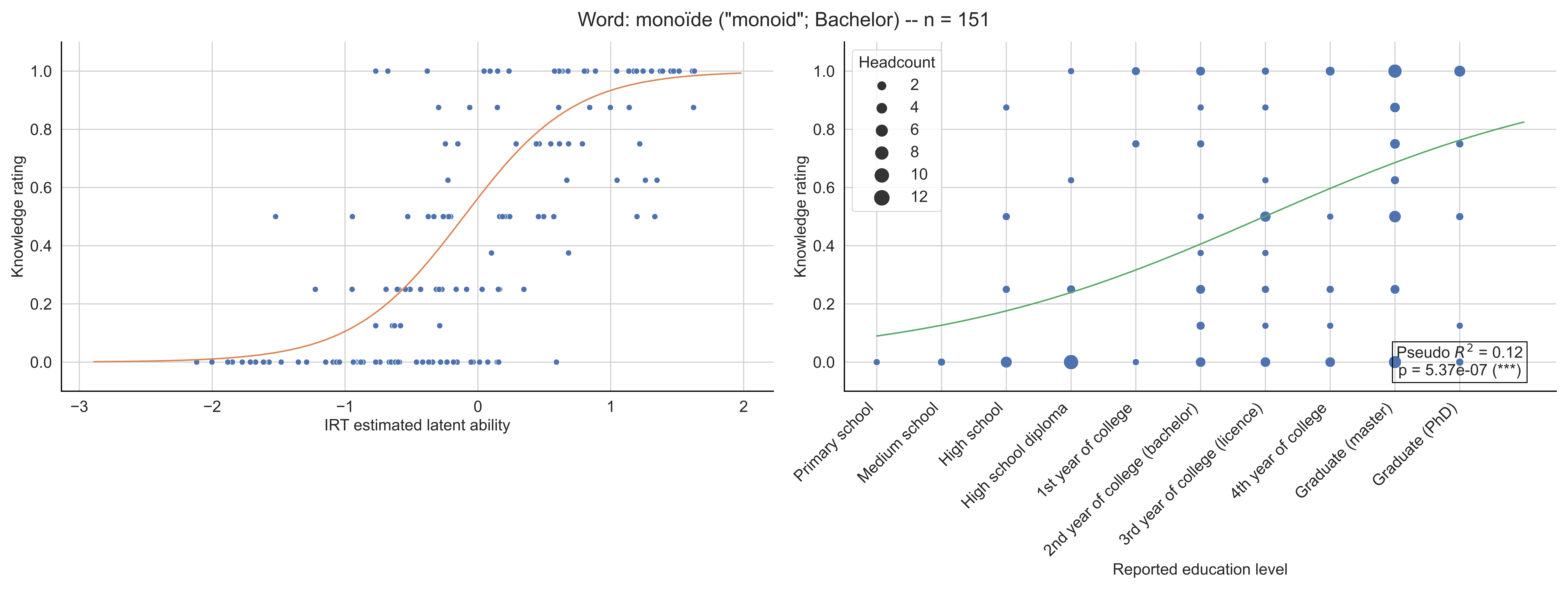
Word: monoïde ("monoid"; Bachelor) -- n = 151
Knowledge rating
IRT estimated latent ability
Headcount
2
4
6
8
10
12
Pseudo $R^2$ = 0.12
p = 5.37e-07 (***)
Primary school
Medium school
High school
High school diploma
1st year of college
2nd year of college (bachelor)
3rd year of college (licence)
4th year of college
Graduate (master)
Graduate (PhD)
Reported education level

Word: monôme ("monomial"; 10th grade) -- n = 132

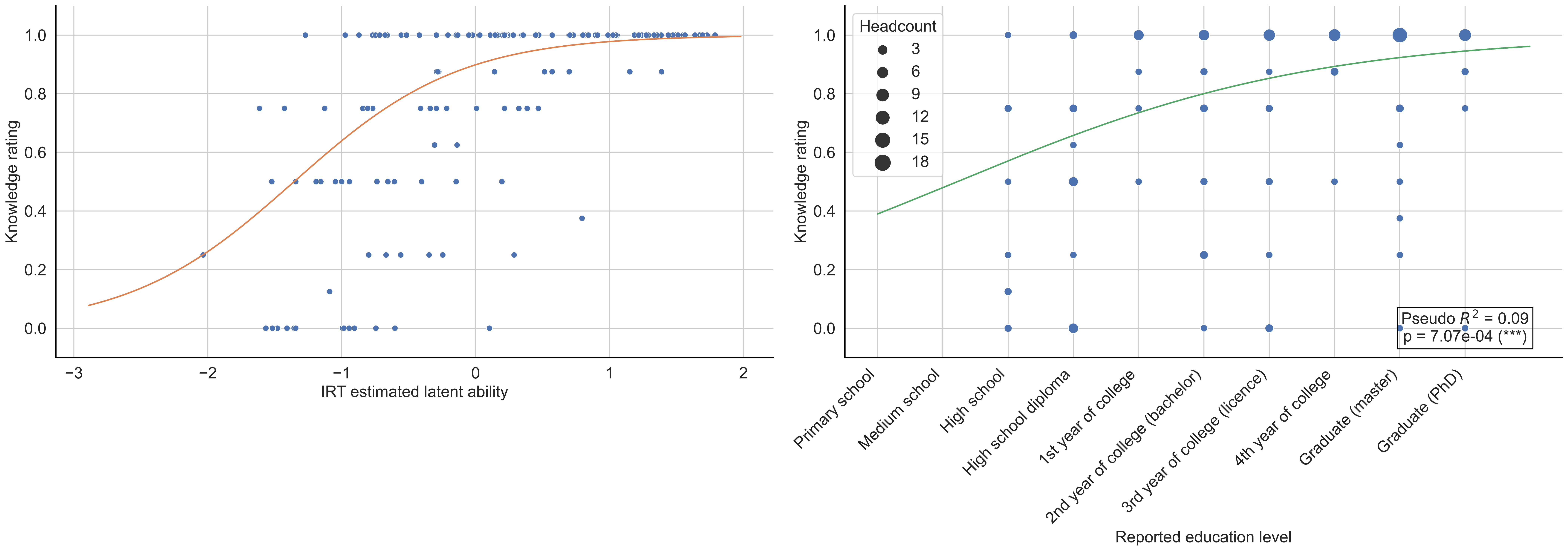

Word: morphisme ("morphism"; Bachelor) -- n = 126

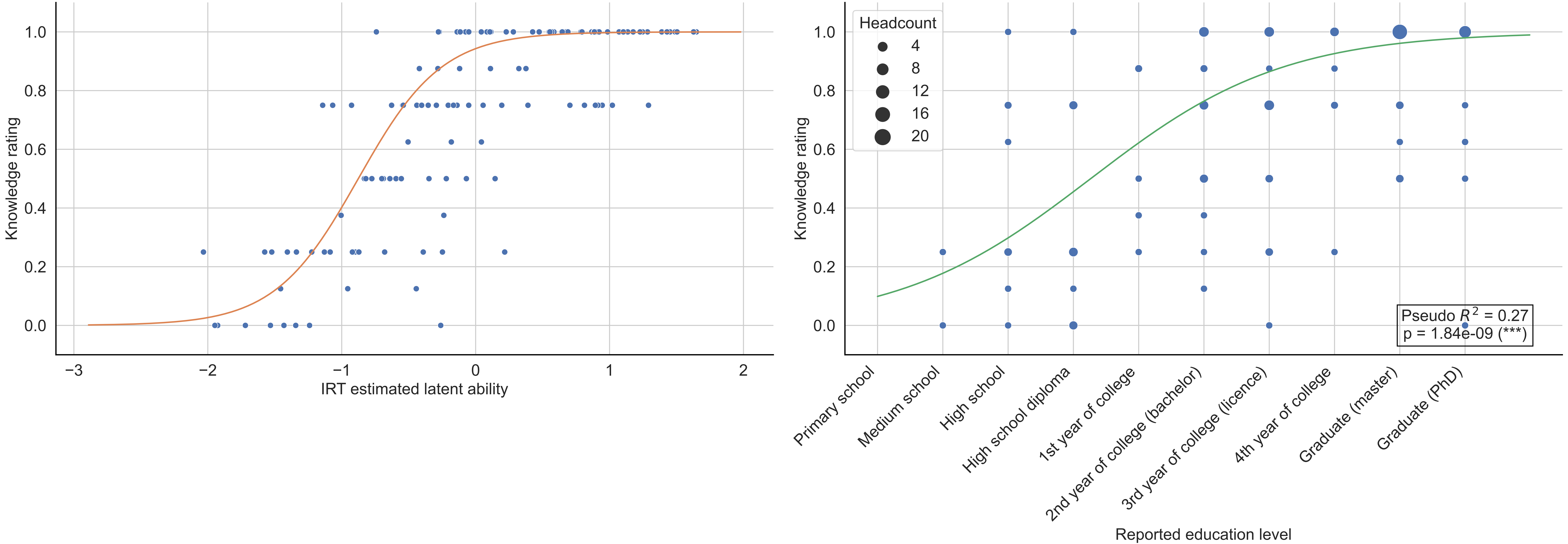

Word: médiatrice ("perpendicular bisector"; Primary school) -- n = 134

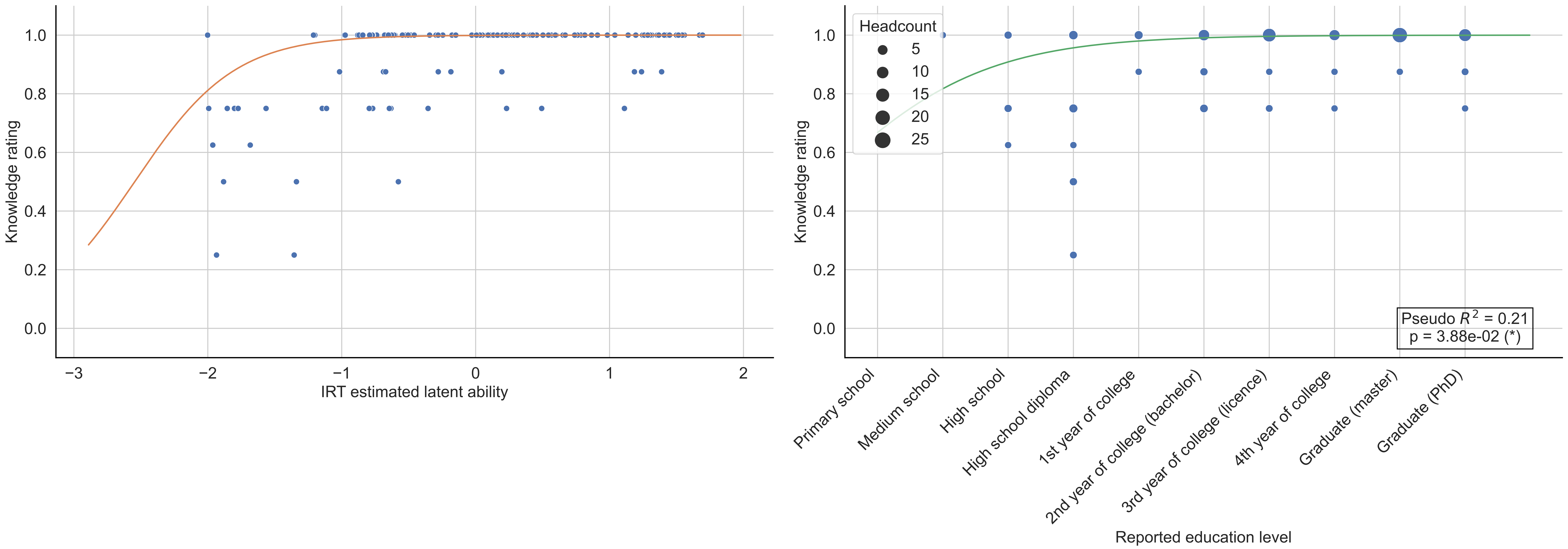

Word: nappe ("nappe"; Licence) -- n = 148

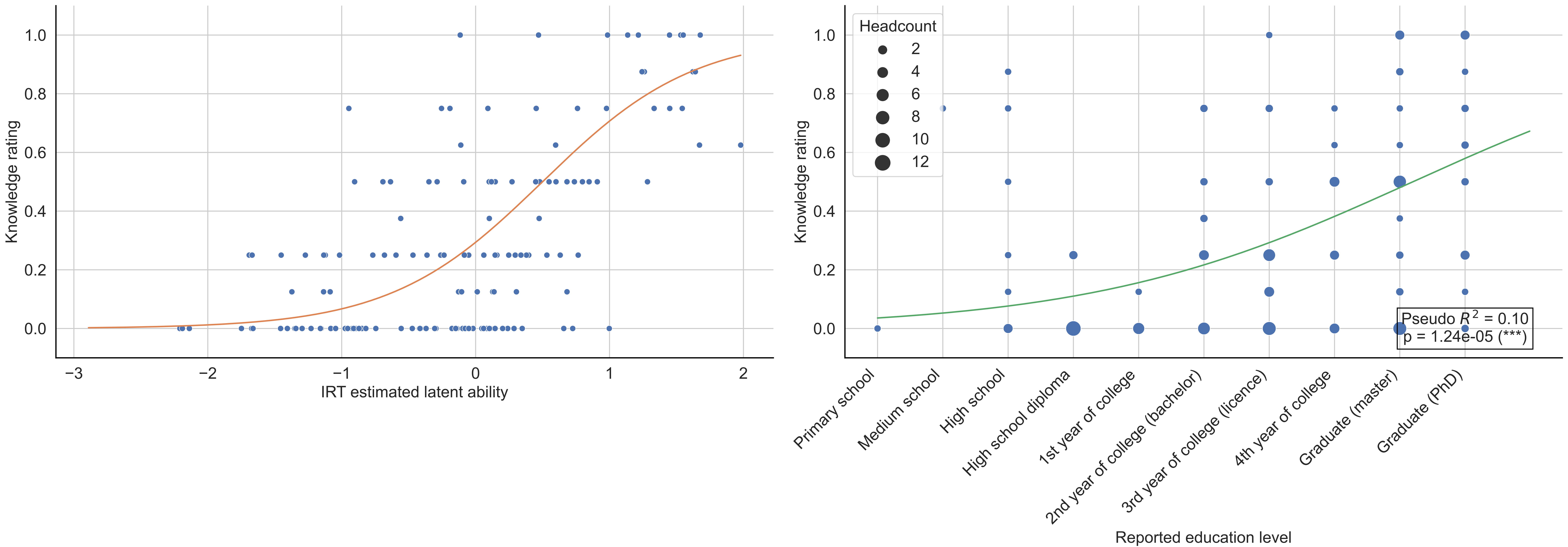

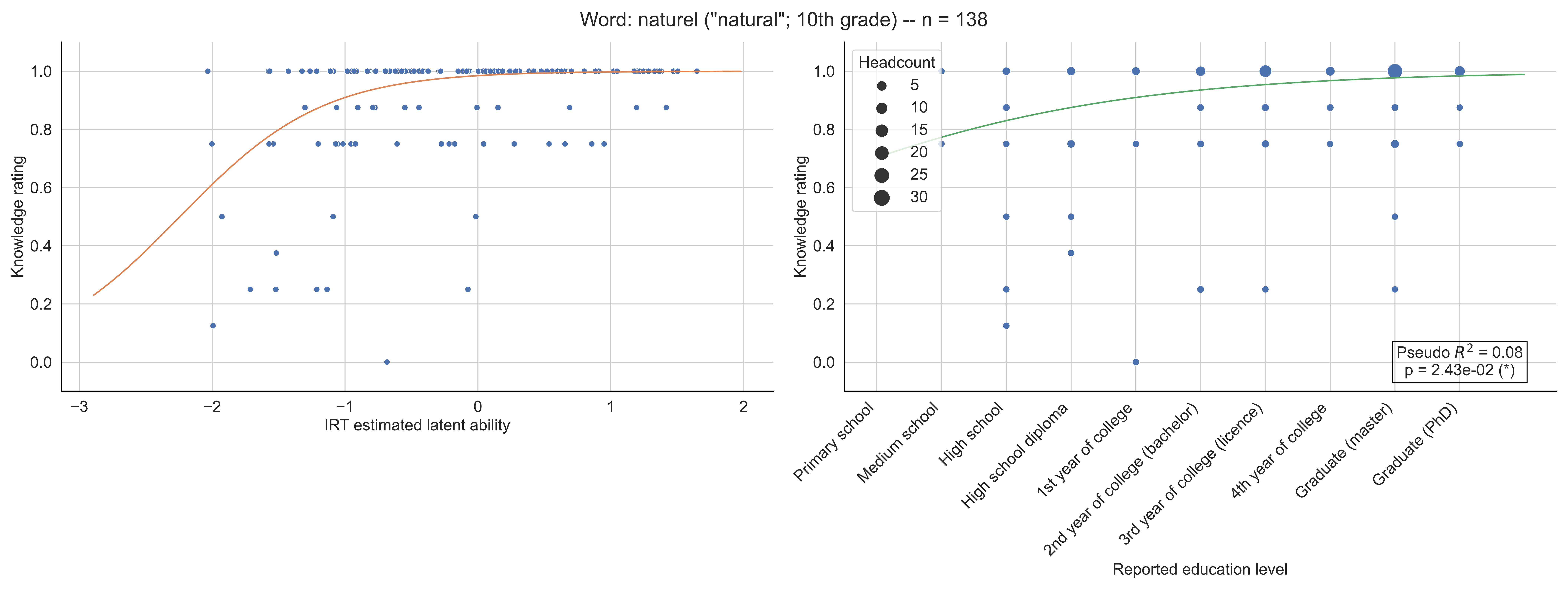

Word: naturel ("natural"; 10th grade) -- n = 138
Knowledge rating
IRT estimated latent ability
Headcount
5
10
15
20
25
30
Pseudo $R^2$ = 0.08
p = 2.43e-02 (*)
Primary school
Medium school
High school
High school diploma
1st year of college
2nd year of college (bachelor)
3rd year of college (licence)
4th year of college
Graduate (master)
Graduate (PhD)
Reported education level

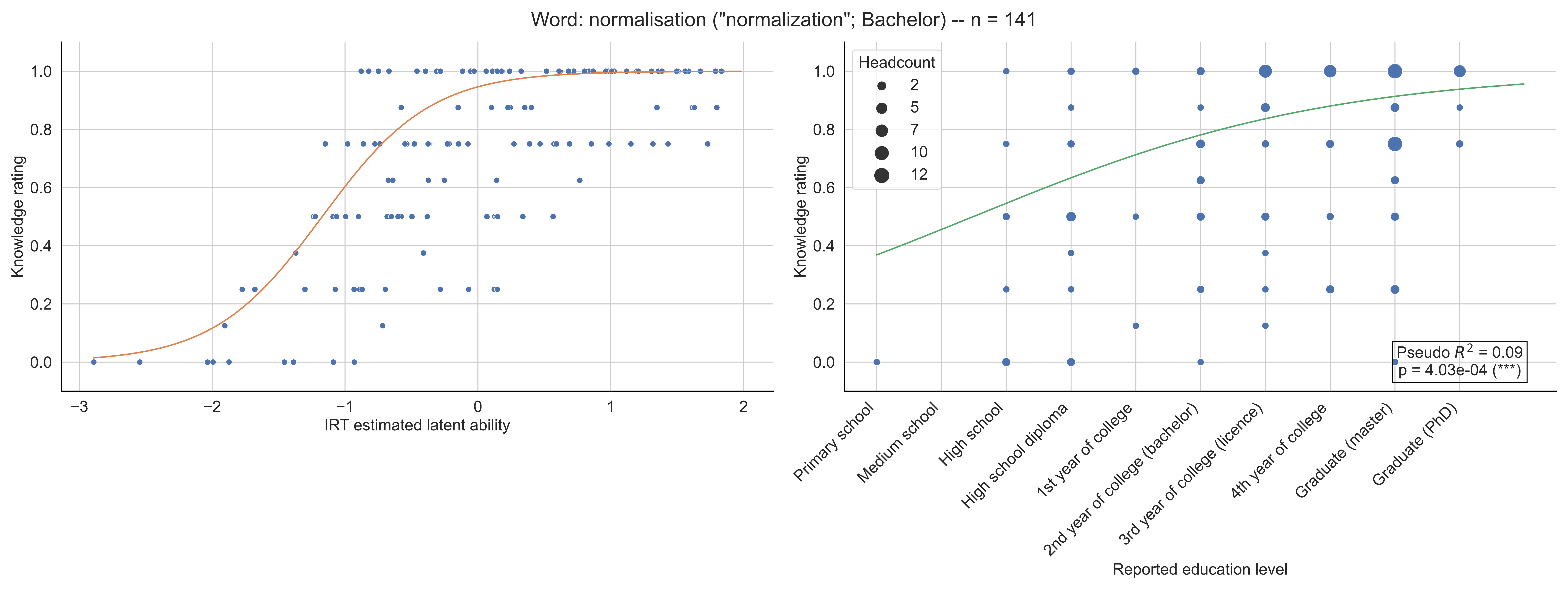

Word: normalisation ("normalization"; Bachelor) -- n = 141
Knowledge rating
IRT estimated latent ability
Headcount
2
5
7
10
12
Pseudo $R^2$ = 0.09
p = 4.03e-04 (***)
Primary school
Medium school
High school
High school diploma
1st year of college
2nd year of college (bachelor)
3rd year of college (licence)
4th year of college
Graduate (master)
Graduate (PhD)
Reported education level

Word: norme ("norm"; Bachelor) -- n = 148

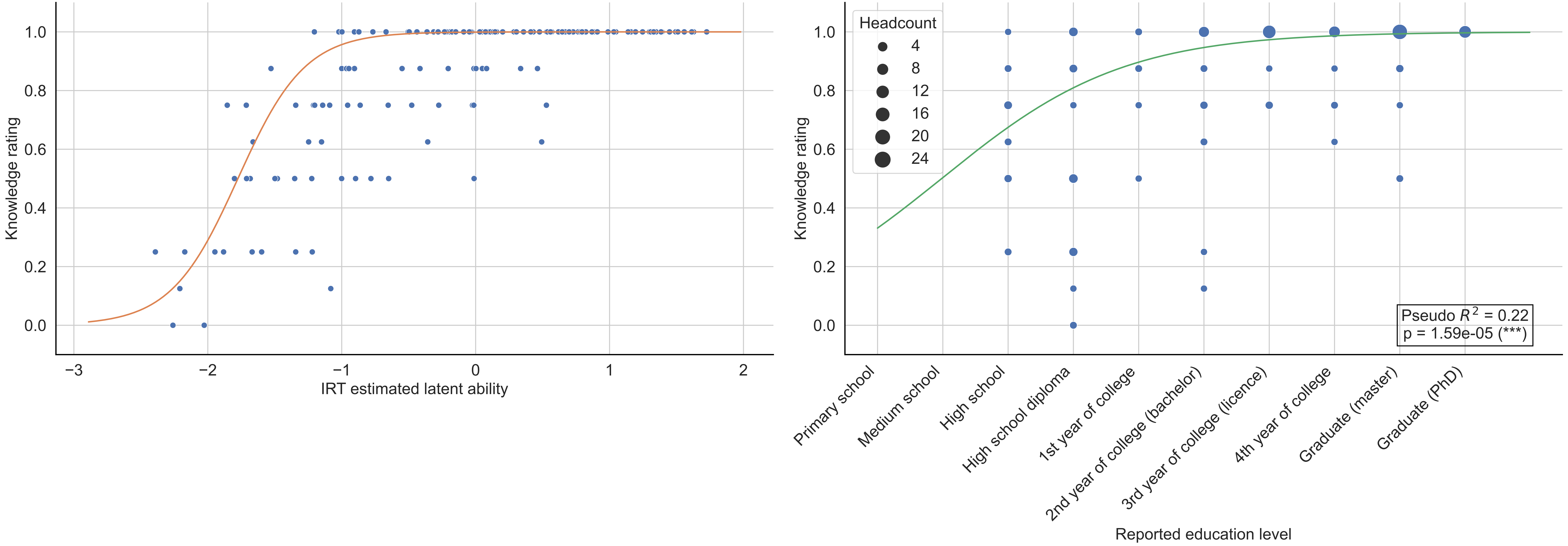

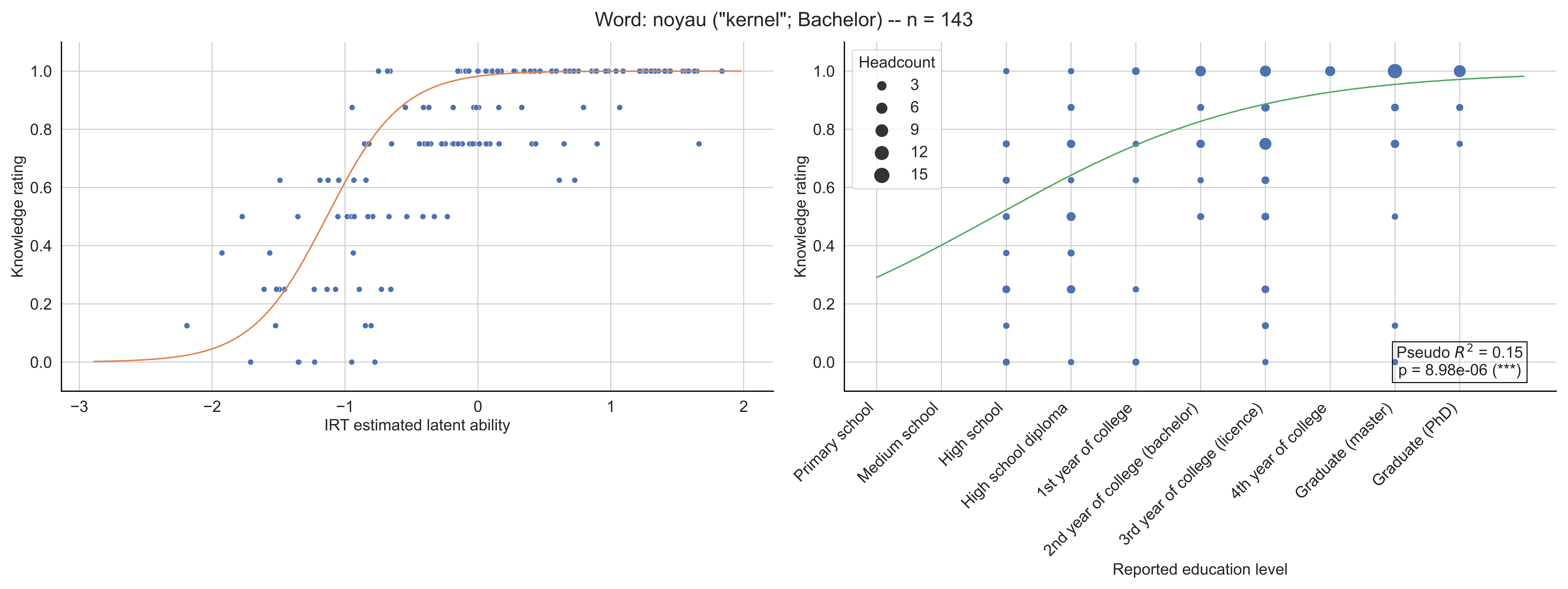

Word: noyau ("kernel"; Bachelor) -- n = 143
Knowledge rating
IRT estimated latent ability
Headcount
3
6
9
12
15
Pseudo $R^2$ = 0.15
p = 8.98e-06 (***)
Primary school
Medium school
High school
High school diploma
1st year of college
2nd year of college (bachelor)
3rd year of college (licence)
4th year of college
Graduate (master)
Graduate (PhD)
Reported education level

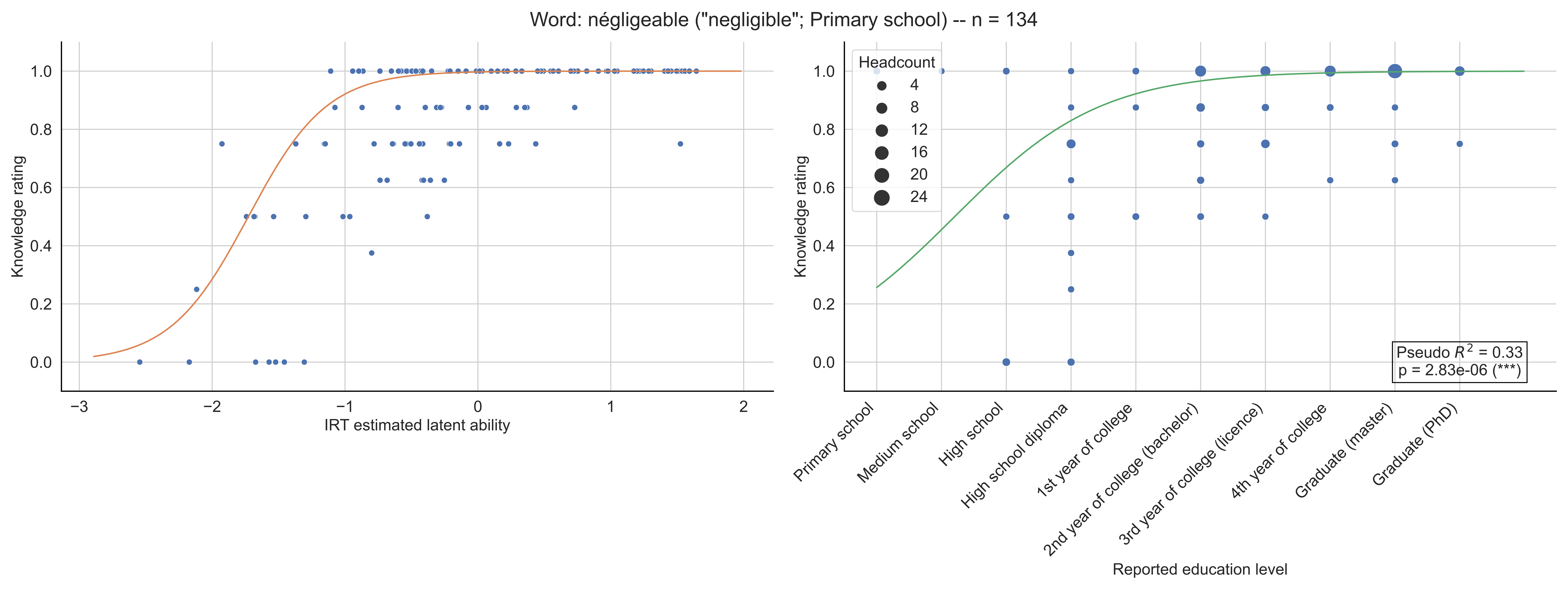

Word: négligeable ("negligible"; Primary school) -- n = 134
Knowledge rating
IRT estimated latent ability
Headcount
4
8
12
16
20
24
Pseudo $R^2$ = 0.33
p = 2.83e-06 (***)
Primary school
Medium school
High school
High school diploma
1st year of college
2nd year of college (bachelor)
3rd year of college (licence)
4th year of college
Graduate (master)
Graduate (PhD)
Reported education level

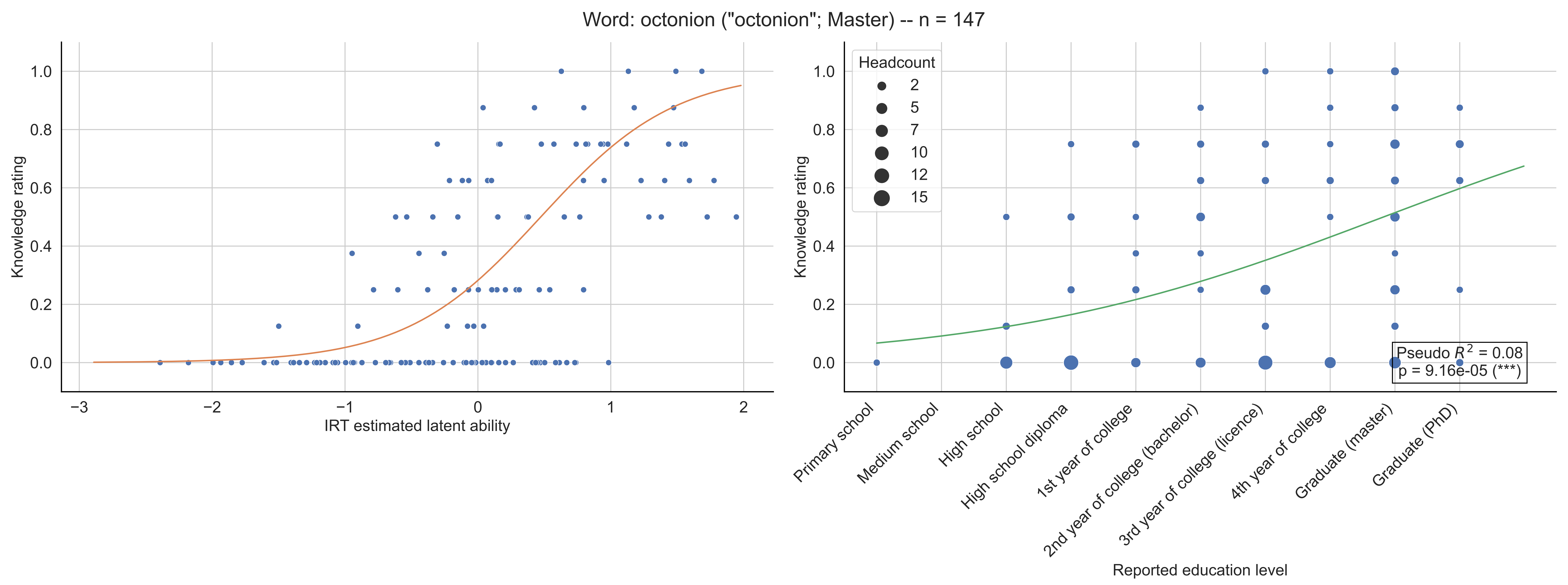
Word: octonion ("octonion"; Master) -- n = 147
Knowledge rating
IRT estimated latent ability
Headcount
2
5
7
10
12
15
Pseudo $R^2$ = 0.08
p = 9.16e-05 (***)
Primary school
Medium school
High school
High school diploma
1st year of college
2nd year of college (bachelor)
3rd year of college (licence)
4th year of college
Graduate (master)
Graduate (PhD)
Reported education level

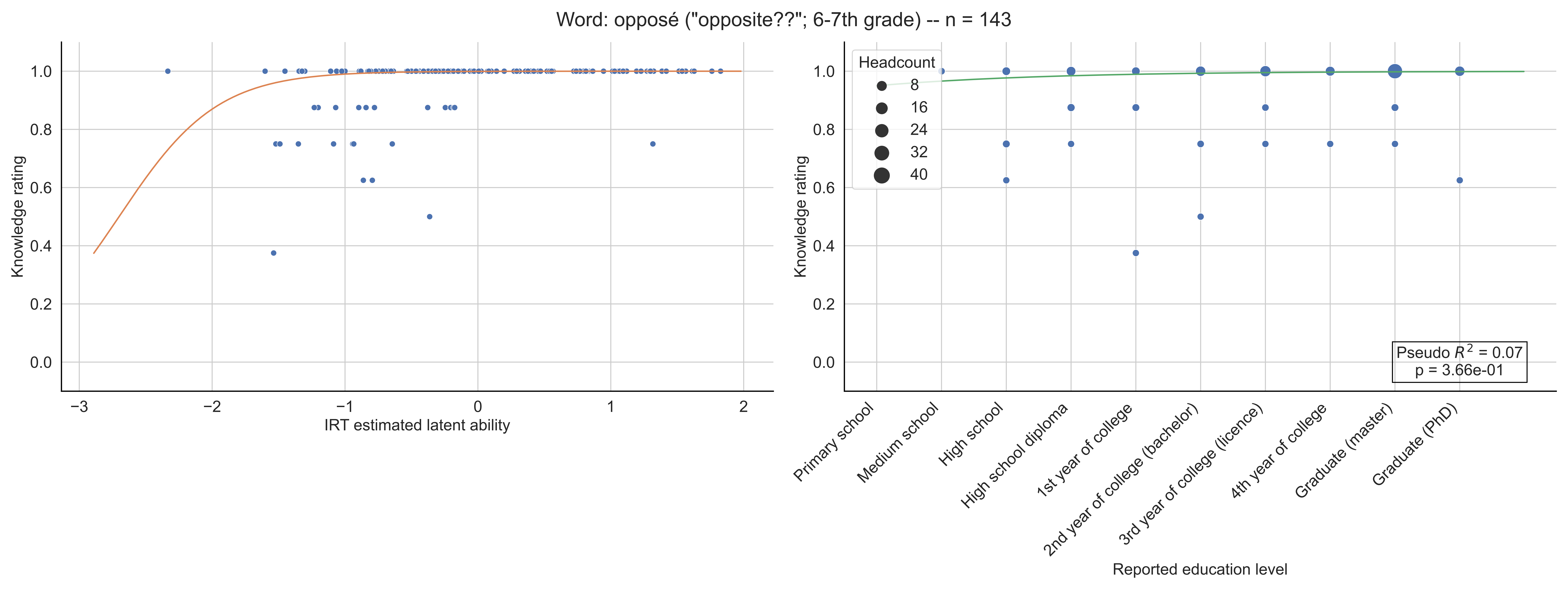
Word: opposé ("opposite??"; 6-7th grade) -- n = 143
Knowledge rating
IRT estimated latent ability
Headcount
8
16
24
32
40
Pseudo $R^2$ = 0.07
p = 3.66e-01
Primary school
Medium school
High school
High school diploma
1st year of college
2nd year of college (bachelor)
3rd year of college (licence)
4th year of college
Graduate (master)
Graduate (PhD)
Reported education level

Word: opérateur ("operator"; Bachelor) -- n = 169

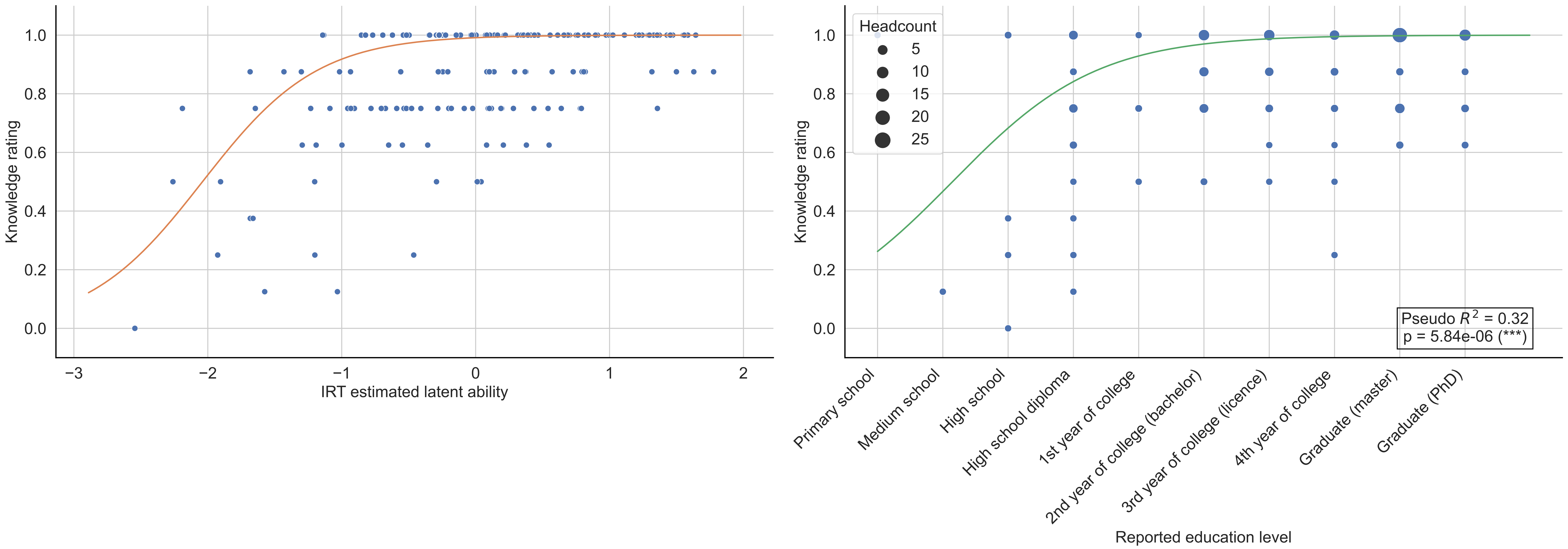

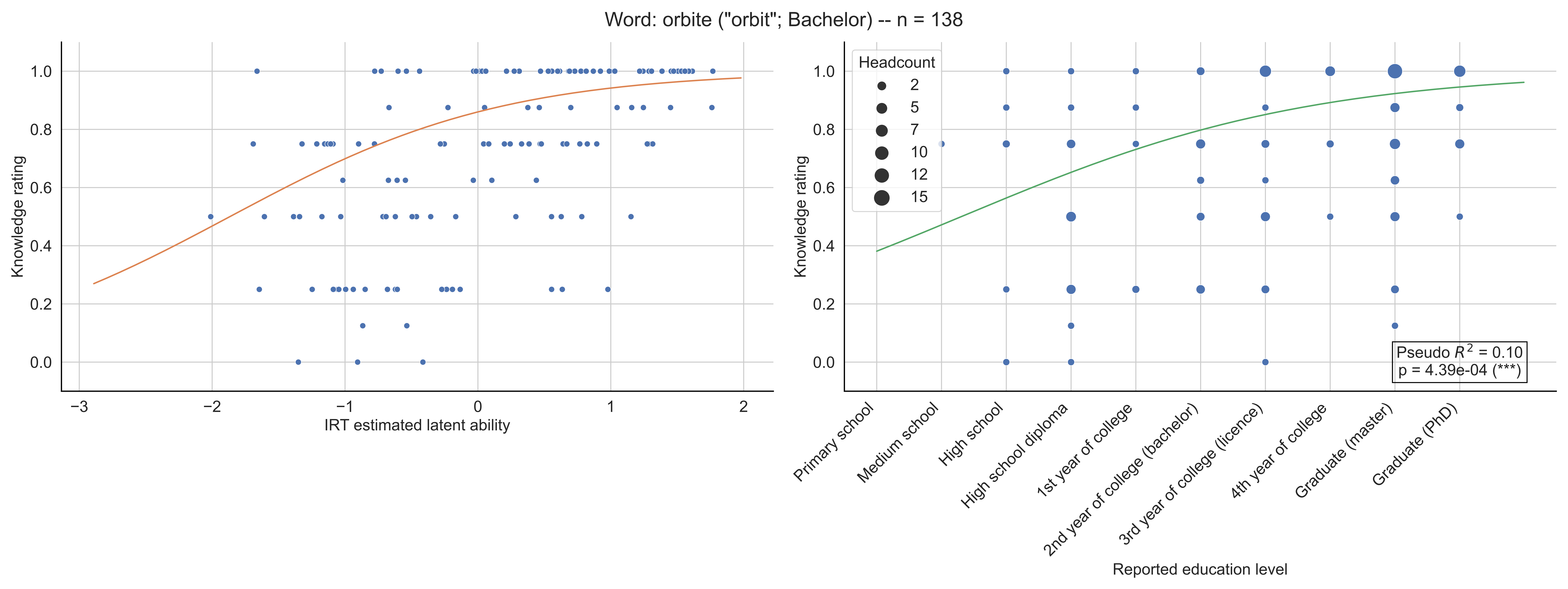

Word: orbite ("orbit"; Bachelor) -- n = 138
Knowledge rating
IRT estimated latent ability
Headcount
2
5
7
10
12
15
Pseudo $R^2$ = 0.10
p = 4.39e-04 (***)
Primary school
Medium school
High school
High school diploma
1st year of college
2nd year of college (bachelor)
3rd year of college (licence)
4th year of college
Graduate (master)
Graduate (PhD)
Reported education level

Word: ordre ("order"; Primary school) -- n = 151

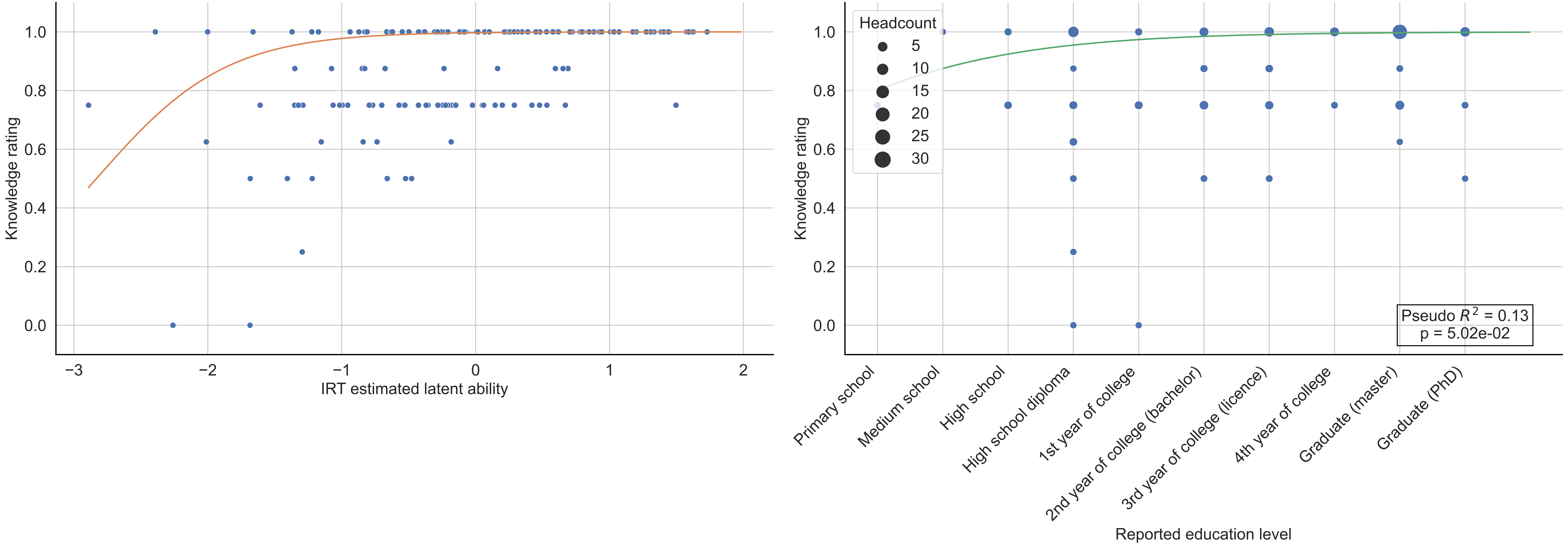

Word: origine ("origin"; 6-7th grade) -- n = 122

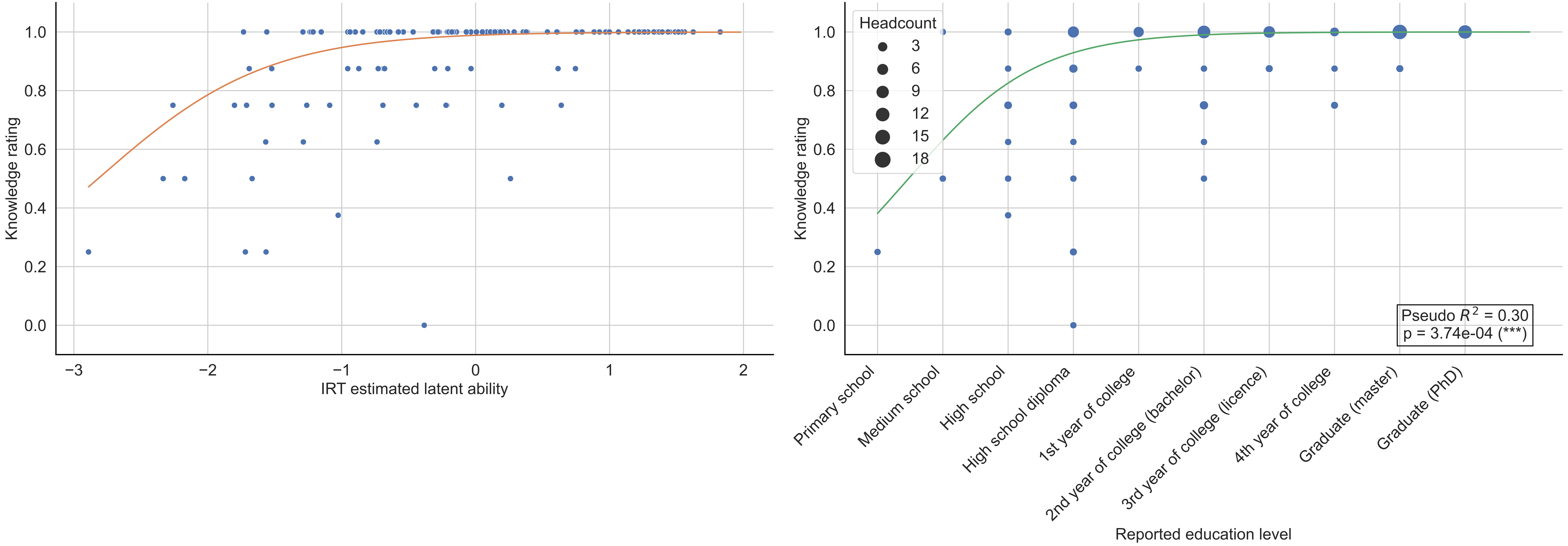

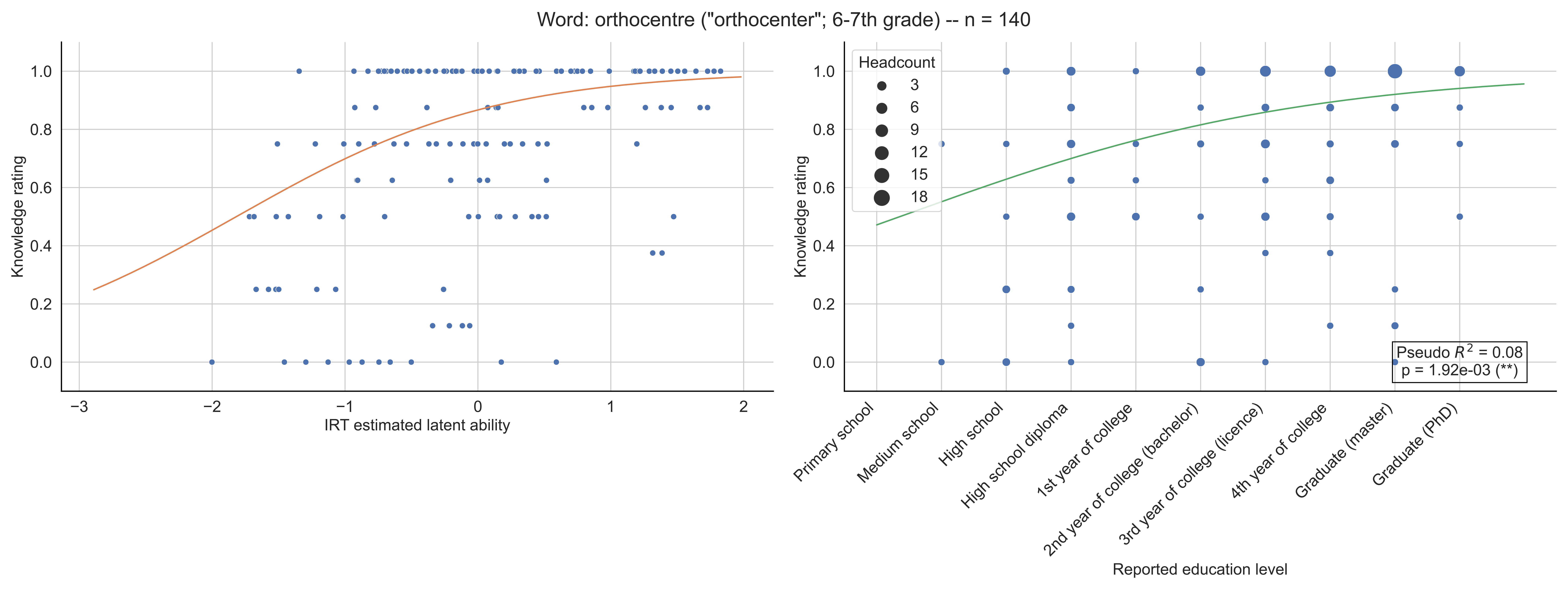

Word: orthocentre ("orthocenter"; 6-7th grade) -- n = 140
Knowledge rating
IRT estimated latent ability
Headcount
3
6
9
12
15
18
Pseudo $R^2$ = 0.08
p = 1.92e-03 (**)
Primary school
Medium school
High school
High school diploma
1st year of college
2nd year of college (bachelor)
3rd year of college (licence)
4th year of college
Graduate (master)
Graduate (PhD)
Reported education level

Word: orthogonalité ("orthogonality"; 11-12th grade) -- n = 122

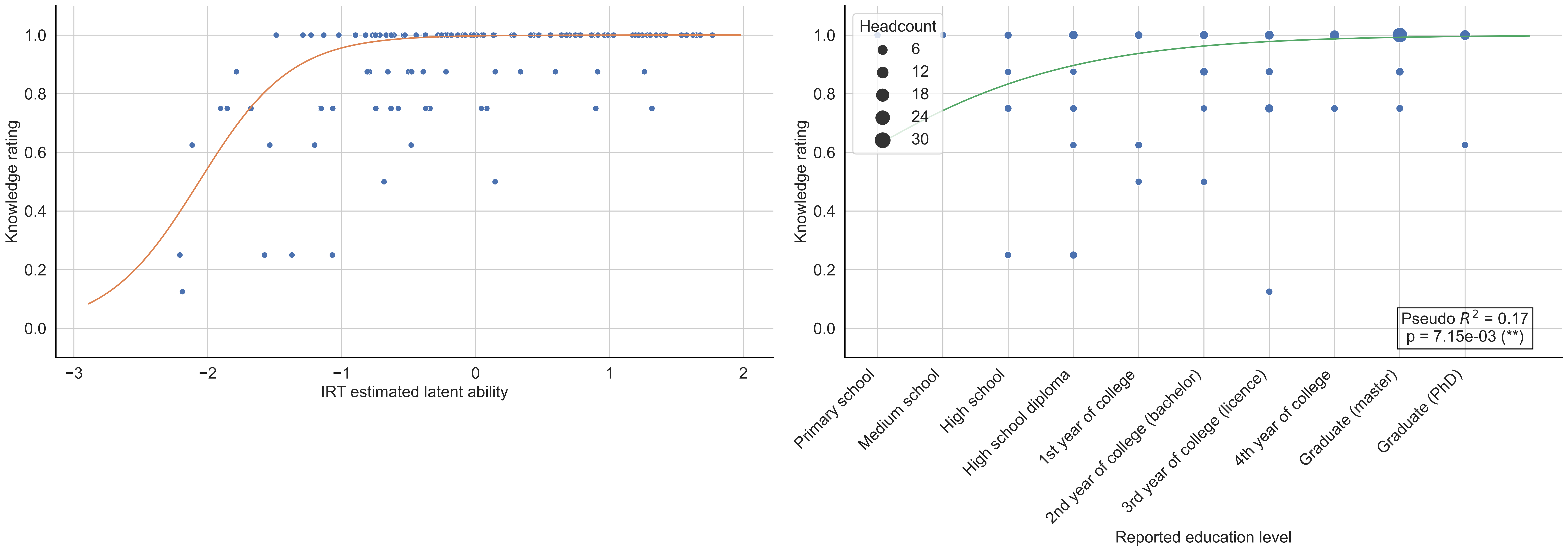

Word: ouvert ("open"; Bachelor) -- n = 155

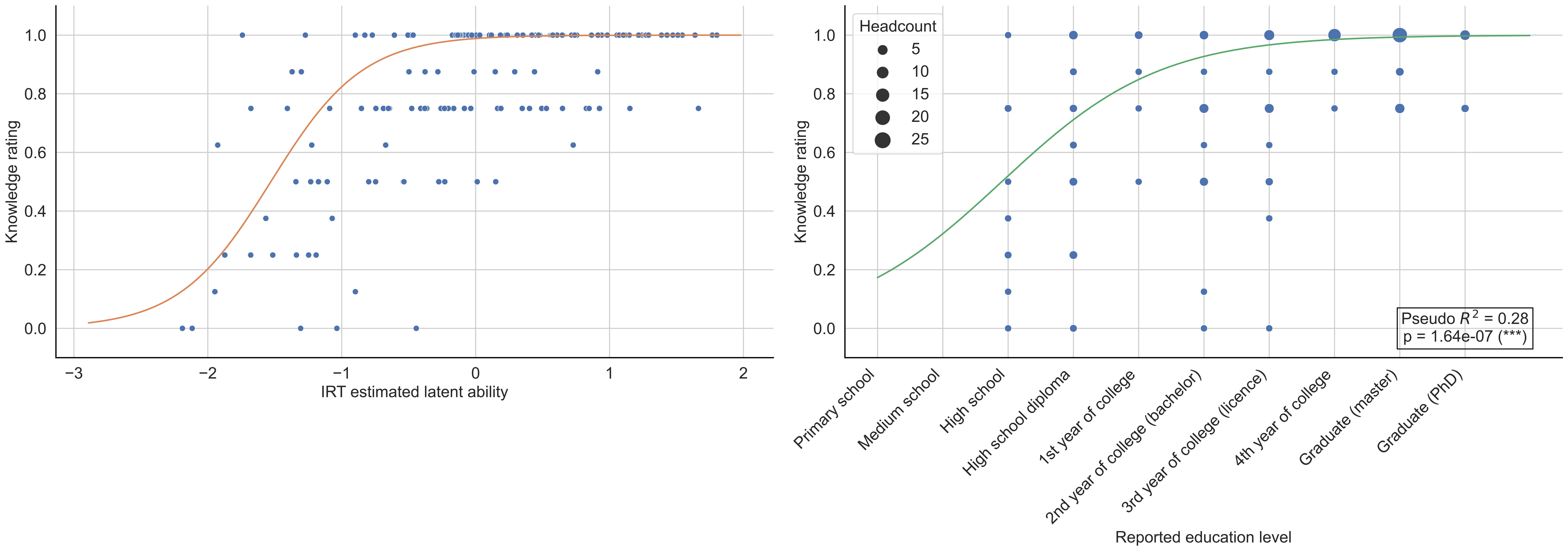

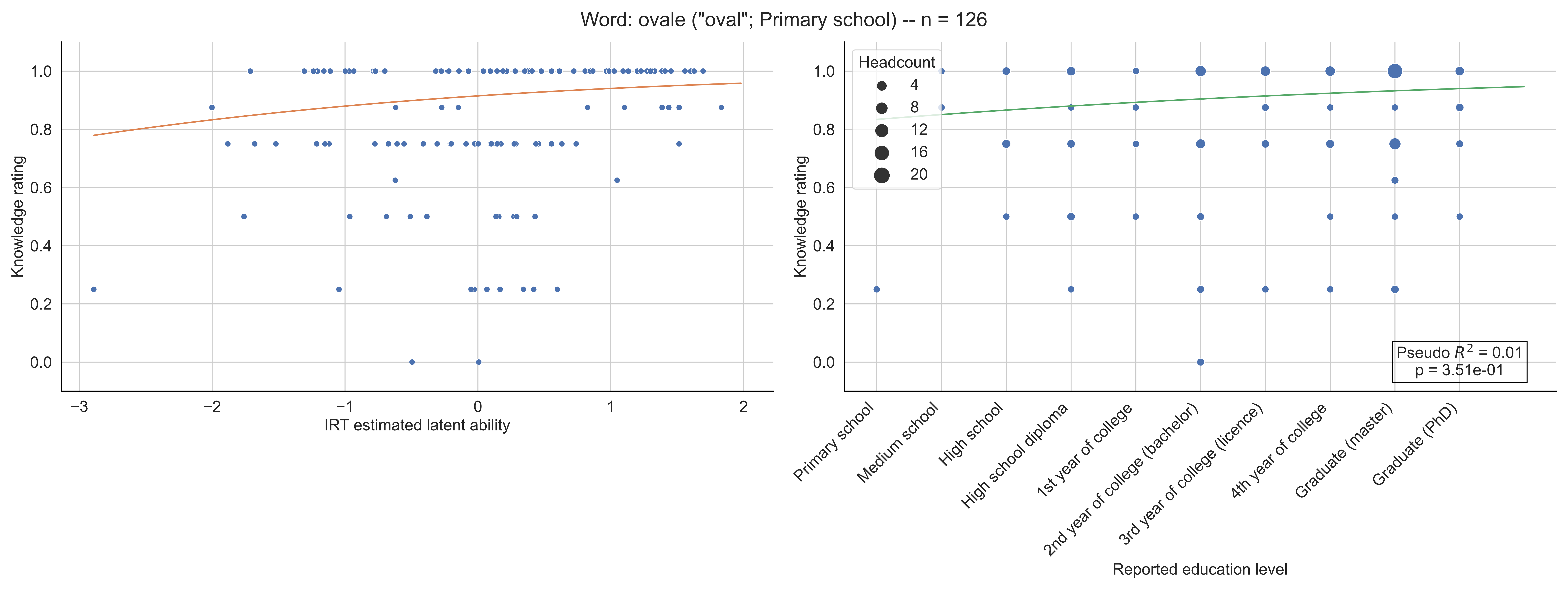

Word: ovale ("oval"; Primary school) -- n = 126
Knowledge rating
IRT estimated latent ability
Headcount
4
8
12
16
20
Pseudo $R^2$ = 0.01
p = 3.51e-01
Primary school
Medium school
High school
High school diploma
1st year of college
2nd year of college (bachelor)
3rd year of college (licence)
4th year of college
Graduate (master)
Graduate (PhD)
Reported education level

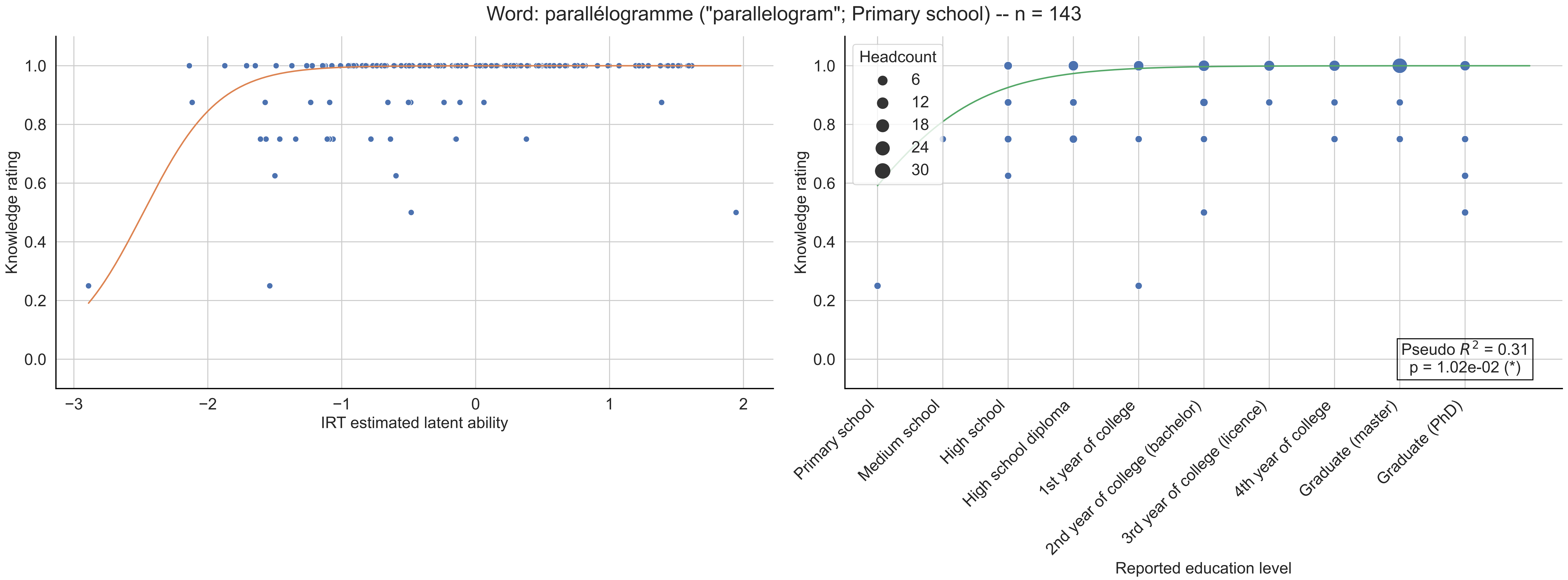

Word: parallélogramme ("parallelogram"; Primary school) -- n = 143
Knowledge rating
IRT estimated latent ability
Headcount
6
12
18
24
30
Pseudo $R^2$ = 0.31
p = 1.02e-02 (*)
Primary school
Medium school
High school
High school diploma
1st year of college
2nd year of college (bachelor)
3rd year of college (licence)
4th year of college
Graduate (master)
Graduate (PhD)
Reported education level

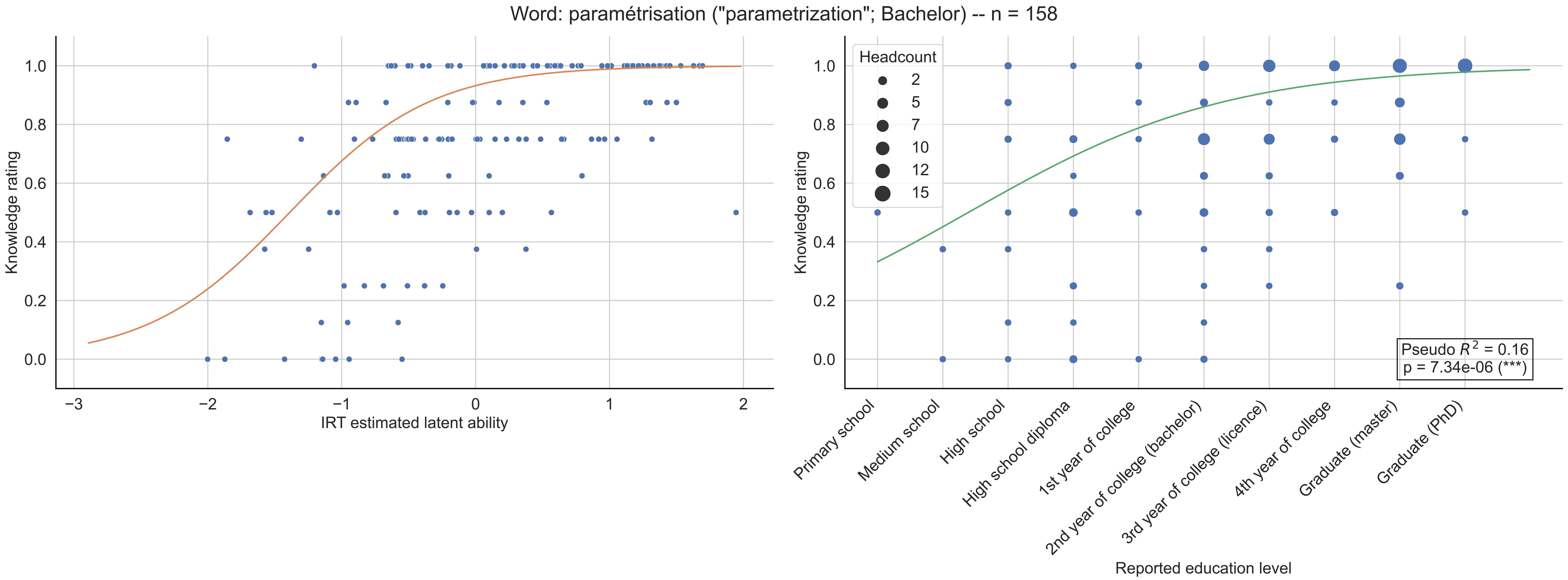

Word: paramétrisation ("parametrization"; Bachelor) -- n = 158
Knowledge rating
IRT estimated latent ability
Headcount
2
5
7
10
12
15
Pseudo $R^2$ = 0.16
p = 7.34e-06 (***)
Primary school
Medium school
High school
High school diploma
1st year of college
2nd year of college (bachelor)
3rd year of college (licence)
4th year of college
Graduate (master)
Graduate (PhD)
Reported education level

Word: partie ("subset"; 10th grade) -- n = 128

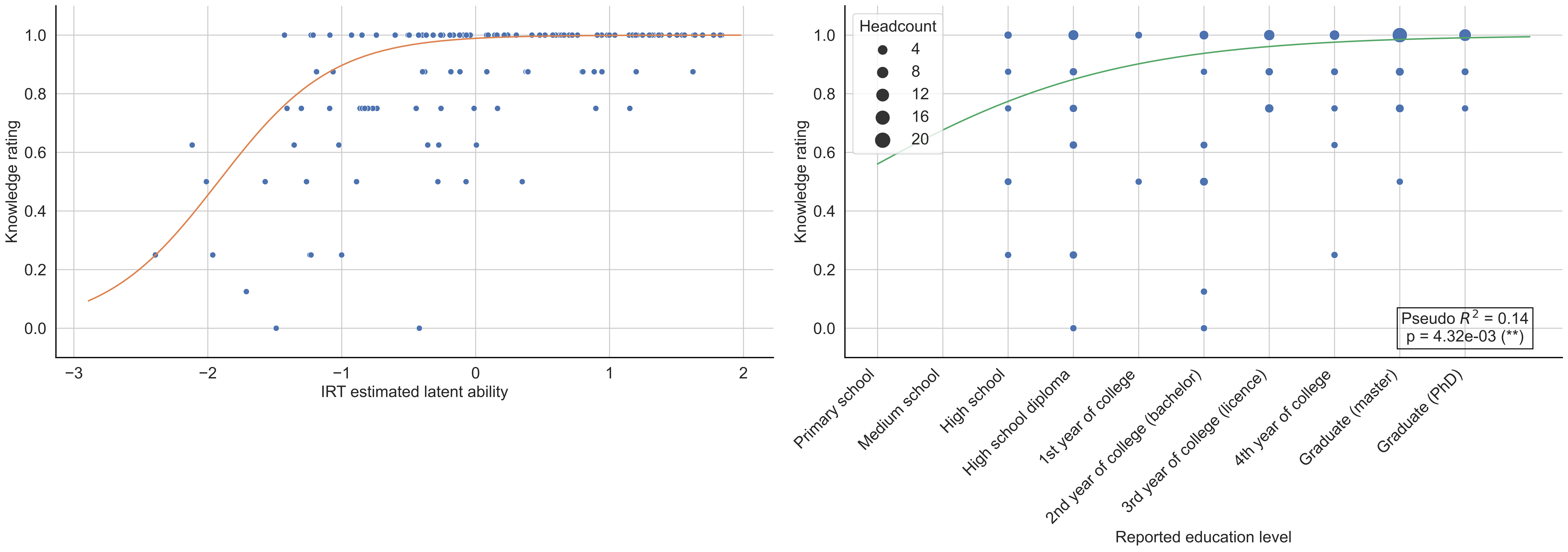

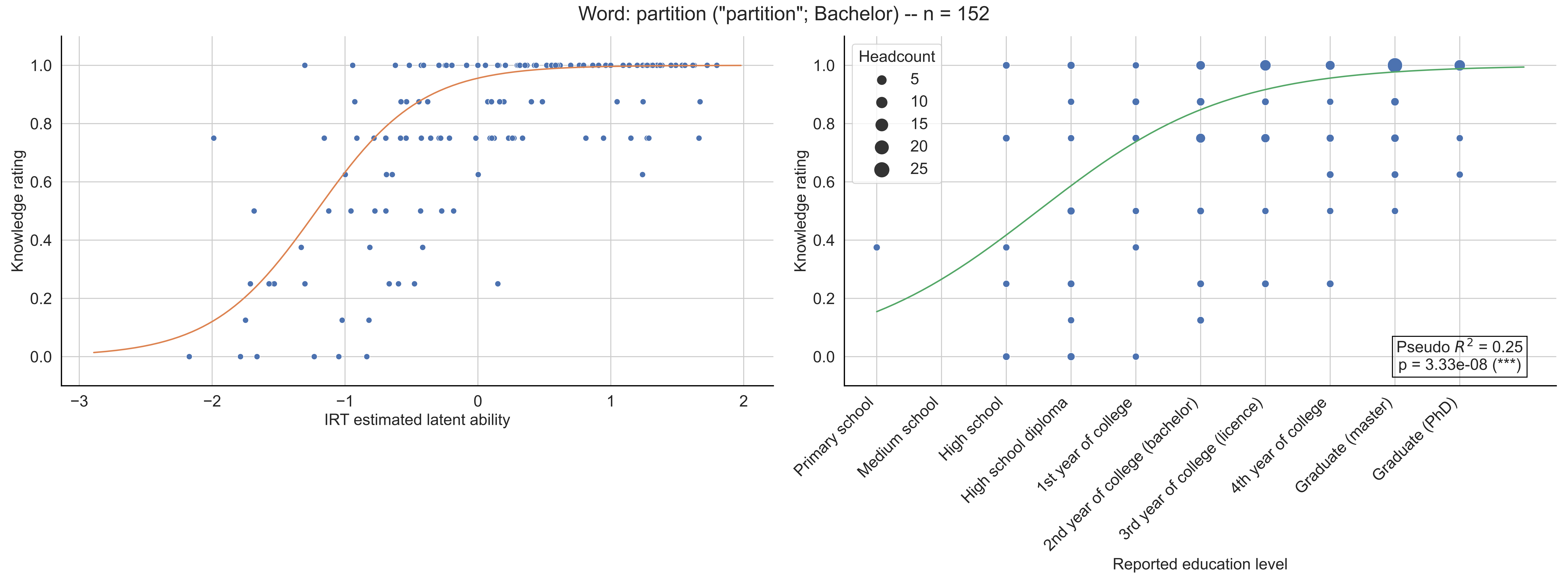

Word: partition ("partition"; Bachelor) -- n = 152
Knowledge rating
IRT estimated latent ability
Headcount
5
10
15
20
25
Pseudo $R^2$ = 0.25
p = 3.33e-08 (***)
Primary school
Medium school
High school
High school diploma
1st year of college
2nd year of college (bachelor)
3rd year of college (licence)
4th year of college
Graduate (master)
Graduate (PhD)
Reported education level

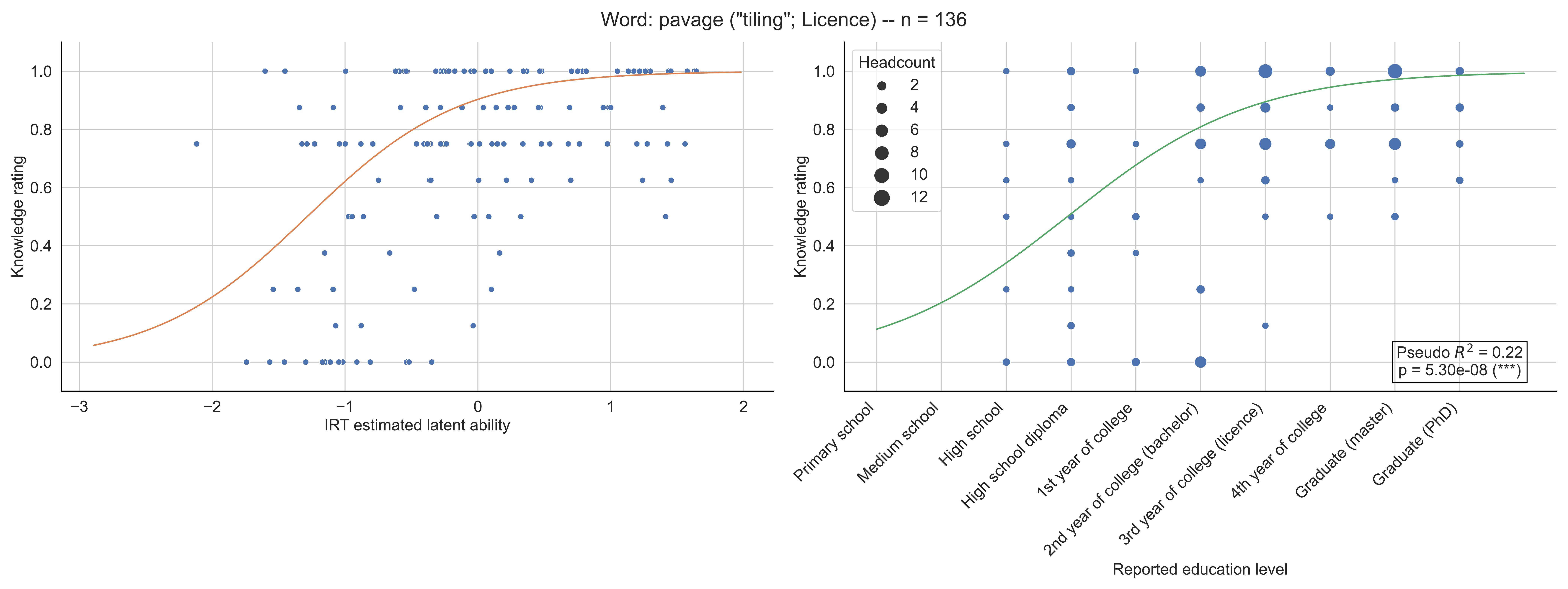

Word: pavage ("tiling"; Licence) -- n = 136
Knowledge rating
IRT estimated latent ability
Headcount
2
4
6
8
10
12
Primary school
Medium school
High school
High school diploma
1st year of college
2nd year of college (bachelor)
3rd year of college (licence)
4th year of college
Graduate (master)
Graduate (PhD)
Reported education level
Pseudo $R^2$ = 0.22
p = 5.30e-08 (***)

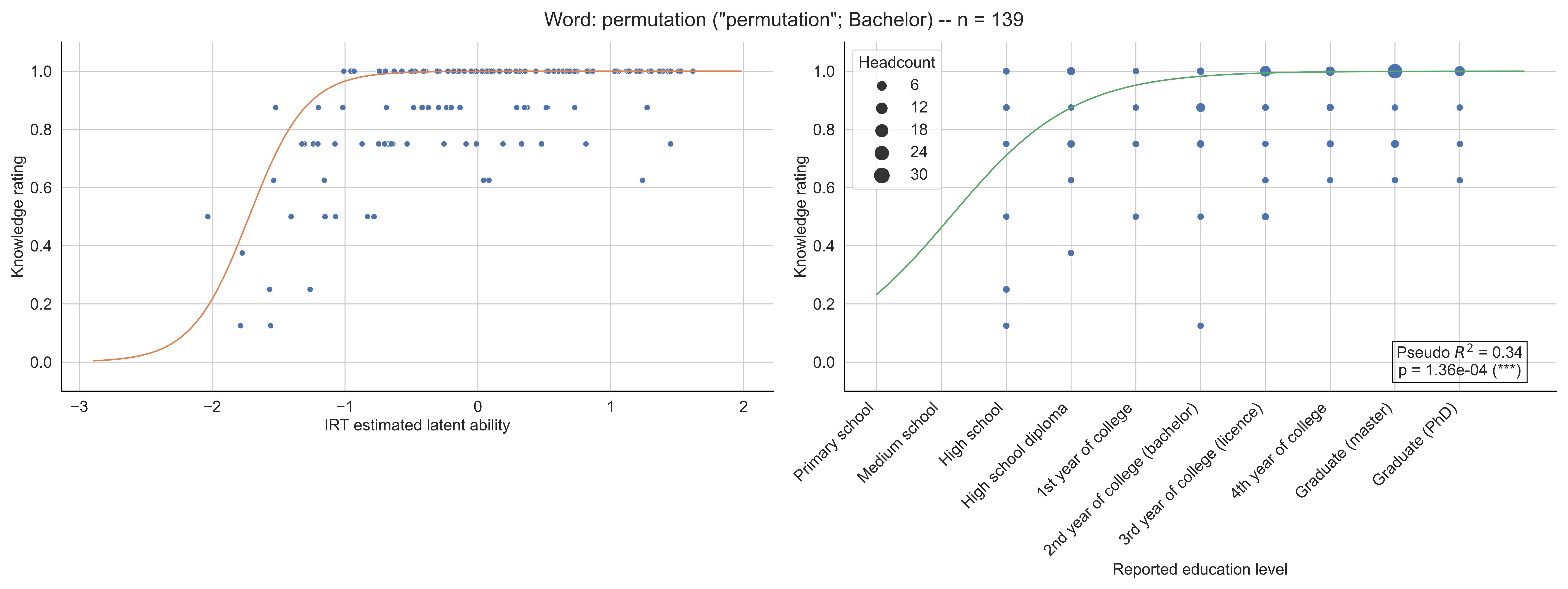

Word: permutation ("permutation"; Bachelor) -- n = 139
Knowledge rating
IRT estimated latent ability
Headcount
6
12
18
24
30
Pseudo $R^2$ = 0.34
p = 1.36e-04 (***)
Primary school
Medium school
High school
High school diploma
1st year of college
2nd year of college (bachelor)
3rd year of college (licence)
4th year of college
Graduate (master)
Graduate (PhD)
Reported education level

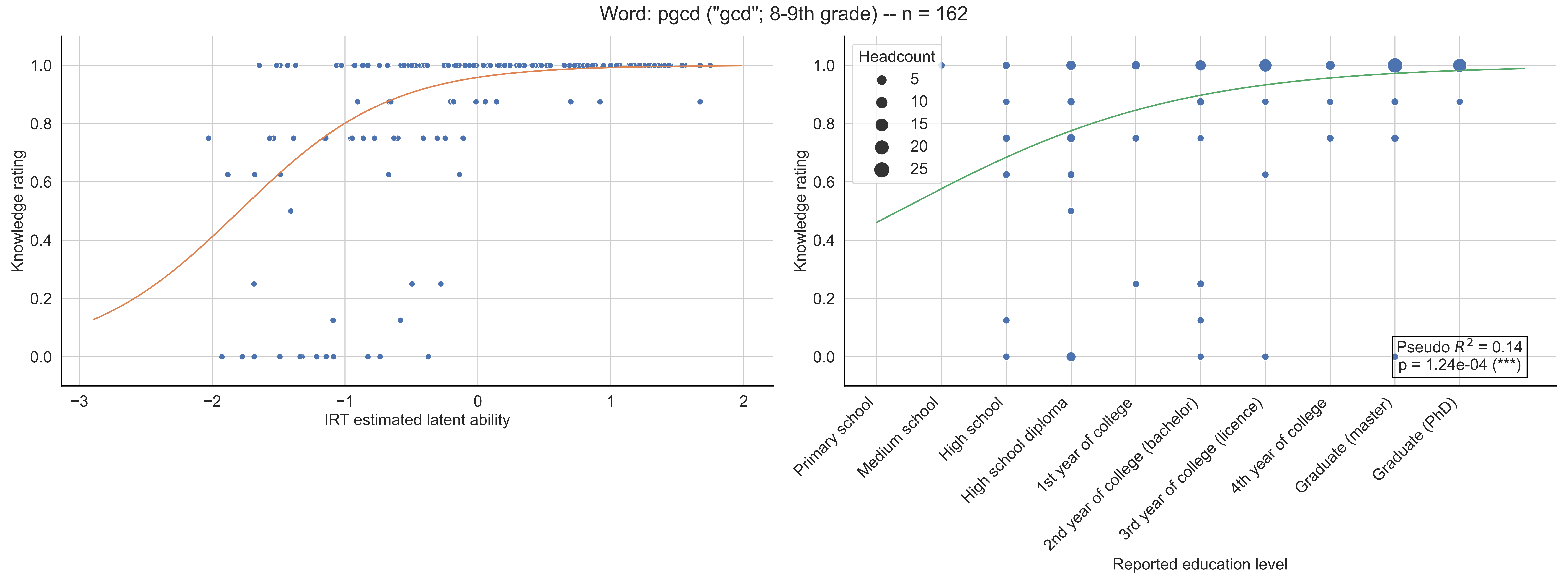

Word: pgcd ("gcd"; 8-9th grade) -- n = 162
Knowledge rating
IRT estimated latent ability
−3
−2
−1
0
1
2
0.0
0.2
0.4
0.6
0.8
1.0
Headcount
5
10
15
20
25
Pseudo $R^2$ = 0.14
p = 1.24e-04 (***)
Primary school
Medium school
High school
High school diploma
1st year of college
2nd year of college (bachelor)
3rd year of college (licence)
4th year of college
Graduate (master)
Graduate (PhD)
Reported education level

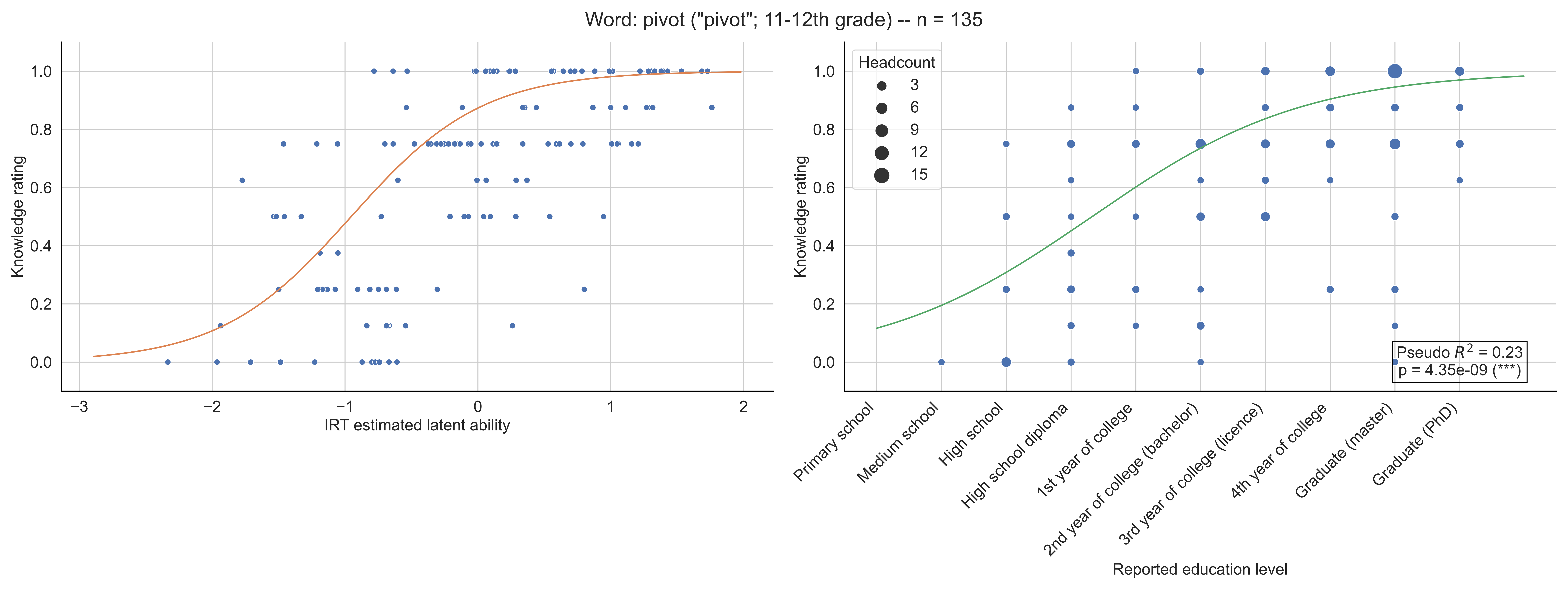

Word: pivot ("pivot"; 11-12th grade) -- n = 135
Knowledge rating
IRT estimated latent ability
Headcount
3
6
9
12
15
Pseudo $R^2$ = 0.23
p = 4.35e-09 (***)
Primary school
Medium school
High school
High school diploma
1st year of college
2nd year of college (bachelor)
3rd year of college (licence)
4th year of college
Graduate (master)
Graduate (PhD)
Reported education level

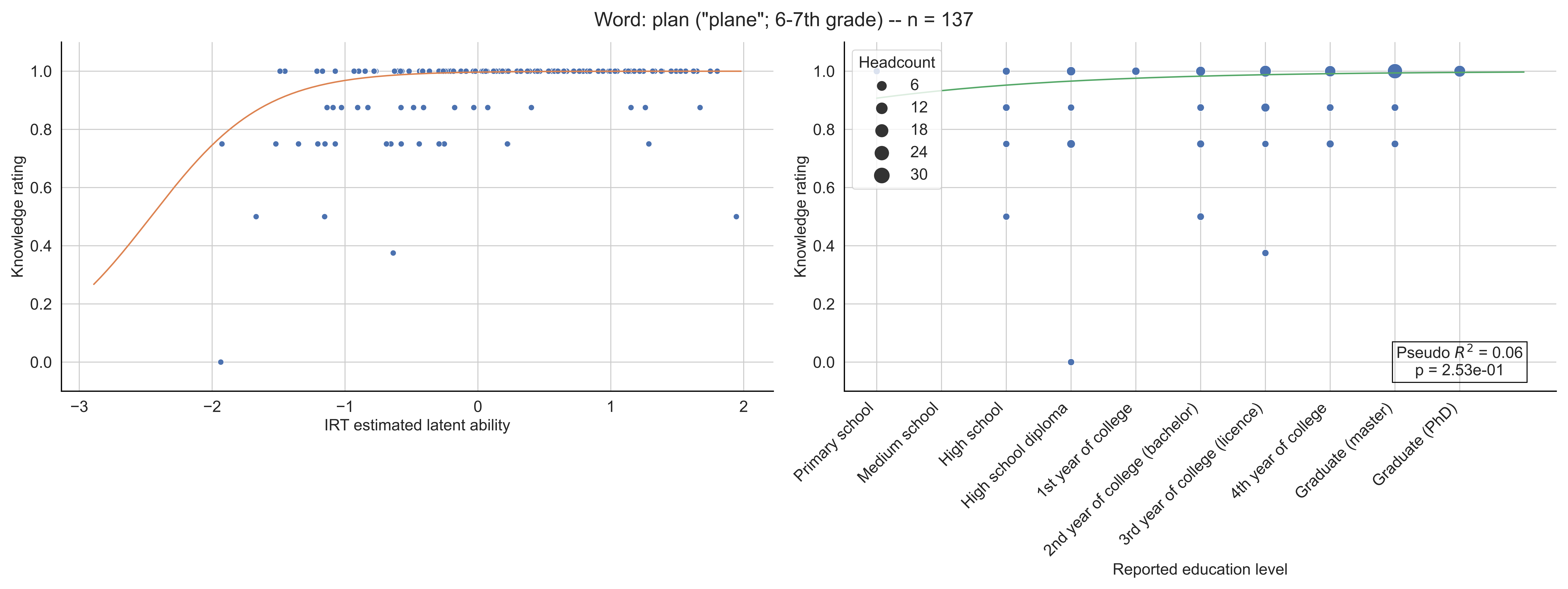
Word: plan ("plane"; 6-7th grade) -- n = 137
Knowledge rating
IRT estimated latent ability
Headcount
6
12
18
24
30
Pseudo $R^2$ = 0.06
p = 2.53e-01
Primary school
Medium school
High school
High school diploma
1st year of college
2nd year of college (bachelor)
3rd year of college (licence)
4th year of college
Graduate (master)
Graduate (PhD)
Reported education level

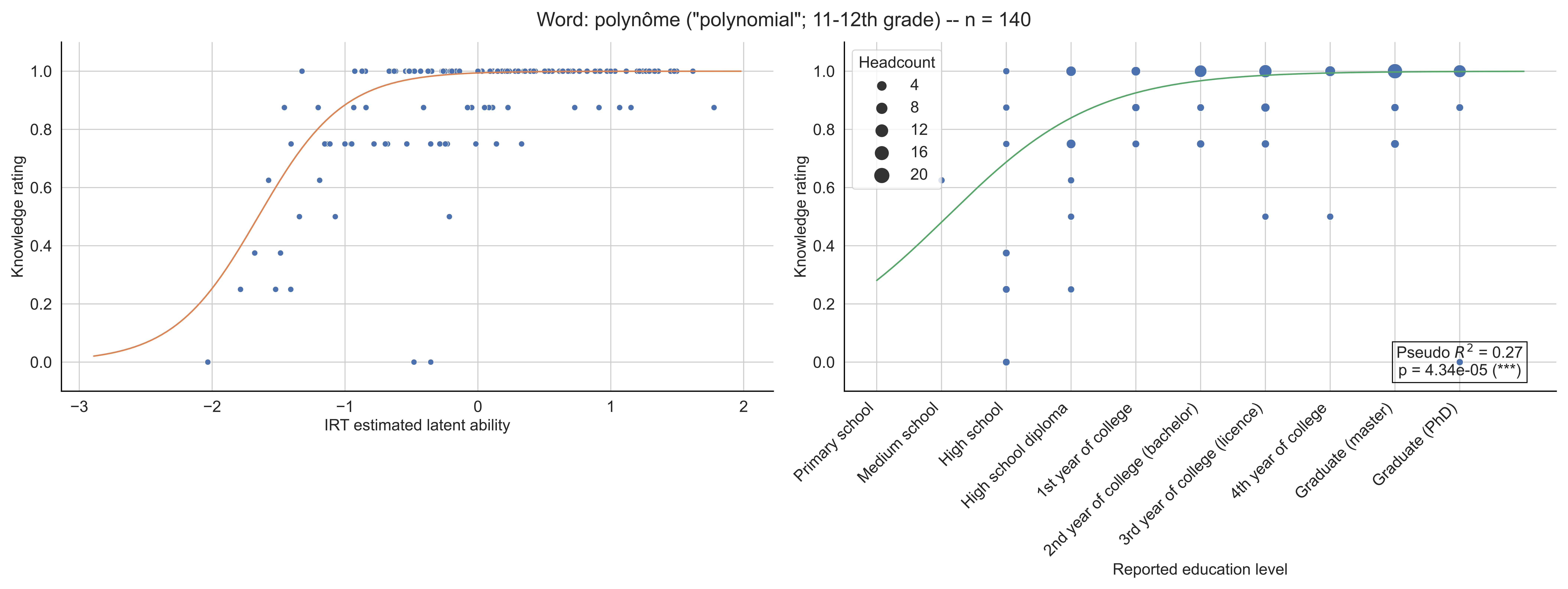
Word: polynôme ("polynomial"; 11-12th grade) -- n = 140
Knowledge rating
IRT estimated latent ability
−3
−2
−1
0
1
2
0.0
0.2
0.4
0.6
0.8
1.0
Headcount
4
8
12
16
20
Pseudo $R^2$ = 0.27
p = 4.34e-05 (***)
Primary school
Medium school
High school
High school diploma
1st year of college
2nd year of college (bachelor)
3rd year of college (licence)
4th year of college
Graduate (master)
Graduate (PhD)
Reported education level

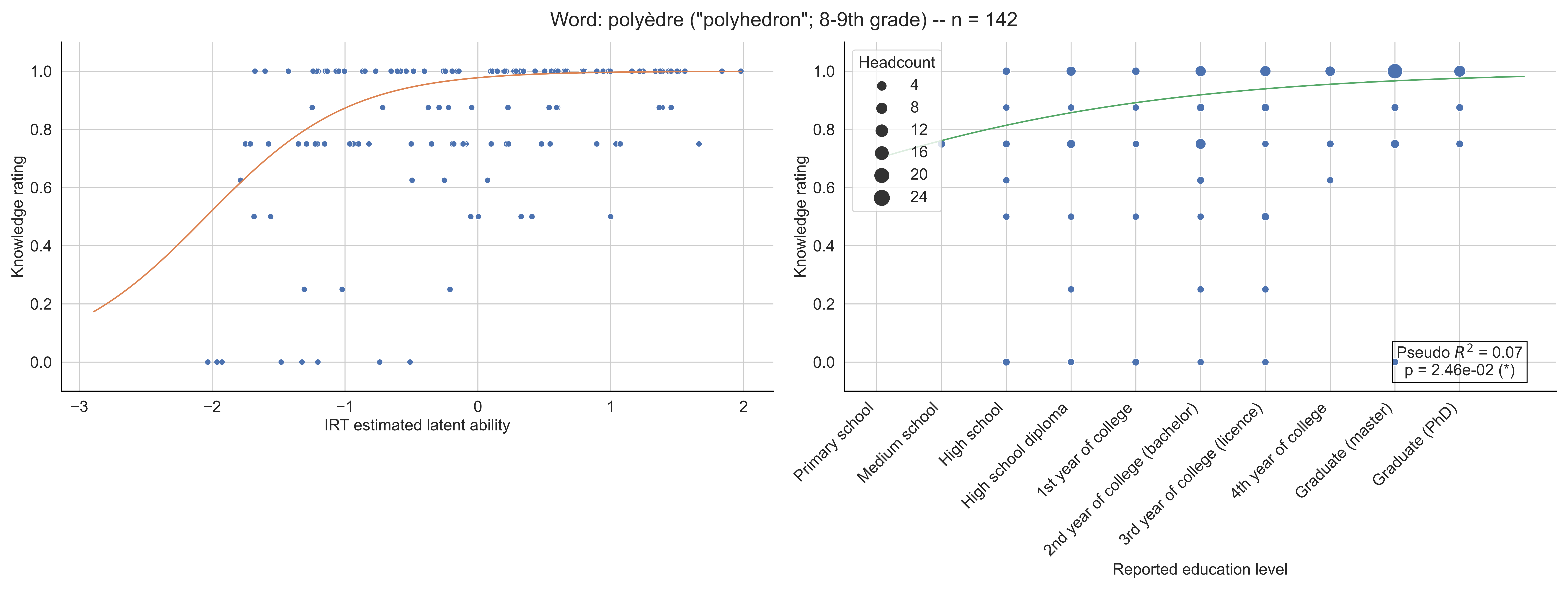

Word: polyèdre ("polyhedron"; 8-9th grade) -- n = 142
Knowledge rating
IRT estimated latent ability
Headcount
4
8
12
16
20
24
Pseudo $R^2$ = 0.07
p = 2.46e-02 (*)
Primary school
Medium school
High school
High school diploma
1st year of college
2nd year of college (bachelor)
3rd year of college (licence)
4th year of college
Graduate (master)
Graduate (PhD)
Reported education level

Word: premier ("prime"; 6-7th grade) -- n = 165

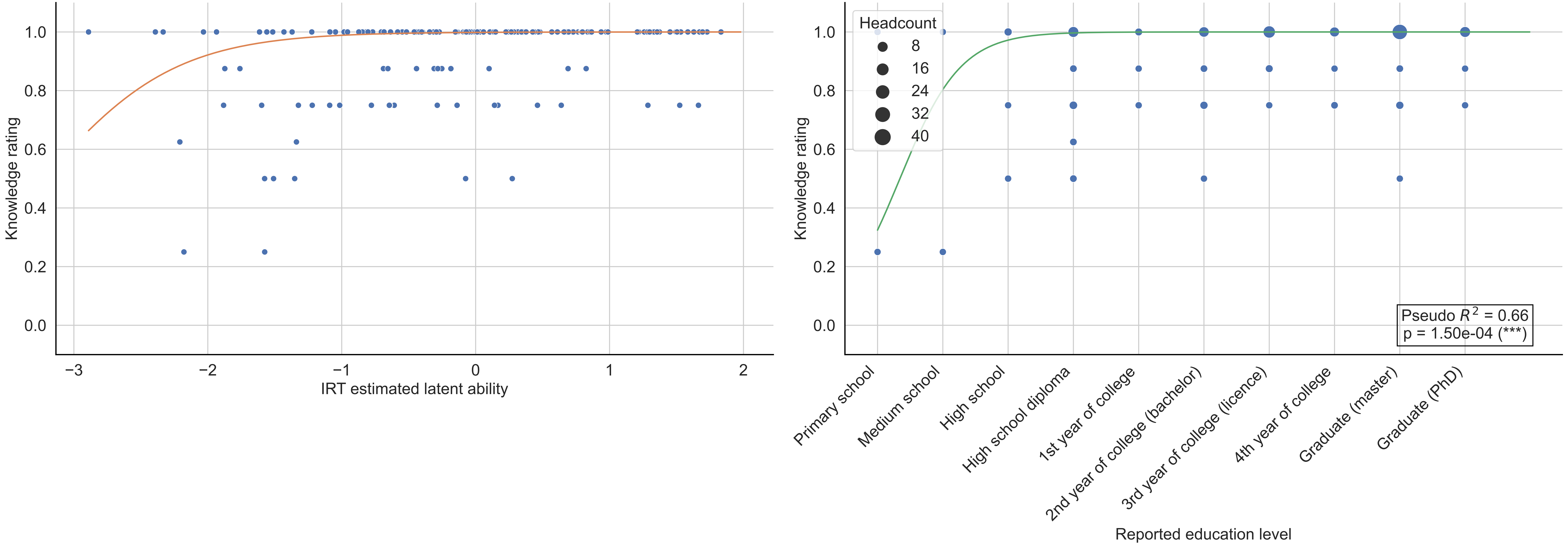

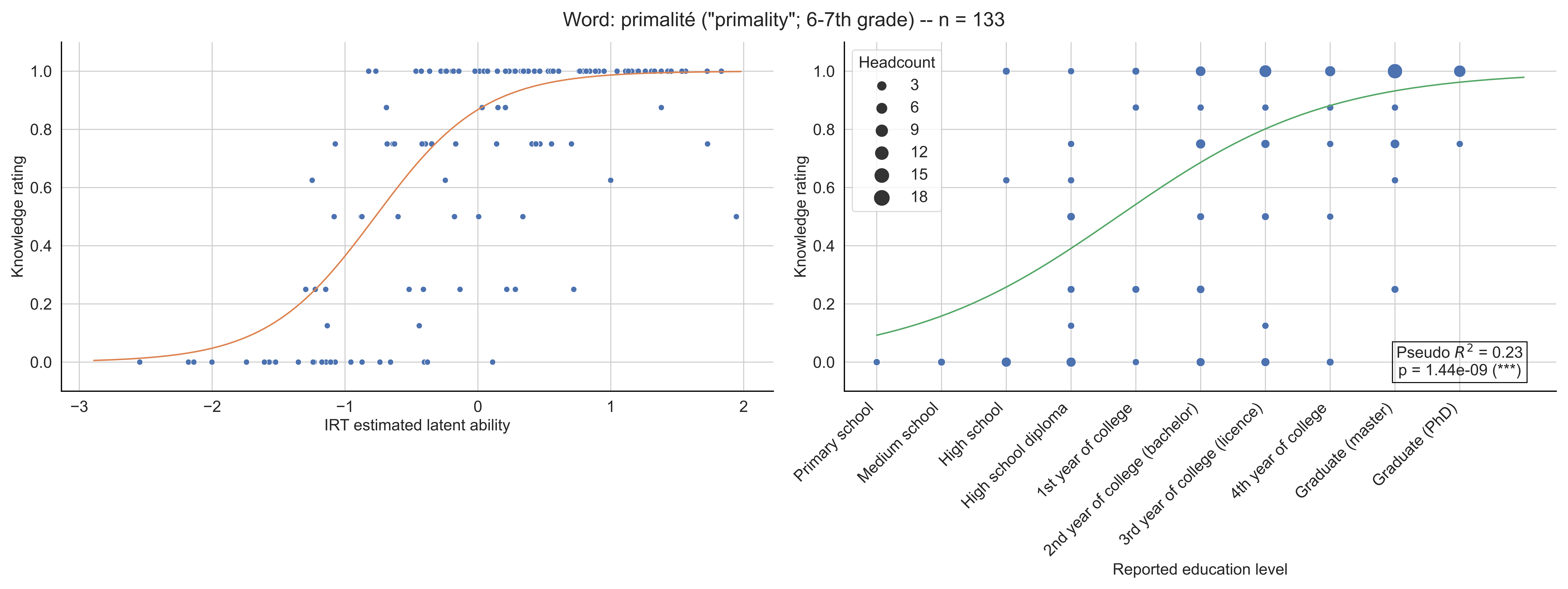
Word: primalité ("primality"; 6-7th grade) -- n = 133
Knowledge rating
IRT estimated latent ability
Headcount
3
6
9
12
15
18
Pseudo $R^2$ = 0.23
p = 1.44e-09 (***)
Primary school
Medium school
High school
High school diploma
1st year of college
2nd year of college (bachelor)
3rd year of college (licence)
4th year of college
Graduate (master)
Graduate (PhD)
Reported education level

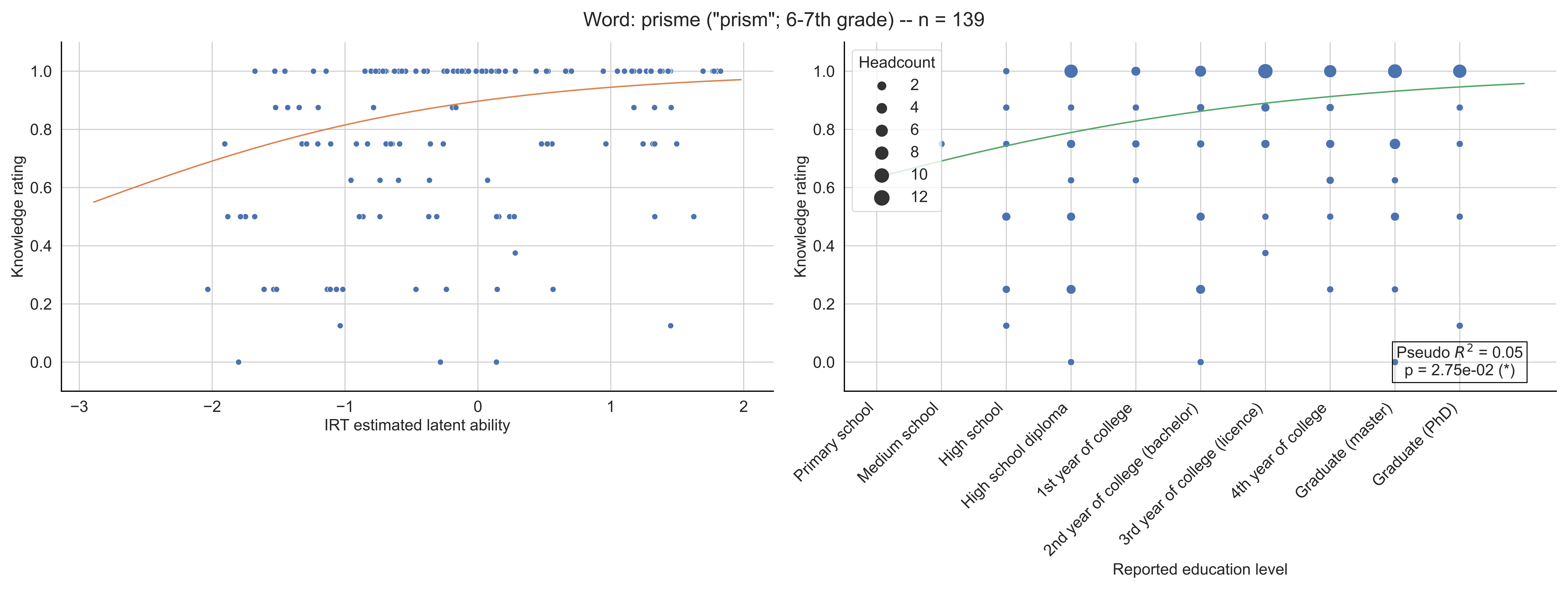
Word: prisme ("prism"; 6-7th grade) -- n = 139
Knowledge rating
IRT estimated latent ability
Headcount
2
4
6
8
10
12
Pseudo $R^2$ = 0.05
p = 2.75e-02 (*)
Primary school
Medium school
High school
High school diploma
1st year of college
2nd year of college (bachelor)
3rd year of college (licence)
4th year of college
Graduate (master)
Graduate (PhD)
Reported education level

Word: profondeur ("depth"; 6-7th grade) -- n = 156

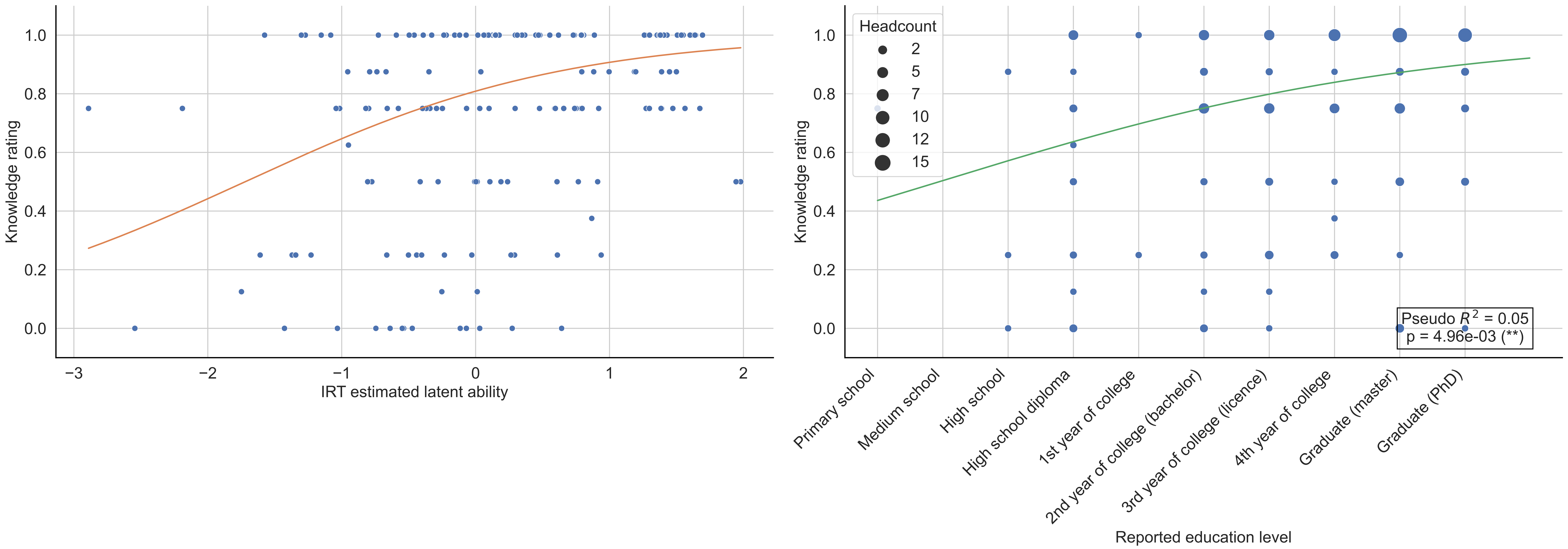

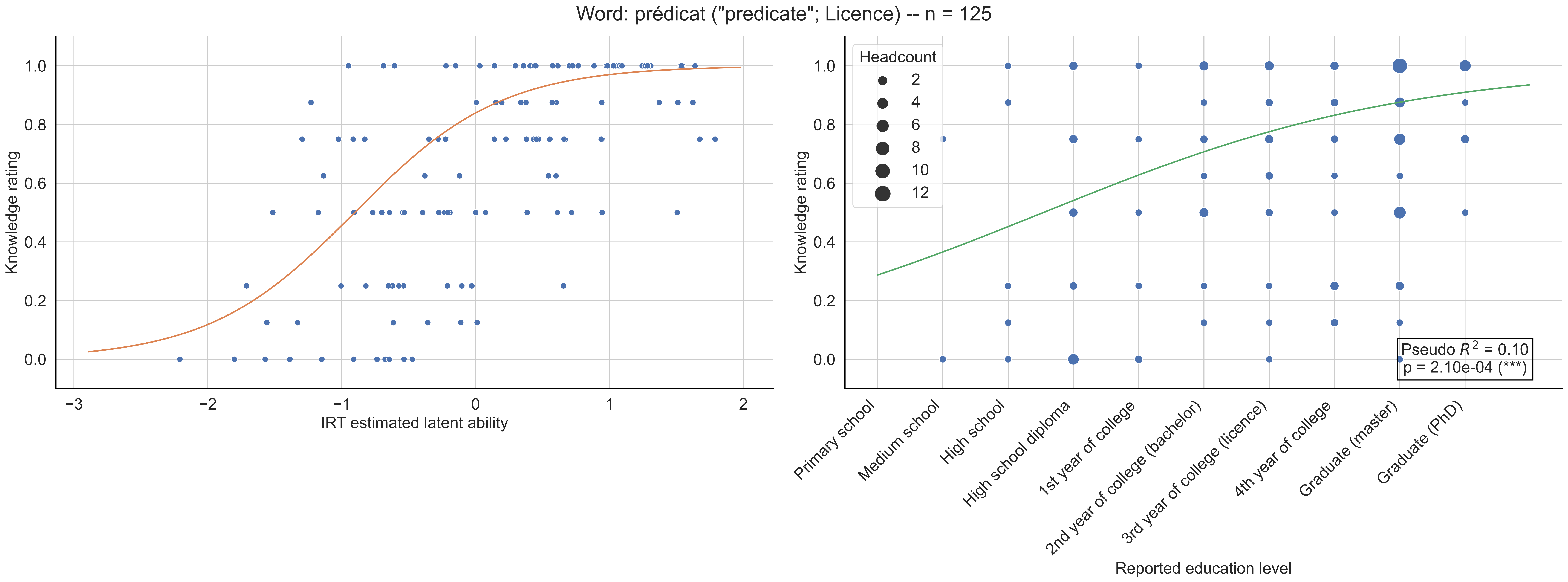
Word: prédicat ("predicate"; Licence) -- n = 125
Knowledge rating
0.0
0.2
0.4
0.6
0.8
1.0
−3
−2
−1
0
1
2
IRT estimated latent ability
Knowledge rating
Headcount
2
4
6
8
10
12
Pseudo $R^2$ = 0.10
p = 2.10e-04 (***)
Primary school
Medium school
High school
High school diploma
1st year of college
2nd year of college (bachelor)
3rd year of college (licence)
4th year of college
Graduate (master)
Graduate (PhD)
Reported education level

Word: préordre ("preorder"; Licence) -- n = 145

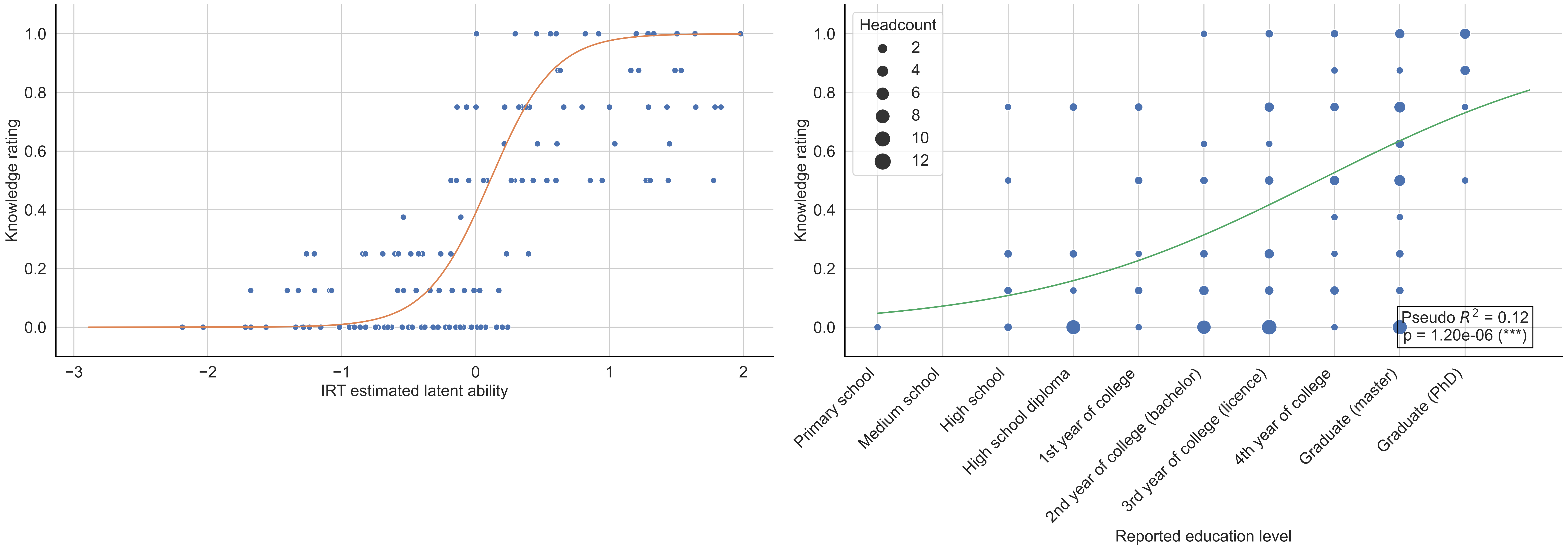

Word: périmètre ("permieter"; Primary school) -- n = 155

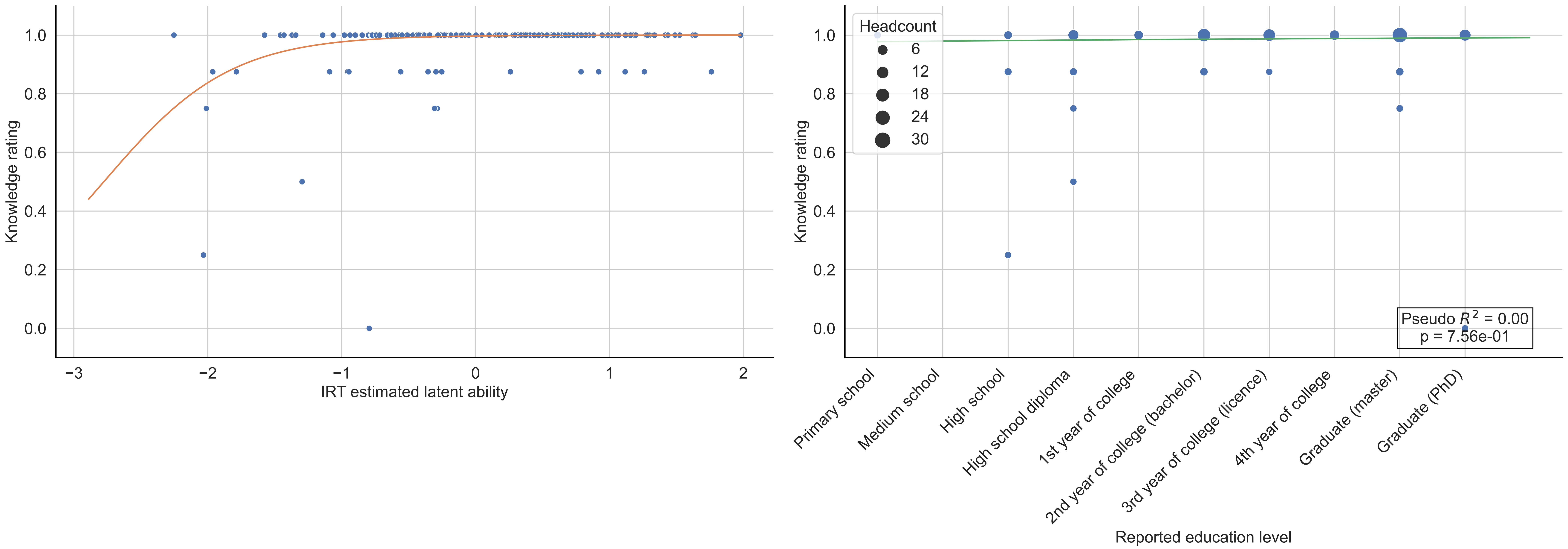

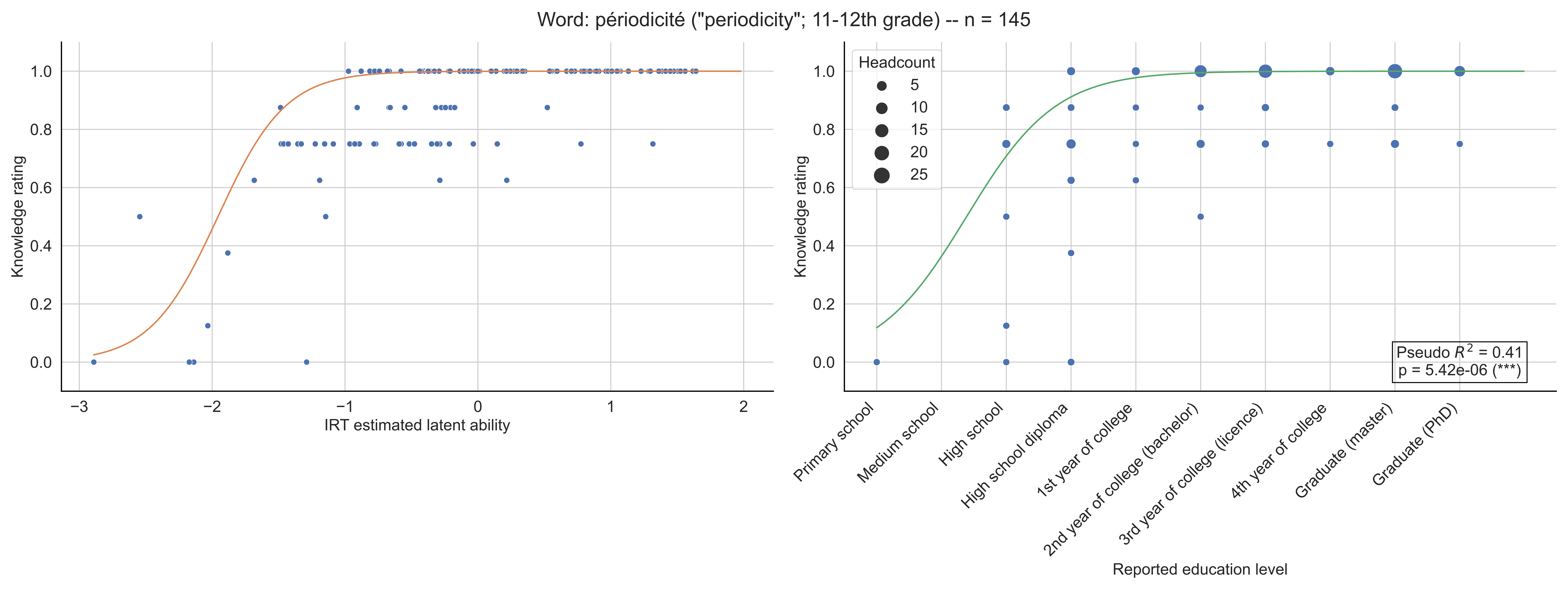
Word: périodicité ("periodicity"; 11-12th grade) -- n = 145
Knowledge rating
IRT estimated latent ability
Headcount
5
10
15
20
25
Pseudo $R^2$ = 0.41
p = 5.42e-06 (***)
Primary school
Medium school
High school
High school diploma
1st year of college
2nd year of college (bachelor)
3rd year of college (licence)
4th year of college
Graduate (master)
Graduate (PhD)
Reported education level

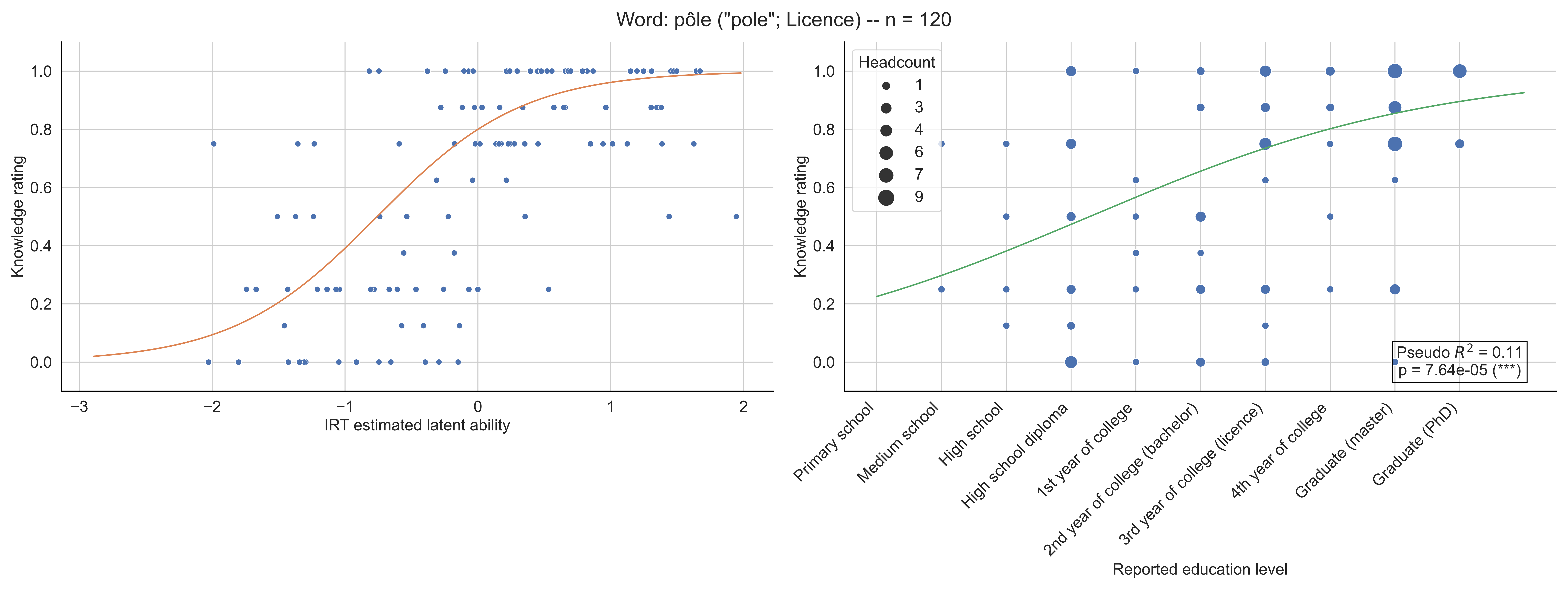

Word: pôle ("pole"; Licence) -- n = 120
Knowledge rating
IRT estimated latent ability
Headcount
1
3
4
6
7
9
Pseudo $R^2$ = 0.11
p = 7.64e-05 (***)
Primary school
Medium school
High school
High school diploma
1st year of college
2nd year of college (bachelor)
3rd year of college (licence)
4th year of college
Graduate (master)
Graduate (PhD)
Reported education level

Word: quadrature ("quadrature"; Master) -- n = 166

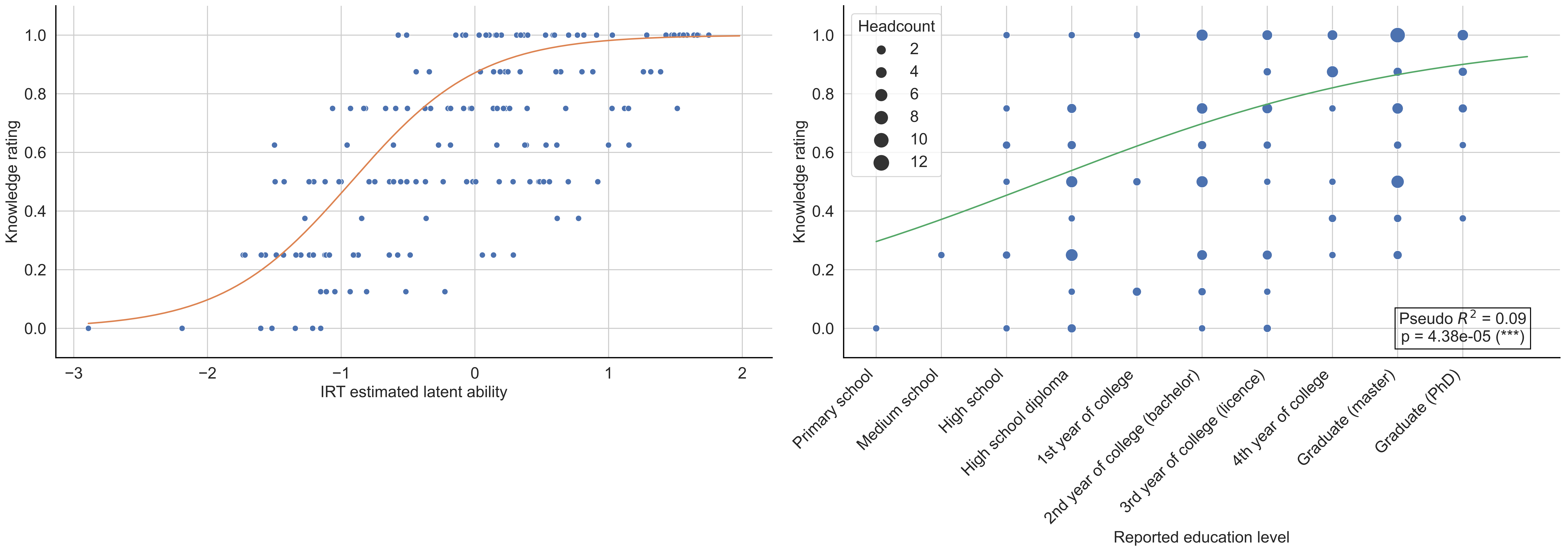

Word: quadrilatère ("quadrilateral"; Primary school) -- n = 140

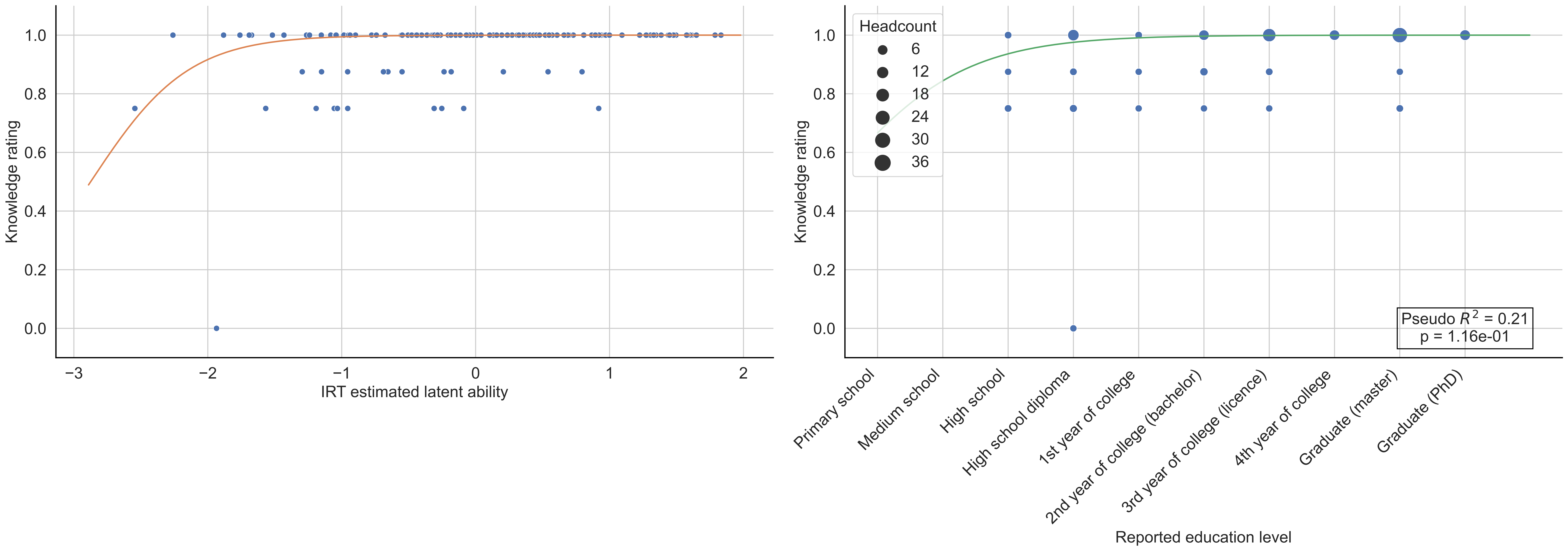

Word: quantificateur ("quantifier"; Bachelor) -- n = 142

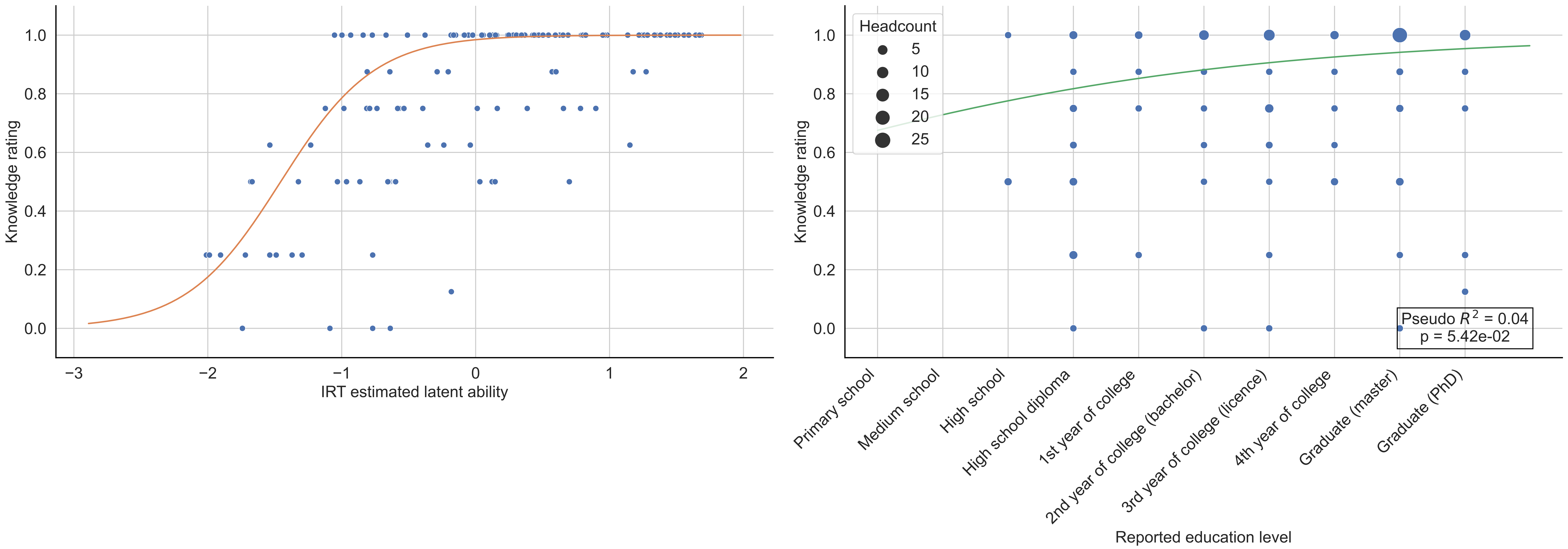

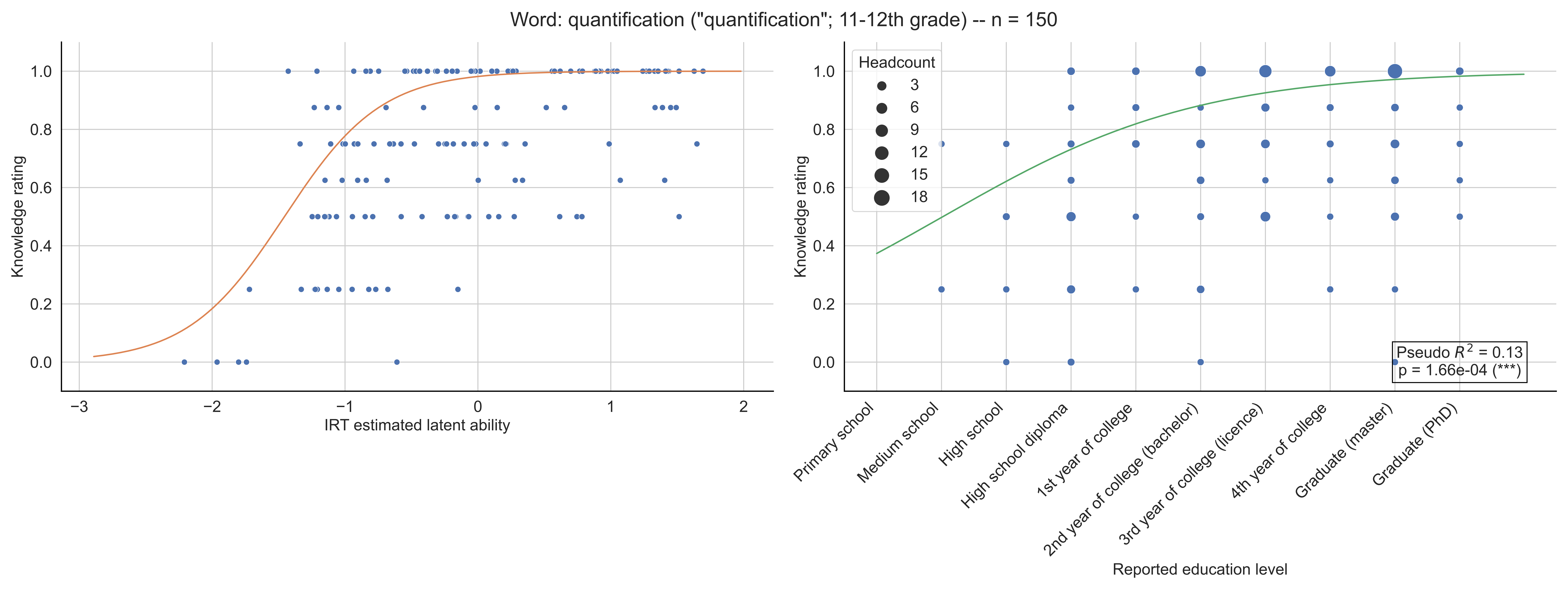

Word: quantification ("quantification"; 11-12th grade) -- n = 150
Knowledge rating
IRT estimated latent ability
Headcount
3
6
9
12
15
18
Pseudo $R^2$ = 0.13
p = 1.66e-04 (***)
Primary school
Medium school
High school
High school diploma
1st year of college
2nd year of college (bachelor)
3rd year of college (licence)
4th year of college
Graduate (master)
Graduate (PhD)
Reported education level

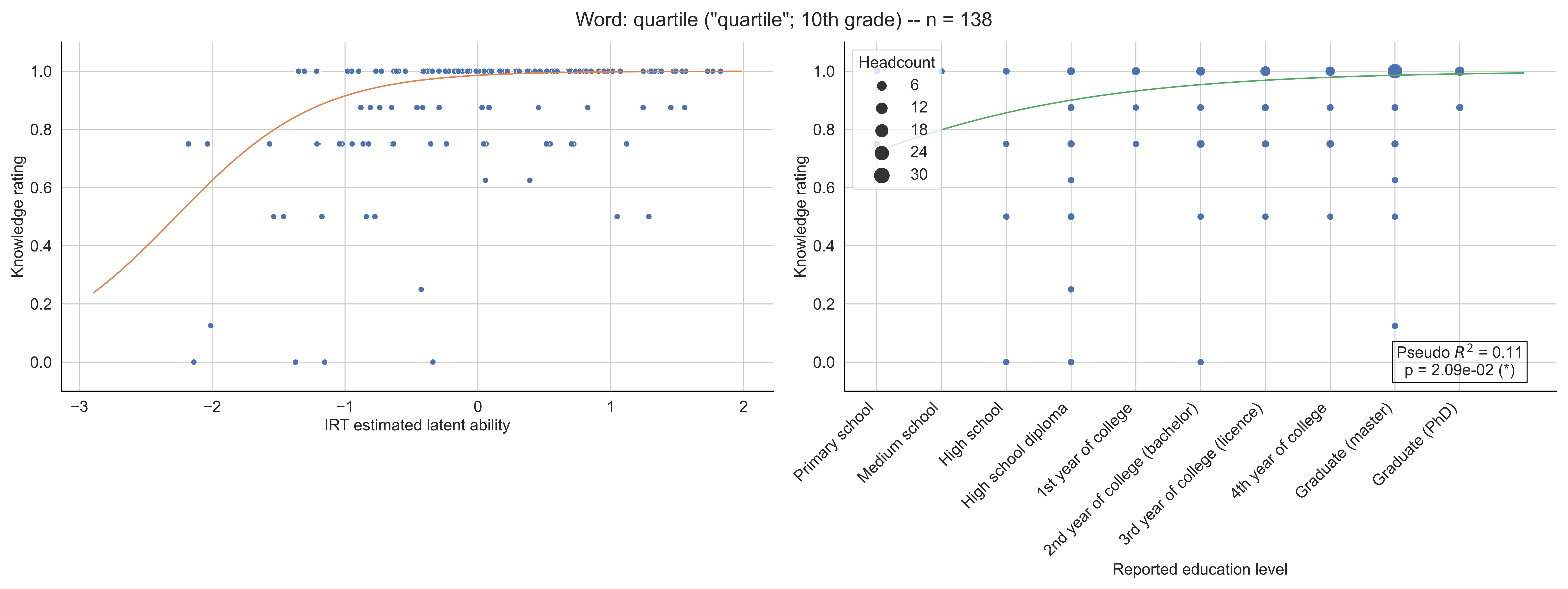

Word: quartile ("quartile"; 10th grade) -- n = 138
Knowledge rating
IRT estimated latent ability
Headcount
6
12
18
24
30
Primary school
Medium school
High school
High school diploma
1st year of college
2nd year of college (bachelor)
3rd year of college (licence)
4th year of college
Graduate (master)
Graduate (PhD)
Reported education level
Pseudo $R^2$ = 0.11
p = 2.09e-02 (*)

Word: quaternion ("quaternion"; Bachelor) -- n = 139

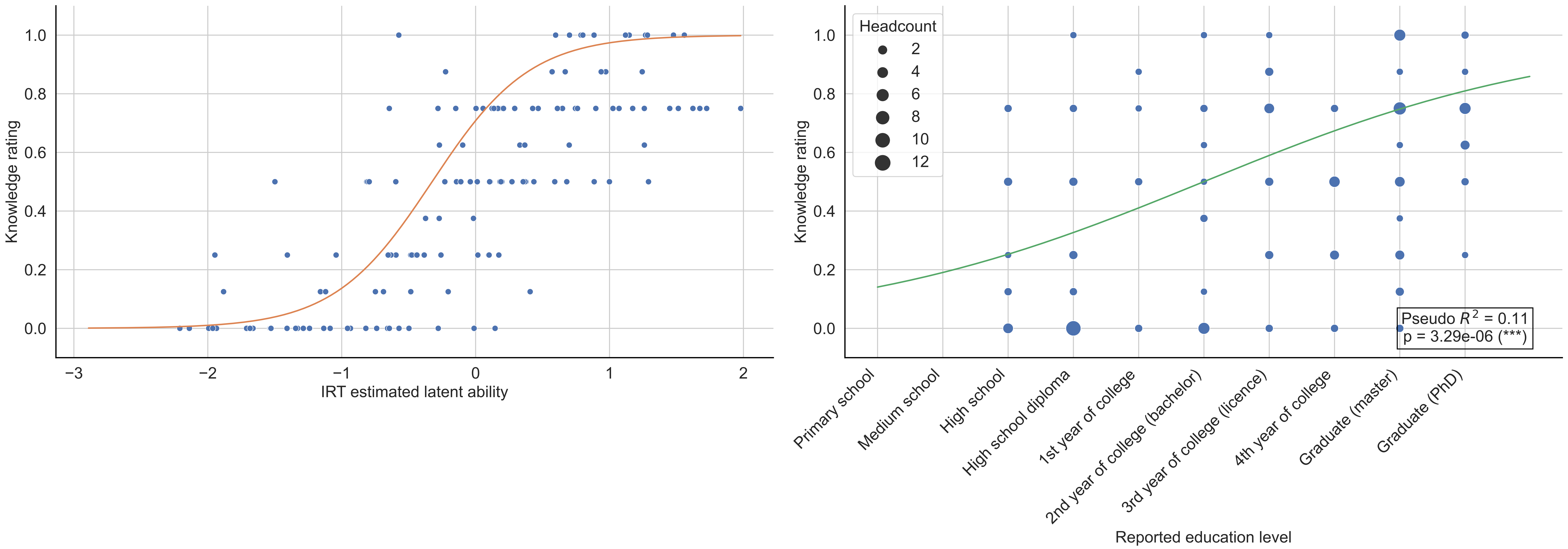

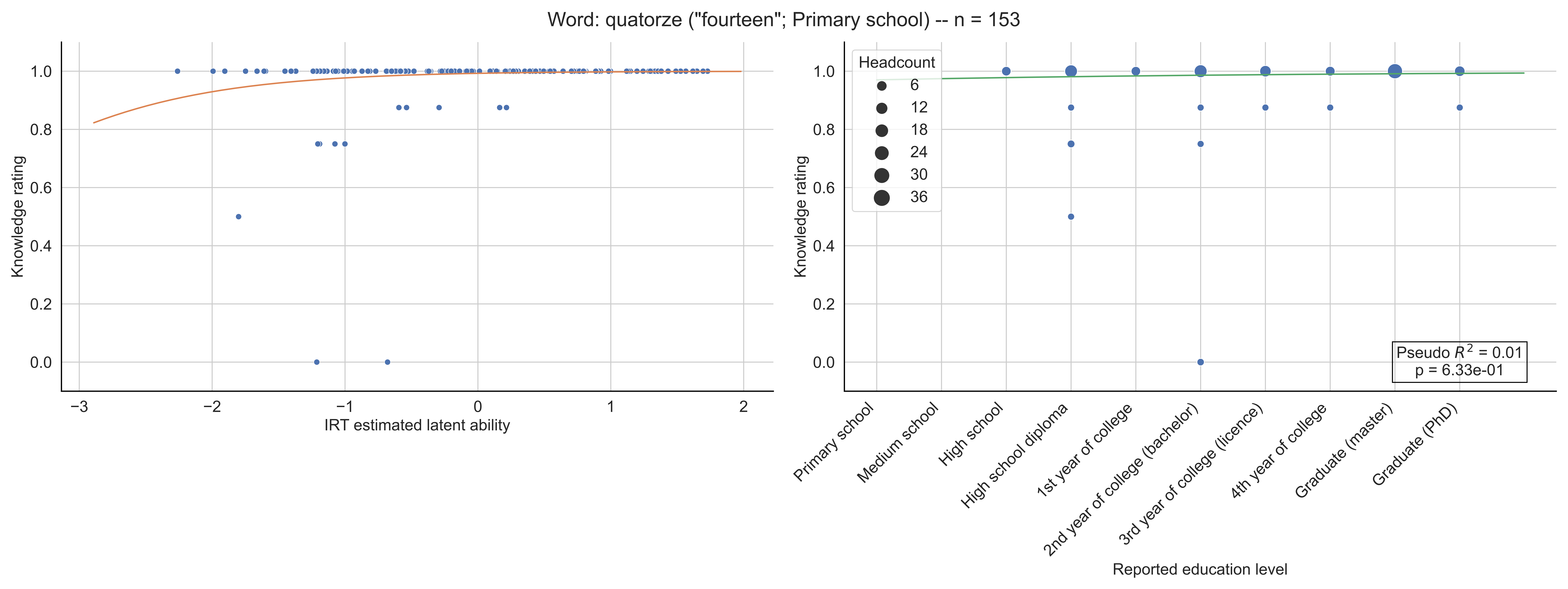
Word: quatorze ("fourteen"; Primary school) -- n = 153
Knowledge rating
IRT estimated latent ability
Headcount
6
12
18
24
30
36
Pseudo $R^2$ = 0.01
p = 6.33e-01
Primary school
Medium school
High school
High school diploma
1st year of college
2nd year of college (bachelor)
3rd year of college (licence)
4th year of college
Graduate (master)
Graduate (PhD)
Reported education level

Word: radian ("radian"; 10th grade) -- n = 159

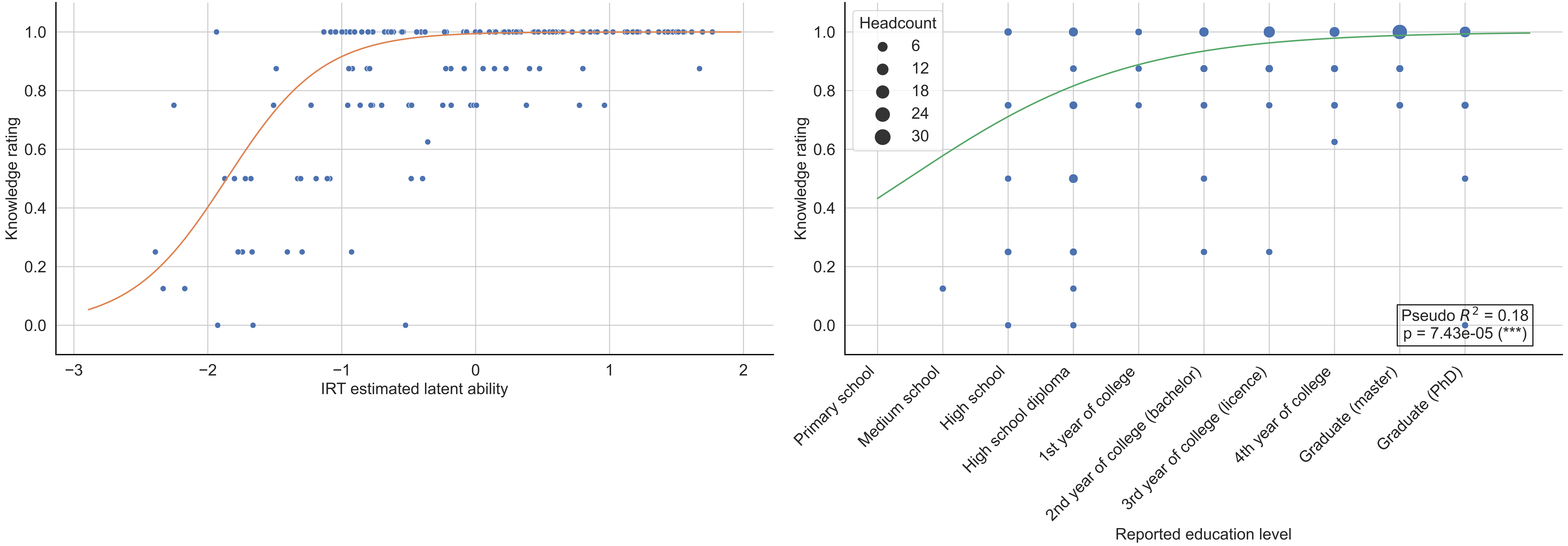

Word: rang ("rank"; Bachelor) -- n = 147

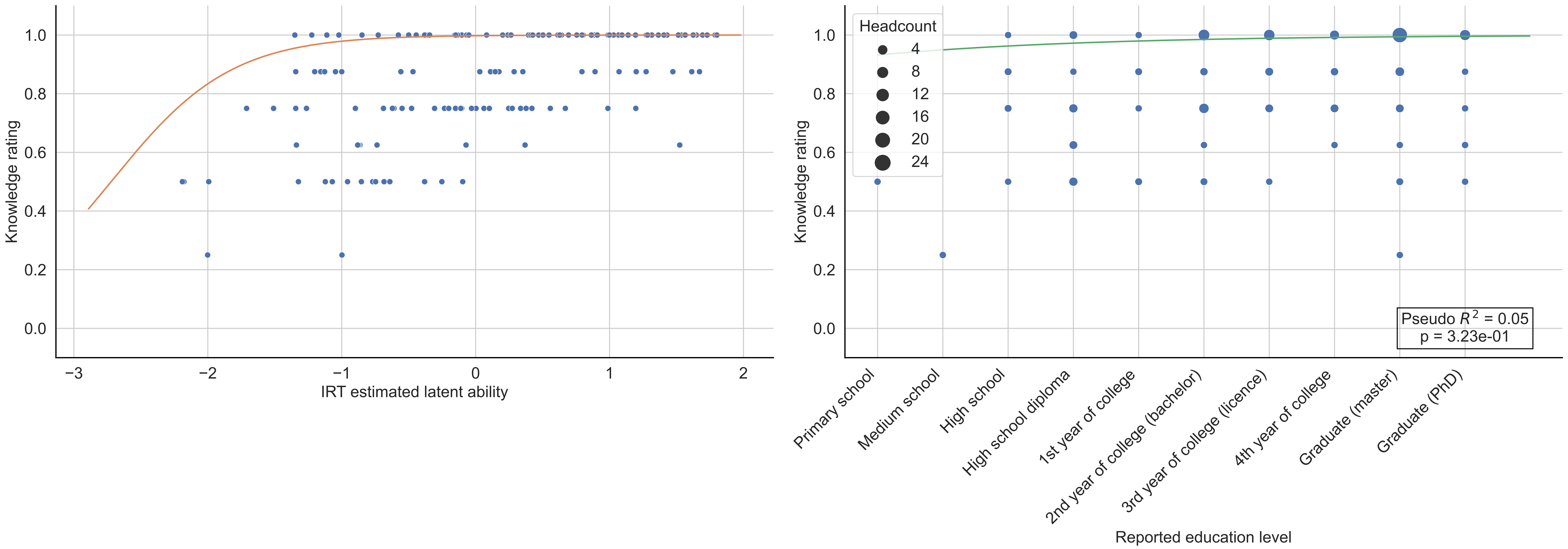

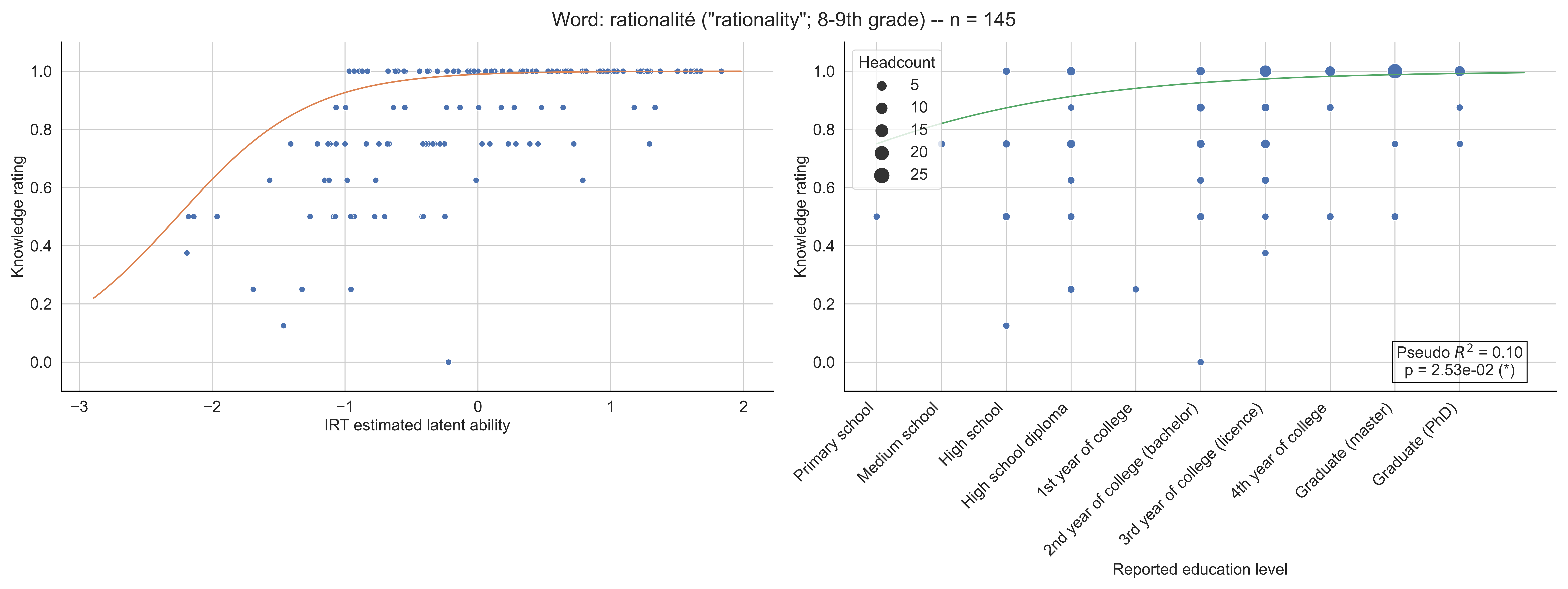

Word: rationalité ("rationality"; 8-9th grade) -- n = 145
Knowledge rating
IRT estimated latent ability
Headcount
5
10
15
20
25
Pseudo $R^2$ = 0.10
p = 2.53e-02 (*)
Primary school
Medium school
High school
High school diploma
1st year of college
2nd year of college (bachelor)
3rd year of college (licence)
4th year of college
Graduate (master)
Graduate (PhD)
Reported education level

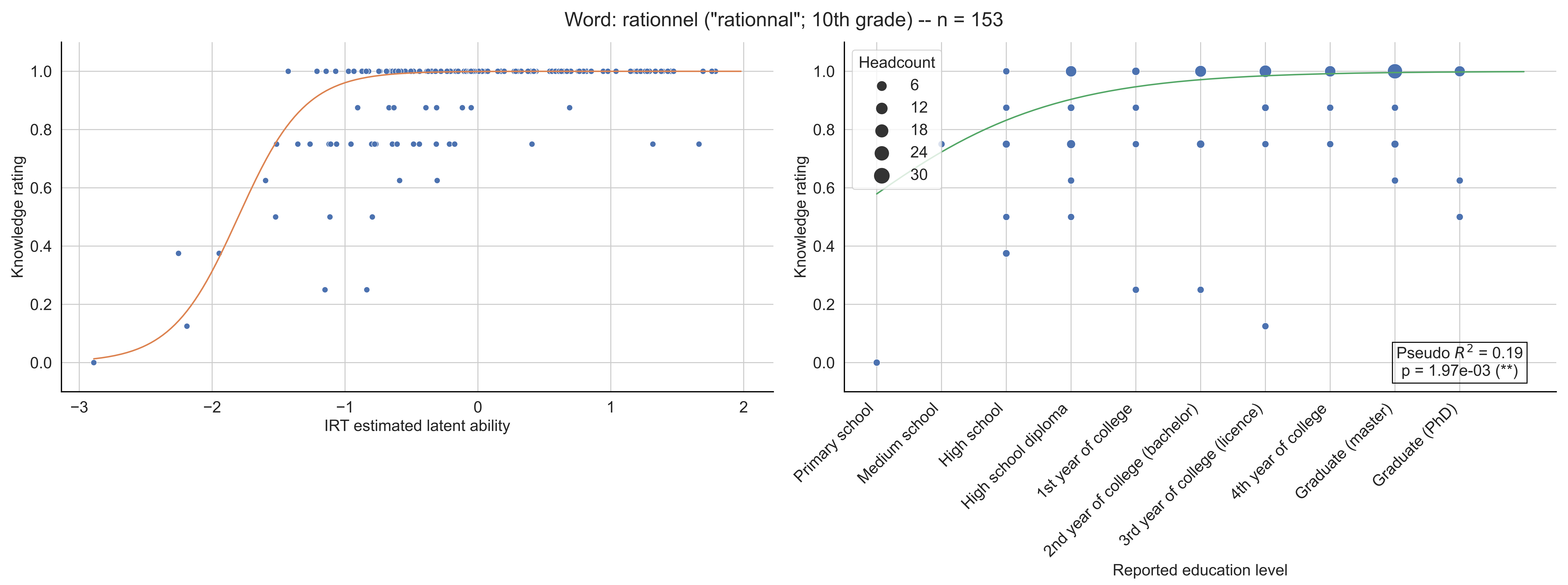
Word: rationnel ("rationnal"; 10th grade) -- n = 153
Knowledge rating
IRT estimated latent ability
Headcount
6
12
18
24
30
Pseudo $R^2$ = 0.19
p = 1.97e-03 (**)
Primary school
Medium school
High school
High school diploma
1st year of college
2nd year of college (bachelor)
3rd year of college (licence)
4th year of college
Graduate (master)
Graduate (PhD)
Reported education level

Word: relatif (6-7th grade) -- n = 150

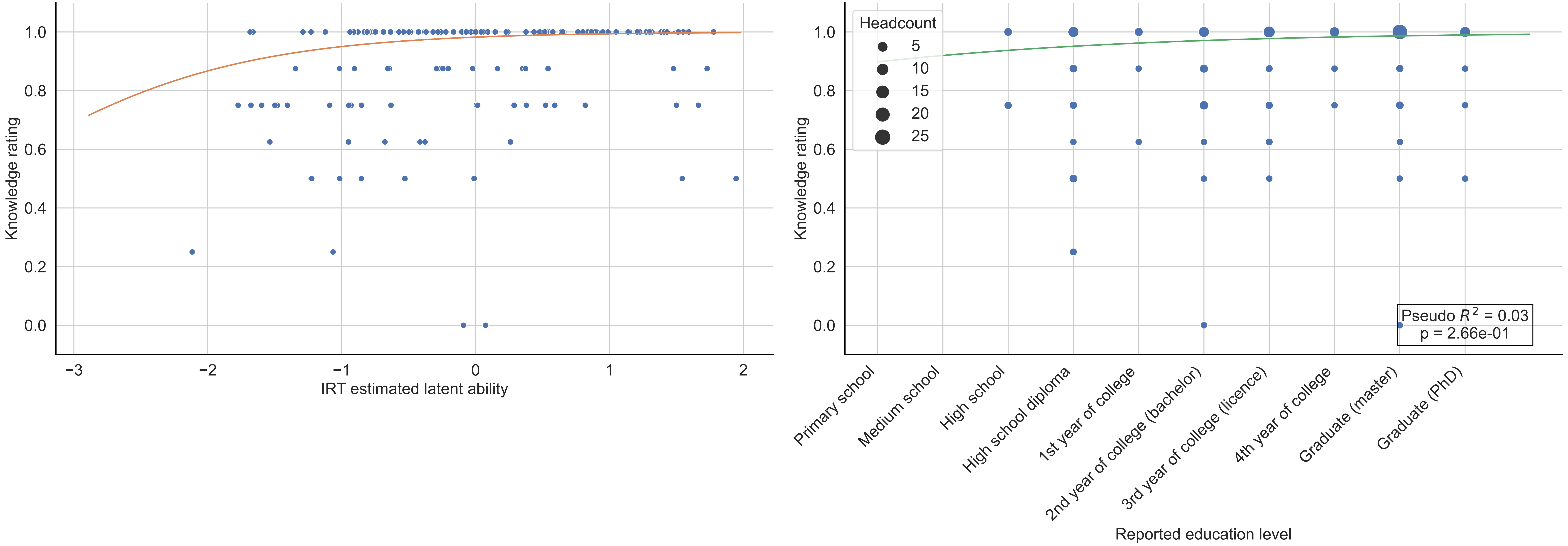

Word: relation ("relation"; 10th grade) -- n = 141

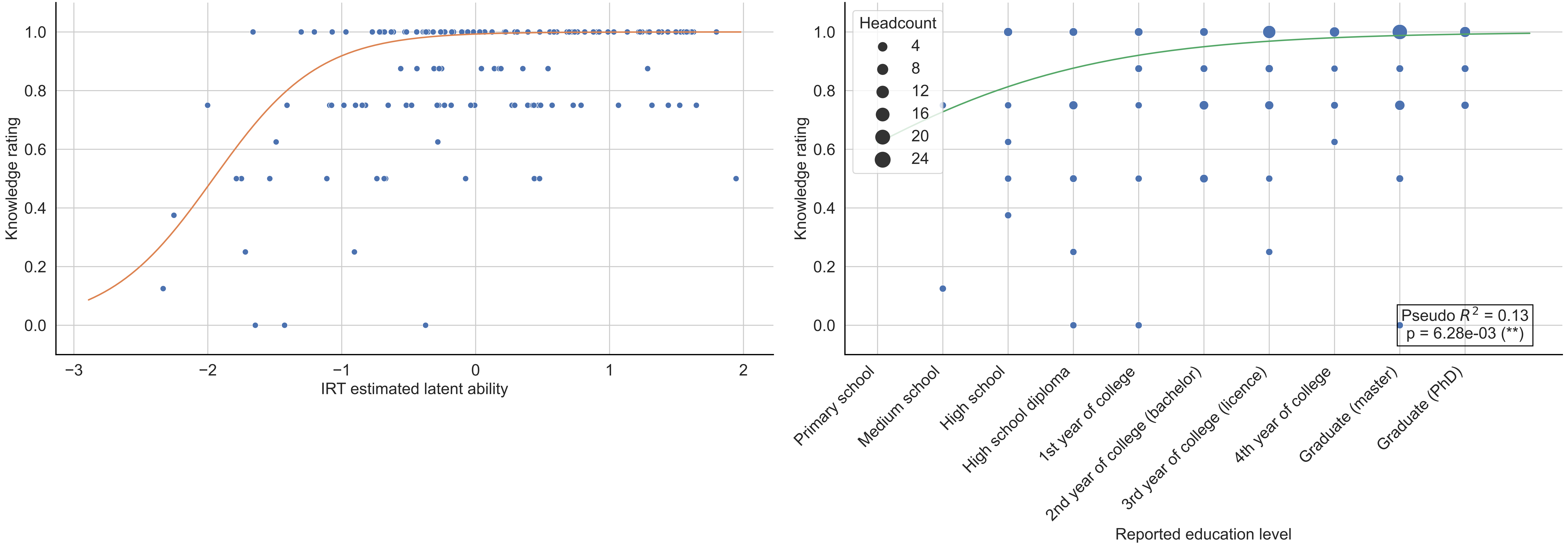

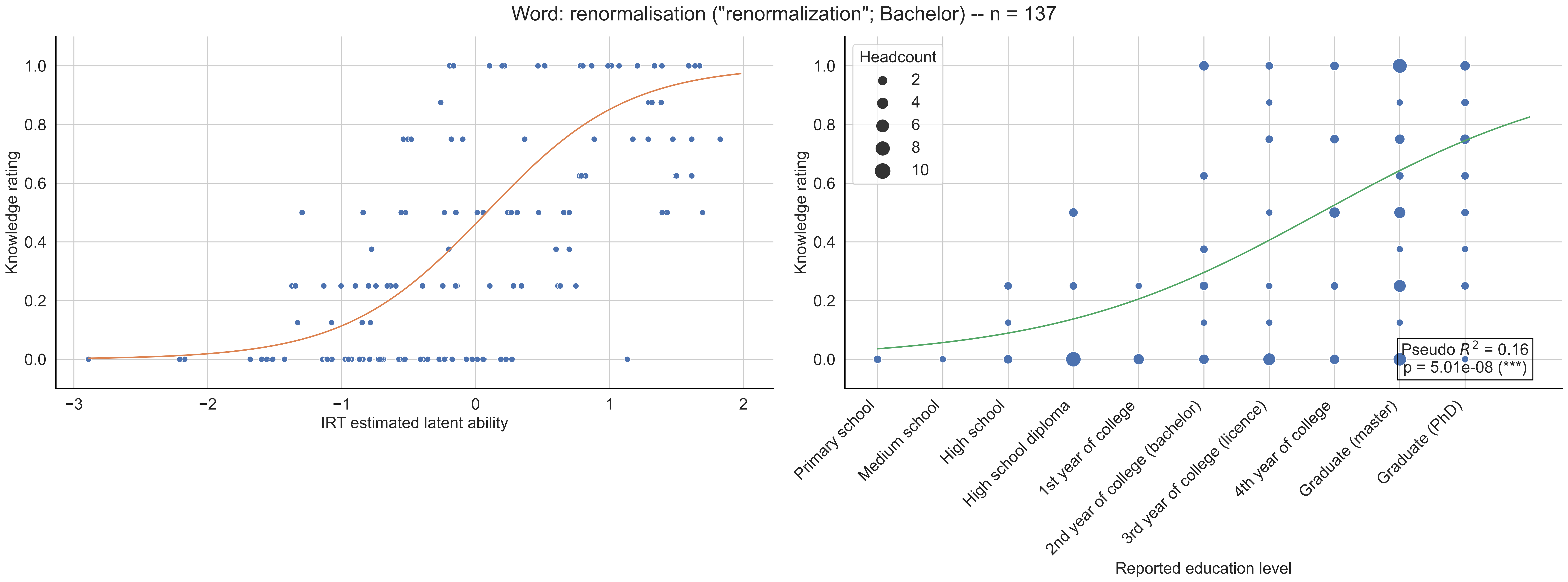
Word: renormalisation ("renormalization"; Bachelor) -- n = 137
Knowledge rating
IRT estimated latent ability
Headcount
2
4
6
8
10
Pseudo $R^2$ = 0.16
p = 5.01e-08 (***)
Primary school
Medium school
High school
High school diploma
1st year of college
2nd year of college (bachelor)
3rd year of college (licence)
4th year of college
Graduate (master)
Graduate (PhD)
Reported education level

Word: représentation ("representation"; 8-9th grade) -- n = 148

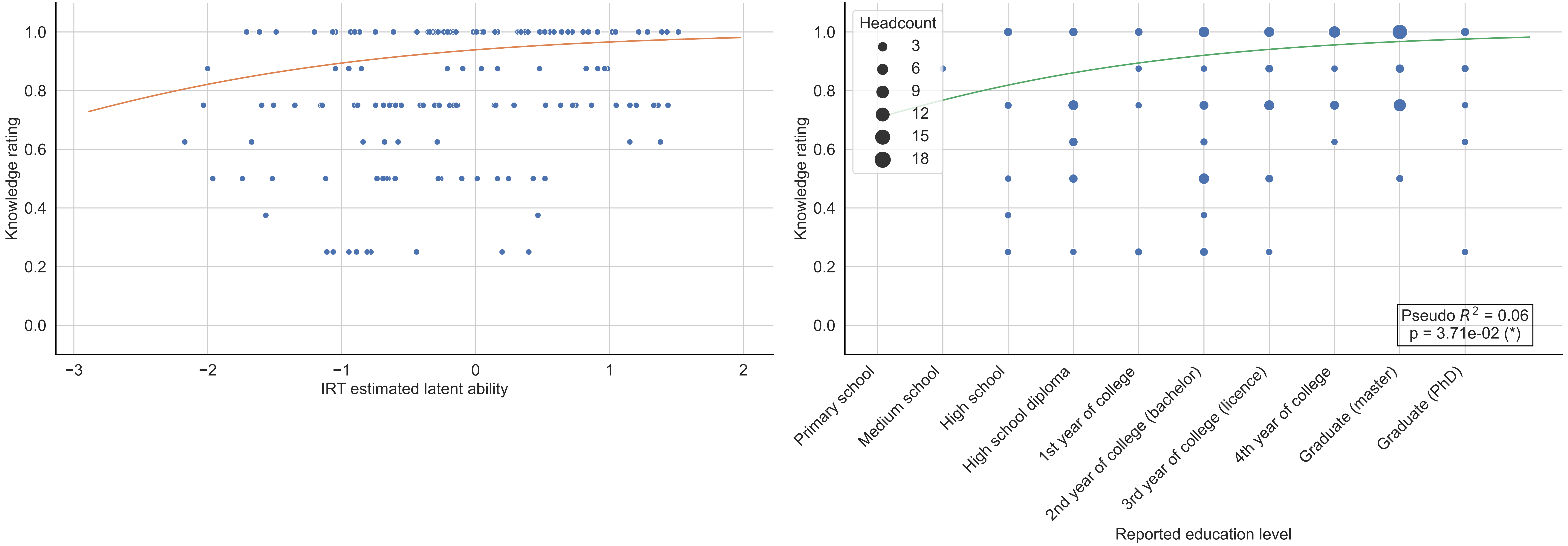

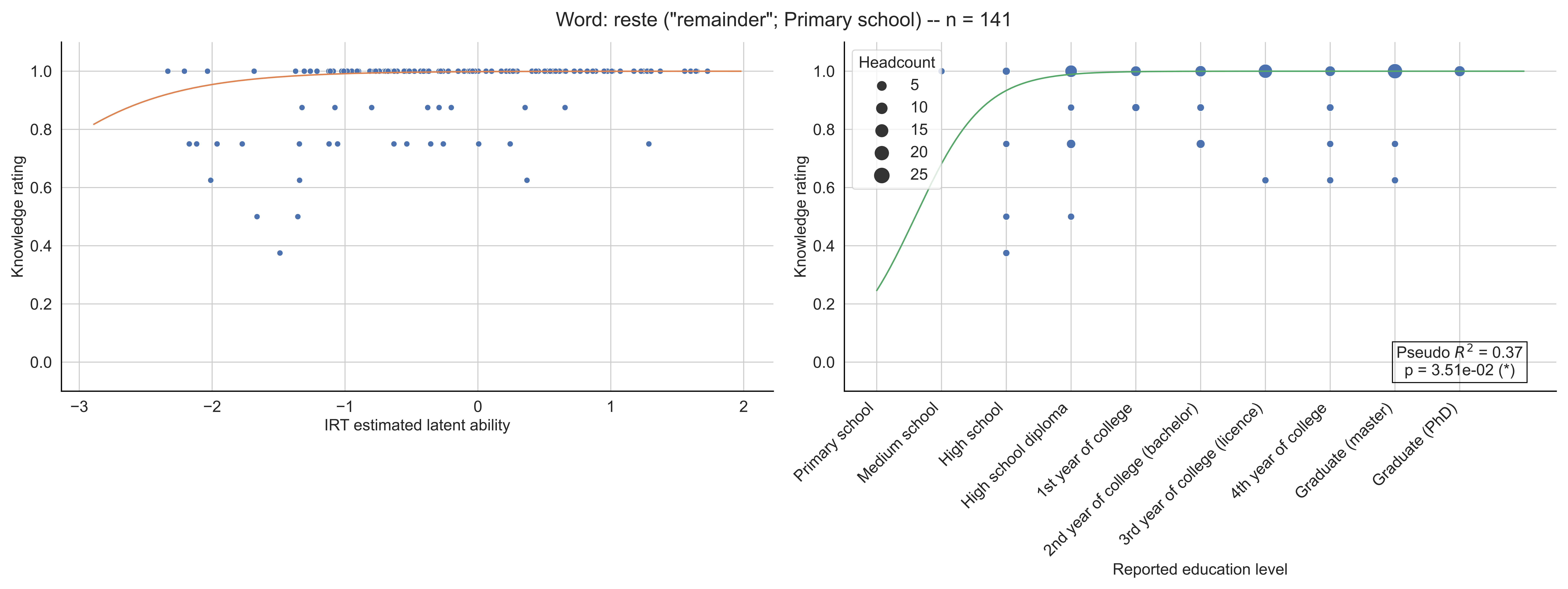
Word: reste ("remainder"; Primary school) -- n = 141
Knowledge rating
IRT estimated latent ability
Headcount
5
10
15
20
25
Pseudo $R^2$ = 0.37
p = 3.51e-02 (*)
Primary school
Medium school
High school
High school diploma
1st year of college
2nd year of college (bachelor)
3rd year of college (licence)
4th year of college
Graduate (master)
Graduate (PhD)
Reported education level

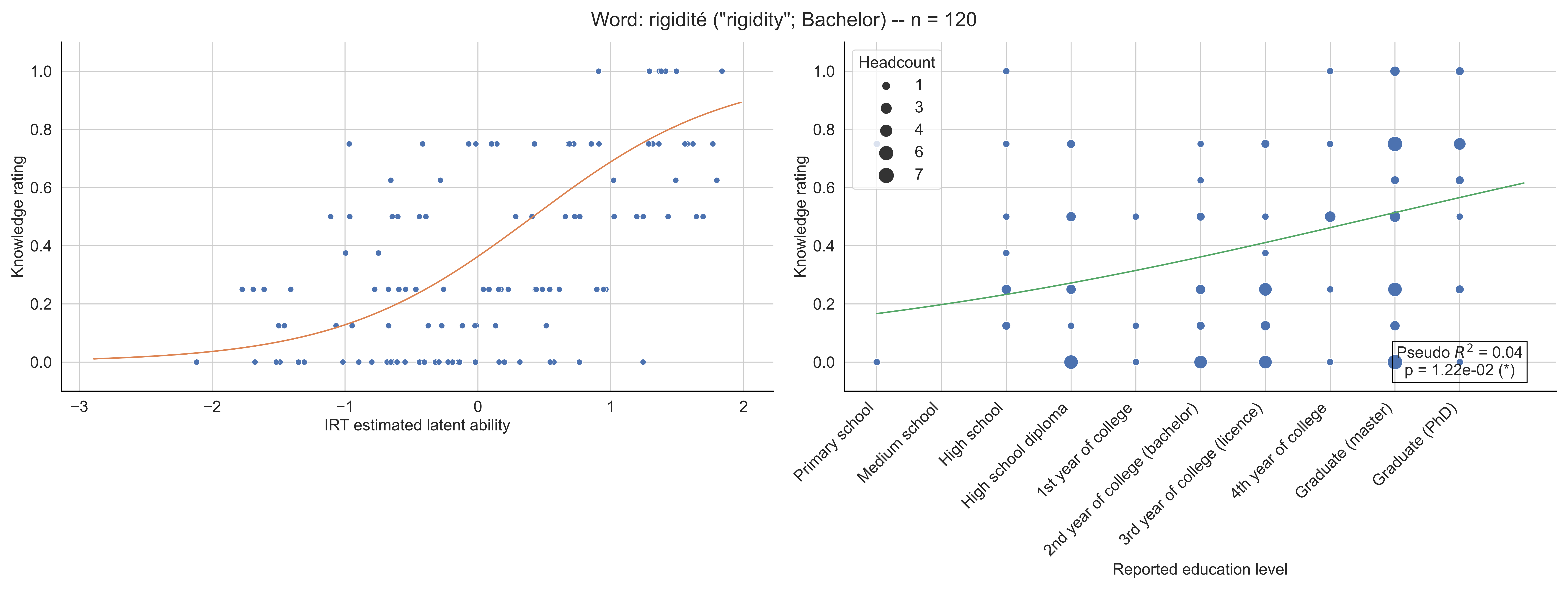

Word: rigidité ("rigidity"; Bachelor) -- n = 120
Knowledge rating
IRT estimated latent ability
Headcount
1
3
4
6
7
Pseudo $R^2$ = 0.04
p = 1.22e-02 (*)
Primary school
Medium school
High school
High school diploma
1st year of college
2nd year of college (bachelor)
3rd year of college (licence)
4th year of college
Graduate (master)
Graduate (PhD)
Reported education level

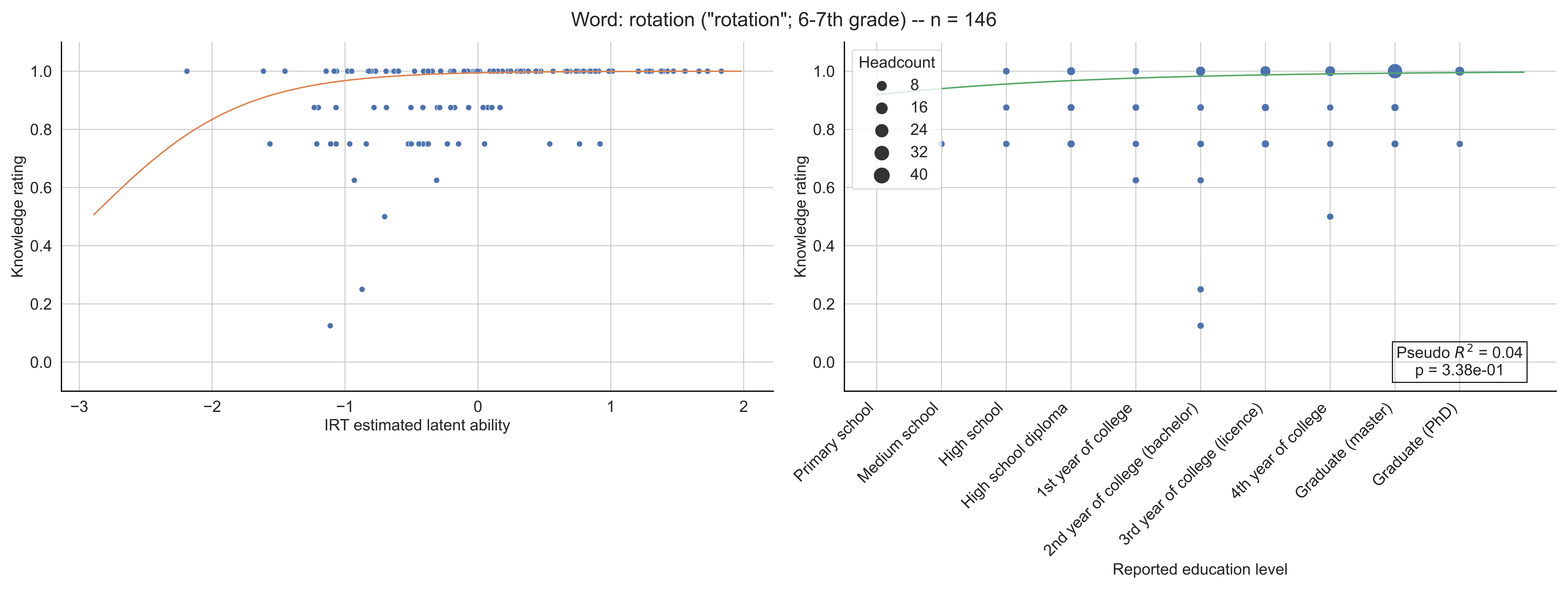
Word: rotation ("rotation"; 6-7th grade) -- n = 146
Knowledge rating
IRT estimated latent ability
Headcount
8
16
24
32
40
Pseudo $R^2$ = 0.04
p = 3.38e-01
Primary school
Medium school
High school
High school diploma
1st year of college
2nd year of college (bachelor)
3rd year of college (licence)
4th year of college
Graduate (master)
Graduate (PhD)
Reported education level

Word: réciprocité ("reciprocity"; Licence) -- n = 162

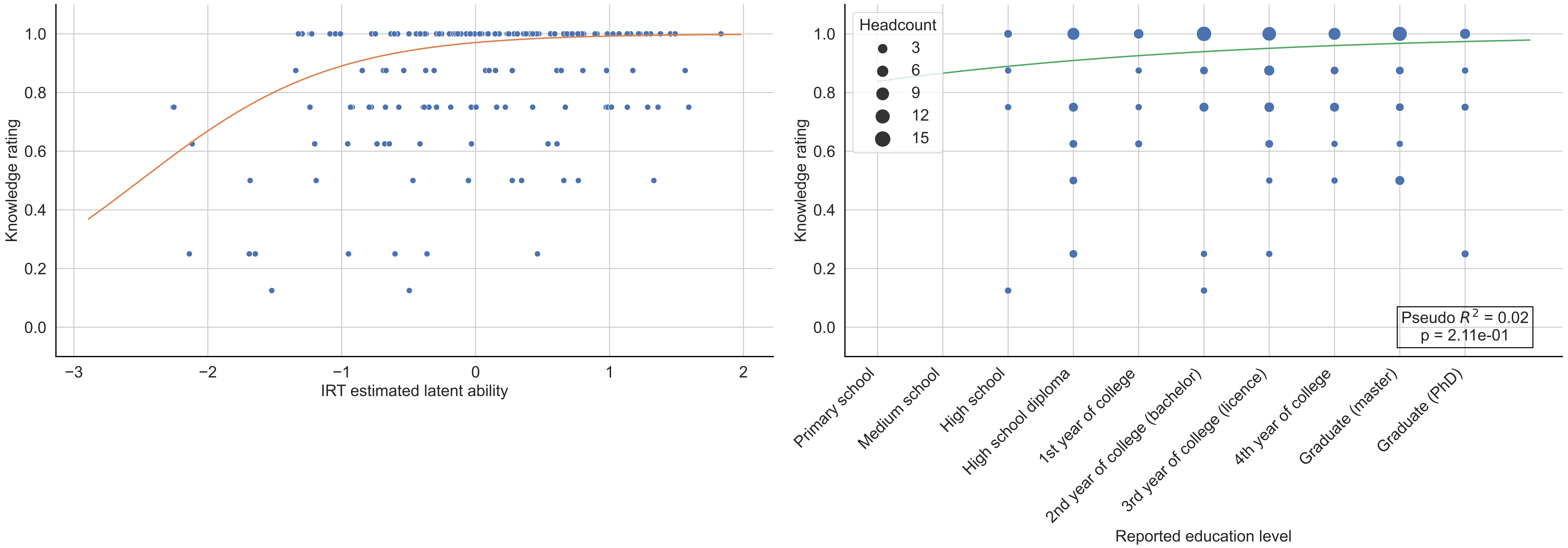

Word: réciproque ("reciprocal"; 8-9th grade) -- n = 150

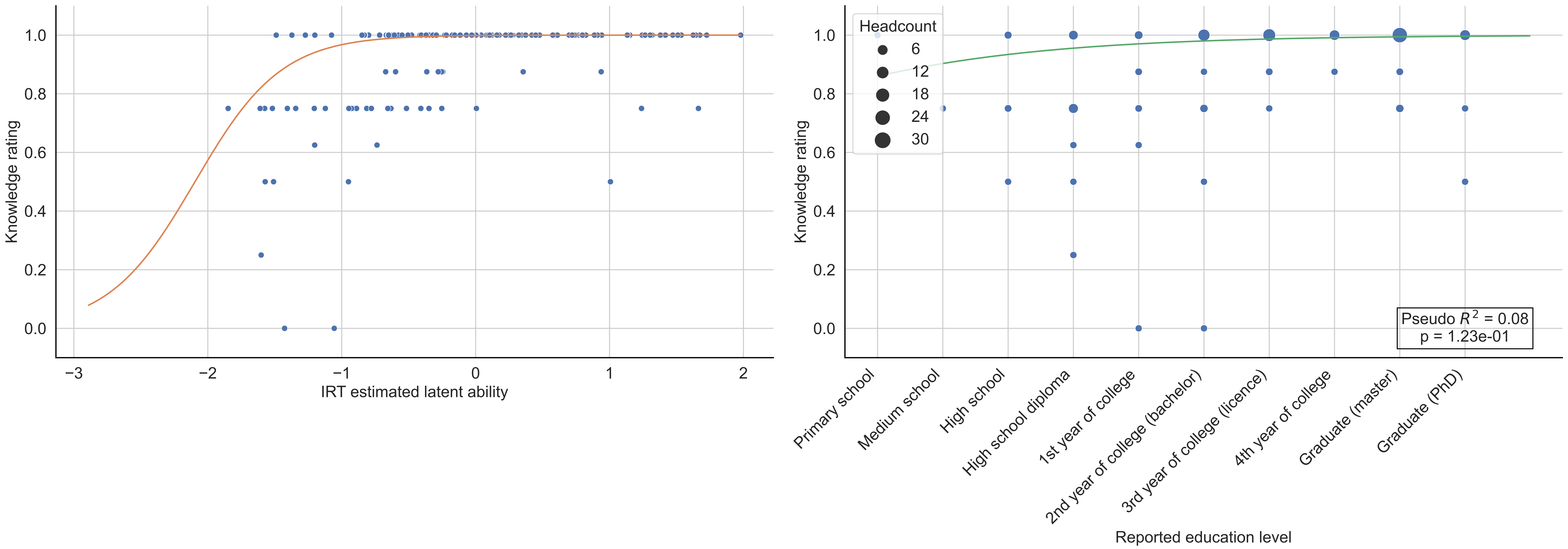

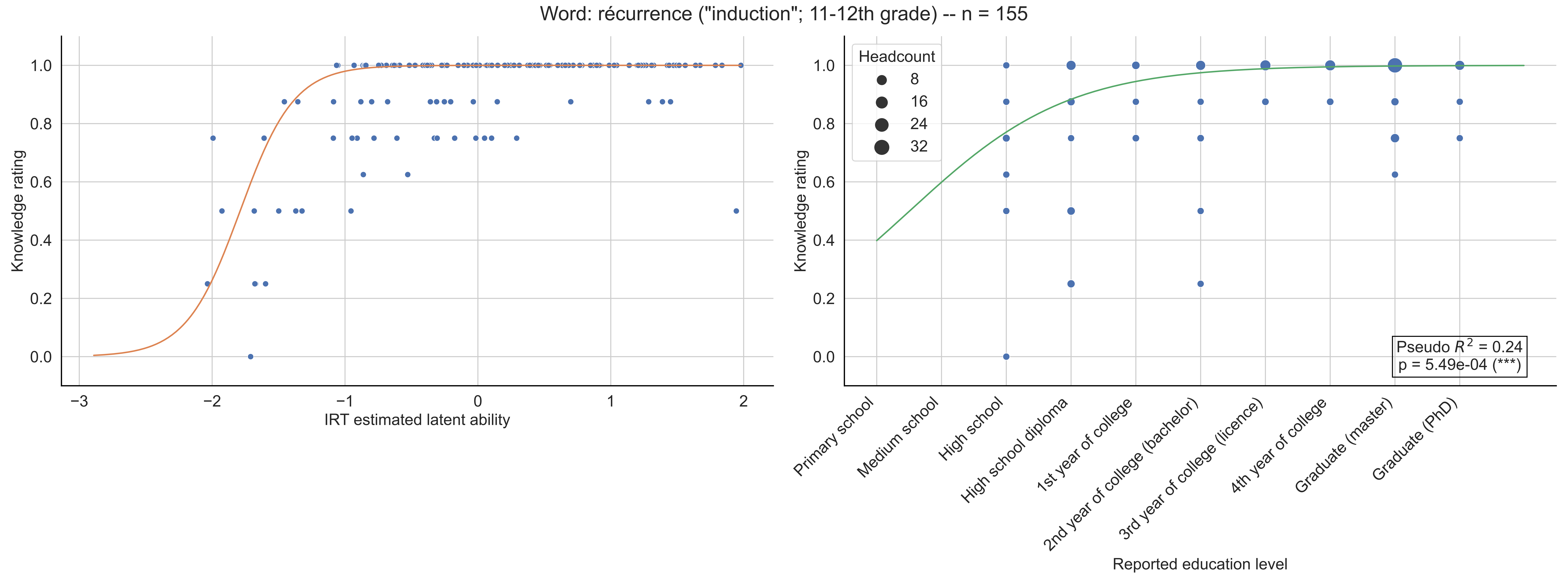

Word: récurrence ("induction"; 11-12th grade) -- n = 155
Knowledge rating
IRT estimated latent ability
Headcount
8
16
24
32
Pseudo $R^2$ = 0.24
p = 5.49e-04 (***)
Primary school
Medium school
High school
High school diploma
1st year of college
2nd year of college (bachelor)
3rd year of college (licence)
4th year of college
Graduate (master)
Graduate (PhD)
Reported education level

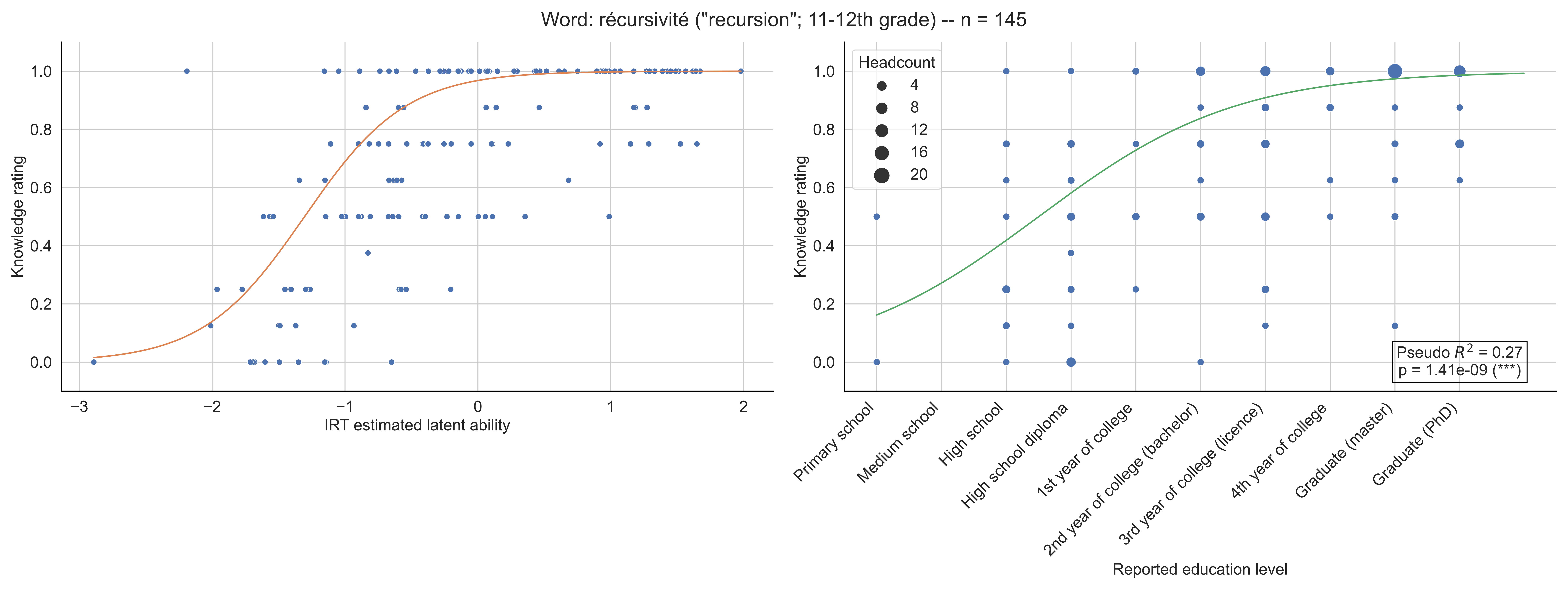

Word: récursivité ("recursion"; 11-12th grade) -- n = 145
Knowledge rating
IRT estimated latent ability
Headcount
4
8
12
16
20
Pseudo $R^2$ = 0.27
p = 1.41e-09 (***)
Primary school
Medium school
High school
High school diploma
1st year of college
2nd year of college (bachelor)
3rd year of college (licence)
4th year of college
Graduate (master)
Graduate (PhD)
Reported education level

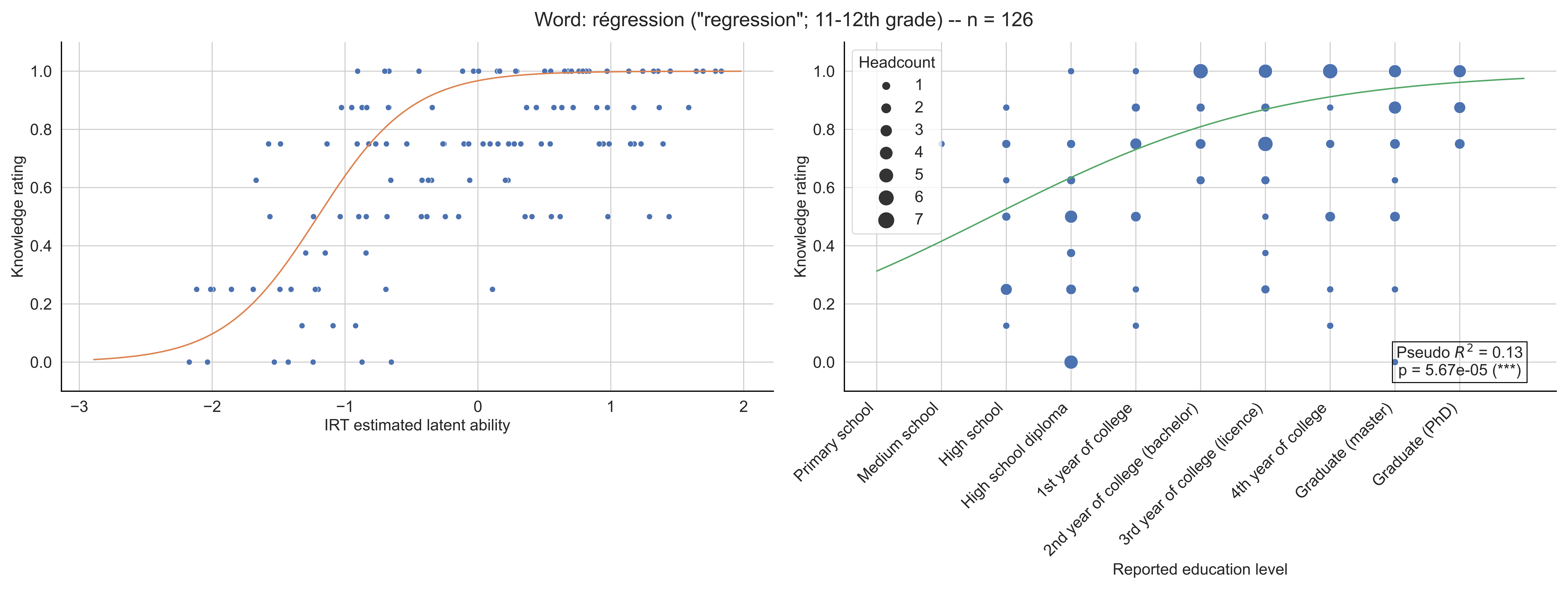

Word: régression ("regression"; 11-12th grade) -- n = 126
Knowledge rating
IRT estimated latent ability
Headcount
1
2
3
4
5
6
7
Pseudo $R^2$ = 0.13
p = 5.67e-05 (***)
Primary school
Medium school
High school
High school diploma
1st year of college
2nd year of college (bachelor)
3rd year of college (licence)
4th year of college
Graduate (master)
Graduate (PhD)
Reported education level

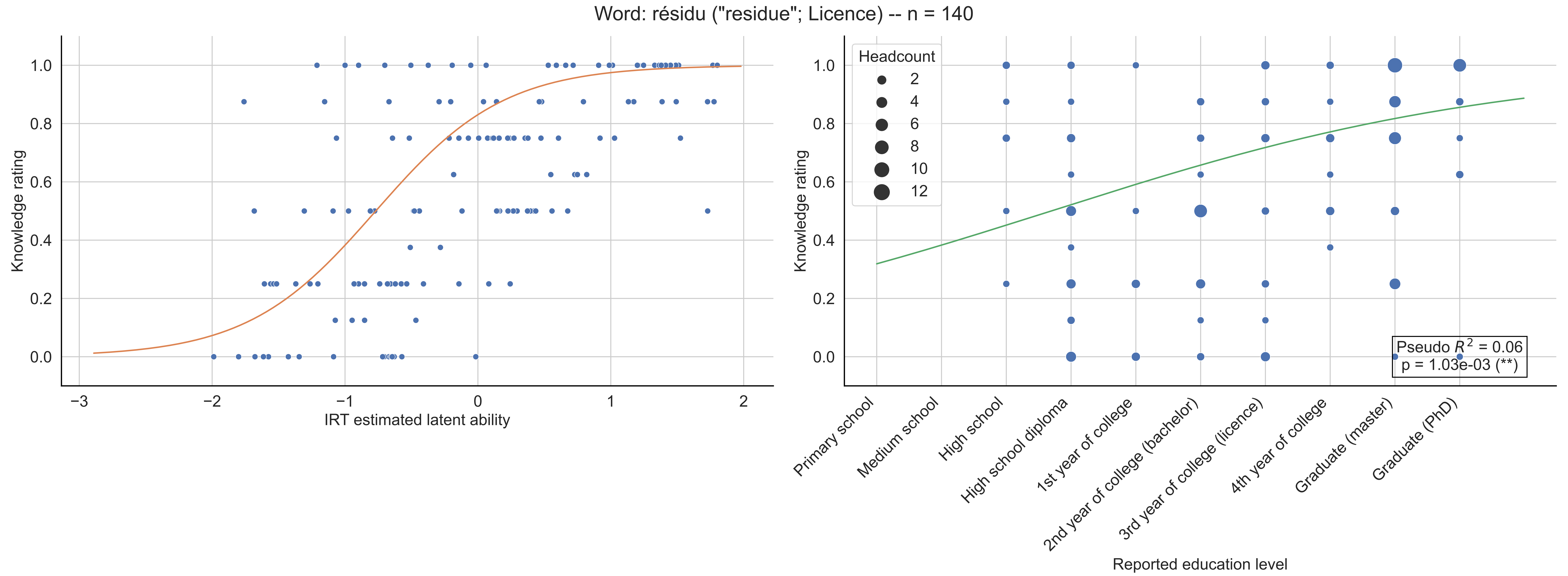
Word: résidu ("residue"; Licence) -- n = 140
Knowledge rating
IRT estimated latent ability
−3
−2
−1
0
1
2
0.0
0.2
0.4
0.6
0.8
1.0
Headcount
2
4
6
8
10
12
Pseudo $R^2$ = 0.06
p = 1.03e-03 (**)
Primary school
Medium school
High school
High school diploma
1st year of college
2nd year of college (bachelor)
3rd year of college (licence)
4th year of college
Graduate (master)
Graduate (PhD)
Reported education level

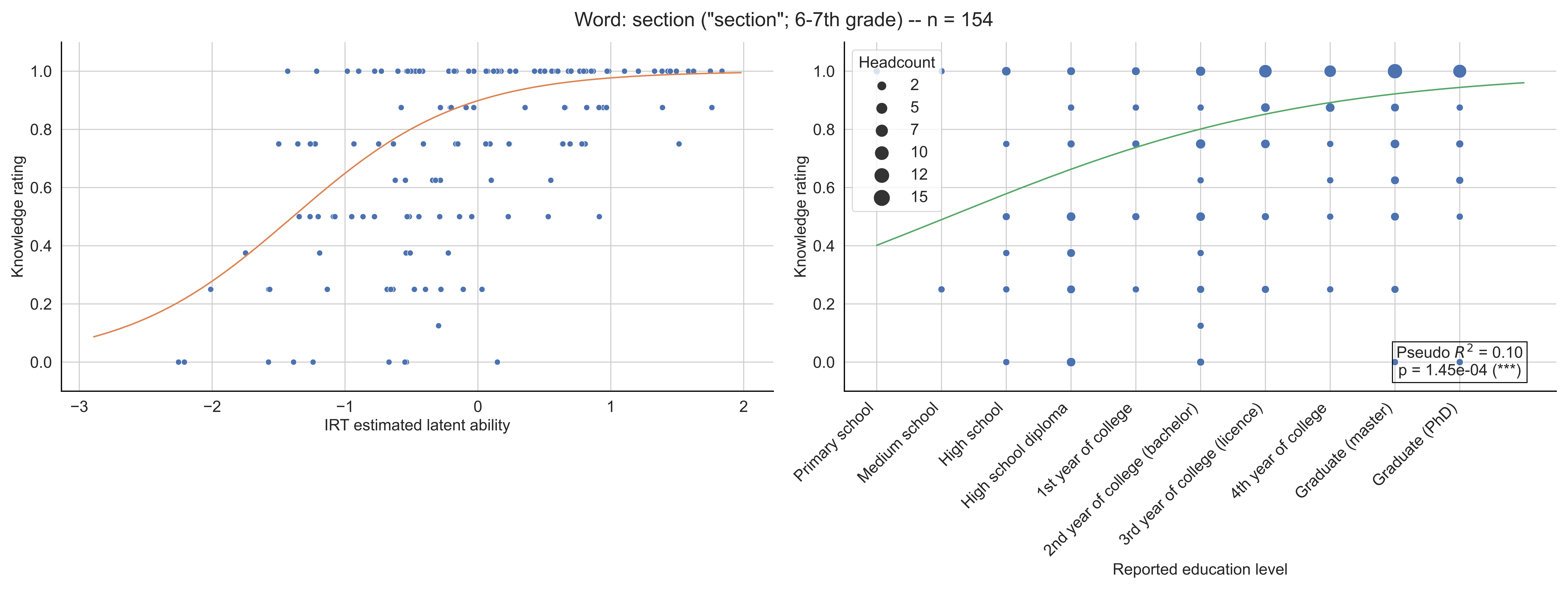

Word: section ("section"; 6-7th grade) -- n = 154
Knowledge rating
IRT estimated latent ability
Headcount
2
5
7
10
12
15
Pseudo $R^2$ = 0.10
p = 1.45e-04 (***)
Primary school
Medium school
High school
High school diploma
1st year of college
2nd year of college (bachelor)
3rd year of college (licence)
4th year of college
Graduate (master)
Graduate (PhD)
Reported education level

Word: sept ("seven"; Primary school) -- n = 157

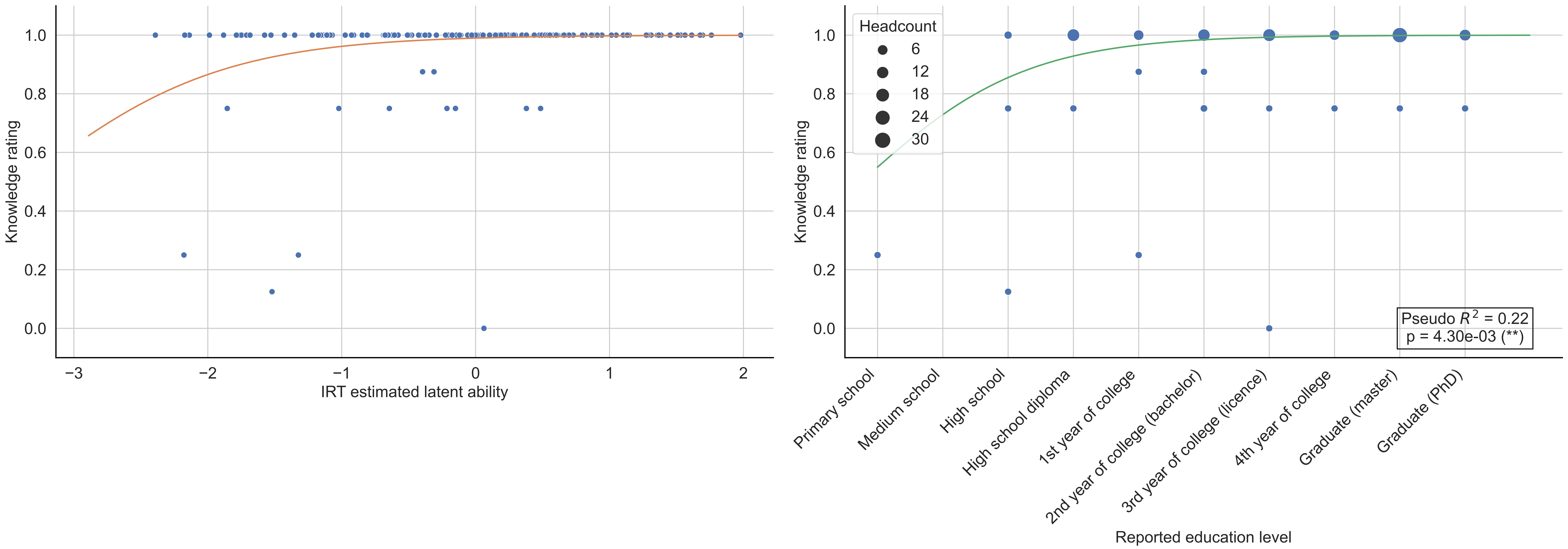

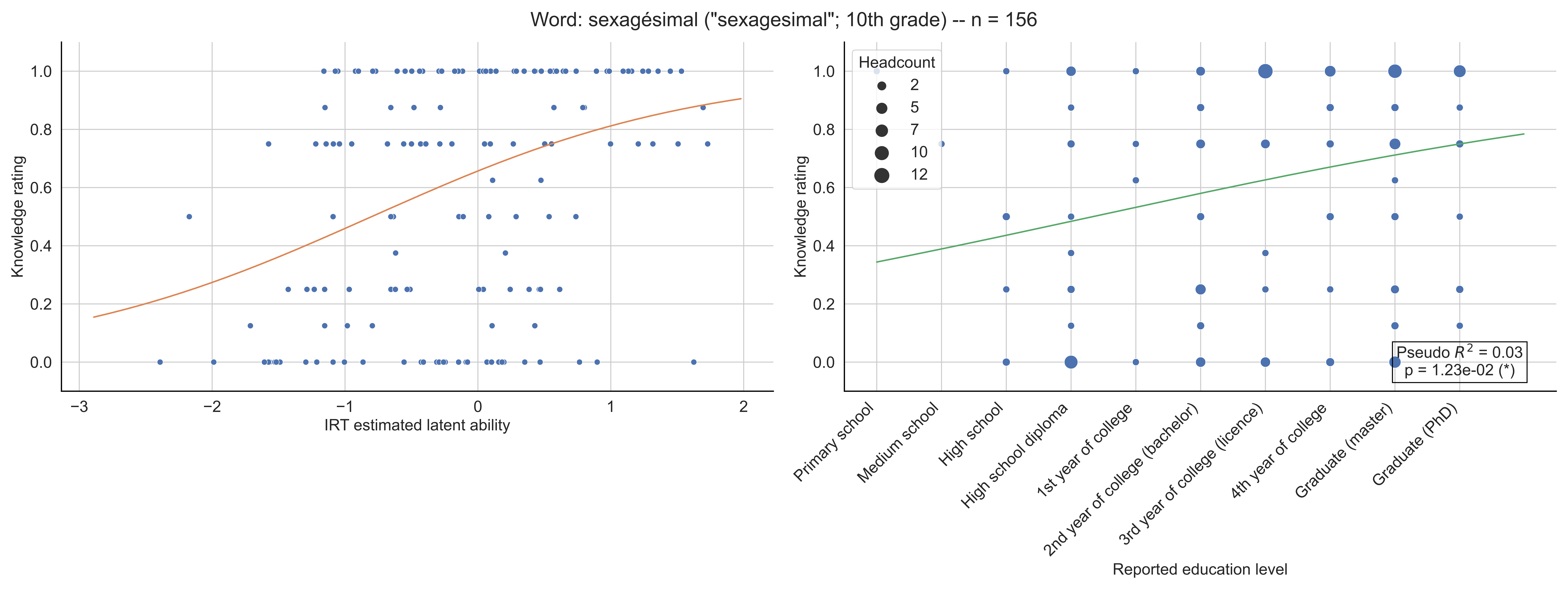
Word: sexagésimal ("sexagesimal"; 10th grade) -- n = 156
Knowledge rating
IRT estimated latent ability
Headcount
2
5
7
10
12
Pseudo $R^2$ = 0.03
p = 1.23e-02 (*)
Primary school
Medium school
High school
High school diploma
1st year of college
2nd year of college (bachelor)
3rd year of college (licence)
4th year of college
Graduate (master)
Graduate (PhD)
Reported education level

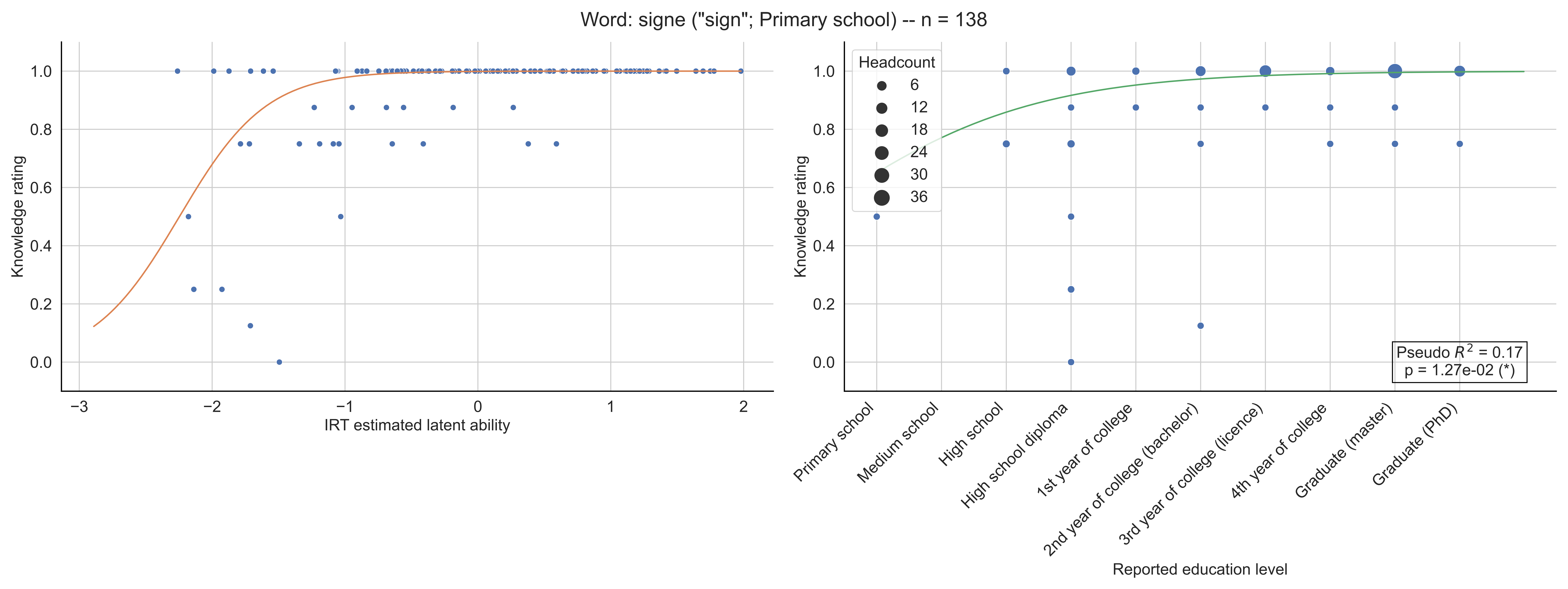

Word: signe ("sign"; Primary school) -- n = 138
Knowledge rating
IRT estimated latent ability
Headcount
6
12
18
24
30
36
Pseudo $R^2$ = 0.17
p = 1.27e-02 (*)
Primary school
Medium school
High school
High school diploma
1st year of college
2nd year of college (bachelor)
3rd year of college (licence)
4th year of college
Graduate (master)
Graduate (PhD)
Reported education level

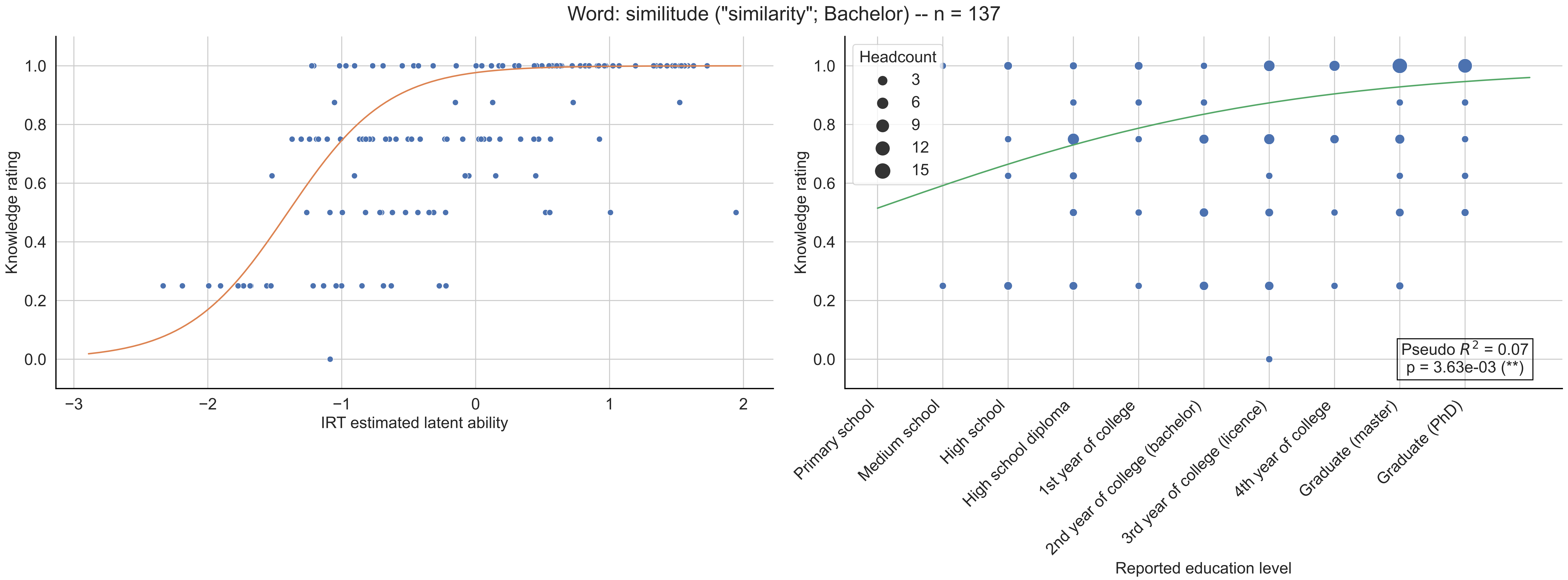

Word: similitude ("similarity"; Bachelor) -- n = 137
Knowledge rating
IRT estimated latent ability
Headcount
3
6
9
12
15
Pseudo $R^2$ = 0.07
p = 3.63e-03 (**)
Primary school
Medium school
High school
High school diploma
1st year of college
2nd year of college (bachelor)
3rd year of college (licence)
4th year of college
Graduate (master)
Graduate (PhD)
Reported education level

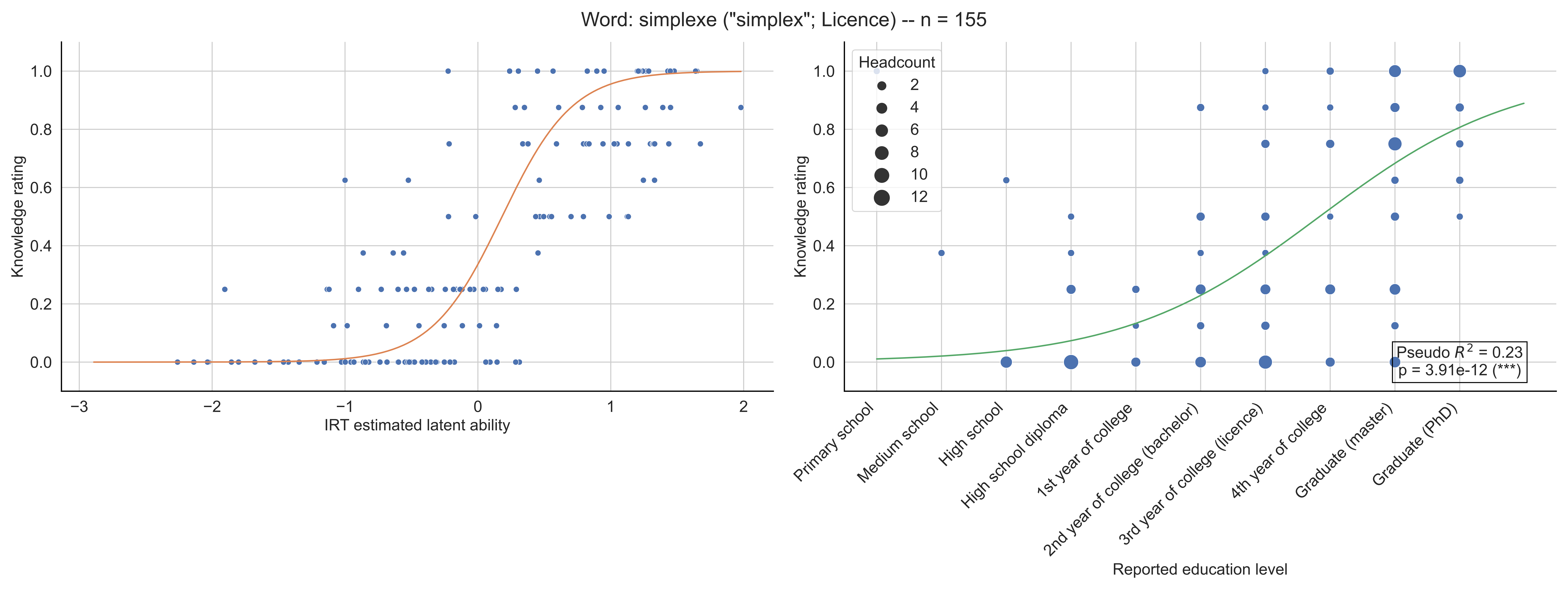

Word: simplexe ("simplex"; Licence) -- n = 155
Knowledge rating
IRT estimated latent ability
Headcount
2
4
6
8
10
12
Pseudo $R^2$ = 0.23
p = 3.91e-12 (***)
Primary school
Medium school
High school
High school diploma
1st year of college
2nd year of college (bachelor)
3rd year of college (licence)
4th year of college
Graduate (master)
Graduate (PhD)
Reported education level

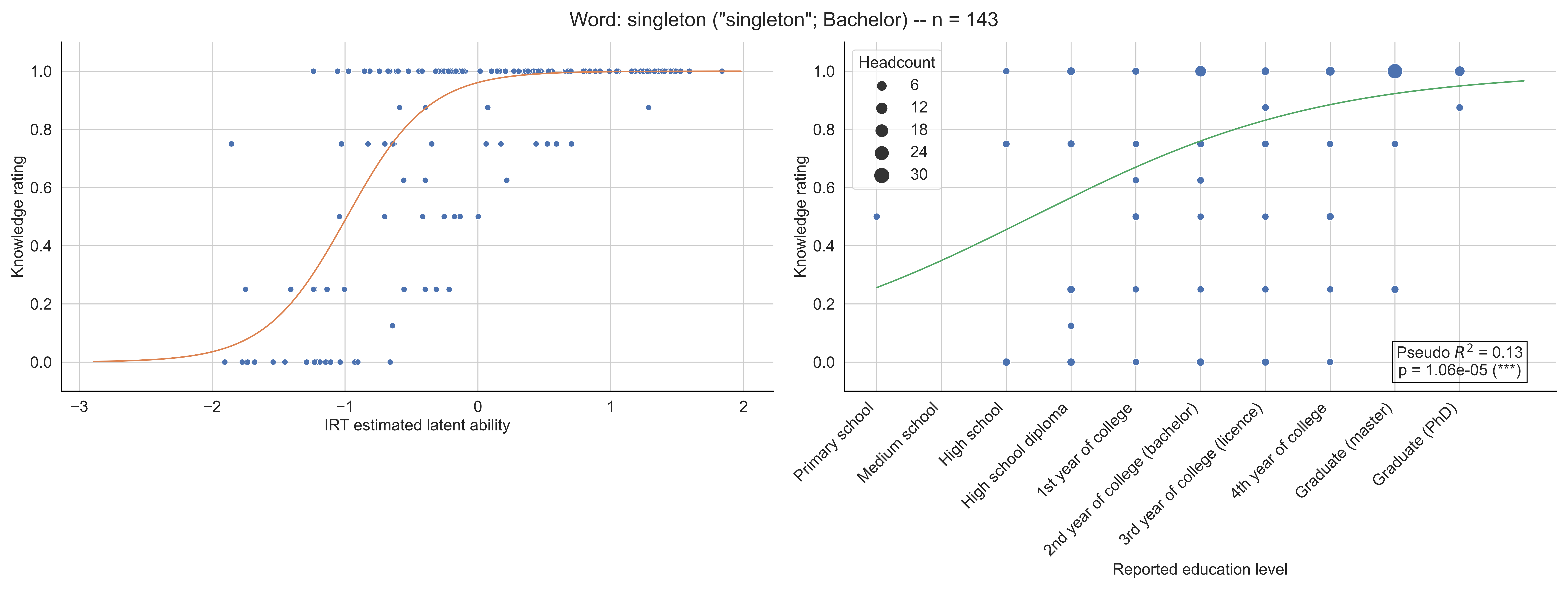

Word: singleton ("singleton"; Bachelor) -- n = 143
Knowledge rating
IRT estimated latent ability
Headcount
6
12
18
24
30
Pseudo $R^2$ = 0.13
p = 1.06e-05 (***)
Primary school
Medium school
High school
High school diploma
1st year of college
2nd year of college (bachelor)
3rd year of college (licence)
4th year of college
Graduate (master)
Graduate (PhD)
Reported education level

Word: singularité ("singularity"; Licence) -- n = 163

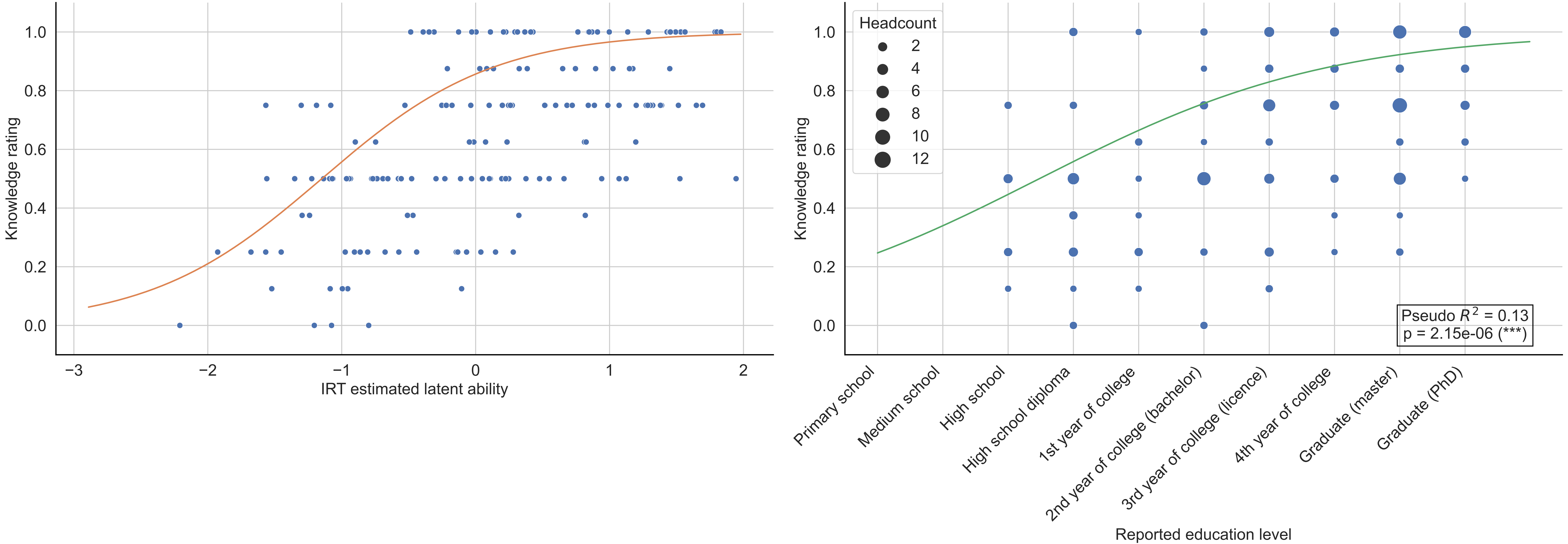

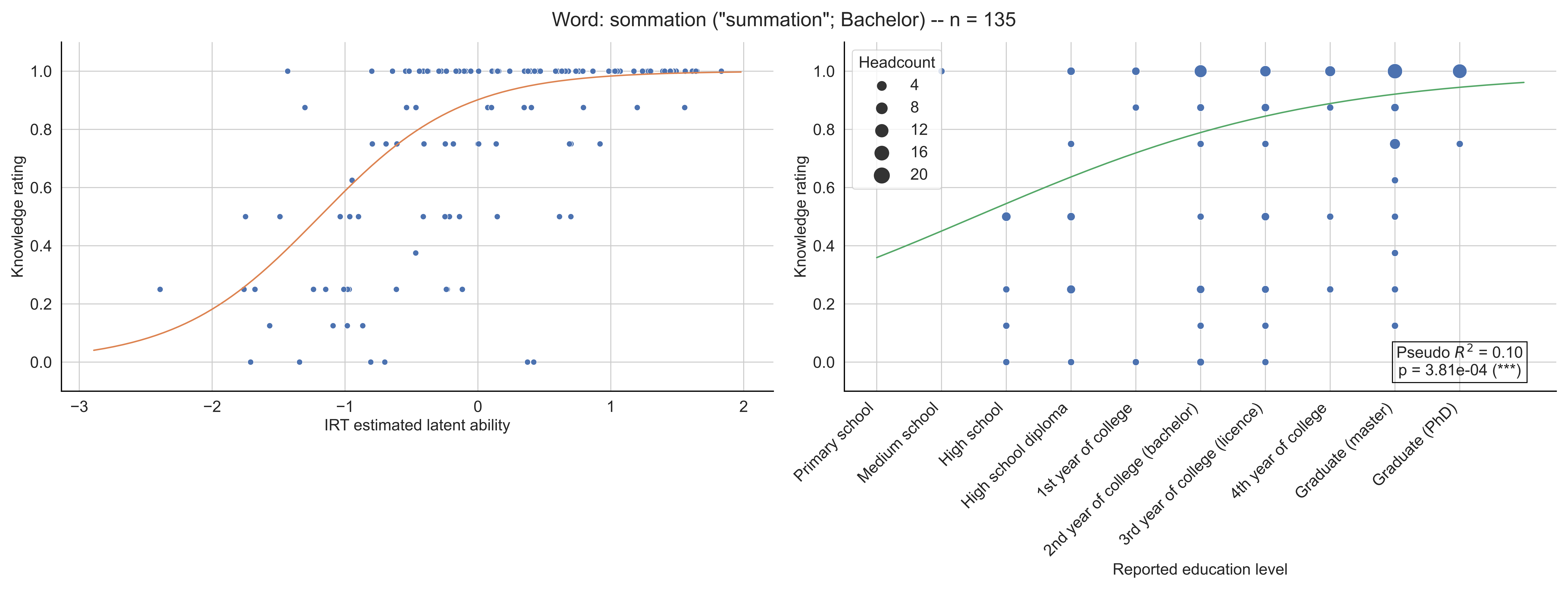

Word: sommation ("summation"; Bachelor) -- n = 135
Knowledge rating
IRT estimated latent ability
Headcount
4
8
12
16
20
Pseudo $R^2$ = 0.10
p = 3.81e-04 (***)
Primary school
Medium school
High school
High school diploma
1st year of college
2nd year of college (bachelor)
3rd year of college (licence)
4th year of college
Graduate (master)
Graduate (PhD)
Reported education level

Word: sommet ("vertice"; 6-7th grade) -- n = 148

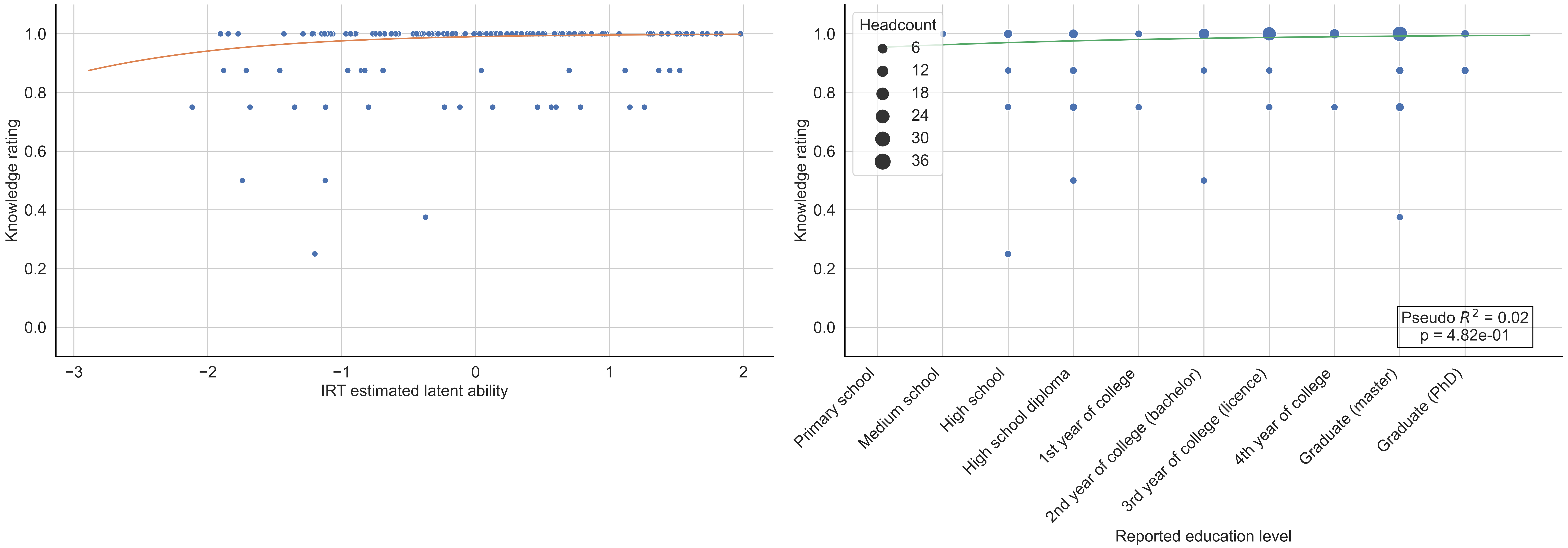

Word: source ("source"; Bachelor) -- n = 137

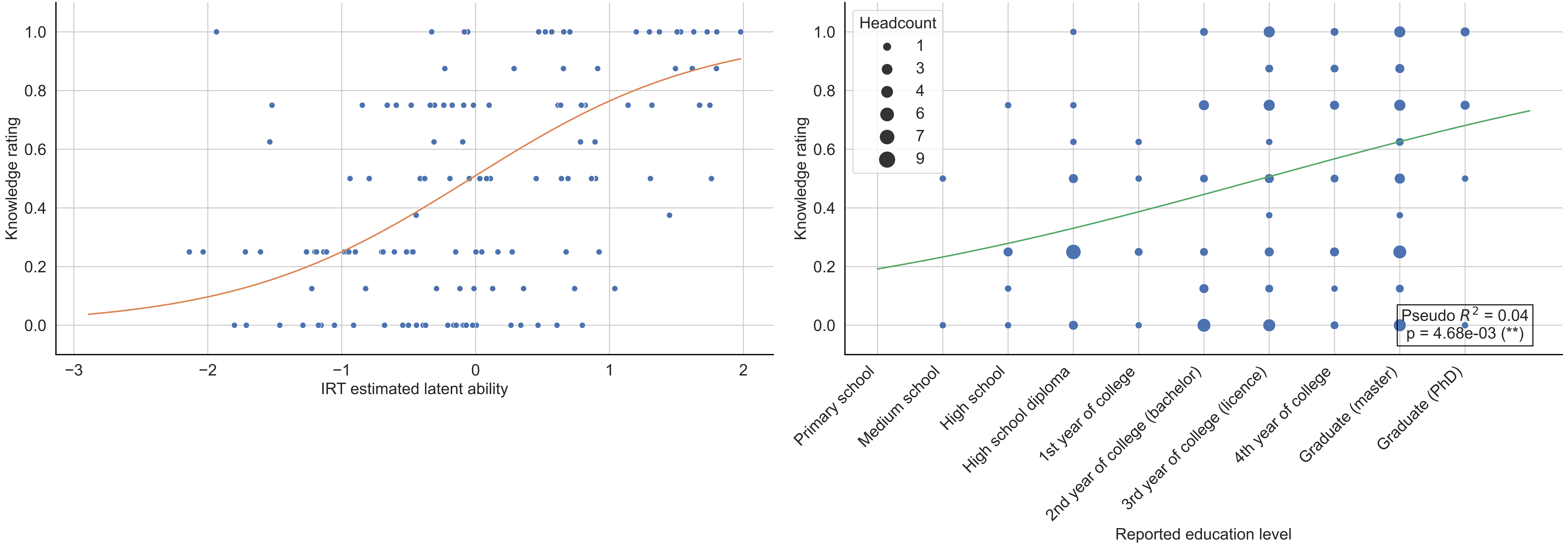

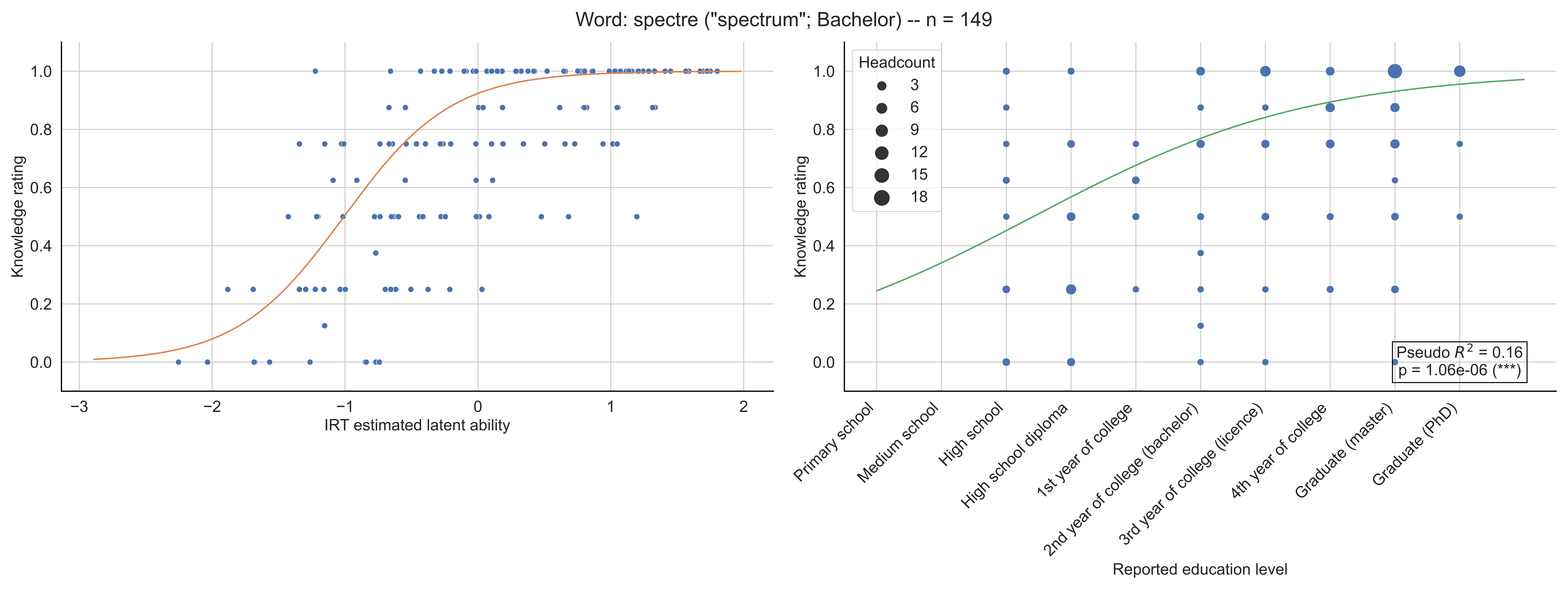

Word: spectre ("spectrum"; Bachelor) -- n = 149
Knowledge rating
IRT estimated latent ability
Headcount
3
6
9
12
15
18
Pseudo $R^2$ = 0.16
p = 1.06e-06 (***)
Primary school
Medium school
High school
High school diploma
1st year of college
2nd year of college (bachelor)
3rd year of college (licence)
4th year of college
Graduate (master)
Graduate (PhD)
Reported education level

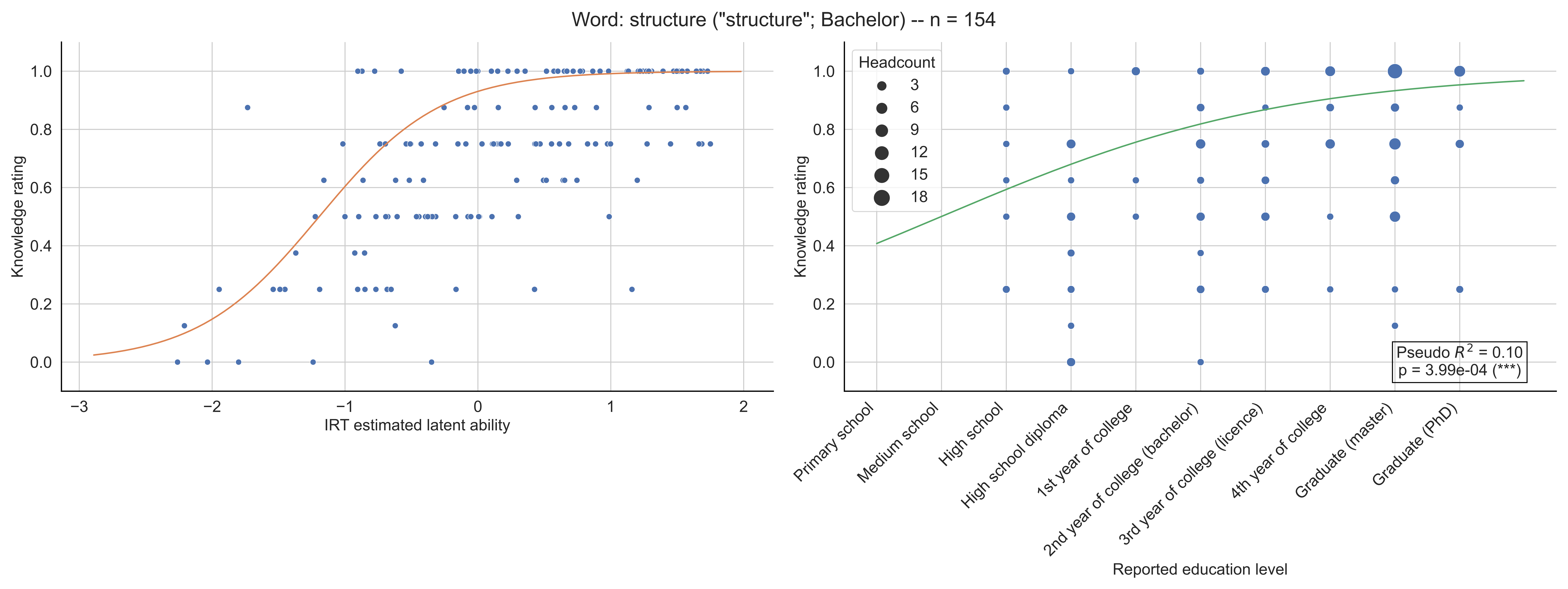

Word: structure ("structure"; Bachelor) -- n = 154
Knowledge rating
IRT estimated latent ability
Headcount
3
6
9
12
15
18
Pseudo $R^2$ = 0.10
p = 3.99e-04 (***)
Primary school
Medium school
High school
High school diploma
1st year of college
2nd year of college (bachelor)
3rd year of college (licence)
4th year of college
Graduate (master)
Graduate (PhD)
Reported education level

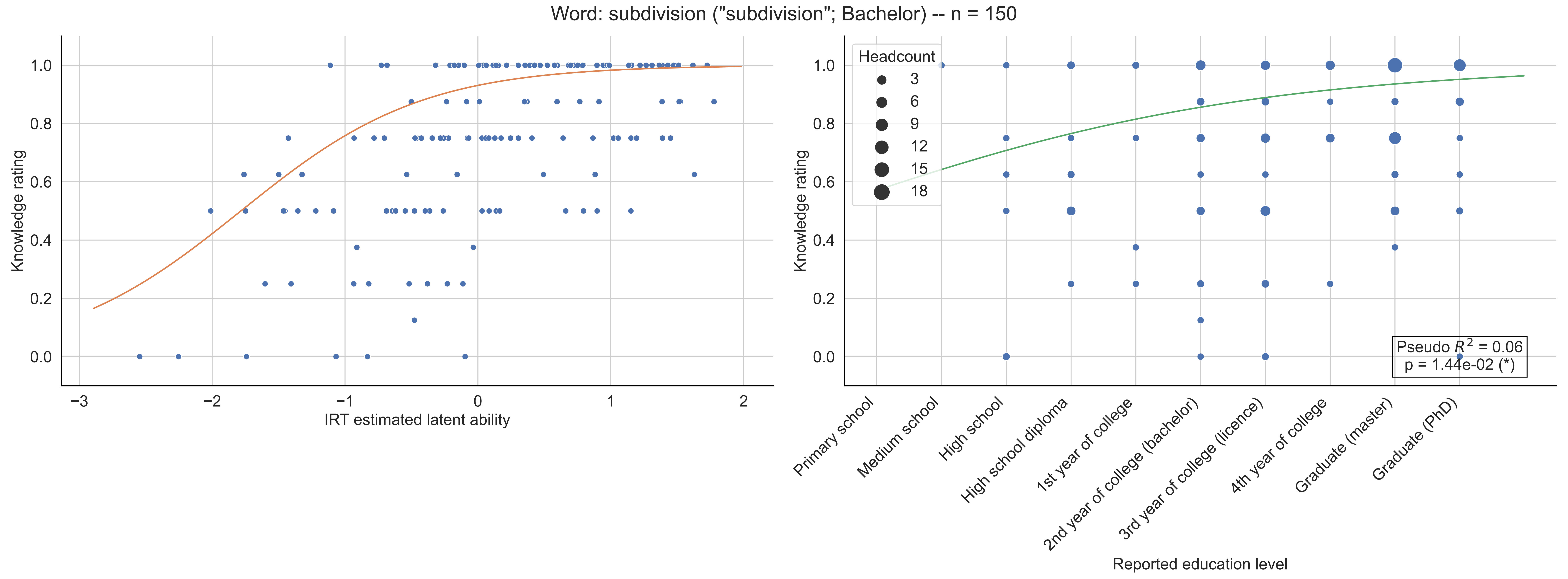

Word: subdivision ("subdivision"; Bachelor) -- n = 150
Knowledge rating
IRT estimated latent ability
Headcount
3
6
9
12
15
18
Pseudo $R^2$ = 0.06
p = 1.44e-02 (*)
Primary school
Medium school
High school
High school diploma
1st year of college
2nd year of college (bachelor)
3rd year of college (licence)
4th year of college
Graduate (master)
Graduate (PhD)
Reported education level

Word: supplémentaire ("complementary"; Bachelor) -- n = 122

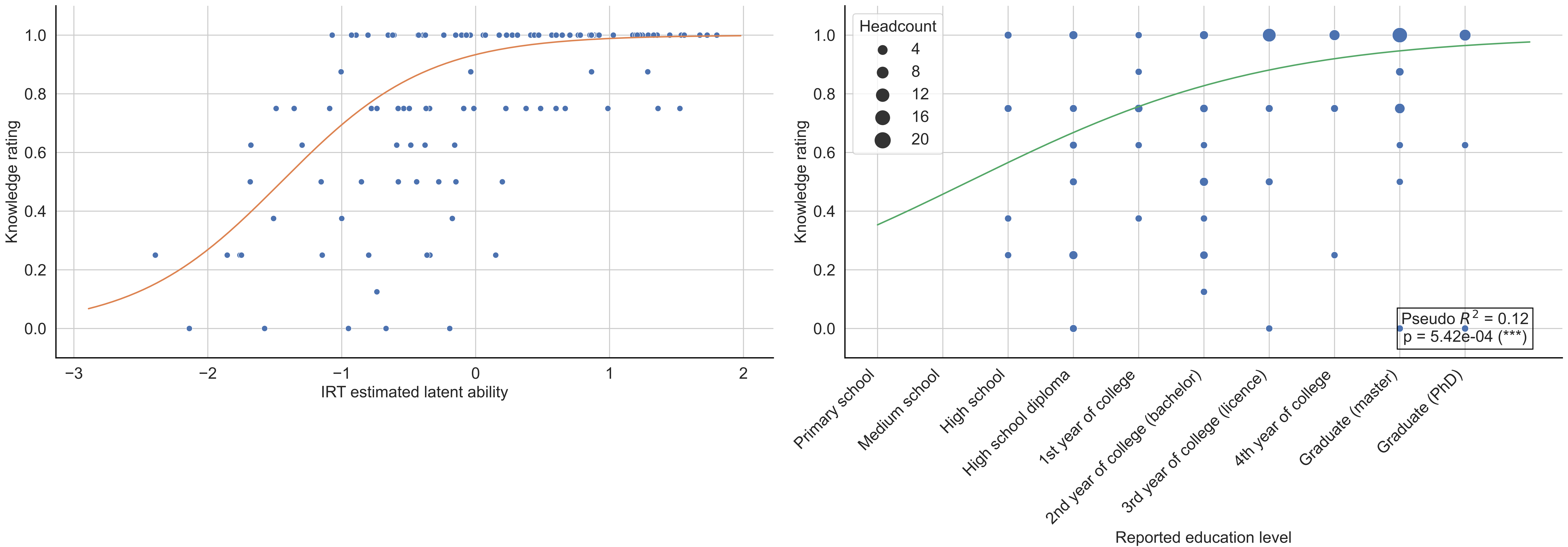

Word: support ("support"; Bachelor) -- n = 142

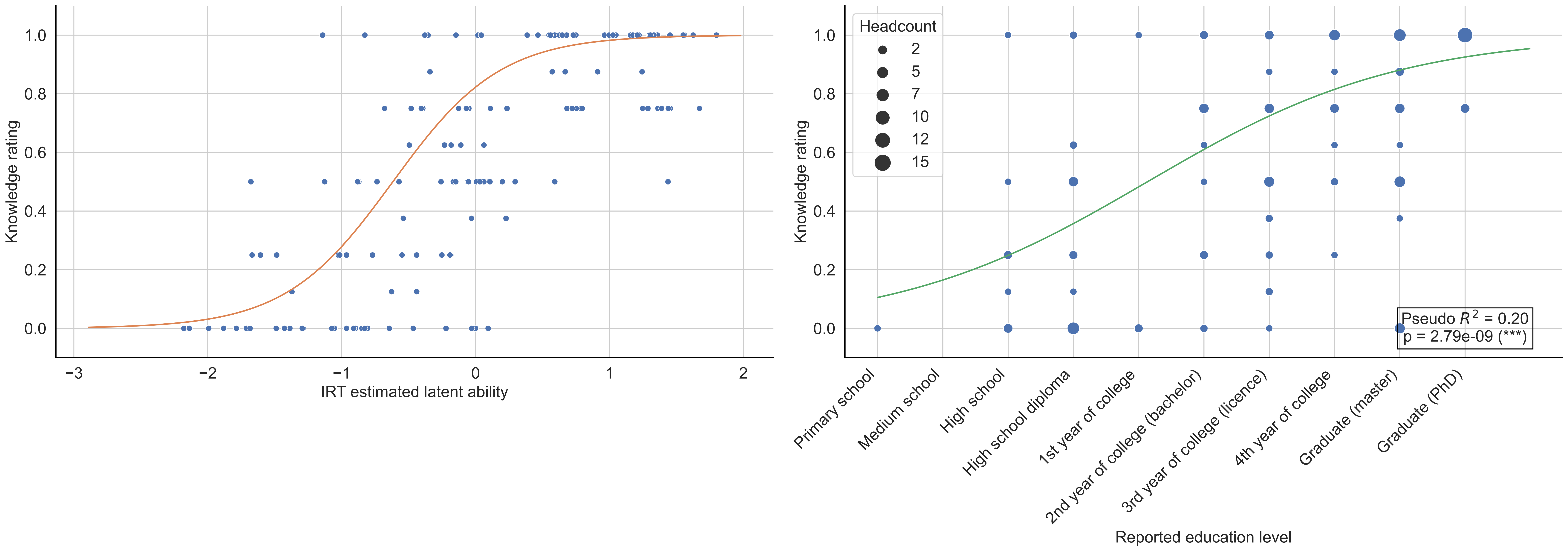

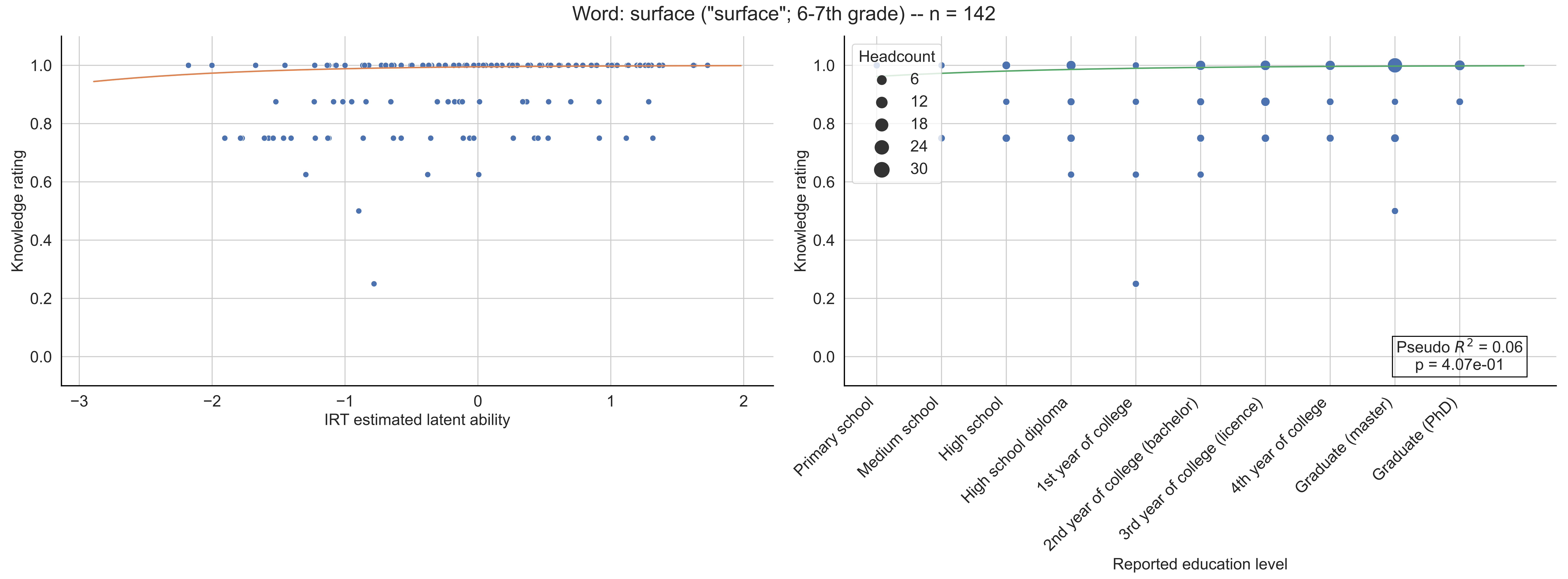
Word: surface ("surface"; 6-7th grade) -- n = 142
Knowledge rating
IRT estimated latent ability
Headcount
6
12
18
24
30
Pseudo $R^2$ = 0.06
p = 4.07e-01
Primary school
Medium school
High school
High school diploma
1st year of college
2nd year of college (bachelor)
3rd year of college (licence)
4th year of college
Graduate (master)
Graduate (PhD)
Reported education level

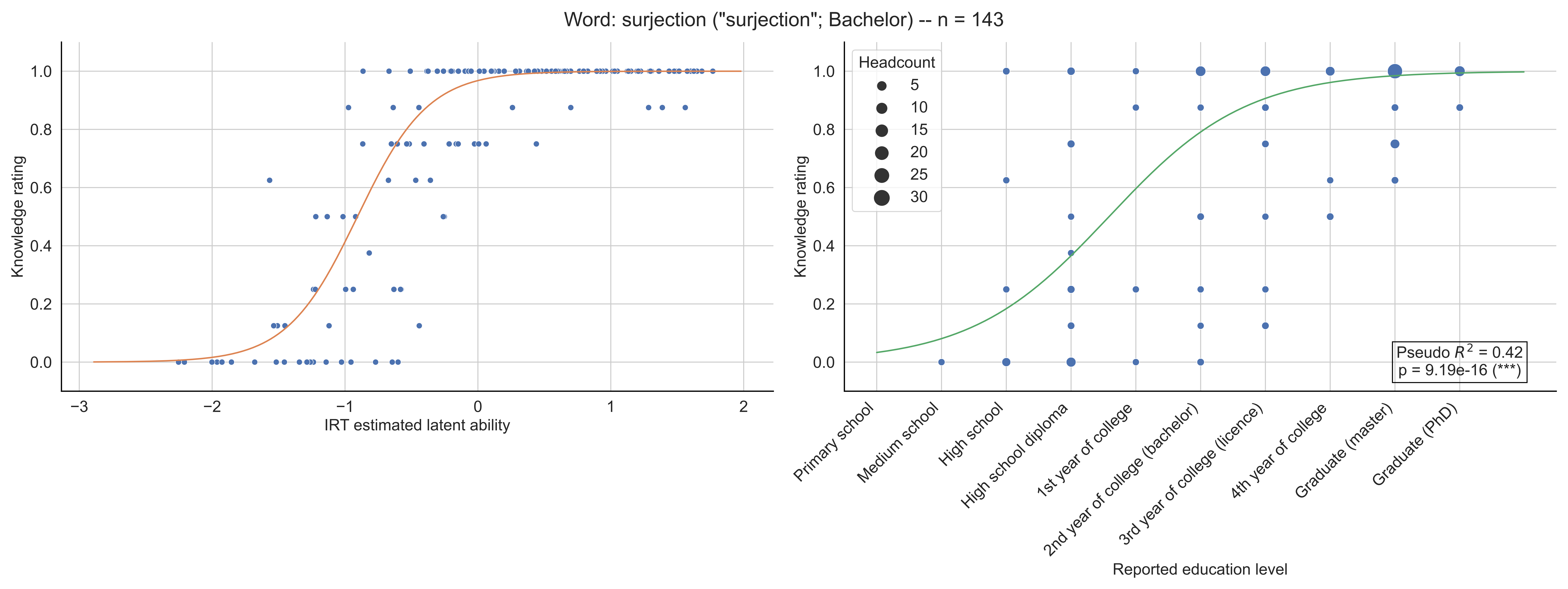
Word: surjection ("surjection"; Bachelor) -- n = 143
Knowledge rating
IRT estimated latent ability
Headcount
5
10
15
20
25
30
Pseudo $R^2$ = 0.42
p = 9.19e-16 (***)
Primary school
Medium school
High school
High school diploma
1st year of college
2nd year of college (bachelor)
3rd year of college (licence)
4th year of college
Graduate (master)
Graduate (PhD)
Reported education level

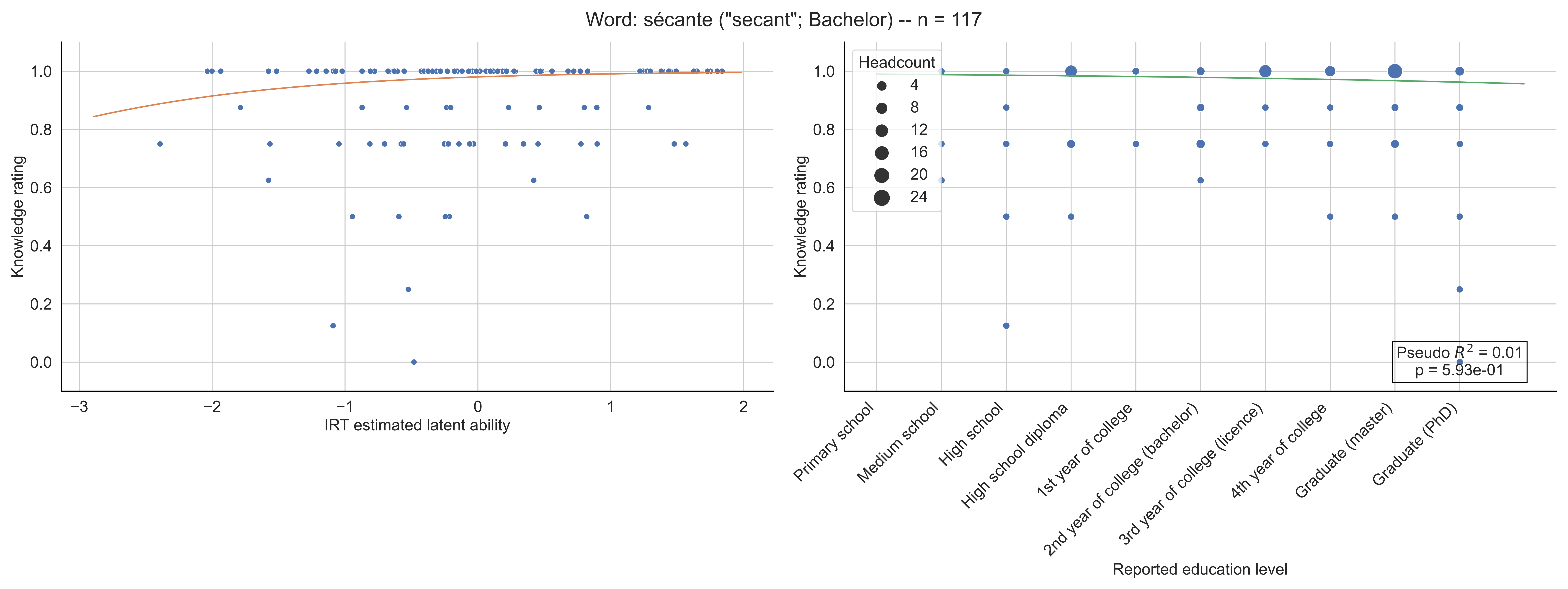

Word: sécante ("secant"; Bachelor) -- n = 117
Knowledge rating
IRT estimated latent ability
Headcount
4
8
12
16
20
24
Pseudo $R^2$ = 0.01
p = 5.93e-01
Primary school
Medium school
High school
High school diploma
1st year of college
2nd year of college (bachelor)
3rd year of college (licence)
4th year of college
Graduate (master)
Graduate (PhD)
Reported education level

Word: séparation ("separation"; Licence) -- n = 144

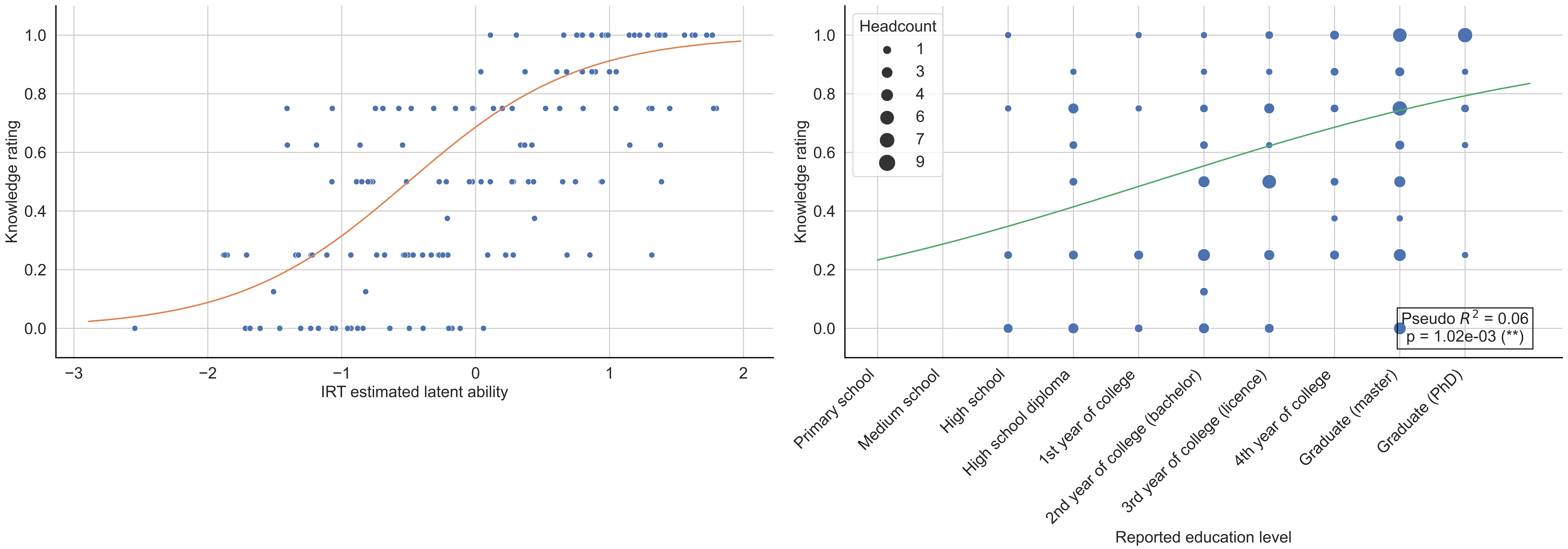

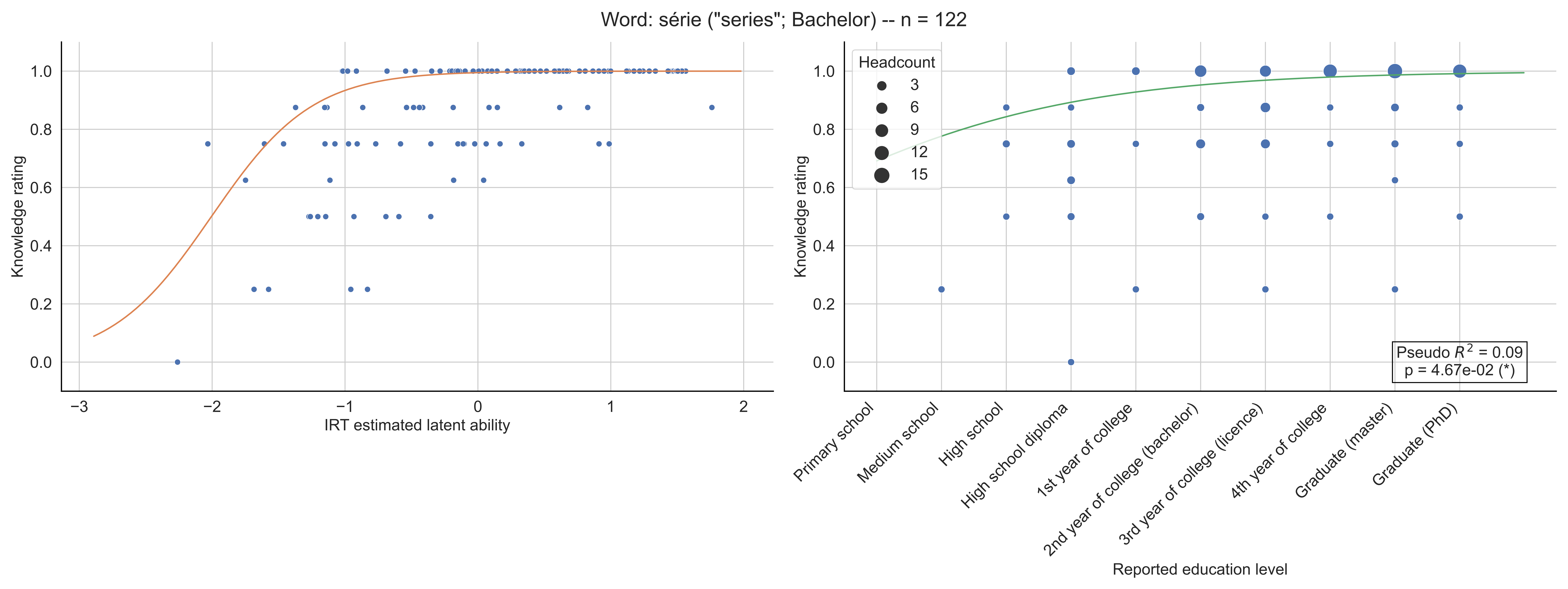

Word: série ("series"; Bachelor) -- n = 122
Knowledge rating
IRT estimated latent ability
Headcount
3
6
9
12
15
Pseudo $R^2$ = 0.09
p = 4.67e-02 (*)
Primary school
Medium school
High school
High school diploma
1st year of college
2nd year of college (bachelor)
3rd year of college (licence)
4th year of college
Graduate (master)
Graduate (PhD)
Reported education level

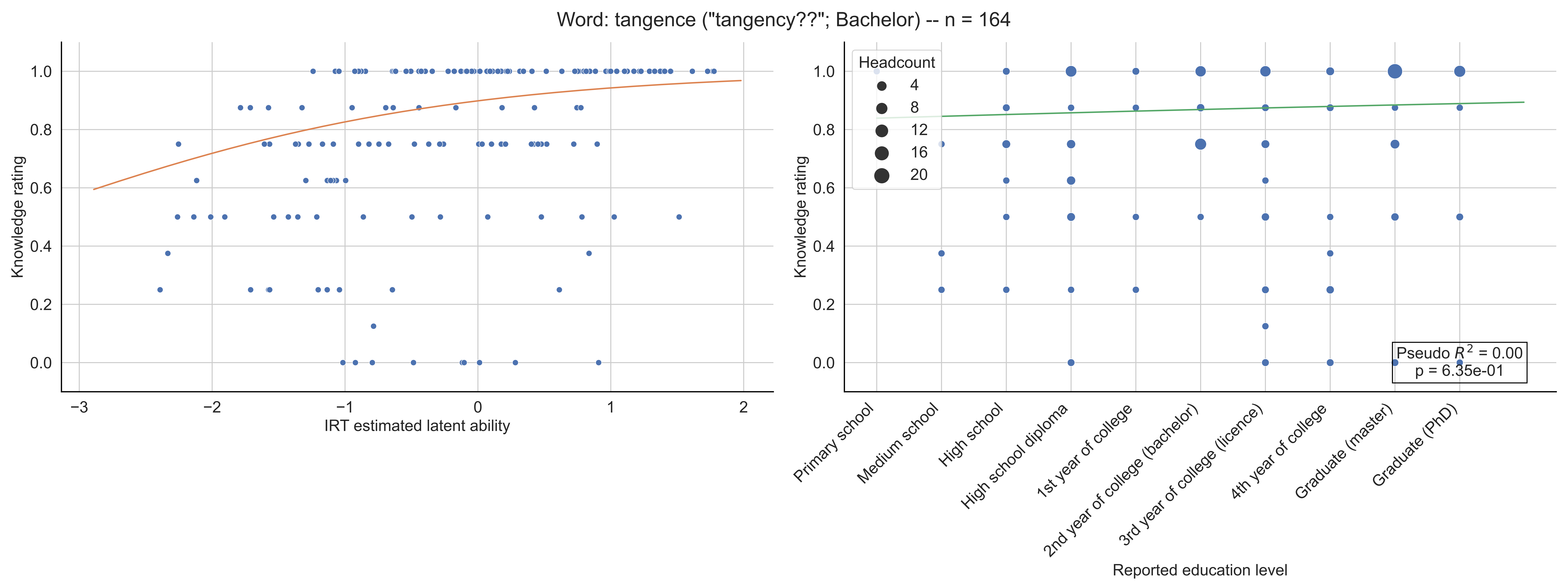

Word: tangence ("tangency??"; Bachelor) -- n = 164
Knowledge rating
IRT estimated latent ability
Headcount
4
8
12
16
20
Pseudo $R^2$ = 0.00
p = 6.35e-01
Primary school
Medium school
High school
High school diploma
1st year of college
2nd year of college (bachelor)
3rd year of college (licence)
4th year of college
Graduate (master)
Graduate (PhD)
Reported education level

Word: tangente ("tangent"; 6-7th grade) -- n = 132

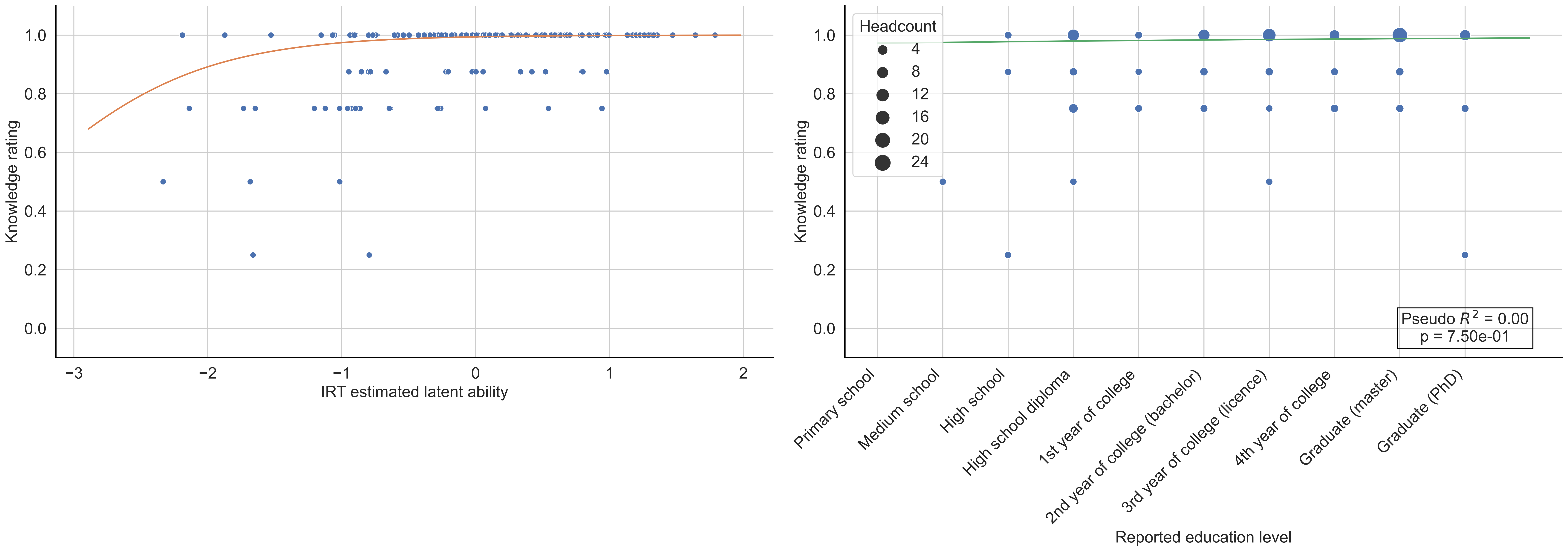

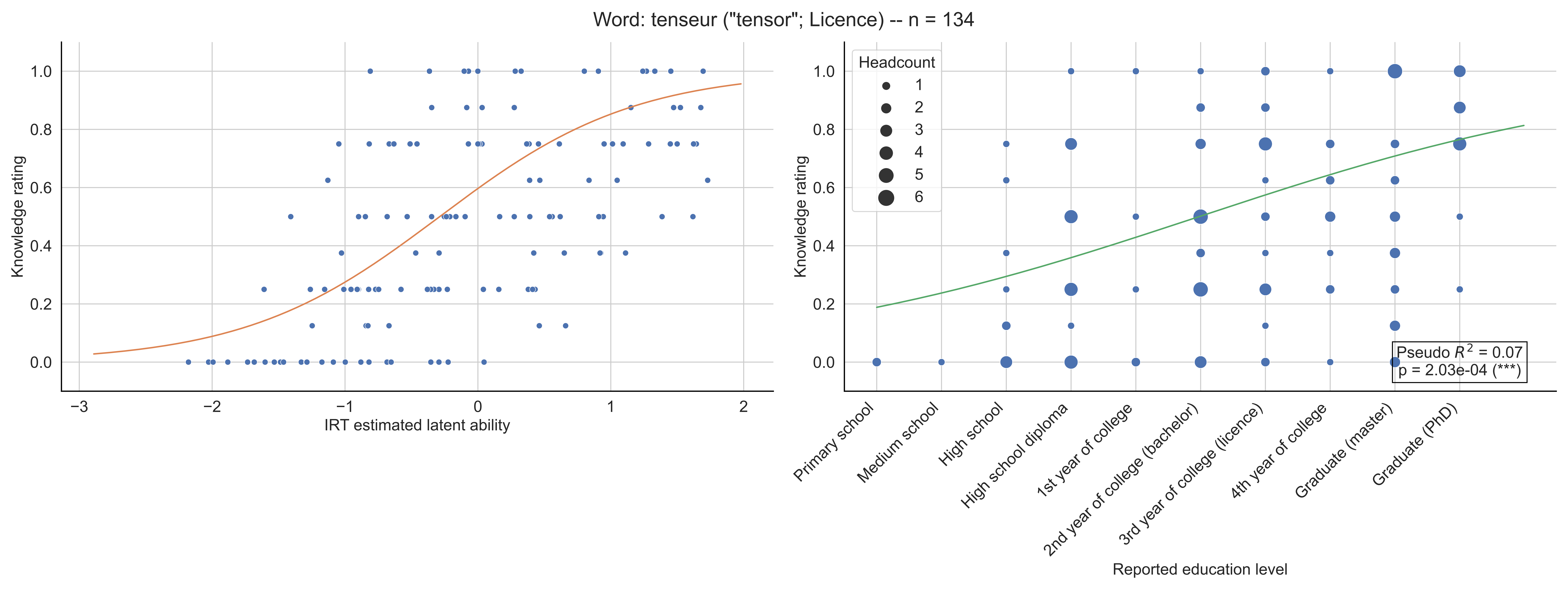

Word: tenseur ("tensor"; Licence) -- n = 134
Knowledge rating
IRT estimated latent ability
Headcount
1
2
3
4
5
6
Pseudo $R^2$ = 0.07
p = 2.03e-04 (***)
Primary school
Medium school
High school
High school diploma
1st year of college
2nd year of college (bachelor)
3rd year of college (licence)
4th year of college
Graduate (master)
Graduate (PhD)
Reported education level

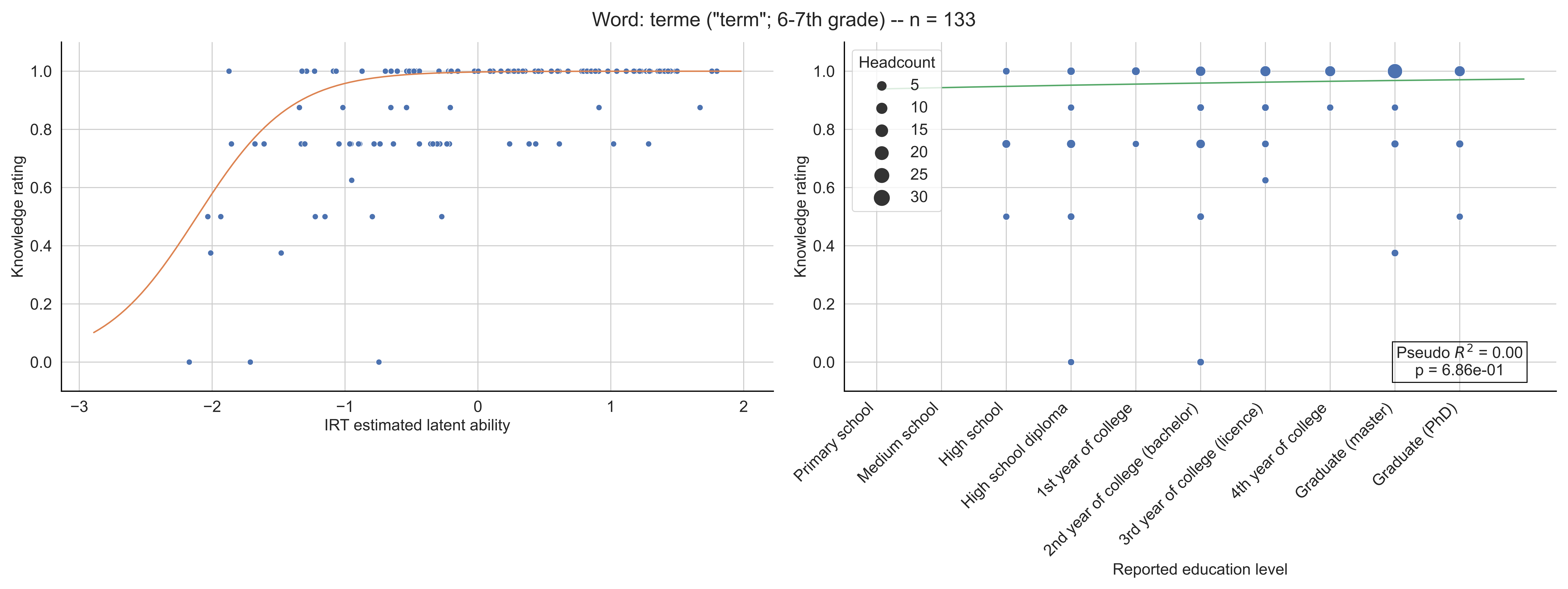

Word: terme ("term"; 6-7th grade) -- n = 133
Knowledge rating
IRT estimated latent ability
Headcount
5
10
15
20
25
30
Pseudo $R^2$ = 0.00
p = 6.86e-01
Primary school
Medium school
High school
High school diploma
1st year of college
2nd year of college (bachelor)
3rd year of college (licence)
4th year of college
Graduate (master)
Graduate (PhD)
Reported education level

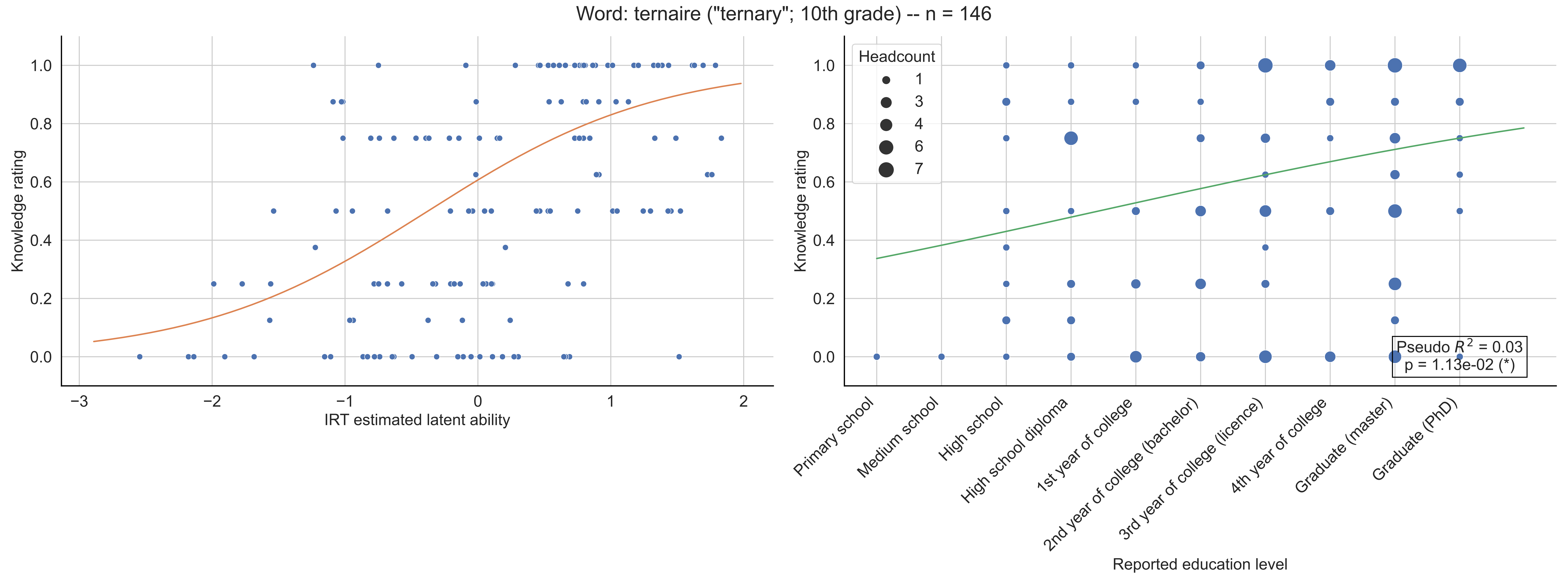
Word: ternaire ("ternary"; 10th grade) -- n = 146
Knowledge rating
IRT estimated latent ability
Headcount
1
3
4
6
7
Pseudo $R^2$ = 0.03
p = 1.13e-02 (*)
Primary school
Medium school
High school
High school diploma
1st year of college
2nd year of college (bachelor)
3rd year of college (licence)
4th year of college
Graduate (master)
Graduate (PhD)
Reported education level

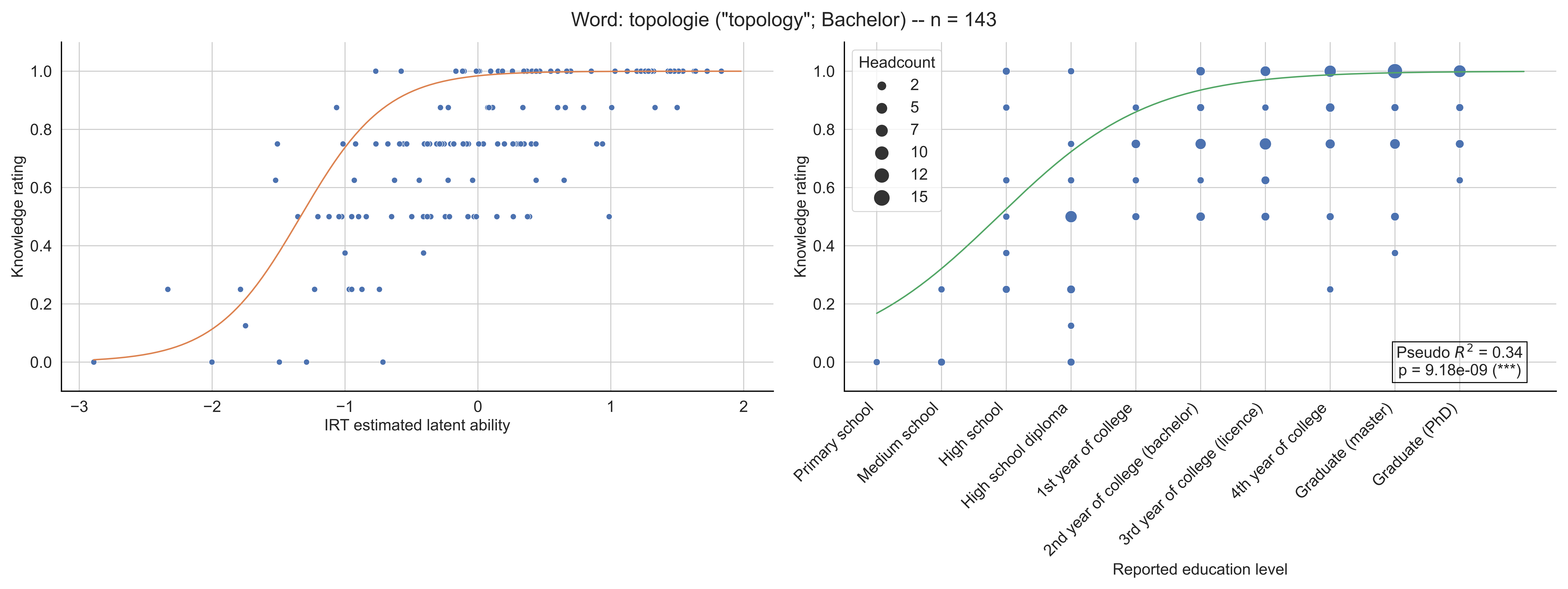

Word: topologie ("topology"; Bachelor) -- n = 143
Knowledge rating
IRT estimated latent ability
Headcount
2
5
7
10
12
15
Pseudo $R^2$ = 0.34
p = 9.18e-09 (***)
Primary school
Medium school
High school
High school diploma
1st year of college
2nd year of college (bachelor)
3rd year of college (licence)
4th year of college
Graduate (master)
Graduate (PhD)
Reported education level

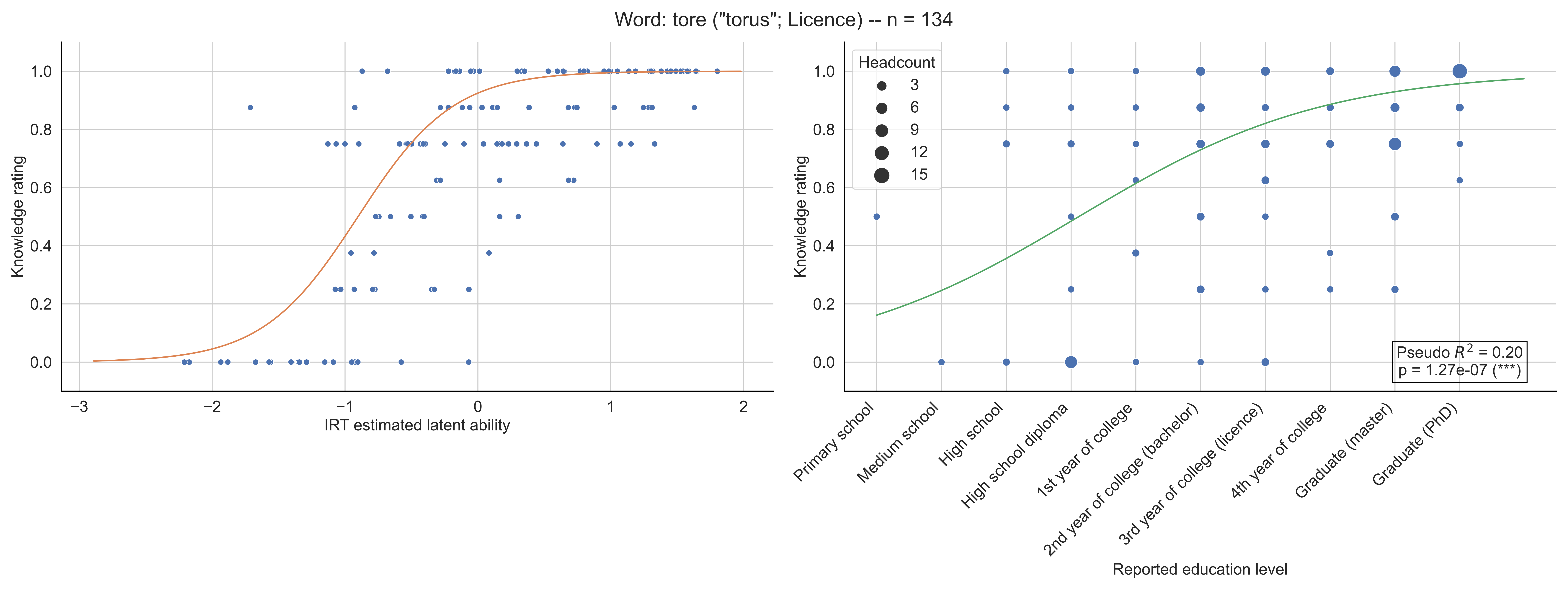

Word: tore ("torus"; Licence) -- n = 134
Knowledge rating
IRT estimated latent ability
Headcount
3
6
9
12
15
Pseudo $R^2$ = 0.20
p = 1.27e-07 (***)
Primary school
Medium school
High school
High school diploma
1st year of college
2nd year of college (bachelor)
3rd year of college (licence)
4th year of college
Graduate (master)
Graduate (PhD)
Reported education level

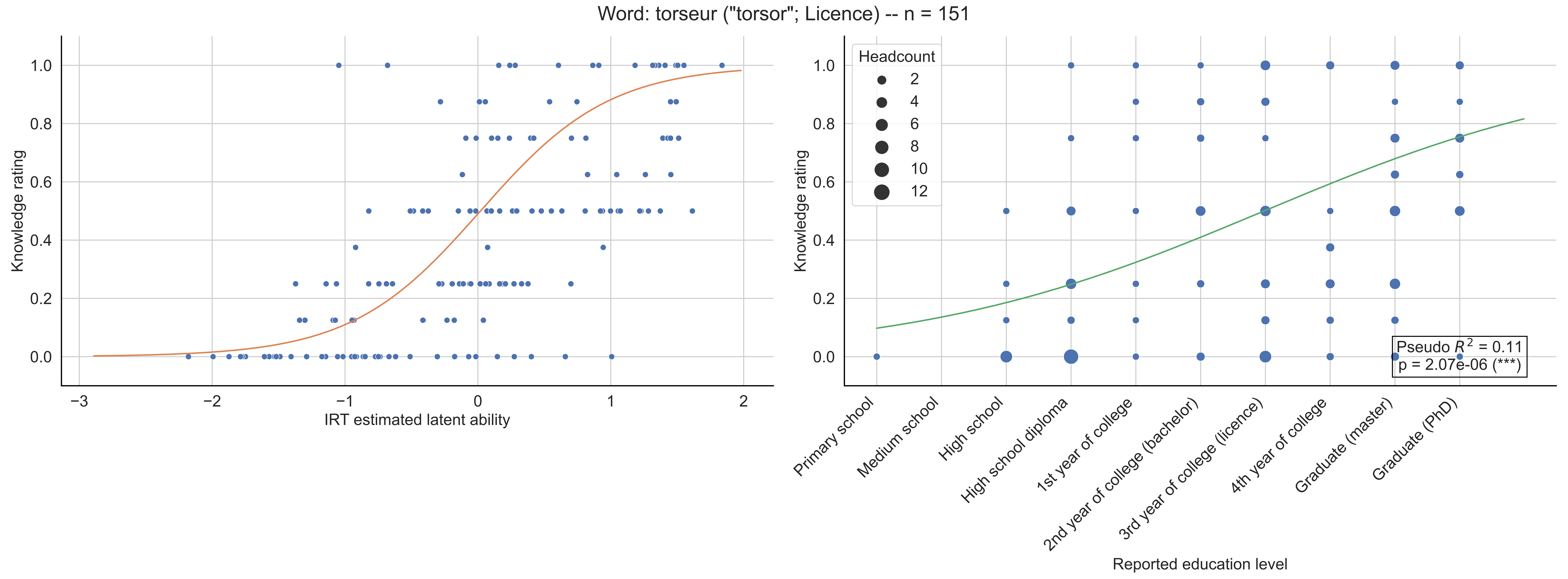

Word: torseur ("torsor"; Licence) -- n = 151
Knowledge rating
IRT estimated latent ability
Headcount
2
4
6
8
10
12
Pseudo $R^2$ = 0.11
p = 2.07e-06 (***)
Primary school
Medium school
High school
High school diploma
1st year of college
2nd year of college (bachelor)
3rd year of college (licence)
4th year of college
Graduate (master)
Graduate (PhD)
Reported education level

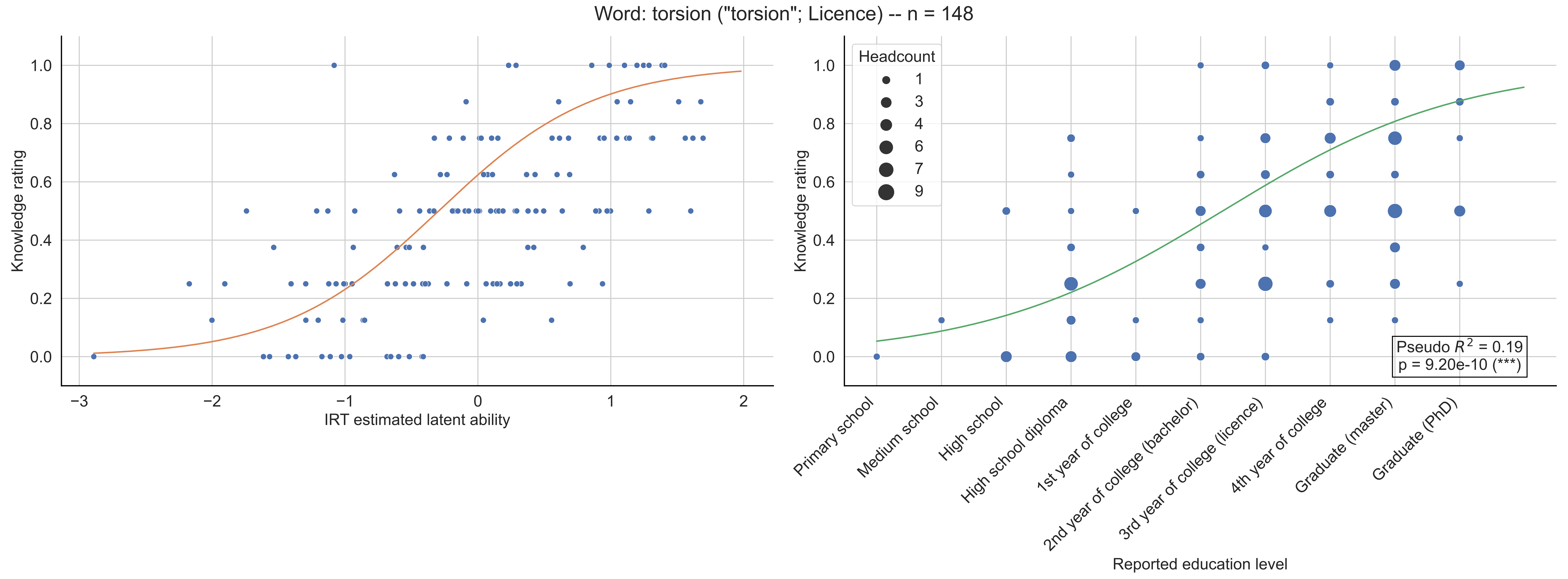

Word: torsion ("torsion"; Licence) -- n = 148
Knowledge rating
IRT estimated latent ability
Headcount
1
3
4
6
7
9
Pseudo $R^2$ = 0.19
p = 9.20e-10 (***)
Primary school
Medium school
High school
High school diploma
1st year of college
2nd year of college (bachelor)
3rd year of college (licence)
4th year of college
Graduate (master)
Graduate (PhD)
Reported education level

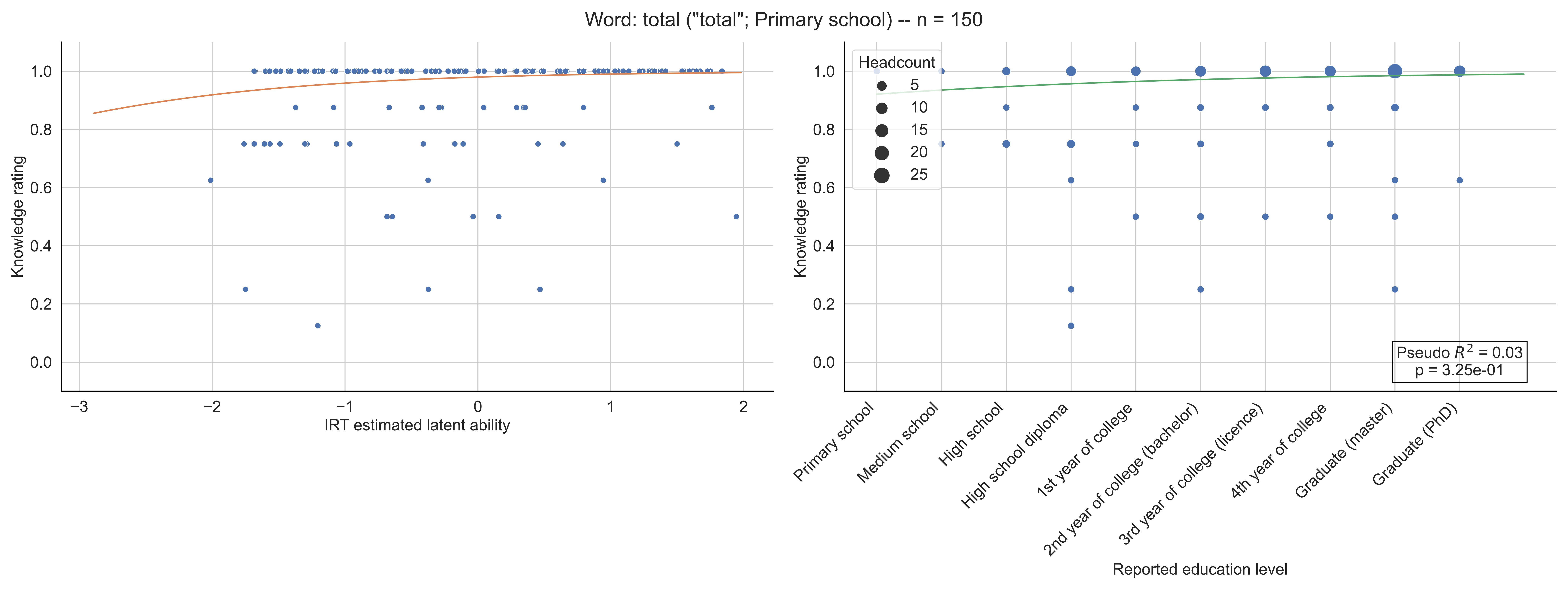

Word: total ("total"; Primary school) -- n = 150
Knowledge rating
IRT estimated latent ability
Headcount
5
10
15
20
25
Pseudo $R^2$ = 0.03
p = 3.25e-01
Primary school
Medium school
High school
High school diploma
1st year of college
2nd year of college (bachelor)
3rd year of college (licence)
4th year of college
Graduate (master)
Graduate (PhD)
Reported education level

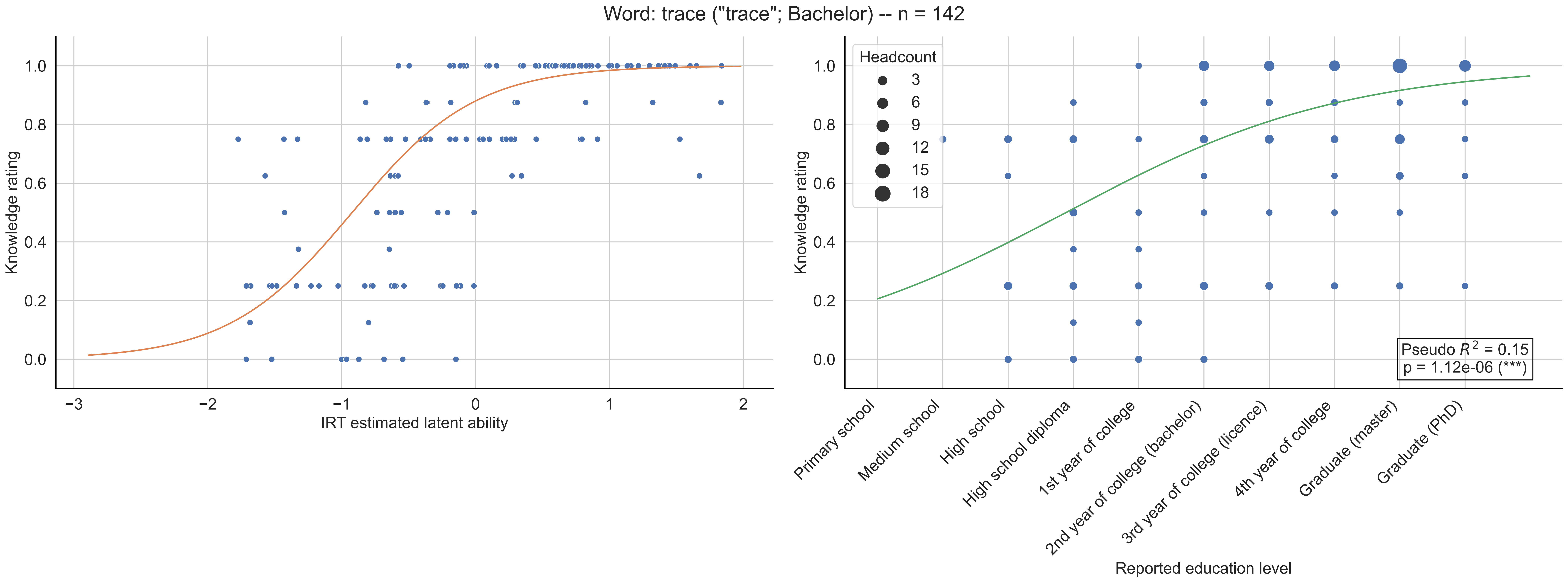

Word: trace ("trace"; Bachelor) -- n = 142
Knowledge rating
IRT estimated latent ability
Headcount
3
6
9
12
15
18
Pseudo $R^2$ = 0.15
p = 1.12e-06 (***)
Primary school
Medium school
High school
High school diploma
1st year of college
2nd year of college (bachelor)
3rd year of college (licence)
4th year of college
Graduate (master)
Graduate (PhD)
Reported education level

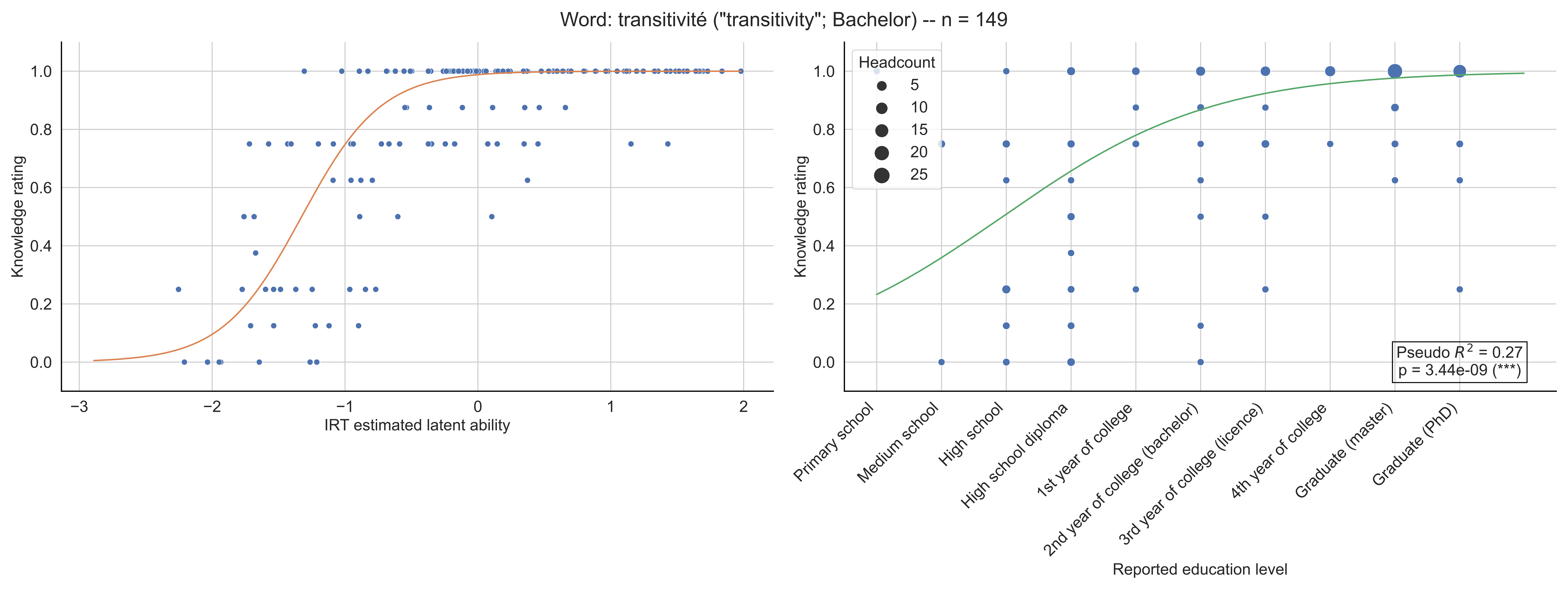

Word: transitivité ("transitivity"; Bachelor) -- n = 149
Knowledge rating
IRT estimated latent ability
Headcount
5
10
15
20
25
Pseudo $R^2$ = 0.27
p = 3.44e-09 (***)
Reported education level
Primary school
Medium school
High school
High school diploma
1st year of college
2nd year of college (bachelor)
3rd year of college (licence)
4th year of college
Graduate (master)
Graduate (PhD)

Word: translation ("translation"; 10th grade) -- n = 138

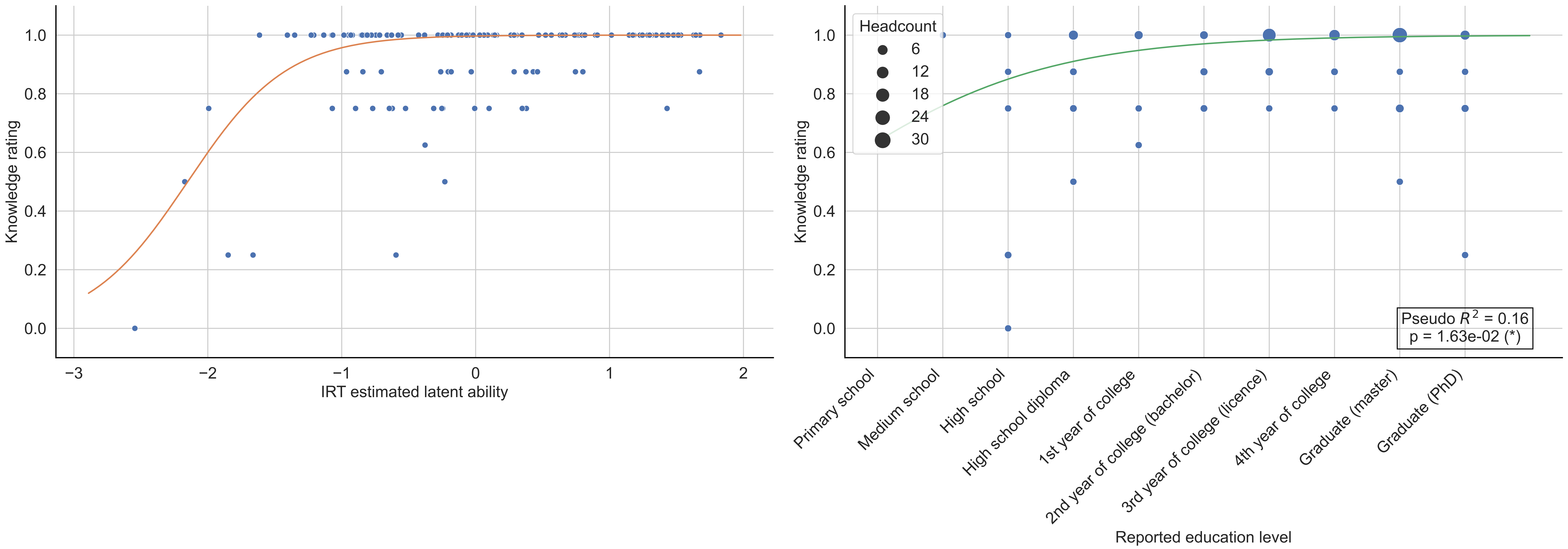

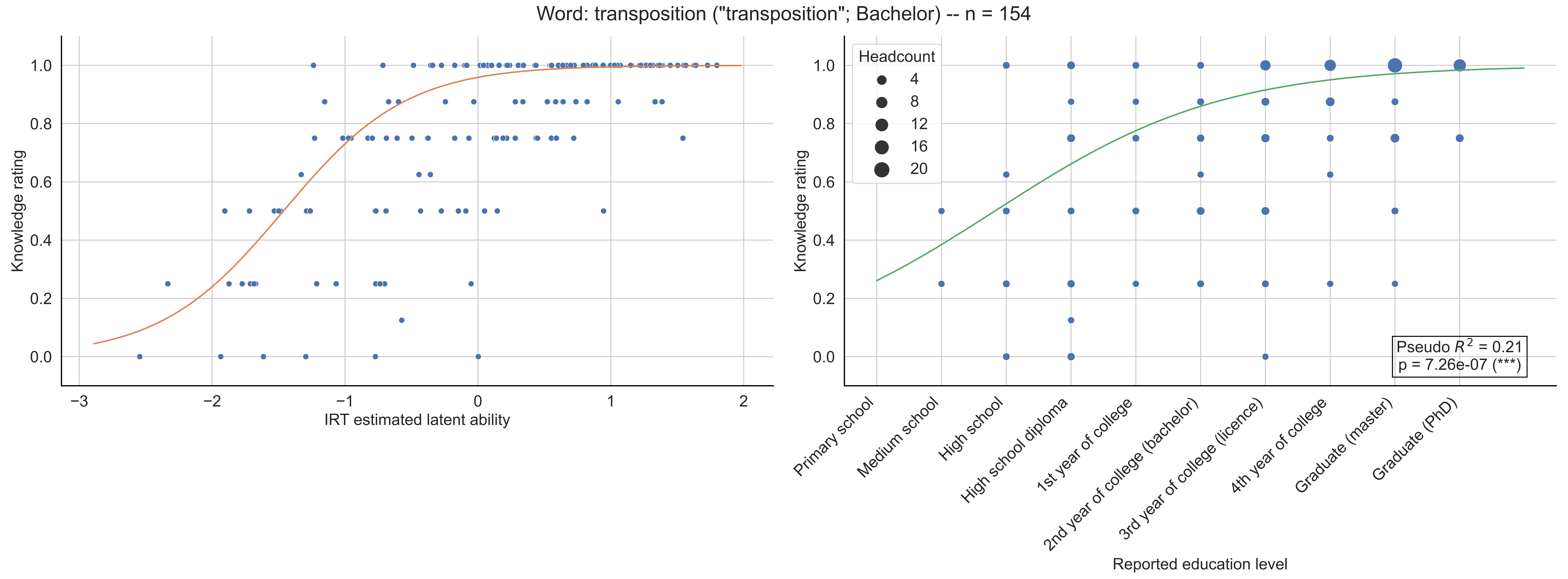
Word: transposition ("transposition"; Bachelor) -- n = 154
Knowledge rating
IRT estimated latent ability
−3
−2
−1
0
1
2
0.0
0.2
0.4
0.6
0.8
1.0
Headcount
4
8
12
16
20
Pseudo $R^2$ = 0.21
p = 7.26e-07 (***)
Primary school
Medium school
High school
High school diploma
1st year of college
2nd year of college (bachelor)
3rd year of college (licence)
4th year of college
Graduate (master)
Graduate (PhD)
Reported education level

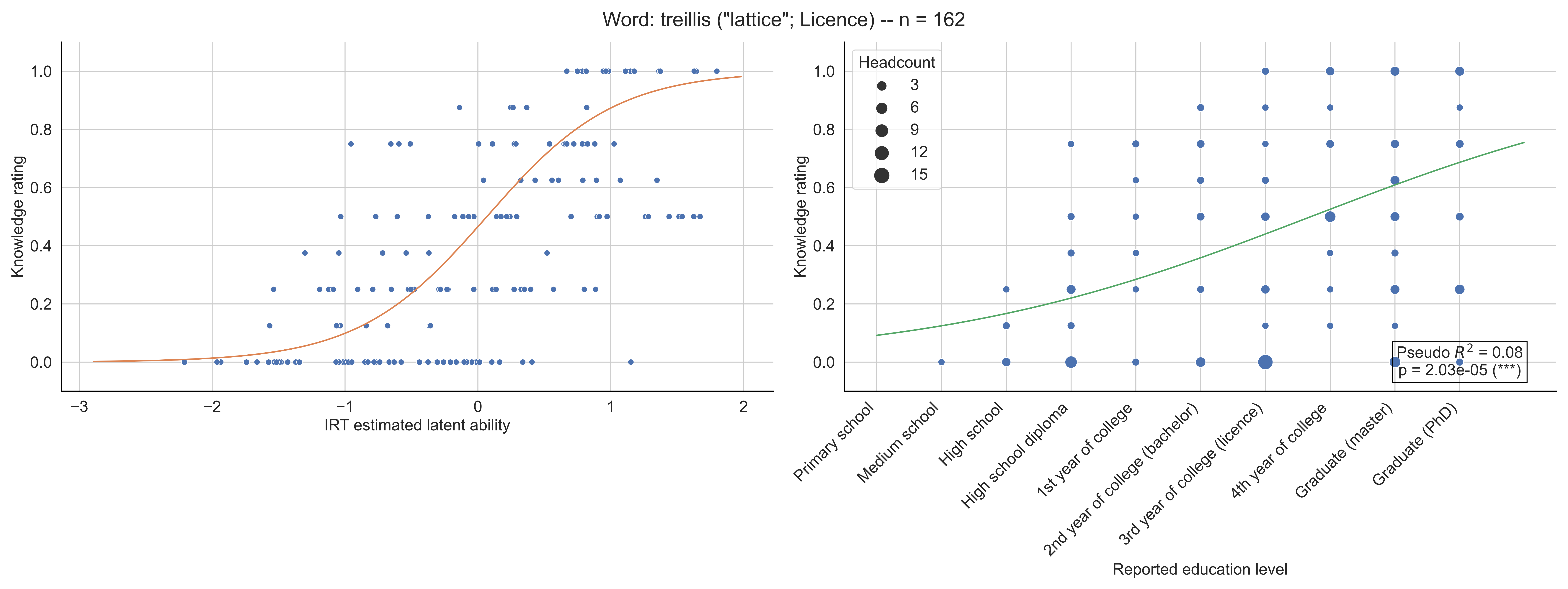

Word: treillis ("lattice"; Licence) -- n = 162
Knowledge rating
IRT estimated latent ability
Headcount
3
6
9
12
15
Pseudo $R^2$ = 0.08
p = 2.03e-05 (***)
Primary school
Medium school
High school
High school diploma
1st year of college
2nd year of college (bachelor)
3rd year of college (licence)
4th year of college
Graduate (master)
Graduate (PhD)
Reported education level

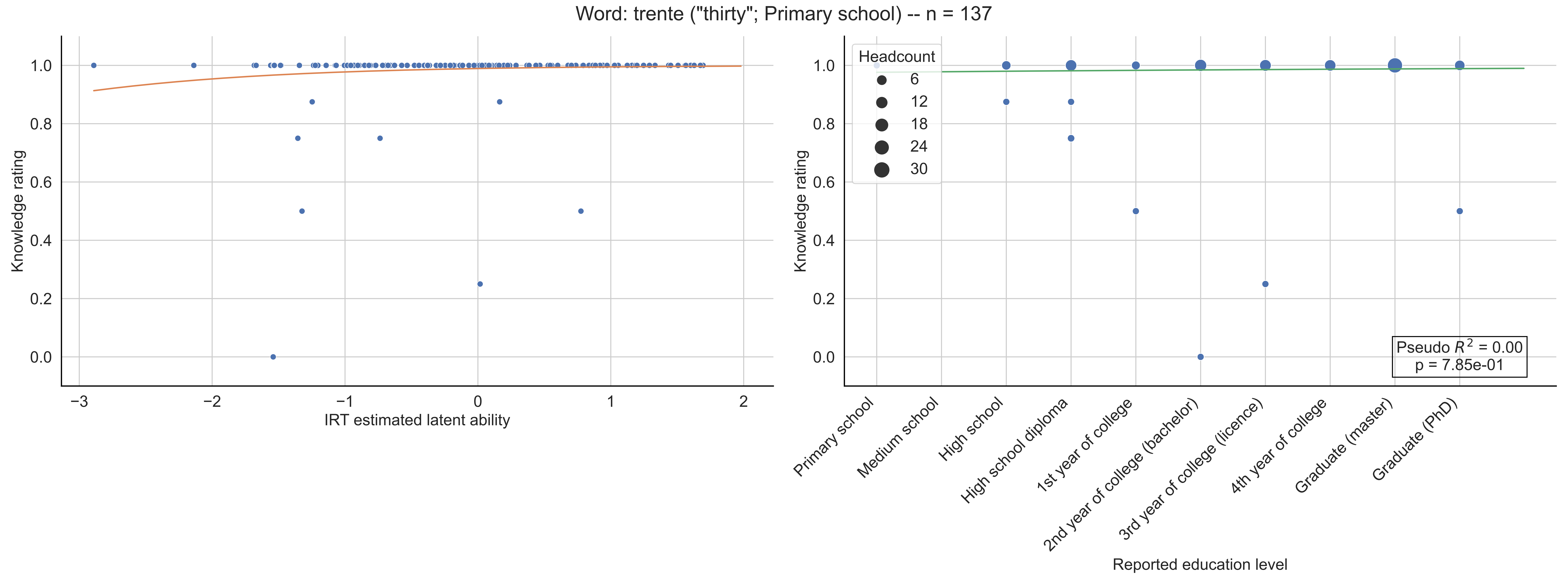

Word: trente ("thirty"; Primary school) -- n = 137
Knowledge rating
IRT estimated latent ability
Headcount
6
12
18
24
30
Pseudo $R^2$ = 0.00
p = 7.85e-01
Primary school
Medium school
High school
High school diploma
1st year of college
2nd year of college (bachelor)
3rd year of college (licence)
4th year of college
Graduate (master)
Graduate (PhD)
Reported education level

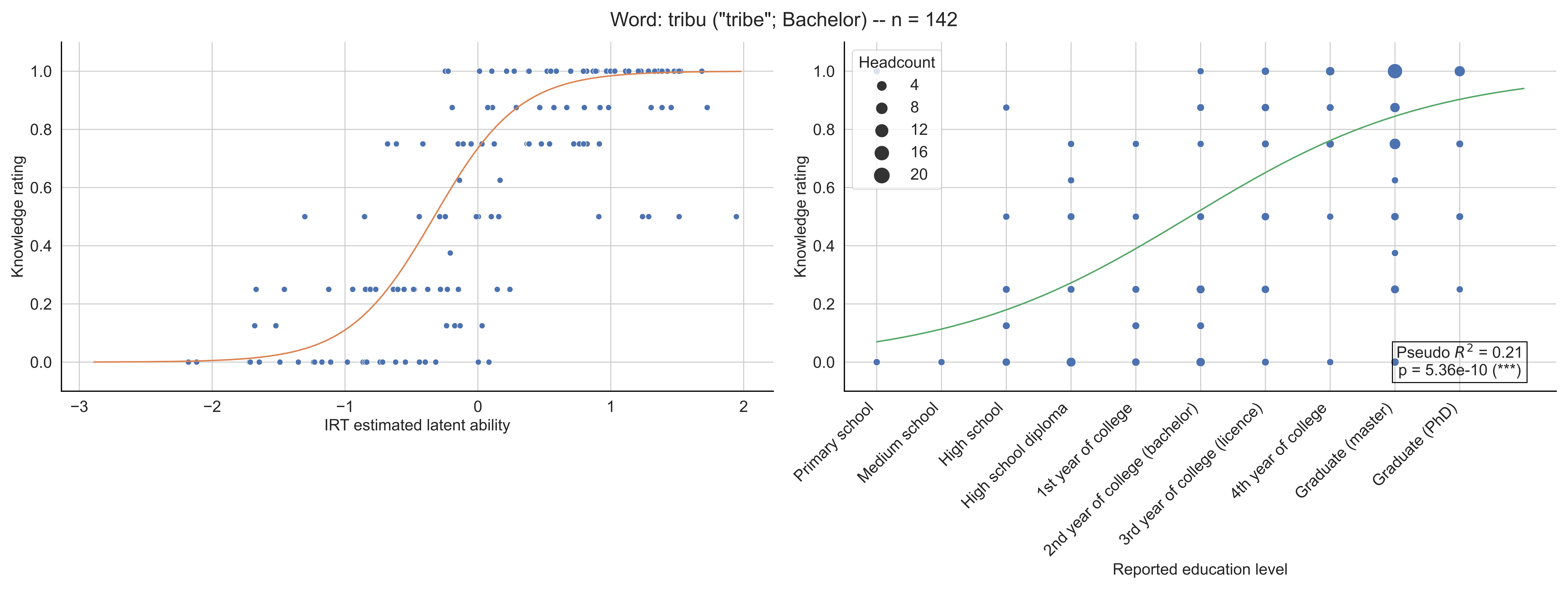

Word: tribu ("tribe"; Bachelor) -- n = 142
Knowledge rating
IRT estimated latent ability
Headcount
4
8
12
16
20
Pseudo $R^2$ = 0.21
p = 5.36e-10 (***)
Primary school
Medium school
High school
High school diploma
1st year of college
2nd year of college (bachelor)
3rd year of college (licence)
4th year of college
Graduate (master)
Graduate (PhD)
Reported education level

Word: trigonométrie ("trigonometry"; 8-9th grade) -- n = 135

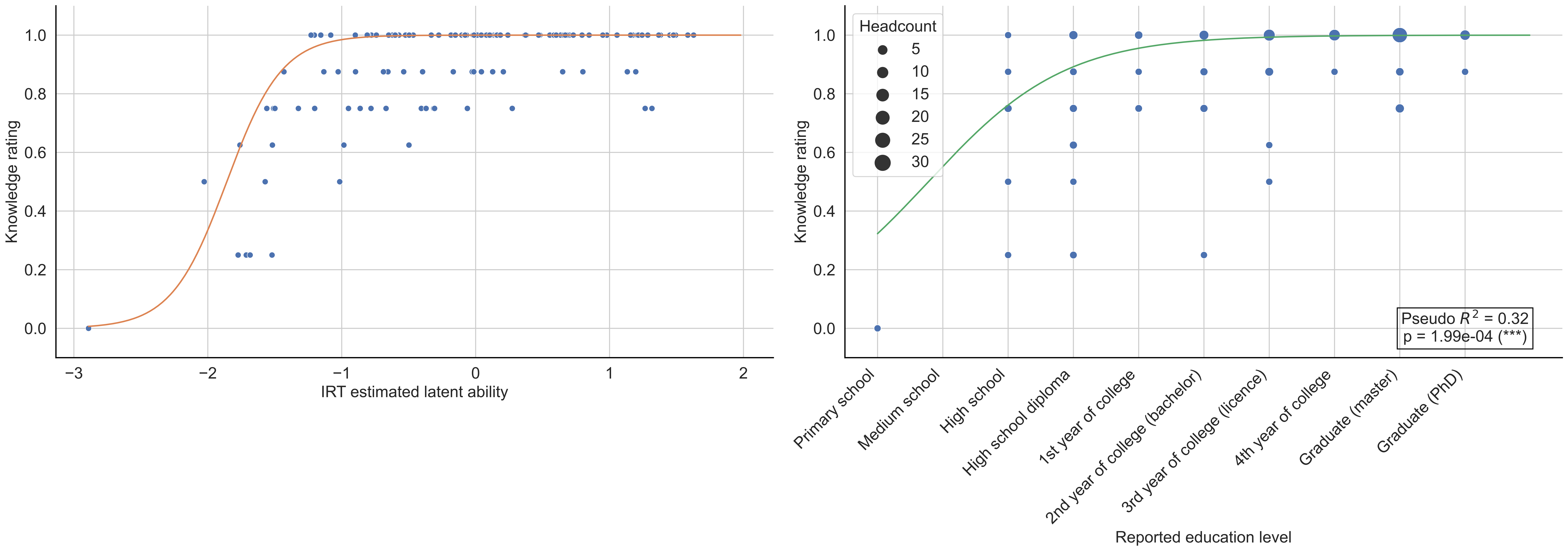

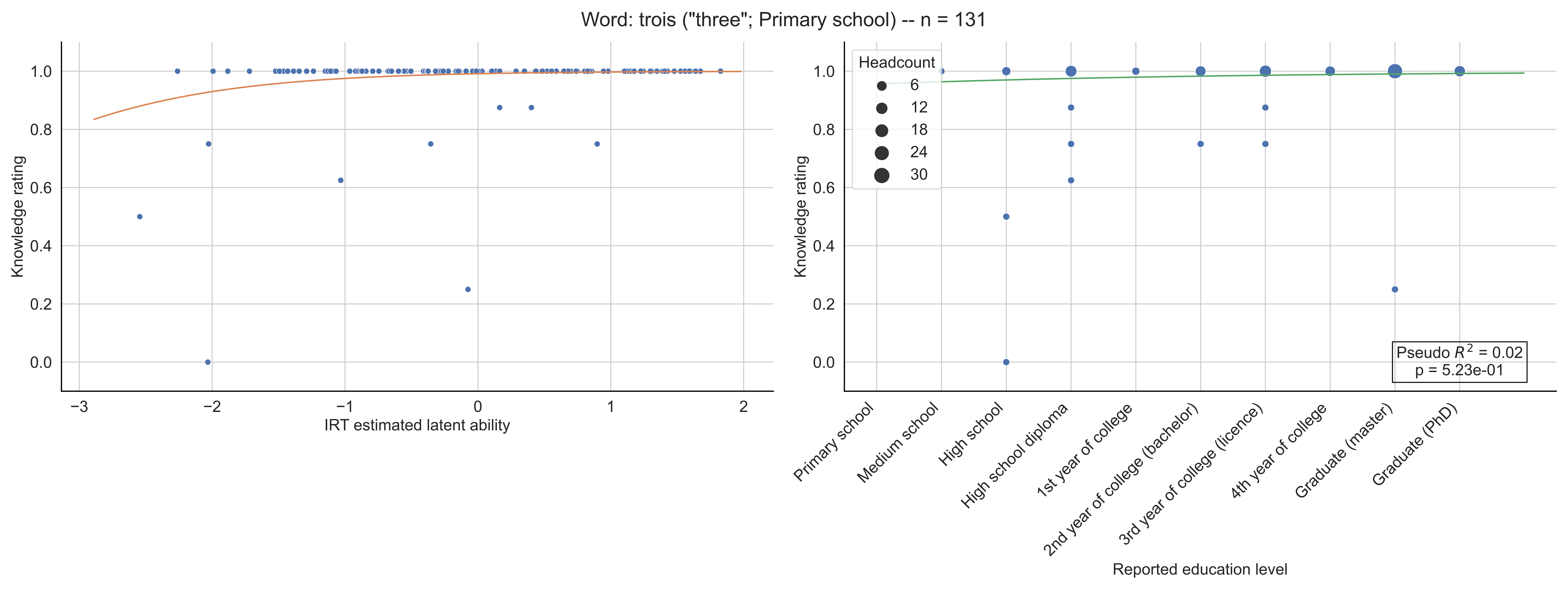

Word: trois ("three"; Primary school) -- n = 131
Knowledge rating
IRT estimated latent ability
Headcount
6
12
18
24
30
Pseudo $R^2$ = 0.02
p = 5.23e-01
Primary school
Medium school
High school
High school diploma
1st year of college
2nd year of college (bachelor)
3rd year of college (licence)
4th year of college
Graduate (master)
Graduate (PhD)
Reported education level

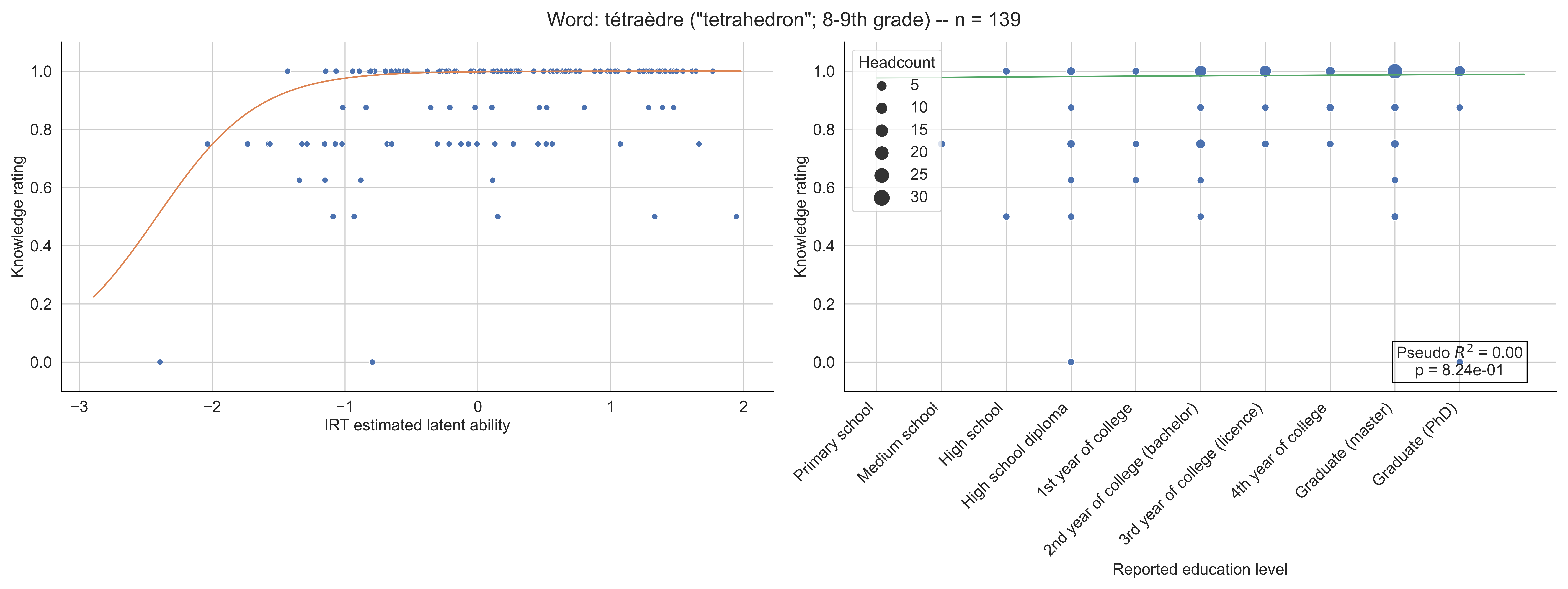

Word: tétraèdre ("tetrahedron"; 8-9th grade) -- n = 139
Knowledge rating
IRT estimated latent ability
Headcount
5
10
15
20
25
30
Pseudo $R^2$ = 0.00
p = 8.24e-01
Primary school
Medium school
High school
High school diploma
1st year of college
2nd year of college (bachelor)
3rd year of college (licence)
4th year of college
Graduate (master)
Graduate (PhD)
Reported education level

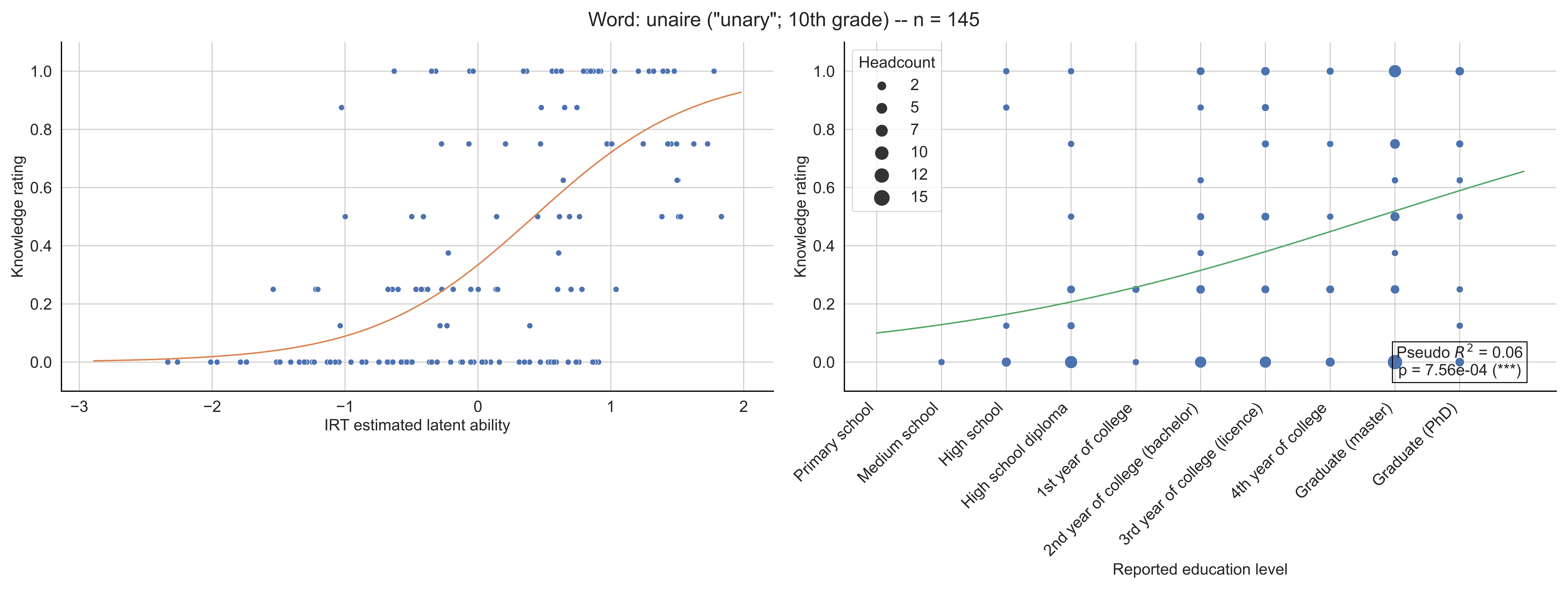

Word: unaire ("unary"; 10th grade) -- n = 145
Knowledge rating
IRT estimated latent ability
Headcount
2
5
7
10
12
15
Pseudo $R^2$ = 0.06
p = 7.56e-04 (***)
Primary school
Medium school
High school
High school diploma
1st year of college
2nd year of college (bachelor)
3rd year of college (licence)
4th year of college
Graduate (master)
Graduate (PhD)
Reported education level

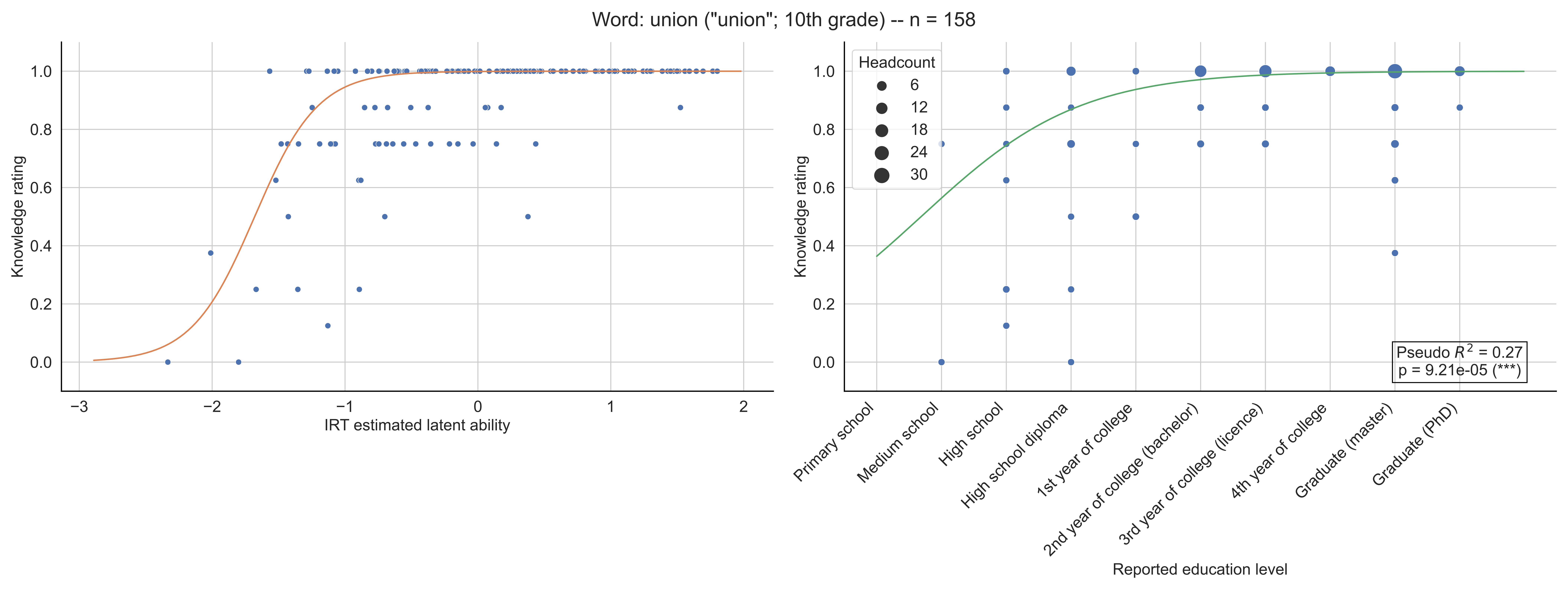
Word: union ("union"; 10th grade) -- n = 158
Knowledge rating
IRT estimated latent ability
Headcount
6
12
18
24
30
Primary school
Medium school
High school
High school diploma
1st year of college
2nd year of college (bachelor)
3rd year of college (licence)
4th year of college
Graduate (master)
Graduate (PhD)
Reported education level
Pseudo $R^2$ = 0.27
p = 9.21e-05 (***)

Word: univers ("universe"; 10th grade) -- n = 149

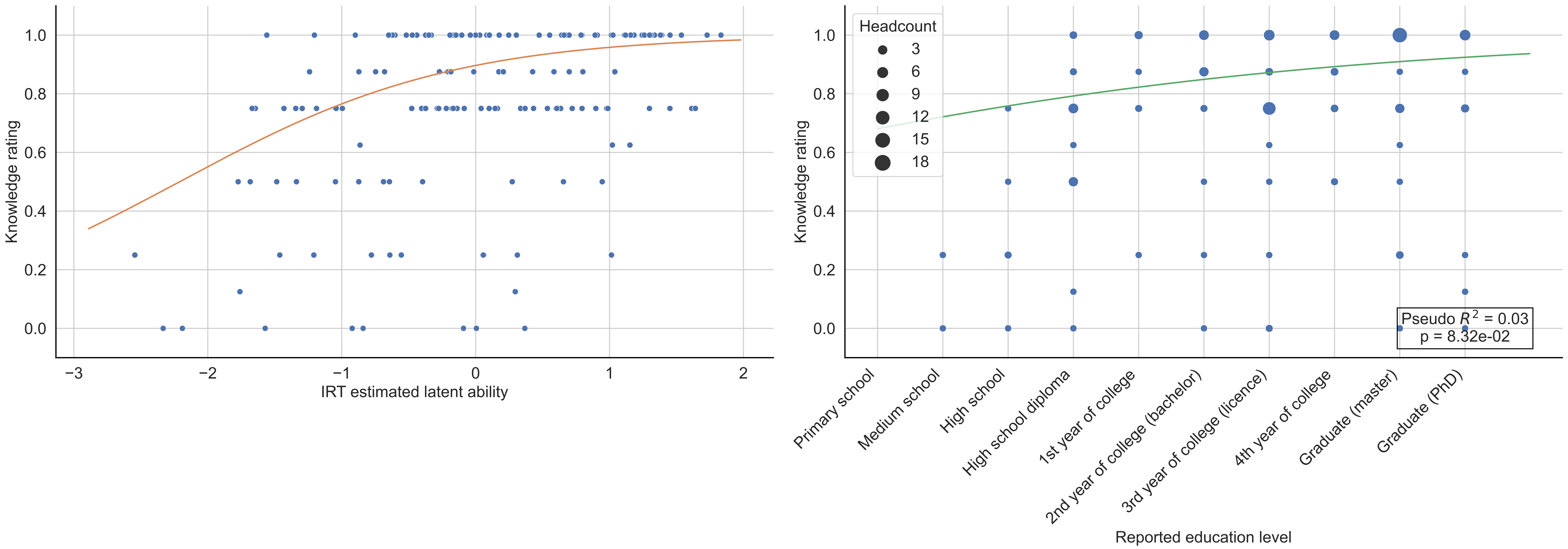

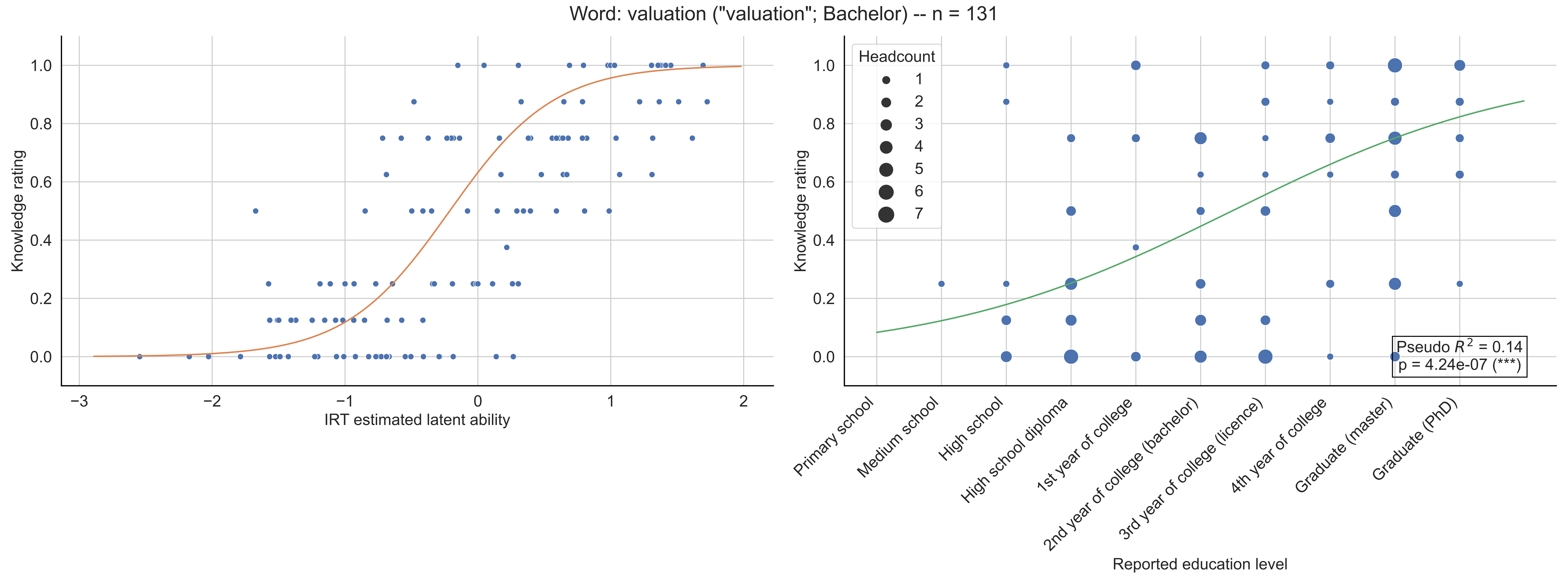

Word: valuation ("valuation"; Bachelor) -- n = 131
Knowledge rating
IRT estimated latent ability
Headcount
1
2
3
4
5
6
7
Pseudo $R^2$ = 0.14
p = 4.24e-07 (***)
Primary school
Medium school
High school
High school diploma
1st year of college
2nd year of college (bachelor)
3rd year of college (licence)
4th year of college
Graduate (master)
Graduate (PhD)
Reported education level

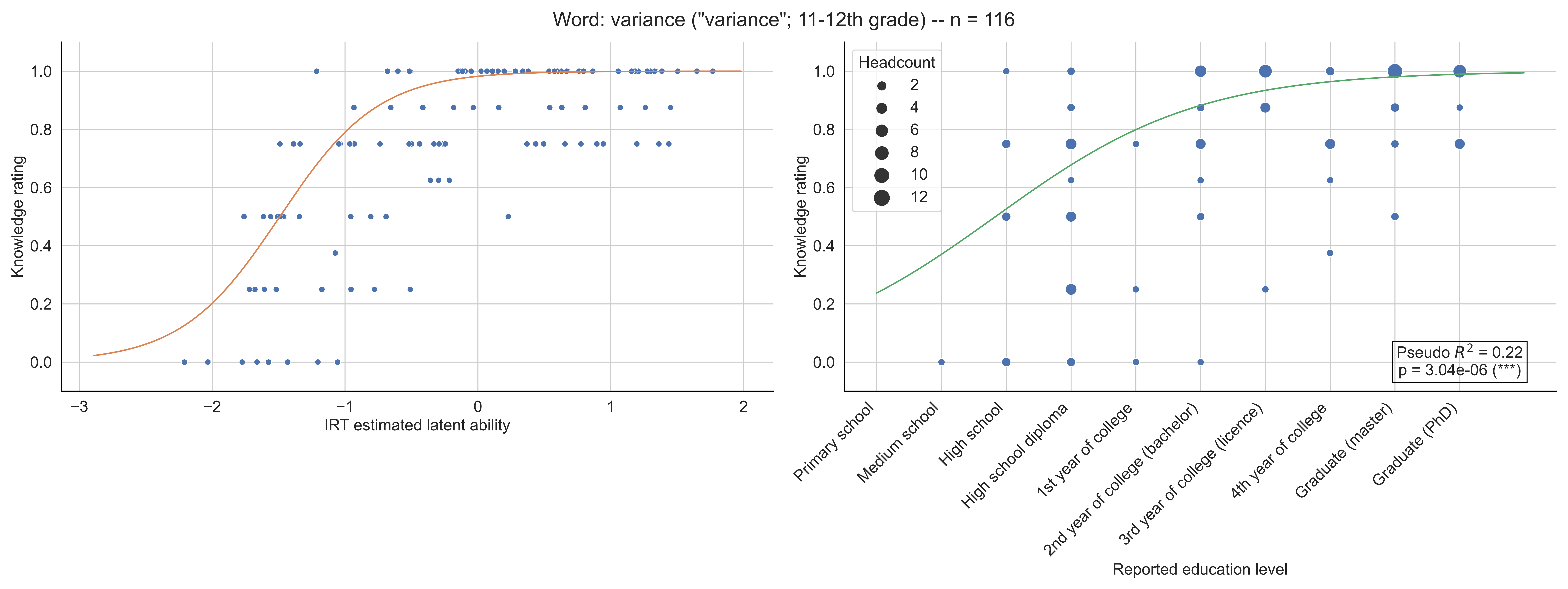

Word: variance ("variance"; 11-12th grade) -- n = 116
Knowledge rating
IRT estimated latent ability
Headcount
2
4
6
8
10
12
Pseudo $R^2$ = 0.22
p = 3.04e-06 (***)
Primary school
Medium school
High school
High school diploma
1st year of college
2nd year of college (bachelor)
3rd year of college (licence)
4th year of college
Graduate (master)
Graduate (PhD)
Reported education level

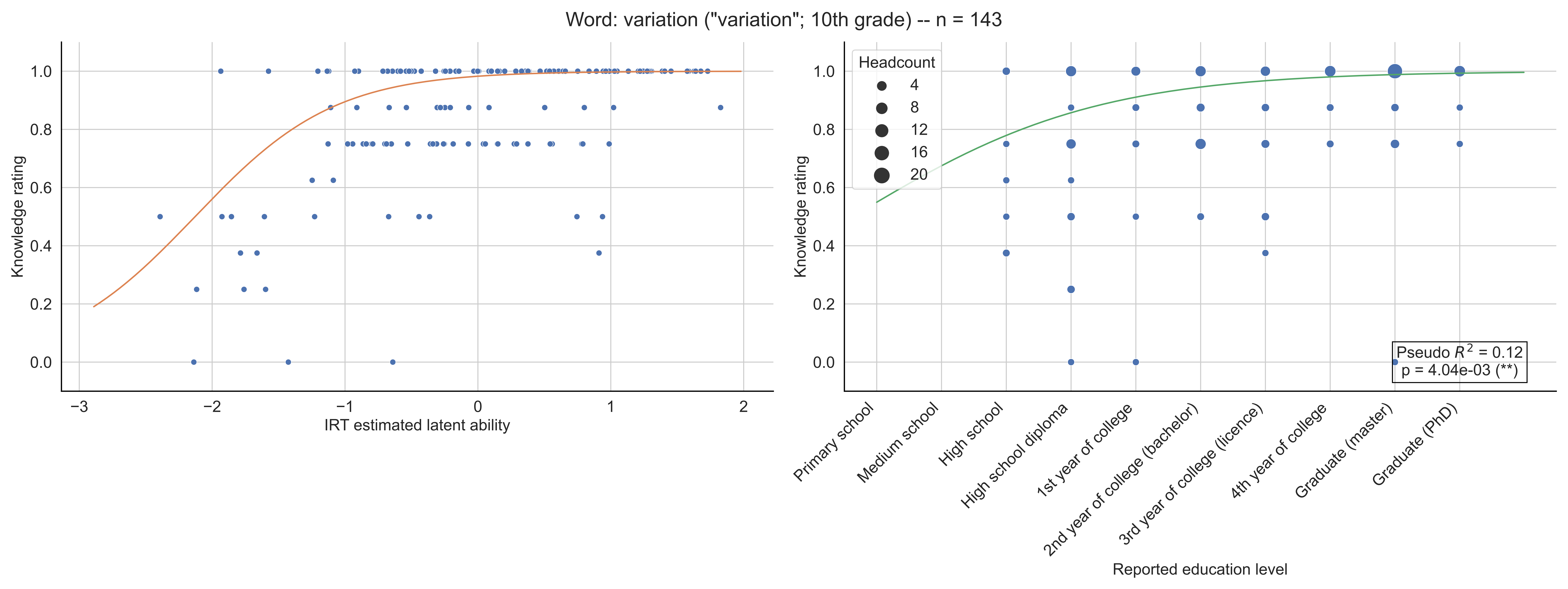

Word: variation ("variation"; 10th grade) -- n = 143
Knowledge rating
IRT estimated latent ability
Headcount
4
8
12
16
20
Pseudo $R^2$ = 0.12
p = 4.04e-03 (**)
Primary school
Medium school
High school
High school diploma
1st year of college
2nd year of college (bachelor)
3rd year of college (licence)
4th year of college
Graduate (master)
Graduate (PhD)
Reported education level

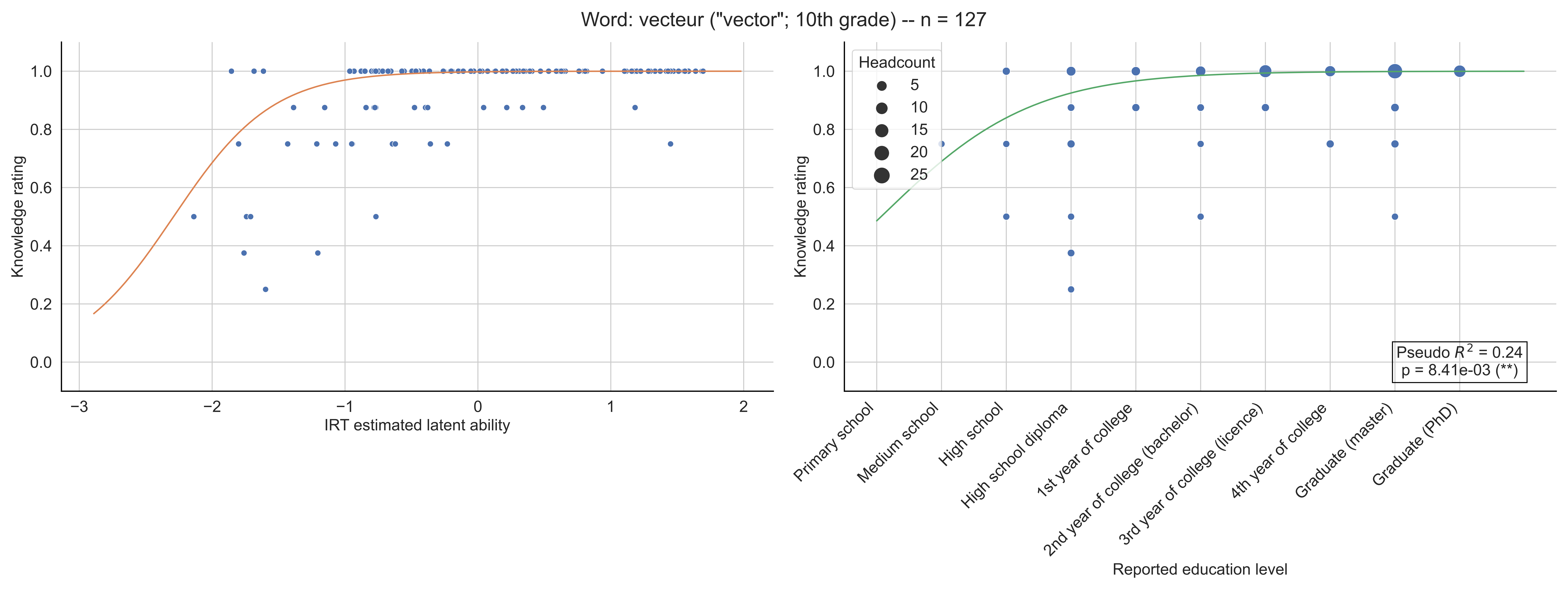

Word: vecteur ("vector"; 10th grade) -- n = 127
Knowledge rating
IRT estimated latent ability
Headcount
5
10
15
20
25
Pseudo $R^2$ = 0.24
p = 8.41e-03 (**)
Primary school
Medium school
High school
High school diploma
1st year of college
2nd year of college (bachelor)
3rd year of college (licence)
4th year of college
Graduate (master)
Graduate (PhD)
Reported education level

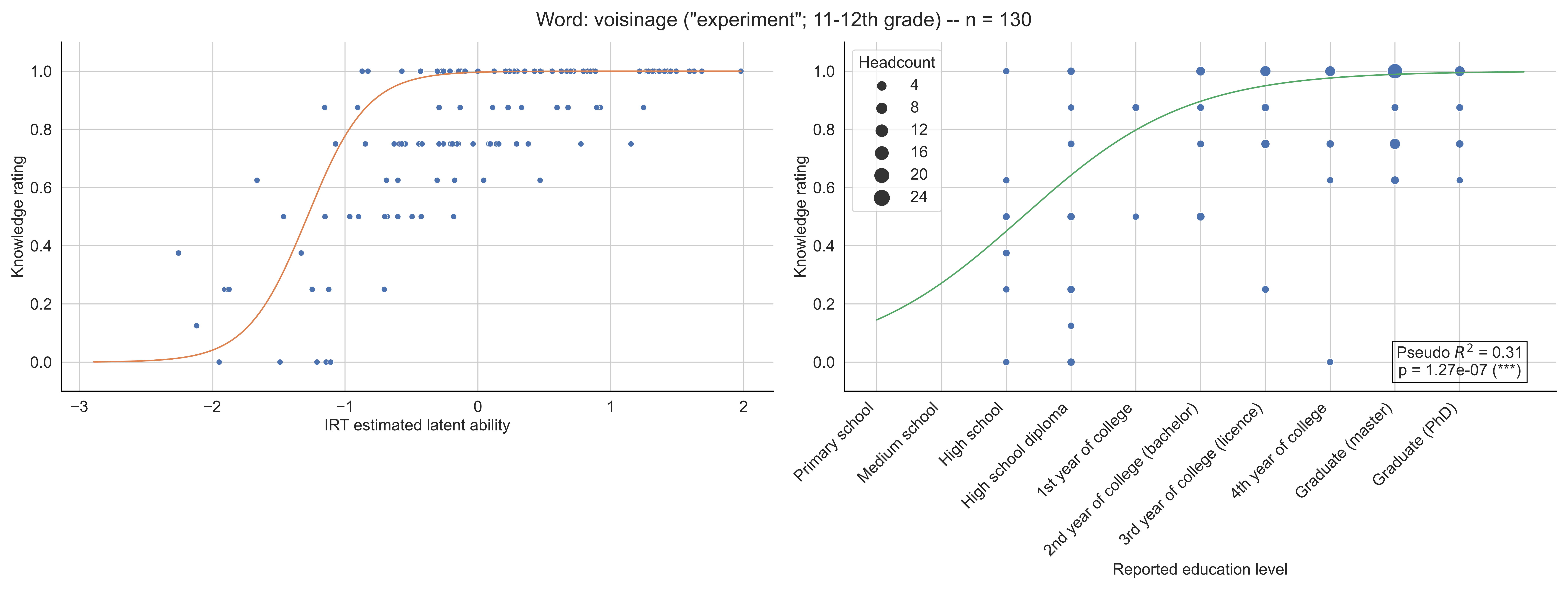

Word: voisinage ("experiment"; 11-12th grade) -- n = 130
Knowledge rating
IRT estimated latent ability
Headcount
4
8
12
16
20
24
Pseudo $R^2$ = 0.31
p = 1.27e-07 (***)
Primary school
Medium school
High school
High school diploma
1st year of college
2nd year of college (bachelor)
3rd year of college (licence)
4th year of college
Graduate (master)
Graduate (PhD)
Reported education level

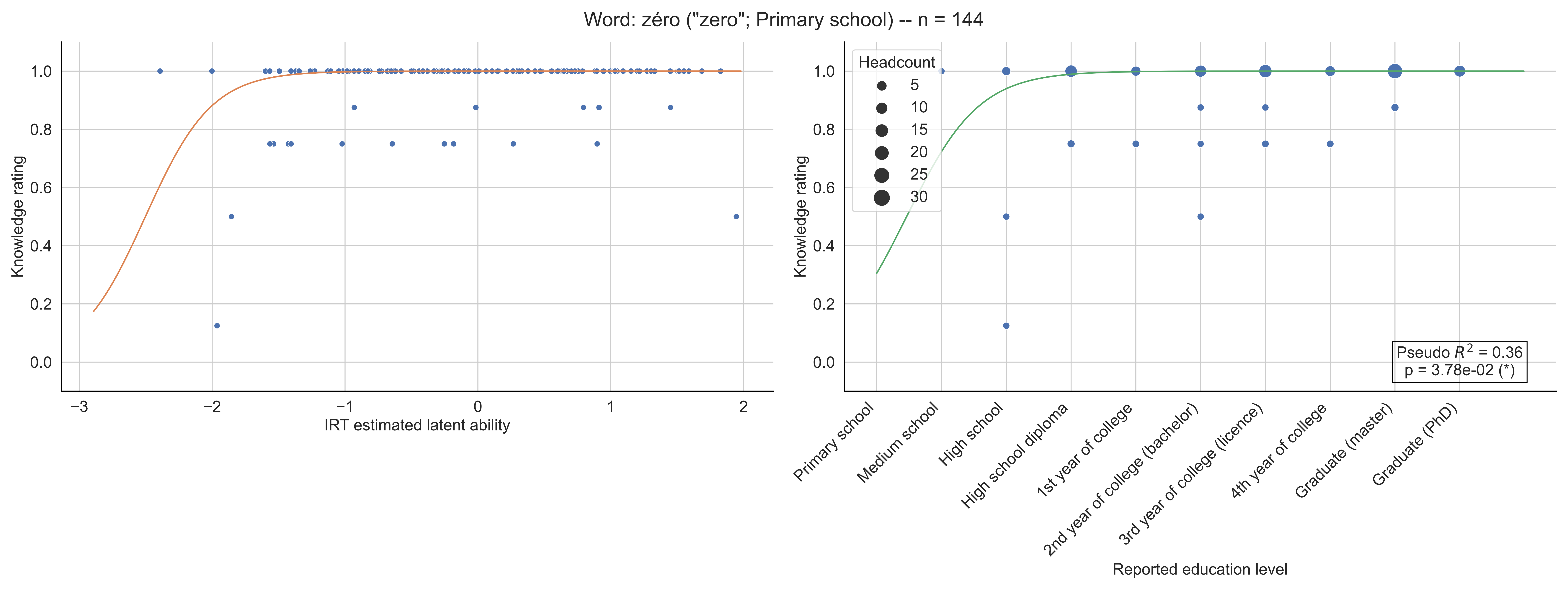

Word: zéro ("zero"; Primary school) -- n = 144
Knowledge rating
IRT estimated latent ability
Headcount
5
10
15
20
25
30
Pseudo $R^2$ = 0.36
p = 3.78e-02 (*)
Primary school
Medium school
High school
High school diploma
1st year of college
2nd year of college (bachelor)
3rd year of college (licence)
4th year of college
Graduate (master)
Graduate (PhD)
Reported education level

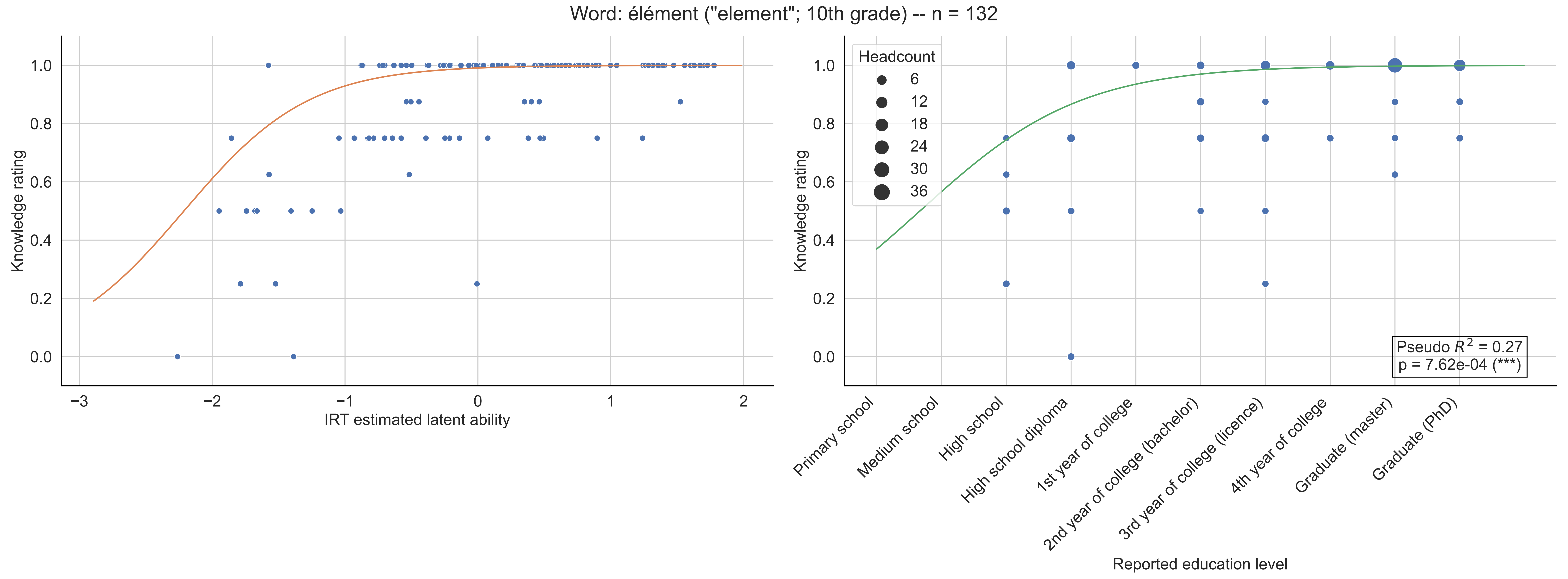
Word: élément ("element"; 10th grade) -- n = 132
Knowledge rating
IRT estimated latent ability
Headcount
6
12
18
24
30
36
Pseudo $R^2$ = 0.27
p = 7.62e-04 (***)
Primary school
Medium school
High school
High school diploma
1st year of college
2nd year of college (bachelor)
3rd year of college (licence)
4th year of college
Graduate (master)
Graduate (PhD)
Reported education level

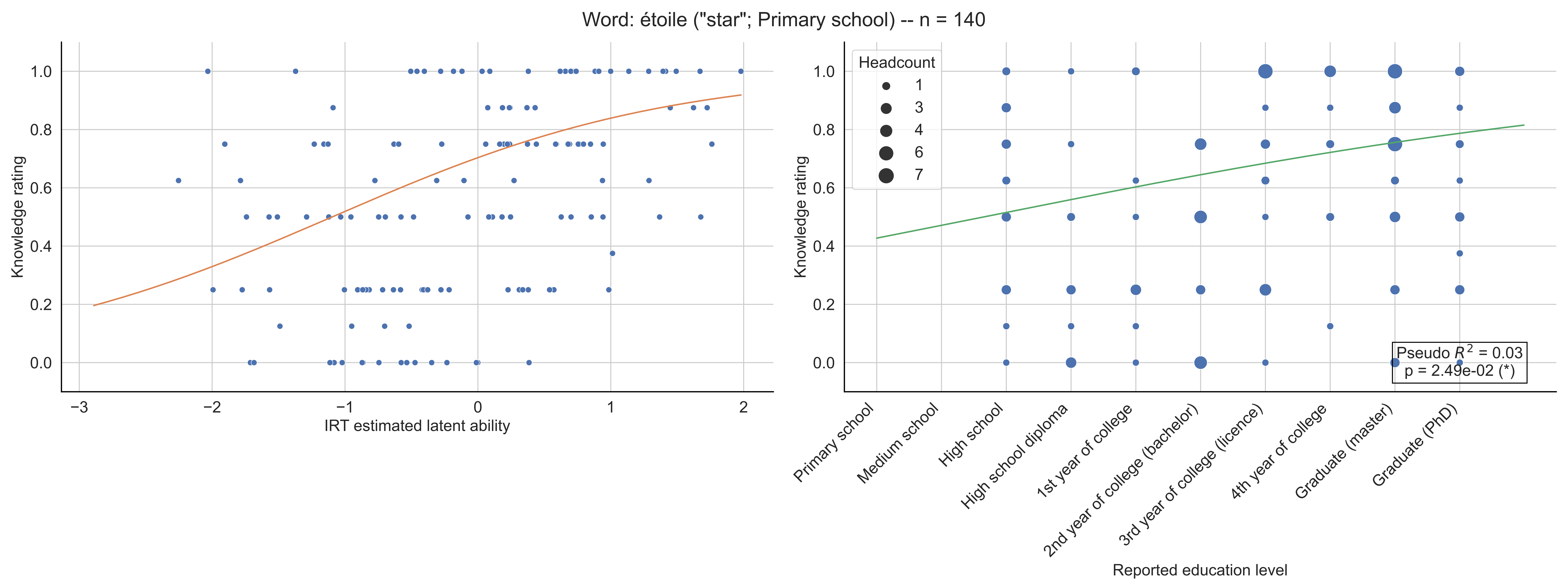

Word: étoile ("star"; Primary school) -- n = 140
Knowledge rating
IRT estimated latent ability
Headcount
1
3
4
6
7
Pseudo $R^2$ = 0.03
p = 2.49e-02 (*)
Primary school
Medium school
High school
High school diploma
1st year of college
2nd year of college (bachelor)
3rd year of college (licence)
4th year of college
Graduate (master)
Graduate (PhD)
Reported education level

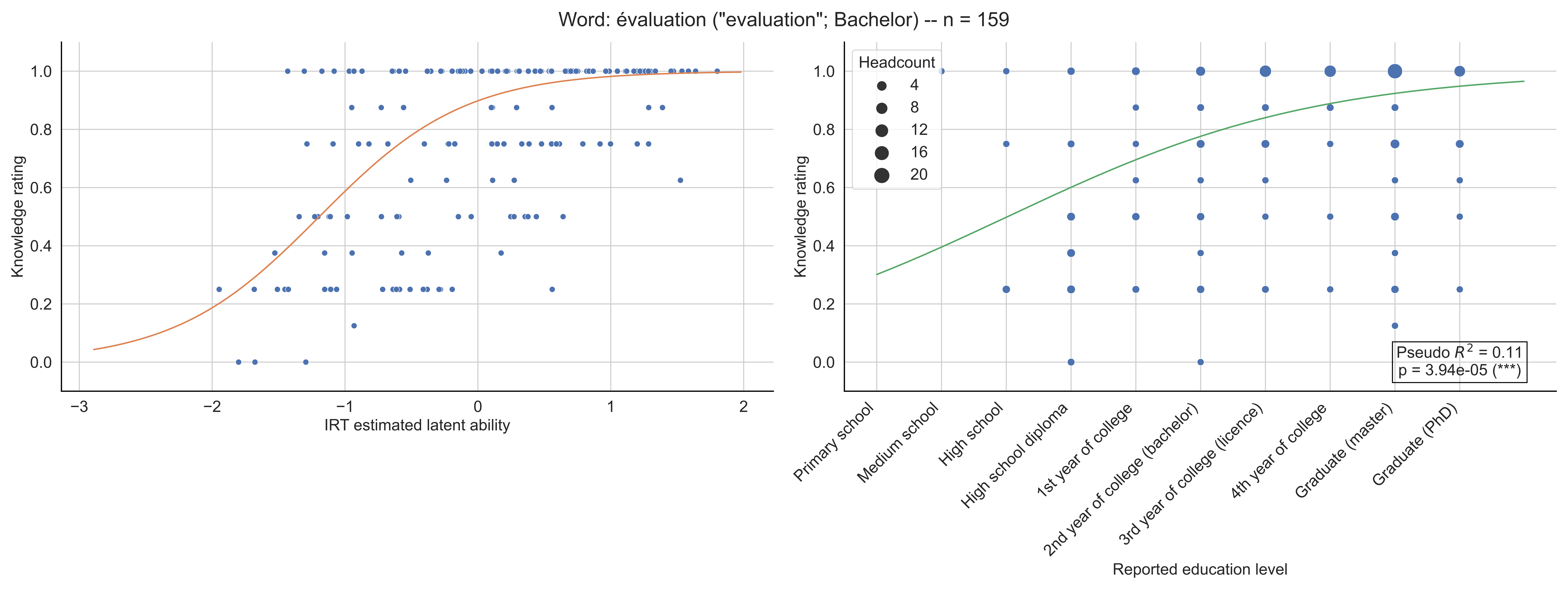
Word: évaluation ("evaluation"; Bachelor) -- n = 159
Knowledge rating
IRT estimated latent ability
Headcount
4
8
12
16
20
Pseudo $R^2$ = 0.11
p = 3.94e-05 (***)
Primary school
Medium school
High school
High school diploma
1st year of college
2nd year of college (bachelor)
3rd year of college (licence)
4th year of college
Graduate (master)
Graduate (PhD)
Reported education level

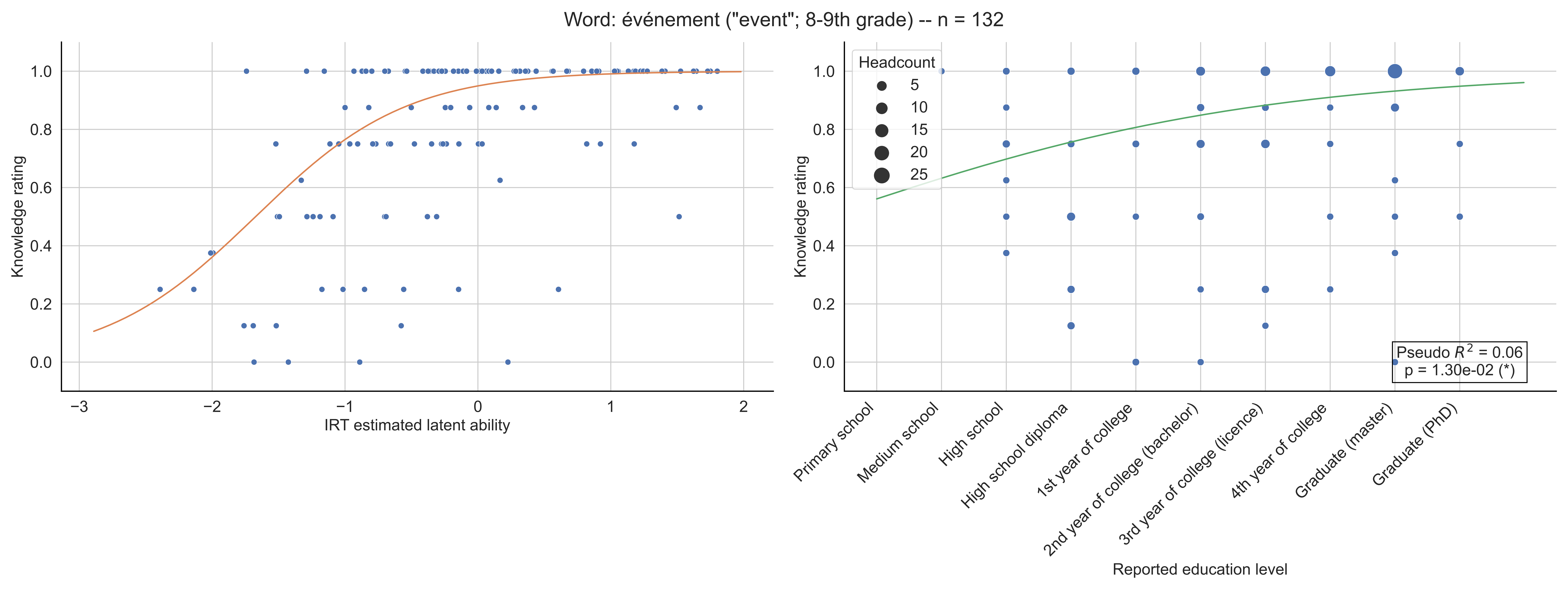
Word: événement ("event"; 8-9th grade) -- n = 132
Knowledge rating
IRT estimated latent ability
Headcount
5
10
15
20
25
Pseudo $R^2$ = 0.06
p = 1.30e-02 (*)
Primary school
Medium school
High school
High school diploma
1st year of college
2nd year of college (bachelor)
3rd year of college (licence)
4th year of college
Graduate (master)
Graduate (PhD)
Reported education level
