## Supplementary PDF 2 for "Mapping and modeling the semantic space of math concepts"

### S2 Table. Number of pairs selected for the similarity rating task for each word grade.

| **Word grade** | **Number of pairs selected for the online experiment** |
|---|---|
| Primary school | 1,021 |
| 6-7th grade | 472 |
| 8-9th grade | 202 |
| 10th grade | 334 |
| 11-12th grade | 433 |
| bachelor | 789 |
| *licence* | 370 |
| master | 135 |
